## Supplemental figures and tables for "M1BP cooperates with CP190 to activate transcription at TAD borders and promote chromatin insulator activity"

**Supplementary Figure 1**

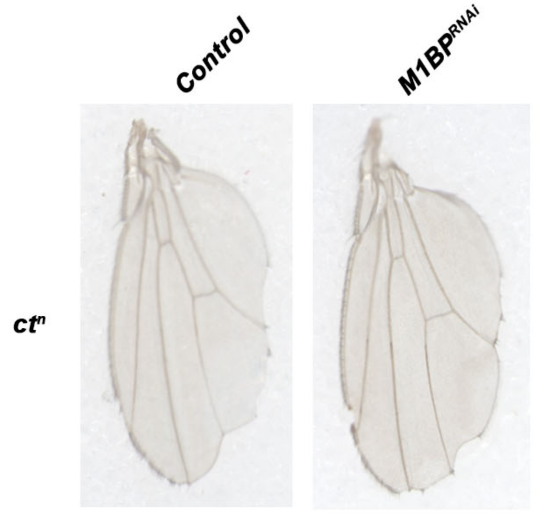

**Supplementary Figure 1. Related to Figure 2. Depletion of M1BP does not affect *gypsy*-independent *ct^n* phenotype.**

Knockdown of *M1BP* does not affect phenotype of the *gypsy*-independent $ct^n$ allele. Wild type (left) and $M1BP^{RNAi}$ lines (right) driven by *Ser-Gal4* display similar wing margin phenotypes for the *gypsy*-independent $ct^n$ loss-of function allele. Fifty flies were assayed for each genotype, and a representative wing is shown.

#### Supplementary Figure 2

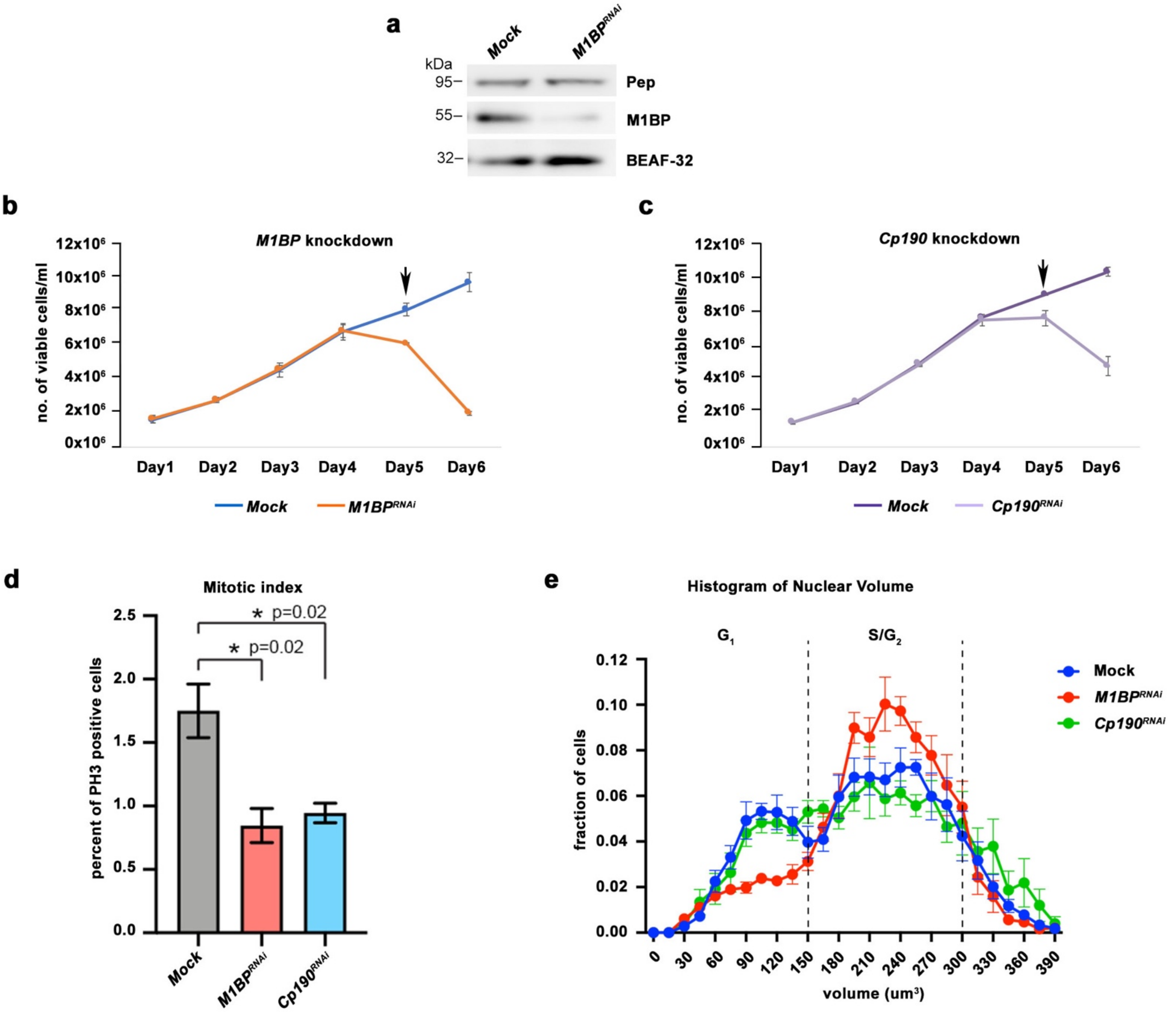

**Supplementary Figure 2. Related to Figure 3 and 6. Functional analysis of M1BP and CP190.**

**a** Western blotting of total lysates from Kc control Mock and *M1BP* knockdown cells showing knockdown efficiency of *M1BP* after 5 days of dsRNA transfection and no effect on protein levels of BEAF-32. Pep is used as loading control. **b** Viable cell counts of Mock or *M1BP* knockdown cells at indicated times after dsRNA transfection in Kc cells. Error bars represent the standard deviation of two biological replicates. **c** Viable cell counts of Mock or *Cp190* knockdown cells at indicated times after dsRNA transfection in Kc cells. Error bars represent the standard deviation of two biological replicates. **d** PH3ser10 positive cells were counted in viable (Tubulin positive) Mock, *M1BP* knockdown and *Cp190* knockdown cells after 5 days of knockdown after transfection. Error bars represent standard deviation of three biological replicates. *P*-values are calculated by t-test; ns, not significant; **P* ≤0.05; ***P* ≤0.01. **e** Graph representing cell cycle stage in Mock, *M1BP* knockdown and *Cp190* knockdown based on nuclear volume.

**Supplementary Figure 3**

**a**

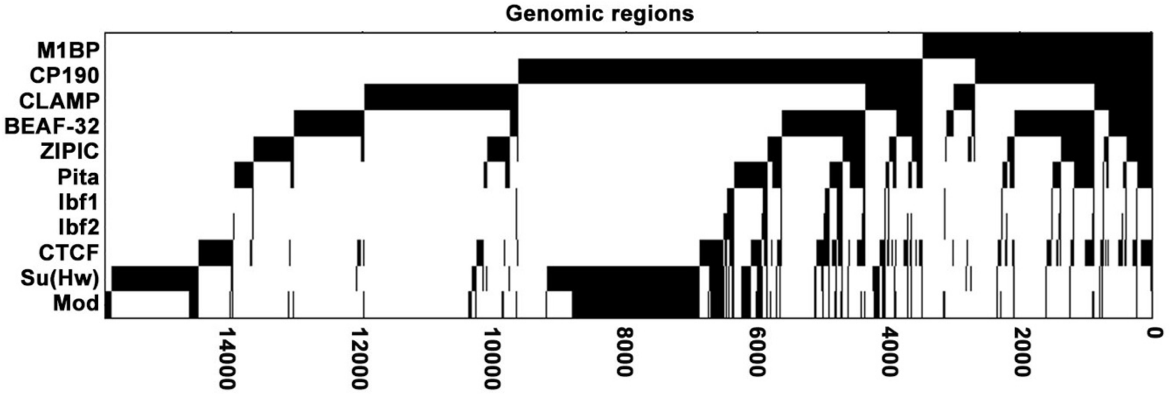

**b**

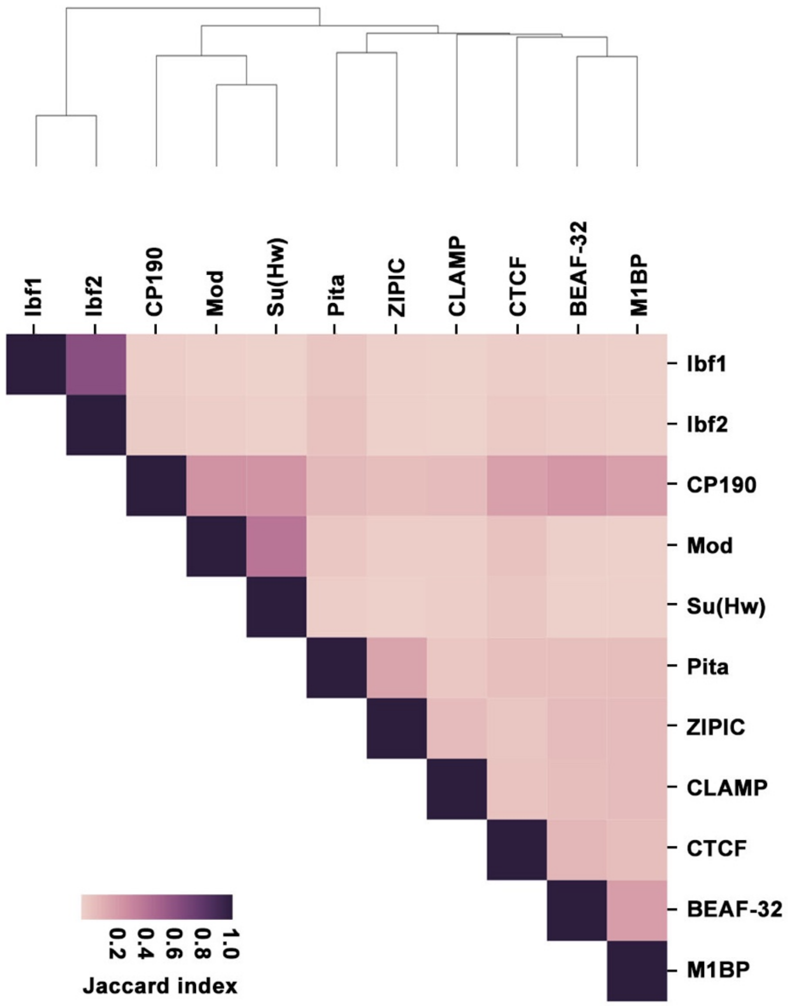

**Supplementary Figure 3. Related to Figure 3. M1BP co-localizes with CP190 and other factors.**

**a** Binary heat map shows comparison of binding of M1BP, CP190, CLAMP, BEAF-32, Su(Hw), Mod(mdg4)67.2, and CTCF in Kc cells and ZIPIC, Pita, Ibf1, and Ibf2 in S2 cells. A black mark in a row indicates presence of the particular factor at that genomic region.

**b** Jaccard heatmap shows heatmap of pairwise comparisons of colocalization of M1BP with *gypsy* components and other factors.

#### Supplementary Figure 4

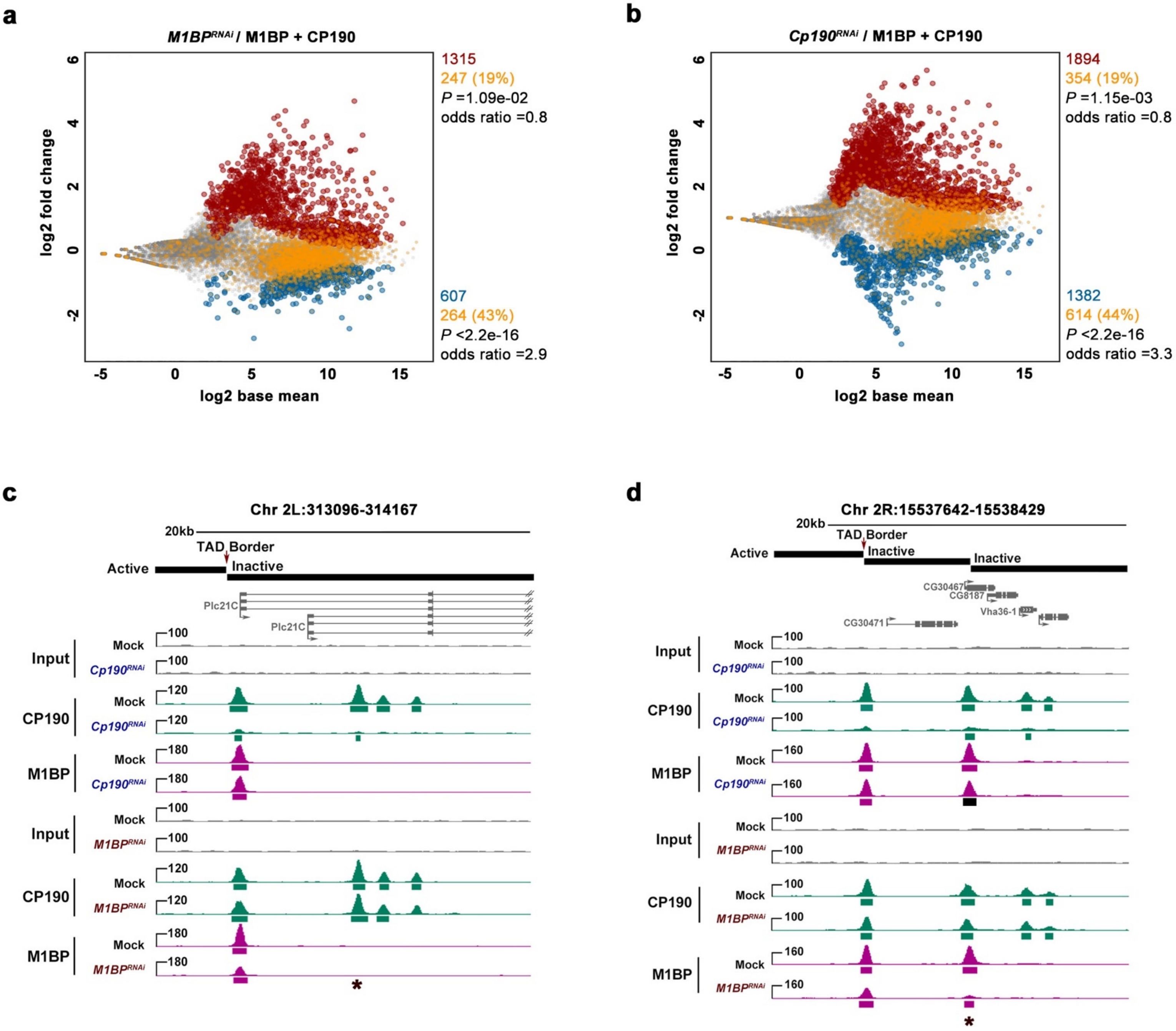

**Supplementary Figure 4. Related to Figure 4 and Figure 6. M1BP and CP190 bind promoters of M1BP- and CP190-activated genes and M1BP promotes CP190 recruitment and vice versa.**

**a** MA plot shows changes in neuRNA levels upon depletion of M1BP. Statistically significant changes include 1315 up-regulated genes (red) and 607 down-regulated genes (blue). Gene promoters containing common overlapping peaks of both M1BP and CP190. FET *P*-values and odds ratios indicate significance of enrichment between affected genes and presence of both M1BP and CP190 peaks at promoters. Peak is additionally colored yellow. **b** MA plot representing affected genes in neuRNA seq after depletion of CP910. Significantly up-regulated genes 1894 (red) and 1382 down-regulated genes (blue) are shown. Common M1BP and CP190 peaks associated with promoters of affected genes are colored yellow. **c** Example screenshot of ChIP-seq profiles showing there is no change in CP190 binding after depletion of M1BP at a site that is not co-bound by M1BP. Asterisk indicates particular peak measured in Figure 6C, site 12. **d** Example screenshot of ChIP-seq profiles showing there is no change in CP190 binding after depletion of M1BP, and M1BP binding after depletion of CP190 at shared site. Asterisk indicates the particular peak measured in Figure 6C, site 11.

**Supplementary Figure 5**

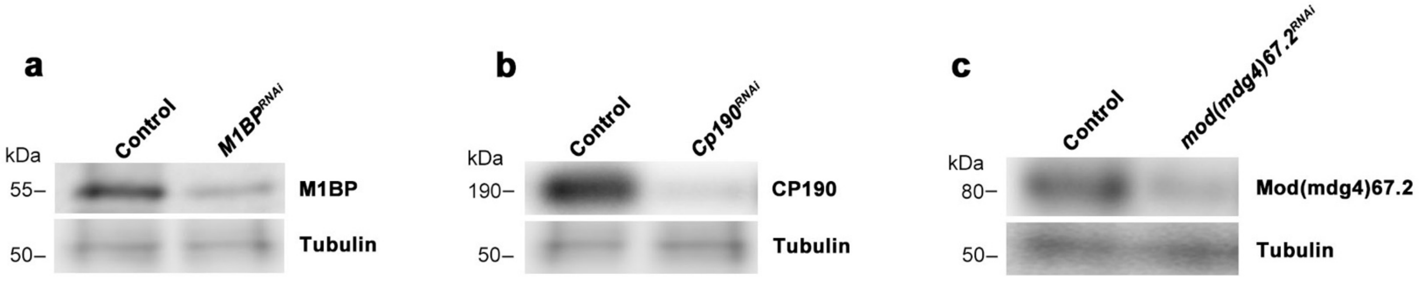

**Supplementary Figure 5. Related to Figure 5. dsRNAs are used to deplete M1BP, CP190 and Mod(mdg4)67.2 in Kc cells.**

**a** Western blot showing the knockdown efficiency of *M1BP*,

**b** *Cp190*,

**c** *mod(mdg4)*.

#### Supplementary Figure 6

**a**

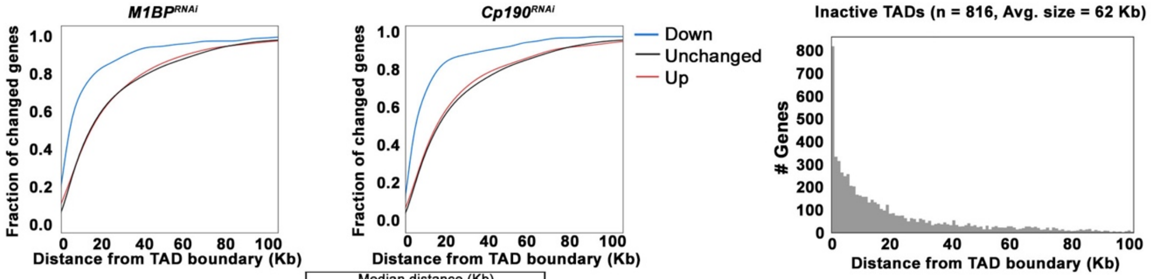

| | | | Median distance (Kb) | | |
|---|---|---|---|---|---|
| Knockdown | Gene changes | # Genes | Affected | Unchanged | FDR |
| $M1BP^{RNAi}$ | Down | 139 | 2.4 | 12.2 | 2.2e-17 |
| | Up | 532 | 12.2 | 12.2 | 0.16 |
| | Unchanged | 5847 | | | |
| $Cp190^{RNAi}$ | Down | 324 | 3.0 | 12.7 | 2.5e-36 |
| | Up | 773 | 11.5 | 12.7 | 0.076 |
| | Unchanged | 5421 | | | |

**b**

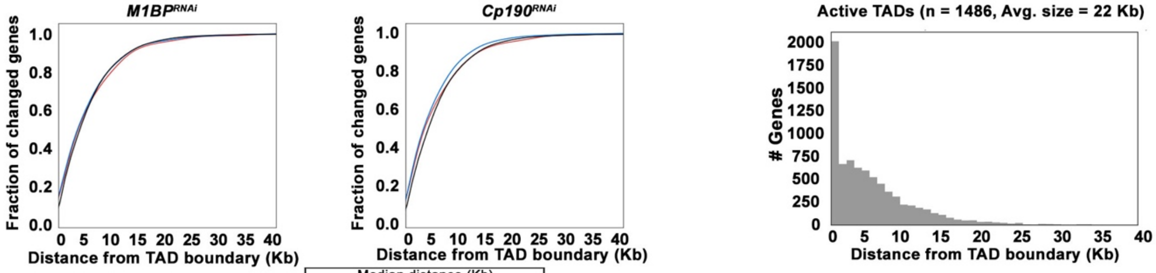

| | | | Median distance (Kb) | | |
|---|---|---|---|---|---|
| Knockdown | Gene changes | # Genes | Affected | Unchanged | FDR |
| $M1BP^{RNAi}$ | Down | 417 | 3.3 | 3.9 | 0.020 |
| | Up | 462 | 3.4 | 3.9 | 0.11 |
| | Unchanged | 6762 | | | |
| $Cp190^{RNAi}$ | Down | 922 | 3.0 | 4.0 | 5.2e-07 |
| | Up | 687 | 3.0 | 4.0 | 3.0e-03 |
| | Unchanged | 6032 | | | |

**c**

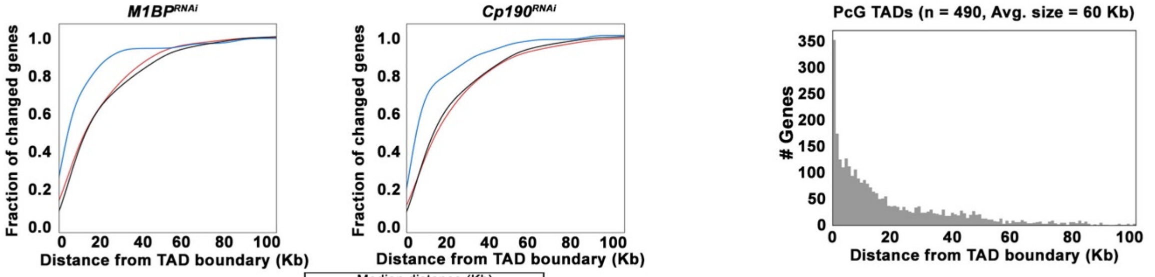

| | | | Median distance (Kb) | | |
|---|---|---|---|---|---|
| Knockdown | Gene changes | # Genes | Affected | Unchanged | FDR |
| $M1BP^{RNAi}$ | Down | 40 | 2.0 | 11.8 | 4.3e-06 |
| | Up | 265 | 10.6 | 11.8 | 0.058 |
| | Unchanged | 2698 | | | |
| $Cp190^{RNAi}$ | Down | 99 | 3.3 | 11.8 | 5.9e-10 |
| | Up | 372 | 12.7 | 11.8 | 0.43 |
| | Unchanged | 2532 | | | |

**d**

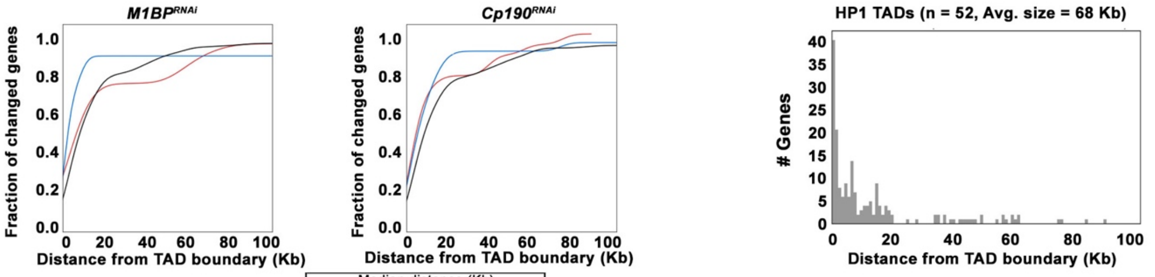

| | | | Median distance (Kb) | | |
|---|---|---|---|---|---|
| Knockdown | Gene changes | # Genes | Affected | Unchanged | FDR |
| $M1BP^{RNAi}$ | Down | 9 | 1.4 | 6.9 | 0.057 |
| | Up | 20 | 4.7 | 6.9 | 0.18 |
| | Unchanged | 168 | | | |
| $Cp190^{RNAi}$ | Down | 23 | 2.1 | 6.9 | 0.057 |
| | Up | 19 | 1.6 | 6.9 | 0.057 |
| | Unchanged | 155 | | | |

**Supplementary Figure 6. Related to Figure 6. M1BP and CP190 transcriptionally activate genes near TAD borders genome-wide independent of chromatin state.** Cumulative histograms of promoter distance from closest TAD border for (**a**) inactive, (**b**) active, (**c**) PcG, and (**d**) HP1 TADs classified by change in nascent expression in *M1BP* (left) or *Cp190* (right) knockdown cells. Downregulated (blue), upregulated (red), or unchanged (black) genes are indicated. Tables below each graph indicate number of changed genes for upregulated, downregulated, and unchanged genes for each knockdown and median distance from closest TAD border. Genes with promoters not located inside a TAD were excluded from the analysis. Mann-Whitney U test for each set of changed genes against unchanged genes are shown. Average sizes and number for each TAD class are also shown. Histogram with number of genes per closest distance to each TAD border is shown. Note that active TADs are the shortest and therefore display the least median distance between borders and all gene promoters yet still display statistically significant differences.

**Supplementary Figure 7**

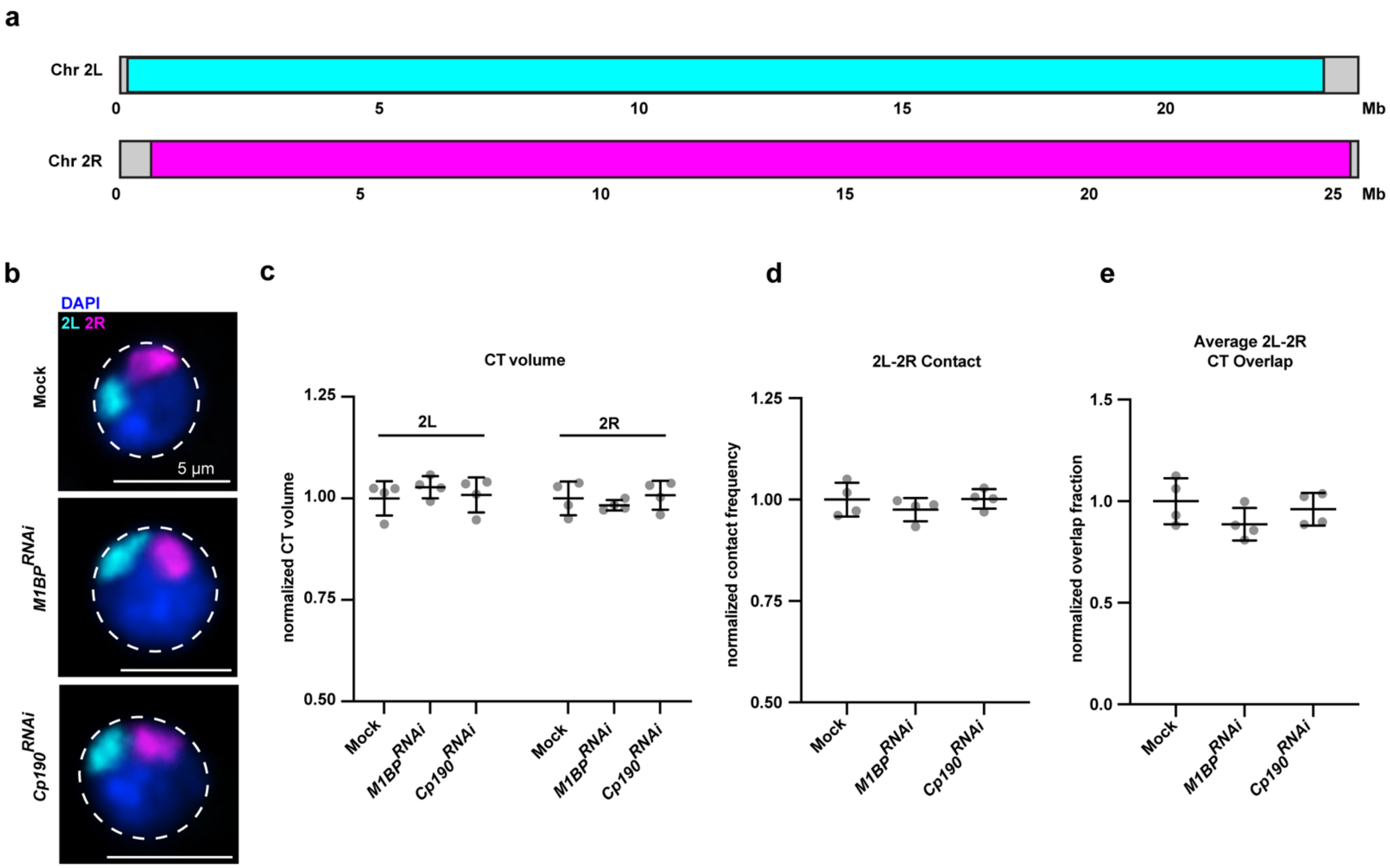

**Supplementary Figure 7. Related to Figure 8. Depletion of M1BP or CP190 does not alter CT formation.**

**a** Schematic of whole chromosome paints used in b-e. **b** Representative images of CTs in control (Mock) or knockdown cells (*M1BP*, *Cp190*). Images are projections of 10 Z stacks. Dashed line represents nuclear edge. **c** Average CT volume shown as a fraction of nuclear volume normalized to controls. 2L, left; 2R, right. **d** Absolute contact frequencies between 2L and 2R (where 100% = contact in all cells) normalized to controls. **e** Average 2L-2R CT intermixing volume as a fraction of 2R volume. All values are normalized to the average of mock controls. All data shown in c-e are from four biological replicates. n>100 cells were measured per replicate. Error bars show standard deviation. Paired t-test of means before normalization to controls for all samples, $P > 0.05$.

#### Supplementary Figure 8

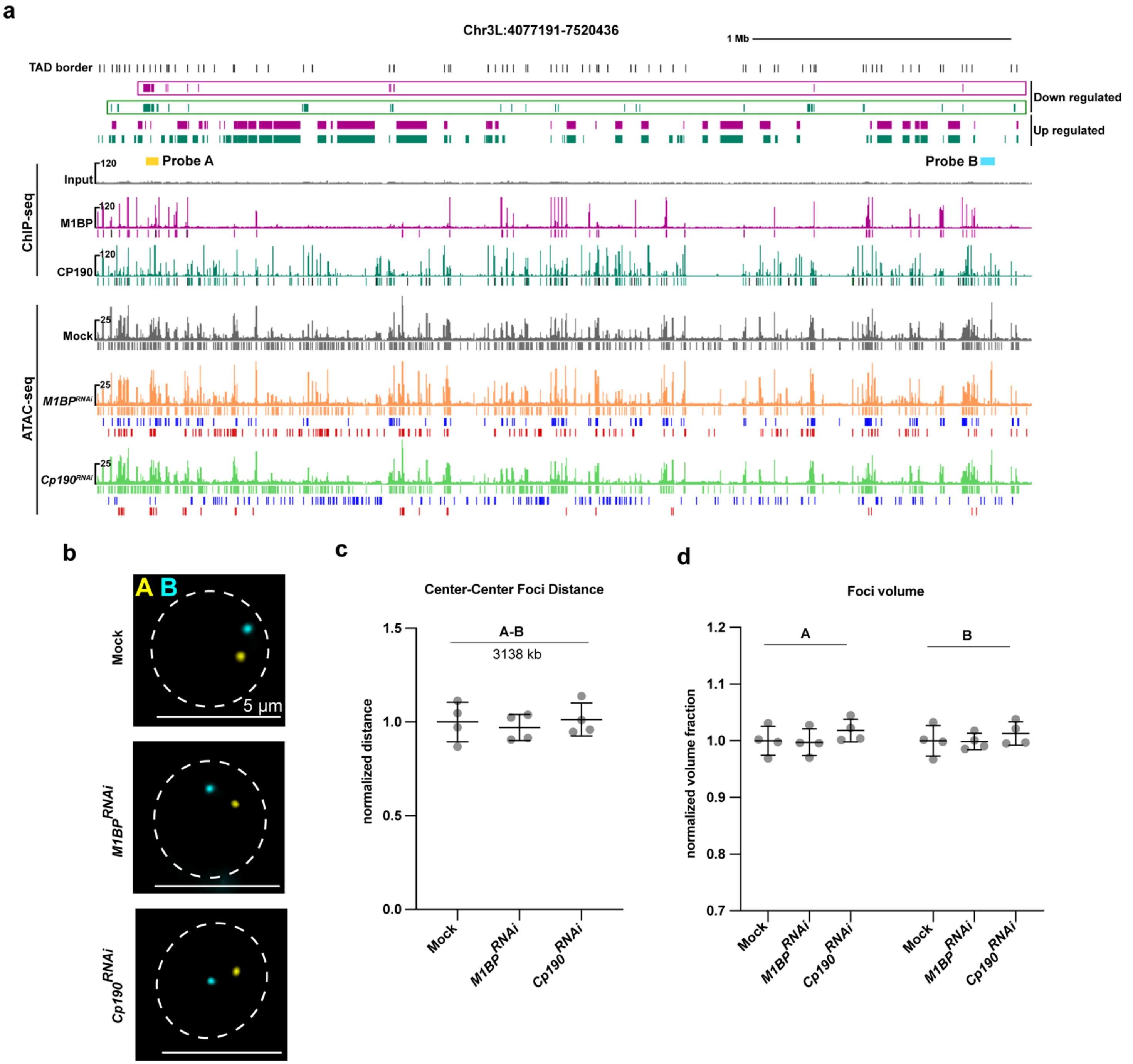

**Supplementary Figure 8. Related to Figure 8. Depletion of M1BP or CP190 does not change long-range intra-chromosomal distance.**

**a** Screenshot of regions detected by A and B probes spaced ~3.1 Mb apart. TAD borders, downregulated (boxed) and upregulated genes in either M1BP (purple) or CP190 (green) knockdowns, ChIP-seq signals of M1BP and CP190, and called peaks are shown. Called ChIP-seq peaks shown in black are decreased in the knockdown of the opposite factor. ATAC-seq signals of mock, M1BP-depleted, and CP190-depleted cells, called peaks, decreased peaks relative to mock (blue bars), and increased peaks (red bars) are also shown. **b** Representative nuclei labeled with probes shown in a. Single Z slices are shown. Dashed line represents nuclear edge. **c** Average center-to-center distance between probes A and B. Single cell distances were normalized to nuclear radius, and all population averages were normalized to their respective controls. **d** Normalized foci volume shown as a fraction of nuclear volume. All averages were normalized to the average of mock controls. All data shown in c-d are from four biological replicates. $n>100$ cells were measured per replicate. Error bars show standard deviation. Paired t-test of means before normalization to controls for all samples, $P > 0.05$.

### Supplementary Figure 9

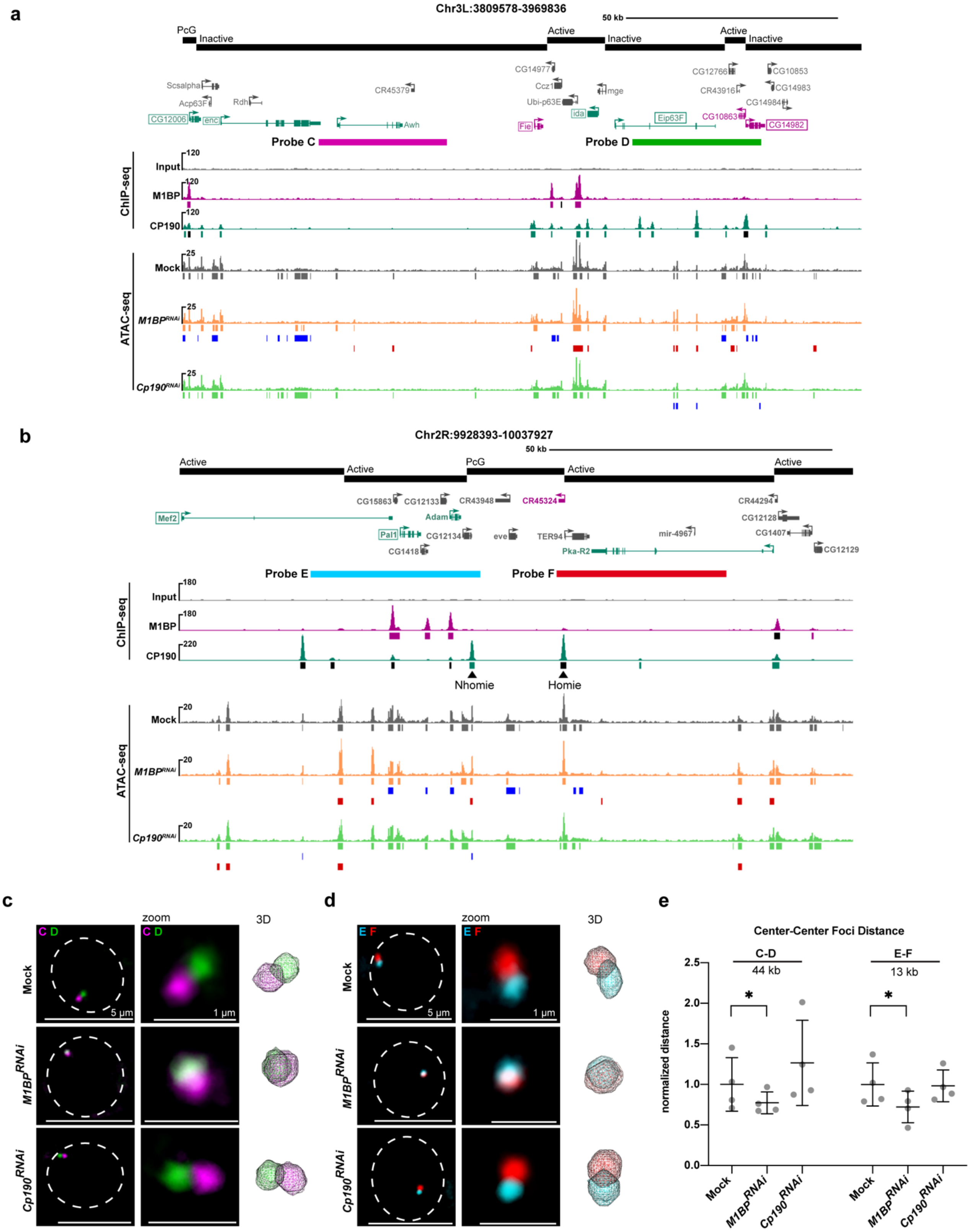

**Supplementary Figure 9. Related to Figure 8. Knockdown of *M1BP* increases local genome compaction.**

**a** Screenshot of regions detected by C and D probes spaced ~44 kb apart. TADs with state classification, longest isoform of genes, ChIP-seq signals of M1BP and CP190, and called peaks are shown. Upregulated genes in either *M1BP* or *Cp190* knockdown are shown in purple or green, respectively. Downregulated genes in either *M1BP* or *Cp190* knockdowns are outlined with a box and shown in purple or green, respectively. Called ChIP-seq peaks shown in black are significantly decreased in the knockdown of the opposite factor. ATAC-seq signals of mock, M1BP-depleted, and CP190-depleted cells, called peaks, decreased peaks relative to mock (blue bars), and increased peaks (red bars) are also shown. **b** Screenshot of regions detected by E and F probes spaced ~13 kb apart. **c**, **d** Left: Representative G1 nuclei labeled with probes shown in **a** and **b**, respectively. Single Z slices are shown. Dashed line represents nuclear edge. Center: Zoom of FISH signals. Right: 3D mesh rendering from TANGO. **e** Average center-to-center distance between probes C-D (left) and E-F (right). Single cell distances were normalized to nuclear radius before population averages were calculated. All averages were normalized to the average of mock controls. *C-D, $P$=0.05, *E-F, $P$=0.03, Paired t-test of means before normalization to controls. Data shown are from G1 cells from four biological replicates. n>100 cells were measured per replicate. Error bars show standard deviation.

#### Supplementary Figure 10

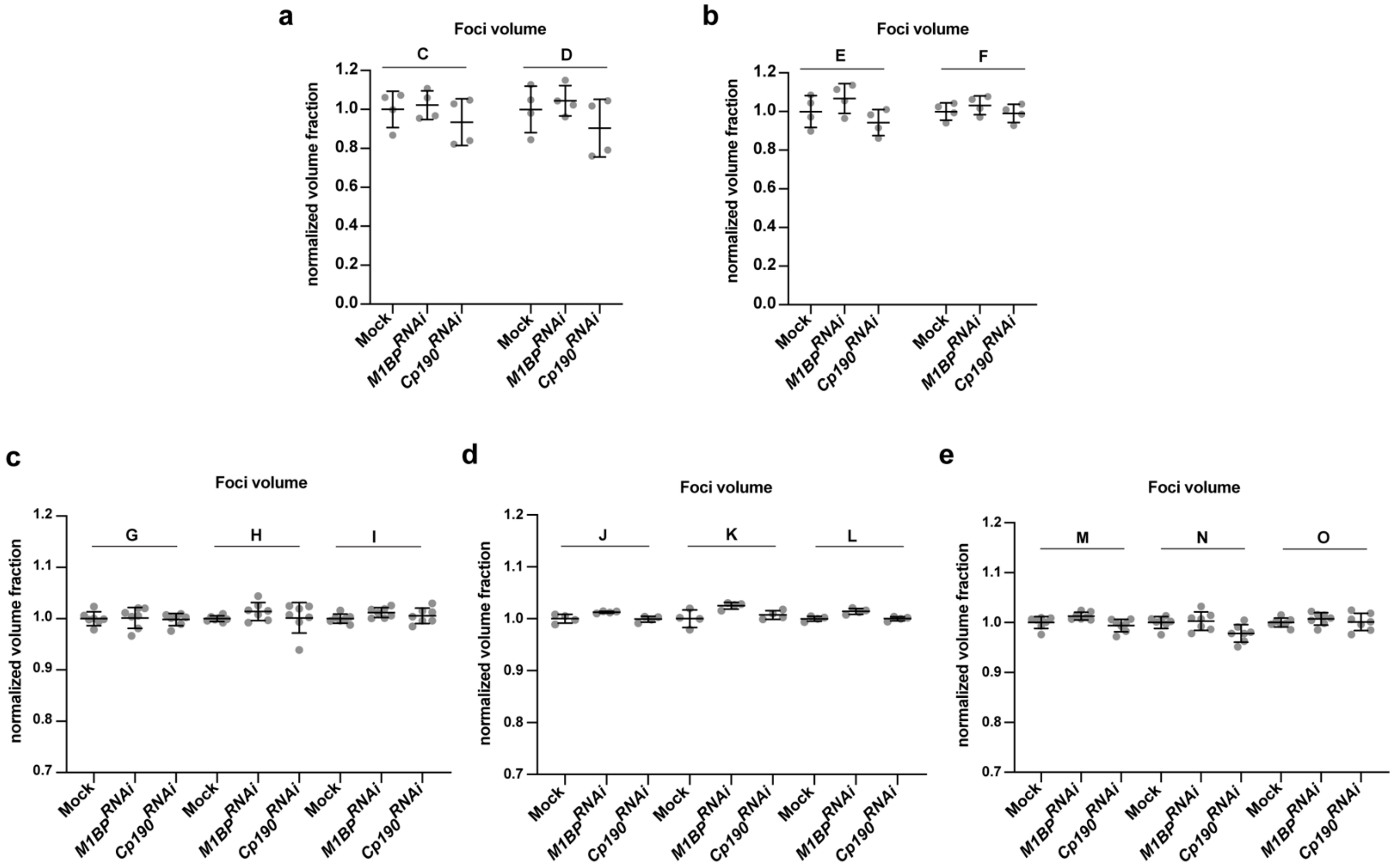

**Supplementary Figure 10. Related to Figure 8. Knockdowns of *M1BP* and *Cp190* do not alter foci volume.**

**a** Normalized foci volume shown as a fraction of nuclear volume of Probes C, D, **b** Probes E, F, **c** Probes G, H, I, **d** Probes J, K, L, **e** M, N and O. All averages were normalized to the average of their respective mock controls. Error bars show standard deviation. Unpaired t-test of means before normalization to controls for all samples, $P > 0.05$.

### Supplementary Figure 11

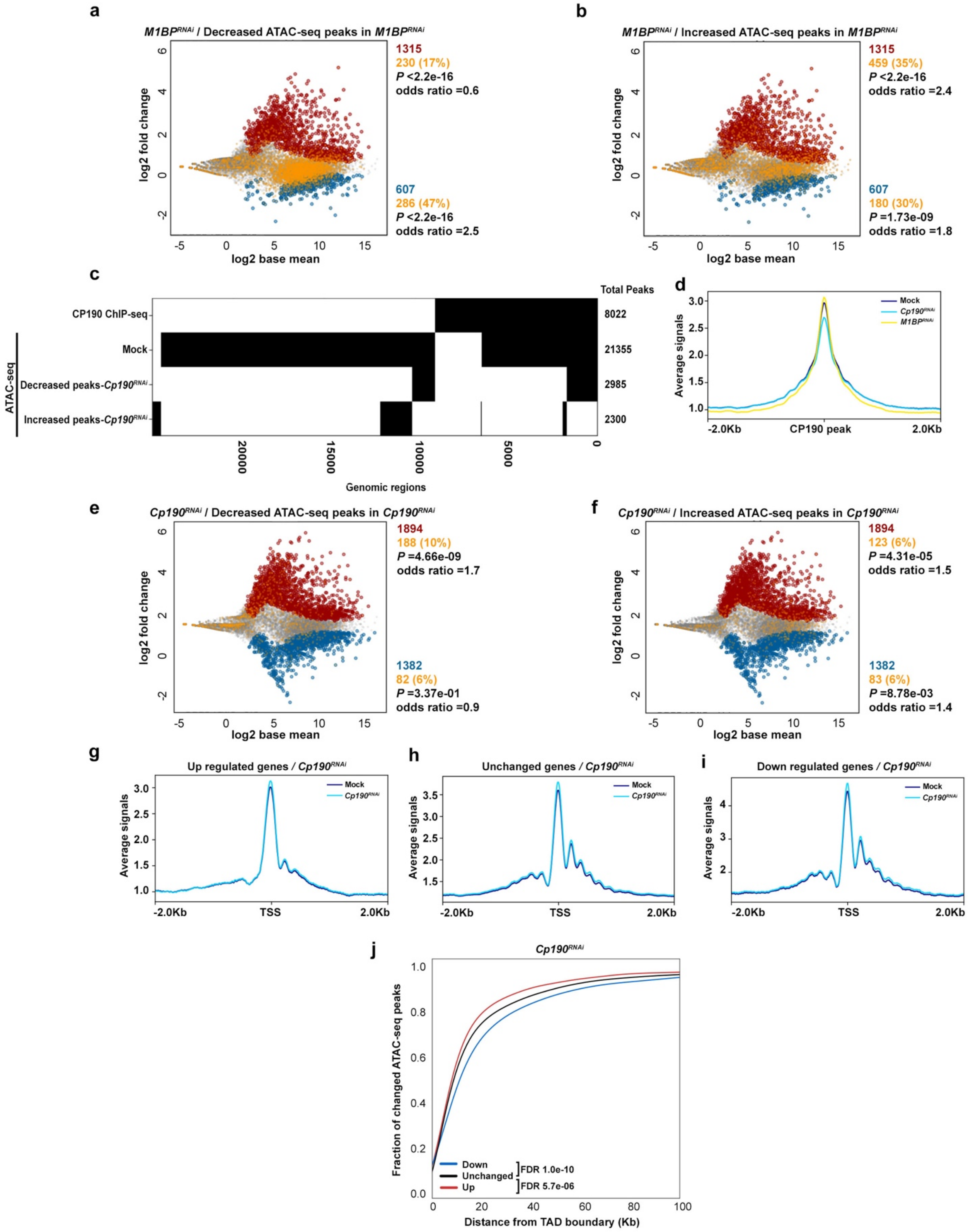

**Supplementary Figure 11. Related to Figure 9. *CP190* knockdown shows mild changes in chromatin accessibility genome-wide.**

**a**, **b** MA plots showing changes in neuRNA levels upon depletion of M1BP. Statistically significant changes include 1315 up-regulated genes (red) and 607 down-regulated genes (blue) using $p_{adj} < 0.05$. Unchanged genes are indicated in grey. Gene promoters containing decreased ATAC-seq peaks (**a**) or increased ATAC-seq peaks in depletion of M1BP (**b**) are additionally colored yellow. **c** Binary heatmap of CP190 ChIP-seq peaks, Mock ATAC-seq peaks, decreased and increased ATAC-seq peaks in *Cp190* knockdown ordered by supervised hierarchical clustering. Each row represents a single independent genomic location, and a black mark in a column represents the presence of a particular factor. **d** Average ATAC-seq signals of Mock, $M1BP^{RNAi}$ and $Cp190^{RNAi}$ from a 4 Kb genomic window centered on CP190 binding sites (ChIP-seq peaks). **e**, **f** MA plots showing changes in neuRNA levels upon depletion of CP190. Statistically significant changes include 1894 up-regulated genes (red) and 1382 down-regulated genes (blue) using $p_{adj} < 0.05$. Unchanged genes are indicated in grey. Gene promoters containing decreased ATAC-seq peaks (**e**) or increased ATAC-seq peaks in depletion of CP190 (**f**) are additionally colored yellow. **g**, **h**, **i** Average signal plots show enrichement of ATAC-seq signals of Mock, $M1BP^{RNAi}$ and $Cp190^{RNAi}$ from a 4 Kb genomic window at TSS of upregulated genes (g) Unchanged genes (h) Downregulated genes (i) in *Cp190* knockdown**. j** Cumulative histograms of $Cp190^{RNAi}$-ATAC-seq peak center distance from closest TAD border classified by change in nascent expression in *Cp190* knockdown cells. Downregulated (blue), upregulated (red), or unchanged (black) genes are indicated. Mann-Whitney U test for each set of changed genes against unchanged genes are shown.

#### Supplementary Table 1

| Ratio H/L Su(Hw) | Score | Intensity | Razor + unique peptides | Protein IDs | Protein names | Gene names |
|---|---|---|---|---|---|---|
| 759.84 | 6.936 | 1031700000 | 1 | E1JJK8;E1JJK9;E1JJK7;E1JJK6;M9PJM8;X2JF93;M9PEJ5;P43125-2;P43125 | Protein retinal degeneration B | rdgB |
| 226.74 | 323.31 | 35903000000 | 18 | A0A0B4K6W1;P08970 | Protein suppressor of hairy wing | su(Hw) |
| 186.37 | 190.53 | 1570700000 | 3 | B7YZR7;E1JGK5;Q9V895;Q9V895-2 | Acidic leucine-rich nuclear phosphoprotein 32 family member A | Mapmodulin;Anp32a |
| 166.26 | 254.18 | 27153000000 | 16 | Q95RQ8;Q9W1N1 | | pita |
| 95.362 | 148.09 | 2715000000 | 5 | Q9V3G3 | Peptidyl-prolyl cis-trans isomerase E | cyp33 |
| 92.033 | 323.31 | 2.65E+11 | 34 | Q24478 | Centrosome-associated zinc finger protein CP190 | Cp190 |
| 71.067 | 323.31 | 19026000000 | 9 | A0A0B4LGV6 | | Ibf1 |
| 70.513 | 183.59 | 9968600000 | 4 | Q9VHG6 | | Ibf2 |
| 68.41 | 61.294 | 160900000 | 2 | Q9NKV0-4;Q9NKV0 | Myeloid leukemia factor | Mlf |
| 60.171 | 323.31 | 24538000000 | 16 | Q86B87;Q86B87-22;Q86B87-26;A0A0B4JDE0 | Modifier of mdg4 | mod(mdg4) |
| 56.354 | 47.126 | 274800000 | 3 | Q9V3E9 | | spag |
| 55.631 | 213.95 | 2588700000 | 4 | Q9VMX3 | | |
| 52.474 | 51.673 | 847930000 | 3 | P38040 | Guanine nucleotide-binding protein subunit gamma-1 | Ggamma1 |
| 51.558 | 10.57 | 4746400000 | 3 | X2JIQ5;Q9W3W8 | 60S ribosomal protein L17 | RpL17 |
| 51.349 | 142.6 | 2793200000 | 5 | A1Z7P7;Q9XYM5;Q5BIG7 | | Su(var)2-10 |
| 47.735 | 323.31 | 14506000000 | 5 | Q94513 | | BEAF-32 |
| 47.512 | 323.31 | 1602500000 | 8 | Q9VVK8 | | |
| 47.453 | 1.7017 | 41188000 | 1 | M9PDV6;M9PE32 | | Fife |
| 46.133 | 323.31 | 2666900000 | 6 | Q24113;O76892 | | nonA-l |
| 44.666 | 124.87 | 557080000 | 2 | Q9VA00 | Protein zwilch | zwilch |
| 43.966 | 28.563 | 358740000 | 3 | Q9VSL0 | | |
| 43.742 | 4.6731 | 433770000 | 2 | Q9VWF8 | | |
| 43.222 | 73.623 | 1252500000 | 5 | Q9V3Z3 | | meso18E |
| 42.278 | 323.31 | 1138600000 | 8 | A0A0B4LHY1;Q8WR53;Q7KRS7;A8JRI1;Q95UI6;A0A0B4K6P4;A0A0B4K6W9;Q8IMF8;Q7KES3;E1JJ45 | | heph |
| 42.132 | 322.83 | 13874000000 | 11 | Q9VHM3 | | M1BP |
| 41.257 | 240.35 | 864600000 | 5 | Q7JW01;A0A0B4KFV4 | | |
| 41.03 | 323.31 | 4513500000 | 7 | Q9VS55 | | CTCF |
| 39.906 | 323.31 | 3067100000 | 7 | Q9V3D6 | Probable cleavage and polyadenylation | Cpsf100 |

| | | | | | | |
|---|---|---|---|---|---|---|
| | | | | | specificity factor subunit 2 | |
| 39.375 | 168.45 | 1758500000 | 5 | Q8IPA8;Q8IPA5 | | |
| 38.81 | 323.31 | 1510700000 | 7 | Q9VDY1 | Putative DNA helicase Ino80 | Ino80 |
| 37.517 | 291.82 | 1457400000 | 4 | A0A0B4KHL4;Q86BR9;Q86NK8;A0A0B4K7C4;A0A0B4K697;Q8IHB0;Q8INB6;Q4AB31;Q59DW8;Q9VEX2 | | gish |
| 37.27 | 168.97 | 1885800000 | 3 | E0R905;Q7KSP5;A0A0B4KGA3 | | l(3)neo38-RA;l(3)neo38 |
| 37.193 | 250.95 | 745160000 | 5 | Q9VQ58 | | CG15356-RA |
| 36.813 | 7.3316 | 566970000 | 3 | P20227 | TATA-box-binding protein | Tbp |
| 35.987 | 78.424 | 758850000 | 3 | M9PBX0;P42003 | Mothers against decapentaplegic homolog;Protein mothers against dpp | Mad |
| 35.88 | 48.547 | 1071300000 | 6 | Q9V3G7 | 26S proteasome non-ATPase regulatory subunit 6 | Rpn7 |
| 34.71 | 323.31 | 16889000000 | 18 | Q6NP69;Q8INS9 | | gfzf |
| 34.659 | 91.305 | 513110000 | 3 | Q9VPX5 | Vacuolar protein sorting-associated protein 29 | Vps29 |
| 34.145 | 230.49 | 409190000 | 3 | Q9W268;Q8MLT6 | | dve |
| 33.454 | 74.954 | 3987600000 | 4 | P07664 | Serendipity locus protein delta | Sry-delta |
| 33.121 | 323.31 | 17919000000 | 29 | Q9VZ00;X2JEU3 | | CG1737-RA |
| 33.043 | 323.31 | 2619700000 | 4 | Q7K4G8 | | |
| 32.819 | 323.31 | 4030300000 | 16 | E1JH59;A1Z9E2;A1Z9E2-2 | Protein lin-54 homolog | mip120 |
| 32.745 | 323.31 | 3565500000 | 17 | Q961B8 | | ADD1 |
| 32.497 | 250.7 | 1841000000 | 7 | Q9VVA6 | | nudC |
| 32.386 | 189.8 | 1239600000 | 4 | Q9VDQ3 | | |
| 32.223 | 194.78 | 2446500000 | 4 | Q9VFP0 | | |
| 32.128 | 323.31 | 48156000000 | 56 | M9PDH1;M9PE12;E1JHV6;Q9W0T1;Q9W0T1-2;M9PBG5 | Nucleosome-remodeling factor subunit NURF301 | E(bx) |
| 31.951 | 130.51 | 431070000 | 3 | M9PF62;P48423 | GTPase-activating protein | RasGAP1 |

**Supplementary Table 1.** Mass-spec with antibody against Su(Hw) using *Drosophila* embryonic nuclear extract. Presence of M1BP is highlighted in yellow.

#### Supplementary Table 2

| Ratio H/L cp190 | Score | Intensity | Razor + unique peptides | Protein IDs | Protein names | Gene names |
|---|---|---|---|---|---|---|
| 1370.2 | 323.31 | 4.514E+09 | 13 | Q9VS55 | | CTCF |
| 849.97 | 36.952 | 1.507E+09 | 5 | Q9W403 | | |
| 622.8 | 296.58 | 2.171E+09 | 31 | M9MS40;Q9W321;M9PH39 | | nej |
| 526.61 | 56.116 | 2.643E+09 | 4 | P12982;Q05547 | Serine/threonine-protein phosphatase alpha-2 isoform;Serine/threonine-protein phosphatase alpha-3 isoform | Pp1-87B;Pp1-13C |
| 523.24 | 323.31 | 1.451E+10 | 7 | Q94513 | | BEAF-32 |
| 477.82 | 118.7 | 466710000 | 3 | Q86B87-12 | Modifier of mdg4 | mod(mdg4) |
| 438.98 | 323.31 | 2.62E+09 | 9 | Q7K4G8 | | |
| 358.8 | 9.4308 | 70798000 | 1 | Q7JWH5 | | Roc2 |
| 342.74 | 74.954 | 3.988E+09 | 8 | P07664 | Serendipity locus protein delta | Sry-delta |
| 321.29 | 322.83 | 1.387E+10 | 14 | Q9VHM3 | | M1BP |
| 317.47 | 183.51 | 4.295E+09 | 6 | Q7K5M2;A0A0B4LFE1 | | |
| 298.11 | 11.881 | 9.867E+09 | 2 | P48588 | 40S ribosomal protein S25 | RpS25 |
| 237.09 | 203.37 | 1.079E+09 | 17 | Q9VKJ1 | MPN domain-containing protein CG4751 | CG4751 |
| 229.62 | 323.31 | 1.465E+09 | 12 | Q9VF00;A0A0B4JDD2 | | GATAe |
| 225.25 | 323.31 | 975430000 | 11 | Q7K2K2;A0A0C4DHG4 | | CG10265-RB |
| 216.59 | 323.31 | 9.912E+10 | 26 | Q9VP57 | | pzg |
| 213.05 | 323.31 | 2.649E+11 | 41 | Q24478 | Centrosome-associated zinc finger protein CP190 | Cp190 |
| 213.01 | 240.88 | 303420000 | 3 | Q9NDJ2-2 | Helicase domino | dom |
| 201.73 | 323.31 | 3.59E+10 | 20 | A0A0B4K6W1;P08970 | Protein suppressor of hairy wing | su(Hw) |
| 197.23 | 323.31 | 4.816E+10 | 65 | M9PDH1;M9PE12;E1JHV6;Q9W0T1;Q9W0T1-2;M9PBG5 | Nucleosome-remodeling factor subunit NURF301 | E(bx) |
| 196.61 | 237.15 | 2.041E+09 | 25 | Q9VUQ5-2;Q9VUQ5;M9PFK7 | Protein argonaute-2 | AGO2 |
| 173.41 | 124.71 | 2.052E+09 | 6 | Q867Z4 | Longitudinals lacking protein, isoforms F/I/K/T | lola |
| 170.98 | 215.43 | 3.55E+09 | 11 | Q7K4B2 | | |
| 170.3 | 323.31 | 6.167E+09 | 24 | M9PID3;M9PG82;M9PG20;Q9VPB0 | | HIPP1 |
| 169.98 | 323.31 | 3.389E+10 | 24 | Q86BS3 | | Chro |
| 167.96 | 323.31 | 2.454E+10 | 17 | Q86B87;Q86B87-22;Q86B87-26;A0A0B4JDE0 | Modifier of mdg4 | mod(mdg4) |
| 167.67 | 183.59 | 9.969E+09 | 7 | Q9VHG6 | | Ibf2 |
| 147.65 | 114.87 | 420980000 | 6 | Q9VK63 | | |

| | | | | | | |
|---|---|---|---|---|---|---|
| 146.68 | 323.31 | 1.689E+10 | 26 | Q6NP69;Q8INS9 | | gfzf |
| 145.58 | 18.381 | 117830000 | 7 | Q9VBA2 | | SIP2-RE |
| 145.13 | 54.312 | 968860000 | 3 | X2JKS9;Q9VXB3 | Histone deacetylase complex subunit SAP30 homolog | Sap30 |
| 143.66 | 81.503 | 1.029E+09 | 9 | Q9VR99 | Cactin | cactin |
| 139.69 | 252.05 | 2.901E+09 | 23 | Q9I7Z8 | | |
| 135.98 | 323.31 | 1.551E+09 | 10 | Q9W3N9 | | |
| 135.29 | 168.97 | 1.886E+09 | 6 | E0R905;Q7KSP5;A0A0B4KGA3 | | l(3)neo38-RA;l(3)neo38 |
| 131.93 | 323.31 | 619740000 | 14 | Q9VE54 | | |
| 130.24 | 156.34 | 802760000 | 10 | Q9V9N4;Q8IGP5 | | Jupiter;Clamp |
| 126.26 | 21.231 | 478820000 | 7 | Q9XYA7;Q8INR0;Q8INQ9 | | stck |
| 123.78 | 323.31 | 1.903E+10 | 11 | A0A0B4LGV6 | | Ibf1 |
| 122.29 | 323.31 | 4.03E+09 | 30 | E1JH59;A1Z9E2;A1Z9E2-2 | Protein lin-54 homolog | mip120 |
| 120.84 | 26.232 | 253560000 | 7 | Q8T921;Q9VRA1 | | |
| 113.05 | 185.8 | 2.189E+09 | 12 | Q7JWH6 | | |
| 113 | 170.43 | 417110000 | 5 | Q9W4V9 | | dwg |
| 111.91 | 66.405 | 88806000 | 4 | Q9VFX8 | | Ravus |
| 110.91 | 84.781 | 615980000 | 10 | P08630;M9PCX8;M9PCD8;P08630-2 | Tyrosine-protein kinase Btk29A;Non-specific protein-tyrosine kinase | Btk29A |
| 109.48 | 323.31 | 658580000 | 16 | X2JEY9;M9PHF1;X2JJP5;A8JUU1 | | Sec16 |
| 105.36 | 291.16 | 760310000 | 11 | Q7K4M4 | Protein teflon | tef |
| 103.69 | 323.31 | 4.119E+09 | 17 | Q9VRI0 | | Hlc |
| 103.13 | 145.88 | 815040000 | 10 | Q9VDS7;A0A0B4LID7 | | CG4538-RA |

**Supplementary Table 2.** Mass-spec with antibody against CP190 using *Drosophila* embryonic nuclear extract. Presence of M1BP is highlighted in yellow.

**Supplementary Table 3**

| Knockdown | *Act5C-Gal4* (ubiquitous) | *Mef2-Gal4* (muscle-specific) | *l(3)31-Gal4* (CNS-enriched) | *Ser-Gal4* (various tissues) |
|---|---|---|---|---|
| *M1BP*$^{RNAi}$ | Late larval lethality (n=150) | Pupal lethality (n=100) | Viable (n=100) | Viable (n=120) |

**Supplementary Table 3**. All viability or lethality levels were 100% unless otherwise indicated. n=total number of flies tested.

#### Supplementary Table 4

| ***Act5C-Gal4>*** | | | | | | |
|---|---|---|---|---|---|---|
| | ***non-ins; ctrl*** | ***non-ins; su(Hw)*** | ***non-ins; M1BP$^{RNAi}$*** | ***ins; ctrl*** | ***ins; su(Hw)*** | ***ins; M1BP$^{RNAi}$*** |
| ***non-ins; su(Hw)*** | 9.62E-01 | | | | | |
| ***non-ins; M1BP$^{RNAi}$*** | 9.74E-01 | 9.99E-01 | | | | |
| ***ins; ctrl*** | 1.46E-12 | 1.46E-12 | 1.46E-12 | | 1.46E-12 | 4.43E-05 |
| ***ins; su(Hw)*** | 1.55-E-02 | 1.12E-01 | 9.48E-02 | 1.46E-12 | | 1.60E-12 |
| ***ins; M1BP$^{RNAi}$*** | 1.46E-12 | 1.46E-12 | 1.46E-12 | 4.43E-05 | 1.60E-12 | |
| ***Mef2-Gal4>*** | | | | | | |
| | ***non-ins; ctrl*** | ***non-ins; su(Hw)*** | ***non-ins; M1BP$^{RNAi}$*** | ***ins; ctrl*** | ***ins; su(Hw)*** | ***ins; M1BP$^{RNAi}$*** |
| ***non-ins; su(Hw)*** | 8.99E-01 | | | | | |
| ***non-ins; M1BP$^{RNAi}$*** | 6.50E-01 | 9.97E-01 | | | | |
| ***ins; ctrl*** | 1.46E-12 | 1.46E-12 | 1.46E-12 | | 1.46E-12 | 2.71E-09 |
| ***ins; su(Hw)*** | 1.00E+00 | 9.01E-01 | 6.48E-01 | 1.46E-12 | | 1.50E-12 |
| ***ins; M1BP$^{RNAi}$*** | 1.51E-12 | 1.47E-12 | 1.46E-12 | 2.71E-09 | 1.50E-12 | |
| ***l(3)31-1-Gal4>*** | | | | | | |
| | ***non-ins; ctrl*** | ***non-ins; su(Hw)*** | ***non-ins; M1BP$^{RNAi}$*** | ***ins; ctrl*** | ***ins; su(Hw)*** | ***ins; M1BP$^{RNAi}$*** |
| ***non-ins; su(Hw)*** | 1.11E-03 | | | | | |
| ***non-ins; M1BP$^{RNAi}$*** | 2.58E-01 | 3.20E-01 | | | | |
| ***ins; ctrl*** | 1.46E-12 | 1.19E-11 | 1.49E-12 | | 1.46E-12 | 1.56E-02 |
| ***ins; su(Hw)*** | 9.87E-01 | 7.70E-05 | 5.55E-02 | 1.46E-12 | | 1.52E-12 |
| ***ins; M1BP$^{RNAi}$*** | 2.08E-12 | 8.14E-06 | 2.10E-02 | 1.56E-02 | 1.52E-12 | |

**Supplementary Table 4.** Luciferase assay *p*-values for *M1BP* knockdown and all other controls using *Act5C-Gal4*, *Mef2-Gal4* and *l(3)31-1-Gal4* in male larvae are indicated. *Gal4* driver lines are highlighted with orange. All genotypes of larvae are indicated with bold and italic.

**Supplementary Table 6**

| Knockdown | Forward primer | Reverse Primer |
|---|---|---|
| *M1BP*$^{RNAi}$ | TAATACGACTCACTATAGGGAAGTTA TTCGAGCGGAGCAA | TAATACGACTCACTATAGGGTAGAC GTACGCGTCGTTGAC |
| *Cp190*$^{RNAi}$ | TAATACGACTCACTATAGGGAGA GTAAACGGACGACCCATTAGCATTC | TAATACGACTCACTATAGGGAGA GTCTGCTCTGGTTCCTGCTC |
| *mod(mdg4)67.2*$^{RNAi}$ | TAATACGACTCACTATAGGGAGA AACAAAGGTTGAGGATCAGAC | TAATACGACTCACTATAGGGAGA TGGGGTCCACTTCTTCCTC |

**Supplementary Table 6.** List of primers for generation of dsRNA against *M1BP*, *Cp190* and *mod(mdg4).*

#### Supplementary Table 7

| | | | | *M1BP^RNAi* | *M1BP^RNAi* | *Cp190^RNAi* | *Cp190^RNAi* | | |
|---|---|---|---|---|---|---|---|---|---|
| **Site name** | **Chromosome site** | **M1BP peaks** | **CP190 peaks** | **M1BP decreased /lost in *M1BP^RNAi*** | **CP190 decreased /lost in *M1BP^RNAi*** | **M1BP decreased/ lost in *Cp190^RNAi*** | **CP190 decreased/ lost in *Cp190^RNAi*** | **Forward primer** | **Reverse primer** |
| 1 | chr2L:285520-286009 | Yes | Yes | Yes | Yes | Yes | Yes | GGGGAACTGCACGTAGTTGT | GGTGCCGTGAAAAATCAATC |
| 2 | chr2L:4955229-4955768 | Yes | Yes | Yes | Yes | Yes | Yes | TCCGAGATAGCGGAGAAGAA | CGTCGGTGATCGGTAGAAAT |
| 3 | chr3L:9360165-9360542 | Yes | Yes | Yes | Yes | Yes | Yes | CATGGGGGTGGTGATAAAAG | GCGGAGATTACCGAATTGAA |
| 4 | chr2L:8082778-8083790 | Yes | Yes | Yes | Yes | Yes | Yes | CTCGGGAACTCAAAGATTCG | GTTCTGGCCACACCTGAAAT |
| 5 | chrX:21339252-21340005 | Yes | Yes | Yes | Yes | Yes | Yes | ACCGTTAAAAGCCATGCAAA | TGGAAATCACAGAAGGCACA |
| 6 | chr2R:19304896 -19305345 | Yes | Yes | Yes | Yes | No | Yes | TACTCCGGTACTCGGTGCTC | TGTGCCTCCACATAGTTTCG |
| 7 | chr2R:20976497-20977464 | Yes | Yes | Yes | Yes | No | Yes | GGCACACGATCCAAAGTACC | TCACTTGCCTTTTCCTTGCT |
| 8 | chr2L:285520-286009 | Yes | Yes | Yes | Yes | No | Yes | GGGGAACTGCACGTAGTTGT | GTTTTGGTGCCGTGAAAAAT |
| 9 | chr3R:31745692-31746393 | Yes | Yes | Yes | Yes | No | Yes | TCAGGAGCTTCATGTGTTCG | CTCAAATTTCCGCTTTGGAA |
| 10 | chrX:17702652-17703572 | Yes | Yes | Yes | No | No | Yes | CGCTACAATTGGGGCATATT | TTCAGATGCGCTACATTTGC |
| 11 | chr2R:15537642-15538429 | Yes | Yes | Yes | No | No | Yes | ACCGAAGTGCGAGAAAGTGT | CAGAATGCAGAATGGCAGTG |
| 12 | chr2L:313096-314167 | No | Yes | No | No | No | Yes | AAATTACCAGCCGAATGCAC | GCCATAAATCGTCTGCCACT |
| *gypsy* | N/A | N/A | N/A | N/A | N/A | N/A | N/A | TCAAAAAATAAGTGCTGCATACTTTTTAG | AGCACAATTGATCGGCTA |
| *TART* | N/A | N/A | N/A | N/A | N/A | N/A | N/A | GTACAACCAAAGTTGACGGG | TGTATCGATATTTCGCGCTTTT |
| *Rint1* | chr3L:7322619-7324619 | Yes | Yes | Yes | Yes | No | Yes | TTGTGTGCTGCTCACCTTTC | GTGTGGCGTCGTTTTCTTTT |
| *CG7872* | chrX:15577205-15579205 | Yes | Yes | Yes | Yes | No | Yes | AATCGGTATTTGGCGAACTG | AACACCGATAACCGCGATAG |
| *CG6454* | chr3R:24348182-24350182 | Yes | Yes | Yes | Yes | No | Yes | GTTGTCATCCTTCAGCGACA | ACTCATTCCCTCCGGAGTTT |
| *tapas* | chr2R:20318180-20320180 | Yes | Yes | Yes | Yes | No | Yes | GCCCACGGTCTCACTAGAAG | TGGGGTTTTCCTGAGAAATG |
| *PDZ-GEF* | chr2L:6322767-6324767 | Yes | Yes | Yes | Yes | Yes | Yes | TAACCGATTCCCAGATCAGC | AGTCCGACGCGTCTTACAGT |

**Supplementary Table 7.** Differential binding of CP190 in *M1BP* knockdown and differential binding of M1BP in *Cp190* knockdown are verified by directed ChIP-qPCR and differential binding of Su(Hw), CP190, Mod(mdg4)67.2 in *M1BP* knockdown and in *Cp190* knockdown are tested by ChIP-qPCR.

**Supplementary Table 8**

| Antibody | Dilution for WB | Dilution for IF | Dilution for ChIP | Resources |
|---|---|---|---|---|
| Guinea pig anti-CP190 | 1:10000 | — | — | Matzat et al., 2012 |
| Rabbit anti-CP190 | — | 1:5000 | 1:333 | Pai et al., 2004 |
| Guinea pig anti-Su(Hw) | 1:5000 | — | 1:333 | Moshkovich and Lei, 2010 |
| Rabbit anti-Mod(mdg4)67.2 | 1:1000 | — | 1:333 | Van Bortle et al., 2014 |
| Rabbit anti-M1BP | 1:1000 | — | 1:200 | Li and Gilmour, 2013 |
| Rabbit anti-Pc | 1:8000 | — | — | Moshkovich et al., 2011 |
| Mouse anti-Pep | 1:1000 | — | — | Amero et al., 1991 |
| Rabbit anti-Histone H3(phospho S10) | — | 1:5000 | — | Abcam Cat No ab5176 |
| Mouse anti-Tubulin | | 1:500 | — | Sigma Cat No T6074 |

**Supplementary Table 8.** List of antibodies used for western blots, immunofluorescence and Chromatin immunoprecipitation.

**Supplementary Table 9**

| Probe | Chromosome | Start | Stop | Domain Size (bp) |
|---|---|---|---|---|
| A | Chr3L | 4204273 | 4254767 | 50494 |
| B | Chr3L | 7393235 | 7447744 | 54509 |
| C | Chr3L | 3841773 | 3872172 | 30399 |
| D | Chr3L | 3915948 | 3946359 | 30411 |
| E | Chr2R | 9944392 | 9974392 | 30000 |
| F | Chr2R | 9987751 | 10017741 | 29990 |
| G | Chr3L | 18906473 | 18938719 | 32246 |
| H | Chr3L | 18953719 | 18985965 | 32246 |
| I | Chr3L | 19000965 | 19033211 | 32246 |
| J | Chr3R | 13402320 | 13435028 | 32708 |
| K | Chr3R | 13450028 | 13482736 | 32708 |
| L | Chr3R | 13497736 | 13530444 | 32708 |
| M | Chr3R | 29636892 | 29669392 | 32500 |
| N | Chr3R | 29684392 | 29716892 | 32500 |
| O | Chr3R | 29731892 | 29764392 | 32500 |

**Supplementary Table 9.** List of probes used for FISH experiments.

**Supplementary Table 5**

| motif_id | motif_alt_id | sequence_name | start | stop | strand | score | p-value | q-value | matched_sequence |
|---|---|---|---|---|---|---|---|---|---|
| 1 | GMTACGGTCACACTG | chrX:18084761-18086761 | 1020 | 1034 | + | 28.743 | 6.44E-10 | 0.0204 | GCTACGGTCACACTG |
| 1 | GMTACGGTCACACTG | chr3R:24869816-24871816 | 1047 | 1061 | - | 23.236 | 2.86E-09 | 0.0204 | GGTACGGTCACACTG |
| 1 | GMTACGGTCACACTG | chr2L:4456674-4458674 | 137 | 151 | - | 22.536 | 1.31E-08 | 0.0204 | GCTACGGTCACACAG |
| 1 | GMTACGGTCACACTG | chr3R:14308089-14310089 | 486 | 500 | - | 22.536 | 1.31E-08 | 0.0204 | GCAACGGTCACACTG |
| 1 | GMTACGGTCACACTG | chr3L:10660475-10662475 | 538 | 552 | - | 22.536 | 1.31E-08 | 0.0204 | GCTACGGTCACTCTG |
| 1 | GMTACGGTCACACTG | chr3R:12428560-12430560 | 715 | 729 | - | 22.536 | 1.31E-08 | 0.0204 | GCTTCGGTCACACTG |
| 1 | GMTACGGTCACACTG | chrX:13128018-13130018 | 868 | 882 | - | 22.536 | 1.31E-08 | 0.0204 | GCTACGGTCACACCG |
| 1 | GMTACGGTCACACTG | chrX:13127952-13129952 | 934 | 948 | - | 22.536 | 1.31E-08 | 0.0204 | GCTACGGTCACACCG |
| 1 | GMTACGGTCACACTG | chr2R:20976056-20978056 | 985 | 999 | - | 22.536 | 1.31E-08 | 0.0204 | GCTGCGGTCACACTG |
| 1 | GMTACGGTCACACTG | chr3R:14307589-14309589 | 986 | 1000 | - | 22.536 | 1.31E-08 | 0.0204 | GCAACGGTCACACTG |
| 1 | GMTACGGTCACACTG | chr3R:12428282-12430282 | 993 | 1007 | - | 22.536 | 1.31E-08 | 0.0204 | GCTTCGGTCACACTG |
| 1 | GMTACGGTCACACTG | chr2L:4966775-4968775 | 998 | 1012 | - | 22.536 | 1.31E-08 | 0.0204 | GCCACGGTCACACTG |
| 1 | GMTACGGTCACACTG | chr3L:10660013-10662013 | 1000 | 1014 | - | 22.536 | 1.31E-08 | 0.0204 | GCTACGGTCACTCTG |
| 1 | GMTACGGTCACACTG | chr2L:4455807-4457807 | 1004 | 1018 | - | 22.536 | 1.31E-08 | 0.0204 | GCTACGGTCACACAG |
| 1 | GMTACGGTCACACTG | chrX:13127794-13129794 | 1092 | 1106 | - | 22.536 | 1.31E-08 | 0.0204 | GCTACGGTCACACCG |
| 1 | GMTACGGTCACACTG | chrX:13127630-13129630 | 1256 | 1270 | - | 22.536 | 1.31E-08 | 0.0204 | GCTACGGTCACACCG |
| 1 | GMTACGGTCACACTG | chr2L:1980176-1982176 | 612 | 626 | + | 22.536 | 1.31E-08 | 0.0204 | GCTGCGGTCACACTG |
| 1 | GMTACGGTCACACTG | chr3L:15967204-15969204 | 936 | 950 | + | 22.536 | 1.31E-08 | 0.0204 | GCTTCGGTCACACTG |
| 1 | GMTACGGTCACACTG | chrX:16609685-16611685 | 953 | 967 | + | 22.536 | 1.31E-08 | 0.0204 | GCAACGGTCACACTG |
| 1 | GMTACGGTCACACTG | chrX:15787897-15789897 | 956 | 970 | + | 22.536 | 1.31E-08 | 0.0204 | GCGACGGTCACACTG |
| 1 | GMTACGGTCACACTG | chr2R:14238932-14240932 | 964 | 978 | + | 22.536 | 1.31E-08 | 0.0204 | GCTTCGGTCACACTG |
| 1 | GMTACGGTCACACTG | chr3L:9072105-9074105 | 980 | 994 | + | 22.536 | 1.31E-08 | 0.0204 | GCTACGGCCACACTG |
| 1 | GMTACGGTCACACTG | chr2R:24132989-24134989 | 981 | 995 | + | 22.536 | 1.31E-08 | 0.0204 | GCTGCGGTCACACTG |
| 1 | GMTACGGTCACACTG | chr3R:20616883-20618883 | 989 | 1003 | + | 22.536 | 1.31E-08 | 0.0204 | GCGACGGTCACACTG |
| 1 | GMTACGGTCACACTG | chr2R:7011254-7013254 | 991 | 1005 | + | 22.536 | 1.31E-08 | 0.0204 | GCCACGGTCACACTG |
| 1 | GMTACGGTCACACTG | chr2L:2150754-2152754 | 998 | 1012 | + | 22.536 | 1.31E-08 | 0.0204 | GCGACGGTCACACTG |
| 1 | GMTACGGTCACACTG | chr2R:17456663-17458663 | 1000 | 1014 | + | 22.536 | 1.31E-08 | 0.0204 | GCCACGGTCACACTG |
| 1 | GMTACGGTCACACTG | chr2L:10975772-10977772 | 1007 | 1021 | + | 22.536 | 1.31E-08 | 0.0204 | GCTTCGGTCACACTG |
| 1 | GMTACGGTCACACTG | chr3L:13043434-13045434 | 1011 | 1025 | + | 22.536 | 1.31E-08 | 0.0204 | GCTCCGGTCACACTG |
| 1 | GMTACGGTCACACTG | chrX:11579772-11581772 | 1024 | 1038 | + | 22.536 | 1.31E-08 | 0.0204 | GCGACGGTCACACTG |

| | | | | | | | | | |
|---|---|---|---|---|---|---|---|---|---|
| 1 | GMTACGGTCACACTG | chr3R:5392922-5394922 | 1024 | 1038 | + | 22.536 | 1.31E-08 | 0.0204 | GCTGCGGTCACACTG |
| 1 | GMTACGGTCACACTG | chrX:16275270-16277270 | 1028 | 1042 | + | 22.536 | 1.31E-08 | 0.0204 | GCTGCGGTCACACTG |
| 1 | GMTACGGTCACACTG | chr2L:5799253-5801253 | 1032 | 1046 | + | 22.536 | 1.31E-08 | 0.0204 | GCTACGGCCACACTG |
| 1 | GMTACGGTCACACTG | chr2L:1979755-1981755 | 1033 | 1047 | + | 22.536 | 1.31E-08 | 0.0204 | GCTGCGGTCACACTG |
| 1 | GMTACGGTCACACTG | chrX:16275180-16277180 | 1118 | 1132 | + | 22.536 | 1.31E-08 | 0.0204 | GCTGCGGTCACACTG |
| 1 | GMTACGGTCACACTG | chr2R:14238703-14240703 | 1193 | 1207 | + | 22.536 | 1.31E-08 | 0.0204 | GCTTCGGTCACACTG |
| 1 | GMTACGGTCACACTG | chr3L:15966872-15968872 | 1268 | 1282 | + | 22.536 | 1.31E-08 | 0.0204 | GCTTCGGTCACACTG |
| 1 | GMTACGGTCACACTG | chr3L:9071731-9073731 | 1354 | 1368 | + | 22.536 | 1.31E-08 | 0.0204 | GCTACGGCCACACTG |
| 1 | GMTACGGTCACACTG | chr2L:10975399-10977399 | 1380 | 1394 | + | 22.536 | 1.31E-08 | 0.0204 | GCTTCGGTCACACTG |
| 1 | GMTACGGTCACACTG | chr2R:24132551-24134551 | 1419 | 1433 | + | 22.536 | 1.31E-08 | 0.0204 | GCTGCGGTCACACTG |
| 1 | GMTACGGTCACACTG | chr3R:30045470-30047470 | 1430 | 1444 | + | 22.536 | 1.31E-08 | 0.0204 | GCGACGGTCACACTG |
| 1 | GMTACGGTCACACTG | chr3L:20493921-20495921 | 1448 | 1462 | + | 22.536 | 1.31E-08 | 0.0204 | GCCACGGTCACACTG |
| 1 | GMTACGGTCACACTG | chr2R:7010739-7012739 | 1506 | 1520 | + | 22.536 | 1.31E-08 | 0.0204 | GCCACGGTCACACTG |
| 1 | GMTACGGTCACACTG | chr3R:30045161-30047161 | 1739 | 1753 | + | 22.536 | 1.31E-08 | 0.0204 | GCGACGGTCACACTG |
| 1 | GMTACGGTCACACTG | chr3R:20615901-20617901 | 1971 | 1985 | + | 22.536 | 1.31E-08 | 0.0204 | GCGACGGTCACACTG |
| 1 | GMTACGGTCACACTG | chrX:1030496-1032496 | 253 | 267 | + | 22.257 | 4.32E-08 | 0.0317 | GATACGGTCACACCG |
| 1 | GMTACGGTCACACTG | chr3L:257176-259176 | 818 | 832 | + | 22.257 | 4.32E-08 | 0.0317 | GATGCGGTCACACTG |
| 1 | GMTACGGTCACACTG | chr3L:16864295-16866295 | 959 | 973 | + | 22.257 | 4.32E-08 | 0.0317 | GCTACGTTCACACTG |
| 1 | GMTACGGTCACACTG | chr3R:29222485-29224485 | 968 | 982 | + | 22.257 | 4.32E-08 | 0.0317 | GCTAGGGTCACACTG |
| 1 | GMTACGGTCACACTG | chr3L:19909300-19911300 | 970 | 984 | + | 22.257 | 4.32E-08 | 0.0317 | GAAACGGTCACACTG |
| 1 | GMTACGGTCACACTG | chrX:10767537-10769537 | 984 | 998 | + | 22.257 | 4.32E-08 | 0.0317 | CCTACGGTCACACTG |
| 1 | GMTACGGTCACACTG | chrX:6371915-6373915 | 985 | 999 | + | 22.257 | 4.32E-08 | 0.0317 | CCTACGGTCACACTG |
| 1 | GMTACGGTCACACTG | chrX:1029758-1031758 | 991 | 1005 | + | 22.257 | 4.32E-08 | 0.0317 | GATACGGTCACACCG |
| 1 | GMTACGGTCACACTG | chr2R:18134426-18136426 | 992 | 1006 | + | 22.257 | 4.32E-08 | 0.0317 | GATGCGGTCACACTG |
| 1 | GMTACGGTCACACTG | chr3R:25880984-25882984 | 993 | 1007 | + | 22.257 | 4.32E-08 | 0.0317 | GAAACGGTCACACTG |
| 1 | GMTACGGTCACACTG | chr2R:10893459-10895459 | 1000 | 1014 | + | 22.257 | 4.32E-08 | 0.0317 | GACACGGTCACACTG |
| 1 | GMTACGGTCACACTG | chr2L:472013-474013 | 1001 | 1015 | + | 22.257 | 4.32E-08 | 0.0317 | GAAACGGTCACACTG |
| 1 | GMTACGGTCACACTG | chrX:18379082-18381082 | 1016 | 1030 | + | 22.257 | 4.32E-08 | 0.0317 | GATACGGTCACACCG |
| 1 | GMTACGGTCACACTG | chr3L:3070133-3072133 | 1021 | 1035 | + | 22.257 | 4.32E-08 | 0.0317 | GATGCGGTCACACTG |
| 1 | GMTACGGTCACACTG | chr3R:29222428-29224428 | 1025 | 1039 | + | 22.257 | 4.32E-08 | 0.0317 | GCTAGGGTCACACTG |
| 1 | GMTACGGTCACACTG | chr3L:256954-258954 | 1040 | 1054 | + | 22.257 | 4.32E-08 | 0.0317 | GATGCGGTCACACTG |
| 1 | GMTACGGTCACACTG | chr2R:10893414-10895414 | 1045 | 1059 | + | 22.257 | 4.32E-08 | 0.0317 | GACACGGTCACACTG |
| 1 | GMTACGGTCACACTG | chr3L:19909215-19911215 | 1055 | 1069 | + | 22.257 | 4.32E-08 | 0.0317 | GAAACGGTCACACTG |

| | | | | | | | | | |
|---|---|---|---|---|---|---|---|---|---|
| 1 | GMTACGGTCACACTG | chr3R:24529576-24531576 | 1075 | 1089 | + | 22.257 | 4.32E-08 | 0.0317 | TCTACGGTCACACTG |
| 1 | GMTACGGTCACACTG | chr3R:24529558-24531558 | 1093 | 1107 | + | 22.257 | 4.32E-08 | 0.0317 | TCTACGGTCACACTG |
| 1 | GMTACGGTCACACTG | chr3R:18241531-18243531 | 1175 | 1189 | + | 22.257 | 4.32E-08 | 0.0317 | GATACGGTCACACCG |
| 1 | GMTACGGTCACACTG | chrX:10767338-10769338 | 1183 | 1197 | + | 22.257 | 4.32E-08 | 0.0317 | CCTACGGTCACACTG |
| 1 | GMTACGGTCACACTG | chrX:1029556-1031556 | 1193 | 1207 | + | 22.257 | 4.32E-08 | 0.0317 | GATACGGTCACACCG |
| 1 | GMTACGGTCACACTG | chrX:18378854-18380854 | 1244 | 1258 | + | 22.257 | 4.32E-08 | 0.0317 | GATACGGTCACACCG |
| 1 | GMTACGGTCACACTG | chr3L:3069839-3071839 | 1315 | 1329 | + | 22.257 | 4.32E-08 | 0.0317 | GATGCGGTCACACTG |
| 1 | GMTACGGTCACACTG | chr3R:18241159-18243159 | 1547 | 1561 | + | 22.257 | 4.32E-08 | 0.0317 | GATACGGTCACACCG |
| 1 | GMTACGGTCACACTG | chr3R:25880427-25882427 | 1550 | 1564 | + | 22.257 | 4.32E-08 | 0.0317 | GAAACGGTCACACTG |
| 1 | GMTACGGTCACACTG | chr3R:28598834-28600834 | 1566 | 1580 | + | 22.257 | 4.32E-08 | 0.0317 | GAAACGGTCACACTG |
| 1 | GMTACGGTCACACTG | chr2R:18133634-18135634 | 1784 | 1798 | + | 22.257 | 4.32E-08 | 0.0317 | GATGCGGTCACACTG |
| 1 | GMTACGGTCACACTG | chr2L:471200-473200 | 1814 | 1828 | + | 22.257 | 4.32E-08 | 0.0317 | GAAACGGTCACACTG |
| 1 | GMTACGGTCACACTG | chr3L:19908410-19910410 | 1860 | 1874 | + | 22.257 | 4.32E-08 | 0.0317 | GAAACGGTCACACTG |
| 1 | GMTACGGTCACACTG | chr3L:19908365-19910365 | 1905 | 1919 | + | 22.257 | 4.32E-08 | 0.0317 | GAAACGGTCACACTG |
| 1 | GMTACGGTCACACTG | chr2R:17572462-17574462 | 327 | 341 | - | 22.257 | 4.32E-08 | 0.0317 | CCTACGGTCACACTG |
| 1 | GMTACGGTCACACTG | chr2R:23746560-23748560 | 401 | 415 | - | 22.257 | 4.32E-08 | 0.0317 | GAAACGGTCACACTG |
| 1 | GMTACGGTCACACTG | chr3L:9722423-9724423 | 737 | 751 | - | 22.257 | 4.32E-08 | 0.0317 | GCTAAGGTCACACTG |
| 1 | GMTACGGTCACACTG | chr2R:19436483-19438483 | 750 | 764 | - | 22.257 | 4.32E-08 | 0.0317 | GATTCGGTCACACTG |
| 1 | GMTACGGTCACACTG | chr3R:29488468-29490468 | 798 | 812 | - | 22.257 | 4.32E-08 | 0.0317 | GAAACGGTCACACTG |
| 1 | GMTACGGTCACACTG | chr3L:12749119-12751119 | 939 | 953 | - | 22.257 | 4.32E-08 | 0.0317 | GCTACGGTCACACTC |
| 1 | GMTACGGTCACACTG | chr3R:27694235-27696235 | 960 | 974 | - | 22.257 | 4.32E-08 | 0.0317 | GCTATGGTCACACTG |
| 1 | GMTACGGTCACACTG | chr2R:17571829-17573829 | 960 | 974 | - | 22.257 | 4.32E-08 | 0.0317 | CCTACGGTCACACTG |
| 1 | GMTACGGTCACACTG | chr3L:9722194-9724194 | 966 | 980 | - | 22.257 | 4.32E-08 | 0.0317 | GCTAAGGTCACACTG |
| 1 | GMTACGGTCACACTG | chr2R:19436259-19438259 | 974 | 988 | - | 22.257 | 4.32E-08 | 0.0317 | GATTCGGTCACACTG |
| 1 | GMTACGGTCACACTG | chr3R:6659100-6661100 | 976 | 990 | - | 22.257 | 4.32E-08 | 0.0317 | CCTACGGTCACACTG |
| 1 | GMTACGGTCACACTG | chrX:12591193-12593193 | 978 | 992 | - | 22.257 | 4.32E-08 | 0.0317 | CCTACGGTCACACTG |
| 1 | GMTACGGTCACACTG | chr3L:7653752-7655752 | 986 | 1000 | - | 22.257 | 4.32E-08 | 0.0317 | GACACGGTCACACTG |
| 1 | GMTACGGTCACACTG | chr3R:29488265-29490265 | 1001 | 1015 | - | 22.257 | 4.32E-08 | 0.0317 | GAAACGGTCACACTG |
| 1 | GMTACGGTCACACTG | chr3L:12749056-12751056 | 1002 | 1016 | - | 22.257 | 4.32E-08 | 0.0317 | GCTACGGTCACACTC |
| 1 | GMTACGGTCACACTG | chr2R:17571784-17573784 | 1005 | 1019 | - | 22.257 | 4.32E-08 | 0.0317 | CCTACGGTCACACTG |
| 1 | GMTACGGTCACACTG | chr2R:23745829-23747829 | 1132 | 1146 | - | 22.257 | 4.32E-08 | 0.0317 | GAAACGGTCACACTG |
| 1 | GMTACGGTCACACTG | chr3R:27693883-27695883 | 1312 | 1326 | - | 22.257 | 4.32E-08 | 0.0317 | GCTATGGTCACACTG |
| 1 | GMTACGGTCACACTG | chr3R:27693318-27695318 | 1877 | 1891 | - | 22.257 | 4.32E-08 | 0.0317 | GCTATGGTCACACTG |

| | | | | | | | | | |
|---|---|---|---|---|---|---|---|---|---|
| 1 | GMTACGGTCACACTG | chr2R:13484617-13486617 | 1002 | 1016 | + | 21.979 | 6.48E-08 | 0.0407 | TATACGGTCACACTG |
| 1 | GMTACGGTCACACTG | chrX:18090259-18092259 | 1005 | 1019 | + | 21.979 | 6.48E-08 | 0.0407 | CATACGGTCACACTG |
| 1 | GMTACGGTCACACTG | chr3R:24057287-24059287 | 1025 | 1039 | + | 21.979 | 6.48E-08 | 0.0407 | GATACGGTCATACTG |
| 1 | GMTACGGTCACACTG | chr3L:1328949-1330949 | 1028 | 1042 | + | 21.979 | 6.48E-08 | 0.0407 | CATACGGTCACACTG |
| 1 | GMTACGGTCACACTG | chr2R:13484391-13486391 | 1228 | 1242 | + | 21.979 | 6.48E-08 | 0.0407 | TATACGGTCACACTG |
| 1 | GMTACGGTCACACTG | chrX:18089509-18091509 | 1755 | 1769 | + | 21.979 | 6.48E-08 | 0.0407 | CATACGGTCACACTG |
| 1 | GMTACGGTCACACTG | chr2R:20286596-20288596 | 475 | 489 | - | 21.979 | 6.48E-08 | 0.0407 | CATACGGTCACACTG |
| 1 | GMTACGGTCACACTG | chr2R:21167930-21169930 | 775 | 789 | - | 21.979 | 6.48E-08 | 0.0407 | CATACGGTCACACTG |
| 1 | GMTACGGTCACACTG | chr3L:21135353-21137353 | 841 | 855 | - | 21.979 | 6.48E-08 | 0.0407 | GATACGGTCACACTT |
| 1 | GMTACGGTCACACTG | chr3R:20251770-20253770 | 889 | 903 | - | 21.979 | 6.48E-08 | 0.0407 | GATACGGTTACACTG |
| 1 | GMTACGGTCACACTG | chr2R:14255493-14257493 | 948 | 962 | - | 21.979 | 6.48E-08 | 0.0407 | GATACGGTCATACTG |
| 1 | GMTACGGTCACACTG | chr2R:20286094-20288094 | 977 | 991 | - | 21.979 | 6.48E-08 | 0.0407 | CATACGGTCACACTG |
| 1 | GMTACGGTCACACTG | chr3R:13675681-13677681 | 987 | 1001 | - | 21.979 | 6.48E-08 | 0.0407 | AATACGGTCACACTG |
| 1 | GMTACGGTCACACTG | chr2R:21167659-21169659 | 1046 | 1060 | - | 21.979 | 6.48E-08 | 0.0407 | CATACGGTCACACTG |
| 1 | GMTACGGTCACACTG | chr3R:13675549-13677549 | 1119 | 1133 | - | 21.979 | 6.48E-08 | 0.0407 | AATACGGTCACACTG |
| 1 | GMTACGGTCACACTG | chr2R:14255192-14257192 | 1249 | 1263 | - | 21.979 | 6.48E-08 | 0.0407 | GATACGGTCATACTG |
| 1 | GMTACGGTCACACTG | chr2R:8951339-8953339 | 991 | 1005 | + | 17.029 | 8.75E-08 | 0.0446 | GGTGCGGTCACACTG |
| 1 | GMTACGGTCACACTG | chr3L:12145519-12147519 | 999 | 1013 | + | 17.029 | 8.75E-08 | 0.0446 | GTCACGGTCACACTG |
| 1 | GMTACGGTCACACTG | chr2L:5999228-6001228 | 1004 | 1018 | + | 17.029 | 8.75E-08 | 0.0446 | GTCACGGTCACACTG |
| 1 | GMTACGGTCACACTG | chr2L:7781865-7783865 | 1012 | 1026 | + | 17.029 | 8.75E-08 | 0.0446 | GGAACGGTCACACTG |
| 1 | GMTACGGTCACACTG | chr3L:4039529-4041529 | 1027 | 1041 | + | 17.029 | 8.75E-08 | 0.0446 | GGTGCGGTCACACTG |
| 1 | GMTACGGTCACACTG | chrX:9255168-9257168 | 1028 | 1042 | + | 17.029 | 8.75E-08 | 0.0446 | GGTGCGGTCACACTG |
| 1 | GMTACGGTCACACTG | chr3L:4039482-4041482 | 1074 | 1088 | + | 17.029 | 8.75E-08 | 0.0446 | GGTGCGGTCACACTG |
| 1 | GMTACGGTCACACTG | chr3L:12145097-12147097 | 1421 | 1435 | + | 17.029 | 8.75E-08 | 0.0446 | GTCACGGTCACACTG |
| 1 | GMTACGGTCACACTG | chr2L:5998501-6000501 | 1731 | 1745 | + | 17.029 | 8.75E-08 | 0.0446 | GTCACGGTCACACTG |
| 1 | GMTACGGTCACACTG | chr2R:8950595-8952595 | 1735 | 1749 | + | 17.029 | 8.75E-08 | 0.0446 | GGTGCGGTCACACTG |
| 1 | GMTACGGTCACACTG | chr3R:30036730-30038730 | 103 | 117 | - | 17.029 | 8.75E-08 | 0.0446 | GGCACGGTCACACTG |
| 1 | GMTACGGTCACACTG | chr2R:5652676-5654676 | 127 | 141 | - | 17.029 | 8.75E-08 | 0.0446 | GTTTCGGTCACACTG |
| 1 | GMTACGGTCACACTG | chr2R:16097948-16099948 | 596 | 610 | - | 17.029 | 8.75E-08 | 0.0446 | GTTGCGGTCACACTG |
| 1 | GMTACGGTCACACTG | chr3R:25264844-25266844 | 870 | 884 | - | 17.029 | 8.75E-08 | 0.0446 | GGAACGGTCACACTG |
| 1 | GMTACGGTCACACTG | chrX:13562823-13564823 | 883 | 897 | - | 17.029 | 8.75E-08 | 0.0446 | GGAACGGTCACACTG |
| 1 | GMTACGGTCACACTG | chrX:13562823-13564823 | 883 | 897 | - | 17.029 | 8.75E-08 | 0.0446 | GGAACGGTCACACTG |
| 1 | GMTACGGTCACACTG | chrX:6290303-6292303 | 914 | 928 | - | 17.029 | 8.75E-08 | 0.0446 | GTTACGGGCACACTG |

| | | | | | | | | | |
|---|---|---|---|---|---|---|---|---|---|
| 1 | GMTACGGTCACACTG | chr3R:8715243-8717243 | 942 | 956 | - | 17.029 | 8.75E-08 | 0.0446 | GTTCCGGTCACACTG |
| 1 | GMTACGGTCACACTG | chr3R:30035870-30037870 | 963 | 977 | - | 17.029 | 8.75E-08 | 0.0446 | GGCACGGTCACACTG |
| 1 | GMTACGGTCACACTG | chrX:6290245-6292245 | 972 | 986 | - | 17.029 | 8.75E-08 | 0.0446 | GTTACGGGCACACTG |
| 1 | GMTACGGTCACACTG | chr2R:16097564-16099564 | 980 | 994 | - | 17.029 | 8.75E-08 | 0.0446 | GTTGCGGTCACACTG |
| 1 | GMTACGGTCACACTG | chr2R:5651815-5653815 | 988 | 1002 | - | 17.029 | 8.75E-08 | 0.0446 | GTTTCGGTCACACTG |
| 1 | GMTACGGTCACACTG | chr3L:12398322-12400322 | 993 | 1007 | - | 17.029 | 8.75E-08 | 0.0446 | GGTACGGCCACACTG |
| 1 | GMTACGGTCACACTG | chr3R:25264721-25266721 | 993 | 1007 | - | 17.029 | 8.75E-08 | 0.0446 | GGAACGGTCACACTG |
| 1 | GMTACGGTCACACTG | chrX:19565019-19567019 | 1087 | 1101 | - | 17.029 | 8.75E-08 | 0.0446 | GGCACGGTCACACTG |
| 1 | GMTACGGTCACACTG | chrX:19564764-19566764 | 1342 | 1356 | - | 17.029 | 8.75E-08 | 0.0446 | GGCACGGTCACACTG |
| 1 | GMTACGGTCACACTG | chrX:15709647-15711647 | 983 | 997 | + | 16.75 | 1.27E-07 | 0.0541 | GTTAAGGTCACACTG |
| 1 | GMTACGGTCACACTG | chr3R:20830127-20832127 | 992 | 1006 | + | 16.75 | 1.27E-07 | 0.0541 | GTTACGGTCACACTA |
| 1 | GMTACGGTCACACTG | chr3L:8985710-8987710 | 1021 | 1035 | + | 16.75 | 1.27E-07 | 0.0541 | ATTACGGTCACACTG |
| 1 | GMTACGGTCACACTG | chrX:15709539-15711539 | 1091 | 1105 | + | 16.75 | 1.27E-07 | 0.0541 | GTTAAGGTCACACTG |
| 1 | GMTACGGTCACACTG | chr3R:19026368-19028368 | 1320 | 1334 | + | 16.75 | 1.27E-07 | 0.0541 | GTTACGGTGACACTG |
| 1 | GMTACGGTCACACTG | chr3R:20829580-20831580 | 1539 | 1553 | + | 16.75 | 1.27E-07 | 0.0541 | GTTACGGTCACACTA |
| 1 | GMTACGGTCACACTG | chr3R:4774053-4776053 | 1540 | 1554 | + | 16.75 | 1.27E-07 | 0.0541 | AGTACGGTCACACTG |
| 1 | GMTACGGTCACACTG | chr3L:8985091-8987091 | 1640 | 1654 | + | 16.75 | 1.27E-07 | 0.0541 | ATTACGGTCACACTG |
| 1 | GMTACGGTCACACTG | chr3R:31219702-31221702 | 85 | 99 | - | 16.75 | 1.27E-07 | 0.0541 | GTTATGGTCACACTG |
| 1 | GMTACGGTCACACTG | chr3L:21432422-21434422 | 674 | 688 | - | 16.75 | 1.27E-07 | 0.0541 | AGTACGGTCACACTG |
| 1 | GMTACGGTCACACTG | chr3L:2261919-2263919 | 708 | 722 | - | 16.75 | 1.27E-07 | 0.0541 | AGTACGGTCACACTG |
| 1 | GMTACGGTCACACTG | chr3R:5367481-5369481 | 801 | 815 | - | 16.75 | 1.27E-07 | 0.0541 | TGTACGGTCACACTG |
| 1 | GMTACGGTCACACTG | chr4:1059969-1061969 | 912 | 926 | - | 16.75 | 1.27E-07 | 0.0541 | GTTAAGGTCACACTG |
| 1 | GMTACGGTCACACTG | chr3L:5751973-5753973 | 941 | 955 | - | 16.75 | 1.27E-07 | 0.0541 | CTTACGGTCACACTG |
| 1 | GMTACGGTCACACTG | chr3L:13441594-13443594 | 957 | 971 | - | 16.75 | 1.27E-07 | 0.0541 | GTTATGGTCACACTG |
| 1 | GMTACGGTCACACTG | chr3L:21432134-21434134 | 962 | 976 | - | 16.75 | 1.27E-07 | 0.0541 | AGTACGGTCACACTG |
| 1 | GMTACGGTCACACTG | chr3L:2261664-2263664 | 963 | 977 | - | 16.75 | 1.27E-07 | 0.0541 | AGTACGGTCACACTG |
| 1 | GMTACGGTCACACTG | chr3L:19099363-19101363 | 967 | 981 | - | 16.75 | 1.27E-07 | 0.0541 | CGTACGGTCACACTG |
| 1 | GMTACGGTCACACTG | chr2L:2144187-2146187 | 971 | 985 | - | 16.75 | 1.27E-07 | 0.0541 | AGTACGGTCACACTG |
| 1 | GMTACGGTCACACTG | chr3R:5367306-5369306 | 976 | 990 | - | 16.75 | 1.27E-07 | 0.0541 | TGTACGGTCACACTG |
| 1 | GMTACGGTCACACTG | chr3R:25044151-25046151 | 983 | 997 | - | 16.75 | 1.27E-07 | 0.0541 | GGTATGGTCACACTG |
| 1 | GMTACGGTCACACTG | chr2L:13020312-13022312 | 987 | 1001 | - | 16.75 | 1.27E-07 | 0.0541 | GGTACGGTAACACTG |
| 1 | GMTACGGTCACACTG | chr3L:6755497-6757497 | 1010 | 1024 | - | 16.75 | 1.27E-07 | 0.0541 | TGTACGGTCACACTG |
| 1 | GMTACGGTCACACTG | chr3L:16590050-16592050 | 1029 | 1043 | - | 16.75 | 1.27E-07 | 0.0541 | AGTACGGTCACACTG |

| | | | | | | | | | |
|---|---|---|---|---|---|---|---|---|---|
| 1 | GMTACGGTCACACTG | chr3R:31218630-31220630 | 1157 | 1171 | - | 16.75 | 1.27E-07 | 0.0541 | GTTATGGTCACACTG |
| 1 | GMTACGGTCACACTG | chr3R:31218372-31220372 | 1415 | 1429 | - | 16.75 | 1.27E-07 | 0.0541 | GTTATGGTCACACTG |
| 1 | GMTACGGTCACACTG | chr3L:5751189-5753189 | 1725 | 1739 | - | 16.75 | 1.27E-07 | 0.0541 | CTTACGGTCACACTG |
| 1 | GMTACGGTCACACTG | chr3R:27947711-27949711 | 293 | 307 | - | 16.329 | 1.94E-07 | 0.0654 | GCTGCGGTCACTCTG |
| 1 | GMTACGGTCACACTG | chr3L:17432868-17434868 | 314 | 328 | - | 16.329 | 1.94E-07 | 0.0654 | GCTGCGGCCACACTG |
| 1 | GMTACGGTCACACTG | chr2L:10262554-10264554 | 321 | 335 | - | 16.329 | 1.94E-07 | 0.0654 | GCTACGGACGCACTG |
| 1 | GMTACGGTCACACTG | chrX:11453786-11455786 | 552 | 566 | - | 16.329 | 1.94E-07 | 0.0654 | GCGACGGTCACACCG |
| 1 | GMTACGGTCACACTG | chr3L:17432621-17434621 | 561 | 575 | - | 16.329 | 1.94E-07 | 0.0654 | GCTGCGGCCACACTG |
| 1 | GMTACGGTCACACTG | chr3R:18298229-18300229 | 639 | 653 | - | 16.329 | 1.94E-07 | 0.0654 | GCCACGGTCACTCTG |
| 1 | GMTACGGTCACACTG | chr2R:17589964-17591964 | 696 | 710 | - | 16.329 | 1.94E-07 | 0.0654 | GCCGCGGTCACACTG |
| 1 | GMTACGGTCACACTG | chr3L:1301944-1303944 | 717 | 731 | - | 16.329 | 1.94E-07 | 0.0654 | GCCGCGGTCACACTG |
| 1 | GMTACGGTCACACTG | chr2R:17589884-17591884 | 776 | 790 | - | 16.329 | 1.94E-07 | 0.0654 | GCCGCGGTCACACTG |
| 1 | GMTACGGTCACACTG | chr2R:23889512-23891512 | 802 | 816 | + | 16.329 | 1.94E-07 | 0.0654 | GCTGCGGTCACACCG |
| 1 | GMTACGGTCACACTG | chr3R:17681669-17683669 | 825 | 839 | - | 16.329 | 1.94E-07 | 0.0654 | GCCACGGTCACTCTG |
| 1 | GMTACGGTCACACTG | chrX:2584980-2586980 | 964 | 978 | + | 16.329 | 1.94E-07 | 0.0654 | GCAACGGCCACACTG |
| 1 | GMTACGGTCACACTG | chrX:11453386-11455386 | 952 | 966 | - | 16.329 | 1.94E-07 | 0.0654 | GCGACGGTCACACCG |
| 1 | GMTACGGTCACACTG | chr3L:1575699-1577699 | 967 | 981 | + | 16.329 | 1.94E-07 | 0.0654 | GCTGCGGCCACACTG |
| 1 | GMTACGGTCACACTG | chr2R:14215627-14217627 | 954 | 968 | - | 16.329 | 1.94E-07 | 0.0654 | GCAACGGTCACTCTG |
| 1 | GMTACGGTCACACTG | chr2L:20655426-20657426 | 973 | 987 | + | 16.329 | 1.94E-07 | 0.0654 | GCTCCGGCCACACTG |
| 1 | GMTACGGTCACACTG | chr2R:13501841-13503841 | 962 | 976 | - | 16.329 | 1.94E-07 | 0.0654 | GCCACGGTCACGCTG |
| 1 | GMTACGGTCACACTG | chr3L:9401460-9403460 | 968 | 982 | - | 16.329 | 1.94E-07 | 0.0654 | GCAACGGCCACACTG |
| 1 | GMTACGGTCACACTG | chr2L:10203317-10205317 | 968 | 982 | - | 16.329 | 1.94E-07 | 0.0654 | GCCGCGGTCACACTG |
| 1 | GMTACGGTCACACTG | chr2R:14980263-14982263 | 992 | 1006 | + | 16.329 | 1.94E-07 | 0.0654 | GCTGCGGTCACACCG |
| 1 | GMTACGGTCACACTG | chr3L:1301683-1303683 | 978 | 992 | - | 16.329 | 1.94E-07 | 0.0654 | GCCGCGGTCACACTG |
| 1 | GMTACGGTCACACTG | chr3R:17681515-17683515 | 979 | 993 | - | 16.329 | 1.94E-07 | 0.0654 | GCCACGGTCACTCTG |
| 1 | GMTACGGTCACACTG | chr2R:8079041-8081041 | 979 | 993 | - | 16.329 | 1.94E-07 | 0.0654 | GCAACGGCCACACTG |
| 1 | GMTACGGTCACACTG | chr2R:6806626-6808626 | 996 | 1010 | + | 16.329 | 1.94E-07 | 0.0654 | GCCACGGCCACACTG |
| 1 | GMTACGGTCACACTG | chr3R:27947021-27949021 | 983 | 997 | - | 16.329 | 1.94E-07 | 0.0654 | GCTGCGGTCACTCTG |
| 1 | GMTACGGTCACACTG | chrX:646304-648304 | 997 | 1011 | + | 16.329 | 1.94E-07 | 0.0654 | GCGGCGGTCACACTG |
| 1 | GMTACGGTCACACTG | chr3R:18297879-18299879 | 989 | 1003 | - | 16.329 | 1.94E-07 | 0.0654 | GCCACGGTCACTCTG |
| 1 | GMTACGGTCACACTG | chr3L:17432190-17434190 | 992 | 1006 | - | 16.329 | 1.94E-07 | 0.0654 | GCTGCGGCCACACTG |
| 1 | GMTACGGTCACACTG | chr3L:8967528-8969528 | 1007 | 1021 | + | 16.329 | 1.94E-07 | 0.0654 | GCGACGGGCACACTG |
| 1 | GMTACGGTCACACTG | chrX:16559016-16561016 | 995 | 1009 | - | 16.329 | 1.94E-07 | 0.0654 | GCCACGGCCACACTG |

| | | | | | | | | | |
|---|---|---|---|---|---|---|---|---|---|
| 1 | GMTACGGTCACACTG | chr3L:19585253-19587253 | 1012 | 1026 | + | 16.329 | 1.94E-07 | 0.0654 | GCAACGGTCACTCTG |
| 1 | GMTACGGTCACACTG | chr2R:14215627-14217627 | 998 | 1012 | - | 16.329 | 1.94E-07 | 0.0654 | GCGGCGGTCACACTG |
| 1 | GMTACGGTCACACTG | chr2L:2364345-2366345 | 1021 | 1035 | + | 16.329 | 1.94E-07 | 0.0654 | GCGACGGGCACACTG |
| 1 | GMTACGGTCACACTG | chr2R:23889276-23891276 | 1038 | 1052 | + | 16.329 | 1.94E-07 | 0.0654 | GCTGCGGTCACACCG |
| 1 | GMTACGGTCACACTG | chr2R:7406819-7408819 | 1040 | 1054 | - | 16.329 | 1.94E-07 | 0.0654 | GCAACGGTCACTCTG |
| 1 | GMTACGGTCACACTG | chr2L:2364273-2366273 | 1093 | 1107 | + | 16.329 | 1.94E-07 | 0.0654 | GCGACGGGCACACTG |
| 1 | GMTACGGTCACACTG | chrX:15319037-15321037 | 1130 | 1144 | - | 16.329 | 1.94E-07 | 0.0654 | GCTGCGGCCACACTG |
| 1 | GMTACGGTCACACTG | chr2L:20655178-20657178 | 1221 | 1235 | + | 16.329 | 1.94E-07 | 0.0654 | GCTCCGGCCACACTG |
| 1 | GMTACGGTCACACTG | chrX:2584699-2586699 | 1245 | 1259 | + | 16.329 | 1.94E-07 | 0.0654 | GCAACGGCCACACTG |
| 1 | GMTACGGTCACACTG | chr3L:19585003-19587003 | 1262 | 1276 | + | 16.329 | 1.94E-07 | 0.0654 | GCAACGGTCACTCTG |
| 1 | GMTACGGTCACACTG | chrX:8587811-8589811 | 1572 | 1586 | + | 16.329 | 1.94E-07 | 0.0654 | GCTACGGACACACGG |
| 1 | GMTACGGTCACACTG | chrX:15318479-15320479 | 1688 | 1702 | - | 16.329 | 1.94E-07 | 0.0654 | GCTGCGGCCACACTG |
| 1 | GMTACGGTCACACTG | chr2R:14214864-14216864 | 1717 | 1731 | - | 16.329 | 1.94E-07 | 0.0654 | GCAACGGTCACTCTG |
| 1 | GMTACGGTCACACTG | chr2R:14214864-14216864 | 1761 | 1775 | - | 16.329 | 1.94E-07 | 0.0654 | GCGGCGGTCACACTG |
| 1 | GMTACGGTCACACTG | chr2L:21262933-21264933 | 25 | 39 | + | 16.05 | 5.57E-07 | 0.102 | ACTGCGGTCACACTG |
| 1 | GMTACGGTCACACTG | chrX:2161196-2163196 | 14 | 28 | - | 16.05 | 5.57E-07 | 0.102 | TCTACGGACACACTG |
| 1 | GMTACGGTCACACTG | chr3L:21533660-21535660 | 26 | 40 | - | 16.05 | 5.57E-07 | 0.102 | GCTTTGGTCACACTG |
| 1 | GMTACGGTCACACTG | chrX:2161180-2163180 | 30 | 44 | - | 16.05 | 5.57E-07 | 0.102 | TCTACGGACACACTG |
| 1 | GMTACGGTCACACTG | chr2L:277234-279234 | 57 | 71 | - | 16.05 | 5.57E-07 | 0.102 | TCGACGGTCACACTG |
| 1 | GMTACGGTCACACTG | chrX:2009300-2011300 | 78 | 92 | - | 16.05 | 5.57E-07 | 0.102 | TCGACGGTCACACTG |
| 1 | GMTACGGTCACACTG | chr3R:12691597-12693597 | 161 | 175 | + | 16.05 | 5.57E-07 | 0.102 | CCTACGGTCACCCTG |
| 1 | GMTACGGTCACACTG | chr3L:16112212-16114212 | 168 | 182 | - | 16.05 | 5.57E-07 | 0.102 | GCTGCGGTCACACTA |
| 1 | GMTACGGTCACACTG | chr3R:8778923-8780923 | 212 | 226 | + | 16.05 | 5.57E-07 | 0.102 | GCAACGGTTACACTG |
| 1 | GMTACGGTCACACTG | chr3L:19519945-19521945 | 255 | 269 | + | 16.05 | 5.57E-07 | 0.102 | GCTACGGTCACGATG |
| 1 | GMTACGGTCACACTG | chr3L:19260394-19262394 | 278 | 292 | - | 16.05 | 5.57E-07 | 0.102 | CCTACGGCCACACTG |
| 1 | GMTACGGTCACACTG | chr3R:21101889-21103889 | 318 | 332 | + | 16.05 | 5.57E-07 | 0.102 | GCCATGGTCACACTG |
| 1 | GMTACGGTCACACTG | chr2R:16997594-16999594 | 309 | 323 | - | 16.05 | 5.57E-07 | 0.102 | GCTACGTTCACCCTG |
| 1 | GMTACGGTCACACTG | chr2R:16997594-16999594 | 309 | 323 | - | 16.05 | 5.57E-07 | 0.102 | GCTACGTTCACCCTG |
| 1 | GMTACGGTCACACTG | chr3R:16453492-16455492 | 320 | 334 | - | 16.05 | 5.57E-07 | 0.102 | GCTGCGGTCACACTT |
| 1 | GMTACGGTCACACTG | chrX:16586873-16588873 | 364 | 378 | - | 16.05 | 5.57E-07 | 0.102 | GAAACGGTCACACCG |
| 1 | GMTACGGTCACACTG | chrX:15319037-15321037 | 390 | 404 | + | 16.05 | 5.57E-07 | 0.102 | CCGACGGTCACACTG |
| 1 | GMTACGGTCACACTG | chrX:2008957-2010957 | 421 | 435 | - | 16.05 | 5.57E-07 | 0.102 | TCGACGGTCACACTG |
| 1 | GMTACGGTCACACTG | chr3L:7911463-7913463 | 456 | 470 | - | 16.05 | 5.57E-07 | 0.102 | GCGACGGTTACACTG |

| | | | | | | | | | |
|---|---|---|---|---|---|---|---|---|---|
| 1 | GMTACGGTCACACTG | chr3L:18800312-18802312 | 497 | 511 | + | 16.05 | 5.57E-07 | 0.102 | GCTACGCACACACTG |
| 1 | GMTACGGTCACACTG | chr3L:18800309-18802309 | 500 | 514 | + | 16.05 | 5.57E-07 | 0.102 | GCTACGCACACACTG |
| 1 | GMTACGGTCACACTG | chr2R:16867222-16869222 | 512 | 526 | + | 16.05 | 5.57E-07 | 0.102 | GCAACGGTAACACTG |
| 1 | GMTACGGTCACACTG | chr2R:13158141-13160141 | 544 | 558 | + | 16.05 | 5.57E-07 | 0.102 | TCCACGGTCACACTG |
| 1 | GMTACGGTCACACTG | chrX:5746747-5748747 | 551 | 565 | - | 16.05 | 5.57E-07 | 0.102 | GATACGGCCCCACTG |
| 1 | GMTACGGTCACACTG | chr2R:21480927-21482927 | 609 | 623 | + | 16.05 | 5.57E-07 | 0.102 | GCTGCGGTTACACTG |
| 1 | GMTACGGTCACACTG | chr3R:19402455-19404455 | 621 | 635 | - | 16.05 | 5.57E-07 | 0.102 | GCTGCGGTCATACTG |
| 1 | GMTACGGTCACACTG | chr2L:10375168-10377168 | 622 | 636 | - | 16.05 | 5.57E-07 | 0.102 | GACACGGTCACGCTG |
| 1 | GMTACGGTCACACTG | chr3R:13010649-13012649 | 626 | 640 | - | 16.05 | 5.57E-07 | 0.102 | ACGACGGTCACACTG |
| 1 | GMTACGGTCACACTG | chr2L:4393196-4395196 | 655 | 669 | - | 16.05 | 5.57E-07 | 0.102 | TCCACGGTCACACTG |
| 1 | GMTACGGTCACACTG | chr3L:18747687-18749687 | 665 | 679 | - | 16.05 | 5.57E-07 | 0.102 | GCTTCGGTCACACTA |
| 1 | GMTACGGTCACACTG | chr2R:6661265-6663265 | 680 | 694 | - | 16.05 | 5.57E-07 | 0.102 | TCTGCGGTCACACTG |
| 1 | GMTACGGTCACACTG | chr2R:14851941-14853941 | 686 | 700 | - | 16.05 | 5.57E-07 | 0.102 | GCCACGGTCACACTA |
| 1 | GMTACGGTCACACTG | chr2L:276582-278582 | 709 | 723 | - | 16.05 | 5.57E-07 | 0.102 | TCGACGGTCACACTG |
| 1 | GMTACGGTCACACTG | chr3L:10637146-10639146 | 711 | 725 | - | 16.05 | 5.57E-07 | 0.102 | ACAACGGTCACACTG |
| 1 | GMTACGGTCACACTG | chr3L:10637130-10639130 | 727 | 741 | - | 16.05 | 5.57E-07 | 0.102 | ACAACGGTCACACTG |
| 1 | GMTACGGTCACACTG | chr3R:24752849-24754849 | 750 | 764 | + | 16.05 | 5.57E-07 | 0.102 | GCAACGTTCACACTG |
| 1 | GMTACGGTCACACTG | chr2R:23889512-23891512 | 751 | 765 | + | 16.05 | 5.57E-07 | 0.102 | GCTCTGGTCACACTG |
| 1 | GMTACGGTCACACTG | chr2R:12518166-12520166 | 742 | 756 | - | 16.05 | 5.57E-07 | 0.102 | GCGACGGTAACACTG |
| 1 | GMTACGGTCACACTG | chr3L:15565013-15567013 | 759 | 773 | + | 16.05 | 5.57E-07 | 0.102 | TCTTCGGTCACACTG |
| 1 | GMTACGGTCACACTG | chr3L:16111626-16113626 | 754 | 768 | - | 16.05 | 5.57E-07 | 0.102 | GCTGCGGTCACACTA |
| 1 | GMTACGGTCACACTG | chrX:22528964-22530964 | 782 | 796 | + | 16.05 | 5.57E-07 | 0.102 | GACACGGTCACACGG |
| 1 | GMTACGGTCACACTG | chr2L:5541480-5543480 | 800 | 814 | - | 16.05 | 5.57E-07 | 0.102 | ACTTCGGTCACACTG |
| 1 | GMTACGGTCACACTG | chr2L:4970702-4972702 | 815 | 829 | + | 16.05 | 5.57E-07 | 0.102 | GCAACGGTCACACTA |
| 1 | GMTACGGTCACACTG | chr2R:6661118-6663118 | 827 | 841 | - | 16.05 | 5.57E-07 | 0.102 | TCTGCGGTCACACTG |
| 1 | GMTACGGTCACACTG | chrX:9318359-9320359 | 872 | 886 | + | 16.05 | 5.57E-07 | 0.102 | ACCACGGTCACACTG |
| 1 | GMTACGGTCACACTG | chrX:10876893-10878893 | 892 | 906 | - | 16.05 | 5.57E-07 | 0.102 | GCCAAGGTCACACTG |
| 1 | GMTACGGTCACACTG | chr2R:23382483-23384483 | 908 | 922 | - | 16.05 | 5.57E-07 | 0.102 | CCTACGGTCACTCTG |
| 1 | GMTACGGTCACACTG | chr2R:23953091-23955091 | 919 | 933 | - | 16.05 | 5.57E-07 | 0.102 | GCTGCGGTCACACTT |
| 1 | GMTACGGTCACACTG | chr2L:3465157-3467157 | 921 | 935 | - | 16.05 | 5.57E-07 | 0.102 | ACTGCGGTCACACTG |
| 1 | GMTACGGTCACACTG | chr3R:9403385-9405385 | 922 | 936 | - | 16.05 | 5.57E-07 | 0.102 | GCTCTGGTCACACTG |
| 1 | GMTACGGTCACACTG | chr3R:9403385-9405385 | 922 | 936 | - | 16.05 | 5.57E-07 | 0.102 | GCTCTGGTCACACTG |
| 1 | GMTACGGTCACACTG | chr3R:19402153-19404153 | 923 | 937 | - | 16.05 | 5.57E-07 | 0.102 | GCTGCGGTCATACTG |

| | | | | | | | | | |
|---|---|---|---|---|---|---|---|---|---|
| 1 | GMTACGGTCACACTG | chrX:15318479-15320479 | 948 | 962 | + | 16.05 | 5.57E-07 | 0.102 | CCGACGGTCACACTG |
| 1 | GMTACGGTCACACTG | chr2R:13553077-13555077 | 957 | 971 | + | 16.05 | 5.57E-07 | 0.102 | GCTGCGGTCACACTA |
| 1 | GMTACGGTCACACTG | chrX:3133870-3135870 | 975 | 989 | + | 16.05 | 5.57E-07 | 0.102 | GACGCGGTCACACTG |
| 1 | GMTACGGTCACACTG | chr3L:11112740-11114740 | 962 | 976 | - | 16.05 | 5.57E-07 | 0.102 | GCTACGGTCACACCA |
| 1 | GMTACGGTCACACTG | chr2R:23953048-23955048 | 962 | 976 | - | 16.05 | 5.57E-07 | 0.102 | GCTGCGGTCACACTT |
| 1 | GMTACGGTCACACTG | chrX:17248502-17250502 | 963 | 977 | - | 16.05 | 5.57E-07 | 0.102 | GAAACGGCCACACTG |
| 1 | GMTACGGTCACACTG | chr3L:7910954-7912954 | 965 | 979 | - | 16.05 | 5.57E-07 | 0.102 | GCGACGGTTACACTG |
| 1 | GMTACGGTCACACTG | chr2R:10896885-10898885 | 965 | 979 | - | 16.05 | 5.57E-07 | 0.102 | ACTGCGGTCACACTG |
| 1 | GMTACGGTCACACTG | chrX:21604264-21606264 | 965 | 979 | - | 16.05 | 5.57E-07 | 0.102 | GCGACGGTCACATTG |
| 1 | GMTACGGTCACACTG | chr3R:13010310-13012310 | 965 | 979 | - | 16.05 | 5.57E-07 | 0.102 | ACGACGGTCACACTG |
| 1 | GMTACGGTCACACTG | chr2R:23376045-23378045 | 979 | 993 | + | 16.05 | 5.57E-07 | 0.102 | TCAACGGTCACACTG |
| 1 | GMTACGGTCACACTG | chr2L:5541314-5543314 | 966 | 980 | - | 16.05 | 5.57E-07 | 0.102 | ACTTCGGTCACACTG |
| 1 | GMTACGGTCACACTG | chr2L:3465112-3467112 | 966 | 980 | - | 16.05 | 5.57E-07 | 0.102 | ACTGCGGTCACACTG |
| 1 | GMTACGGTCACACTG | chr2R:23382424-23384424 | 967 | 981 | - | 16.05 | 5.57E-07 | 0.102 | CCTACGGTCACTCTG |
| 1 | GMTACGGTCACACTG | chr3R:16995446-16997446 | 985 | 999 | + | 16.05 | 5.57E-07 | 0.102 | GCGACGGTCACACTA |
| 1 | GMTACGGTCACACTG | chr2R:23889276-23891276 | 987 | 1001 | + | 16.05 | 5.57E-07 | 0.102 | GCTCTGGTCACACTG |
| 1 | GMTACGGTCACACTG | chr3L:21532713-21534713 | 973 | 987 | - | 16.05 | 5.57E-07 | 0.102 | GCTTTGGTCACACTG |
| 1 | GMTACGGTCACACTG | chr2R:14851653-14853653 | 974 | 988 | - | 16.05 | 5.57E-07 | 0.102 | GCCACGGTCACACTA |
| 1 | GMTACGGTCACACTG | chr2R:10251417-10253417 | 975 | 989 | - | 16.05 | 5.57E-07 | 0.102 | TCAACGGTCACACTG |
| 1 | GMTACGGTCACACTG | chr2R:9399329-9401329 | 975 | 989 | - | 16.05 | 5.57E-07 | 0.102 | TCTACGGTCACGCTG |
| 1 | GMTACGGTCACACTG | chr2L:4392876-4394876 | 975 | 989 | - | 16.05 | 5.57E-07 | 0.102 | TCCACGGTCACACTG |
| 1 | GMTACGGTCACACTG | chrX:17888547-17890547 | 990 | 1004 | + | 16.05 | 5.57E-07 | 0.102 | GCTACGCACACACTG |
| 1 | GMTACGGTCACACTG | chrX:17898702-17900702 | 990 | 1004 | + | 16.05 | 5.57E-07 | 0.102 | GCTACGCACACACTG |
| 1 | GMTACGGTCACACTG | chr3R:13381106-13383106 | 991 | 1005 | + | 16.05 | 5.57E-07 | 0.102 | GCTCTGGTCACACTG |
| 1 | GMTACGGTCACACTG | chr3R:27920089-27922089 | 977 | 991 | - | 16.05 | 5.57E-07 | 0.102 | GCTACGGTCACACCA |
| 1 | GMTACGGTCACACTG | chr2R:17517223-17519223 | 991 | 1005 | + | 16.05 | 5.57E-07 | 0.102 | GCTCTGGTCACACTG |
| 1 | GMTACGGTCACACTG | chr3R:29130193-29132193 | 993 | 1007 | + | 16.05 | 5.57E-07 | 0.102 | GCCACGGTCACACTA |
| 1 | GMTACGGTCACACTG | chr2L:3451466-3453466 | 995 | 1009 | + | 16.05 | 5.57E-07 | 0.102 | CCTACGGTCACACCG |
| 1 | GMTACGGTCACACTG | chr2L:20757144-20759144 | 995 | 1009 | + | 16.05 | 5.57E-07 | 0.102 | GCTGCGGTCACACTA |
| 1 | GMTACGGTCACACTG | chr2L:6466698-6468698 | 982 | 996 | - | 16.05 | 5.57E-07 | 0.102 | GAAACGGTCACGCTG |
| 1 | GMTACGGTCACACTG | chrX:17906692-17908692 | 996 | 1010 | + | 16.05 | 5.57E-07 | 0.102 | GCTGTGGTCACACTG |
| 1 | GMTACGGTCACACTG | chr3R:14308089-14310089 | 996 | 1010 | + | 16.05 | 5.57E-07 | 0.102 | GCCACGGTCACACTT |
| 1 | GMTACGGTCACACTG | chr3L:14775129-14777129 | 984 | 998 | - | 16.05 | 5.57E-07 | 0.102 | GAAACGGCCACACTG |

| | | | | | | | | | |
|---|---|---|---|---|---|---|---|---|---|
| 1 | GMTACGGTCACACTG | chr3L:18747367-18749367 | 985 | 999 | - | 16.05 | 5.57E-07 | 0.102 | GCTTCGGTCACACTA |
| 1 | GMTACGGTCACACTG | chr3L:21496494-21498494 | 986 | 1000 | - | 16.05 | 5.57E-07 | 0.102 | ACTGCGGTCACACTG |
| 1 | GMTACGGTCACACTG | chr2L:21261956-21263956 | 1002 | 1016 | + | 16.05 | 5.57E-07 | 0.102 | ACTGCGGTCACACTG |
| 1 | GMTACGGTCACACTG | chrX:5420533-5422533 | 1002 | 1016 | + | 16.05 | 5.57E-07 | 0.102 | GCAACGGTCACACTT |
| 1 | GMTACGGTCACACTG | chr2R:19282961-19284961 | 990 | 1004 | - | 16.05 | 5.57E-07 | 0.102 | ACAACGGTCACACTG |
| 1 | GMTACGGTCACACTG | chr3L:9854285-9856285 | 990 | 1004 | - | 16.05 | 5.57E-07 | 0.102 | GCTACGGTCACTCTA |
| 1 | GMTACGGTCACACTG | chr3L:17424477-17426477 | 1004 | 1018 | + | 16.05 | 5.57E-07 | 0.102 | ACAACGGTCACACTG |
| 1 | GMTACGGTCACACTG | chr3R:16452821-16454821 | 991 | 1005 | - | 16.05 | 5.57E-07 | 0.102 | GCTGCGGTCACACTT |
| 1 | GMTACGGTCACACTG | chr3R:29856491-29858491 | 991 | 1005 | - | 16.05 | 5.57E-07 | 0.102 | GCTGTGGTCACACTG |
| 1 | GMTACGGTCACACTG | chr3R:13271011-13273011 | 992 | 1006 | - | 16.05 | 5.57E-07 | 0.102 | TCAACGGTCACACTG |
| 1 | GMTACGGTCACACTG | chr3L:16112212-16114212 | 1006 | 1020 | + | 16.05 | 5.57E-07 | 0.102 | CCAACGGTCACACTG |
| 1 | GMTACGGTCACACTG | chr3L:19259679-19261679 | 993 | 1007 | - | 16.05 | 5.57E-07 | 0.102 | CCTACGGCCACACTG |
| 1 | GMTACGGTCACACTG | chr2R:7394932-7396932 | 993 | 1007 | - | 16.05 | 5.57E-07 | 0.102 | GCGAGGGTCACACTG |
| 1 | GMTACGGTCACACTG | chr3R:21101199-21103199 | 1008 | 1022 | + | 16.05 | 5.57E-07 | 0.102 | GCCATGGTCACACTG |
| 1 | GMTACGGTCACACTG | chr2R:6970637-6972637 | 994 | 1008 | - | 16.05 | 5.57E-07 | 0.102 | GCGATGGTCACACTG |
| 1 | GMTACGGTCACACTG | chr2R:14768423-14770423 | 994 | 1008 | - | 16.05 | 5.57E-07 | 0.102 | TCCACGGTCACACTG |
| 1 | GMTACGGTCACACTG | chrX:9318223-9320223 | 1008 | 1022 | + | 16.05 | 5.57E-07 | 0.102 | ACCACGGTCACACTG |
| 1 | GMTACGGTCACACTG | chr2R:17004451-17006451 | 1008 | 1022 | + | 16.05 | 5.57E-07 | 0.102 | GCTGCGGTCACACTA |
| 1 | GMTACGGTCACACTG | chr2R:6970637-6972637 | 994 | 1008 | - | 16.05 | 5.57E-07 | 0.102 | GCGATGGTCACACTG |
| 1 | GMTACGGTCACACTG | chr2R:12517912-12519912 | 996 | 1010 | - | 16.05 | 5.57E-07 | 0.102 | GCGACGGTAACACTG |
| 1 | GMTACGGTCACACTG | chr3R:31610046-31612046 | 1010 | 1024 | + | 16.05 | 5.57E-07 | 0.102 | GAAACGGTCACTCTG |
| 1 | GMTACGGTCACACTG | chr3L:5809973-5811973 | 996 | 1010 | - | 16.05 | 5.57E-07 | 0.102 | GCTACGGCCACACTT |
| 1 | GMTACGGTCACACTG | chr3L:3302708-3304708 | 1011 | 1025 | + | 16.05 | 5.57E-07 | 0.102 | GCTGCAGTCACACTG |
| 1 | GMTACGGTCACACTG | chr3R:11215535-11217535 | 1011 | 1025 | + | 16.05 | 5.57E-07 | 0.102 | GCAACGGTCACACTC |
| 1 | GMTACGGTCACACTG | chrX:21357286-21359286 | 1011 | 1025 | + | 16.05 | 5.57E-07 | 0.102 | GCTGCGGTCACACTA |
| 1 | GMTACGGTCACACTG | chr3R:13387709-13389709 | 999 | 1013 | - | 16.05 | 5.57E-07 | 0.102 | GCTCCGGTCACACTT |
| 1 | GMTACGGTCACACTG | chr3R:12690743-12692743 | 1015 | 1029 | + | 16.05 | 5.57E-07 | 0.102 | CCTACGGTCACCCTG |
| 1 | GMTACGGTCACACTG | chr2L:14354145-14356145 | 1001 | 1015 | - | 16.05 | 5.57E-07 | 0.102 | CCTGCGGTCACACTG |
| 1 | GMTACGGTCACACTG | chrX:18656755-18658755 | 1017 | 1031 | + | 16.05 | 5.57E-07 | 0.102 | GCTACGGTCATACCG |
| 1 | GMTACGGTCACACTG | chr3L:1856165-1858165 | 1019 | 1033 | + | 16.05 | 5.57E-07 | 0.102 | GCTCTGGTCACACTG |
| 1 | GMTACGGTCACACTG | chr2R:13157666-13159666 | 1019 | 1033 | + | 16.05 | 5.57E-07 | 0.102 | TCCACGGTCACACTG |
| 1 | GMTACGGTCACACTG | chr2R:16866714-16868714 | 1020 | 1034 | + | 16.05 | 5.57E-07 | 0.102 | GCAACGGTAACACTG |
| 1 | GMTACGGTCACACTG | chr3R:9236682-9238682 | 1008 | 1022 | - | 16.05 | 5.57E-07 | 0.102 | GCTGCGGTCACACTA |

| | | | | | | | | | |
|---|---|---|---|---|---|---|---|---|---|
| 1 | GMTACGGTCACACTG | chr2R:5670741-5672741 | 1023 | 1037 | + | 16.05 | 5.57E-07 | 0.102 | GCGACGGTCACACTT |
| 1 | GMTACGGTCACACTG | chr3L:19814938-19816938 | 1013 | 1027 | - | 16.05 | 5.57E-07 | 0.102 | GCCACGTTCACACTG |
| 1 | GMTACGGTCACACTG | chr2R:17517186-17519186 | 1028 | 1042 | + | 16.05 | 5.57E-07 | 0.102 | GCTCTGGTCACACTG |
| 1 | GMTACGGTCACACTG | chrX:2008364-2010364 | 1014 | 1028 | - | 16.05 | 5.57E-07 | 0.102 | TCGACGGTCACACTG |
| 1 | GMTACGGTCACACTG | chr3L:19260394-19262394 | 1029 | 1043 | + | 16.05 | 5.57E-07 | 0.102 | GCTTTGGTCACACTG |
| 1 | GMTACGGTCACACTG | chrX:13844866-13846866 | 1017 | 1031 | - | 16.05 | 5.57E-07 | 0.102 | GCTCTGGTCACACTG |
| 1 | GMTACGGTCACACTG | chr2R:8946344-8948344 | 1021 | 1035 | - | 16.05 | 5.57E-07 | 0.102 | GCGAGGGTCACACTG |
| 1 | GMTACGGTCACACTG | chr2R:18183090-18185090 | 1041 | 1055 | + | 16.05 | 5.57E-07 | 0.102 | GCTGCGGTCACACTA |
| 1 | GMTACGGTCACACTG | chr2L:6472904-6474904 | 1032 | 1046 | - | 16.05 | 5.57E-07 | 0.102 | TCAACGGTCACACTG |
| 1 | GMTACGGTCACACTG | chr2L:10374757-10376757 | 1033 | 1047 | - | 16.05 | 5.57E-07 | 0.102 | GACACGGTCACGCTG |
| 1 | GMTACGGTCACACTG | chr2L:10841597-10843597 | 1035 | 1049 | - | 16.05 | 5.57E-07 | 0.102 | GCTACGGCCACACTA |
| 1 | GMTACGGTCACACTG | chr3L:5809931-5811931 | 1038 | 1052 | - | 16.05 | 5.57E-07 | 0.102 | GCTACGGCCACACTT |
| 1 | GMTACGGTCACACTG | chr3R:9403269-9405269 | 1038 | 1052 | - | 16.05 | 5.57E-07 | 0.102 | GCTCTGGTCACACTG |
| 1 | GMTACGGTCACACTG | chrX:5746254-5748254 | 1044 | 1058 | - | 16.05 | 5.57E-07 | 0.102 | GATACGGCCCCACTG |
| 1 | GMTACGGTCACACTG | chr3R:21348601-21350601 | 1057 | 1071 | - | 16.05 | 5.57E-07 | 0.102 | GCTGCGGTCATACTG |
| 1 | GMTACGGTCACACTG | chr2L:14354085-14356085 | 1061 | 1075 | - | 16.05 | 5.57E-07 | 0.102 | CCTGCGGTCACACTG |
| 1 | GMTACGGTCACACTG | chrX:16586172-16588172 | 1065 | 1079 | - | 16.05 | 5.57E-07 | 0.102 | GAAACGGTCACACCG |
| 1 | GMTACGGTCACACTG | chrX:15943897-15945897 | 1089 | 1103 | + | 16.05 | 5.57E-07 | 0.102 | TCAACGGTCACACTG |
| 1 | GMTACGGTCACACTG | chr3R:21855266-21857266 | 1095 | 1109 | + | 16.05 | 5.57E-07 | 0.102 | GAAACGGTCACTCTG |
| 1 | GMTACGGTCACACTG | chr2R:12756546-12758546 | 1101 | 1115 | + | 16.05 | 5.57E-07 | 0.102 | GAAGCGGTCACACTG |
| 1 | GMTACGGTCACACTG | chr2R:12756529-12758529 | 1118 | 1132 | + | 16.05 | 5.57E-07 | 0.102 | GAAGCGGTCACACTG |
| 1 | GMTACGGTCACACTG | chr3R:16642605-16644605 | 1137 | 1151 | + | 16.05 | 5.57E-07 | 0.102 | TCAACGGTCACACTG |
| 1 | GMTACGGTCACACTG | chr2L:20756985-20758985 | 1154 | 1168 | + | 16.05 | 5.57E-07 | 0.102 | GCTGCGGTCACACTA |
| 1 | GMTACGGTCACACTG | chr3L:22074867-22076867 | 1193 | 1207 | + | 16.05 | 5.57E-07 | 0.102 | GAAGCGGTCACACTG |
| 1 | GMTACGGTCACACTG | chr3R:29129972-29131972 | 1214 | 1228 | + | 16.05 | 5.57E-07 | 0.102 | GCCACGGTCACACTA |
| 1 | GMTACGGTCACACTG | chr3R:11215287-11217287 | 1259 | 1273 | + | 16.05 | 5.57E-07 | 0.102 | GCAACGGTCACACTC |
| 1 | GMTACGGTCACACTG | chr3L:17424217-17426217 | 1264 | 1278 | + | 16.05 | 5.57E-07 | 0.102 | ACAACGGTCACACTG |
| 1 | GMTACGGTCACACTG | chr3R:23355278-23357278 | 1282 | 1296 | + | 16.05 | 5.57E-07 | 0.102 | GCTCAGGTCACACTG |
| 1 | GMTACGGTCACACTG | chr3R:13380809-13382809 | 1288 | 1302 | + | 16.05 | 5.57E-07 | 0.102 | GCTCTGGTCACACTG |
| 1 | GMTACGGTCACACTG | chr3R:12690468-12692468 | 1290 | 1304 | + | 16.05 | 5.57E-07 | 0.102 | CCTACGGTCACCCTG |
| 1 | GMTACGGTCACACTG | chr3L:22074756-22076756 | 1304 | 1318 | + | 16.05 | 5.57E-07 | 0.102 | GAAGCGGTCACACTG |
| 1 | GMTACGGTCACACTG | chr2R:17004149-17006149 | 1310 | 1324 | + | 16.05 | 5.57E-07 | 0.102 | GCTGCGGTCACACTA |
| 1 | GMTACGGTCACACTG | chr3L:17473761-17475761 | 1312 | 1326 | + | 16.05 | 5.57E-07 | 0.102 | TCCACGGTCACACTG |

| | | | | | | | | | |
|---|---|---|---|---|---|---|---|---|---|
| 1 | GMTACGGTCACACTG | chr3L:11444161-11446161 | 1345 | 1359 | + | 16.05 | 5.57E-07 | 0.102 | GCGACGGTAACACTG |
| 1 | GMTACGGTCACACTG | chr2R:21480176-21482176 | 1360 | 1374 | + | 16.05 | 5.57E-07 | 0.102 | GCTGCGGTTACACTG |
| 1 | GMTACGGTCACACTG | chr3R:16995071-16997071 | 1360 | 1374 | + | 16.05 | 5.57E-07 | 0.102 | GCGACGGTCACACTA |
| 1 | GMTACGGTCACACTG | chr3L:11444104-11446104 | 1402 | 1416 | + | 16.05 | 5.57E-07 | 0.102 | GCGACGGTAACACTG |
| 1 | GMTACGGTCACACTG | chr2R:9115122-9117122 | 1405 | 1419 | - | 16.05 | 5.57E-07 | 0.102 | TCTGCGGTCACACTG |
| 1 | GMTACGGTCACACTG | chrX:17906261-17908261 | 1427 | 1441 | + | 16.05 | 5.57E-07 | 0.102 | GCTGTGGTCACACTG |
| 1 | GMTACGGTCACACTG | chr2R:15379988-15381988 | 1452 | 1466 | + | 16.05 | 5.57E-07 | 0.102 | GCTGTGGTCACACTG |
| 1 | GMTACGGTCACACTG | chrX:5420533-5422533 | 1458 | 1472 | + | 16.05 | 5.57E-07 | 0.102 | GCAACGGTCACACTT |
| 1 | GMTACGGTCACACTG | chrX:18656286-18658286 | 1486 | 1500 | + | 16.05 | 5.57E-07 | 0.102 | GCTACGGTCATACCG |
| 1 | GMTACGGTCACACTG | chr3R:21100718-21102718 | 1489 | 1503 | + | 16.05 | 5.57E-07 | 0.102 | GCCATGGTCACACTG |
| 1 | GMTACGGTCACACTG | chr3R:23355064-23357064 | 1496 | 1510 | + | 16.05 | 5.57E-07 | 0.102 | GCTCAGGTCACACTG |
| 1 | GMTACGGTCACACTG | chr3R:14307589-14309589 | 1496 | 1510 | + | 16.05 | 5.57E-07 | 0.102 | GCCACGGTCACACTT |
| 1 | GMTACGGTCACACTG | chr3L:19518649-19520649 | 1551 | 1565 | + | 16.05 | 5.57E-07 | 0.102 | GCTACGGTCACGATG |
| 1 | GMTACGGTCACACTG | chr3R:32051060-32053060 | 1554 | 1568 | + | 16.05 | 5.57E-07 | 0.102 | ACAACGGTCACACTG |
| 1 | GMTACGGTCACACTG | chr3L:16111626-16113626 | 1592 | 1606 | + | 16.05 | 5.57E-07 | 0.102 | CCAACGGTCACACTG |
| 1 | GMTACGGTCACACTG | chr3L:3302112-3304112 | 1607 | 1621 | + | 16.05 | 5.57E-07 | 0.102 | GCTGCAGTCACACTG |
| 1 | GMTACGGTCACACTG | chr3L:15987236-15989236 | 1617 | 1631 | + | 16.05 | 5.57E-07 | 0.102 | GAAACGGTCACCCTG |
| 1 | GMTACGGTCACACTG | chr3R:32050993-32052993 | 1621 | 1635 | + | 16.05 | 5.57E-07 | 0.102 | ACAACGGTCACACTG |
| 1 | GMTACGGTCACACTG | chr3L:19259040-19261040 | 1632 | 1646 | - | 16.05 | 5.57E-07 | 0.102 | CCTACGGCCACACTG |
| 1 | GMTACGGTCACACTG | chrX:5419867-5421867 | 1668 | 1682 | + | 16.05 | 5.57E-07 | 0.102 | GCAACGGTCACACTT |
| 1 | GMTACGGTCACACTG | chr2R:15379724-15381724 | 1716 | 1730 | + | 16.05 | 5.57E-07 | 0.102 | GCTGTGGTCACACTG |
| 1 | GMTACGGTCACACTG | chr3L:19259679-19261679 | 1744 | 1758 | + | 16.05 | 5.57E-07 | 0.102 | GCTTTGGTCACACTG |
| 1 | GMTACGGTCACACTG | chr2R:13552240-13554240 | 1794 | 1808 | + | 16.05 | 5.57E-07 | 0.102 | GCTGCGGTCACACTA |
| 1 | GMTACGGTCACACTG | chr2R:7393996-7395996 | 1929 | 1943 | - | 16.05 | 5.57E-07 | 0.102 | GCGAGGGTCACACTG |
| 1 | GMTACGGTCACACTG | chr3R:29745083-29747083 | 1 | 15 | - | 15.771 | 1.11E-06 | 0.148 | GAAACGGTCATACTG |
| 1 | GMTACGGTCACACTG | chr3R:15924629-15926629 | 281 | 295 | + | 15.771 | 1.11E-06 | 0.148 | GCTACGATCACATTG |
| 1 | GMTACGGTCACACTG | chr3R:21073007-21075007 | 313 | 327 | - | 15.771 | 1.11E-06 | 0.148 | GATACGTGCACACTG |
| 1 | GMTACGGTCACACTG | chr2L:2884929-2886929 | 357 | 371 | + | 15.771 | 1.11E-06 | 0.148 | CATACGGCCACACTG |
| 1 | GMTACGGTCACACTG | chr3L:22721091-22723091 | 357 | 371 | + | 15.771 | 1.11E-06 | 0.148 | GAGACGGTCAGACTG |
| 1 | GMTACGGTCACACTG | chr3R:22725967-22727967 | 433 | 447 | + | 15.771 | 1.11E-06 | 0.148 | AAAACGGTCACACTG |
| 1 | GMTACGGTCACACTG | chr2L:2884850-2886850 | 436 | 450 | + | 15.771 | 1.11E-06 | 0.148 | CATACGGCCACACTG |
| 1 | GMTACGGTCACACTG | chr3R:15507886-15509886 | 449 | 463 | - | 15.771 | 1.11E-06 | 0.148 | GATACGGTCACTGTG |
| 1 | GMTACGGTCACACTG | chr3R:19650018-19652018 | 490 | 504 | + | 15.771 | 1.11E-06 | 0.148 | GATACGGGCACACTT |

| | | | | | | | | | |
|---|---|---|---|---|---|---|---|---|---|
| 1 | GMTACGGTCACACTG | chr2L:22168668-22170668 | 500 | 514 | - | 15.771 | 1.11E-06 | 0.148 | GAAACGGTCACACTT |
| 1 | GMTACGGTCACACTG | chr2L:10210143-10212143 | 523 | 537 | - | 15.771 | 1.11E-06 | 0.148 | GATCCGGTCACACTA |
| 1 | GMTACGGTCACACTG | chr2L:16737850-16739850 | 531 | 545 | - | 15.771 | 1.11E-06 | 0.148 | AAGACGGTCACACTG |
| 1 | GMTACGGTCACACTG | chrX:15803370-15805370 | 549 | 563 | - | 15.771 | 1.11E-06 | 0.148 | AATACGGTCACACCG |
| 1 | GMTACGGTCACACTG | chr3L:17845916-17847916 | 568 | 582 | - | 15.771 | 1.11E-06 | 0.148 | AATGCGGTCACACTG |
| 1 | GMTACGGTCACACTG | chr2R:8723258-8725258 | 592 | 606 | + | 15.771 | 1.11E-06 | 0.148 | AACACGGTCACACTG |
| 1 | GMTACGGTCACACTG | chr3L:16131028-16133028 | 584 | 598 | - | 15.771 | 1.11E-06 | 0.148 | GATTTGGTCACACTG |
| 1 | GMTACGGTCACACTG | chr4:787978-789978 | 606 | 620 | + | 15.771 | 1.11E-06 | 0.148 | GATCCGTTCACACTG |
| 1 | GMTACGGTCACACTG | chrX:16005147-16007147 | 625 | 639 | - | 15.771 | 1.11E-06 | 0.148 | ACTACGTTCACACTG |
| 1 | GMTACGGTCACACTG | chr2L:8402924-8404924 | 648 | 662 | + | 15.771 | 1.11E-06 | 0.148 | CATACGGGCACACTG |
| 1 | GMTACGGTCACACTG | chr2R:11306879-11308879 | 636 | 650 | - | 15.771 | 1.11E-06 | 0.148 | GATTCAGTCACACTG |
| 1 | GMTACGGTCACACTG | chr3L:21727932-21729932 | 714 | 728 | + | 15.771 | 1.11E-06 | 0.148 | TAGACGGTCACACTG |
| 1 | GMTACGGTCACACTG | chr2L:9911911-9913911 | 723 | 737 | + | 15.771 | 1.11E-06 | 0.148 | AATGCGGTCACACTG |
| 1 | GMTACGGTCACACTG | chr3R:31579951-31581951 | 718 | 732 | - | 15.771 | 1.11E-06 | 0.148 | CAAACGGTCACACTG |
| 1 | GMTACGGTCACACTG | chrX:13743928-13745928 | 720 | 734 | - | 15.771 | 1.11E-06 | 0.148 | GATTCGGTAACACTG |
| 1 | GMTACGGTCACACTG | chr2R:21142545-21144545 | 755 | 769 | + | 15.771 | 1.11E-06 | 0.148 | TAGACGGTCACACTG |
| 1 | GMTACGGTCACACTG | chr3R:29744337-29746337 | 747 | 761 | - | 15.771 | 1.11E-06 | 0.148 | GAAACGGTCATACTG |
| 1 | GMTACGGTCACACTG | chr3L:2243478-2245478 | 756 | 770 | - | 15.771 | 1.11E-06 | 0.148 | AATACGGCCACACTG |
| 1 | GMTACGGTCACACTG | chr3R:29932287-29934287 | 765 | 779 | - | 15.771 | 1.11E-06 | 0.148 | GACACGGTCATACTG |
| 1 | GMTACGGTCACACTG | chr3L:2588414-2590414 | 779 | 793 | + | 15.771 | 1.11E-06 | 0.148 | ACTACGGTCACACTA |
| 1 | GMTACGGTCACACTG | chr3R:29932287-29934287 | 765 | 779 | - | 15.771 | 1.11E-06 | 0.148 | GACACGGTCATACTG |
| 1 | GMTACGGTCACACTG | chr3L:17041797-17043797 | 767 | 781 | - | 15.771 | 1.11E-06 | 0.148 | GATATGGGCACACTG |
| 1 | GMTACGGTCACACTG | chr2L:20952-22952 | 790 | 804 | - | 15.771 | 1.11E-06 | 0.148 | GACATGGTCACACTG |
| 1 | GMTACGGTCACACTG | chr3L:19585253-19587253 | 809 | 823 | + | 15.771 | 1.11E-06 | 0.148 | GAGACGTTCACACTG |
| 1 | GMTACGGTCACACTG | chrX:16883581-16885581 | 799 | 813 | - | 15.771 | 1.11E-06 | 0.148 | GCTATGGTCACACTA |
| 1 | GMTACGGTCACACTG | chr2L:22168668-22170668 | 822 | 836 | + | 15.771 | 1.11E-06 | 0.148 | GCTAAGGTAACACTG |
| 1 | GMTACGGTCACACTG | chr3L:21727806-21729806 | 840 | 854 | + | 15.771 | 1.11E-06 | 0.148 | TAGACGGTCACACTG |
| 1 | GMTACGGTCACACTG | chr2L:15515727-15517727 | 867 | 881 | + | 15.771 | 1.11E-06 | 0.148 | GATAAGGTCACTCTG |
| 1 | GMTACGGTCACACTG | chr4:1229837-1231837 | 862 | 876 | - | 15.771 | 1.11E-06 | 0.148 | AACACGGTCACACTG |
| 1 | GMTACGGTCACACTG | chrX:6287100-6289100 | 884 | 898 | + | 15.771 | 1.11E-06 | 0.148 | AATACGGTCTCACTG |
| 1 | GMTACGGTCACACTG | chr3R:22420161-22422161 | 888 | 902 | - | 15.771 | 1.11E-06 | 0.148 | GCTATGGTCACACTT |
| 1 | GMTACGGTCACACTG | chr2R:20869791-20871791 | 935 | 949 | + | 15.771 | 1.11E-06 | 0.148 | GAAAGGGTCACACTG |
| 1 | GMTACGGTCACACTG | chr3R:24031157-24033157 | 967 | 981 | + | 15.771 | 1.11E-06 | 0.148 | AACACGGTCACACTG |

| | | | | | | | | | |
|---|---|---|---|---|---|---|---|---|---|
| 1 | GMTACGGTCACACTG | chr3L:21033535-21035535 | 967 | 981 | + | 15.771 | 1.11E-06 | 0.148 | TCTATGGTCACACTG |
| 1 | GMTACGGTCACACTG | chr3R:26911886-26913886 | 957 | 971 | - | 15.771 | 1.11E-06 | 0.148 | TACACGGTCACACTG |
| 1 | GMTACGGTCACACTG | chr3R:15833648-15835648 | 959 | 973 | - | 15.771 | 1.11E-06 | 0.148 | GAAATGGTCACACTG |
| 1 | GMTACGGTCACACTG | chr3R:27947711-27949711 | 974 | 988 | + | 15.771 | 1.11E-06 | 0.148 | AATACGGTCACTCTG |
| 1 | GMTACGGTCACACTG | chrX:16004812-16006812 | 960 | 974 | - | 15.771 | 1.11E-06 | 0.148 | ACTACGTTCACACTG |
| 1 | GMTACGGTCACACTG | chr3L:14244993-14246993 | 960 | 974 | - | 15.771 | 1.11E-06 | 0.148 | ACTACGGTCACACTA |
| 1 | GMTACGGTCACACTG | chr2R:20256650-20258650 | 978 | 992 | + | 15.771 | 1.11E-06 | 0.148 | AAAACGGTCACACTG |
| 1 | GMTACGGTCACACTG | chr2L:10055900-10057900 | 979 | 993 | + | 15.771 | 1.11E-06 | 0.148 | CCTACGTTCACACTG |
| 1 | GMTACGGTCACACTG | chrX:13743683-13745683 | 965 | 979 | - | 15.771 | 1.11E-06 | 0.148 | GATTCGGTAACACTG |
| 1 | GMTACGGTCACACTG | chr3L:6188745-6190745 | 981 | 995 | + | 15.771 | 1.11E-06 | 0.148 | GAAACGGTCAGACTG |
| 1 | GMTACGGTCACACTG | chr3L:360472-362472 | 969 | 983 | - | 15.771 | 1.11E-06 | 0.148 | TATTCGGTCACACTG |
| 1 | GMTACGGTCACACTG | chr3R:31579699-31581699 | 970 | 984 | - | 15.771 | 1.11E-06 | 0.148 | CAAACGGTCACACTG |
| 1 | GMTACGGTCACACTG | chr3R:22725414-22727414 | 986 | 1000 | + | 15.771 | 1.11E-06 | 0.148 | AAAACGGTCACACTG |
| 1 | GMTACGGTCACACTG | chr2L:16309956-16311956 | 987 | 1001 | + | 15.771 | 1.11E-06 | 0.148 | GATGCGGTCACACTA |
| 1 | GMTACGGTCACACTG | chr3L:19585253-19587253 | 975 | 989 | - | 15.771 | 1.11E-06 | 0.148 | GAGATGGTCACACTG |
| 1 | GMTACGGTCACACTG | chrX:15768664-15770664 | 992 | 1006 | + | 15.771 | 1.11E-06 | 0.148 | GACATGGTCACACTG |
| 1 | GMTACGGTCACACTG | chr3L:16130632-16132632 | 980 | 994 | - | 15.771 | 1.11E-06 | 0.148 | GATTTGGTCACACTG |
| 1 | GMTACGGTCACACTG | chrX:6286990-6288990 | 994 | 1008 | + | 15.771 | 1.11E-06 | 0.148 | AATACGGTCTCACTG |
| 1 | GMTACGGTCACACTG | chr2L:6023539-6025539 | 981 | 995 | - | 15.771 | 1.11E-06 | 0.148 | AAAACGGTCACACTG |
| 1 | GMTACGGTCACACTG | chr2R:16782671-16784671 | 995 | 1009 | + | 15.771 | 1.11E-06 | 0.148 | CACACGGTCACACTG |
| 1 | GMTACGGTCACACTG | chr3R:9019022-9021022 | 981 | 995 | - | 15.771 | 1.11E-06 | 0.148 | ACTACGGTCACACTT |
| 1 | GMTACGGTCACACTG | chr3R:22420068-22422068 | 981 | 995 | - | 15.771 | 1.11E-06 | 0.148 | GCTATGGTCACACTT |
| 1 | GMTACGGTCACACTG | chr3L:11108690-11110690 | 996 | 1010 | + | 15.771 | 1.11E-06 | 0.148 | AAAACGGTCACACTG |
| 1 | GMTACGGTCACACTG | chr2R:24564554-24566554 | 996 | 1010 | + | 15.771 | 1.11E-06 | 0.148 | CCTACGGTCACACTA |
| 1 | GMTACGGTCACACTG | chr3L:17845501-17847501 | 983 | 997 | - | 15.771 | 1.11E-06 | 0.148 | AATGCGGTCACACTG |
| 1 | GMTACGGTCACACTG | chr3R:8307645-8309645 | 984 | 998 | - | 15.771 | 1.11E-06 | 0.148 | GCTACAGTCACATTG |
| 1 | GMTACGGTCACACTG | chr2L:10200004-10202004 | 984 | 998 | - | 15.771 | 1.11E-06 | 0.148 | CAAACGGTCACACTG |
| 1 | GMTACGGTCACACTG | chr2L:8402573-8404573 | 999 | 1013 | + | 15.771 | 1.11E-06 | 0.148 | CATACGGGCACACTG |
| 1 | GMTACGGTCACACTG | chr2L:10209681-10211681 | 985 | 999 | - | 15.771 | 1.11E-06 | 0.148 | GATCCGGTCACACTA |
| 1 | GMTACGGTCACACTG | chr4:1229713-1231713 | 986 | 1000 | - | 15.771 | 1.11E-06 | 0.148 | AACACGGTCACACTG |
| 1 | GMTACGGTCACACTG | chr2L:11998654-12000654 | 1000 | 1014 | + | 15.771 | 1.11E-06 | 0.148 | AATACGGTCCCACTG |
| 1 | GMTACGGTCACACTG | chr2R:11306528-11308528 | 987 | 1001 | - | 15.771 | 1.11E-06 | 0.148 | GATTCAGTCACACTG |
| 1 | GMTACGGTCACACTG | chr2L:11994740-11996740 | 1003 | 1017 | + | 15.771 | 1.11E-06 | 0.148 | AATACGGTCCCACTG |

| | | | | | | | | | |
|---|---|---|---|---|---|---|---|---|---|
| 1 | GMTACGGTCACACTG | chr3L:17041574-17043574 | 990 | 1004 | - | 15.771 | 1.11E-06 | 0.148 | GATATGGGCACACTG |
| 1 | GMTACGGTCACACTG | chr3R:29744092-29746092 | 992 | 1006 | - | 15.771 | 1.11E-06 | 0.148 | GAAACGGTCATACTG |
| 1 | GMTACGGTCACACTG | chr2L:16737388-16739388 | 993 | 1007 | - | 15.771 | 1.11E-06 | 0.148 | AAGACGGTCACACTG |
| 1 | GMTACGGTCACACTG | chr3R:7472781-7474781 | 993 | 1007 | - | 15.771 | 1.11E-06 | 0.148 | TACACGGTCACACTG |
| 1 | GMTACGGTCACACTG | chr3R:29932057-29934057 | 995 | 1009 | - | 15.771 | 1.11E-06 | 0.148 | GACACGGTCATACTG |
| 1 | GMTACGGTCACACTG | chr3L:7322619-7324619 | 1011 | 1025 | + | 15.771 | 1.11E-06 | 0.148 | GACATGGTCACACTG |
| 1 | GMTACGGTCACACTG | chr2R:8722839-8724839 | 1011 | 1025 | + | 15.771 | 1.11E-06 | 0.148 | AACACGGTCACACTG |
| 1 | GMTACGGTCACACTG | chr3R:12430321-12432321 | 1012 | 1026 | + | 15.771 | 1.11E-06 | 0.148 | CATTCGGTCACACTG |
| 1 | GMTACGGTCACACTG | chr2R:18818467-18820467 | 1012 | 1026 | + | 15.771 | 1.11E-06 | 0.148 | CATACGGTCACACCG |
| 1 | GMTACGGTCACACTG | chr2L:12171981-12173981 | 1012 | 1026 | + | 15.771 | 1.11E-06 | 0.148 | CACACGGTCACACTG |
| 1 | GMTACGGTCACACTG | chrX:15996016-15998016 | 1013 | 1027 | + | 15.771 | 1.11E-06 | 0.148 | AATACGGCCACACTG |
| 1 | GMTACGGTCACACTG | chr3R:21072319-21074319 | 1001 | 1015 | - | 15.771 | 1.11E-06 | 0.148 | GATACGTGCACACTG |
| 1 | GMTACGGTCACACTG | chr3R:22744620-22746620 | 1016 | 1030 | + | 15.771 | 1.11E-06 | 0.148 | CAAACGGTCACACTG |
| 1 | GMTACGGTCACACTG | chr3L:3218156-3220156 | 1021 | 1035 | + | 15.771 | 1.11E-06 | 0.148 | CAGACGGTCACACTG |
| 1 | GMTACGGTCACACTG | chr3L:8997184-8999184 | 1024 | 1038 | + | 15.771 | 1.11E-06 | 0.148 | GAGACGGTGACACTG |
| 1 | GMTACGGTCACACTG | chr3L:14050206-14052206 | 1025 | 1039 | + | 15.771 | 1.11E-06 | 0.148 | AAGACGGTCACACTG |
| 1 | GMTACGGTCACACTG | chr2R:11639941-11641941 | 1011 | 1025 | - | 15.771 | 1.11E-06 | 0.148 | GAGAGGGTCACACTG |
| 1 | GMTACGGTCACACTG | chr2R:16263652-16265652 | 1025 | 1039 | + | 15.771 | 1.11E-06 | 0.148 | CAGACGGTCACACTG |
| 1 | GMTACGGTCACACTG | chr2L:9911608-9913608 | 1026 | 1040 | + | 15.771 | 1.11E-06 | 0.148 | AATGCGGTCACACTG |
| 1 | GMTACGGTCACACTG | chr2L:10199974-10201974 | 1014 | 1028 | - | 15.771 | 1.11E-06 | 0.148 | CAAACGGTCACACTG |
| 1 | GMTACGGTCACACTG | chrX:16883365-16885365 | 1015 | 1029 | - | 15.771 | 1.11E-06 | 0.148 | GCTATGGTCACACTA |
| 1 | GMTACGGTCACACTG | chr2L:22168143-22170143 | 1025 | 1039 | - | 15.771 | 1.11E-06 | 0.148 | GAAACGGTCACACTT |
| 1 | GMTACGGTCACACTG | chr3L:9854285-9856285 | 1031 | 1045 | - | 15.771 | 1.11E-06 | 0.148 | GCTAAGGTCACACTA |
| 1 | GMTACGGTCACACTG | chrX:15802880-15804880 | 1039 | 1053 | - | 15.771 | 1.11E-06 | 0.148 | AATACGGTCACACCG |
| 1 | GMTACGGTCACACTG | chr3R:22744581-22746581 | 1055 | 1069 | + | 15.771 | 1.11E-06 | 0.148 | CAAACGGTCACACTG |
| 1 | GMTACGGTCACACTG | chr2R:16263619-16265619 | 1058 | 1072 | + | 15.771 | 1.11E-06 | 0.148 | CAGACGGTCACACTG |
| 1 | GMTACGGTCACACTG | chr3L:19585003-19587003 | 1059 | 1073 | + | 15.771 | 1.11E-06 | 0.148 | GAGACGTTCACACTG |
| 1 | GMTACGGTCACACTG | chrX:21362028-21364028 | 1061 | 1075 | + | 15.771 | 1.11E-06 | 0.148 | CATACGGTCACACGG |
| 1 | GMTACGGTCACACTG | chrX:21362028-21364028 | 1061 | 1075 | + | 15.771 | 1.11E-06 | 0.148 | CATACGGTCACACGG |
| 1 | GMTACGGTCACACTG | chr3L:360383-362383 | 1058 | 1072 | - | 15.771 | 1.11E-06 | 0.148 | TATTCGGTCACACTG |
| 1 | GMTACGGTCACACTG | chr3R:8307485-8309485 | 1144 | 1158 | - | 15.771 | 1.11E-06 | 0.148 | GCTACAGTCACATTG |
| 1 | GMTACGGTCACACTG | chr3R:7127087-7129087 | 1158 | 1172 | + | 15.771 | 1.11E-06 | 0.148 | CAAACGGTCACACTG |
| 1 | GMTACGGTCACACTG | chr3L:21500421-21502421 | 1161 | 1175 | + | 15.771 | 1.11E-06 | 0.148 | GCTACGGTGACAGTG |

| | | | | | | | | | |
|---|---|---|---|---|---|---|---|---|---|
| 1 | GMTACGGTCACACTG | chr3L:11108505-11110505 | 1181 | 1195 | + | 15.771 | 1.11E-06 | 0.148 | AAAACGGTCACACTG |
| 1 | GMTACGGTCACACTG | chr2L:6496053-6498053 | 1183 | 1197 | + | 15.771 | 1.11E-06 | 0.148 | GATCCGGTGACACTG |
| 1 | GMTACGGTCACACTG | chr3R:15833425-15835425 | 1182 | 1196 | - | 15.771 | 1.11E-06 | 0.148 | GAAATGGTCACACTG |
| 1 | GMTACGGTCACACTG | chr2L:16309726-16311726 | 1217 | 1231 | + | 15.771 | 1.11E-06 | 0.148 | GATGCGGTCACACTA |
| 1 | GMTACGGTCACACTG | chr3R:7127027-7129027 | 1218 | 1232 | + | 15.771 | 1.11E-06 | 0.148 | CAAACGGTCACACTG |
| 1 | GMTACGGTCACACTG | chr3L:8996989-8998989 | 1219 | 1233 | + | 15.771 | 1.11E-06 | 0.148 | GAGACGGTGACACTG |
| 1 | GMTACGGTCACACTG | chr3L:7322411-7324411 | 1219 | 1233 | + | 15.771 | 1.11E-06 | 0.148 | GACATGGTCACACTG |
| 1 | GMTACGGTCACACTG | chr3L:3902492-3904492 | 1223 | 1237 | + | 15.771 | 1.11E-06 | 0.148 | GATACGGACACATTG |
| 1 | GMTACGGTCACACTG | chr2R:20256390-20258390 | 1238 | 1252 | + | 15.771 | 1.11E-06 | 0.148 | AAAACGGTCACACTG |
| 1 | GMTACGGTCACACTG | chr3L:19585003-19587003 | 1225 | 1239 | - | 15.771 | 1.11E-06 | 0.148 | GAGATGGTCACACTG |
| 1 | GMTACGGTCACACTG | chr3L:22720203-22722203 | 1245 | 1259 | + | 15.771 | 1.11E-06 | 0.148 | GAGACGGTCAGACTG |
| 1 | GMTACGGTCACACTG | chr3L:3217920-3219920 | 1257 | 1271 | + | 15.771 | 1.11E-06 | 0.148 | CAGACGGTCACACTG |
| 1 | GMTACGGTCACACTG | chr3L:14244691-14246691 | 1262 | 1276 | - | 15.771 | 1.11E-06 | 0.148 | ACTACGGTCACACTA |
| 1 | GMTACGGTCACACTG | chr2L:8402280-8404280 | 1292 | 1306 | + | 15.771 | 1.11E-06 | 0.148 | CATACGGGCACACTG |
| 1 | GMTACGGTCACACTG | chr3R:24030827-24032827 | 1297 | 1311 | + | 15.771 | 1.11E-06 | 0.148 | AACACGGTCACACTG |
| 1 | GMTACGGTCACACTG | chr3L:11033802-11035802 | 1284 | 1298 | - | 15.771 | 1.11E-06 | 0.148 | GATACGCACACACTG |
| 1 | GMTACGGTCACACTG | chr3L:21033203-21035203 | 1299 | 1313 | + | 15.771 | 1.11E-06 | 0.148 | TCTATGGTCACACTG |
| 1 | GMTACGGTCACACTG | chr3R:9038471-9040471 | 1310 | 1324 | + | 15.771 | 1.11E-06 | 0.148 | CCTACGGTCACACTC |
| 1 | GMTACGGTCACACTG | chr2L:11994432-11996432 | 1311 | 1325 | + | 15.771 | 1.11E-06 | 0.148 | AATACGGTCCCACTG |
| 1 | GMTACGGTCACACTG | chr2L:11998340-12000340 | 1314 | 1328 | + | 15.771 | 1.11E-06 | 0.148 | AATACGGTCCCACTG |
| 1 | GMTACGGTCACACTG | chr2R:20869392-20871392 | 1334 | 1348 | + | 15.771 | 1.11E-06 | 0.148 | GAAAGGGTCACACTG |
| 1 | GMTACGGTCACACTG | chr2L:22168143-22170143 | 1347 | 1361 | + | 15.771 | 1.11E-06 | 0.148 | GCTAAGGTAACACTG |
| 1 | GMTACGGTCACACTG | chr2L:20376-22376 | 1366 | 1380 | - | 15.771 | 1.11E-06 | 0.148 | GACATGGTCACACTG |
| 1 | GMTACGGTCACACTG | chr3L:12749119-12751119 | 1382 | 1396 | + | 15.771 | 1.11E-06 | 0.148 | GATACGGTCGCACTA |
| 1 | GMTACGGTCACACTG | chr2L:12171586-12173586 | 1407 | 1421 | + | 15.771 | 1.11E-06 | 0.148 | CACACGGTCACACTG |
| 1 | GMTACGGTCACACTG | chr3L:12749056-12751056 | 1445 | 1459 | + | 15.771 | 1.11E-06 | 0.148 | GATACGGTCGCACTA |
| 1 | GMTACGGTCACACTG | chr2L:10819705-10821705 | 1480 | 1494 | - | 15.771 | 1.11E-06 | 0.148 | GCTATGGTCACAATG |
| 1 | GMTACGGTCACACTG | chr3L:11000489-11002489 | 1504 | 1518 | - | 15.771 | 1.11E-06 | 0.148 | GATACGCTCACGCTG |
| 1 | GMTACGGTCACACTG | chr3L:3902184-3904184 | 1531 | 1545 | + | 15.771 | 1.11E-06 | 0.148 | GATACGGACACATTG |
| 1 | GMTACGGTCACACTG | chr3R:7472222-7474222 | 1552 | 1566 | - | 15.771 | 1.11E-06 | 0.148 | TACACGGTCACACTG |
| 1 | GMTACGGTCACACTG | chr3R:16971236-16973236 | 1593 | 1607 | + | 15.771 | 1.11E-06 | 0.148 | GATACGGATACACTG |
| 1 | GMTACGGTCACACTG | chr3L:6188123-6190123 | 1603 | 1617 | + | 15.771 | 1.11E-06 | 0.148 | GAAACGGTCAGACTG |
| 1 | GMTACGGTCACACTG | chr3R:27947021-27949021 | 1664 | 1678 | + | 15.771 | 1.11E-06 | 0.148 | AATACGGTCACTCTG |

| | | | | | | | | | |
|---|---|---|---|---|---|---|---|---|---|
| 1 | GMTACGGTCACACTG | chr3R:15923187-15925187 | 1723 | 1737 | + | 15.771 | 1.11E-06 | 0.148 | GCTACGATCACATTG |
| 1 | GMTACGGTCACACTG | chr3L:23182913-23184913 | 1864 | 1878 | - | 15.771 | 1.11E-06 | 0.148 | GATACGCTCTCACTG |
| 1 | GMTACGGTCACACTG | chr2R:21141368-21143368 | 1932 | 1946 | + | 15.771 | 1.11E-06 | 0.148 | TAGACGGTCACACTG |
| 1 | GMTACGGTCACACTG | chr2L:10208744-10210744 | 1922 | 1936 | - | 15.771 | 1.11E-06 | 0.148 | GATCCGGTCACACTA |
| 1 | GMTACGGTCACACTG | chr2L:20720914-20722914 | 179 | 193 | + | 15.493 | 1.37E-06 | 0.17 | GATACCGTCACACTT |
| 1 | GMTACGGTCACACTG | chr2R:22102177-22104177 | 985 | 999 | + | 15.493 | 1.37E-06 | 0.17 | CATACGGTCACACTT |
| 1 | GMTACGGTCACACTG | chr2R:24762167-24764167 | 989 | 1003 | + | 15.493 | 1.37E-06 | 0.17 | CATACGGTCACACTT |
| 1 | GMTACGGTCACACTG | chr3R:24654160-24656160 | 993 | 1007 | + | 15.493 | 1.37E-06 | 0.17 | CATACGGTCATACTG |
| 1 | GMTACGGTCACACTG | chr2L:14361069-14363069 | 993 | 1007 | + | 15.493 | 1.37E-06 | 0.17 | AATACGGTCACACTT |
| 1 | GMTACGGTCACACTG | chr3R:23125422-23127422 | 1032 | 1046 | + | 15.493 | 1.37E-06 | 0.17 | AATACAGTCACACTG |
| 1 | GMTACGGTCACACTG | chr2R:24762094-24764094 | 1062 | 1076 | + | 15.493 | 1.37E-06 | 0.17 | CATACGGTCACACTT |
| 1 | GMTACGGTCACACTG | chr2R:21096305-21098305 | 1068 | 1082 | + | 15.493 | 1.37E-06 | 0.17 | TATACGGTCACACTA |
| 1 | GMTACGGTCACACTG | chr2L:5054058-5056058 | 1069 | 1083 | + | 15.493 | 1.37E-06 | 0.17 | TATAGGGTCACACTG |
| 1 | GMTACGGTCACACTG | chr2L:5054016-5056016 | 1111 | 1125 | + | 15.493 | 1.37E-06 | 0.17 | TATAGGGTCACACTG |
| 1 | GMTACGGTCACACTG | chr2R:20423338-20425338 | 1237 | 1251 | + | 15.493 | 1.37E-06 | 0.17 | GATAAGGTCAAACTG |
| 1 | GMTACGGTCACACTG | chr3R:24653820-24655820 | 1333 | 1347 | + | 15.493 | 1.37E-06 | 0.17 | CATACGGTCATACTG |
| 1 | GMTACGGTCACACTG | chr2L:14360303-14362303 | 1759 | 1773 | + | 15.493 | 1.37E-06 | 0.17 | AATACGGTCACACTT |
| 1 | GMTACGGTCACACTG | chr3R:5624367-5626367 | 16 | 30 | - | 15.493 | 1.37E-06 | 0.17 | TATACGGTCACATTG |
| 1 | GMTACGGTCACACTG | chr2L:15060656-15062656 | 125 | 139 | - | 15.493 | 1.37E-06 | 0.17 | AATACGGTAACACTG |
| 1 | GMTACGGTCACACTG | chr2L:15060656-15062656 | 125 | 139 | - | 15.493 | 1.37E-06 | 0.17 | AATACGGTAACACTG |
| 1 | GMTACGGTCACACTG | chr3R:5624178-5626178 | 205 | 219 | - | 15.493 | 1.37E-06 | 0.17 | TATACGGTCACATTG |
| 1 | GMTACGGTCACACTG | chr3L:17015536-17017536 | 479 | 493 | - | 15.493 | 1.37E-06 | 0.17 | TATACGGTCACACTT |
| 1 | GMTACGGTCACACTG | chr3L:5779051-5781051 | 525 | 539 | - | 15.493 | 1.37E-06 | 0.17 | GATATGGTCACACTC |
| 1 | GMTACGGTCACACTG | chr3L:5779051-5781051 | 525 | 539 | - | 15.493 | 1.37E-06 | 0.17 | GATATGGTCACACTC |
| 1 | GMTACGGTCACACTG | chrX:7932742-7934742 | 596 | 610 | - | 15.493 | 1.37E-06 | 0.17 | CATATGGTCACACTG |
| 1 | GMTACGGTCACACTG | chrX:1379914-1381914 | 725 | 739 | - | 15.493 | 1.37E-06 | 0.17 | GATACGCTTACACTG |
| 1 | GMTACGGTCACACTG | chrX:9246342-9248342 | 793 | 807 | - | 15.493 | 1.37E-06 | 0.17 | TATACGGTCACACTC |
| 1 | GMTACGGTCACACTG | chrX:9246336-9248336 | 799 | 813 | - | 15.493 | 1.37E-06 | 0.17 | TATACGGTCACACTC |
| 1 | GMTACGGTCACACTG | chr3L:17015180-17017180 | 835 | 849 | - | 15.493 | 1.37E-06 | 0.17 | TATACGGTCACACTT |
| 1 | GMTACGGTCACACTG | chrX:4105113-4107113 | 872 | 886 | - | 15.493 | 1.37E-06 | 0.17 | TATATGGTCACACTG |
| 1 | GMTACGGTCACACTG | chr2R:12759999-12761999 | 891 | 905 | - | 15.493 | 1.37E-06 | 0.17 | CATACGTTCACACTG |
| 1 | GMTACGGTCACACTG | chr3R:24624671-24626671 | 892 | 906 | - | 15.493 | 1.37E-06 | 0.17 | CATAAGGTCACACTG |
| 1 | GMTACGGTCACACTG | chr3R:5623474-5625474 | 909 | 923 | - | 15.493 | 1.37E-06 | 0.17 | TATACGGTCACATTG |

| | | | | | | | | | |
|---|---|---|---|---|---|---|---|---|---|
| 1 | GMTACGGTCACACTG | chr3L:5778591-5780591 | 985 | 999 | - | 15.493 | 1.37E-06 | 0.17 | GATATGGTCACACTC |
| 1 | GMTACGGTCACACTG | chrX:7932351-7934351 | 987 | 1001 | - | 15.493 | 1.37E-06 | 0.17 | CATATGGTCACACTG |
| 1 | GMTACGGTCACACTG | chr2R:17732238-17734238 | 1001 | 1015 | - | 15.493 | 1.37E-06 | 0.17 | AATATGGTCACACTG |
| 1 | GMTACGGTCACACTG | chr3R:24624562-24626562 | 1001 | 1015 | - | 15.493 | 1.37E-06 | 0.17 | CATAAGGTCACACTG |
| 1 | GMTACGGTCACACTG | chr3L:13389006-13391006 | 1012 | 1026 | - | 15.493 | 1.37E-06 | 0.17 | AATACGGTTACACTG |
| 1 | GMTACGGTCACACTG | chr2L:15059757-15061757 | 1024 | 1038 | - | 15.493 | 1.37E-06 | 0.17 | AATACGGTAACACTG |
| 1 | GMTACGGTCACACTG | chrX:1379480-1381480 | 1159 | 1173 | - | 15.493 | 1.37E-06 | 0.17 | GATACGCTTACACTG |
| 1 | GMTACGGTCACACTG | chr3R:5623125-5625125 | 1258 | 1272 | - | 15.493 | 1.37E-06 | 0.17 | TATACGGTCACATTG |
| 1 | GMTACGGTCACACTG | chr3R:5623125-5625125 | 1258 | 1272 | - | 15.493 | 1.37E-06 | 0.17 | TATACGGTCACATTG |
| 1 | GMTACGGTCACACTG | chr3R:30624592-30626592 | 217 | 231 | - | 10.821 | 1.52E-06 | 0.176 | GTTACGGACACGCTG |
| 1 | GMTACGGTCACACTG | chr3R:16612400-16614400 | 452 | 466 | + | 10.821 | 1.52E-06 | 0.176 | GGTCCGGCCACACTG |
| 1 | GMTACGGTCACACTG | chr2R:8599415-8601415 | 522 | 536 | - | 10.821 | 1.52E-06 | 0.176 | GGCACGGTCACGCTG |
| 1 | GMTACGGTCACACTG | chr3R:24718697-24720697 | 642 | 656 | - | 10.821 | 1.52E-06 | 0.176 | GGTGCGGTCACGCTG |
| 1 | GMTACGGTCACACTG | chr3L:3471447-3473447 | 661 | 675 | - | 10.821 | 1.52E-06 | 0.176 | GGAACGGTCACTCTG |
| 1 | GMTACGGTCACACTG | chr3R:21218077-21220077 | 685 | 699 | - | 10.821 | 1.52E-06 | 0.176 | GGTGCGGCCACACTG |
| 1 | GMTACGGTCACACTG | chr3R:9245649-9247649 | 724 | 738 | + | 10.821 | 1.52E-06 | 0.176 | GTAACGGGCACACTG |
| 1 | GMTACGGTCACACTG | chr3R:28844270-28846270 | 881 | 895 | - | 10.821 | 1.52E-06 | 0.176 | GTGACGGTCACACCG |
| 1 | GMTACGGTCACACTG | chr3R:17810541-17812541 | 922 | 936 | - | 10.821 | 1.52E-06 | 0.176 | GGTACGGTCACTCGG |
| 1 | GMTACGGTCACACTG | chrX:17656372-17658372 | 939 | 953 | + | 10.821 | 1.52E-06 | 0.176 | GGCACGGCCACACTG |
| 1 | GMTACGGTCACACTG | chrX:19597031-19599031 | 968 | 982 | + | 10.821 | 1.52E-06 | 0.176 | GGTCCGGCCACACTG |
| 1 | GMTACGGTCACACTG | chr3R:12407316-12409316 | 972 | 986 | + | 10.821 | 1.52E-06 | 0.176 | GGCACGGTCACACCG |
| 1 | GMTACGGTCACACTG | chr3R:31060210-31062210 | 962 | 976 | - | 10.821 | 1.52E-06 | 0.176 | GGCGCGGTCACACTG |
| 1 | GMTACGGTCACACTG | chr2L:418952-420952 | 977 | 991 | + | 10.821 | 1.52E-06 | 0.176 | GGCGCGGTCACACTG |
| 1 | GMTACGGTCACACTG | chr3R:21217791-21219791 | 971 | 985 | - | 10.821 | 1.52E-06 | 0.176 | GGTGCGGCCACACTG |
| 1 | GMTACGGTCACACTG | chr2L:2579292-2581292 | 985 | 999 | + | 10.821 | 1.52E-06 | 0.176 | GGTTCGGCCACACTG |
| 1 | GMTACGGTCACACTG | chr3R:14285903-14287903 | 974 | 988 | - | 10.821 | 1.52E-06 | 0.176 | GGCGCGGTCACACTG |
| 1 | GMTACGGTCACACTG | chr2R:8598962-8600962 | 975 | 989 | - | 10.821 | 1.52E-06 | 0.176 | GGCACGGTCACGCTG |
| 1 | GMTACGGTCACACTG | chr3R:24718364-24720364 | 975 | 989 | - | 10.821 | 1.52E-06 | 0.176 | GGTGCGGTCACGCTG |
| 1 | GMTACGGTCACACTG | chr3L:3164181-3166181 | 993 | 1007 | + | 10.821 | 1.52E-06 | 0.176 | GTAACGGTCACTCTG |
| 1 | GMTACGGTCACACTG | chr3L:11997138-11999138 | 979 | 993 | - | 10.821 | 1.52E-06 | 0.176 | GGTGCGGTCACACCG |
| 1 | GMTACGGTCACACTG | chr3L:2651088-2653088 | 993 | 1007 | + | 10.821 | 1.52E-06 | 0.176 | GGCCCGGTCACACTG |
| 1 | GMTACGGTCACACTG | chr3L:2651088-2653088 | 993 | 1007 | + | 10.821 | 1.52E-06 | 0.176 | GGCCCGGTCACACTG |
| 1 | GMTACGGTCACACTG | chr2R:9284097-9286097 | 981 | 995 | - | 10.821 | 1.52E-06 | 0.176 | GGCGCGGTCACACTG |

| | | | | | | | | | |
|---|---|---|---|---|---|---|---|---|---|
| 1 | GMTACGGTCACACTG | chr3R:16049801-16051801 | 1006 | 1020 | + | 10.821 | 1.52E-06 | 0.176 | GTAGCGGTCACACTG |
| 1 | GMTACGGTCACACTG | chr3L:3471113-3473113 | 995 | 1009 | - | 10.821 | 1.52E-06 | 0.176 | GGAACGGTCACTCTG |
| 1 | GMTACGGTCACACTG | chrX:21344358-21346358 | 1020 | 1034 | + | 10.821 | 1.52E-06 | 0.176 | GGCGCGGTCACACTG |
| 1 | GMTACGGTCACACTG | chr2L:6851315-6853315 | 1024 | 1038 | + | 10.821 | 1.52E-06 | 0.176 | GGGGCGGTCACACTG |
| 1 | GMTACGGTCACACTG | chr3R:23722331-23724331 | 1027 | 1041 | + | 10.821 | 1.52E-06 | 0.176 | GGGACGGTCGCACTG |
| 1 | GMTACGGTCACACTG | chr2L:2579172-2581172 | 1105 | 1119 | + | 10.821 | 1.52E-06 | 0.176 | GGTTCGGCCACACTG |
| 1 | GMTACGGTCACACTG | chr3L:2650850-2652850 | 1231 | 1245 | + | 10.821 | 1.52E-06 | 0.176 | GGCCCGGTCACACTG |
| 1 | GMTACGGTCACACTG | chr2R:9283845-9285845 | 1233 | 1247 | - | 10.821 | 1.52E-06 | 0.176 | GGCGCGGTCACACTG |
| 1 | GMTACGGTCACACTG | chr2L:418652-420652 | 1277 | 1291 | + | 10.821 | 1.52E-06 | 0.176 | GGCGCGGTCACACTG |
| 1 | GMTACGGTCACACTG | chr3R:17810134-17812134 | 1329 | 1343 | - | 10.821 | 1.52E-06 | 0.176 | GGTACGGTCACTCGG |
| 1 | GMTACGGTCACACTG | chr3R:28843748-28845748 | 1403 | 1417 | - | 10.821 | 1.52E-06 | 0.176 | GTGACGGTCACACCG |
| 1 | GMTACGGTCACACTG | chrX:17655818-17657818 | 1493 | 1507 | + | 10.821 | 1.52E-06 | 0.176 | GGCACGGCCACACTG |
| 1 | GMTACGGTCACACTG | chr3R:9244869-9246869 | 1504 | 1518 | + | 10.821 | 1.52E-06 | 0.176 | GTAACGGGCACACTG |
| 1 | GMTACGGTCACACTG | chr2R:21170001-21172001 | 1515 | 1529 | + | 10.821 | 1.52E-06 | 0.176 | GTTACGGACTCACTG |
| 1 | GMTACGGTCACACTG | chr3L:3470309-3472309 | 1799 | 1813 | - | 10.821 | 1.52E-06 | 0.176 | GGAACGGTCACTCTG |
| 1 | GMTACGGTCACACTG | chr2R:21169672-21171672 | 1844 | 1858 | + | 10.821 | 1.52E-06 | 0.176 | GTTACGGACTCACTG |
| 1 | GMTACGGTCACACTG | chr3R:30033907-30035907 | 1937 | 1951 | + | 10.821 | 1.52E-06 | 0.176 | GTTCCGGGCACACTG |
| 1 | GMTACGGTCACACTG | chr2L:18199336-18201336 | 1975 | 1989 | + | 10.821 | 1.52E-06 | 0.176 | GGCACGGCCACACTG |
| 1 | GMTACGGTCACACTG | chrX:9329487-9331487 | 122 | 136 | + | 10.543 | 2.14E-06 | 0.191 | GGTCAGGTCACACTG |
| 1 | GMTACGGTCACACTG | chr2R:14617118-14619118 | 203 | 217 | + | 10.543 | 2.14E-06 | 0.191 | GGTACGGTCAGCCTG |
| 1 | GMTACGGTCACACTG | chr3R:27272273-27274273 | 223 | 237 | + | 10.543 | 2.14E-06 | 0.191 | AGTACGGCCACACTG |
| 1 | GMTACGGTCACACTG | chrX:18136784-18138784 | 371 | 385 | + | 10.543 | 2.14E-06 | 0.191 | AGAACGGTCACACTG |
| 1 | GMTACGGTCACACTG | chrX:475984-477984 | 457 | 471 | + | 10.543 | 2.14E-06 | 0.191 | GTTACGGTCACTGTG |
| 1 | GMTACGGTCACACTG | chrX:18656755-18658755 | 570 | 584 | + | 10.543 | 2.14E-06 | 0.191 | GGTTTGGTCACACTG |
| 1 | GMTACGGTCACACTG | chr2R:13167498-13169498 | 578 | 592 | + | 10.543 | 2.14E-06 | 0.191 | CGCACGGTCACACTG |
| 1 | GMTACGGTCACACTG | chrX:10330818-10332818 | 607 | 621 | + | 10.543 | 2.14E-06 | 0.191 | GTGACGGTCACATTG |
| 1 | GMTACGGTCACACTG | chr2L:8365038-8367038 | 656 | 670 | + | 10.543 | 2.14E-06 | 0.191 | GTTATGGTCACACAG |
| 1 | GMTACGGTCACACTG | chr3R:24056310-24058310 | 693 | 707 | + | 10.543 | 2.14E-06 | 0.191 | GGTACGGTTACCCTG |
| 1 | GMTACGGTCACACTG | chr2R:24880817-24882817 | 745 | 759 | + | 10.543 | 2.14E-06 | 0.191 | AGCACGGTCACACTG |
| 1 | GMTACGGTCACACTG | chr2R:13823858-13825858 | 787 | 801 | + | 10.543 | 2.14E-06 | 0.191 | GTTCCGTTCACACTG |
| 1 | GMTACGGTCACACTG | chr2L:6064997-6066997 | 807 | 821 | + | 10.543 | 2.14E-06 | 0.191 | GTTACGGCCAAACTG |
| 1 | GMTACGGTCACACTG | chr2R:23329068-23331068 | 815 | 829 | + | 10.543 | 2.14E-06 | 0.191 | GTTATGGTCTCACTG |
| 1 | GMTACGGTCACACTG | chr2L:6064953-6066953 | 851 | 865 | + | 10.543 | 2.14E-06 | 0.191 | GTTACGGCCAAACTG |

| | | | | | | | | | |
|---|---|---|---|---|---|---|---|---|---|
| 1 | GMTACGGTCACACTG | chrX:475579-477579 | 862 | 876 | + | 10.543 | 2.14E-06 | 0.191 | GTTACGGTCACTGTG |
| 1 | GMTACGGTCACACTG | chr2R:9841686-9843686 | 876 | 890 | + | 10.543 | 2.14E-06 | 0.191 | CGCACGGTCACACTG |
| 1 | GMTACGGTCACACTG | chrX:19200979-19202979 | 928 | 942 | + | 10.543 | 2.14E-06 | 0.191 | GGCAGGGTCACACTG |
| 1 | GMTACGGTCACACTG | chr3R:21126573-21128573 | 934 | 948 | + | 10.543 | 2.14E-06 | 0.191 | TGAACGGTCACACTG |
| 1 | GMTACGGTCACACTG | chrX:6224647-6226647 | 954 | 968 | + | 10.543 | 2.14E-06 | 0.191 | GGTATGGACACACTG |
| 1 | GMTACGGTCACACTG | chr3R:31212458-31214458 | 959 | 973 | + | 10.543 | 2.14E-06 | 0.191 | GTGAAGGTCACACTG |
| 1 | GMTACGGTCACACTG | chr2R:11306879-11308879 | 969 | 983 | + | 10.543 | 2.14E-06 | 0.191 | TTTTCGGTCACACTG |
| 1 | GMTACGGTCACACTG | chrX:11815898-11817898 | 973 | 987 | + | 10.543 | 2.14E-06 | 0.191 | TGCACGGTCACACTG |
| 1 | GMTACGGTCACACTG | chrX:6300285-6302285 | 978 | 992 | + | 10.543 | 2.14E-06 | 0.191 | GGCACGGTCACACTT |
| 1 | GMTACGGTCACACTG | chr3L:18744856-18746856 | 986 | 1000 | + | 10.543 | 2.14E-06 | 0.191 | GTTCTGGTCACACTG |
| 1 | GMTACGGTCACACTG | chr3L:11213505-11215505 | 987 | 1001 | + | 10.543 | 2.14E-06 | 0.191 | GGCACGGTCACACTC |
| 1 | GMTACGGTCACACTG | chr3L:3895900-3897900 | 989 | 1003 | + | 10.543 | 2.14E-06 | 0.191 | AGTCCGGTCACACTG |
| 1 | GMTACGGTCACACTG | chr3R:13010649-13012649 | 989 | 1003 | + | 10.543 | 2.14E-06 | 0.191 | TGTGCGGTCACACTG |
| 1 | GMTACGGTCACACTG | chr3L:8544001-8546001 | 990 | 1004 | + | 10.543 | 2.14E-06 | 0.191 | CGCACGGTCACACTG |
| 1 | GMTACGGTCACACTG | chrX:9141493-9143493 | 992 | 1006 | + | 10.543 | 2.14E-06 | 0.191 | CGGACGGTCACACTG |
| 1 | GMTACGGTCACACTG | chr2R:24880570-24882570 | 992 | 1006 | + | 10.543 | 2.14E-06 | 0.191 | AGCACGGTCACACTG |
| 1 | GMTACGGTCACACTG | chr2R:8420033-8422033 | 993 | 1007 | + | 10.543 | 2.14E-06 | 0.191 | TGAACGGTCACACTG |
| 1 | GMTACGGTCACACTG | chr3R:14121654-14123654 | 993 | 1007 | + | 10.543 | 2.14E-06 | 0.191 | TTTGCGGTCACACTG |
| 1 | GMTACGGTCACACTG | chr3L:14052215-14054215 | 996 | 1010 | + | 10.543 | 2.14E-06 | 0.191 | AGTTCGGTCACACTG |
| 1 | GMTACGGTCACACTG | chr3R:10261123-10263123 | 996 | 1010 | + | 10.543 | 2.14E-06 | 0.191 | CGCACGGTCACACTG |
| 1 | GMTACGGTCACACTG | chr3R:29724830-29726830 | 1001 | 1015 | + | 10.543 | 2.14E-06 | 0.191 | GGTGTGGTCACACTG |
| 1 | GMTACGGTCACACTG | chr2L:10506081-10508081 | 1003 | 1017 | + | 10.543 | 2.14E-06 | 0.191 | AGTTCGGTCACACTG |
| 1 | GMTACGGTCACACTG | chr2R:12632589-12634589 | 1004 | 1018 | + | 10.543 | 2.14E-06 | 0.191 | AGTACGGTCACTCTG |
| 1 | GMTACGGTCACACTG | chr2L:10221657-10223657 | 1004 | 1018 | + | 10.543 | 2.14E-06 | 0.191 | AGTTCGGTCACACTG |
| 1 | GMTACGGTCACACTG | chr2R:22173922-22175922 | 1005 | 1019 | + | 10.543 | 2.14E-06 | 0.191 | AGCACGGTCACACTG |
| 1 | GMTACGGTCACACTG | chr3R:5354092-5356092 | 1007 | 1021 | + | 10.543 | 2.14E-06 | 0.191 | AGTACGGTCACTCTG |
| 1 | GMTACGGTCACACTG | chr2R:8892613-8894613 | 1007 | 1021 | + | 10.543 | 2.14E-06 | 0.191 | AGCACGGTCACACTG |
| 1 | GMTACGGTCACACTG | chr3R:4484008-4486008 | 1008 | 1022 | + | 10.543 | 2.14E-06 | 0.191 | GGAATGGTCACACTG |
| 1 | GMTACGGTCACACTG | chr4:307093-309093 | 1010 | 1024 | + | 10.543 | 2.14E-06 | 0.191 | TGTGCGGTCACACTG |
| 1 | GMTACGGTCACACTG | chr2L:19053078-19055078 | 1010 | 1024 | + | 10.543 | 2.14E-06 | 0.191 | TTAACGGTCACACTG |
| 1 | GMTACGGTCACACTG | chr2L:6322767-6324767 | 1010 | 1024 | + | 10.543 | 2.14E-06 | 0.191 | GGTGTGGTCACACTG |
| 1 | GMTACGGTCACACTG | chr2R:23701477-23703477 | 1014 | 1028 | + | 10.543 | 2.14E-06 | 0.191 | AGCACGGTCACACTG |
| 1 | GMTACGGTCACACTG | chr2L:3448683-3450683 | 1015 | 1029 | + | 10.543 | 2.14E-06 | 0.191 | GGTCTGGTCACACTG |

| | | | | | | | | | |
|---|---|---|---|---|---|---|---|---|---|
| 1 | GMTACGGTCACACTG | chr2R:10837727-10839727 | 1015 | 1029 | + | 10.543 | 2.14E-06 | 0.191 | CTTGCGGTCACACTG |
| 1 | GMTACGGTCACACTG | chr3R:9558760-9560760 | 1015 | 1029 | + | 10.543 | 2.14E-06 | 0.191 | AGTACGGCCACACTG |
| 1 | GMTACGGTCACACTG | chrX:15785047-15787047 | 1018 | 1032 | + | 10.543 | 2.14E-06 | 0.191 | CGTGCGGTCACACTG |
| 1 | GMTACGGTCACACTG | chr3L:21616623-21618623 | 1019 | 1033 | + | 10.543 | 2.14E-06 | 0.191 | GTTCTGGTCACACTG |
| 1 | GMTACGGTCACACTG | chr3L:6192480-6194480 | 1019 | 1033 | + | 10.543 | 2.14E-06 | 0.191 | TGCACGGTCACACTG |
| 1 | GMTACGGTCACACTG | chr3L:16089454-16091454 | 1021 | 1035 | + | 10.543 | 2.14E-06 | 0.191 | GGTCTGGTCACACTG |
| 1 | GMTACGGTCACACTG | chr3R:21126485-21128485 | 1022 | 1036 | + | 10.543 | 2.14E-06 | 0.191 | TGAACGGTCACACTG |
| 1 | GMTACGGTCACACTG | chr3L:1864430-1866430 | 1025 | 1039 | + | 10.543 | 2.14E-06 | 0.191 | GGAACGGTCACATTG |
| 1 | GMTACGGTCACACTG | chr3L:11792514-11794514 | 1027 | 1041 | + | 10.543 | 2.14E-06 | 0.191 | AGTGCGGTCACACTG |
| 1 | GMTACGGTCACACTG | chr2R:23066196-23068196 | 1028 | 1042 | + | 10.543 | 2.14E-06 | 0.191 | CGAACGGTCACACTG |
| 1 | GMTACGGTCACACTG | chrX:18656286-18658286 | 1039 | 1053 | + | 10.543 | 2.14E-06 | 0.191 | GGTTTGGTCACACTG |
| 1 | GMTACGGTCACACTG | chr3R:31212376-31214376 | 1041 | 1055 | + | 10.543 | 2.14E-06 | 0.191 | GTGAAGGTCACACTG |
| 1 | GMTACGGTCACACTG | chr2R:13167032-13169032 | 1044 | 1058 | + | 10.543 | 2.14E-06 | 0.191 | CGCACGGTCACACTG |
| 1 | GMTACGGTCACACTG | chr2R:11296520-11298520 | 1048 | 1062 | + | 10.543 | 2.14E-06 | 0.191 | GGTCCGGTCACACTA |
| 1 | GMTACGGTCACACTG | chrX:18787464-18789464 | 1057 | 1071 | + | 10.543 | 2.14E-06 | 0.191 | GGTCTGGTCACACTG |
| 1 | GMTACGGTCACACTG | chr2R:9841502-9843502 | 1060 | 1074 | + | 10.543 | 2.14E-06 | 0.191 | CGCACGGTCACACTG |
| 1 | GMTACGGTCACACTG | chr3L:318884-320884 | 1070 | 1084 | + | 10.543 | 2.14E-06 | 0.191 | TGTGCGGTCACACTG |
| 1 | GMTACGGTCACACTG | chr3R:15094631-15096631 | 1070 | 1084 | + | 10.543 | 2.14E-06 | 0.191 | GTTGCGGTCACACTT |
| 1 | GMTACGGTCACACTG | chr2R:24268179-24270179 | 1075 | 1089 | + | 10.543 | 2.14E-06 | 0.191 | GTTAAGGCCACACTG |
| 1 | GMTACGGTCACACTG | chr3L:16089392-16091392 | 1083 | 1097 | + | 10.543 | 2.14E-06 | 0.191 | GGTCTGGTCACACTG |
| 1 | GMTACGGTCACACTG | chr2R:23701406-23703406 | 1085 | 1099 | + | 10.543 | 2.14E-06 | 0.191 | AGCACGGTCACACTG |
| 1 | GMTACGGTCACACTG | chr2R:13823553-13825553 | 1092 | 1106 | + | 10.543 | 2.14E-06 | 0.191 | GTTCCGTTCACACTG |
| 1 | GMTACGGTCACACTG | chrX:4103342-4105342 | 1103 | 1117 | + | 10.543 | 2.14E-06 | 0.191 | ATCACGGTCACACTG |
| 1 | GMTACGGTCACACTG | chr3L:14052079-14054079 | 1132 | 1146 | + | 10.543 | 2.14E-06 | 0.191 | AGTTCGGTCACACTG |
| 1 | GMTACGGTCACACTG | chr3R:21137270-21139270 | 1142 | 1156 | + | 10.543 | 2.14E-06 | 0.191 | CTAACGGTCACACTG |
| 1 | GMTACGGTCACACTG | chr3L:1864240-1866240 | 1215 | 1229 | + | 10.543 | 2.14E-06 | 0.191 | GGAACGGTCACATTG |
| 1 | GMTACGGTCACACTG | chr3R:14121420-14123420 | 1227 | 1241 | + | 10.543 | 2.14E-06 | 0.191 | TTTGCGGTCACACTG |
| 1 | GMTACGGTCACACTG | chr2R:22173691-22175691 | 1236 | 1250 | + | 10.543 | 2.14E-06 | 0.191 | AGCACGGTCACACTG |
| 1 | GMTACGGTCACACTG | chrX:4103197-4105197 | 1248 | 1262 | + | 10.543 | 2.14E-06 | 0.191 | ATCACGGTCACACTG |
| 1 | GMTACGGTCACACTG | chr3R:9558525-9560525 | 1250 | 1264 | + | 10.543 | 2.14E-06 | 0.191 | AGTACGGCCACACTG |
| 1 | GMTACGGTCACACTG | chr3L:11792283-11794283 | 1258 | 1272 | + | 10.543 | 2.14E-06 | 0.191 | AGTGCGGTCACACTG |
| 1 | GMTACGGTCACACTG | chr3L:21616376-21618376 | 1266 | 1280 | + | 10.543 | 2.14E-06 | 0.191 | GTTCTGGTCACACTG |
| 1 | GMTACGGTCACACTG | chr3R:10260849-10262849 | 1270 | 1284 | + | 10.543 | 2.14E-06 | 0.191 | CGCACGGTCACACTG |

| | | | | | | | | | |
|---|---|---|---|---|---|---|---|---|---|
| 1 | GMTACGGTCACACTG | chr3L:3895603-3897603 | 1286 | 1300 | + | 10.543 | 2.14E-06 | 0.191 | AGTCCGGTCACACTG |
| 1 | GMTACGGTCACACTG | chr2R:24113398-24115398 | 1298 | 1312 | + | 10.543 | 2.14E-06 | 0.191 | AGTGCGGTCACACTG |
| 1 | GMTACGGTCACACTG | chr2L:10505779-10507779 | 1305 | 1319 | + | 10.543 | 2.14E-06 | 0.191 | AGTTCGGTCACACTG |
| 1 | GMTACGGTCACACTG | chrX:18787210-18789210 | 1311 | 1325 | + | 10.543 | 2.14E-06 | 0.191 | GGTCTGGTCACACTG |
| 1 | GMTACGGTCACACTG | chrX:11815551-11817551 | 1320 | 1334 | + | 10.543 | 2.14E-06 | 0.191 | TGCACGGTCACACTG |
| 1 | GMTACGGTCACACTG | chr3L:6192179-6194179 | 1320 | 1334 | + | 10.543 | 2.14E-06 | 0.191 | TGCACGGTCACACTG |
| 1 | GMTACGGTCACACTG | chr2R:11306528-11308528 | 1320 | 1334 | + | 10.543 | 2.14E-06 | 0.191 | TTTTCGGTCACACTG |
| 1 | GMTACGGTCACACTG | chr3R:13010310-13012310 | 1328 | 1342 | + | 10.543 | 2.14E-06 | 0.191 | TGTGCGGTCACACTG |
| 1 | GMTACGGTCACACTG | chr3R:5353747-5355747 | 1352 | 1366 | + | 10.543 | 2.14E-06 | 0.191 | AGTACGGTCACTCTG |
| 1 | GMTACGGTCACACTG | chr2R:24113292-24115292 | 1404 | 1418 | + | 10.543 | 2.14E-06 | 0.191 | AGTGCGGTCACACTG |
| 1 | GMTACGGTCACACTG | chrX:6224140-6226140 | 1461 | 1475 | + | 10.543 | 2.14E-06 | 0.191 | GGTATGGACACACTG |
| 1 | GMTACGGTCACACTG | chrX:15784583-15786583 | 1482 | 1496 | + | 10.543 | 2.14E-06 | 0.191 | CGTGCGGTCACACTG |
| 1 | GMTACGGTCACACTG | chrX:9140965-9142965 | 1520 | 1534 | + | 10.543 | 2.14E-06 | 0.191 | CGGACGGTCACACTG |
| 1 | GMTACGGTCACACTG | chrX:16349315-16351315 | 1679 | 1693 | + | 10.543 | 2.14E-06 | 0.191 | GTTAAGGACACACTG |
| 1 | GMTACGGTCACACTG | chr2L:10505301-10507301 | 1783 | 1797 | + | 10.543 | 2.14E-06 | 0.191 | AGTTCGGTCACACTG |
| 1 | GMTACGGTCACACTG | chr3R:29724027-29726027 | 1804 | 1818 | + | 10.543 | 2.14E-06 | 0.191 | GGTGTGGTCACACTG |
| 1 | GMTACGGTCACACTG | chr2L:8363855-8365855 | 1839 | 1853 | + | 10.543 | 2.14E-06 | 0.191 | GTTATGGTCACACAG |
| 1 | GMTACGGTCACACTG | chr3R:29723950-29725950 | 1881 | 1895 | + | 10.543 | 2.14E-06 | 0.191 | GGTGTGGTCACACTG |
| 1 | GMTACGGTCACACTG | chr2R:12631674-12633674 | 1919 | 1933 | + | 10.543 | 2.14E-06 | 0.191 | AGTACGGTCACTCTG |
| 1 | GMTACGGTCACACTG | chr3L:1246817-1248817 | 80 | 94 | - | 10.543 | 2.14E-06 | 0.191 | GTTGCGGTCACACTA |
| 1 | GMTACGGTCACACTG | chr2R:24875691-24877691 | 165 | 179 | - | 10.543 | 2.14E-06 | 0.191 | GTTTCGGTCATACTG |
| 1 | GMTACGGTCACACTG | chr3L:1246677-1248677 | 220 | 234 | - | 10.543 | 2.14E-06 | 0.191 | GTTGCGGTCACACTA |
| 1 | GMTACGGTCACACTG | chr3L:3460317-3462317 | 247 | 261 | - | 10.543 | 2.14E-06 | 0.191 | CGCACGGTCACACTG |
| 1 | GMTACGGTCACACTG | chrX:11148323-11150323 | 282 | 296 | - | 10.543 | 2.14E-06 | 0.191 | CGCACGGTCACACTG |
| 1 | GMTACGGTCACACTG | chr3R:30427820-30429820 | 340 | 354 | - | 10.543 | 2.14E-06 | 0.191 | GGCATGGTCACACTG |
| 1 | GMTACGGTCACACTG | chrX:10082154-10084154 | 362 | 376 | - | 10.543 | 2.14E-06 | 0.191 | AGCACGGTCACACTG |
| 1 | GMTACGGTCACACTG | chr2L:15750682-15752682 | 402 | 416 | - | 10.543 | 2.14E-06 | 0.191 | AGTGCGGTCACACTG |
| 1 | GMTACGGTCACACTG | chr2L:19795365-19797365 | 450 | 464 | - | 10.543 | 2.14E-06 | 0.191 | CGAACGGTCACACTG |
| 1 | GMTACGGTCACACTG | chrX:18656755-18658755 | 526 | 540 | - | 10.543 | 2.14E-06 | 0.191 | GGTACGGTCACCCTA |
| 1 | GMTACGGTCACACTG | chr3L:4363550-4365550 | 534 | 548 | - | 10.543 | 2.14E-06 | 0.191 | ATTGCGGTCACACTG |
| 1 | GMTACGGTCACACTG | chr2L:8082420-8084420 | 550 | 564 | - | 10.543 | 2.14E-06 | 0.191 | CGAACGGTCACACTG |
| 1 | GMTACGGTCACACTG | chr3L:3459977-3461977 | 587 | 601 | - | 10.543 | 2.14E-06 | 0.191 | CGCACGGTCACACTG |
| 1 | GMTACGGTCACACTG | chr2L:3158950-3160950 | 676 | 690 | - | 10.543 | 2.14E-06 | 0.191 | GTCATGGTCACACTG |

| | | | | | | | | | |
|---|---|---|---|---|---|---|---|---|---|
| 1 | GMTACGGTCACACTG | chrX:13328824-13330824 | 680 | 694 | - | 10.543 | 2.14E-06 | 0.191 | TTAACGGTCACACTG |
| 1 | GMTACGGTCACACTG | chrX:940075-942075 | 723 | 737 | - | 10.543 | 2.14E-06 | 0.191 | AGGACGGTCACACTG |
| 1 | GMTACGGTCACACTG | chr2L:13191201-13193201 | 752 | 766 | - | 10.543 | 2.14E-06 | 0.191 | TTTACGGCCACACTG |
| 1 | GMTACGGTCACACTG | chr3R:13873916-13875916 | 814 | 828 | - | 10.543 | 2.14E-06 | 0.191 | GGTACGGTCACGCTC |
| 1 | GMTACGGTCACACTG | chrX:13412242-13414242 | 827 | 841 | - | 10.543 | 2.14E-06 | 0.191 | GGCATGGTCACACTG |
| 1 | GMTACGGTCACACTG | chr2L:12973322-12975322 | 835 | 849 | - | 10.543 | 2.14E-06 | 0.191 | GGTGGGGTCACACTG |
| 1 | GMTACGGTCACACTG | chr3L:11122348-11124348 | 843 | 857 | - | 10.543 | 2.14E-06 | 0.191 | GTTCTGGTCACACTG |
| 1 | GMTACGGTCACACTG | chr2R:16271114-16273114 | 849 | 863 | - | 10.543 | 2.14E-06 | 0.191 | AGAACGGTCACACTG |
| 1 | GMTACGGTCACACTG | chr2L:21147546-21149546 | 861 | 875 | - | 10.543 | 2.14E-06 | 0.191 | CTCACGGTCACACTG |
| 1 | GMTACGGTCACACTG | chrX:18862950-18864950 | 863 | 877 | - | 10.543 | 2.14E-06 | 0.191 | ATGACGGTCACACTG |
| 1 | GMTACGGTCACACTG | chr2L:6126775-6128775 | 869 | 883 | - | 10.543 | 2.14E-06 | 0.191 | CGCACGGTCACACTG |
| 1 | GMTACGGTCACACTG | chr3L:10663739-10665739 | 882 | 896 | - | 10.543 | 2.14E-06 | 0.191 | GGTACGGTCATACCG |
| 1 | GMTACGGTCACACTG | chr3R:30427276-30429276 | 884 | 898 | - | 10.543 | 2.14E-06 | 0.191 | GGCATGGTCACACTG |
| 1 | GMTACGGTCACACTG | chrX:11147716-11149716 | 889 | 903 | - | 10.543 | 2.14E-06 | 0.191 | CGCACGGTCACACTG |
| 1 | GMTACGGTCACACTG | chrX:4105113-4107113 | 906 | 920 | - | 10.543 | 2.14E-06 | 0.191 | TGCACGGTCACACTG |
| 1 | GMTACGGTCACACTG | chr3L:12503292-12505292 | 933 | 947 | - | 10.543 | 2.14E-06 | 0.191 | CGAACGGTCACACTG |
| 1 | GMTACGGTCACACTG | chr3R:24185359-24187359 | 954 | 968 | - | 10.543 | 2.14E-06 | 0.191 | GTGATGGTCACACTG |
| 1 | GMTACGGTCACACTG | chr2R:21383391-21385391 | 960 | 974 | - | 10.543 | 2.14E-06 | 0.191 | AGTGCGGTCACACTG |
| 1 | GMTACGGTCACACTG | chr3R:11818359-11820359 | 963 | 977 | - | 10.543 | 2.14E-06 | 0.191 | ATTGCGGTCACACTG |
| 1 | GMTACGGTCACACTG | chr2L:19794851-19796851 | 964 | 978 | - | 10.543 | 2.14E-06 | 0.191 | CGAACGGTCACACTG |
| 1 | GMTACGGTCACACTG | chr3L:22074867-22076867 | 965 | 979 | - | 10.543 | 2.14E-06 | 0.191 | GGTAGGGTCACTCTG |
| 1 | GMTACGGTCACACTG | chr2R:17864963-17866963 | 969 | 983 | - | 10.543 | 2.14E-06 | 0.191 | TGGACGGTCACACTG |
| 1 | GMTACGGTCACACTG | chr2L:13190979-13192979 | 974 | 988 | - | 10.543 | 2.14E-06 | 0.191 | TTTACGGCCACACTG |
| 1 | GMTACGGTCACACTG | chr3L:1245923-1247923 | 974 | 988 | - | 10.543 | 2.14E-06 | 0.191 | GTTGCGGTCACACTA |
| 1 | GMTACGGTCACACTG | chr3L:11122216-11124216 | 975 | 989 | - | 10.543 | 2.14E-06 | 0.191 | GTTCTGGTCACACTG |
| 1 | GMTACGGTCACACTG | chrX:15856823-15858823 | 975 | 989 | - | 10.543 | 2.14E-06 | 0.191 | GTAACGGTCACACTA |
| 1 | GMTACGGTCACACTG | chr2L:8300079-8302079 | 977 | 991 | - | 10.543 | 2.14E-06 | 0.191 | AGTGCGGTCACACTG |
| 1 | GMTACGGTCACACTG | chr3L:11710386-11712386 | 982 | 996 | - | 10.543 | 2.14E-06 | 0.191 | TTTGCGGTCACACTG |
| 1 | GMTACGGTCACACTG | chr3L:4363102-4365102 | 982 | 996 | - | 10.543 | 2.14E-06 | 0.191 | ATTGCGGTCACACTG |
| 1 | GMTACGGTCACACTG | chr3L:18999608-19001608 | 982 | 996 | - | 10.543 | 2.14E-06 | 0.191 | TGGACGGTCACACTG |
| 1 | GMTACGGTCACACTG | chr2L:3158643-3160643 | 983 | 997 | - | 10.543 | 2.14E-06 | 0.191 | GTCATGGTCACACTG |
| 1 | GMTACGGTCACACTG | chr3R:11818339-11820339 | 983 | 997 | - | 10.543 | 2.14E-06 | 0.191 | ATTGCGGTCACACTG |
| 1 | GMTACGGTCACACTG | chr3R:9724615-9726615 | 983 | 997 | - | 10.543 | 2.14E-06 | 0.191 | AGTACGGTCACTCTG |

| | | | | | | | | | |
|---|---|---|---|---|---|---|---|---|---|
| 1 | GMTACGGTCACACTG | chr2R:13482229-13484229 | 983 | 997 | - | 10.543 | 2.14E-06 | 0.191 | AGTGCGGTCACACTG |
| 1 | GMTACGGTCACACTG | chr2R:14857640-14859640 | 985 | 999 | - | 10.543 | 2.14E-06 | 0.191 | GTCATGGTCACACTG |
| 1 | GMTACGGTCACACTG | chr2L:1613160-1615160 | 985 | 999 | - | 10.543 | 2.14E-06 | 0.191 | AGTGCGGTCACACTG |
| 1 | GMTACGGTCACACTG | chr2L:8081982-8083982 | 988 | 1002 | - | 10.543 | 2.14E-06 | 0.191 | CGAACGGTCACACTG |
| 1 | GMTACGGTCACACTG | chr2R:24874865-24876865 | 991 | 1005 | - | 10.543 | 2.14E-06 | 0.191 | GTTTCGGTCATACTG |
| 1 | GMTACGGTCACACTG | chr2R:16270971-16272971 | 992 | 1006 | - | 10.543 | 2.14E-06 | 0.191 | AGAACGGTCACACTG |
| 1 | GMTACGGTCACACTG | chrX:939805-941805 | 993 | 1007 | - | 10.543 | 2.14E-06 | 0.191 | AGGACGGTCACACTG |
| 1 | GMTACGGTCACACTG | chr2R:24786411-24788411 | 993 | 1007 | - | 10.543 | 2.14E-06 | 0.191 | ATTGCGGTCACACTG |
| 1 | GMTACGGTCACACTG | chr3R:24104763-24106763 | 993 | 1007 | - | 10.543 | 2.14E-06 | 0.191 | GGCACGGTCACACTC |
| 1 | GMTACGGTCACACTG | chr3L:1244910-1246910 | 994 | 1008 | - | 10.543 | 2.14E-06 | 0.191 | GTTGCGGTCACACTA |
| 1 | GMTACGGTCACACTG | chrX:13412074-13414074 | 995 | 1009 | - | 10.543 | 2.14E-06 | 0.191 | GGCATGGTCACACTG |
| 1 | GMTACGGTCACACTG | chrX:18656286-18658286 | 995 | 1009 | - | 10.543 | 2.14E-06 | 0.191 | GGTACGGTCACCCTA |
| 1 | GMTACGGTCACACTG | chr2L:6683364-6685364 | 995 | 1009 | - | 10.543 | 2.14E-06 | 0.191 | CGTGCGGTCACACTG |
| 1 | GMTACGGTCACACTG | chr2L:15750078-15752078 | 1006 | 1020 | - | 10.543 | 2.14E-06 | 0.191 | AGTGCGGTCACACTG |
| 1 | GMTACGGTCACACTG | chr3L:9398227-9400227 | 1013 | 1027 | - | 10.543 | 2.14E-06 | 0.191 | AGAACGGTCACACTG |
| 1 | GMTACGGTCACACTG | chr2L:19507764-19509764 | 1013 | 1027 | - | 10.543 | 2.14E-06 | 0.191 | GTCATGGTCACACTG |
| 1 | GMTACGGTCACACTG | chrX:10081492-10083492 | 1024 | 1038 | - | 10.543 | 2.14E-06 | 0.191 | AGCACGGTCACACTG |
| 1 | GMTACGGTCACACTG | chrX:18862776-18864776 | 1037 | 1051 | - | 10.543 | 2.14E-06 | 0.191 | ATGACGGTCACACTG |
| 1 | GMTACGGTCACACTG | chr3R:29858865-29860865 | 1043 | 1057 | - | 10.543 | 2.14E-06 | 0.191 | GTTGCGGTCACACTC |
| 1 | GMTACGGTCACACTG | chrX:12896327-12898327 | 1074 | 1088 | - | 10.543 | 2.14E-06 | 0.191 | TGCACGGTCACACTG |
| 1 | GMTACGGTCACACTG | chr3L:22074756-22076756 | 1076 | 1090 | - | 10.543 | 2.14E-06 | 0.191 | GGTAGGGTCACTCTG |
| 1 | GMTACGGTCACACTG | chr2L:8081859-8083859 | 1111 | 1125 | - | 10.543 | 2.14E-06 | 0.191 | CGAACGGTCACACTG |
| 1 | GMTACGGTCACACTG | chrX:2157983-2159983 | 1112 | 1126 | - | 10.543 | 2.14E-06 | 0.191 | GTCATGGTCACACTG |
| 1 | GMTACGGTCACACTG | chr3R:27503101-27505101 | 1254 | 1268 | - | 10.543 | 2.14E-06 | 0.191 | GGCATGGTCACACTG |
| 1 | GMTACGGTCACACTG | chrX:6471378-6473378 | 1300 | 1314 | - | 10.543 | 2.14E-06 | 0.191 | GGTTTGGTCACACTG |
| 1 | GMTACGGTCACACTG | chr3L:22748444-22750444 | 1314 | 1328 | - | 10.543 | 2.14E-06 | 0.191 | GGCACGTTCACACTG |
| 1 | GMTACGGTCACACTG | chr3L:12502894-12504894 | 1331 | 1345 | - | 10.543 | 2.14E-06 | 0.191 | CGAACGGTCACACTG |
| 1 | GMTACGGTCACACTG | chrX:9107691-9109691 | 1401 | 1415 | - | 10.543 | 2.14E-06 | 0.191 | TGGACGGTCACACTG |
| 1 | GMTACGGTCACACTG | chr2L:22018208-22020208 | 1441 | 1455 | - | 10.543 | 2.14E-06 | 0.191 | GTTGCGGTCAAACTG |
| 1 | GMTACGGTCACACTG | chr2L:22018208-22020208 | 1441 | 1455 | - | 10.543 | 2.14E-06 | 0.191 | GTTGCGGTCAAACTG |
| 1 | GMTACGGTCACACTG | chrX:22546097-22548097 | 1447 | 1461 | - | 10.543 | 2.14E-06 | 0.191 | GTTACGATCCCACTG |
| 1 | GMTACGGTCACACTG | chr2R:9588050-9590050 | 1453 | 1467 | - | 10.543 | 2.14E-06 | 0.191 | CGTTCGGTCACACTG |
| 1 | GMTACGGTCACACTG | chr3R:17800512-17802512 | 1456 | 1470 | - | 10.543 | 2.14E-06 | 0.191 | GTTACGGTCAAACAG |

| | | | | | | | | | |
|---|---|---|---|---|---|---|---|---|---|
| 1 | GMTACGGTCACACTG | chr2R:24873912-24875912 | 1944 | 1958 | - | 10.543 | 2.14E-06 | 0.191 | GTTTCGGTCATACTG |
| 1 | GMTACGGTCACACTG | chr3R:29857936-29859936 | 1972 | 1986 | - | 10.543 | 2.14E-06 | 0.191 | GTTGCGGTCACACTC |
| 1 | GMTACGGTCACACTG | chr2L:115488-117488 | 19 | 33 | + | 10.264 | 2.62E-06 | 0.219 | TTTACTGTCACACTG |
| 1 | GMTACGGTCACACTG | chr2L:8302429-8304429 | 118 | 132 | + | 10.264 | 2.62E-06 | 0.219 | GGTACGATGACACTG |
| 1 | GMTACGGTCACACTG | chr3L:16586805-16588805 | 298 | 312 | + | 10.264 | 2.62E-06 | 0.219 | AGTACGGTCACATTG |
| 1 | GMTACGGTCACACTG | chr3R:24112286-24114286 | 327 | 341 | + | 10.264 | 2.62E-06 | 0.219 | TGTACGGTCACACTT |
| 1 | GMTACGGTCACACTG | chr3R:11221405-11223405 | 364 | 378 | - | 10.264 | 2.62E-06 | 0.219 | AGTACGGTAACACTG |
| 1 | GMTACGGTCACACTG | chr2L:2286933-2288933 | 381 | 395 | + | 10.264 | 2.62E-06 | 0.219 | GGTAAAGTCACACTG |
| 1 | GMTACGGTCACACTG | chr3L:8600473-8602473 | 380 | 394 | - | 10.264 | 2.62E-06 | 0.219 | GTTACAGTCACACTA |
| 1 | GMTACGGTCACACTG | chrX:11603194-11605194 | 561 | 575 | + | 10.264 | 2.62E-06 | 0.219 | AGTACGGTCACAATG |
| 1 | GMTACGGTCACACTG | chr2L:8301985-8303985 | 562 | 576 | + | 10.264 | 2.62E-06 | 0.219 | GGTACGATGACACTG |
| 1 | GMTACGGTCACACTG | chr3R:24111964-24113964 | 649 | 663 | + | 10.264 | 2.62E-06 | 0.219 | TGTACGGTCACACTT |
| 1 | GMTACGGTCACACTG | chr2R:22126808-22128808 | 684 | 698 | - | 10.264 | 2.62E-06 | 0.219 | CGTACGGTCATACTG |
| 1 | GMTACGGTCACACTG | chr2R:18125347-18127347 | 726 | 740 | + | 10.264 | 2.62E-06 | 0.219 | CTTAAGGTCACACTG |
| 1 | GMTACGGTCACACTG | chr3L:21538862-21540862 | 741 | 755 | + | 10.264 | 2.62E-06 | 0.219 | GGTATGGTCACAATG |
| 1 | GMTACGGTCACACTG | chr2L:16309956-16311956 | 815 | 829 | + | 10.264 | 2.62E-06 | 0.219 | TGTACGGTCACATTG |
| 1 | GMTACGGTCACACTG | chr3L:8600006-8602006 | 847 | 861 | - | 10.264 | 2.62E-06 | 0.219 | GTTACAGTCACACTA |
| 1 | GMTACGGTCACACTG | chr3R:11220894-11222894 | 875 | 889 | - | 10.264 | 2.62E-06 | 0.219 | AGTACGGTAACACTG |
| 1 | GMTACGGTCACACTG | chrX:16650376-16652376 | 911 | 925 | + | 10.264 | 2.62E-06 | 0.219 | GTTACGGTCATACTA |
| 1 | GMTACGGTCACACTG | chrX:16650376-16652376 | 911 | 925 | + | 10.264 | 2.62E-06 | 0.219 | GTTACGGTCATACTA |
| 1 | GMTACGGTCACACTG | chr3L:3168375-3170375 | 913 | 927 | + | 10.264 | 2.62E-06 | 0.219 | TGTATGGTCACACTG |
| 1 | GMTACGGTCACACTG | chrX:15243913-15245913 | 914 | 928 | + | 10.264 | 2.62E-06 | 0.219 | GGTACGTTCATACTG |
| 1 | GMTACGGTCACACTG | chrX:11153421-11155421 | 907 | 921 | - | 10.264 | 2.62E-06 | 0.219 | GGTATGGTCACACTT |
| 1 | GMTACGGTCACACTG | chr3L:16586157-16588157 | 946 | 960 | + | 10.264 | 2.62E-06 | 0.219 | AGTACGGTCACATTG |
| 1 | GMTACGGTCACACTG | chr3R:20559865-20561865 | 936 | 950 | - | 10.264 | 2.62E-06 | 0.219 | GTTACGGTCATACTT |
| 1 | GMTACGGTCACACTG | chr3L:8599879-8601879 | 974 | 988 | - | 10.264 | 2.62E-06 | 0.219 | GTTACAGTCACACTA |
| 1 | GMTACGGTCACACTG | chr3L:4253776-4255776 | 991 | 1005 | + | 10.264 | 2.62E-06 | 0.219 | GTTAAGGTCACACTC |
| 1 | GMTACGGTCACACTG | chr3R:25505275-25507275 | 994 | 1008 | + | 10.264 | 2.62E-06 | 0.219 | GTTAAGGTAACACTG |
| 1 | GMTACGGTCACACTG | chr3R:21776847-21778847 | 983 | 997 | - | 10.264 | 2.62E-06 | 0.219 | GTTACGCTCATACTG |
| 1 | GMTACGGTCACACTG | chr3L:3168286-3170286 | 1002 | 1016 | + | 10.264 | 2.62E-06 | 0.219 | TGTATGGTCACACTG |
| 1 | GMTACGGTCACACTG | chr3L:8314502-8316502 | 989 | 1003 | - | 10.264 | 2.62E-06 | 0.219 | TTTACAGTCACACTG |
| 1 | GMTACGGTCACACTG | chr3L:10878085-10880085 | 1003 | 1017 | + | 10.264 | 2.62E-06 | 0.219 | ATTATGGTCACACTG |
| 1 | GMTACGGTCACACTG | chr3R:13263364-13265364 | 1004 | 1018 | + | 10.264 | 2.62E-06 | 0.219 | AGTACGGTCACATTG |

| | | | | | | | | | |
|---|---|---|---|---|---|---|---|---|---|
| 1 | GMTACGGTCACACTG | chr2R:22126502-22128502 | 990 | 1004 | - | 10.264 | 2.62E-06 | 0.219 | CGTACGGTCATACTG |
| 1 | GMTACGGTCACACTG | chr2L:9329763-9331763 | 1007 | 1021 | + | 10.264 | 2.62E-06 | 0.219 | AGTACGGTCACACTT |
| 1 | GMTACGGTCACACTG | chr3R:5482906-5484906 | 994 | 1008 | - | 10.264 | 2.62E-06 | 0.219 | GGTATGGTCACACTT |
| 1 | GMTACGGTCACACTG | chr2R:19291529-19293529 | 1010 | 1024 | + | 10.264 | 2.62E-06 | 0.219 | TTTATGGTCACACTG |
| 1 | GMTACGGTCACACTG | chr2L:16309726-16311726 | 1045 | 1059 | + | 10.264 | 2.62E-06 | 0.219 | TGTACGGTCACATTG |
| 1 | GMTACGGTCACACTG | chr3L:16586057-16588057 | 1046 | 1060 | + | 10.264 | 2.62E-06 | 0.219 | AGTACGGTCACATTG |
| 1 | GMTACGGTCACACTG | chr3L:10878037-10880037 | 1051 | 1065 | + | 10.264 | 2.62E-06 | 0.219 | ATTATGGTCACACTG |
| 1 | GMTACGGTCACACTG | chr3R:11220639-11222639 | 1130 | 1144 | - | 10.264 | 2.62E-06 | 0.219 | AGTACGGTAACACTG |
| 1 | GMTACGGTCACACTG | chr3L:21538447-21540447 | 1156 | 1170 | + | 10.264 | 2.62E-06 | 0.219 | GGTATGGTCACAATG |
| 1 | GMTACGGTCACACTG | chr3R:24111389-24113389 | 1224 | 1238 | + | 10.264 | 2.62E-06 | 0.219 | TGTACGGTCACACTT |
| 1 | GMTACGGTCACACTG | chr2R:18124779-18126779 | 1294 | 1308 | + | 10.264 | 2.62E-06 | 0.219 | CTTAAGGTCACACTG |
| 1 | GMTACGGTCACACTG | chr3R:13263061-13265061 | 1307 | 1321 | + | 10.264 | 2.62E-06 | 0.219 | AGTACGGTCACATTG |
| 1 | GMTACGGTCACACTG | chr2R:19291196-19293196 | 1343 | 1357 | + | 10.264 | 2.62E-06 | 0.219 | TTTATGGTCACACTG |
| 1 | GMTACGGTCACACTG | chrX:11152990-11154990 | 1338 | 1352 | - | 10.264 | 2.62E-06 | 0.219 | GGTATGGTCACACTT |
| 1 | GMTACGGTCACACTG | chr2L:2285873-2287873 | 1441 | 1455 | + | 10.264 | 2.62E-06 | 0.219 | GGTAAAGTCACACTG |
| 1 | GMTACGGTCACACTG | chr3L:4253306-4255306 | 1461 | 1475 | + | 10.264 | 2.62E-06 | 0.219 | GTTAAGGTCACACTC |
| 1 | GMTACGGTCACACTG | chr3R:20559280-20561280 | 1521 | 1535 | - | 10.264 | 2.62E-06 | 0.219 | GTTACGGTCATACTT |
| 1 | GMTACGGTCACACTG | chr2R:16016737-16018737 | 1592 | 1606 | + | 10.264 | 2.62E-06 | 0.219 | GTTACGATCACAATG |
| 1 | GMTACGGTCACACTG | chr2L:113726-115726 | 1781 | 1795 | + | 10.264 | 2.62E-06 | 0.219 | TTTACTGTCACACTG |
| 1 | GMTACGGTCACACTG | chr2L:8082420-8084420 | 1825 | 1839 | + | 10.264 | 2.62E-06 | 0.219 | GTTACTGTCACAATG |
| 1 | GMTACGGTCACACTG | chr2L:10326943-10328943 | 57 | 71 | + | 10.121 | 2.85E-06 | 0.23 | GCTGCGGTCGCTCTG |
| 1 | GMTACGGTCACACTG | chr3L:5779863-5781863 | 192 | 206 | + | 10.121 | 2.85E-06 | 0.23 | GCTGCGGCCGCACTG |
| 1 | GMTACGGTCACACTG | chr3R:26092722-26094722 | 245 | 259 | - | 10.121 | 2.85E-06 | 0.23 | GCTGCGGTCGCTCTG |
| 1 | GMTACGGTCACACTG | chrX:14852269-14854269 | 279 | 293 | + | 10.121 | 2.85E-06 | 0.23 | GCATCGGGCACACTG |
| 1 | GMTACGGTCACACTG | chr3R:32014520-32016520 | 383 | 397 | + | 10.121 | 2.85E-06 | 0.23 | GCAGCGGGCACACTG |
| 1 | GMTACGGTCACACTG | chr3R:24658621-24660621 | 404 | 418 | - | 10.121 | 2.85E-06 | 0.23 | GCTGCGGTCCCGCTG |
| 1 | GMTACGGTCACACTG | chr3R:27321325-27323325 | 532 | 546 | + | 10.121 | 2.85E-06 | 0.23 | GCCACGGGCACTCTG |
| 1 | GMTACGGTCACACTG | chr3R:12997089-12999089 | 967 | 981 | + | 10.121 | 2.85E-06 | 0.23 | GCCACGGGCACACCG |
| 1 | GMTACGGTCACACTG | chr2L:6919704-6921704 | 977 | 991 | + | 10.121 | 2.85E-06 | 0.23 | GCGGCGGTCACACAG |
| 1 | GMTACGGTCACACTG | chr3R:19878719-19880719 | 984 | 998 | + | 10.121 | 2.85E-06 | 0.23 | GCAGCGGTCACACAG |
| 1 | GMTACGGTCACACTG | chr2R:8599415-8601415 | 990 | 1004 | + | 10.121 | 2.85E-06 | 0.23 | GCCTCGGCCACACTG |
| 1 | GMTACGGTCACACTG | chr3L:5779051-5781051 | 1004 | 1018 | + | 10.121 | 2.85E-06 | 0.23 | GCTGCGGCCGCACTG |
| 1 | GMTACGGTCACACTG | chr3L:5779051-5781051 | 1004 | 1018 | + | 10.121 | 2.85E-06 | 0.23 | GCTGCGGCCGCACTG |

| | | | | | | | | | |
|---|---|---|---|---|---|---|---|---|---|
| 1 | GMTACGGTCACACTG | chrX:2072483-2074483 | 1052 | 1066 | + | 10.121 | 2.85E-06 | 0.23 | GCGTCGGGCACACTG |
| 1 | GMTACGGTCACACTG | chr2L:15911558-15913558 | 1082 | 1096 | - | 10.121 | 2.85E-06 | 0.23 | GCTGCGGCCACTCTG |
| 1 | GMTACGGTCACACTG | chr3R:15972298-15974298 | 1144 | 1158 | - | 10.121 | 2.85E-06 | 0.23 | GCGACGGACACACGG |
| 1 | GMTACGGTCACACTG | chr2R:24401680-24403680 | 1317 | 1331 | + | 10.121 | 2.85E-06 | 0.23 | GCGTCGGCCACACTG |
| 1 | GMTACGGTCACACTG | chr3R:12996726-12998726 | 1330 | 1344 | + | 10.121 | 2.85E-06 | 0.23 | GCCACGGGCACACCG |
| 1 | GMTACGGTCACACTG | chr3R:12996726-12998726 | 1330 | 1344 | + | 10.121 | 2.85E-06 | 0.23 | GCCACGGGCACACCG |
| 1 | GMTACGGTCACACTG | chrX:1921981-1923981 | 1380 | 1394 | + | 10.121 | 2.85E-06 | 0.23 | GCCACGGTCGCGCTG |
| 1 | GMTACGGTCACACTG | chr3R:24657605-24659605 | 1420 | 1434 | - | 10.121 | 2.85E-06 | 0.23 | GCTGCGGTCCCGCTG |
| 1 | GMTACGGTCACACTG | chr2R:8598962-8600962 | 1443 | 1457 | + | 10.121 | 2.85E-06 | 0.23 | GCCTCGGCCACACTG |
| 1 | GMTACGGTCACACTG | chr3L:5778591-5780591 | 1464 | 1478 | + | 10.121 | 2.85E-06 | 0.23 | GCTGCGGCCGCACTG |
| 1 | GMTACGGTCACACTG | chr2L:6919210-6921210 | 1471 | 1485 | + | 10.121 | 2.85E-06 | 0.23 | GCGGCGGTCACACAG |
| 1 | GMTACGGTCACACTG | chrX:1921874-1923874 | 1487 | 1501 | + | 10.121 | 2.85E-06 | 0.23 | GCCACGGTCGCGCTG |
| 1 | GMTACGGTCACACTG | chr3L:22711155-22713155 | 1483 | 1497 | - | 10.121 | 2.85E-06 | 0.23 | GCTACGGTCTCTCCG |
| 1 | GMTACGGTCACACTG | chr3R:15971913-15973913 | 1529 | 1543 | - | 10.121 | 2.85E-06 | 0.23 | GCGACGGACACACGG |
| 1 | GMTACGGTCACACTG | chr2R:9344349-9346349 | 1655 | 1669 | + | 10.121 | 2.85E-06 | 0.23 | GCAACGGACACCCTG |
| 1 | GMTACGGTCACACTG | chr3L:18474298-18476298 | 1908 | 1922 | + | 10.121 | 2.85E-06 | 0.23 | GCTGCGGTCCCGCTG |
| 1 | GMTACGGTCACACTG | chr2R:20985500-20987500 | 1909 | 1923 | + | 10.121 | 2.85E-06 | 0.23 | GCTACGGGCGCACAG |
| 1 | GMTACGGTCACACTG | chr2L:15910691-15912691 | 1949 | 1963 | - | 10.121 | 2.85E-06 | 0.23 | GCTGCGGCCACTCTG |
| 1 | GMTACGGTCACACTG | chr3L:1651750-1653750 | 32 | 46 | + | 9.8429 | 5.00E-06 | 0.286 | GCCACGGGCACAATG |
| 1 | GMTACGGTCACACTG | chrX:19624705-19626705 | 147 | 161 | + | 9.8429 | 5.00E-06 | 0.286 | GCTTCGGTTACGCTG |
| 1 | GMTACGGTCACACTG | chr3R:10855807-10857807 | 152 | 166 | + | 9.8429 | 5.00E-06 | 0.286 | GCTACGGTCCCATCG |
| 1 | GMTACGGTCACACTG | chrX:1341106-1343106 | 161 | 175 | + | 9.8429 | 5.00E-06 | 0.286 | GCGAAGGTCGCACTG |
| 1 | GMTACGGTCACACTG | chr3R:8291335-8293335 | 169 | 183 | + | 9.8429 | 5.00E-06 | 0.286 | GCCAAGGACACACTG |
| 1 | GMTACGGTCACACTG | chr3R:15218256-15220256 | 265 | 279 | + | 9.8429 | 5.00E-06 | 0.286 | GCTCCAGTCACCCTG |
| 1 | GMTACGGTCACACTG | chr3R:13630942-13632942 | 282 | 296 | + | 9.8429 | 5.00E-06 | 0.286 | GCTGCTGCCACACTG |
| 1 | GMTACGGTCACACTG | chr3L:20200638-20202638 | 287 | 301 | + | 9.8429 | 5.00E-06 | 0.286 | GCCACGCCCACACTG |
| 1 | GMTACGGTCACACTG | chrX:19211082-19213082 | 300 | 314 | + | 9.8429 | 5.00E-06 | 0.286 | GCTACGGGGACTCTG |
| 1 | GMTACGGTCACACTG | chr3L:3250022-3252022 | 345 | 359 | + | 9.8429 | 5.00E-06 | 0.286 | GCAACGTTCTCACTG |
| 1 | GMTACGGTCACACTG | chr3L:22740793-22742793 | 358 | 372 | + | 9.8429 | 5.00E-06 | 0.286 | GCTTCGGTCGCACTT |
| 1 | GMTACGGTCACACTG | chr2L:5099877-5101877 | 390 | 404 | + | 9.8429 | 5.00E-06 | 0.286 | TCAGCGGTCACACTG |
| 1 | GMTACGGTCACACTG | chr2R:23041943-23043943 | 513 | 527 | + | 9.8429 | 5.00E-06 | 0.286 | GCTACGGACACAACG |
| 1 | GMTACGGTCACACTG | chrX:6967750-6969750 | 530 | 544 | + | 9.8429 | 5.00E-06 | 0.286 | TCTGCGGACACACTG |
| 1 | GMTACGGTCACACTG | chrX:5878869-5880869 | 546 | 560 | + | 9.8429 | 5.00E-06 | 0.286 | GCTACCGTCACTCCG |

| | | | | | | | | | |
|---|---|---|---|---|---|---|---|---|---|
| 1 | GMTACGGTCACACTG | chrX:9231378-9233378 | 549 | 563 | + | 9.8429 | 5.00E-06 | 0.286 | GCGACGGTCATTCTG |
| 1 | GMTACGGTCACACTG | chr3R:14658438-14660438 | 573 | 587 | + | 9.8429 | 5.00E-06 | 0.286 | GCTACGGCAACGCTG |
| 1 | GMTACGGTCACACTG | chr2R:23511225-23513225 | 575 | 589 | + | 9.8429 | 5.00E-06 | 0.286 | GCCACGGTCTCAATG |
| 1 | GMTACGGTCACACTG | chr3L:22836070-22838070 | 598 | 612 | + | 9.8429 | 5.00E-06 | 0.286 | GCGACAGTCACTCTG |
| 1 | GMTACGGTCACACTG | chr3R:24876045-24878045 | 599 | 613 | + | 9.8429 | 5.00E-06 | 0.286 | GCTTCGGCCACACTA |
| 1 | GMTACGGTCACACTG | chr3R:24876045-24878045 | 599 | 613 | + | 9.8429 | 5.00E-06 | 0.286 | GCTTCGGCCACACTA |
| 1 | GMTACGGTCACACTG | chr3L:22836068-22838068 | 600 | 614 | + | 9.8429 | 5.00E-06 | 0.286 | GCGACAGTCACTCTG |
| 1 | GMTACGGTCACACTG | chr3R:10055083-10057083 | 620 | 634 | + | 9.8429 | 5.00E-06 | 0.286 | GATTCGGGCACACGG |
| 1 | GMTACGGTCACACTG | chrX:5878787-5880787 | 628 | 642 | + | 9.8429 | 5.00E-06 | 0.286 | GCTACCGTCACTCCG |
| 1 | GMTACGGTCACACTG | chrX:10328872-10330872 | 646 | 660 | + | 9.8429 | 5.00E-06 | 0.286 | GAGCCGGGCACACTG |
| 1 | GMTACGGTCACACTG | chr2R:6039647-6041647 | 647 | 661 | + | 9.8429 | 5.00E-06 | 0.286 | GCGACGTTCTCACTG |
| 1 | GMTACGGTCACACTG | chr3R:15217874-15219874 | 647 | 661 | + | 9.8429 | 5.00E-06 | 0.286 | GCTCCAGTCACCCTG |
| 1 | GMTACGGTCACACTG | chrX:1340587-1342587 | 680 | 694 | + | 9.8429 | 5.00E-06 | 0.286 | GCGAAGGTCGCACTG |
| 1 | GMTACGGTCACACTG | chr2R:23539773-23541773 | 756 | 770 | + | 9.8429 | 5.00E-06 | 0.286 | GCGGCGATCACACTG |
| 1 | GMTACGGTCACACTG | chr3L:22740388-22742388 | 763 | 777 | + | 9.8429 | 5.00E-06 | 0.286 | GCTTCGGTCGCACTT |
| 1 | GMTACGGTCACACTG | chr2R:11344119-11346119 | 765 | 779 | + | 9.8429 | 5.00E-06 | 0.286 | GCGGCGGTCACAATG |
| 1 | GMTACGGTCACACTG | chr2L:318259-320259 | 766 | 780 | + | 9.8429 | 5.00E-06 | 0.286 | GCGGCGGTCACAATG |
| 1 | GMTACGGTCACACTG | chr3R:26946830-26948830 | 789 | 803 | + | 9.8429 | 5.00E-06 | 0.286 | GCTGCGGGCACACTT |
| 1 | GMTACGGTCACACTG | chr3R:11136177-11138177 | 823 | 837 | + | 9.8429 | 5.00E-06 | 0.286 | GCTACGGTCGAACAG |
| 1 | GMTACGGTCACACTG | chr3L:4625903-4627903 | 827 | 841 | + | 9.8429 | 5.00E-06 | 0.286 | TCCGCGGTCACACTG |
| 1 | GMTACGGTCACACTG | chr3R:13630379-13632379 | 845 | 859 | + | 9.8429 | 5.00E-06 | 0.286 | GCTGCTGCCACACTG |
| 1 | GMTACGGTCACACTG | chr3L:11832578-11834578 | 858 | 872 | + | 9.8429 | 5.00E-06 | 0.286 | GCTACGCTCGCACGG |
| 1 | GMTACGGTCACACTG | chr2L:5070109-5072109 | 876 | 890 | + | 9.8429 | 5.00E-06 | 0.286 | GCGGCGGTCACACTT |
| 1 | GMTACGGTCACACTG | chrX:12630605-12632605 | 926 | 940 | + | 9.8429 | 5.00E-06 | 0.286 | GCTTTGGTCTCACTG |
| 1 | GMTACGGTCACACTG | chrX:17702352-17704352 | 957 | 971 | + | 9.8429 | 5.00E-06 | 0.286 | CCCACGGTCACTCTG |
| 1 | GMTACGGTCACACTG | chr3L:5793192-5795192 | 959 | 973 | + | 9.8429 | 5.00E-06 | 0.286 | GCTGCGGTCACACCT |
| 1 | GMTACGGTCACACTG | chr2L:6488090-6490090 | 960 | 974 | + | 9.8429 | 5.00E-06 | 0.286 | GCAATGGTCACTCTG |
| 1 | GMTACGGTCACACTG | chr3R:18905609-18907609 | 962 | 976 | + | 9.8429 | 5.00E-06 | 0.286 | CCACCGGTCACACTG |
| 1 | GMTACGGTCACACTG | chr2R:12150057-12152057 | 964 | 978 | + | 9.8429 | 5.00E-06 | 0.286 | GCTACGGACACACAC |
| 1 | GMTACGGTCACACTG | chr2R:8599415-8601415 | 970 | 984 | + | 9.8429 | 5.00E-06 | 0.286 | GCGATGGCCACACTG |
| 1 | GMTACGGTCACACTG | chr3L:10881890-10883890 | 976 | 990 | + | 9.8429 | 5.00E-06 | 0.286 | ACTGCGGACACACTG |
| 1 | GMTACGGTCACACTG | chr2L:5099288-5101288 | 979 | 993 | + | 9.8429 | 5.00E-06 | 0.286 | TCAGCGGTCACACTG |
| 1 | GMTACGGTCACACTG | chrX:17106582-17108582 | 979 | 993 | + | 9.8429 | 5.00E-06 | 0.286 | GCGCTGGTCACACTG |

| | | | | | | | | | |
|---|---|---|---|---|---|---|---|---|---|
| 1 | GMTACGGTCACACTG | chr3L:9697336-9699336 | 980 | 994 | + | 9.8429 | 5.00E-06 | 0.286 | GCGGCGGTCACACTA |
| 1 | GMTACGGTCACACTG | chrX:17148001-17150001 | 981 | 995 | + | 9.8429 | 5.00E-06 | 0.286 | TCTCCGGCCACACTG |
| 1 | GMTACGGTCACACTG | chr3R:19904164-19906164 | 982 | 996 | + | 9.8429 | 5.00E-06 | 0.286 | GCTGCGGTCACACCC |
| 1 | GMTACGGTCACACTG | chr2R:7805484-7807484 | 982 | 996 | + | 9.8429 | 5.00E-06 | 0.286 | GCCCTGGTCACACTG |
| 1 | GMTACGGTCACACTG | chrX:6778659-6780659 | 983 | 997 | + | 9.8429 | 5.00E-06 | 0.286 | GCGACCGTCGCACTG |
| 1 | GMTACGGTCACACTG | chr3R:9590309-9592309 | 985 | 999 | + | 9.8429 | 5.00E-06 | 0.286 | CCGACGGTCACACAG |
| 1 | GMTACGGTCACACTG | chr3R:4386742-4388742 | 986 | 1000 | + | 9.8429 | 5.00E-06 | 0.286 | TCCTCGGTCACACTG |
| 1 | GMTACGGTCACACTG | chrX:8408272-8410272 | 986 | 1000 | + | 9.8429 | 5.00E-06 | 0.286 | ACCACGGTCACGCTG |
| 1 | GMTACGGTCACACTG | chr3R:24875654-24877654 | 990 | 1004 | + | 9.8429 | 5.00E-06 | 0.286 | GCTTCGGCCACACTA |
| 1 | GMTACGGTCACACTG | chr2L:7062986-7064986 | 993 | 1007 | + | 9.8429 | 5.00E-06 | 0.286 | GCTGCGGTCACACAA |
| 1 | GMTACGGTCACACTG | chr2R:16867222-16869222 | 993 | 1007 | + | 9.8429 | 5.00E-06 | 0.286 | GCGAAGGTCACTCTG |
| 1 | GMTACGGTCACACTG | chr3R:11627759-11629759 | 994 | 1008 | + | 9.8429 | 5.00E-06 | 0.286 | CCAACGGTCACACAG |
| 1 | GMTACGGTCACACTG | chr2L:13202762-13204762 | 994 | 1008 | + | 9.8429 | 5.00E-06 | 0.286 | GATGCGGCCACTCTG |
| 1 | GMTACGGTCACACTG | chr3L:4625734-4627734 | 996 | 1010 | + | 9.8429 | 5.00E-06 | 0.286 | TCCGCGGTCACACTG |
| 1 | GMTACGGTCACACTG | chr2L:6497647-6499647 | 997 | 1011 | + | 9.8429 | 5.00E-06 | 0.286 | GCAACGGTCACTTTG |
| 1 | GMTACGGTCACACTG | chr2R:9102238-9104238 | 1000 | 1014 | + | 9.8429 | 5.00E-06 | 0.286 | GCGGTGGTCACACTG |
| 1 | GMTACGGTCACACTG | chr2R:21010970-21012970 | 1001 | 1015 | + | 9.8429 | 5.00E-06 | 0.286 | ACGTCGGTCACACTG |
| 1 | GMTACGGTCACACTG | chrX:19622114-19624114 | 1001 | 1015 | + | 9.8429 | 5.00E-06 | 0.286 | GCCGCGGTCACACTA |
| 1 | GMTACGGTCACACTG | chr2R:18439965-18441965 | 1005 | 1019 | + | 9.8429 | 5.00E-06 | 0.286 | TCCACGGCCACACTG |
| 1 | GMTACGGTCACACTG | chrX:15709647-15711647 | 1006 | 1020 | + | 9.8429 | 5.00E-06 | 0.286 | GCGAGGGGCACACTG |
| 1 | GMTACGGTCACACTG | chr2L:6556800-6558800 | 1007 | 1021 | + | 9.8429 | 5.00E-06 | 0.286 | ACCGCGGTCACACTG |
| 1 | GMTACGGTCACACTG | chr3L:9445864-9447864 | 1008 | 1022 | + | 9.8429 | 5.00E-06 | 0.286 | GCCATGGTCACTCTG |
| 1 | GMTACGGTCACACTG | chr2L:19185043-19187043 | 1009 | 1023 | + | 9.8429 | 5.00E-06 | 0.286 | GCAACGTGCACACTG |
| 1 | GMTACGGTCACACTG | chr2R:21684283-21686283 | 1010 | 1024 | + | 9.8429 | 5.00E-06 | 0.286 | ACTGCGGTCACACCG |
| 1 | GMTACGGTCACACTG | chr3R:11966708-11968708 | 1011 | 1025 | + | 9.8429 | 5.00E-06 | 0.286 | GAAACGGTCACTCCG |
| 1 | GMTACGGTCACACTG | chr3R:24649125-24651125 | 1012 | 1026 | + | 9.8429 | 5.00E-06 | 0.286 | CCAACGGTCACTCTG |
| 1 | GMTACGGTCACACTG | chr3L:8190270-8192270 | 1012 | 1026 | + | 9.8429 | 5.00E-06 | 0.286 | ACCTCGGTCACACTG |
| 1 | GMTACGGTCACACTG | chr3L:11707979-11709979 | 1014 | 1028 | + | 9.8429 | 5.00E-06 | 0.286 | GCTGCGGCCACACTA |
| 1 | GMTACGGTCACACTG | chr3R:9684760-9686760 | 1014 | 1028 | + | 9.8429 | 5.00E-06 | 0.286 | GCTTTGGACACACTG |
| 1 | GMTACGGTCACACTG | chr3L:11707979-11709979 | 1014 | 1028 | + | 9.8429 | 5.00E-06 | 0.286 | GCTGCGGCCACACTA |
| 1 | GMTACGGTCACACTG | chr3L:20435250-20437250 | 1014 | 1028 | + | 9.8429 | 5.00E-06 | 0.286 | GCTGCGACCACACTG |
| 1 | GMTACGGTCACACTG | chr3R:6345111-6347111 | 1014 | 1028 | + | 9.8429 | 5.00E-06 | 0.286 | GCCTTGGTCACACTG |
| 1 | GMTACGGTCACACTG | chr3R:9692079-9694079 | 1015 | 1029 | + | 9.8429 | 5.00E-06 | 0.286 | GCTTTGGACACACTG |

| | | | | | | | | | |
|---|---|---|---|---|---|---|---|---|---|
| 1 | GMTACGGTCACACTG | chrX:2623804-2625804 | 1019 | 1033 | + | 9.8429 | 5.00E-06 | 0.286 | GCGCTGGTCACACTG |
| 1 | GMTACGGTCACACTG | chr3R:17681669-17683669 | 1025 | 1039 | + | 9.8429 | 5.00E-06 | 0.286 | GCCATGGTCGCACTG |
| 1 | GMTACGGTCACACTG | chr2L:13826772-13828772 | 1026 | 1040 | + | 9.8429 | 5.00E-06 | 0.286 | GCGGCGGTCACATTG |
| 1 | GMTACGGTCACACTG | chr3R:11966691-11968691 | 1028 | 1042 | + | 9.8429 | 5.00E-06 | 0.286 | GAAACGGTCACTCCG |
| 1 | GMTACGGTCACACTG | chr2L:5069957-5071957 | 1028 | 1042 | + | 9.8429 | 5.00E-06 | 0.286 | GCGGCGGTCACACTT |
| 1 | GMTACGGTCACACTG | chr3L:17464945-17466945 | 1032 | 1046 | + | 9.8429 | 5.00E-06 | 0.286 | GACACGGTCACTCCG |
| 1 | GMTACGGTCACACTG | chr3R:26946583-26948583 | 1036 | 1050 | + | 9.8429 | 5.00E-06 | 0.286 | GCTGCGGGCACACTT |
| 1 | GMTACGGTCACACTG | chrX:19622074-19624074 | 1041 | 1055 | + | 9.8429 | 5.00E-06 | 0.286 | GCCGCGGTCACACTA |
| 1 | GMTACGGTCACACTG | chr2L:19053078-19055078 | 1046 | 1060 | + | 9.8429 | 5.00E-06 | 0.286 | GCCACGGTCACACAA |
| 1 | GMTACGGTCACACTG | chr2R:23401309-23403309 | 1048 | 1062 | + | 9.8429 | 5.00E-06 | 0.286 | GCAACGGTTTCACTG |
| 1 | GMTACGGTCACACTG | chr2R:21010921-21012921 | 1050 | 1064 | + | 9.8429 | 5.00E-06 | 0.286 | ACGTCGGTCACACTG |
| 1 | GMTACGGTCACACTG | chr3L:20435209-20437209 | 1055 | 1069 | + | 9.8429 | 5.00E-06 | 0.286 | GCTGCGACCACACTG |
| 1 | GMTACGGTCACACTG | chrX:5763553-5765553 | 1072 | 1086 | + | 9.8429 | 5.00E-06 | 0.286 | GCCACGGCCAAACTG |
| 1 | GMTACGGTCACACTG | chrX:19703259-19705259 | 1076 | 1090 | + | 9.8429 | 5.00E-06 | 0.286 | CCCGCGGTCACACTG |
| 1 | GMTACGGTCACACTG | chr3R:14028228-14030228 | 1080 | 1094 | + | 9.8429 | 5.00E-06 | 0.286 | GCTGCAGTCACGCTG |
| 1 | GMTACGGTCACACTG | chr2R:7805385-7807385 | 1081 | 1095 | + | 9.8429 | 5.00E-06 | 0.286 | GCCCTGGTCACACTG |
| 1 | GMTACGGTCACACTG | chr3R:24649043-24651043 | 1094 | 1108 | + | 9.8429 | 5.00E-06 | 0.286 | CCAACGGTCACTCTG |
| 1 | GMTACGGTCACACTG | chrX:7290298-7292298 | 1095 | 1109 | + | 9.8429 | 5.00E-06 | 0.286 | GCAGGGGTCACACTG |
| 1 | GMTACGGTCACACTG | chrX:15709539-15711539 | 1114 | 1128 | + | 9.8429 | 5.00E-06 | 0.286 | GCGAGGGGCACACTG |
| 1 | GMTACGGTCACACTG | chr2R:20617503-20619503 | 1116 | 1130 | + | 9.8429 | 5.00E-06 | 0.286 | CCTGCGGTCTCACTG |
| 1 | GMTACGGTCACACTG | chr3R:27093533-27095533 | 1133 | 1147 | + | 9.8429 | 5.00E-06 | 0.286 | GCTACGGCCGCACTT |
| 1 | GMTACGGTCACACTG | chrX:19703176-19705176 | 1159 | 1173 | + | 9.8429 | 5.00E-06 | 0.286 | CCCGCGGTCACACTG |
| 1 | GMTACGGTCACACTG | chr3R:27871390-27873390 | 1165 | 1179 | + | 9.8429 | 5.00E-06 | 0.286 | GCTTTGGGCACACTG |
| 1 | GMTACGGTCACACTG | chr3L:16583124-16585124 | 1170 | 1184 | + | 9.8429 | 5.00E-06 | 0.286 | GCGACGGCCACACTA |
| 1 | GMTACGGTCACACTG | chr3R:17681515-17683515 | 1179 | 1193 | + | 9.8429 | 5.00E-06 | 0.286 | GCCATGGTCGCACTG |
| 1 | GMTACGGTCACACTG | chr2R:8357538-8359538 | 1209 | 1223 | + | 9.8429 | 5.00E-06 | 0.286 | GATCCGGACGCACTG |
| 1 | GMTACGGTCACACTG | chr3R:6344911-6346911 | 1214 | 1228 | + | 9.8429 | 5.00E-06 | 0.286 | GCCTTGGTCACACTG |
| 1 | GMTACGGTCACACTG | chrX:2623575-2625575 | 1248 | 1262 | + | 9.8429 | 5.00E-06 | 0.286 | GCGCTGGTCACACTG |
| 1 | GMTACGGTCACACTG | chr3R:8290240-8292240 | 1264 | 1278 | + | 9.8429 | 5.00E-06 | 0.286 | GCCAAGGACACACTG |
| 1 | GMTACGGTCACACTG | chr2L:19033576-19035576 | 1274 | 1288 | + | 9.8429 | 5.00E-06 | 0.286 | GCAACGGTGTCACTG |
| 1 | GMTACGGTCACACTG | chr3R:19903868-19905868 | 1278 | 1292 | + | 9.8429 | 5.00E-06 | 0.286 | GCTGCGGTCACACCC |
| 1 | GMTACGGTCACACTG | chr3R:26946334-26948334 | 1285 | 1299 | + | 9.8429 | 5.00E-06 | 0.286 | GCTGCGGGCACACTT |
| 1 | GMTACGGTCACACTG | chrX:17702024-17704024 | 1285 | 1299 | + | 9.8429 | 5.00E-06 | 0.286 | CCCACGGTCACTCTG |

| | | | | | | | | | |
|---|---|---|---|---|---|---|---|---|---|
| 1 | GMTACGGTCACACTG | chr2L:18866161-18868161 | 1287 | 1301 | + | 9.8429 | 5.00E-06 | 0.286 | GACTCGGACACACTG |
| 1 | GMTACGGTCACACTG | chr3R:9590002-9592002 | 1292 | 1306 | + | 9.8429 | 5.00E-06 | 0.286 | CCGACGGTCACACAG |
| 1 | GMTACGGTCACACTG | chr2L:6487756-6489756 | 1294 | 1308 | + | 9.8429 | 5.00E-06 | 0.286 | GCAATGGTCACTCTG |
| 1 | GMTACGGTCACACTG | chr3R:9684476-9686476 | 1298 | 1312 | + | 9.8429 | 5.00E-06 | 0.286 | GCTTTGGACACACTG |
| 1 | GMTACGGTCACACTG | chr3R:9691794-9693794 | 1300 | 1314 | + | 9.8429 | 5.00E-06 | 0.286 | GCTTTGGACACACTG |
| 1 | GMTACGGTCACACTG | chr2L:6556502-6558502 | 1305 | 1319 | + | 9.8429 | 5.00E-06 | 0.286 | ACCGCGGTCACACTG |
| 1 | GMTACGGTCACACTG | chr3R:9691775-9693775 | 1319 | 1333 | + | 9.8429 | 5.00E-06 | 0.286 | GCTTTGGACACACTG |
| 1 | GMTACGGTCACACTG | chr3L:272391-274391 | 1324 | 1338 | + | 9.8429 | 5.00E-06 | 0.286 | GCTCGGGGCACACTG |
| 1 | GMTACGGTCACACTG | chr2L:19184726-19186726 | 1326 | 1340 | + | 9.8429 | 5.00E-06 | 0.286 | GCAACGTGCACACTG |
| 1 | GMTACGGTCACACTG | chrX:19622114-19624114 | 1371 | 1385 | + | 9.8429 | 5.00E-06 | 0.286 | GATTCGGTCCCGCTG |
| 1 | GMTACGGTCACACTG | chrX:17202112-17204112 | 1381 | 1395 | + | 9.8429 | 5.00E-06 | 0.286 | GCGGCGGTTACACTG |
| 1 | GMTACGGTCACACTG | chr2R:20617217-20619217 | 1402 | 1416 | + | 9.8429 | 5.00E-06 | 0.286 | CCTGCGGTCTCACTG |
| 1 | GMTACGGTCACACTG | chr2L:19033444-19035444 | 1406 | 1420 | + | 9.8429 | 5.00E-06 | 0.286 | GCAACGGTGTCACTG |
| 1 | GMTACGGTCACACTG | chrX:19622074-19624074 | 1411 | 1425 | + | 9.8429 | 5.00E-06 | 0.286 | GATTCGGTCCCGCTG |
| 1 | GMTACGGTCACACTG | chr2L:3170092-3172092 | 1417 | 1431 | + | 9.8429 | 5.00E-06 | 0.286 | GCACCGATCACACTG |
| 1 | GMTACGGTCACACTG | chr2R:13622044-13624044 | 1420 | 1434 | + | 9.8429 | 5.00E-06 | 0.286 | GCTGCGGTTTCACTG |
| 1 | GMTACGGTCACACTG | chrX:5763203-5765203 | 1422 | 1436 | + | 9.8429 | 5.00E-06 | 0.286 | GCCACGGCCAAACTG |
| 1 | GMTACGGTCACACTG | chr2R:8598962-8600962 | 1423 | 1437 | + | 9.8429 | 5.00E-06 | 0.286 | GCGATGGCCACACTG |
| 1 | GMTACGGTCACACTG | chr3L:11707553-11709553 | 1440 | 1454 | + | 9.8429 | 5.00E-06 | 0.286 | GCTGCGGCCACACTA |
| 1 | GMTACGGTCACACTG | chr3R:16651319-16653319 | 1454 | 1468 | + | 9.8429 | 5.00E-06 | 0.286 | GCTGTGGGCACACTG |
| 1 | GMTACGGTCACACTG | chr2L:13202295-13204295 | 1461 | 1475 | + | 9.8429 | 5.00E-06 | 0.286 | GATGCGGCCACTCTG |
| 1 | GMTACGGTCACACTG | chr3R:7248795-7250795 | 1475 | 1489 | + | 9.8429 | 5.00E-06 | 0.286 | GACTCGGTCCCACTG |
| 1 | GMTACGGTCACACTG | chr3L:10881386-10883386 | 1480 | 1494 | + | 9.8429 | 5.00E-06 | 0.286 | ACTGCGGACACACTG |
| 1 | GMTACGGTCACACTG | chr2R:18439475-18441475 | 1495 | 1509 | + | 9.8429 | 5.00E-06 | 0.286 | TCCACGGCCACACTG |
| 1 | GMTACGGTCACACTG | chr2R:16866714-16868714 | 1501 | 1515 | + | 9.8429 | 5.00E-06 | 0.286 | GCGAAGGTCACTCTG |
| 1 | GMTACGGTCACACTG | chrX:14316150-14318150 | 1503 | 1517 | + | 9.8429 | 5.00E-06 | 0.286 | GCGACGGCCACACTT |
| 1 | GMTACGGTCACACTG | chr3R:7248738-7250738 | 1532 | 1546 | + | 9.8429 | 5.00E-06 | 0.286 | GACTCGGTCCCACTG |
| 1 | GMTACGGTCACACTG | chr3L:4625196-4627196 | 1534 | 1548 | + | 9.8429 | 5.00E-06 | 0.286 | TCCGCGGTCACACTG |
| 1 | GMTACGGTCACACTG | chr2L:3844716-3846716 | 1535 | 1549 | + | 9.8429 | 5.00E-06 | 0.286 | GCTACGGGCGCAGTG |
| 1 | GMTACGGTCACACTG | chrX:17841161-17843161 | 1535 | 1549 | + | 9.8429 | 5.00E-06 | 0.286 | GCTACTGTCTCACCG |
| 1 | GMTACGGTCACACTG | chr3R:16651319-16653319 | 1568 | 1582 | + | 9.8429 | 5.00E-06 | 0.286 | GCCACTGTCACCCTG |
| 1 | GMTACGGTCACACTG | chr3R:10778569-10780569 | 1622 | 1636 | + | 9.8429 | 5.00E-06 | 0.286 | GCTACGGCCGCAGTG |
| 1 | GMTACGGTCACACTG | chr2R:13506557-13508557 | 1623 | 1637 | + | 9.8429 | 5.00E-06 | 0.286 | ACTACGGCCACACAG |

| | | | | | | | | | |
|---|---|---|---|---|---|---|---|---|---|
| 1 | GMTACGGTCACACTG | chr3R:9554785-9556785 | 1658 | 1672 | + | 9.8429 | 5.00E-06 | 0.286 | GCTACGGCAACACGG |
| 1 | GMTACGGTCACACTG | chrX:17147264-17149264 | 1718 | 1732 | + | 9.8429 | 5.00E-06 | 0.286 | TCTCCGGCCACACTG |
| 1 | GMTACGGTCACACTG | chr2L:8205171-8207171 | 1720 | 1734 | + | 9.8429 | 5.00E-06 | 0.286 | GCTACGGTGGCACAG |
| 1 | GMTACGGTCACACTG | chr2L:4793549-4795549 | 1728 | 1742 | + | 9.8429 | 5.00E-06 | 0.286 | GCTACGGTCACGCCT |
| 1 | GMTACGGTCACACTG | chr2R:8088482-8090482 | 1775 | 1789 | + | 9.8429 | 5.00E-06 | 0.286 | GCAAGGGTCACGCTG |
| 1 | GMTACGGTCACACTG | chr3L:22549226-22551226 | 1798 | 1812 | + | 9.8429 | 5.00E-06 | 0.286 | GCTACGAGCCCACTG |
| 1 | GMTACGGTCACACTG | chr3L:20199093-20201093 | 1832 | 1846 | + | 9.8429 | 5.00E-06 | 0.286 | GCCACGCCCACACTG |
| 1 | GMTACGGTCACACTG | chr2L:8205044-8207044 | 1847 | 1861 | + | 9.8429 | 5.00E-06 | 0.286 | GCTACGGTGGCACAG |
| 1 | GMTACGGTCACACTG | chrX:17201598-17203598 | 1895 | 1909 | + | 9.8429 | 5.00E-06 | 0.286 | GCGGCGGTTACACTG |
| 1 | GMTACGGTCACACTG | chr3R:18904655-18906655 | 1916 | 1930 | + | 9.8429 | 5.00E-06 | 0.286 | CCACCGGTCACACTG |
| 1 | GMTACGGTCACACTG | chr3R:8289578-8291578 | 1926 | 1940 | + | 9.8429 | 5.00E-06 | 0.286 | GCCAAGGACACACTG |
| 1 | GMTACGGTCACACTG | chr2R:11890135-11892135 | 1929 | 1943 | + | 9.8429 | 5.00E-06 | 0.286 | GCTTCGGTCACACCA |
| 1 | GMTACGGTCACACTG | chr3L:20220814-20222814 | 1931 | 1945 | + | 9.8429 | 5.00E-06 | 0.286 | GCGACGGTCGGACTG |
| 1 | GMTACGGTCACACTG | chr3L:11828030-11830030 | 4 | 18 | - | 9.8429 | 5.00E-06 | 0.286 | GCTTCAGTCGCACTG |
| 1 | GMTACGGTCACACTG | chr2L:9937790-9939790 | 5 | 19 | - | 9.8429 | 5.00E-06 | 0.286 | GCGACCGACACACTG |
| 1 | GMTACGGTCACACTG | chr3R:17131662-17133662 | 27 | 41 | - | 9.8429 | 5.00E-06 | 0.286 | TCCACGGTCACTCTG |
| 1 | GMTACGGTCACACTG | chr3L:20836488-20838488 | 46 | 60 | - | 9.8429 | 5.00E-06 | 0.286 | GCAACTGTCTCACTG |
| 1 | GMTACGGTCACACTG | chrX:16885818-16887818 | 87 | 101 | - | 9.8429 | 5.00E-06 | 0.286 | GATGCGGGCCCACTG |
| 1 | GMTACGGTCACACTG | chrX:15939451-15941451 | 88 | 102 | - | 9.8429 | 5.00E-06 | 0.286 | GCTTCGGTCACTATG |
| 1 | GMTACGGTCACACTG | chr3R:15766438-15768438 | 99 | 113 | - | 9.8429 | 5.00E-06 | 0.286 | GCTGCTGTCCCACTG |
| 1 | GMTACGGTCACACTG | chr2R:15379988-15381988 | 101 | 115 | - | 9.8429 | 5.00E-06 | 0.286 | GATCCGGACGCACTG |
| 1 | GMTACGGTCACACTG | chr2L:19078028-19080028 | 101 | 115 | - | 9.8429 | 5.00E-06 | 0.286 | GCTACGGCCAATCTG |
| 1 | GMTACGGTCACACTG | chr2L:21142229-21144229 | 125 | 139 | - | 9.8429 | 5.00E-06 | 0.286 | GCGACGGTCACCCTC |
| 1 | GMTACGGTCACACTG | chr2R:13197337-13199337 | 149 | 163 | - | 9.8429 | 5.00E-06 | 0.286 | GCTCCGGTAACACCG |
| 1 | GMTACGGTCACACTG | chrX:21318296-21320296 | 170 | 184 | - | 9.8429 | 5.00E-06 | 0.286 | GAGTCGGTCCCACTG |
| 1 | GMTACGGTCACACTG | chrX:21318296-21320296 | 170 | 184 | - | 9.8429 | 5.00E-06 | 0.286 | GAGTCGGTCCCACTG |
| 1 | GMTACGGTCACACTG | chr3L:12804902-12806902 | 189 | 203 | - | 9.8429 | 5.00E-06 | 0.286 | CCTGCGGCCACACTG |
| 1 | GMTACGGTCACACTG | chr2L:5980764-5982764 | 213 | 227 | - | 9.8429 | 5.00E-06 | 0.286 | TCGACGGTCACTCTG |
| 1 | GMTACGGTCACACTG | chrX:6526888-6528888 | 224 | 238 | - | 9.8429 | 5.00E-06 | 0.286 | GCAACGGTCACAAAG |
| 1 | GMTACGGTCACACTG | chr2R:17725914-17727914 | 225 | 239 | - | 9.8429 | 5.00E-06 | 0.286 | GCTCCCGGCACACTG |
| 1 | GMTACGGTCACACTG | chr2R:17725914-17727914 | 225 | 239 | - | 9.8429 | 5.00E-06 | 0.286 | GCTCCCGGCACACTG |
| 1 | GMTACGGTCACACTG | chr2R:8723905-8725905 | 262 | 276 | - | 9.8429 | 5.00E-06 | 0.286 | GCACCGGTCACACTT |
| 1 | GMTACGGTCACACTG | chr3R:22725967-22727967 | 285 | 299 | - | 9.8429 | 5.00E-06 | 0.286 | GCTACGGACACGGTG |

| | | | | | | | | | |
|---|---|---|---|---|---|---|---|---|---|
| 1 | GMTACGGTCACACTG | chr2R:17725847-17727847 | 292 | 306 | - | 9.8429 | 5.00E-06 | 0.286 | GCTCCCGGCACACTG |
| 1 | GMTACGGTCACACTG | chr2L:13175423-13177423 | 307 | 321 | - | 9.8429 | 5.00E-06 | 0.286 | GATGCGGCCACTCTG |
| 1 | GMTACGGTCACACTG | chrX:16885560-16887560 | 345 | 359 | - | 9.8429 | 5.00E-06 | 0.286 | GATGCGGGCCCACTG |
| 1 | GMTACGGTCACACTG | chr2R:15379724-15381724 | 365 | 379 | - | 9.8429 | 5.00E-06 | 0.286 | GATCCGGACGCACTG |
| 1 | GMTACGGTCACACTG | chr2L:18158272-18160272 | 378 | 392 | - | 9.8429 | 5.00E-06 | 0.286 | GCGATGGTCACACAG |
| 1 | GMTACGGTCACACTG | chr3L:8844814-8846814 | 398 | 412 | - | 9.8429 | 5.00E-06 | 0.286 | GACACGGTCACCCAG |
| 1 | GMTACGGTCACACTG | chr3L:20836135-20838135 | 399 | 413 | - | 9.8429 | 5.00E-06 | 0.286 | GCAACTGTCTCACTG |
| 1 | GMTACGGTCACACTG | chrX:11089210-11091210 | 400 | 414 | - | 9.8429 | 5.00E-06 | 0.286 | GATGCGGACGCACTG |
| 1 | GMTACGGTCACACTG | chrX:17709144-17711144 | 409 | 423 | - | 9.8429 | 5.00E-06 | 0.286 | GCCACGGTCACTCTA |
| 1 | GMTACGGTCACACTG | chr2R:23605654-23607654 | 410 | 424 | - | 9.8429 | 5.00E-06 | 0.286 | GATTCGGTCTCACCG |
| 1 | GMTACGGTCACACTG | chrX:15066007-15068007 | 414 | 428 | - | 9.8429 | 5.00E-06 | 0.286 | TCTGCGGGCACACTG |
| 1 | GMTACGGTCACACTG | chr3R:5052738-5054738 | 444 | 458 | - | 9.8429 | 5.00E-06 | 0.286 | ACTACGGACTCACTG |
| 1 | GMTACGGTCACACTG | chr2R:11218771-11220771 | 456 | 470 | - | 9.8429 | 5.00E-06 | 0.286 | GCTACTGCCCCACTG |
| 1 | GMTACGGTCACACTG | chr2L:6919704-6921704 | 462 | 476 | - | 9.8429 | 5.00E-06 | 0.286 | TCCACGGTCACTCTG |
| 1 | GMTACGGTCACACTG | chr2R:9163967-9165967 | 468 | 482 | - | 9.8429 | 5.00E-06 | 0.286 | GCACAGGTCACACTG |
| 1 | GMTACGGTCACACTG | chr2L:7222081-7224081 | 489 | 503 | - | 9.8429 | 5.00E-06 | 0.286 | GCCACGTACACACTG |
| 1 | GMTACGGTCACACTG | chr3R:17131198-17133198 | 491 | 505 | - | 9.8429 | 5.00E-06 | 0.286 | TCCACGGTCACTCTG |
| 1 | GMTACGGTCACACTG | chr3L:4321259-4323259 | 498 | 512 | - | 9.8429 | 5.00E-06 | 0.286 | GATCCGGTCGCCCTG |
| 1 | GMTACGGTCACACTG | chr2L:13202762-13204762 | 508 | 522 | - | 9.8429 | 5.00E-06 | 0.286 | CCCACGGTCACACAG |
| 1 | GMTACGGTCACACTG | chr3L:4253776-4255776 | 511 | 525 | - | 9.8429 | 5.00E-06 | 0.286 | GCGGTGGTCACACTG |
| 1 | GMTACGGTCACACTG | chr3L:16711126-16713126 | 526 | 540 | - | 9.8429 | 5.00E-06 | 0.286 | GCTGCGGTCACTCTA |
| 1 | GMTACGGTCACACTG | chr3L:16711126-16713126 | 551 | 565 | - | 9.8429 | 5.00E-06 | 0.286 | GCAACGGTCACTTTG |
| 1 | GMTACGGTCACACTG | chr2L:6479930-6481930 | 565 | 579 | - | 9.8429 | 5.00E-06 | 0.286 | CCAACGGCCACACTG |
| 1 | GMTACGGTCACACTG | chr2L:6479929-6481929 | 566 | 580 | - | 9.8429 | 5.00E-06 | 0.286 | CCAACGGCCACACTG |
| 1 | GMTACGGTCACACTG | chr3L:15987236-15989236 | 571 | 585 | - | 9.8429 | 5.00E-06 | 0.286 | GCTGCGGTCACTCTA |
| 1 | GMTACGGTCACACTG | chr3L:21941221-21943221 | 591 | 605 | - | 9.8429 | 5.00E-06 | 0.286 | GCCACGGCCACACTC |
| 1 | GMTACGGTCACACTG | chr2L:15006813-15008813 | 593 | 607 | - | 9.8429 | 5.00E-06 | 0.286 | GCAACGGTCACTTTG |
| 1 | GMTACGGTCACACTG | chr3R:5684855-5686855 | 594 | 608 | - | 9.8429 | 5.00E-06 | 0.286 | GCCATGGTCGCACTG |
| 1 | GMTACGGTCACACTG | chr3L:23149499-23151499 | 604 | 618 | - | 9.8429 | 5.00E-06 | 0.286 | GCGACTGTCACACCG |
| 1 | GMTACGGTCACACTG | chr3L:9072105-9074105 | 611 | 625 | - | 9.8429 | 5.00E-06 | 0.286 | GCGACGGTCACTCTA |
| 1 | GMTACGGTCACACTG | chrX:15703125-15705125 | 618 | 632 | - | 9.8429 | 5.00E-06 | 0.286 | GCTGCGGTCACACCA |
| 1 | GMTACGGTCACACTG | chr3R:31771219-31773219 | 630 | 644 | - | 9.8429 | 5.00E-06 | 0.286 | GCGCTGGTCACACTG |
| 1 | GMTACGGTCACACTG | chr2L:6047002-6049002 | 642 | 656 | - | 9.8429 | 5.00E-06 | 0.286 | GAGACGGTCACGCAG |

| | | | | | | | | | |
|---|---|---|---|---|---|---|---|---|---|
| 1 | GMTACGGTCACACTG | chr3L:9441267-9443267 | 655 | 669 | - | 9.8429 | 5.00E-06 | 0.286 | GACACGGACACACAG |
| 1 | GMTACGGTCACACTG | chr3L:9441267-9443267 | 655 | 669 | - | 9.8429 | 5.00E-06 | 0.286 | GACACGGACACACAG |
| 1 | GMTACGGTCACACTG | chr2L:11516391-11518391 | 659 | 673 | - | 9.8429 | 5.00E-06 | 0.286 | TCCGCGGTCACACTG |
| 1 | GMTACGGTCACACTG | chr2L:9937134-9939134 | 661 | 675 | - | 9.8429 | 5.00E-06 | 0.286 | GCGACCGACACACTG |
| 1 | GMTACGGTCACACTG | chr3R:5600433-5602433 | 664 | 678 | - | 9.8429 | 5.00E-06 | 0.286 | GCCACGGTCATACCG |
| 1 | GMTACGGTCACACTG | chr3R:24031157-24033157 | 664 | 678 | - | 9.8429 | 5.00E-06 | 0.286 | GCTCTGGCCACACTG |
| 1 | GMTACGGTCACACTG | chr2L:20296299-20298299 | 683 | 697 | - | 9.8429 | 5.00E-06 | 0.286 | GCTTCTGACACACTG |
| 1 | GMTACGGTCACACTG | chr2R:11218539-11220539 | 688 | 702 | - | 9.8429 | 5.00E-06 | 0.286 | GCTACTGCCCCACTG |
| 1 | GMTACGGTCACACTG | chr3R:27111566-27113566 | 693 | 707 | - | 9.8429 | 5.00E-06 | 0.286 | GCGCTGGTCACACTG |
| 1 | GMTACGGTCACACTG | chr2L:18968028-18970028 | 704 | 718 | - | 9.8429 | 5.00E-06 | 0.286 | GCCCTGGTCACACTG |
| 1 | GMTACGGTCACACTG | chr2L:11987270-11989270 | 729 | 743 | - | 9.8429 | 5.00E-06 | 0.286 | ACTGCGGTCACACCG |
| 1 | GMTACGGTCACACTG | chr3R:30745290-30747290 | 737 | 751 | - | 9.8429 | 5.00E-06 | 0.286 | GATGCGGACACCCTG |
| 1 | GMTACGGTCACACTG | chr2R:20256650-20258650 | 737 | 751 | - | 9.8429 | 5.00E-06 | 0.286 | GCACTGGTCACACTG |
| 1 | GMTACGGTCACACTG | chr3R:8779789-8781789 | 742 | 756 | - | 9.8429 | 5.00E-06 | 0.286 | GCTACGGAGGCACTG |
| 1 | GMTACGGTCACACTG | chr3R:27287769-27289769 | 747 | 761 | - | 9.8429 | 5.00E-06 | 0.286 | GCGACAGTCACACCG |
| 1 | GMTACGGTCACACTG | chrX:16799930-16801930 | 752 | 766 | - | 9.8429 | 5.00E-06 | 0.286 | CCGCCGGTCACACTG |
| 1 | GMTACGGTCACACTG | chr3R:31579951-31581951 | 771 | 785 | - | 9.8429 | 5.00E-06 | 0.286 | GCTTTGGCCACACTG |
| 1 | GMTACGGTCACACTG | chr2R:22936279-22938279 | 777 | 791 | - | 9.8429 | 5.00E-06 | 0.286 | ACCACGGACACACTG |
| 1 | GMTACGGTCACACTG | chr3L:16669741-16671741 | 800 | 814 | - | 9.8429 | 5.00E-06 | 0.286 | GCTCCGCTCCCACTG |
| 1 | GMTACGGTCACACTG | chr2R:7495466-7497466 | 808 | 822 | - | 9.8429 | 5.00E-06 | 0.286 | GCTGCTGCCACACTG |
| 1 | GMTACGGTCACACTG | chr3R:22725414-22727414 | 838 | 852 | - | 9.8429 | 5.00E-06 | 0.286 | GCTACGGACACGGTG |
| 1 | GMTACGGTCACACTG | chr2R:13221459-13223459 | 844 | 858 | - | 9.8429 | 5.00E-06 | 0.286 | CCAACGGTCACCCTG |
| 1 | GMTACGGTCACACTG | chr2L:1076948-1078948 | 858 | 872 | - | 9.8429 | 5.00E-06 | 0.286 | GCCGCGGTCACACTC |
| 1 | GMTACGGTCACACTG | chr2R:13221433-13223433 | 870 | 884 | - | 9.8429 | 5.00E-06 | 0.286 | CCAACGGTCACCCTG |
| 1 | GMTACGGTCACACTG | chr2R:8173165-8175165 | 870 | 884 | - | 9.8429 | 5.00E-06 | 0.286 | ACTCCGGTCACACCG |
| 1 | GMTACGGTCACACTG | chr2R:23073823-23075823 | 880 | 894 | - | 9.8429 | 5.00E-06 | 0.286 | GCTGCGGCCACACTC |
| 1 | GMTACGGTCACACTG | chr3L:20200638-20202638 | 893 | 907 | - | 9.8429 | 5.00E-06 | 0.286 | GAGCCGGTCACGCTG |
| 1 | GMTACGGTCACACTG | chrX:17708653-17710653 | 900 | 914 | - | 9.8429 | 5.00E-06 | 0.286 | GCCACGGTCACTCTA |
| 1 | GMTACGGTCACACTG | chr2R:8723258-8725258 | 909 | 923 | - | 9.8429 | 5.00E-06 | 0.286 | GCACCGGTCACACTT |
| 1 | GMTACGGTCACACTG | chrX:16639968-16641968 | 910 | 924 | - | 9.8429 | 5.00E-06 | 0.286 | GCCGGGGTCACACTG |
| 1 | GMTACGGTCACACTG | chr3R:10851724-10853724 | 913 | 927 | - | 9.8429 | 5.00E-06 | 0.286 | GCCTCGGTCACACTC |
| 1 | GMTACGGTCACACTG | chr3R:15315471-15317471 | 917 | 931 | - | 9.8429 | 5.00E-06 | 0.286 | GCTACCGTCGCACCG |
| 1 | GMTACGGTCACACTG | chr2L:6479575-6481575 | 920 | 934 | - | 9.8429 | 5.00E-06 | 0.286 | CCAACGGCCACACTG |

| | | | | | | | | | |
|---|---|---|---|---|---|---|---|---|---|
| 1 | GMTACGGTCACACTG | chr2L:8071895-8073895 | 937 | 951 | - | 9.8429 | 5.00E-06 | 0.286 | GCACTGGTCACACTG |
| 1 | GMTACGGTCACACTG | chrX:8060645-8062645 | 941 | 955 | - | 9.8429 | 5.00E-06 | 0.286 | GCTCTGGTCACACCG |
| 1 | GMTACGGTCACACTG | chr3L:7868042-7870042 | 943 | 957 | - | 9.8429 | 5.00E-06 | 0.286 | GCTTGGGTCACCCTG |
| 1 | GMTACGGTCACACTG | chr3L:211709-213709 | 947 | 961 | - | 9.8429 | 5.00E-06 | 0.286 | GCTTCGGCCACACTC |
| 1 | GMTACGGTCACACTG | chr2L:6919210-6921210 | 956 | 970 | - | 9.8429 | 5.00E-06 | 0.286 | TCCACGGTCACTCTG |
| 1 | GMTACGGTCACACTG | chr2L:5980018-5982018 | 959 | 973 | - | 9.8429 | 5.00E-06 | 0.286 | TCGACGGTCACTCTG |
| 1 | GMTACGGTCACACTG | chr2L:11987039-11989039 | 960 | 974 | - | 9.8429 | 5.00E-06 | 0.286 | ACTGCGGTCACACCG |
| 1 | GMTACGGTCACACTG | chr3R:5600136-5602136 | 961 | 975 | - | 9.8429 | 5.00E-06 | 0.286 | GCCACGGTCATACCG |
| 1 | GMTACGGTCACACTG | chrX:3670801-3672801 | 963 | 977 | - | 9.8429 | 5.00E-06 | 0.286 | GCAACGGACACAGTG |
| 1 | GMTACGGTCACACTG | chr2L:11516081-11518081 | 969 | 983 | - | 9.8429 | 5.00E-06 | 0.286 | TCCGCGGTCACACTG |
| 1 | GMTACGGTCACACTG | chr3L:211687-213687 | 969 | 983 | - | 9.8429 | 5.00E-06 | 0.286 | GCTTCGGCCACACTC |
| 1 | GMTACGGTCACACTG | chrX:17626895-17628895 | 971 | 985 | - | 9.8429 | 5.00E-06 | 0.286 | CCTTCGGCCACACTG |
| 1 | GMTACGGTCACACTG | chrX:16799710-16801710 | 972 | 986 | - | 9.8429 | 5.00E-06 | 0.286 | CCGCCGGTCACACTG |
| 1 | GMTACGGTCACACTG | chrX:15702770-15704770 | 973 | 987 | - | 9.8429 | 5.00E-06 | 0.286 | GCTGCGGTCACACCA |
| 1 | GMTACGGTCACACTG | chr3L:16710677-16712677 | 975 | 989 | - | 9.8429 | 5.00E-06 | 0.286 | GCTGCGGTCACTCTA |
| 1 | GMTACGGTCACACTG | chr3R:4811096-4813096 | 975 | 989 | - | 9.8429 | 5.00E-06 | 0.286 | GACGCGGTCTCACTG |
| 1 | GMTACGGTCACACTG | chr3R:4811096-4813096 | 975 | 989 | - | 9.8429 | 5.00E-06 | 0.286 | GACGCGGTCTCACTG |
| 1 | GMTACGGTCACACTG | chr3L:16710677-16712677 | 975 | 989 | - | 9.8429 | 5.00E-06 | 0.286 | GCTGCGGTCACTCTA |
| 1 | GMTACGGTCACACTG | chr2L:15006431-15008431 | 975 | 989 | - | 9.8429 | 5.00E-06 | 0.286 | GCAACGGTCACTTTG |
| 1 | GMTACGGTCACACTG | chr2L:13202295-13204295 | 975 | 989 | - | 9.8429 | 5.00E-06 | 0.286 | CCCACGGTCACACAG |
| 1 | GMTACGGTCACACTG | chr3L:12804116-12806116 | 975 | 989 | - | 9.8429 | 5.00E-06 | 0.286 | CCTGCGGCCACACTG |
| 1 | GMTACGGTCACACTG | chr2R:23073725-23075725 | 978 | 992 | - | 9.8429 | 5.00E-06 | 0.286 | GCTGCGGCCACACTC |
| 1 | GMTACGGTCACACTG | chrX:13829317-13831317 | 978 | 992 | - | 9.8429 | 5.00E-06 | 0.286 | GCGGCGGTCACACTT |
| 1 | GMTACGGTCACACTG | chr2R:22936077-22938077 | 979 | 993 | - | 9.8429 | 5.00E-06 | 0.286 | ACCACGGACACACTG |
| 1 | GMTACGGTCACACTG | chr3R:5216456-5218456 | 980 | 994 | - | 9.8429 | 5.00E-06 | 0.286 | CCTGCGGTCACACCG |
| 1 | GMTACGGTCACACTG | chr3L:4253306-4255306 | 981 | 995 | - | 9.8429 | 5.00E-06 | 0.286 | GCGGTGGTCACACTG |
| 1 | GMTACGGTCACACTG | chr2R:7532506-7534506 | 982 | 996 | - | 9.8429 | 5.00E-06 | 0.286 | CCGGCGGTCACACTG |
| 1 | GMTACGGTCACACTG | chr2L:17472215-17474215 | 983 | 997 | - | 9.8429 | 5.00E-06 | 0.286 | CCTGCGGTCGCACTG |
| 1 | GMTACGGTCACACTG | chr3L:14801168-14803168 | 984 | 998 | - | 9.8429 | 5.00E-06 | 0.286 | TCCGCGGTCACACTG |
| 1 | GMTACGGTCACACTG | chr2L:8071848-8073848 | 984 | 998 | - | 9.8429 | 5.00E-06 | 0.286 | GCACTGGTCACACTG |
| 1 | GMTACGGTCACACTG | chr3R:25032465-25034465 | 985 | 999 | - | 9.8429 | 5.00E-06 | 0.286 | CCACCGGTCACACTG |
| 1 | GMTACGGTCACACTG | chr3L:9071731-9073731 | 985 | 999 | - | 9.8429 | 5.00E-06 | 0.286 | GCGACGGTCACTCTA |
| 1 | GMTACGGTCACACTG | chr3R:23818237-23820237 | 986 | 1000 | - | 9.8429 | 5.00E-06 | 0.286 | GCAACGGTCACTCTT |

| | | | | | | | | | |
|---|---|---|---|---|---|---|---|---|---|
| 1 | GMTACGGTCACACTG | chr3R:23818237-23820237 | 986 | 1000 | - | 9.8429 | 5.00E-06 | 0.286 | GCAACGGTCACTCTT |
| 1 | GMTACGGTCACACTG | chr3L:12205022-12207022 | 986 | 1000 | - | 9.8429 | 5.00E-06 | 0.286 | GCCTTGGTCACACTG |
| 1 | GMTACGGTCACACTG | chr2L:18967744-18969744 | 988 | 1002 | - | 9.8429 | 5.00E-06 | 0.286 | GCCCTGGTCACACTG |
| 1 | GMTACGGTCACACTG | chr3R:27111270-27113270 | 989 | 1003 | - | 9.8429 | 5.00E-06 | 0.286 | GCGCTGGTCACACTG |
| 1 | GMTACGGTCACACTG | chrX:6362310-6364310 | 989 | 1003 | - | 9.8429 | 5.00E-06 | 0.286 | ACTTCGGCCACACTG |
| 1 | GMTACGGTCACACTG | chrX:15065432-15067432 | 989 | 1003 | - | 9.8429 | 5.00E-06 | 0.286 | TCTGCGGGCACACTG |
| 1 | GMTACGGTCACACTG | chr3R:27287526-27289526 | 990 | 1004 | - | 9.8429 | 5.00E-06 | 0.286 | GCGACAGTCACACCG |
| 1 | GMTACGGTCACACTG | chr2L:1076815-1078815 | 991 | 1005 | - | 9.8429 | 5.00E-06 | 0.286 | GCCGCGGTCACACTC |
| 1 | GMTACGGTCACACTG | chr2R:7532497-7534497 | 991 | 1005 | - | 9.8429 | 5.00E-06 | 0.286 | CCGGCGGTCACACTG |
| 1 | GMTACGGTCACACTG | chr3L:15986815-15988815 | 992 | 1006 | - | 9.8429 | 5.00E-06 | 0.286 | GCTGCGGTCACTCTA |
| 1 | GMTACGGTCACACTG | chrX:6275724-6277724 | 993 | 1007 | - | 9.8429 | 5.00E-06 | 0.286 | GAGCCGGCCACACTG |
| 1 | GMTACGGTCACACTG | chr3R:24030827-24032827 | 994 | 1008 | - | 9.8429 | 5.00E-06 | 0.286 | GCTCTGGCCACACTG |
| 1 | GMTACGGTCACACTG | chr2L:850071-852071 | 995 | 1009 | - | 9.8429 | 5.00E-06 | 0.286 | CCAACGGTCACTCTG |
| 1 | GMTACGGTCACACTG | chr3L:1533034-1535034 | 995 | 1009 | - | 9.8429 | 5.00E-06 | 0.286 | CCTGCGGTCACTCTG |
| 1 | GMTACGGTCACACTG | chr2R:20256390-20258390 | 997 | 1011 | - | 9.8429 | 5.00E-06 | 0.286 | GCACTGGTCACACTG |
| 1 | GMTACGGTCACACTG | chr3L:16710677-16712677 | 1000 | 1014 | - | 9.8429 | 5.00E-06 | 0.286 | GCAACGGTCACTTTG |
| 1 | GMTACGGTCACACTG | chr3L:16710677-16712677 | 1000 | 1014 | - | 9.8429 | 5.00E-06 | 0.286 | GCAACGGTCACTTTG |
| 1 | GMTACGGTCACACTG | chr3L:21940812-21942812 | 1000 | 1014 | - | 9.8429 | 5.00E-06 | 0.286 | GCCACGGCCACACTC |
| 1 | GMTACGGTCACACTG | chr3R:31770848-31772848 | 1001 | 1015 | - | 9.8429 | 5.00E-06 | 0.286 | GCGCTGGTCACACTG |
| 1 | GMTACGGTCACACTG | chr2L:21141347-21143347 | 1007 | 1021 | - | 9.8429 | 5.00E-06 | 0.286 | GCGACGGTCACCCTC |
| 1 | GMTACGGTCACACTG | chr3R:5684434-5686434 | 1015 | 1029 | - | 9.8429 | 5.00E-06 | 0.286 | GCCATGGTCGCACTG |
| 1 | GMTACGGTCACACTG | chr3R:31579699-31581699 | 1023 | 1037 | - | 9.8429 | 5.00E-06 | 0.286 | GCTTTGGCCACACTG |
| 1 | GMTACGGTCACACTG | chr3L:15986784-15988784 | 1023 | 1037 | - | 9.8429 | 5.00E-06 | 0.286 | GCTGCGGTCACTCTA |
| 1 | GMTACGGTCACACTG | chr2R:12199815-12201815 | 1026 | 1040 | - | 9.8429 | 5.00E-06 | 0.286 | GCGACGGTCCCACTT |
| 1 | GMTACGGTCACACTG | chrX:5321696-5323696 | 1027 | 1041 | - | 9.8429 | 5.00E-06 | 0.286 | GCACTGGTCACACTG |
| 1 | GMTACGGTCACACTG | chr3R:6007579-6009579 | 1030 | 1044 | - | 9.8429 | 5.00E-06 | 0.286 | GCTGCGGTCACACCA |
| 1 | GMTACGGTCACACTG | chr3R:6214832-6216832 | 1035 | 1049 | - | 9.8429 | 5.00E-06 | 0.286 | GCTACGGTCTTGCTG |
| 1 | GMTACGGTCACACTG | chr3R:13433984-13435984 | 1047 | 1061 | - | 9.8429 | 5.00E-06 | 0.286 | TCGACGGTCACACAG |
| 1 | GMTACGGTCACACTG | chr3R:4402076-4404076 | 1057 | 1071 | - | 9.8429 | 5.00E-06 | 0.286 | GCTACGGCTACACAG |
| 1 | GMTACGGTCACACTG | chr3R:12673608-12675608 | 1058 | 1072 | - | 9.8429 | 5.00E-06 | 0.286 | GCAACGAACACACTG |
| 1 | GMTACGGTCACACTG | chr3L:6963758-6965758 | 1083 | 1097 | - | 9.8429 | 5.00E-06 | 0.286 | CCCGCGGTCACACTG |
| 1 | GMTACGGTCACACTG | chr2R:21673901-21675901 | 1142 | 1156 | - | 9.8429 | 5.00E-06 | 0.286 | GCCACGTCCACACTG |
| 1 | GMTACGGTCACACTG | chrX:8060436-8062436 | 1150 | 1164 | - | 9.8429 | 5.00E-06 | 0.286 | GCTCTGGTCACACCG |

| | | | | | | | | | |
|---|---|---|---|---|---|---|---|---|---|
| 1 | GMTACGGTCACACTG | chr3L:7455149-7457149 | 1159 | 1173 | - | 9.8429 | 5.00E-06 | 0.286 | GATACGGTCTCCCGG |
| 1 | GMTACGGTCACACTG | chr2R:7949339-7951339 | 1172 | 1186 | - | 9.8429 | 5.00E-06 | 0.286 | GCCTCGGTCAAACTG |
| 1 | GMTACGGTCACACTG | chrX:6680767-6682767 | 1200 | 1214 | - | 9.8429 | 5.00E-06 | 0.286 | GCCACTGTCGCACTG |
| 1 | GMTACGGTCACACTG | chr3R:4401928-4403928 | 1205 | 1219 | - | 9.8429 | 5.00E-06 | 0.286 | GCTACGGCTACACAG |
| 1 | GMTACGGTCACACTG | chr3R:27503101-27505101 | 1216 | 1230 | - | 9.8429 | 5.00E-06 | 0.286 | GCTTTGGTCACACAG |
| 1 | GMTACGGTCACACTG | chr3L:7629508-7631508 | 1234 | 1248 | - | 9.8429 | 5.00E-06 | 0.286 | GCGACGGCGACACTG |
| 1 | GMTACGGTCACACTG | chr2R:14667391-14669391 | 1258 | 1272 | - | 9.8429 | 5.00E-06 | 0.286 | CCGACGGCCACACTG |
| 1 | GMTACGGTCACACTG | chrX:3670502-3672502 | 1262 | 1276 | - | 9.8429 | 5.00E-06 | 0.286 | GCAACGGACACAGTG |
| 1 | GMTACGGTCACACTG | chr3R:12696322-12698322 | 1274 | 1288 | - | 9.8429 | 5.00E-06 | 0.286 | GCGGCGGTCACACTT |
| 1 | GMTACGGTCACACTG | chr2L:12026438-12028438 | 1280 | 1294 | - | 9.8429 | 5.00E-06 | 0.286 | GCCACGGTTACACGG |
| 1 | GMTACGGTCACACTG | chr3R:9585942-9587942 | 1292 | 1306 | - | 9.8429 | 5.00E-06 | 0.286 | TCTACGGTCGCACCG |
| 1 | GMTACGGTCACACTG | chr3R:19729715-19731715 | 1307 | 1321 | - | 9.8429 | 5.00E-06 | 0.286 | GACACGGACACACAG |
| 1 | GMTACGGTCACACTG | chr3R:20730631-20732631 | 1317 | 1331 | - | 9.8429 | 5.00E-06 | 0.286 | GCTGCGGTCACAACG |
| 1 | GMTACGGTCACACTG | chrX:11088290-11090290 | 1320 | 1334 | - | 9.8429 | 5.00E-06 | 0.286 | GATGCGGACGCACTG |
| 1 | GMTACGGTCACACTG | chr2L:9911911-9913911 | 1325 | 1339 | - | 9.8429 | 5.00E-06 | 0.286 | GCCACCGGCACACTG |
| 1 | GMTACGGTCACACTG | chr2R:8722839-8724839 | 1328 | 1342 | - | 9.8429 | 5.00E-06 | 0.286 | GCACCGGTCACACTT |
| 1 | GMTACGGTCACACTG | chr3L:5779863-5781863 | 1332 | 1346 | - | 9.8429 | 5.00E-06 | 0.286 | GCCACGGGAACACTG |
| 1 | GMTACGGTCACACTG | chr3L:20835182-20837182 | 1352 | 1366 | - | 9.8429 | 5.00E-06 | 0.286 | GCAACTGTCTCACTG |
| 1 | GMTACGGTCACACTG | chr3R:10734957-10736957 | 1356 | 1370 | - | 9.8429 | 5.00E-06 | 0.286 | CCTGCGGTCACACCG |
| 1 | GMTACGGTCACACTG | chr2L:849702-851702 | 1364 | 1378 | - | 9.8429 | 5.00E-06 | 0.286 | CCAACGGTCACTCTG |
| 1 | GMTACGGTCACACTG | chr3L:20139607-20141607 | 1372 | 1386 | - | 9.8429 | 5.00E-06 | 0.286 | TCATCGGTCACACTG |
| 1 | GMTACGGTCACACTG | chr2R:7664828-7666828 | 1375 | 1389 | - | 9.8429 | 5.00E-06 | 0.286 | GCACCGGTCACAATG |
| 1 | GMTACGGTCACACTG | chr2R:8172613-8174613 | 1422 | 1436 | - | 9.8429 | 5.00E-06 | 0.286 | ACTCCGGTCACACCG |
| 1 | GMTACGGTCACACTG | chrX:16639419-16641419 | 1459 | 1473 | - | 9.8429 | 5.00E-06 | 0.286 | GCCGGGGTCACACTG |
| 1 | GMTACGGTCACACTG | chr3L:3883801-3885801 | 1459 | 1473 | - | 9.8429 | 5.00E-06 | 0.286 | GCTACGGCCACACAA |
| 1 | GMTACGGTCACACTG | chr2R:23382483-23384483 | 1471 | 1485 | - | 9.8429 | 5.00E-06 | 0.286 | GCTGGGGTCACACCG |
| 1 | GMTACGGTCACACTG | chr2R:7531981-7533981 | 1507 | 1521 | - | 9.8429 | 5.00E-06 | 0.286 | CCGGCGGTCACACTG |
| 1 | GMTACGGTCACACTG | chr2R:15895427-15897427 | 1529 | 1543 | - | 9.8429 | 5.00E-06 | 0.286 | GAGACGGACCCACTG |
| 1 | GMTACGGTCACACTG | chr2R:23382424-23384424 | 1530 | 1544 | - | 9.8429 | 5.00E-06 | 0.286 | GCTGGGGTCACACCG |
| 1 | GMTACGGTCACACTG | chr2L:9936232-9938232 | 1563 | 1577 | - | 9.8429 | 5.00E-06 | 0.286 | GCGACCGACACACTG |
| 1 | GMTACGGTCACACTG | chr3L:23148530-23150530 | 1573 | 1587 | - | 9.8429 | 5.00E-06 | 0.286 | GCGACTGTCACACCG |
| 1 | GMTACGGTCACACTG | chr2L:20857695-20859695 | 1598 | 1612 | - | 9.8429 | 5.00E-06 | 0.286 | GCCACGGCCAAACTG |
| 1 | GMTACGGTCACACTG | chr3R:8778923-8780923 | 1608 | 1622 | - | 9.8429 | 5.00E-06 | 0.286 | GCTACGGAGGCACTG |

| | | | | | | | | | |
|---|---|---|---|---|---|---|---|---|---|
| 1 | GMTACGGTCACACTG | chr3R:4810444-4812444 | 1627 | 1641 | - | 9.8429 | 5.00E-06 | 0.286 | GACGCGGTCTCACTG |
| 1 | GMTACGGTCACACTG | chr2L:9911608-9913608 | 1628 | 1642 | - | 9.8429 | 5.00E-06 | 0.286 | GCCACCGGCACACTG |
| 1 | GMTACGGTCACACTG | chr2L:8302429-8304429 | 1647 | 1661 | - | 9.8429 | 5.00E-06 | 0.286 | GCTGCGGTCGCAGTG |
| 1 | GMTACGGTCACACTG | chr2R:7948839-7950839 | 1672 | 1686 | - | 9.8429 | 5.00E-06 | 0.286 | GCCTCGGTCAAACTG |
| 1 | GMTACGGTCACACTG | chrX:17300929-17302929 | 1715 | 1729 | - | 9.8429 | 5.00E-06 | 0.286 | GCGACGGTCACCATG |
| 1 | GMTACGGTCACACTG | chr2R:23072968-23074968 | 1735 | 1749 | - | 9.8429 | 5.00E-06 | 0.286 | GCTGCGGCCACACTC |
| 1 | GMTACGGTCACACTG | chr3R:30744263-30746263 | 1764 | 1778 | - | 9.8429 | 5.00E-06 | 0.286 | GATGCGGACACCCTG |
| 1 | GMTACGGTCACACTG | chr2R:20848939-20850939 | 1911 | 1925 | - | 9.8429 | 5.00E-06 | 0.286 | TCTACGGTCTCACAG |
| 1 | GMTACGGTCACACTG | chrX:15471164-15473164 | 1933 | 1947 | - | 9.8429 | 5.00E-06 | 0.286 | GCTCCGGTTACACAG |
| 1 | GMTACGGTCACACTG | chrX:21316522-21318522 | 1944 | 1958 | - | 9.8429 | 5.00E-06 | 0.286 | GAGTCGGTCCCACTG |
| 1 | GMTACGGTCACACTG | chrX:18482772-18484772 | 1949 | 1963 | - | 9.8429 | 5.00E-06 | 0.286 | GCCACCGTCACACCG |
| 1 | GMTACGGTCACACTG | chr2R:5759862-5761862 | 1971 | 1985 | - | 9.8429 | 5.00E-06 | 0.286 | GCTCCGGTGGCACTG |
| 1 | GMTACGGTCACACTG | chrX:15471119-15473119 | 1978 | 1992 | - | 9.8429 | 5.00E-06 | 0.286 | GCTCCGGTTACACAG |
| 1 | GMTACGGTCACACTG | chr2R:8236271-8238271 | 1980 | 1994 | - | 9.8429 | 5.00E-06 | 0.286 | GCCACTGGCACACTG |
| 1 | GMTACGGTCACACTG | chr3L:8411825-8413825 | 12 | 26 | - | 9.5643 | 1.06E-05 | 0.4 | GCTTTGGTCATACTG |
| 1 | GMTACGGTCACACTG | chr2R:23125042-23127042 | 28 | 42 | - | 9.5643 | 1.06E-05 | 0.4 | CCTACGGACAAACTG |
| 1 | GMTACGGTCACACTG | chr2R:25250927-25252927 | 47 | 61 | - | 9.5643 | 1.06E-05 | 0.4 | GCCACGCTCACACTT |
| 1 | GMTACGGTCACACTG | chr3L:8963769-8965769 | 48 | 62 | - | 9.5643 | 1.06E-05 | 0.4 | GCTACGATCACAAAG |
| 1 | GMTACGGTCACACTG | chr2R:11590485-11592485 | 51 | 65 | - | 9.5643 | 1.06E-05 | 0.4 | GATCCGGTGACCCTG |
| 1 | GMTACGGTCACACTG | chrX:1574132-1576132 | 53 | 67 | - | 9.5643 | 1.06E-05 | 0.4 | CATACGGACACACAG |
| 1 | GMTACGGTCACACTG | chr2L:11113014-11115014 | 57 | 71 | - | 9.5643 | 1.06E-05 | 0.4 | GATACAGACACACCG |
| 1 | GMTACGGTCACACTG | chr2L:13256477-13258477 | 58 | 72 | - | 9.5643 | 1.06E-05 | 0.4 | GCTGCGGTCAGAATG |
| 1 | GMTACGGTCACACTG | chr3R:10787226-10789226 | 62 | 76 | - | 9.5643 | 1.06E-05 | 0.4 | GATAGGGGCACGCTG |
| 1 | GMTACGGTCACACTG | chrX:1348357-1350357 | 83 | 97 | - | 9.5643 | 1.06E-05 | 0.4 | GCTAAGATCCCACTG |
| 1 | GMTACGGTCACACTG | chr2L:10093088-10095088 | 95 | 109 | - | 9.5643 | 1.06E-05 | 0.4 | ACTGCAGTCACACTG |
| 1 | GMTACGGTCACACTG | chr3R:28302571-28304571 | 105 | 119 | - | 9.5643 | 1.06E-05 | 0.4 | GCTGCGGTCAAAATG |
| 1 | GMTACGGTCACACTG | chrX:3241084-3243084 | 111 | 125 | - | 9.5643 | 1.06E-05 | 0.4 | GCTACGTACACAATG |
| 1 | GMTACGGTCACACTG | chr2R:22666155-22668155 | 114 | 128 | - | 9.5643 | 1.06E-05 | 0.4 | GATAAGGCCCCACTG |
| 1 | GMTACGGTCACACTG | chr3L:16086525-16088525 | 114 | 128 | - | 9.5643 | 1.06E-05 | 0.4 | GATACGGACGAACTG |
| 1 | GMTACGGTCACACTG | chr2R:22666155-22668155 | 114 | 128 | - | 9.5643 | 1.06E-05 | 0.4 | GATAAGGCCCCACTG |
| 1 | GMTACGGTCACACTG | chr3L:12533590-12535590 | 126 | 140 | - | 9.5643 | 1.06E-05 | 0.4 | GCTCTGGTTACACTG |
| 1 | GMTACGGTCACACTG | chr2R:21847478-21849478 | 127 | 141 | - | 9.5643 | 1.06E-05 | 0.4 | GATAAGGACCCACTG |
| 1 | GMTACGGTCACACTG | chr2L:18453649-18455649 | 128 | 142 | - | 9.5643 | 1.06E-05 | 0.4 | GATCCGCTCACACAG |

| | | | | | | | | | |
|---|---|---|---|---|---|---|---|---|---|
| 1 | GMTACGGTCACACTG | chr3L:10318785-10320785 | 138 | 152 | - | 9.5643 | 1.06E-05 | 0.4 | GATACGTTCGCTCTG |
| 1 | GMTACGGTCACACTG | chr3R:5735355-5737355 | 146 | 160 | - | 9.5643 | 1.06E-05 | 0.4 | GATAAGGCCACCCTG |
| 1 | GMTACGGTCACACTG | chr4:776146-778146 | 150 | 164 | - | 9.5643 | 1.06E-05 | 0.4 | GATTCGGTCCCATTG |
| 1 | GMTACGGTCACACTG | chr2R:16868402-16870402 | 152 | 166 | - | 9.5643 | 1.06E-05 | 0.4 | GCCACGTTCACATTG |
| 1 | GMTACGGTCACACTG | chr2R:8951339-8953339 | 154 | 168 | - | 9.5643 | 1.06E-05 | 0.4 | AAAACGGTCACACCG |
| 1 | GMTACGGTCACACTG | chr3R:27953917-27955917 | 159 | 173 | - | 9.5643 | 1.06E-05 | 0.4 | GCTACGGTCACATCA |
| 1 | GMTACGGTCACACTG | chr2L:12711952-12713952 | 159 | 173 | - | 9.5643 | 1.06E-05 | 0.4 | GCTACGGTGGCACTT |
| 1 | GMTACGGTCACACTG | chr2R:17747935-17749935 | 168 | 182 | - | 9.5643 | 1.06E-05 | 0.4 | TCAACGGTCACACTA |
| 1 | GMTACGGTCACACTG | chr4:42778-44778 | 169 | 183 | - | 9.5643 | 1.06E-05 | 0.4 | GCTACGCTCACGATG |
| 1 | GMTACGGTCACACTG | chr2L:1707452-1709452 | 171 | 185 | - | 9.5643 | 1.06E-05 | 0.4 | GCTGCGCTGACACTG |
| 1 | GMTACGGTCACACTG | chr3R:12194294-12196294 | 187 | 201 | - | 9.5643 | 1.06E-05 | 0.4 | GATTCGGTCGCAATG |
| 1 | GMTACGGTCACACTG | chr2R:6141444-6143444 | 188 | 202 | - | 9.5643 | 1.06E-05 | 0.4 | GACACAGTCACACAG |
| 1 | GMTACGGTCACACTG | chr2L:9919160-9921160 | 188 | 202 | - | 9.5643 | 1.06E-05 | 0.4 | GAAACGCTCTCACTG |
| 1 | GMTACGGTCACACTG | chr2L:18453587-18455587 | 190 | 204 | - | 9.5643 | 1.06E-05 | 0.4 | GATCCGCTCACACAG |
| 1 | GMTACGGTCACACTG | chr2L:19130954-19132954 | 217 | 231 | - | 9.5643 | 1.06E-05 | 0.4 | GATGCGCTCACCCTG |
| 1 | GMTACGGTCACACTG | chr3L:9455339-9457339 | 223 | 237 | - | 9.5643 | 1.06E-05 | 0.4 | GATACAGGCACACAG |
| 1 | GMTACGGTCACACTG | chr3L:9455318-9457318 | 244 | 258 | - | 9.5643 | 1.06E-05 | 0.4 | GATACAGGCACACAG |
| 1 | GMTACGGTCACACTG | chr3R:29145893-29147893 | 244 | 258 | - | 9.5643 | 1.06E-05 | 0.4 | CATACGGACGCACTG |
| 1 | GMTACGGTCACACTG | chr2R:16282435-16284435 | 253 | 267 | - | 9.5643 | 1.06E-05 | 0.4 | AATACGGTCTCTCTG |
| 1 | GMTACGGTCACACTG | chr2R:10098677-10100677 | 271 | 285 | - | 9.5643 | 1.06E-05 | 0.4 | GAAACGGTTACCCTG |
| 1 | GMTACGGTCACACTG | chr3L:9455281-9457281 | 281 | 295 | - | 9.5643 | 1.06E-05 | 0.4 | GATACAGGCACACAG |
| 1 | GMTACGGTCACACTG | chrX:20093398-20095398 | 286 | 300 | - | 9.5643 | 1.06E-05 | 0.4 | GCCACGCTCACACTT |
| 1 | GMTACGGTCACACTG | chr3L:11609135-11611135 | 306 | 320 | - | 9.5643 | 1.06E-05 | 0.4 | ACTGCGGTAACACTG |
| 1 | GMTACGGTCACACTG | chr3R:21888539-21890539 | 329 | 343 | - | 9.5643 | 1.06E-05 | 0.4 | GATATGGCCGCACTG |
| 1 | GMTACGGTCACACTG | chr2R:19356532-19358532 | 337 | 351 | - | 9.5643 | 1.06E-05 | 0.4 | GCTACGAAAACACTG |
| 1 | GMTACGGTCACACTG | chr3R:10120773-10122773 | 347 | 361 | - | 9.5643 | 1.06E-05 | 0.4 | GAAACGGACAAACTG |
| 1 | GMTACGGTCACACTG | chr2R:10098590-10100590 | 358 | 372 | - | 9.5643 | 1.06E-05 | 0.4 | GAAACGGTTACCCTG |
| 1 | GMTACGGTCACACTG | chr2R:23684543-23686543 | 361 | 375 | - | 9.5643 | 1.06E-05 | 0.4 | GACACGGTCACATCG |
| 1 | GMTACGGTCACACTG | chr3R:24529576-24531576 | 371 | 385 | - | 9.5643 | 1.06E-05 | 0.4 | GATACAGTCTCGCTG |
| 1 | GMTACGGTCACACTG | chr2R:9131480-9133480 | 375 | 389 | - | 9.5643 | 1.06E-05 | 0.4 | CCGACGGTTACACTG |
| 1 | GMTACGGTCACACTG | chr2R:7509357-7511357 | 379 | 393 | - | 9.5643 | 1.06E-05 | 0.4 | CCTACGCTCACACGG |
| 1 | GMTACGGTCACACTG | chr3R:24529558-24531558 | 389 | 403 | - | 9.5643 | 1.06E-05 | 0.4 | GATACAGTCTCGCTG |
| 1 | GMTACGGTCACACTG | chrX:21034469-21036469 | 398 | 412 | - | 9.5643 | 1.06E-05 | 0.4 | GATCTGGTCACACCG |

| | | | | | | | | | |
|---|---|---|---|---|---|---|---|---|---|
| 1 | GMTACGGTCACACTG | chr2R:7698500-7700500 | 399 | 413 | - | 9.5643 | 1.06E-05 | 0.4 | ACTCTGGTCACACTG |
| 1 | GMTACGGTCACACTG | chr3R:9014446-9016446 | 425 | 439 | - | 9.5643 | 1.06E-05 | 0.4 | CAAACGGTCGCACTG |
| 1 | GMTACGGTCACACTG | chr2R:18854087-18856087 | 432 | 446 | - | 9.5643 | 1.06E-05 | 0.4 | GCTACGCACACACTT |
| 1 | GMTACGGTCACACTG | chr2R:22475314-22477314 | 447 | 461 | - | 9.5643 | 1.06E-05 | 0.4 | GCTACTTGCACACTG |
| 1 | GMTACGGTCACACTG | chrX:4314059-4316059 | 449 | 463 | - | 9.5643 | 1.06E-05 | 0.4 | GATGCGCACACACTG |
| 1 | GMTACGGTCACACTG | chr3L:5996208-5998208 | 462 | 476 | - | 9.5643 | 1.06E-05 | 0.4 | GCTAAGGATACACTG |
| 1 | GMTACGGTCACACTG | chr2R:6141165-6143165 | 467 | 481 | - | 9.5643 | 1.06E-05 | 0.4 | GACACAGTCACACAG |
| 1 | GMTACGGTCACACTG | chr3R:12194011-12196011 | 470 | 484 | - | 9.5643 | 1.06E-05 | 0.4 | GATTCGGTCGCAATG |
| 1 | GMTACGGTCACACTG | chr3L:20627498-20629498 | 470 | 484 | - | 9.5643 | 1.06E-05 | 0.4 | GCTACGGATACACTT |
| 1 | GMTACGGTCACACTG | chrX:21318296-21320296 | 473 | 487 | - | 9.5643 | 1.06E-05 | 0.4 | GATACGGACACCGTG |
| 1 | GMTACGGTCACACTG | chrX:21318296-21320296 | 473 | 487 | - | 9.5643 | 1.06E-05 | 0.4 | GATACGGACACCGTG |
| 1 | GMTACGGTCACACTG | chr2R:23063769-23065769 | 483 | 497 | - | 9.5643 | 1.06E-05 | 0.4 | CCTGCGGTCACACTT |
| 1 | GMTACGGTCACACTG | chr3R:30036730-30038730 | 494 | 508 | - | 9.5643 | 1.06E-05 | 0.4 | GCTACGATCAACCTG |
| 1 | GMTACGGTCACACTG | chr2L:7993798-7995798 | 498 | 512 | - | 9.5643 | 1.06E-05 | 0.4 | GCAATGGTAACACTG |
| 1 | GMTACGGTCACACTG | chr2R:20326270-20328270 | 512 | 526 | - | 9.5643 | 1.06E-05 | 0.4 | TCCACGGTCACACTT |
| 1 | GMTACGGTCACACTG | chrX:11704300-11706300 | 530 | 544 | - | 9.5643 | 1.06E-05 | 0.4 | GCCAAGGTCAAACTG |
| 1 | GMTACGGTCACACTG | chr3R:9014341-9016341 | 530 | 544 | - | 9.5643 | 1.06E-05 | 0.4 | CAAACGGTCGCACTG |
| 1 | GMTACGGTCACACTG | chrX:10325464-10327464 | 533 | 547 | - | 9.5643 | 1.06E-05 | 0.4 | GATCCGGTAGCACTG |
| 1 | GMTACGGTCACACTG | chr3R:26005023-26007023 | 552 | 566 | - | 9.5643 | 1.06E-05 | 0.4 | GCTACGTTCATGCTG |
| 1 | GMTACGGTCACACTG | chr2R:7539111-7541111 | 561 | 575 | - | 9.5643 | 1.06E-05 | 0.4 | AAAACGGTCACTCTG |
| 1 | GMTACGGTCACACTG | chr2L:12429892-12431892 | 572 | 586 | - | 9.5643 | 1.06E-05 | 0.4 | GCTACGATCACACCC |
| 1 | GMTACGGTCACACTG | chr2L:16731550-16733550 | 574 | 588 | - | 9.5643 | 1.06E-05 | 0.4 | CATACGGCCACTCTG |
| 1 | GMTACGGTCACACTG | chrX:16942290-16944290 | 575 | 589 | - | 9.5643 | 1.06E-05 | 0.4 | GCTCTGGTAACACTG |
| 1 | GMTACGGTCACACTG | chr3L:4363550-4365550 | 578 | 592 | - | 9.5643 | 1.06E-05 | 0.4 | TCTCTGGTCACACTG |
| 1 | GMTACGGTCACACTG | chr3L:20808669-20810669 | 580 | 594 | - | 9.5643 | 1.06E-05 | 0.4 | GAAACGGTCACGTTG |
| 1 | GMTACGGTCACACTG | chr3R:27915908-27917908 | 592 | 606 | - | 9.5643 | 1.06E-05 | 0.4 | ACTGCGGTCACACTT |
| 1 | GMTACGGTCACACTG | chrX:7706775-7708775 | 596 | 610 | - | 9.5643 | 1.06E-05 | 0.4 | GCTATGGTCACTCTA |
| 1 | GMTACGGTCACACTG | chr3L:14785791-14787791 | 596 | 610 | - | 9.5643 | 1.06E-05 | 0.4 | AATGCGGTCACACCG |
| 1 | GMTACGGTCACACTG | chrX:19637767-19639767 | 597 | 611 | - | 9.5643 | 1.06E-05 | 0.4 | AAAACGGCCACACTG |
| 1 | GMTACGGTCACACTG | chr2L:2237251-2239251 | 601 | 615 | - | 9.5643 | 1.06E-05 | 0.4 | GCTACTGTGACGCTG |
| 1 | GMTACGGTCACACTG | chr3L:10166023-10168023 | 601 | 615 | - | 9.5643 | 1.06E-05 | 0.4 | GATCTGGCCACACTG |
| 1 | GMTACGGTCACACTG | chrY:3041787-3043787 | 619 | 633 | - | 9.5643 | 1.06E-05 | 0.4 | GAGAAGGTCACTCTG |
| 1 | GMTACGGTCACACTG | chr3R:24192832-24194832 | 622 | 636 | - | 9.5643 | 1.06E-05 | 0.4 | GACCTGGTCACACTG |

| | | | | | | | | | |
|---|---|---|---|---|---|---|---|---|---|
| 1 | GMTACGGTCACACTG | chr2R:7698253-7700253 | 646 | 660 | - | 9.5643 | 1.06E-05 | 0.4 | ACTCTGGTCACACTG |
| 1 | GMTACGGTCACACTG | chr3L:15977524-15979524 | 646 | 660 | - | 9.5643 | 1.06E-05 | 0.4 | GCTTCGGTCAGAATG |
| 1 | GMTACGGTCACACTG | chr2R:23842286-23844286 | 652 | 666 | - | 9.5643 | 1.06E-05 | 0.4 | GCTACGTTCACCGTG |
| 1 | GMTACGGTCACACTG | chr2L:6963421-6965421 | 654 | 668 | - | 9.5643 | 1.06E-05 | 0.4 | AATACGGGCACTCTG |
| 1 | GMTACGGTCACACTG | chr3R:24168562-24170562 | 660 | 674 | - | 9.5643 | 1.06E-05 | 0.4 | GATCCTGCCACACTG |
| 1 | GMTACGGTCACACTG | chr2R:6985043-6987043 | 661 | 675 | - | 9.5643 | 1.06E-05 | 0.4 | TCTGCGGTCACACTA |
| 1 | GMTACGGTCACACTG | chr2R:22665605-22667605 | 664 | 678 | - | 9.5643 | 1.06E-05 | 0.4 | GATAAGGCCCCACTG |
| 1 | GMTACGGTCACACTG | chr2R:6195320-6197320 | 665 | 679 | - | 9.5643 | 1.06E-05 | 0.4 | GATACGGTCTCAAAG |
| 1 | GMTACGGTCACACTG | chr2R:23308816-23310816 | 672 | 686 | - | 9.5643 | 1.06E-05 | 0.4 | GATCCTGTCGCACTG |
| 1 | GMTACGGTCACACTG | chr3L:7914437-7916437 | 676 | 690 | - | 9.5643 | 1.06E-05 | 0.4 | CAACCGGTCACACTG |
| 1 | GMTACGGTCACACTG | chr2L:5341234-5343234 | 686 | 700 | - | 9.5643 | 1.06E-05 | 0.4 | GCCAAGGTCACAGTG |
| 1 | GMTACGGTCACACTG | chrY:1121988-1123988 | 693 | 707 | - | 9.5643 | 1.06E-05 | 0.4 | GCTTTGGTCACACTA |
| 1 | GMTACGGTCACACTG | chrX:19682808-19684808 | 698 | 712 | - | 9.5643 | 1.06E-05 | 0.4 | TAAGCGGTCACACTG |
| 1 | GMTACGGTCACACTG | chr2R:8248595-8250595 | 701 | 715 | - | 9.5643 | 1.06E-05 | 0.4 | GCAACGGTAACACTC |
| 1 | GMTACGGTCACACTG | chr3L:3070133-3072133 | 706 | 720 | - | 9.5643 | 1.06E-05 | 0.4 | GATTTGGTCACACCG |
| 1 | GMTACGGTCACACTG | chr3R:9790483-9792483 | 706 | 720 | - | 9.5643 | 1.06E-05 | 0.4 | GACTTGGTCACACTG |
| 1 | GMTACGGTCACACTG | chrX:8164708-8166708 | 710 | 724 | - | 9.5643 | 1.06E-05 | 0.4 | GATGCGGTCATTCTG |
| 1 | GMTACGGTCACACTG | chr3L:9718328-9720328 | 713 | 727 | - | 9.5643 | 1.06E-05 | 0.4 | GATCTGGTCACACCG |
| 1 | GMTACGGTCACACTG | chr3L:12532996-12534996 | 720 | 734 | - | 9.5643 | 1.06E-05 | 0.4 | GCTCTGGTTACACTG |
| 1 | GMTACGGTCACACTG | chr3R:10120400-10122400 | 720 | 734 | - | 9.5643 | 1.06E-05 | 0.4 | GAAACGGACAAACTG |
| 1 | GMTACGGTCACACTG | chr3R:24813262-24815262 | 734 | 748 | - | 9.5643 | 1.06E-05 | 0.4 | GATGCGGTCAATCTG |
| 1 | GMTACGGTCACACTG | chr3R:20056748-20058748 | 734 | 748 | - | 9.5643 | 1.06E-05 | 0.4 | CACGCGGTCACACTG |
| 1 | GMTACGGTCACACTG | chr2R:16281946-16283946 | 742 | 756 | - | 9.5643 | 1.06E-05 | 0.4 | AATACGGTCTCTCTG |
| 1 | GMTACGGTCACACTG | chr3R:23362574-23364574 | 748 | 762 | - | 9.5643 | 1.06E-05 | 0.4 | GCTACGATCACACCC |
| 1 | GMTACGGTCACACTG | chr2R:20643036-20645036 | 748 | 762 | - | 9.5643 | 1.06E-05 | 0.4 | TACGCGGTCACACTG |
| 1 | GMTACGGTCACACTG | chr2L:8996605-8998605 | 750 | 764 | - | 9.5643 | 1.06E-05 | 0.4 | CAAACGGTCACACAG |
| 1 | GMTACGGTCACACTG | chr3R:4588308-4590308 | 751 | 765 | - | 9.5643 | 1.06E-05 | 0.4 | GCTACTGCCACATTG |
| 1 | GMTACGGTCACACTG | chr3R:8823411-8825411 | 757 | 771 | - | 9.5643 | 1.06E-05 | 0.4 | AAAACGGGCACACTG |
| 1 | GMTACGGTCACACTG | chr3R:25261741-25263741 | 760 | 774 | - | 9.5643 | 1.06E-05 | 0.4 | TCAACGGTCACACTT |
| 1 | GMTACGGTCACACTG | chr2R:18403605-18405605 | 762 | 776 | - | 9.5643 | 1.06E-05 | 0.4 | GATACGGTCTAACGG |
| 1 | GMTACGGTCACACTG | chrY:3027019-3029019 | 783 | 797 | - | 9.5643 | 1.06E-05 | 0.4 | GAGAAGGTCACTCTG |
| 1 | GMTACGGTCACACTG | chrY:3034310-3036310 | 783 | 797 | - | 9.5643 | 1.06E-05 | 0.4 | GAGAAGGTCACTCTG |
| 1 | GMTACGGTCACACTG | chr3R:18662738-18664738 | 787 | 801 | - | 9.5643 | 1.06E-05 | 0.4 | ACTTTGGTCACACTG |

| | | | | | | | | | |
|---|---|---|---|---|---|---|---|---|---|
| 1 | GMTACGGTCACACTG | chrX:19694250-19696250 | 791 | 805 | - | 9.5643 | 1.06E-05 | 0.4 | GAGGCAGTCACACTG |
| 1 | GMTACGGTCACACTG | chrX:17860002-17862002 | 795 | 809 | - | 9.5643 | 1.06E-05 | 0.4 | ACCACGGTCACATTG |
| 1 | GMTACGGTCACACTG | chr3L:1722153-1724153 | 795 | 809 | - | 9.5643 | 1.06E-05 | 0.4 | GATCCGGTCATGCTG |
| 1 | GMTACGGTCACACTG | chr2L:9946648-9948648 | 798 | 812 | - | 9.5643 | 1.06E-05 | 0.4 | GCTACCGTCATACAG |
| 1 | GMTACGGTCACACTG | chr3R:30176877-30178877 | 817 | 831 | - | 9.5643 | 1.06E-05 | 0.4 | ACTCAGGTCACACTG |
| 1 | GMTACGGTCACACTG | chr2R:9131030-9133030 | 825 | 839 | - | 9.5643 | 1.06E-05 | 0.4 | CCGACGGTTACACTG |
| 1 | GMTACGGTCACACTG | chr2R:24598947-24600947 | 835 | 849 | - | 9.5643 | 1.06E-05 | 0.4 | GCTTTGGTCATACTG |
| 1 | GMTACGGTCACACTG | chr2R:12759999-12761999 | 837 | 851 | - | 9.5643 | 1.06E-05 | 0.4 | GCGACAGTCACACTC |
| 1 | GMTACGGTCACACTG | chr2R:8630962-8632962 | 837 | 851 | - | 9.5643 | 1.06E-05 | 0.4 | ACTCTGGTCACACTG |
| 1 | GMTACGGTCACACTG | chr2R:6882280-6884280 | 840 | 854 | - | 9.5643 | 1.06E-05 | 0.4 | TATTCGGTCTCACTG |
| 1 | GMTACGGTCACACTG | chr2R:18631045-18633045 | 848 | 862 | - | 9.5643 | 1.06E-05 | 0.4 | CAAACGGGCACACTG |
| 1 | GMTACGGTCACACTG | chrX:9157622-9159622 | 873 | 887 | - | 9.5643 | 1.06E-05 | 0.4 | TATTCGGCCACACTG |
| 1 | GMTACGGTCACACTG | chrX:6523464-6525464 | 876 | 890 | - | 9.5643 | 1.06E-05 | 0.4 | GAGCTGGTCACACTG |
| 1 | GMTACGGTCACACTG | chrX:8573306-8575306 | 884 | 898 | - | 9.5643 | 1.06E-05 | 0.4 | GCTCCGGTTACACTA |
| 1 | GMTACGGTCACACTG | chr3L:9718144-9720144 | 897 | 911 | - | 9.5643 | 1.06E-05 | 0.4 | GATCTGGTCACACCG |
| 1 | GMTACGGTCACACTG | chr2R:8950595-8952595 | 898 | 912 | - | 9.5643 | 1.06E-05 | 0.4 | AAAACGGTCACACCG |
| 1 | GMTACGGTCACACTG | chr2R:13861002-13863002 | 903 | 917 | - | 9.5643 | 1.06E-05 | 0.4 | GCGACGGTCATACTA |
| 1 | GMTACGGTCACACTG | chr3R:20592139-20594139 | 905 | 919 | - | 9.5643 | 1.06E-05 | 0.4 | GCTGAGGTCACACTA |
| 1 | GMTACGGTCACACTG | chr3R:11740890-11742890 | 906 | 920 | - | 9.5643 | 1.06E-05 | 0.4 | CCGACGGTCACACTA |
| 1 | GMTACGGTCACACTG | chr2R:13860997-13862997 | 908 | 922 | - | 9.5643 | 1.06E-05 | 0.4 | GCGACGGTCATACTA |
| 1 | GMTACGGTCACACTG | chr2R:22665359-22667359 | 910 | 924 | - | 9.5643 | 1.06E-05 | 0.4 | GATAAGGCCCCACTG |
| 1 | GMTACGGTCACACTG | chr3L:15831464-15833464 | 914 | 928 | - | 9.5643 | 1.06E-05 | 0.4 | GAAGCGTTCACACTG |
| 1 | GMTACGGTCACACTG | chr2L:10269051-10271051 | 916 | 930 | - | 9.5643 | 1.06E-05 | 0.4 | GAAACGGGAACACTG |
| 1 | GMTACGGTCACACTG | chrX:6523464-6525464 | 917 | 931 | - | 9.5643 | 1.06E-05 | 0.4 | GCTGGGGTGACACTG |
| 1 | GMTACGGTCACACTG | chr3R:24332433-24334433 | 933 | 947 | - | 9.5643 | 1.06E-05 | 0.4 | GCAATGGTCACAGTG |
| 1 | GMTACGGTCACACTG | chr3L:21538862-21540862 | 948 | 962 | - | 9.5643 | 1.06E-05 | 0.4 | GAAATGGTCACCCTG |
| 1 | GMTACGGTCACACTG | chr3L:16377566-16379566 | 957 | 971 | - | 9.5643 | 1.06E-05 | 0.4 | CCTGCGGTCATACTG |
| 1 | GMTACGGTCACACTG | chr3L:1884657-1886657 | 961 | 975 | - | 9.5643 | 1.06E-05 | 0.4 | GATACAGACACACAG |
| 1 | GMTACGGTCACACTG | chr2R:24598821-24600821 | 961 | 975 | - | 9.5643 | 1.06E-05 | 0.4 | GCTTTGGTCATACTG |
| 1 | GMTACGGTCACACTG | chr3R:9013909-9015909 | 962 | 976 | - | 9.5643 | 1.06E-05 | 0.4 | CAAACGGTCGCACTG |
| 1 | GMTACGGTCACACTG | chr3R:15485050-15487050 | 963 | 977 | - | 9.5643 | 1.06E-05 | 0.4 | AAAGCGGTCACACTG |
| 1 | GMTACGGTCACACTG | chr3R:27953112-27955112 | 964 | 978 | - | 9.5643 | 1.06E-05 | 0.4 | GCTACGGTCACATCA |
| 1 | GMTACGGTCACACTG | chr3R:21757764-21759764 | 965 | 979 | - | 9.5643 | 1.06E-05 | 0.4 | GATTCGGACACACTC |

| | | | | | | | | | |
|---|---|---|---|---|---|---|---|---|---|
| 1 | GMTACGGTCACACTG | chr2R:12183310-12185310 | 966 | 980 | - | 9.5643 | 1.06E-05 | 0.4 | GCTGTGGTCACACTA |
| 1 | GMTACGGTCACACTG | chr3R:9790221-9792221 | 968 | 982 | - | 9.5643 | 1.06E-05 | 0.4 | GACTTGGTCACACTG |
| 1 | GMTACGGTCACACTG | chr2L:18681075-18683075 | 971 | 985 | - | 9.5643 | 1.06E-05 | 0.4 | ACTTTGGTCACACTG |
| 1 | GMTACGGTCACACTG | chr3R:17808222-17810222 | 971 | 985 | - | 9.5643 | 1.06E-05 | 0.4 | ACAACGGTCACACTT |
| 1 | GMTACGGTCACACTG | chr3L:17241128-17243128 | 972 | 986 | - | 9.5643 | 1.06E-05 | 0.4 | GACGCGGTCACACTT |
| 1 | GMTACGGTCACACTG | chr2R:15939073-15941073 | 972 | 986 | - | 9.5643 | 1.06E-05 | 0.4 | GATTCAGCCACACTG |
| 1 | GMTACGGTCACACTG | chr3R:18662552-18664552 | 973 | 987 | - | 9.5643 | 1.06E-05 | 0.4 | ACTTTGGTCACACTG |
| 1 | GMTACGGTCACACTG | chr2L:14354145-14356145 | 973 | 987 | - | 9.5643 | 1.06E-05 | 0.4 | CCAAGGGTCACACTG |
| 1 | GMTACGGTCACACTG | chr2L:11129737-11131737 | 974 | 988 | - | 9.5643 | 1.06E-05 | 0.4 | CCCACGGTCACACTT |
| 1 | GMTACGGTCACACTG | chr3L:14785412-14787412 | 975 | 989 | - | 9.5643 | 1.06E-05 | 0.4 | AATGCGGTCACACCG |
| 1 | GMTACGGTCACACTG | chr3L:16350991-16352991 | 975 | 989 | - | 9.5643 | 1.06E-05 | 0.4 | GCAATGGTCATACTG |
| 1 | GMTACGGTCACACTG | chr2L:8996379-8998379 | 976 | 990 | - | 9.5643 | 1.06E-05 | 0.4 | CAAACGGTCACACAG |
| 1 | GMTACGGTCACACTG | chr3L:11608465-11610465 | 976 | 990 | - | 9.5643 | 1.06E-05 | 0.4 | ACTGCGGTAACACTG |
| 1 | GMTACGGTCACACTG | chr2L:9575769-9577769 | 976 | 990 | - | 9.5643 | 1.06E-05 | 0.4 | AACGCGGTCACACTG |
| 1 | GMTACGGTCACACTG | chr2R:20325806-20327806 | 976 | 990 | - | 9.5643 | 1.06E-05 | 0.4 | TCCACGGTCACACTT |
| 1 | GMTACGGTCACACTG | chr2R:7538696-7540696 | 976 | 990 | - | 9.5643 | 1.06E-05 | 0.4 | AAAACGGTCACTCTG |
| 1 | GMTACGGTCACACTG | chr3L:5764846-5766846 | 977 | 991 | - | 9.5643 | 1.06E-05 | 0.4 | CAAGCGGTCACACTG |
| 1 | GMTACGGTCACACTG | chr3L:20808272-20810272 | 977 | 991 | - | 9.5643 | 1.06E-05 | 0.4 | GAAACGGTCACGTTG |
| 1 | GMTACGGTCACACTG | chr2L:1709462-1711462 | 979 | 993 | - | 9.5643 | 1.06E-05 | 0.4 | TCCATGGTCACACTG |
| 1 | GMTACGGTCACACTG | chr3R:23125422-23127422 | 980 | 994 | - | 9.5643 | 1.06E-05 | 0.4 | TCTGCGGTCACATTG |
| 1 | GMTACGGTCACACTG | chrX:9157515-9159515 | 980 | 994 | - | 9.5643 | 1.06E-05 | 0.4 | TATTCGGCCACACTG |
| 1 | GMTACGGTCACACTG | chr3L:11707979-11709979 | 983 | 997 | - | 9.5643 | 1.06E-05 | 0.4 | CAAGCGGTCACACTG |
| 1 | GMTACGGTCACACTG | chr3L:11707979-11709979 | 983 | 997 | - | 9.5643 | 1.06E-05 | 0.4 | CAAGCGGTCACACTG |
| 1 | GMTACGGTCACACTG | chr3R:27915516-27917516 | 984 | 998 | - | 9.5643 | 1.06E-05 | 0.4 | ACTGCGGTCACACTT |
| 1 | GMTACGGTCACACTG | chr3L:1730107-1732107 | 984 | 998 | - | 9.5643 | 1.06E-05 | 0.4 | AAAGCGGTCACACTG |
| 1 | GMTACGGTCACACTG | chr3R:24813012-24815012 | 984 | 998 | - | 9.5643 | 1.06E-05 | 0.4 | GATGCGGTCAATCTG |
| 1 | GMTACGGTCACACTG | chr3L:8410851-8412851 | 986 | 1000 | - | 9.5643 | 1.06E-05 | 0.4 | GCTTTGGTCATACTG |
| 1 | GMTACGGTCACACTG | chr3L:8513921-8515921 | 986 | 1000 | - | 9.5643 | 1.06E-05 | 0.4 | GCAACGGTCATACTT |
| 1 | GMTACGGTCACACTG | chr2R:9881357-9883357 | 987 | 1001 | - | 9.5643 | 1.06E-05 | 0.4 | CCAACGGTCACACTA |
| 1 | GMTACGGTCACACTG | chrX:19682518-19684518 | 988 | 1002 | - | 9.5643 | 1.06E-05 | 0.4 | TAAGCGGTCACACTG |
| 1 | GMTACGGTCACACTG | chr3L:15013062-15015062 | 989 | 1003 | - | 9.5643 | 1.06E-05 | 0.4 | TCTGCGCTCACACTG |
| 1 | GMTACGGTCACACTG | chr3R:8722174-8724174 | 990 | 1004 | - | 9.5643 | 1.06E-05 | 0.4 | GATGCGCACACACTG |
| 1 | GMTACGGTCACACTG | chr2L:16731133-16733133 | 991 | 1005 | - | 9.5643 | 1.06E-05 | 0.4 | CATACGGCCACTCTG |

| | | | | | | | | | |
|---|---|---|---|---|---|---|---|---|---|
| 1 | GMTACGGTCACACTG | chr3R:8823177-8825177 | 991 | 1005 | - | 9.5643 | 1.06E-05 | 0.4 | AAAACGGGCACACTG |
| 1 | GMTACGGTCACACTG | chr2L:20925147-20927147 | 992 | 1006 | - | 9.5643 | 1.06E-05 | 0.4 | CCTACGGCCACACTA |
| 1 | GMTACGGTCACACTG | chr3R:24356938-24358938 | 992 | 1006 | - | 9.5643 | 1.06E-05 | 0.4 | GATATGGGCACACAG |
| 1 | GMTACGGTCACACTG | chr2R:23063260-23065260 | 992 | 1006 | - | 9.5643 | 1.06E-05 | 0.4 | CCTGCGGTCACACTT |
| 1 | GMTACGGTCACACTG | chr2R:6984712-6986712 | 992 | 1006 | - | 9.5643 | 1.06E-05 | 0.4 | TCTGCGGTCACACTA |
| 1 | GMTACGGTCACACTG | chr2R:6984712-6986712 | 992 | 1006 | - | 9.5643 | 1.06E-05 | 0.4 | TCTGCGGTCACACTA |
| 1 | GMTACGGTCACACTG | chr3L:12532724-12534724 | 992 | 1006 | - | 9.5643 | 1.06E-05 | 0.4 | GCTCTGGTTACACTG |
| 1 | GMTACGGTCACACTG | chr2R:4470060-4472060 | 993 | 1007 | - | 9.5643 | 1.06E-05 | 0.4 | TCCACGGTCACACTA |
| 1 | GMTACGGTCACACTG | chr3L:16609418-16611418 | 993 | 1007 | - | 9.5643 | 1.06E-05 | 0.4 | ACAACGGTCACATTG |
| 1 | GMTACGGTCACACTG | chr3L:7914120-7916120 | 993 | 1007 | - | 9.5643 | 1.06E-05 | 0.4 | CAACCGGTCACACTG |
| 1 | GMTACGGTCACACTG | chrX:16941871-16943871 | 994 | 1008 | - | 9.5643 | 1.06E-05 | 0.4 | GCTCTGGTAACACTG |
| 1 | GMTACGGTCACACTG | chr3R:30176700-30178700 | 994 | 1008 | - | 9.5643 | 1.06E-05 | 0.4 | ACTCAGGTCACACTG |
| 1 | GMTACGGTCACACTG | chrX:19637369-19639369 | 995 | 1009 | - | 9.5643 | 1.06E-05 | 0.4 | AAAACGGCCACACTG |
| 1 | GMTACGGTCACACTG | chrX:7097053-7099053 | 995 | 1009 | - | 9.5643 | 1.06E-05 | 0.4 | GAAACGTGCACACTG |
| 1 | GMTACGGTCACACTG | chr3L:4135038-4137038 | 996 | 1010 | - | 9.5643 | 1.06E-05 | 0.4 | GAAACGGCCACACTA |
| 1 | GMTACGGTCACACTG | chr2R:12199815-12201815 | 997 | 1011 | - | 9.5643 | 1.06E-05 | 0.4 | CAAACGGTCACTCTG |
| 1 | GMTACGGTCACACTG | chrX:17859800-17861800 | 997 | 1011 | - | 9.5643 | 1.06E-05 | 0.4 | ACCACGGTCACATTG |
| 1 | GMTACGGTCACACTG | chr2R:7697902-7699902 | 997 | 1011 | - | 9.5643 | 1.06E-05 | 0.4 | ACTCTGGTCACACTG |
| 1 | GMTACGGTCACACTG | chr2R:20642785-20644785 | 999 | 1013 | - | 9.5643 | 1.06E-05 | 0.4 | TACGCGGTCACACTG |
| 1 | GMTACGGTCACACTG | chr2R:20642785-20644785 | 999 | 1013 | - | 9.5643 | 1.06E-05 | 0.4 | TACGCGGTCACACTG |
| 1 | GMTACGGTCACACTG | chr3L:3069839-3071839 | 1000 | 1014 | - | 9.5643 | 1.06E-05 | 0.4 | GATTTGGTCACACCG |
| 1 | GMTACGGTCACACTG | chr2R:12150057-12152057 | 1000 | 1014 | - | 9.5643 | 1.06E-05 | 0.4 | CCTACGGTCACACCA |
| 1 | GMTACGGTCACACTG | chr2L:6963074-6965074 | 1001 | 1015 | - | 9.5643 | 1.06E-05 | 0.4 | AATACGGGCACTCTG |
| 1 | GMTACGGTCACACTG | chrX:8141279-8143279 | 1001 | 1015 | - | 9.5643 | 1.06E-05 | 0.4 | CCAACGGTCACACTA |
| 1 | GMTACGGTCACACTG | chr3R:25261498-25263498 | 1003 | 1017 | - | 9.5643 | 1.06E-05 | 0.4 | TCAACGGTCACACTT |
| 1 | GMTACGGTCACACTG | chr2R:24602572-24604572 | 1004 | 1018 | - | 9.5643 | 1.06E-05 | 0.4 | AATCCGGACACACTG |
| 1 | GMTACGGTCACACTG | chr3R:27958245-27960245 | 1005 | 1019 | - | 9.5643 | 1.06E-05 | 0.4 | AACGCGGTCACACTG |
| 1 | GMTACGGTCACACTG | chr2L:10974293-10976293 | 1005 | 1019 | - | 9.5643 | 1.06E-05 | 0.4 | ACTGCGGTCACACTT |
| 1 | GMTACGGTCACACTG | chr3R:27958245-27960245 | 1005 | 1019 | - | 9.5643 | 1.06E-05 | 0.4 | AACGCGGTCACACTG |
| 1 | GMTACGGTCACACTG | chr3R:17020120-17022120 | 1006 | 1020 | - | 9.5643 | 1.06E-05 | 0.4 | AAGGCGGTCACACTG |
| 1 | GMTACGGTCACACTG | chr2L:15112480-15114480 | 1008 | 1022 | - | 9.5643 | 1.06E-05 | 0.4 | GAAACGGTACCACTG |
| 1 | GMTACGGTCACACTG | chr2L:11112061-11114061 | 1010 | 1024 | - | 9.5643 | 1.06E-05 | 0.4 | GATACAGACACACCG |
| 1 | GMTACGGTCACACTG | chrX:1456054-1458054 | 1011 | 1025 | - | 9.5643 | 1.06E-05 | 0.4 | GAAACGGTCACAAAG |

| | | | | | | | | | |
|---|---|---|---|---|---|---|---|---|---|
| 1 | GMTACGGTCACACTG | chr3R:11740783-11742783 | 1013 | 1027 | - | 9.5643 | 1.06E-05 | 0.4 | CCGACGGTCACACTA |
| 1 | GMTACGGTCACACTG | chr3L:16350951-16352951 | 1015 | 1029 | - | 9.5643 | 1.06E-05 | 0.4 | GCAATGGTCATACTG |
| 1 | GMTACGGTCACACTG | chr3R:5216456-5218456 | 1016 | 1030 | - | 9.5643 | 1.06E-05 | 0.4 | TCTGCGGTCATACTG |
| 1 | GMTACGGTCACACTG | chrX:6523323-6525323 | 1017 | 1031 | - | 9.5643 | 1.06E-05 | 0.4 | GAGCTGGTCACACTG |
| 1 | GMTACGGTCACACTG | chr3L:1730107-1732107 | 1019 | 1033 | - | 9.5643 | 1.06E-05 | 0.4 | GCTAGGGCCACACTA |
| 1 | GMTACGGTCACACTG | chr2L:1709421-1711421 | 1020 | 1034 | - | 9.5643 | 1.06E-05 | 0.4 | TCCATGGTCACACTG |
| 1 | GMTACGGTCACACTG | chr2R:17747078-17749078 | 1025 | 1039 | - | 9.5643 | 1.06E-05 | 0.4 | TCAACGGTCACACTA |
| 1 | GMTACGGTCACACTG | chr3L:4363102-4365102 | 1026 | 1040 | - | 9.5643 | 1.06E-05 | 0.4 | TCTCTGGTCACACTG |
| 1 | GMTACGGTCACACTG | chrX:12896327-12898327 | 1029 | 1043 | - | 9.5643 | 1.06E-05 | 0.4 | TAGCCGGTCACACTG |
| 1 | GMTACGGTCACACTG | chr3R:24192423-24194423 | 1031 | 1045 | - | 9.5643 | 1.06E-05 | 0.4 | GACCTGGTCACACTG |
| 1 | GMTACGGTCACACTG | chr2L:14354085-14356085 | 1033 | 1047 | - | 9.5643 | 1.06E-05 | 0.4 | CCAAGGGTCACACTG |
| 1 | GMTACGGTCACACTG | chr3R:31680019-31682019 | 1033 | 1047 | - | 9.5643 | 1.06E-05 | 0.4 | GCCACAATCACACTG |
| 1 | GMTACGGTCACACTG | chrX:10324958-10326958 | 1039 | 1053 | - | 9.5643 | 1.06E-05 | 0.4 | GATCCGGTAGCACTG |
| 1 | GMTACGGTCACACTG | chr3L:21705365-21707365 | 1045 | 1059 | - | 9.5643 | 1.06E-05 | 0.4 | GCTGCCGTCAAACTG |
| 1 | GMTACGGTCACACTG | chrX:17300929-17302929 | 1046 | 1060 | - | 9.5643 | 1.06E-05 | 0.4 | AAAACGGGCACACTG |
| 1 | GMTACGGTCACACTG | chr3R:24332316-24334316 | 1050 | 1064 | - | 9.5643 | 1.06E-05 | 0.4 | GCAATGGTCACAGTG |
| 1 | GMTACGGTCACACTG | chrX:6523323-6525323 | 1058 | 1072 | - | 9.5643 | 1.06E-05 | 0.4 | GCTGGGGTGACACTG |
| 1 | GMTACGGTCACACTG | chr2L:9699810-9701810 | 1062 | 1076 | - | 9.5643 | 1.06E-05 | 0.4 | GCTAAGTTCACACCG |
| 1 | GMTACGGTCACACTG | chr2L:11129645-11131645 | 1066 | 1080 | - | 9.5643 | 1.06E-05 | 0.4 | CCCACGGTCACACTT |
| 1 | GMTACGGTCACACTG | chr3R:27960821-27962821 | 1075 | 1089 | - | 9.5643 | 1.06E-05 | 0.4 | AACGCGGTCACACTG |
| 1 | GMTACGGTCACACTG | chr3L:12416580-12418580 | 1080 | 1094 | - | 9.5643 | 1.06E-05 | 0.4 | ACTTTGGTCACACTG |
| 1 | GMTACGGTCACACTG | chr2L:7993184-7995184 | 1112 | 1126 | - | 9.5643 | 1.06E-05 | 0.4 | GCAATGGTAACACTG |
| 1 | GMTACGGTCACACTG | chr3L:5764702-5766702 | 1121 | 1135 | - | 9.5643 | 1.06E-05 | 0.4 | CAAGCGGTCACACTG |
| 1 | GMTACGGTCACACTG | chr3R:24270233-24272233 | 1141 | 1155 | - | 9.5643 | 1.06E-05 | 0.4 | TCTCTGGTCACACTG |
| 1 | GMTACGGTCACACTG | chr2R:23358215-23360215 | 1141 | 1155 | - | 9.5643 | 1.06E-05 | 0.4 | GCTGCGCTTACACTG |
| 1 | GMTACGGTCACACTG | chr3R:22013697-22015697 | 1149 | 1163 | - | 9.5643 | 1.06E-05 | 0.4 | GCTGAGGTCACACTT |
| 1 | GMTACGGTCACACTG | chrX:23010303-23012303 | 1153 | 1167 | - | 9.5643 | 1.06E-05 | 0.4 | GCTACGCCCACACTT |
| 1 | GMTACGGTCACACTG | chr2R:12651805-12653805 | 1159 | 1173 | - | 9.5643 | 1.06E-05 | 0.4 | GATAGGGTCGCTCTG |
| 1 | GMTACGGTCACACTG | chr3R:16213278-16215278 | 1160 | 1174 | - | 9.5643 | 1.06E-05 | 0.4 | GCCACGTTCAGACTG |
| 1 | GMTACGGTCACACTG | chrX:1050525-1052525 | 1173 | 1187 | - | 9.5643 | 1.06E-05 | 0.4 | GCCAAGGTCACATTG |
| 1 | GMTACGGTCACACTG | chr4:41774-43774 | 1173 | 1187 | - | 9.5643 | 1.06E-05 | 0.4 | GCTACGCTCACGATG |
| 1 | GMTACGGTCACACTG | chr2L:16793211-16795211 | 1174 | 1188 | - | 9.5643 | 1.06E-05 | 0.4 | ACCACGGTCACATTG |
| 1 | GMTACGGTCACACTG | chr2R:22033844-22035844 | 1190 | 1204 | - | 9.5643 | 1.06E-05 | 0.4 | GACACGGTCACCTTG |

| | | | | | | | | | |
|---|---|---|---|---|---|---|---|---|---|
| 1 | GMTACGGTCACACTG | chr3R:27606879-27608879 | 1228 | 1242 | - | 9.5643 | 1.06E-05 | 0.4 | GCAACTGTCAAACTG |
| 1 | GMTACGGTCACACTG | chr3R:30806117-30808117 | 1233 | 1247 | - | 9.5643 | 1.06E-05 | 0.4 | GCTACGGTGACCTTG |
| 1 | GMTACGGTCACACTG | chr2R:9711321-9713321 | 1235 | 1249 | - | 9.5643 | 1.06E-05 | 0.4 | GATTCGAACACACTG |
| 1 | GMTACGGTCACACTG | chrX:16082232-16084232 | 1236 | 1250 | - | 9.5643 | 1.06E-05 | 0.4 | GCTACGGTCAACATG |
| 1 | GMTACGGTCACACTG | chr2L:6708086-6710086 | 1245 | 1259 | - | 9.5643 | 1.06E-05 | 0.4 | GATCTGGCCACACTG |
| 1 | GMTACGGTCACACTG | chr2L:13020312-13022312 | 1266 | 1280 | - | 9.5643 | 1.06E-05 | 0.4 | GATACGGCAACTCTG |
| 1 | GMTACGGTCACACTG | chr3R:26324294-26326294 | 1286 | 1300 | - | 9.5643 | 1.06E-05 | 0.4 | GCTAAGGTCGCACTT |
| 1 | GMTACGGTCACACTG | chr2L:727362-729362 | 1307 | 1321 | - | 9.5643 | 1.06E-05 | 0.4 | TCCACGATCACACTG |
| 1 | GMTACGGTCACACTG | chr3L:16555823-16557823 | 1307 | 1321 | - | 9.5643 | 1.06E-05 | 0.4 | CCTAGGGCCACACTG |
| 1 | GMTACGGTCACACTG | chr3R:27339622-27341622 | 1313 | 1327 | - | 9.5643 | 1.06E-05 | 0.4 | GCGACTGTGACACTG |
| 1 | GMTACGGTCACACTG | chr3R:25261182-25263182 | 1319 | 1333 | - | 9.5643 | 1.06E-05 | 0.4 | TCAACGGTCACACTT |
| 1 | GMTACGGTCACACTG | chr2R:11372607-11374607 | 1322 | 1336 | - | 9.5643 | 1.06E-05 | 0.4 | GCAACGGTGACAATG |
| 1 | GMTACGGTCACACTG | chr2R:16867222-16869222 | 1332 | 1346 | - | 9.5643 | 1.06E-05 | 0.4 | GCCACGTTCACATTG |
| 1 | GMTACGGTCACACTG | chr2R:19355525-19357525 | 1344 | 1358 | - | 9.5643 | 1.06E-05 | 0.4 | GCTACGAAAACACTG |
| 1 | GMTACGGTCACACTG | chr3R:30035870-30037870 | 1354 | 1368 | - | 9.5643 | 1.06E-05 | 0.4 | GCTACGATCAACCTG |
| 1 | GMTACGGTCACACTG | chr3L:21538447-21540447 | 1363 | 1377 | - | 9.5643 | 1.06E-05 | 0.4 | GAAATGGTCACCCTG |
| 1 | GMTACGGTCACACTG | chr3R:20591671-20593671 | 1373 | 1387 | - | 9.5643 | 1.06E-05 | 0.4 | GCTGAGGTCACACTA |
| 1 | GMTACGGTCACACTG | chr3L:691919-693919 | 1376 | 1390 | - | 9.5643 | 1.06E-05 | 0.4 | GACAAGGCCACACTG |
| 1 | GMTACGGTCACACTG | chr3L:1476619-1478619 | 1395 | 1409 | - | 9.5643 | 1.06E-05 | 0.4 | ACCACGGTCACAGTG |
| 1 | GMTACGGTCACACTG | chr2R:23357961-23359961 | 1395 | 1409 | - | 9.5643 | 1.06E-05 | 0.4 | GCTGCGCTTACACTG |
| 1 | GMTACGGTCACACTG | chr3L:211709-213709 | 1396 | 1410 | - | 9.5643 | 1.06E-05 | 0.4 | GATACTGGCACACCG |
| 1 | GMTACGGTCACACTG | chr2R:18877391-18879391 | 1402 | 1416 | - | 9.5643 | 1.06E-05 | 0.4 | CATTCGGCCACACTG |
| 1 | GMTACGGTCACACTG | chr2R:8950086-8952086 | 1407 | 1421 | - | 9.5643 | 1.06E-05 | 0.4 | AAAACGGTCACACCG |
| 1 | GMTACGGTCACACTG | chr2R:8950086-8952086 | 1407 | 1421 | - | 9.5643 | 1.06E-05 | 0.4 | AAAACGGTCACACCG |
| 1 | GMTACGGTCACACTG | chr3L:11707553-11709553 | 1409 | 1423 | - | 9.5643 | 1.06E-05 | 0.4 | CAAGCGGTCACACTG |
| 1 | GMTACGGTCACACTG | chr3L:211687-213687 | 1418 | 1432 | - | 9.5643 | 1.06E-05 | 0.4 | GATACTGGCACACCG |
| 1 | GMTACGGTCACACTG | chr2L:13285777-13287777 | 1442 | 1456 | - | 9.5643 | 1.06E-05 | 0.4 | GACACAGACACACTG |
| 1 | GMTACGGTCACACTG | chr2R:16229680-16231680 | 1460 | 1474 | - | 9.5643 | 1.06E-05 | 0.4 | GCGACCGTAACACTG |
| 1 | GMTACGGTCACACTG | chr3R:9354495-9356495 | 1475 | 1489 | - | 9.5643 | 1.06E-05 | 0.4 | GCTACTGCCAAACTG |
| 1 | GMTACGGTCACACTG | chr2R:12651487-12653487 | 1477 | 1491 | - | 9.5643 | 1.06E-05 | 0.4 | GATAGGGTCGCTCTG |
| 1 | GMTACGGTCACACTG | chrX:23009972-23011972 | 1484 | 1498 | - | 9.5643 | 1.06E-05 | 0.4 | GCTACGCCCACACTT |
| 1 | GMTACGGTCACACTG | chr2R:12228310-12230310 | 1489 | 1503 | - | 9.5643 | 1.06E-05 | 0.4 | GCTATGGTCACTATG |
| 1 | GMTACGGTCACACTG | chr3L:8197207-8199207 | 1508 | 1522 | - | 9.5643 | 1.06E-05 | 0.4 | GCTGTGGTCACAATG |

| | | | | | | | | | |
|---|---|---|---|---|---|---|---|---|---|
| 1 | GMTACGGTCACACTG | chr2L:6565181-6567181 | 1533 | 1547 | - | 9.5643 | 1.06E-05 | 0.4 | GCAACAGTCAGACTG |
| 1 | GMTACGGTCACACTG | chr3R:29199454-29201454 | 1589 | 1603 | - | 9.5643 | 1.06E-05 | 0.4 | TCTACGGTCTCATTG |
| 1 | GMTACGGTCACACTG | chr2L:10676198-10678198 | 1591 | 1605 | - | 9.5643 | 1.06E-05 | 0.4 | GCTACGTTCACTTTG |
| 1 | GMTACGGTCACACTG | chrX:8572593-8574593 | 1597 | 1611 | - | 9.5643 | 1.06E-05 | 0.4 | GCTCCGGTTACACTA |
| 1 | GMTACGGTCACACTG | chr3L:1884019-1886019 | 1599 | 1613 | - | 9.5643 | 1.06E-05 | 0.4 | GATACAGACACACAG |
| 1 | GMTACGGTCACACTG | chr3R:15484396-15486396 | 1617 | 1631 | - | 9.5643 | 1.06E-05 | 0.4 | AAAGCGGTCACACTG |
| 1 | GMTACGGTCACACTG | chr2R:22664646-22666646 | 1623 | 1637 | - | 9.5643 | 1.06E-05 | 0.4 | GATAAGGCCCCACTG |
| 1 | GMTACGGTCACACTG | chrX:3239571-3241571 | 1624 | 1638 | - | 9.5643 | 1.06E-05 | 0.4 | GCTACGTACACAATG |
| 1 | GMTACGGTCACACTG | chr2L:20530066-20532066 | 1654 | 1668 | - | 9.5643 | 1.06E-05 | 0.4 | ACTACGATCACGCTG |
| 1 | GMTACGGTCACACTG | chr3R:9354313-9356313 | 1657 | 1671 | - | 9.5643 | 1.06E-05 | 0.4 | GCTACTGCCAAACTG |
| 1 | GMTACGGTCACACTG | chr2R:25249316-25251316 | 1658 | 1672 | - | 9.5643 | 1.06E-05 | 0.4 | GCCACGCTCACACTT |
| 1 | GMTACGGTCACACTG | chr3L:16688268-16690268 | 1676 | 1690 | - | 9.5643 | 1.06E-05 | 0.4 | GCTACGTACATACTG |
| 1 | GMTACGGTCACACTG | chr3R:27606396-27608396 | 1711 | 1725 | - | 9.5643 | 1.06E-05 | 0.4 | GCAACTGTCAAACTG |
| 1 | GMTACGGTCACACTG | chr3L:8196987-8198987 | 1728 | 1742 | - | 9.5643 | 1.06E-05 | 0.4 | GCTGTGGTCACAATG |
| 1 | GMTACGGTCACACTG | chr2L:9917618-9919618 | 1730 | 1744 | - | 9.5643 | 1.06E-05 | 0.4 | GAAACGCTCTCACTG |
| 1 | GMTACGGTCACACTG | chr3R:30805619-30807619 | 1731 | 1745 | - | 9.5643 | 1.06E-05 | 0.4 | GCTACGGTGACCTTG |
| 1 | GMTACGGTCACACTG | chr3L:16128617-16130617 | 1733 | 1747 | - | 9.5643 | 1.06E-05 | 0.4 | GCTTCTGTAACACTG |
| 1 | GMTACGGTCACACTG | chrX:2875533-2877533 | 1742 | 1756 | - | 9.5643 | 1.06E-05 | 0.4 | GAGACGGACAAACTG |
| 1 | GMTACGGTCACACTG | chr2R:23357612-23359612 | 1744 | 1758 | - | 9.5643 | 1.06E-05 | 0.4 | GCTGCGCTTACACTG |
| 1 | GMTACGGTCACACTG | chr2L:13285462-13287462 | 1757 | 1771 | - | 9.5643 | 1.06E-05 | 0.4 | GACACAGACACACTG |
| 1 | GMTACGGTCACACTG | chr2R:6970637-6972637 | 1766 | 1780 | - | 9.5643 | 1.06E-05 | 0.4 | GATACGCTCCCTCTG |
| 1 | GMTACGGTCACACTG | chr2R:6970637-6972637 | 1766 | 1780 | - | 9.5643 | 1.06E-05 | 0.4 | GATACGCTCCCTCTG |
| 1 | GMTACGGTCACACTG | chr3L:5764046-5766046 | 1777 | 1791 | - | 9.5643 | 1.06E-05 | 0.4 | CAAGCGGTCACACTG |
| 1 | GMTACGGTCACACTG | chr2L:10091389-10093389 | 1794 | 1808 | - | 9.5643 | 1.06E-05 | 0.4 | ACTGCAGTCACACTG |
| 1 | GMTACGGTCACACTG | chr3R:5550168-5552168 | 1809 | 1823 | - | 9.5643 | 1.06E-05 | 0.4 | GCTACGGCCACATTC |
| 1 | GMTACGGTCACACTG | chr3R:26184251-26186251 | 1809 | 1823 | - | 9.5643 | 1.06E-05 | 0.4 | CCCACGGTCAAACTG |
| 1 | GMTACGGTCACACTG | chr3L:9403797-9405797 | 1810 | 1824 | - | 9.5643 | 1.06E-05 | 0.4 | GATCCGGTCACTTTG |
| 1 | GMTACGGTCACACTG | chrX:2192909-2194909 | 1820 | 1834 | - | 9.5643 | 1.06E-05 | 0.4 | CACTCGGTCACACTG |
| 1 | GMTACGGTCACACTG | chr2R:16866714-16868714 | 1840 | 1854 | - | 9.5643 | 1.06E-05 | 0.4 | GCCACGTTCACATTG |
| 1 | GMTACGGTCACACTG | chr2R:6208468-6210468 | 1842 | 1856 | - | 9.5643 | 1.06E-05 | 0.4 | GATGCGGCCAAACTG |
| 1 | GMTACGGTCACACTG | chrX:4312626-4314626 | 1882 | 1896 | - | 9.5643 | 1.06E-05 | 0.4 | GATGCGCACACACTG |
| 1 | GMTACGGTCACACTG | chr3R:6858192-6860192 | 1903 | 1917 | - | 9.5643 | 1.06E-05 | 0.4 | GCTCCTGTCACACTT |
| 1 | GMTACGGTCACACTG | chr2R:16229198-16231198 | 1942 | 1956 | - | 9.5643 | 1.06E-05 | 0.4 | GCGACCGTAACACTG |

| | | | | | | | | | |
|---|---|---|---|---|---|---|---|---|---|
| 1 | GMTACGGTCACACTG | chr2R:9880396-9882396 | 1948 | 1962 | - | 9.5643 | 1.06E-05 | 0.4 | CCAACGGTCACACTA |
| 1 | GMTACGGTCACACTG | chr3R:16212488-16214488 | 1950 | 1964 | - | 9.5643 | 1.06E-05 | 0.4 | GCCACGTTCAGACTG |
| 1 | GMTACGGTCACACTG | chr2R:15049355-15051355 | 1952 | 1966 | - | 9.5643 | 1.06E-05 | 0.4 | GATGCGGTTACCCTG |
| 1 | GMTACGGTCACACTG | chr2L:16798025-16800025 | 1968 | 1982 | - | 9.5643 | 1.06E-05 | 0.4 | ACGACGGTCACATTG |
| 1 | GMTACGGTCACACTG | chrX:13533400-13535400 | 5 | 19 | + | 9.5643 | 1.06E-05 | 0.4 | GAGACGCCCACACTG |
| 1 | GMTACGGTCACACTG | chr3R:22395880-22397880 | 46 | 60 | + | 9.5643 | 1.06E-05 | 0.4 | GCTACCGGCACACTC |
| 1 | GMTACGGTCACACTG | chr2L:20927824-20929824 | 53 | 67 | + | 9.5643 | 1.06E-05 | 0.4 | GAAACGGCCACACTT |
| 1 | GMTACGGTCACACTG | chr3R:29815451-29817451 | 58 | 72 | + | 9.5643 | 1.06E-05 | 0.4 | GCTGAGGTGACACTG |
| 1 | GMTACGGTCACACTG | chrX:19589170-19591170 | 72 | 86 | + | 9.5643 | 1.06E-05 | 0.4 | GAAACGATCACTCTG |
| 1 | GMTACGGTCACACTG | chr3L:6035490-6037490 | 89 | 103 | + | 9.5643 | 1.06E-05 | 0.4 | ACTACGGTGCCACTG |
| 1 | GMTACGGTCACACTG | chr3R:5484543-5486543 | 102 | 116 | + | 9.5643 | 1.06E-05 | 0.4 | GCTATGGTTACACAG |
| 1 | GMTACGGTCACACTG | chrX:4616564-4618564 | 103 | 117 | + | 9.5643 | 1.06E-05 | 0.4 | GATACGGATACACGG |
| 1 | GMTACGGTCACACTG | chr3L:9627676-9629676 | 106 | 120 | + | 9.5643 | 1.06E-05 | 0.4 | GCTATGGTCACAAAG |
| 1 | GMTACGGTCACACTG | chr3L:22274716-22276716 | 110 | 124 | + | 9.5643 | 1.06E-05 | 0.4 | GATGCGGTCTAACTG |
| 1 | GMTACGGTCACACTG | chr2R:24735657-24737657 | 117 | 131 | + | 9.5643 | 1.06E-05 | 0.4 | GCTGCGTTGACACTG |
| 1 | GMTACGGTCACACTG | chr2L:18589891-18591891 | 121 | 135 | + | 9.5643 | 1.06E-05 | 0.4 | GCTAAGGACACACTT |
| 1 | GMTACGGTCACACTG | chr3R:11786014-11788014 | 128 | 142 | + | 9.5643 | 1.06E-05 | 0.4 | GCTGCAGTCACATTG |
| 1 | GMTACGGTCACACTG | chrX:19958403-19960403 | 131 | 145 | + | 9.5643 | 1.06E-05 | 0.4 | GCTATGGTCACGGTG |
| 1 | GMTACGGTCACACTG | chr2R:12751936-12753936 | 137 | 151 | + | 9.5643 | 1.06E-05 | 0.4 | GCTATGGTCTCATTG |
| 1 | GMTACGGTCACACTG | chr2L:13398831-13400831 | 152 | 166 | + | 9.5643 | 1.06E-05 | 0.4 | GATGCGGTCACATCG |
| 1 | GMTACGGTCACACTG | chr2R:9163967-9165967 | 173 | 187 | + | 9.5643 | 1.06E-05 | 0.4 | GAATAGGTCACACTG |
| 1 | GMTACGGTCACACTG | chr2L:18011380-18013380 | 182 | 196 | + | 9.5643 | 1.06E-05 | 0.4 | CCAAAGGTCACACTG |
| 1 | GMTACGGTCACACTG | chrX:11476058-11478058 | 197 | 211 | + | 9.5643 | 1.06E-05 | 0.4 | GCTACTGTAGCACTG |
| 1 | GMTACGGTCACACTG | chr3L:3256878-3258878 | 205 | 219 | + | 9.5643 | 1.06E-05 | 0.4 | GAAACGCTCACACAG |
| 1 | GMTACGGTCACACTG | chrX:2326128-2328128 | 206 | 220 | + | 9.5643 | 1.06E-05 | 0.4 | GCTACAGTCAAACGG |
| 1 | GMTACGGTCACACTG | chr3R:13405998-13407998 | 221 | 235 | + | 9.5643 | 1.06E-05 | 0.4 | GCTACTGTGCCACTG |
| 1 | GMTACGGTCACACTG | chrX:6820016-6822016 | 245 | 259 | + | 9.5643 | 1.06E-05 | 0.4 | GCCACCGTCACACTC |
| 1 | GMTACGGTCACACTG | chr3L:10885471-10887471 | 252 | 266 | + | 9.5643 | 1.06E-05 | 0.4 | GCTAAGGGCAGACTG |
| 1 | GMTACGGTCACACTG | chrX:18357881-18359881 | 254 | 268 | + | 9.5643 | 1.06E-05 | 0.4 | GCTAAGCTCCCACTG |
| 1 | GMTACGGTCACACTG | chr2R:12923567-12925567 | 310 | 324 | + | 9.5643 | 1.06E-05 | 0.4 | GCTCCGGTCAAACTT |
| 1 | GMTACGGTCACACTG | chr3R:21272817-21274817 | 315 | 329 | + | 9.5643 | 1.06E-05 | 0.4 | GCTACGTTCCCATTG |
| 1 | GMTACGGTCACACTG | chr2R:23889512-23891512 | 333 | 347 | + | 9.5643 | 1.06E-05 | 0.4 | GATGCGGTCGCACTC |
| 1 | GMTACGGTCACACTG | chrX:3174018-3176018 | 336 | 350 | + | 9.5643 | 1.06E-05 | 0.4 | TCTACGGAAACACTG |

| | | | | | | | | | |
|---|---|---|---|---|---|---|---|---|---|
| 1 | GMTACGGTCACACTG | chrX:6819919-6821919 | 342 | 356 | + | 9.5643 | 1.06E-05 | 0.4 | GCCACCGTCACACTC |
| 1 | GMTACGGTCACACTG | chr3L:9072105-9074105 | 350 | 364 | + | 9.5643 | 1.06E-05 | 0.4 | GATACGGACACACCC |
| 1 | GMTACGGTCACACTG | chr3R:20563067-20565067 | 371 | 385 | + | 9.5643 | 1.06E-05 | 0.4 | CATACGGTCCCACAG |
| 1 | GMTACGGTCACACTG | chr3R:29147248-29149248 | 388 | 402 | + | 9.5643 | 1.06E-05 | 0.4 | GATAGGGTCGCACCG |
| 1 | GMTACGGTCACACTG | chr3R:22395528-22397528 | 398 | 412 | + | 9.5643 | 1.06E-05 | 0.4 | GCTACCGGCACACTC |
| 1 | GMTACGGTCACACTG | chr2R:21756345-21758345 | 436 | 450 | + | 9.5643 | 1.06E-05 | 0.4 | GACACGGACACACTC |
| 1 | GMTACGGTCACACTG | chrX:2946094-2948094 | 441 | 455 | + | 9.5643 | 1.06E-05 | 0.4 | GCTACTGCCACACTT |
| 1 | GMTACGGTCACACTG | chr2L:7993798-7995798 | 459 | 473 | + | 9.5643 | 1.06E-05 | 0.4 | GCTACGGTGACTGTG |
| 1 | GMTACGGTCACACTG | chr3R:29147172-29149172 | 464 | 478 | + | 9.5643 | 1.06E-05 | 0.4 | GATAGGGTCGCACCG |
| 1 | GMTACGGTCACACTG | chr3L:14074772-14076772 | 464 | 478 | + | 9.5643 | 1.06E-05 | 0.4 | GATCCGGTCGCATTG |
| 1 | GMTACGGTCACACTG | chr3L:9718328-9720328 | 471 | 485 | + | 9.5643 | 1.06E-05 | 0.4 | GATTCGGTGTCACTG |
| 1 | GMTACGGTCACACTG | chr3R:13041770-13043770 | 494 | 508 | + | 9.5643 | 1.06E-05 | 0.4 | GCTACGGTTACAACG |
| 1 | GMTACGGTCACACTG | chrX:9867568-9869568 | 507 | 521 | + | 9.5643 | 1.06E-05 | 0.4 | TCGACGGTCACATTG |
| 1 | GMTACGGTCACACTG | chr3L:4159932-4161932 | 509 | 523 | + | 9.5643 | 1.06E-05 | 0.4 | GCTACGGACAGATTG |
| 1 | GMTACGGTCACACTG | chr2R:10246814-10248814 | 518 | 532 | + | 9.5643 | 1.06E-05 | 0.4 | GCTATGGTCACCATG |
| 1 | GMTACGGTCACACTG | chr3R:20830127-20832127 | 518 | 532 | + | 9.5643 | 1.06E-05 | 0.4 | GCGATGGTAACACTG |
| 1 | GMTACGGTCACACTG | chrX:19960297-19962297 | 525 | 539 | + | 9.5643 | 1.06E-05 | 0.4 | GATACGGACAGACGG |
| 1 | GMTACGGTCACACTG | chr3R:19389270-19391270 | 538 | 552 | + | 9.5643 | 1.06E-05 | 0.4 | GATATGGTCGCACCG |
| 1 | GMTACGGTCACACTG | chr2L:13191201-13193201 | 562 | 576 | + | 9.5643 | 1.06E-05 | 0.4 | GCTACGCACACACTC |
| 1 | GMTACGGTCACACTG | chr3R:29122370-29124370 | 569 | 583 | + | 9.5643 | 1.06E-05 | 0.4 | GAGCCGTTCACACTG |
| 1 | GMTACGGTCACACTG | chr2R:23889276-23891276 | 569 | 583 | + | 9.5643 | 1.06E-05 | 0.4 | GATGCGGTCGCACTC |
| 1 | GMTACGGTCACACTG | chr3R:8315984-8317984 | 597 | 611 | + | 9.5643 | 1.06E-05 | 0.4 | GATCCGGTGGCACTG |
| 1 | GMTACGGTCACACTG | chr3L:22605369-22607369 | 605 | 619 | + | 9.5643 | 1.06E-05 | 0.4 | GCTTCTGTCACACTT |
| 1 | GMTACGGTCACACTG | chr3L:5905699-5907699 | 622 | 636 | + | 9.5643 | 1.06E-05 | 0.4 | GAAACGGCCAAACTG |
| 1 | GMTACGGTCACACTG | chrX:14236988-14238988 | 633 | 647 | + | 9.5643 | 1.06E-05 | 0.4 | GCTATGGTCACGTTG |
| 1 | GMTACGGTCACACTG | chr3L:9718144-9720144 | 655 | 669 | + | 9.5643 | 1.06E-05 | 0.4 | GATTCGGTGTCACTG |
| 1 | GMTACGGTCACACTG | chr2R:23906654-23908654 | 667 | 681 | + | 9.5643 | 1.06E-05 | 0.4 | GCTGTGGTCACACTA |
| 1 | GMTACGGTCACACTG | chr3R:25246658-25248658 | 668 | 682 | + | 9.5643 | 1.06E-05 | 0.4 | GCTCCGTTTACACTG |
| 1 | GMTACGGTCACACTG | chr2L:18589342-18591342 | 670 | 684 | + | 9.5643 | 1.06E-05 | 0.4 | GCTAAGGACACACTT |
| 1 | GMTACGGTCACACTG | chr2L:418952-420952 | 685 | 699 | + | 9.5643 | 1.06E-05 | 0.4 | GCTATGGGCACACTA |
| 1 | GMTACGGTCACACTG | chrX:14236923-14238923 | 698 | 712 | + | 9.5643 | 1.06E-05 | 0.4 | GCTATGGTCACGTTG |
| 1 | GMTACGGTCACACTG | chr2L:8300079-8302079 | 702 | 716 | + | 9.5643 | 1.06E-05 | 0.4 | GAGACGCTCACACAG |
| 1 | GMTACGGTCACACTG | chr2R:12995501-12997501 | 711 | 725 | + | 9.5643 | 1.06E-05 | 0.4 | TACACGGGCACACTG |

| | | | | | | | | | |
|---|---|---|---|---|---|---|---|---|---|
| 1 | GMTACGGTCACACTG | chr2L:13018434-13020434 | 722 | 736 | + | 9.5643 | 1.06E-05 | 0.4 | CCGACAGTCACACTG |
| 1 | GMTACGGTCACACTG | chr3L:9071731-9073731 | 724 | 738 | + | 9.5643 | 1.06E-05 | 0.4 | GATACGGACACACCC |
| 1 | GMTACGGTCACACTG | chr3R:5590763-5592763 | 741 | 755 | + | 9.5643 | 1.06E-05 | 0.4 | ACTCCGATCACACTG |
| 1 | GMTACGGTCACACTG | chr2L:261203-263203 | 741 | 755 | + | 9.5643 | 1.06E-05 | 0.4 | GATACGGGCTCAATG |
| 1 | GMTACGGTCACACTG | chr3L:16196868-16198868 | 745 | 759 | + | 9.5643 | 1.06E-05 | 0.4 | GATACGGACAAACGG |
| 1 | GMTACGGTCACACTG | chr2R:23356660-23358660 | 754 | 768 | + | 9.5643 | 1.06E-05 | 0.4 | GAAACGGGCACACTA |
| 1 | GMTACGGTCACACTG | chrX:20933869-20935869 | 770 | 784 | + | 9.5643 | 1.06E-05 | 0.4 | GATAAGGTCCCTCTG |
| 1 | GMTACGGTCACACTG | chr2R:10246560-10248560 | 772 | 786 | + | 9.5643 | 1.06E-05 | 0.4 | GCTATGGTCACCATG |
| 1 | GMTACGGTCACACTG | chrX:21155621-21157621 | 777 | 791 | + | 9.5643 | 1.06E-05 | 0.4 | GCTATGGTCACGGTG |
| 1 | GMTACGGTCACACTG | chr2L:13190979-13192979 | 784 | 798 | + | 9.5643 | 1.06E-05 | 0.4 | GCTACGCACACACTC |
| 1 | GMTACGGTCACACTG | chrX:19914685-19916685 | 789 | 803 | + | 9.5643 | 1.06E-05 | 0.4 | GCTATGGTCACGGTG |
| 1 | GMTACGGTCACACTG | chrX:19938396-19940396 | 789 | 803 | + | 9.5643 | 1.06E-05 | 0.4 | GCTATGGTCACGGTG |
| 1 | GMTACGGTCACACTG | chrX:19909521-19911521 | 789 | 803 | + | 9.5643 | 1.06E-05 | 0.4 | GCTATGGTCACGGTG |
| 1 | GMTACGGTCACACTG | chrX:19928580-19930580 | 789 | 803 | + | 9.5643 | 1.06E-05 | 0.4 | GCTATGGTCACGGTG |
| 1 | GMTACGGTCACACTG | chr3L:2710690-2712690 | 802 | 816 | + | 9.5643 | 1.06E-05 | 0.4 | GCTACGCACACACTC |
| 1 | GMTACGGTCACACTG | chr3R:25246514-25248514 | 812 | 826 | + | 9.5643 | 1.06E-05 | 0.4 | GCTCCGTTTACACTG |
| 1 | GMTACGGTCACACTG | chr2R:23356565-23358565 | 849 | 863 | + | 9.5643 | 1.06E-05 | 0.4 | GAAACGGGCACACTA |
| 1 | GMTACGGTCACACTG | chr2R:11370335-11372335 | 853 | 867 | + | 9.5643 | 1.06E-05 | 0.4 | TCGATGGTCACACTG |
| 1 | GMTACGGTCACACTG | chr2L:7422820-7424820 | 857 | 871 | + | 9.5643 | 1.06E-05 | 0.4 | GAAACGGCAACACTG |
| 1 | GMTACGGTCACACTG | chr2R:7406819-7408819 | 864 | 878 | + | 9.5643 | 1.06E-05 | 0.4 | GATTCGAGCACACTG |
| 1 | GMTACGGTCACACTG | chrX:19212254-19214254 | 877 | 891 | + | 9.5643 | 1.06E-05 | 0.4 | GATACGGACATACGG |
| 1 | GMTACGGTCACACTG | chrX:3173474-3175474 | 880 | 894 | + | 9.5643 | 1.06E-05 | 0.4 | TCTACGGAAACACTG |
| 1 | GMTACGGTCACACTG | chr3L:210457-212457 | 881 | 895 | + | 9.5643 | 1.06E-05 | 0.4 | GATGCGCGCACACTG |
| 1 | GMTACGGTCACACTG | chr3R:7919431-7921431 | 883 | 897 | + | 9.5643 | 1.06E-05 | 0.4 | GAAGCGTTCACACTG |
| 1 | GMTACGGTCACACTG | chr3L:17529202-17531202 | 889 | 903 | + | 9.5643 | 1.06E-05 | 0.4 | TACACGGTCACTCTG |
| 1 | GMTACGGTCACACTG | chr3R:31212458-31214458 | 899 | 913 | + | 9.5643 | 1.06E-05 | 0.4 | TAAACGGTCACACAG |
| 1 | GMTACGGTCACACTG | chr3R:25246422-25248422 | 904 | 918 | + | 9.5643 | 1.06E-05 | 0.4 | GCTCCGTTTACACTG |
| 1 | GMTACGGTCACACTG | chrX:19943417-19945417 | 920 | 934 | + | 9.5643 | 1.06E-05 | 0.4 | GCTATGGTCACGGTG |
| 1 | GMTACGGTCACACTG | chrX:19933565-19935565 | 920 | 934 | + | 9.5643 | 1.06E-05 | 0.4 | GCTATGGTCACGGTG |
| 1 | GMTACGGTCACACTG | chrX:19948147-19950147 | 920 | 934 | + | 9.5643 | 1.06E-05 | 0.4 | GCTATGGTCACGGTG |
| 1 | GMTACGGTCACACTG | chrX:19919708-19921708 | 920 | 934 | + | 9.5643 | 1.06E-05 | 0.4 | GCTATGGTCACGGTG |
| 1 | GMTACGGTCACACTG | chrX:19952882-19954882 | 920 | 934 | + | 9.5643 | 1.06E-05 | 0.4 | GCTATGGTCACGGTG |
| 1 | GMTACGGTCACACTG | chr2L:6716625-6718625 | 941 | 955 | + | 9.5643 | 1.06E-05 | 0.4 | TAAGCGGTCACACTG |

| | | | | | | | | | |
|---|---|---|---|---|---|---|---|---|---|
| 1 | GMTACGGTCACACTG | chrX:18829591-18831591 | 947 | 961 | + | 9.5643 | 1.06E-05 | 0.4 | GCTGCAGTCACACTT |
| 1 | GMTACGGTCACACTG | chr2R:21322430-21324430 | 950 | 964 | + | 9.5643 | 1.06E-05 | 0.4 | GAAATGGTCACACCG |
| 1 | GMTACGGTCACACTG | chr2R:16233936-16235936 | 951 | 965 | + | 9.5643 | 1.06E-05 | 0.4 | GATACGGTCTCGGTG |
| 1 | GMTACGGTCACACTG | chrX:9355634-9357634 | 952 | 966 | + | 9.5643 | 1.06E-05 | 0.4 | CCAATGGTCACACTG |
| 1 | GMTACGGTCACACTG | chr2R:12205092-12207092 | 966 | 980 | + | 9.5643 | 1.06E-05 | 0.4 | GCCAAGGTTACACTG |
| 1 | GMTACGGTCACACTG | chr3L:9426010-9428010 | 967 | 981 | + | 9.5643 | 1.06E-05 | 0.4 | GCTGCGGTTACACTT |
| 1 | GMTACGGTCACACTG | chrX:18485921-18487921 | 968 | 982 | + | 9.5643 | 1.06E-05 | 0.4 | GAAACGGTCAGGCTG |
| 1 | GMTACGGTCACACTG | chr2L:1883622-1885622 | 971 | 985 | + | 9.5643 | 1.06E-05 | 0.4 | CCTCTGGTCACACTG |
| 1 | GMTACGGTCACACTG | chr2R:21281298-21283298 | 972 | 986 | + | 9.5643 | 1.06E-05 | 0.4 | TCTGTGGTCACACTG |
| 1 | GMTACGGTCACACTG | chr2R:12922904-12924904 | 973 | 987 | + | 9.5643 | 1.06E-05 | 0.4 | GCTCCGGTCAAACTT |
| 1 | GMTACGGTCACACTG | chr3R:28695242-28697242 | 980 | 994 | + | 9.5643 | 1.06E-05 | 0.4 | GCTCTGGTAACACTG |
| 1 | GMTACGGTCACACTG | chr3L:4002474-4004474 | 980 | 994 | + | 9.5643 | 1.06E-05 | 0.4 | GCTATGGTCACACAA |
| 1 | GMTACGGTCACACTG | chr3R:31212376-31214376 | 981 | 995 | + | 9.5643 | 1.06E-05 | 0.4 | TAAACGGTCACACAG |
| 1 | GMTACGGTCACACTG | chr2R:17019027-17021027 | 981 | 995 | + | 9.5643 | 1.06E-05 | 0.4 | AAAACGGTCACACAG |
| 1 | GMTACGGTCACACTG | chr3R:17079244-17081244 | 982 | 996 | + | 9.5643 | 1.06E-05 | 0.4 | GCTAAGGTCCCACTA |
| 1 | GMTACGGTCACACTG | chr2L:418652-420652 | 985 | 999 | + | 9.5643 | 1.06E-05 | 0.4 | GCTATGGGCACACTA |
| 1 | GMTACGGTCACACTG | chr3R:20831799-20833799 | 986 | 1000 | + | 9.5643 | 1.06E-05 | 0.4 | GAAACGGTCACACCT |
| 1 | GMTACGGTCACACTG | chr2R:18687132-18689132 | 986 | 1000 | + | 9.5643 | 1.06E-05 | 0.4 | TACACGGCCACACTG |
| 1 | GMTACGGTCACACTG | chrX:11368665-11370665 | 986 | 1000 | + | 9.5643 | 1.06E-05 | 0.4 | CCTCTGGTCACACTG |
| 1 | GMTACGGTCACACTG | chr2L:3513872-3515872 | 987 | 1001 | + | 9.5643 | 1.06E-05 | 0.4 | GCGACTGTCACACTA |
| 1 | GMTACGGTCACACTG | chr3R:25850632-25852632 | 988 | 1002 | + | 9.5643 | 1.06E-05 | 0.4 | GAAACTGTCTCACTG |
| 1 | GMTACGGTCACACTG | chr2R:14962292-14964292 | 988 | 1002 | + | 9.5643 | 1.06E-05 | 0.4 | GCTGCGATCACACTA |
| 1 | GMTACGGTCACACTG | chr2R:5753517-5755517 | 991 | 1005 | + | 9.5643 | 1.06E-05 | 0.4 | CCTGCGGTCACACTA |
| 1 | GMTACGGTCACACTG | chr2L:21260168-21262168 | 992 | 1006 | + | 9.5643 | 1.06E-05 | 0.4 | ACTACAGCCACACTG |
| 1 | GMTACGGTCACACTG | chrX:13562823-13564823 | 992 | 1006 | + | 9.5643 | 1.06E-05 | 0.4 | CCGACGGTCACACTC |
| 1 | GMTACGGTCACACTG | chr2L:13397991-13399991 | 992 | 1006 | + | 9.5643 | 1.06E-05 | 0.4 | GATGCGGTCACATCG |
| 1 | GMTACGGTCACACTG | chr3L:20520009-20522009 | 992 | 1006 | + | 9.5643 | 1.06E-05 | 0.4 | ACCACGGTCACACTA |
| 1 | GMTACGGTCACACTG | chrX:13562823-13564823 | 992 | 1006 | + | 9.5643 | 1.06E-05 | 0.4 | CCGACGGTCACACTC |
| 1 | GMTACGGTCACACTG | chr2L:6423336-6425336 | 993 | 1007 | + | 9.5643 | 1.06E-05 | 0.4 | GAAACGGTCACACGT |
| 1 | GMTACGGTCACACTG | chr3R:27581840-27583840 | 993 | 1007 | + | 9.5643 | 1.06E-05 | 0.4 | CAATCGGTCACACTG |
| 1 | GMTACGGTCACACTG | chr2L:21178933-21180933 | 993 | 1007 | + | 9.5643 | 1.06E-05 | 0.4 | TACCCGGTCACACTG |
| 1 | GMTACGGTCACACTG | chr3R:24850601-24852601 | 993 | 1007 | + | 9.5643 | 1.06E-05 | 0.4 | GAAACGGTCACACGC |
| 1 | GMTACGGTCACACTG | chr3L:3810263-3812263 | 993 | 1007 | + | 9.5643 | 1.06E-05 | 0.4 | CCTCTGGTCACACTG |

| | | | | | | | | | |
|---|---|---|---|---|---|---|---|---|---|
| 1 | GMTACGGTCACACTG | chr3L:9429312-9431312 | 994 | 1008 | + | 9.5643 | 1.06E-05 | 0.4 | CCCATGGTCACACTG |
| 1 | GMTACGGTCACACTG | chr3L:7368843-7370843 | 994 | 1008 | + | 9.5643 | 1.06E-05 | 0.4 | CATGCGGTCACTCTG |
| 1 | GMTACGGTCACACTG | chr3R:11881790-11883790 | 996 | 1010 | + | 9.5643 | 1.06E-05 | 0.4 | ACAACGGTCACACTA |
| 1 | GMTACGGTCACACTG | chr3R:4905622-4907622 | 999 | 1013 | + | 9.5643 | 1.06E-05 | 0.4 | GACGCGGTTACACTG |
| 1 | GMTACGGTCACACTG | chr3R:25892495-25894495 | 1000 | 1014 | + | 9.5643 | 1.06E-05 | 0.4 | GCAATGGTCACATTG |
| 1 | GMTACGGTCACACTG | chr3L:3046087-3048087 | 1000 | 1014 | + | 9.5643 | 1.06E-05 | 0.4 | CCGACGGTAACACTG |
| 1 | GMTACGGTCACACTG | chr2L:6657517-6659517 | 1000 | 1014 | + | 9.5643 | 1.06E-05 | 0.4 | CCAACGGTCACACTA |
| 1 | GMTACGGTCACACTG | chrX:16942290-16944290 | 1001 | 1015 | + | 9.5643 | 1.06E-05 | 0.4 | CCAACGGTCACACTA |
| 1 | GMTACGGTCACACTG | chr2R:24036943-24038943 | 1002 | 1016 | + | 9.5643 | 1.06E-05 | 0.4 | ACTCTGGTCACACTG |
| 1 | GMTACGGTCACACTG | chr2R:16177826-16179826 | 1002 | 1016 | + | 9.5643 | 1.06E-05 | 0.4 | ACTGCGGTCACACTA |
| 1 | GMTACGGTCACACTG | chr3R:18365638-18367638 | 1007 | 1021 | + | 9.5643 | 1.06E-05 | 0.4 | GCTCTGGTCACACTT |
| 1 | GMTACGGTCACACTG | chr2L:2856164-2858164 | 1008 | 1022 | + | 9.5643 | 1.06E-05 | 0.4 | AACGCGGTCACACTG |
| 1 | GMTACGGTCACACTG | chr2R:18854087-18856087 | 1009 | 1023 | + | 9.5643 | 1.06E-05 | 0.4 | GCTACGATTCCACTG |
| 1 | GMTACGGTCACACTG | chrX:9148309-9150309 | 1009 | 1023 | + | 9.5643 | 1.06E-05 | 0.4 | GATACGGCCACACCT |
| 1 | GMTACGGTCACACTG | chr3L:17529081-17531081 | 1010 | 1024 | + | 9.5643 | 1.06E-05 | 0.4 | TACACGGTCACTCTG |
| 1 | GMTACGGTCACACTG | chr3R:6645889-6647889 | 1010 | 1024 | + | 9.5643 | 1.06E-05 | 0.4 | TCGACGGTCACACTT |
| 1 | GMTACGGTCACACTG | chr3L:3142808-3144808 | 1011 | 1025 | + | 9.5643 | 1.06E-05 | 0.4 | ACCACGGTCACACTT |
| 1 | GMTACGGTCACACTG | chr2L:8196193-8198193 | 1011 | 1025 | + | 9.5643 | 1.06E-05 | 0.4 | GCTATGGTCACTTTG |
| 1 | GMTACGGTCACACTG | chr2R:8921785-8923785 | 1012 | 1026 | + | 9.5643 | 1.06E-05 | 0.4 | CACACGGTCCCACTG |
| 1 | GMTACGGTCACACTG | chr2R:20089795-20091795 | 1015 | 1029 | + | 9.5643 | 1.06E-05 | 0.4 | GCTGCGGTCACATTA |
| 1 | GMTACGGTCACACTG | chr3R:7919299-7921299 | 1015 | 1029 | + | 9.5643 | 1.06E-05 | 0.4 | GAAGCGTTCACACTG |
| 1 | GMTACGGTCACACTG | chr2L:22248876-22250876 | 1017 | 1031 | + | 9.5643 | 1.06E-05 | 0.4 | TCTATGGCCACACTG |
| 1 | GMTACGGTCACACTG | chr2L:20926860-20928860 | 1017 | 1031 | + | 9.5643 | 1.06E-05 | 0.4 | GAAACGGCCACACTT |
| 1 | GMTACGGTCACACTG | chr3L:12529171-12531171 | 1018 | 1032 | + | 9.5643 | 1.06E-05 | 0.4 | CAAGCGGTCACACTG |
| 1 | GMTACGGTCACACTG | chrX:15761956-15763956 | 1018 | 1032 | + | 9.5643 | 1.06E-05 | 0.4 | GAAGCGGTCACACTA |
| 1 | GMTACGGTCACACTG | chr3R:19876737-19878737 | 1021 | 1035 | + | 9.5643 | 1.06E-05 | 0.4 | CCTCTGGTCACACTG |
| 1 | GMTACGGTCACACTG | chrX:6371915-6373915 | 1022 | 1036 | + | 9.5643 | 1.06E-05 | 0.4 | GCAACGGTTATACTG |
| 1 | GMTACGGTCACACTG | chr3R:11619090-11621090 | 1023 | 1037 | + | 9.5643 | 1.06E-05 | 0.4 | AATGCGGTCACACAG |
| 1 | GMTACGGTCACACTG | chr2L:6676625-6678625 | 1023 | 1037 | + | 9.5643 | 1.06E-05 | 0.4 | ACTTCGGTCACACTA |
| 1 | GMTACGGTCACACTG | chr3L:3049323-3051323 | 1024 | 1038 | + | 9.5643 | 1.06E-05 | 0.4 | CAGCCGGTCACACTG |
| 1 | GMTACGGTCACACTG | chr3L:1664138-1666138 | 1026 | 1040 | + | 9.5643 | 1.06E-05 | 0.4 | GAAACGGGCACACTA |
| 1 | GMTACGGTCACACTG | chr2R:23906294-23908294 | 1027 | 1041 | + | 9.5643 | 1.06E-05 | 0.4 | GCTGTGGTCACACTA |
| 1 | GMTACGGTCACACTG | chr2R:24036915-24038915 | 1030 | 1044 | + | 9.5643 | 1.06E-05 | 0.4 | ACTCTGGTCACACTG |

| | | | | | | | | | |
|---|---|---|---|---|---|---|---|---|---|
| 1 | GMTACGGTCACACTG | chr2R:12874164-12876164 | 1031 | 1045 | + | 9.5643 | 1.06E-05 | 0.4 | GCTAAGGTGGCACTG |
| 1 | GMTACGGTCACACTG | chrX:800479-802479 | 1037 | 1051 | + | 9.5643 | 1.06E-05 | 0.4 | CATGCGGTCACACCG |
| 1 | GMTACGGTCACACTG | chr3R:8315543-8317543 | 1038 | 1052 | + | 9.5643 | 1.06E-05 | 0.4 | GATCCGGTGGCACTG |
| 1 | GMTACGGTCACACTG | chr2R:8921755-8923755 | 1042 | 1056 | + | 9.5643 | 1.06E-05 | 0.4 | CACACGGTCCCACTG |
| 1 | GMTACGGTCACACTG | chr3L:11628505-11630505 | 1045 | 1059 | + | 9.5643 | 1.06E-05 | 0.4 | GAAACGTTCACTCTG |
| 1 | GMTACGGTCACACTG | chr3R:5590459-5592459 | 1045 | 1059 | + | 9.5643 | 1.06E-05 | 0.4 | ACTCCGATCACACTG |
| 1 | GMTACGGTCACACTG | chr2R:10781895-10783895 | 1048 | 1062 | + | 9.5643 | 1.06E-05 | 0.4 | CATGCGGCCACACTG |
| 1 | GMTACGGTCACACTG | chr3R:10705974-10707974 | 1050 | 1064 | + | 9.5643 | 1.06E-05 | 0.4 | GATCCGGTCATACGG |
| 1 | GMTACGGTCACACTG | chr2L:5538196-5540196 | 1056 | 1070 | + | 9.5643 | 1.06E-05 | 0.4 | GAAACGGACACACTA |
| 1 | GMTACGGTCACACTG | chr3R:29121879-29123879 | 1060 | 1074 | + | 9.5643 | 1.06E-05 | 0.4 | GAGCCGTTCACACTG |
| 1 | GMTACGGTCACACTG | chr2R:21322320-21324320 | 1060 | 1074 | + | 9.5643 | 1.06E-05 | 0.4 | GAAATGGTCACACCG |
| 1 | GMTACGGTCACACTG | chr3R:20829580-20831580 | 1065 | 1079 | + | 9.5643 | 1.06E-05 | 0.4 | GCGATGGTAACACTG |
| 1 | GMTACGGTCACACTG | chr2L:7993184-7995184 | 1073 | 1087 | + | 9.5643 | 1.06E-05 | 0.4 | GCTACGGTGACTGTG |
| 1 | GMTACGGTCACACTG | chr2L:15440601-15442601 | 1075 | 1089 | + | 9.5643 | 1.06E-05 | 0.4 | GCTACGGACAAACTA |
| 1 | GMTACGGTCACACTG | chr2L:18010475-18012475 | 1087 | 1101 | + | 9.5643 | 1.06E-05 | 0.4 | CCAAAGGTCACACTG |
| 1 | GMTACGGTCACACTG | chr2L:10849848-10851848 | 1096 | 1110 | + | 9.5643 | 1.06E-05 | 0.4 | GCAATGGTCACACTT |
| 1 | GMTACGGTCACACTG | chr3L:1664064-1666064 | 1100 | 1114 | + | 9.5643 | 1.06E-05 | 0.4 | GAAACGGGCACACTA |
| 1 | GMTACGGTCACACTG | chr3R:22549680-22551680 | 1101 | 1115 | + | 9.5643 | 1.06E-05 | 0.4 | GCTACGGTCGTAGTG |
| 1 | GMTACGGTCACACTG | chr3R:7246090-7248090 | 1117 | 1131 | + | 9.5643 | 1.06E-05 | 0.4 | GCAATGGTCACACTT |
| 1 | GMTACGGTCACACTG | chrX:2325217-2327217 | 1117 | 1131 | + | 9.5643 | 1.06E-05 | 0.4 | GCTACAGTCAAACGG |
| 1 | GMTACGGTCACACTG | chr2L:820289-822289 | 1139 | 1153 | + | 9.5643 | 1.06E-05 | 0.4 | GAACTGGTCACACTG |
| 1 | GMTACGGTCACACTG | chr3L:16233140-16235140 | 1140 | 1154 | + | 9.5643 | 1.06E-05 | 0.4 | GCCACGGTCACAATA |
| 1 | GMTACGGTCACACTG | chr3L:3810114-3812114 | 1142 | 1156 | + | 9.5643 | 1.06E-05 | 0.4 | CCTCTGGTCACACTG |
| 1 | GMTACGGTCACACTG | chr2L:7425866-7427866 | 1143 | 1157 | + | 9.5643 | 1.06E-05 | 0.4 | TCTGCGTTCACACTG |
| 1 | GMTACGGTCACACTG | chr3R:17027693-17029693 | 1147 | 1161 | + | 9.5643 | 1.06E-05 | 0.4 | GAGACGGGCAGACTG |
| 1 | GMTACGGTCACACTG | chr3R:4415438-4417438 | 1155 | 1169 | + | 9.5643 | 1.06E-05 | 0.4 | GAAACCGTCACCCTG |
| 1 | GMTACGGTCACACTG | chr3R:10705863-10707863 | 1161 | 1175 | + | 9.5643 | 1.06E-05 | 0.4 | GATCCGGTCATACGG |
| 1 | GMTACGGTCACACTG | chr2R:9706536-9708536 | 1162 | 1176 | + | 9.5643 | 1.06E-05 | 0.4 | GACACGGCGACACTG |
| 1 | GMTACGGTCACACTG | chr2R:8365688-8367688 | 1174 | 1188 | + | 9.5643 | 1.06E-05 | 0.4 | GCTACGCCCACACTT |
| 1 | GMTACGGTCACACTG | chr2L:17362496-17364496 | 1176 | 1190 | + | 9.5643 | 1.06E-05 | 0.4 | GAAACGGACACAGTG |
| 1 | GMTACGGTCACACTG | chr3R:13287813-13289813 | 1176 | 1190 | + | 9.5643 | 1.06E-05 | 0.4 | GATGCCGGCACACTG |
| 1 | GMTACGGTCACACTG | chr2R:12874012-12876012 | 1183 | 1197 | + | 9.5643 | 1.06E-05 | 0.4 | GCTAAGGTGGCACTG |
| 1 | GMTACGGTCACACTG | chr2L:709568-711568 | 1189 | 1203 | + | 9.5643 | 1.06E-05 | 0.4 | GATACGGCCAGTCTG |

| | | | | | | | | | |
|---|---|---|---|---|---|---|---|---|---|
| 1 | GMTACGGTCACACTG | chr2R:13109769-13111769 | 1189 | 1203 | + | 9.5643 | 1.06E-05 | 0.4 | AATTCGGACACACTG |
| 1 | GMTACGGTCACACTG | chrX:19943147-19945147 | 1190 | 1204 | + | 9.5643 | 1.06E-05 | 0.4 | GCTATGGTCACGGTG |
| 1 | GMTACGGTCACACTG | chrX:19933294-19935294 | 1191 | 1205 | + | 9.5643 | 1.06E-05 | 0.4 | GCTATGGTCACGGTG |
| 1 | GMTACGGTCACACTG | chrX:19928178-19930178 | 1191 | 1205 | + | 9.5643 | 1.06E-05 | 0.4 | GCTATGGTCACGGTG |
| 1 | GMTACGGTCACACTG | chrX:19937994-19939994 | 1191 | 1205 | + | 9.5643 | 1.06E-05 | 0.4 | GCTATGGTCACGGTG |
| 1 | GMTACGGTCACACTG | chrX:19957342-19959342 | 1192 | 1206 | + | 9.5643 | 1.06E-05 | 0.4 | GCTATGGTCACGGTG |
| 1 | GMTACGGTCACACTG | chrX:19919435-19921435 | 1193 | 1207 | + | 9.5643 | 1.06E-05 | 0.4 | GCTATGGTCACGGTG |
| 1 | GMTACGGTCACACTG | chrX:19909117-19911117 | 1193 | 1207 | + | 9.5643 | 1.06E-05 | 0.4 | GCTATGGTCACGGTG |
| 1 | GMTACGGTCACACTG | chrX:19914281-19916281 | 1193 | 1207 | + | 9.5643 | 1.06E-05 | 0.4 | GCTATGGTCACGGTG |
| 1 | GMTACGGTCACACTG | chrX:19952608-19954608 | 1194 | 1208 | + | 9.5643 | 1.06E-05 | 0.4 | GCTATGGTCACGGTG |
| 1 | GMTACGGTCACACTG | chrX:19947873-19949873 | 1194 | 1208 | + | 9.5643 | 1.06E-05 | 0.4 | GCTATGGTCACGGTG |
| 1 | GMTACGGTCACACTG | chr3L:1794392-1796392 | 1196 | 1210 | + | 9.5643 | 1.06E-05 | 0.4 | CCTGCGGTCACACTA |
| 1 | GMTACGGTCACACTG | chr2L:9937863-9939863 | 1203 | 1217 | + | 9.5643 | 1.06E-05 | 0.4 | GAGACCGGCACACTG |
| 1 | GMTACGGTCACACTG | chr3R:4730878-4732878 | 1203 | 1217 | + | 9.5643 | 1.06E-05 | 0.4 | GCTCTGGTAACACTG |
| 1 | GMTACGGTCACACTG | chr2L:3513656-3515656 | 1203 | 1217 | + | 9.5643 | 1.06E-05 | 0.4 | GCGACTGTCACACTA |
| 1 | GMTACGGTCACACTG | chr3R:12711299-12713299 | 1205 | 1219 | + | 9.5643 | 1.06E-05 | 0.4 | GCTAAGGTCCCACTA |
| 1 | GMTACGGTCACACTG | chr3R:11618906-11620906 | 1207 | 1221 | + | 9.5643 | 1.06E-05 | 0.4 | AATGCGGTCACACAG |
| 1 | GMTACGGTCACACTG | chr3R:15856470-15858470 | 1212 | 1226 | + | 9.5643 | 1.06E-05 | 0.4 | GAAACGGTCACGCTA |
| 1 | GMTACGGTCACACTG | chr3R:13041050-13043050 | 1214 | 1228 | + | 9.5643 | 1.06E-05 | 0.4 | GCTACGGTTACAACG |
| 1 | GMTACGGTCACACTG | chr3L:9425758-9427758 | 1219 | 1233 | + | 9.5643 | 1.06E-05 | 0.4 | GCTGCGGTTACACTT |
| 1 | GMTACGGTCACACTG | chrX:21155173-21157173 | 1225 | 1239 | + | 9.5643 | 1.06E-05 | 0.4 | GCTATGGTCACGGTG |
| 1 | GMTACGGTCACACTG | chr3L:20519772-20521772 | 1229 | 1243 | + | 9.5643 | 1.06E-05 | 0.4 | ACCACGGTCACACTA |
| 1 | GMTACGGTCACACTG | chr3L:17528861-17530861 | 1230 | 1244 | + | 9.5643 | 1.06E-05 | 0.4 | TACACGGTCACTCTG |
| 1 | GMTACGGTCACACTG | chr3R:7245969-7247969 | 1238 | 1252 | + | 9.5643 | 1.06E-05 | 0.4 | GCAATGGTCACACTT |
| 1 | GMTACGGTCACACTG | chrX:21323046-21325046 | 1240 | 1254 | + | 9.5643 | 1.06E-05 | 0.4 | ACTACGGTCACACCA |
| 1 | GMTACGGTCACACTG | chr2L:21178683-21180683 | 1243 | 1257 | + | 9.5643 | 1.06E-05 | 0.4 | TACCCGGTCACACTG |
| 1 | GMTACGGTCACACTG | chr3R:19876514-19878514 | 1244 | 1258 | + | 9.5643 | 1.06E-05 | 0.4 | CCTCTGGTCACACTG |
| 1 | GMTACGGTCACACTG | chrX:9148071-9150071 | 1247 | 1261 | + | 9.5643 | 1.06E-05 | 0.4 | GATACGGCCACACCT |
| 1 | GMTACGGTCACACTG | chr3R:11881534-11883534 | 1252 | 1266 | + | 9.5643 | 1.06E-05 | 0.4 | ACAACGGTCACACTA |
| 1 | GMTACGGTCACACTG | chr2R:4241934-4243934 | 1258 | 1272 | + | 9.5643 | 1.06E-05 | 0.4 | GAGCCGTTCACACTG |
| 1 | GMTACGGTCACACTG | chr3L:1794328-1796328 | 1260 | 1274 | + | 9.5643 | 1.06E-05 | 0.4 | CCTGCGGTCACACTA |
| 1 | GMTACGGTCACACTG | chr3L:1794328-1796328 | 1260 | 1274 | + | 9.5643 | 1.06E-05 | 0.4 | CCTGCGGTCACACTA |
| 1 | GMTACGGTCACACTG | chr2L:22168668-22170668 | 1262 | 1276 | + | 9.5643 | 1.06E-05 | 0.4 | GCTTCAGTCAAACTG |

| | | | | | | | | | |
|---|---|---|---|---|---|---|---|---|---|
| 1 | GMTACGGTCACACTG | chr3L:3045812-3047812 | 1275 | 1289 | + | 9.5643 | 1.06E-05 | 0.4 | CCGACGGTAACACTG |
| 1 | GMTACGGTCACACTG | chr2L:9937790-9939790 | 1276 | 1290 | + | 9.5643 | 1.06E-05 | 0.4 | GAGACCGGCACACTG |
| 1 | GMTACGGTCACACTG | chrX:21323009-21325009 | 1277 | 1291 | + | 9.5643 | 1.06E-05 | 0.4 | ACTACGGTCACACCA |
| 1 | GMTACGGTCACACTG | chr3R:17078945-17080945 | 1281 | 1295 | + | 9.5643 | 1.06E-05 | 0.4 | GCTAAGGTCCCACTA |
| 1 | GMTACGGTCACACTG | chr2L:8195918-8197918 | 1286 | 1300 | + | 9.5643 | 1.06E-05 | 0.4 | GCTATGGTCACTTTG |
| 1 | GMTACGGTCACACTG | chr3R:32014941-32016941 | 1298 | 1312 | + | 9.5643 | 1.06E-05 | 0.4 | GCTGAGGTTACACTG |
| 1 | GMTACGGTCACACTG | chr2R:21690295-21692295 | 1301 | 1315 | + | 9.5643 | 1.06E-05 | 0.4 | TCTCTGGTCACACTG |
| 1 | GMTACGGTCACACTG | chr2R:21690295-21692295 | 1301 | 1315 | + | 9.5643 | 1.06E-05 | 0.4 | TCTCTGGTCACACTG |
| 1 | GMTACGGTCACACTG | chr2R:14961969-14963969 | 1311 | 1325 | + | 9.5643 | 1.06E-05 | 0.4 | GCTGCGATCACACTA |
| 1 | GMTACGGTCACACTG | chr2R:19502068-19504068 | 1317 | 1331 | + | 9.5643 | 1.06E-05 | 0.4 | GATACGGTCACGCAA |
| 1 | GMTACGGTCACACTG | chr2L:13397664-13399664 | 1319 | 1333 | + | 9.5643 | 1.06E-05 | 0.4 | GATGCGGTCACATCG |
| 1 | GMTACGGTCACACTG | chr2R:15834055-15836055 | 1322 | 1336 | + | 9.5643 | 1.06E-05 | 0.4 | GACACGGGCACAGTG |
| 1 | GMTACGGTCACACTG | chr3R:25850291-25852291 | 1329 | 1343 | + | 9.5643 | 1.06E-05 | 0.4 | GAAACTGTCTCACTG |
| 1 | GMTACGGTCACACTG | chr3L:14981835-14983835 | 1331 | 1345 | + | 9.5643 | 1.06E-05 | 0.4 | GCTACGGTCACTTTC |
| 1 | GMTACGGTCACACTG | chr3R:12711299-12713299 | 1334 | 1348 | + | 9.5643 | 1.06E-05 | 0.4 | GCTAAGGTCCCACTA |
| 1 | GMTACGGTCACACTG | chr2R:23720007-23722007 | 1340 | 1354 | + | 9.5643 | 1.06E-05 | 0.4 | GCTACGGTGACGATG |
| 1 | GMTACGGTCACACTG | chr3R:12376128-12378128 | 1341 | 1355 | + | 9.5643 | 1.06E-05 | 0.4 | GATATGGGCACCCTG |
| 1 | GMTACGGTCACACTG | chr3R:12377500-12379500 | 1355 | 1369 | + | 9.5643 | 1.06E-05 | 0.4 | GATATGGGCACCCTG |
| 1 | GMTACGGTCACACTG | chrX:9355227-9357227 | 1359 | 1373 | + | 9.5643 | 1.06E-05 | 0.4 | CCAATGGTCACACTG |
| 1 | GMTACGGTCACACTG | chr3L:3142457-3144457 | 1362 | 1376 | + | 9.5643 | 1.06E-05 | 0.4 | ACCACGGTCACACTT |
| 1 | GMTACGGTCACACTG | chr3L:11727348-11729348 | 1362 | 1376 | + | 9.5643 | 1.06E-05 | 0.4 | GATCCGGTGCCACTG |
| 1 | GMTACGGTCACACTG | chr3L:6034216-6036216 | 1363 | 1377 | + | 9.5643 | 1.06E-05 | 0.4 | ACTACGGTGCCACTG |
| 1 | GMTACGGTCACACTG | chr2L:6657152-6659152 | 1365 | 1379 | + | 9.5643 | 1.06E-05 | 0.4 | CCAACGGTCACACTA |
| 1 | GMTACGGTCACACTG | chr2R:13666736-13668736 | 1378 | 1392 | + | 9.5643 | 1.06E-05 | 0.4 | GATACAGACACCCTG |
| 1 | GMTACGGTCACACTG | chr3R:27238163-27240163 | 1393 | 1407 | + | 9.5643 | 1.06E-05 | 0.4 | GATACGAGCTCACTG |
| 1 | GMTACGGTCACACTG | chr3L:16196219-16198219 | 1394 | 1408 | + | 9.5643 | 1.06E-05 | 0.4 | GATACGGACAAACGG |
| 1 | GMTACGGTCACACTG | chr3R:15149375-15151375 | 1416 | 1430 | + | 9.5643 | 1.06E-05 | 0.4 | GCTACGAGCACACTA |
| 1 | GMTACGGTCACACTG | chr2L:3108672-3110672 | 1417 | 1431 | + | 9.5643 | 1.06E-05 | 0.4 | GAAACGATCACCCTG |
| 1 | GMTACGGTCACACTG | chrX:16941871-16943871 | 1420 | 1434 | + | 9.5643 | 1.06E-05 | 0.4 | CCAACGGTCACACTA |
| 1 | GMTACGGTCACACTG | chr2L:16447469-16449469 | 1421 | 1435 | + | 9.5643 | 1.06E-05 | 0.4 | GCTGGGCTCACACTG |
| 1 | GMTACGGTCACACTG | chr2R:23693003-23695003 | 1451 | 1465 | + | 9.5643 | 1.06E-05 | 0.4 | GATCCGGACATACTG |
| 1 | GMTACGGTCACACTG | chr2R:7586079-7588079 | 1455 | 1469 | + | 9.5643 | 1.06E-05 | 0.4 | GAGACGATCACGCTG |
| 1 | GMTACGGTCACACTG | chr2R:21690139-21692139 | 1457 | 1471 | + | 9.5643 | 1.06E-05 | 0.4 | TCTCTGGTCACACTG |

| | | | | | | | | | |
|---|---|---|---|---|---|---|---|---|---|
| 1 | GMTACGGTCACACTG | chr2R:21690139-21692139 | 1457 | 1471 | + | 9.5643 | 1.06E-05 | 0.4 | TCTCTGGTCACACTG |
| 1 | GMTACGGTCACACTG | chr3R:12711299-12713299 | 1463 | 1477 | + | 9.5643 | 1.06E-05 | 0.4 | GCTAAGGTCCCACTA |
| 1 | GMTACGGTCACACTG | chr3R:27238085-27240085 | 1471 | 1485 | + | 9.5643 | 1.06E-05 | 0.4 | GATACGAGCTCACTG |
| 1 | GMTACGGTCACACTG | chr3R:18365162-18367162 | 1483 | 1497 | + | 9.5643 | 1.06E-05 | 0.4 | GCTCTGGTCACACTT |
| 1 | GMTACGGTCACACTG | chr2R:15537175-15539175 | 1484 | 1498 | + | 9.5643 | 1.06E-05 | 0.4 | GAAGTGGTCACACTG |
| 1 | GMTACGGTCACACTG | chr3L:9428821-9430821 | 1485 | 1499 | + | 9.5643 | 1.06E-05 | 0.4 | CCCATGGTCACACTG |
| 1 | GMTACGGTCACACTG | chr2R:17018522-17020522 | 1486 | 1500 | + | 9.5643 | 1.06E-05 | 0.4 | AAAACGGTCACACAG |
| 1 | GMTACGGTCACACTG | chr3R:6685728-6687728 | 1496 | 1510 | + | 9.5643 | 1.06E-05 | 0.4 | GCTACCGTCGCATTG |
| 1 | GMTACGGTCACACTG | chr3R:25126502-25128502 | 1512 | 1526 | + | 9.5643 | 1.06E-05 | 0.4 | GAGAAGGTCACTCTG |
| 1 | GMTACGGTCACACTG | chr2L:16487148-16489148 | 1512 | 1526 | + | 9.5643 | 1.06E-05 | 0.4 | GCTACGGTCTAACTA |
| 1 | GMTACGGTCACACTG | chr2R:16177307-16179307 | 1521 | 1535 | + | 9.5643 | 1.06E-05 | 0.4 | ACTGCGGTCACACTA |
| 1 | GMTACGGTCACACTG | chr2R:23555672-23557672 | 1530 | 1544 | + | 9.5643 | 1.06E-05 | 0.4 | GATACGGACAAGCTG |
| 1 | GMTACGGTCACACTG | chr3R:13699838-13701838 | 1531 | 1545 | + | 9.5643 | 1.06E-05 | 0.4 | GACACGGAAACACTG |
| 1 | GMTACGGTCACACTG | chr2R:8236271-8238271 | 1532 | 1546 | + | 9.5643 | 1.06E-05 | 0.4 | TCCAGGGTCACACTG |
| 1 | GMTACGGTCACACTG | chr2R:18829331-18831331 | 1534 | 1548 | + | 9.5643 | 1.06E-05 | 0.4 | GCAAAGGTGACACTG |
| 1 | GMTACGGTCACACTG | chr3L:7368260-7370260 | 1577 | 1591 | + | 9.5643 | 1.06E-05 | 0.4 | CATGCGGTCACTCTG |
| 1 | GMTACGGTCACACTG | chr3R:27581241-27583241 | 1592 | 1606 | + | 9.5643 | 1.06E-05 | 0.4 | CAATCGGTCACACTG |
| 1 | GMTACGGTCACACTG | chr2R:21280656-21282656 | 1614 | 1628 | + | 9.5643 | 1.06E-05 | 0.4 | TCTGTGGTCACACTG |
| 1 | GMTACGGTCACACTG | chr2L:1163443-1165443 | 1619 | 1633 | + | 9.5643 | 1.06E-05 | 0.4 | GACAAGGTCTCACTG |
| 1 | GMTACGGTCACACTG | chr3R:7638434-7640434 | 1627 | 1641 | + | 9.5643 | 1.06E-05 | 0.4 | TCTTTGGTCACACTG |
| 1 | GMTACGGTCACACTG | chr2L:990300-992300 | 1631 | 1645 | + | 9.5643 | 1.06E-05 | 0.4 | GATAGGGACACACAG |
| 1 | GMTACGGTCACACTG | chr2L:1107540-1109540 | 1635 | 1649 | + | 9.5643 | 1.06E-05 | 0.4 | GAGACGGTCAAGCTG |
| 1 | GMTACGGTCACACTG | chrX:7377140-7379140 | 1639 | 1653 | + | 9.5643 | 1.06E-05 | 0.4 | GCTACTGTTACCCTG |
| 1 | GMTACGGTCACACTG | chr3R:27282780-27284780 | 1643 | 1657 | + | 9.5643 | 1.06E-05 | 0.4 | GAAACGCTCACGCTG |
| 1 | GMTACGGTCACACTG | chr2R:24474386-24476386 | 1645 | 1659 | + | 9.5643 | 1.06E-05 | 0.4 | GCAAAGGTCACAGTG |
| 1 | GMTACGGTCACACTG | chr3R:25126359-25128359 | 1655 | 1669 | + | 9.5643 | 1.06E-05 | 0.4 | GAGAAGGTCACTCTG |
| 1 | GMTACGGTCACACTG | chr3R:25126359-25128359 | 1655 | 1669 | + | 9.5643 | 1.06E-05 | 0.4 | GAGAAGGTCACTCTG |
| 1 | GMTACGGTCACACTG | chr3L:901898-903898 | 1688 | 1702 | + | 9.5643 | 1.06E-05 | 0.4 | GCTACGATTACACAG |
| 1 | GMTACGGTCACACTG | chrX:20142188-20144188 | 1703 | 1717 | + | 9.5643 | 1.06E-05 | 0.4 | GCCACGCTCACACTT |
| 1 | GMTACGGTCACACTG | chr3R:32014520-32016520 | 1719 | 1733 | + | 9.5643 | 1.06E-05 | 0.4 | GCTGAGGTTACACTG |
| 1 | GMTACGGTCACACTG | chr3R:5482906-5484906 | 1739 | 1753 | + | 9.5643 | 1.06E-05 | 0.4 | GCTATGGTTACACAG |
| 1 | GMTACGGTCACACTG | chr3R:29145893-29147893 | 1743 | 1757 | + | 9.5643 | 1.06E-05 | 0.4 | GATAGGGTCGCACCG |
| 1 | GMTACGGTCACACTG | chr2R:18686372-18688372 | 1746 | 1760 | + | 9.5643 | 1.06E-05 | 0.4 | TACACGGCCACACTG |

| | | | | | | | | | |
|---|---|---|---|---|---|---|---|---|---|
| 1 | GMTACGGTCACACTG | chrX:15088682-15090682 | 1749 | 1763 | + | 9.5643 | 1.06E-05 | 0.4 | GCTGCGGTCACAATA |
| 1 | GMTACGGTCACACTG | chrX:18369118-18371118 | 1757 | 1771 | + | 9.5643 | 1.06E-05 | 0.4 | GCCAGGGTCACACTT |
| 1 | GMTACGGTCACACTG | chr2L:22168143-22170143 | 1787 | 1801 | + | 9.5643 | 1.06E-05 | 0.4 | GCTTCAGTCAAACTG |
| 1 | GMTACGGTCACACTG | chr3L:9428518-9430518 | 1788 | 1802 | + | 9.5643 | 1.06E-05 | 0.4 | CCCATGGTCACACTG |
| 1 | GMTACGGTCACACTG | chr2L:1107373-1109373 | 1802 | 1816 | + | 9.5643 | 1.06E-05 | 0.4 | GAGACGGTCAAGCTG |
| 1 | GMTACGGTCACACTG | chr2L:10849141-10851141 | 1803 | 1817 | + | 9.5643 | 1.06E-05 | 0.4 | GCAATGGTCACACTT |
| 1 | GMTACGGTCACACTG | chr3R:13699565-13701565 | 1804 | 1818 | + | 9.5643 | 1.06E-05 | 0.4 | GACACGGAAACACTG |
| 1 | GMTACGGTCACACTG | chrX:15586733-15588733 | 1821 | 1835 | + | 9.5643 | 1.06E-05 | 0.4 | GCTACCGGTACACTG |
| 1 | GMTACGGTCACACTG | chr2R:7971175-7973175 | 1825 | 1839 | + | 9.5643 | 1.06E-05 | 0.4 | GATGCGGCCACAGTG |
| 1 | GMTACGGTCACACTG | chr2L:12007153-12009153 | 1838 | 1852 | + | 9.5643 | 1.06E-05 | 0.4 | GCTACGGACAGATTG |
| 1 | GMTACGGTCACACTG | chr3L:22272985-22274985 | 1841 | 1855 | + | 9.5643 | 1.06E-05 | 0.4 | GATGCGGTCTAACTG |
| 1 | GMTACGGTCACACTG | chr3R:6284200-6286200 | 1842 | 1856 | + | 9.5643 | 1.06E-05 | 0.4 | GCTACAGTCACTCTC |
| 1 | GMTACGGTCACACTG | chr2R:7405837-7407837 | 1846 | 1860 | + | 9.5643 | 1.06E-05 | 0.4 | GATTCGAGCACACTG |
| 1 | GMTACGGTCACACTG | chr3R:28339800-28341800 | 1854 | 1868 | + | 9.5643 | 1.06E-05 | 0.4 | GCTAGCGTCCCACTG |
| 1 | GMTACGGTCACACTG | chr2R:5980844-5982844 | 1874 | 1888 | + | 9.5643 | 1.06E-05 | 0.4 | GCTACGTTCACACGC |
| 1 | GMTACGGTCACACTG | chr3R:11430738-11432738 | 1893 | 1907 | + | 9.5643 | 1.06E-05 | 0.4 | GAGACGGACACAGTG |
| 1 | GMTACGGTCACACTG | chr3L:3034196-3036196 | 1902 | 1916 | + | 9.5643 | 1.06E-05 | 0.4 | GATTCGGTCCCATTG |
| 1 | GMTACGGTCACACTG | chrX:946500-948500 | 1917 | 1931 | + | 9.5643 | 1.06E-05 | 0.4 | GACGAGGTCACACTG |
| 1 | GMTACGGTCACACTG | chr2L:9937134-9939134 | 1932 | 1946 | + | 9.5643 | 1.06E-05 | 0.4 | GAGACCGGCACACTG |
| 1 | GMTACGGTCACACTG | chrX:14192214-14194214 | 1943 | 1957 | + | 9.5643 | 1.06E-05 | 0.4 | GATTCGGTGACACCG |
| 1 | GMTACGGTCACACTG | chr2R:21309146-21311146 | 1950 | 1964 | + | 9.5643 | 1.06E-05 | 0.4 | GCTACGGCCAGACTA |
| 1 | GMTACGGTCACACTG | chr3R:24849627-24851627 | 1967 | 1981 | + | 9.5643 | 1.06E-05 | 0.4 | GAAACGGTCACACGC |
| 1 | GMTACGGTCACACTG | chr3L:6033611-6035611 | 1968 | 1982 | + | 9.5643 | 1.06E-05 | 0.4 | ACTACGGTGCCACTG |
| 1 | GMTACGGTCACACTG | chr3R:18544586-18546586 | 1982 | 1996 | + | 9.5643 | 1.06E-05 | 0.4 | GATACGGCCTCATTG |
| 1 | GMTACGGTCACACTG | chr2R:23555217-23557217 | 1985 | 1999 | + | 9.5643 | 1.06E-05 | 0.4 | GATACGGACAAGCTG |
| 1 | GMTACGGTCACACTG | chr2R:16347527-16349527 | 8 | 22 | - | 9.2857 | 1.62E-05 | 0.515 | GATGTGGTCACACTT |
| 1 | GMTACGGTCACACTG | chr2L:22104512-22106512 | 27 | 41 | - | 9.2857 | 1.62E-05 | 0.515 | TATACGGTCAGACGG |
| 1 | GMTACGGTCACACTG | chr2R:23701477-23703477 | 31 | 45 | - | 9.2857 | 1.62E-05 | 0.515 | GATACGGTCAGAGGG |
| 1 | GMTACGGTCACACTG | chr3R:28339800-28341800 | 63 | 77 | - | 9.2857 | 1.62E-05 | 0.515 | AATACGGTCACACAA |
| 1 | GMTACGGTCACACTG | chr3R:18222722-18224722 | 81 | 95 | - | 9.2857 | 1.62E-05 | 0.515 | GAGACCCTCACACTG |
| 1 | GMTACGGTCACACTG | chr2R:10121574-10123574 | 98 | 112 | - | 9.2857 | 1.62E-05 | 0.515 | GATGGGGTCACACTT |
| 1 | GMTACGGTCACACTG | chr2R:23701406-23703406 | 102 | 116 | - | 9.2857 | 1.62E-05 | 0.515 | GATACGGTCAGAGGG |
| 1 | GMTACGGTCACACTG | chr3R:22463985-22465985 | 121 | 135 | - | 9.2857 | 1.62E-05 | 0.515 | GATGCGGTAACACTT |

| | | | | | | | | |
|---|---|---|---|---|---|---|---|---|
| 1 | GMTACGGTCACACTG | chr3R:18222682-18224682 | 121 | 135 | - | 9.2857 | 1.62E-05 | 0.515 GAGACCCTCACACTG |
| 1 | GMTACGGTCACACTG | chr2R:7811028-7813028 | 155 | 169 | - | 9.2857 | 1.62E-05 | 0.515 GATACTGTTCCACTG |
| 1 | GMTACGGTCACACTG | chr2R:5936400-5938400 | 160 | 174 | - | 9.2857 | 1.62E-05 | 0.515 GAAACGGTTACATTG |
| 1 | GMTACGGTCACACTG | chrX:646304-648304 | 171 | 185 | - | 9.2857 | 1.62E-05 | 0.515 CATACGGTCAGACGG |
| 1 | GMTACGGTCACACTG | chr3L:260953-262953 | 195 | 209 | - | 9.2857 | 1.62E-05 | 0.515 GATAACGCCACACTG |
| 1 | GMTACGGTCACACTG | chr3R:12673608-12675608 | 204 | 218 | - | 9.2857 | 1.62E-05 | 0.515 GATAAAGTCTCACTG |
| 1 | GMTACGGTCACACTG | chr3L:679326-681326 | 222 | 236 | - | 9.2857 | 1.62E-05 | 0.515 GATATGGAAACACTG |
| 1 | GMTACGGTCACACTG | chr3L:13841240-13843240 | 255 | 269 | - | 9.2857 | 1.62E-05 | 0.515 GATACGGTCATCTTG |
| 1 | GMTACGGTCACACTG | chr2R:17766786-17768786 | 281 | 295 | - | 9.2857 | 1.62E-05 | 0.515 CATACGGTCACTCTT |
| 1 | GMTACGGTCACACTG | chr3R:12385249-12387249 | 299 | 313 | - | 9.2857 | 1.62E-05 | 0.515 GCTACTGTAACACTA |
| 1 | GMTACGGTCACACTG | chrX:21338741-21340741 | 332 | 346 | - | 9.2857 | 1.62E-05 | 0.515 AAAACGGTCACATTG |
| 1 | GMTACGGTCACACTG | chr3R:28025615-28027615 | 332 | 346 | - | 9.2857 | 1.62E-05 | 0.515 CATACGGACACACTC |
| 1 | GMTACGGTCACACTG | chr3R:31736035-31738035 | 347 | 361 | - | 9.2857 | 1.62E-05 | 0.515 TCTATGATCACACTG |
| 1 | GMTACGGTCACACTG | chr2L:15008939-15010939 | 378 | 392 | - | 9.2857 | 1.62E-05 | 0.515 CCTACGGTCAAAGTG |
| 1 | GMTACGGTCACACTG | chr2L:5027191-5029191 | 378 | 392 | - | 9.2857 | 1.62E-05 | 0.515 ACTACACTCACACTG |
| 1 | GMTACGGTCACACTG | chr2L:869465-871465 | 396 | 410 | - | 9.2857 | 1.62E-05 | 0.515 GATATGGTCACACGC |
| 1 | GMTACGGTCACACTG | chrX:21338676-21340676 | 397 | 411 | - | 9.2857 | 1.62E-05 | 0.515 AAAACGGTCACATTG |
| 1 | GMTACGGTCACACTG | chr2L:7780465-7782465 | 403 | 417 | - | 9.2857 | 1.62E-05 | 0.515 GATGCGATCATACTG |
| 1 | GMTACGGTCACACTG | chr2R:5662909-5664909 | 405 | 419 | - | 9.2857 | 1.62E-05 | 0.515 GCTACGTTAACACTT |
| 1 | GMTACGGTCACACTG | chr3R:31735963-31737963 | 419 | 433 | - | 9.2857 | 1.62E-05 | 0.515 TCTATGATCACACTG |
| 1 | GMTACGGTCACACTG | chr3R:15371212-15373212 | 422 | 436 | - | 9.2857 | 1.62E-05 | 0.515 GAAACGCTCACAATG |
| 1 | GMTACGGTCACACTG | chr2L:18158272-18160272 | 425 | 439 | - | 9.2857 | 1.62E-05 | 0.515 TAAACGGTCACATTG |
| 1 | GMTACGGTCACACTG | chr3L:16050034-16052034 | 439 | 453 | - | 9.2857 | 1.62E-05 | 0.515 GATACGTTATCACTG |
| 1 | GMTACGGTCACACTG | chr2R:12635671-12637671 | 457 | 471 | - | 9.2857 | 1.62E-05 | 0.515 AATACGGTCACACAA |
| 1 | GMTACGGTCACACTG | chr3L:9713324-9715324 | 465 | 479 | - | 9.2857 | 1.62E-05 | 0.515 GAAACGGTAACACTT |
| 1 | GMTACGGTCACACTG | chr2L:6350034-6352034 | 476 | 490 | - | 9.2857 | 1.62E-05 | 0.515 GATACTATCACTCTG |
| 1 | GMTACGGTCACACTG | chr2R:8018176-8020176 | 490 | 504 | - | 9.2857 | 1.62E-05 | 0.515 GATACGGTTACAAGG |
| 1 | GMTACGGTCACACTG | chr3R:28339334-28341334 | 529 | 543 | - | 9.2857 | 1.62E-05 | 0.515 AATACGGTCACACAA |
| 1 | GMTACGGTCACACTG | chr2L:10857536-10859536 | 550 | 564 | - | 9.2857 | 1.62E-05 | 0.515 CATGCGGTCACACTA |
| 1 | GMTACGGTCACACTG | chr2L:12171981-12173981 | 575 | 589 | - | 9.2857 | 1.62E-05 | 0.515 CATATGGTCACACAG |
| 1 | GMTACGGTCACACTG | chr3R:14897739-14899739 | 592 | 606 | - | 9.2857 | 1.62E-05 | 0.515 CATCTGGTCACACTG |
| 1 | GMTACGGTCACACTG | chr2L:15008724-15010724 | 593 | 607 | - | 9.2857 | 1.62E-05 | 0.515 CCTACGGTCAAAGTG |
| 1 | GMTACGGTCACACTG | chr3R:5793705-5795705 | 598 | 612 | - | 9.2857 | 1.62E-05 | 0.515 CACATGGTCACACTG |

| | | | | | | | | | |
|---|---|---|---|---|---|---|---|---|---|
| 1 | GMTACGGTCACACTG | chr2L:10986806-10988806 | 607 | 621 | - | 9.2857 | 1.62E-05 | 0.515 | CATTTGGTCACACTG |
| 1 | GMTACGGTCACACTG | chr3R:13766549-13768549 | 609 | 623 | - | 9.2857 | 1.62E-05 | 0.515 | GATATGGTTACGCTG |
| 1 | GMTACGGTCACACTG | chr3R:29806127-29808127 | 634 | 648 | - | 9.2857 | 1.62E-05 | 0.515 | CAAATGGTCACACTG |
| 1 | GMTACGGTCACACTG | chr2R:22074308-22076308 | 649 | 663 | - | 9.2857 | 1.62E-05 | 0.515 | GATACGCTGACACGG |
| 1 | GMTACGGTCACACTG | chr3L:16711126-16713126 | 661 | 675 | - | 9.2857 | 1.62E-05 | 0.515 | TAGATGGTCACACTG |
| 1 | GMTACGGTCACACTG | chr3R:13381106-13383106 | 668 | 682 | - | 9.2857 | 1.62E-05 | 0.515 | TAAATGGTCACACTG |
| 1 | GMTACGGTCACACTG | chr3L:16289950-16291950 | 674 | 688 | - | 9.2857 | 1.62E-05 | 0.515 | GATACCGTCACCGTG |
| 1 | GMTACGGTCACACTG | chrX:16883581-16885581 | 676 | 690 | - | 9.2857 | 1.62E-05 | 0.515 | GATGCGGTGATACTG |
| 1 | GMTACGGTCACACTG | chr3R:15370957-15372957 | 677 | 691 | - | 9.2857 | 1.62E-05 | 0.515 | GAAACGCTCACAATG |
| 1 | GMTACGGTCACACTG | chr2L:3844716-3846716 | 678 | 692 | - | 9.2857 | 1.62E-05 | 0.515 | GATACCCTCACACAG |
| 1 | GMTACGGTCACACTG | chrX:11492135-11494135 | 689 | 703 | - | 9.2857 | 1.62E-05 | 0.515 | CACATGGTCACACTG |
| 1 | GMTACGGTCACACTG | chr3R:14325947-14327947 | 704 | 718 | - | 9.2857 | 1.62E-05 | 0.515 | TAAACGTTCACACTG |
| 1 | GMTACGGTCACACTG | chr3R:18198857-18200857 | 714 | 728 | - | 9.2857 | 1.62E-05 | 0.515 | CAAACGGTCACACTC |
| 1 | GMTACGGTCACACTG | chr2L:22129142-22131142 | 743 | 757 | - | 9.2857 | 1.62E-05 | 0.515 | GACATGGTAACACTG |
| 1 | GMTACGGTCACACTG | chr3L:7322619-7324619 | 771 | 785 | - | 9.2857 | 1.62E-05 | 0.515 | CAAACGGTAACACTG |
| 1 | GMTACGGTCACACTG | chrX:6364949-6366949 | 773 | 787 | - | 9.2857 | 1.62E-05 | 0.515 | GAAATGGTCACACTA |
| 1 | GMTACGGTCACACTG | chr2R:23808236-23810236 | 775 | 789 | - | 9.2857 | 1.62E-05 | 0.515 | CATACGGTCACTCTA |
| 1 | GMTACGGTCACACTG | chr3L:539851-541851 | 820 | 834 | - | 9.2857 | 1.62E-05 | 0.515 | GAAATGGTCACACTC |
| 1 | GMTACGGTCACACTG | chrX:11491998-11493998 | 826 | 840 | - | 9.2857 | 1.62E-05 | 0.515 | CACATGGTCACACTG |
| 1 | GMTACGGTCACACTG | chrX:15220176-15222176 | 828 | 842 | - | 9.2857 | 1.62E-05 | 0.515 | AACACGCTCACACTG |
| 1 | GMTACGGTCACACTG | chr2L:1176690-1178690 | 838 | 852 | - | 9.2857 | 1.62E-05 | 0.515 | TAAACGGTCACACTT |
| 1 | GMTACGGTCACACTG | chr3R:29062040-29064040 | 848 | 862 | - | 9.2857 | 1.62E-05 | 0.515 | GCTACGATCAAAATG |
| 1 | GMTACGGTCACACTG | chr2L:8402924-8404924 | 883 | 897 | - | 9.2857 | 1.62E-05 | 0.515 | CATACGCACACACTG |
| 1 | GMTACGGTCACACTG | chr3R:23009562-23011562 | 887 | 901 | - | 9.2857 | 1.62E-05 | 0.515 | GATACGATCACAAAG |
| 1 | GMTACGGTCACACTG | chrX:16883365-16885365 | 892 | 906 | - | 9.2857 | 1.62E-05 | 0.515 | GATGCGGTGATACTG |
| 1 | GMTACGGTCACACTG | chr2R:22836361-22838361 | 898 | 912 | - | 9.2857 | 1.62E-05 | 0.515 | GATAAGGTCATACAG |
| 1 | GMTACGGTCACACTG | chr3L:1664138-1666138 | 902 | 916 | - | 9.2857 | 1.62E-05 | 0.515 | CATCTGGTCACACTG |
| 1 | GMTACGGTCACACTG | chr3R:22210824-22212824 | 908 | 922 | - | 9.2857 | 1.62E-05 | 0.515 | TCTAATGTCACACTG |
| 1 | GMTACGGTCACACTG | chr3R:29061975-29063975 | 913 | 927 | - | 9.2857 | 1.62E-05 | 0.515 | GCTACGATCAAAATG |
| 1 | GMTACGGTCACACTG | chrX:12028032-12030032 | 920 | 934 | - | 9.2857 | 1.62E-05 | 0.515 | GATACGTTAACGCTG |
| 1 | GMTACGGTCACACTG | chr2R:10882547-10884547 | 927 | 941 | - | 9.2857 | 1.62E-05 | 0.515 | GATAAGCACACACTG |
| 1 | GMTACGGTCACACTG | chrX:23095771-23097771 | 937 | 951 | - | 9.2857 | 1.62E-05 | 0.515 | TATGCGGTCACATTG |
| 1 | GMTACGGTCACACTG | chrX:15577205-15579205 | 942 | 956 | - | 9.2857 | 1.62E-05 | 0.515 | GATATGGTCACTCTA |

| | | | | | | | | | |
|---|---|---|---|---|---|---|---|---|---|
| 1 | GMTACGGTCACACTG | chr3R:29805811-29807811 | 950 | 964 | - | 9.2857 | 1.62E-05 | 0.515 | CAAATGGTCACACTG |
| 1 | GMTACGGTCACACTG | chrX:16285288-16287288 | 950 | 964 | - | 9.2857 | 1.62E-05 | 0.515 | AATACAGACACACTG |
| 1 | GMTACGGTCACACTG | chr2R:8626900-8628900 | 951 | 965 | - | 9.2857 | 1.62E-05 | 0.515 | GATACGGTCTTACTT |
| 1 | GMTACGGTCACACTG | chr3R:22744620-22746620 | 954 | 968 | - | 9.2857 | 1.62E-05 | 0.515 | AAAACGGTCACACTC |
| 1 | GMTACGGTCACACTG | chr3L:19927743-19929743 | 957 | 971 | - | 9.2857 | 1.62E-05 | 0.515 | CATCCGTTCACACTG |
| 1 | GMTACGGTCACACTG | chr3R:14897372-14899372 | 959 | 973 | - | 9.2857 | 1.62E-05 | 0.515 | CATCTGGTCACACTG |
| 1 | GMTACGGTCACACTG | chr3L:3238578-3240578 | 961 | 975 | - | 9.2857 | 1.62E-05 | 0.515 | GACACGTTCACACTT |
| 1 | GMTACGGTCACACTG | chr3R:14325689-14327689 | 962 | 976 | - | 9.2857 | 1.62E-05 | 0.515 | TAAACGTTCACACTG |
| 1 | GMTACGGTCACACTG | chr3L:9712827-9714827 | 962 | 976 | - | 9.2857 | 1.62E-05 | 0.515 | GAAACGGTAACACTT |
| 1 | GMTACGGTCACACTG | chr2R:21684283-21686283 | 962 | 976 | - | 9.2857 | 1.62E-05 | 0.515 | AATGCGGTCACATTG |
| 1 | GMTACGGTCACACTG | chr3R:18298229-18300229 | 963 | 977 | - | 9.2857 | 1.62E-05 | 0.515 | GATCTGGTCAGACTG |
| 1 | GMTACGGTCACACTG | chr3R:13380809-13382809 | 965 | 979 | - | 9.2857 | 1.62E-05 | 0.515 | TAAATGGTCACACTG |
| 1 | GMTACGGTCACACTG | chr3L:16010812-16012812 | 968 | 982 | - | 9.2857 | 1.62E-05 | 0.515 | ACTACGGTCACATTC |
| 1 | GMTACGGTCACACTG | chr2L:12171586-12173586 | 970 | 984 | - | 9.2857 | 1.62E-05 | 0.515 | CATATGGTCACACAG |
| 1 | GMTACGGTCACACTG | chr3R:25328316-25330316 | 973 | 987 | - | 9.2857 | 1.62E-05 | 0.515 | AAGACCGTCACACTG |
| 1 | GMTACGGTCACACTG | chrX:23095735-23097735 | 973 | 987 | - | 9.2857 | 1.62E-05 | 0.515 | TATGCGGTCACATTG |
| 1 | GMTACGGTCACACTG | chr3L:1495643-1497643 | 973 | 987 | - | 9.2857 | 1.62E-05 | 0.515 | CATCTGGTCACACTG |
| 1 | GMTACGGTCACACTG | chrX:1926683-1928683 | 974 | 988 | - | 9.2857 | 1.62E-05 | 0.515 | GATCTGGTTACACTG |
| 1 | GMTACGGTCACACTG | chr3R:13433984-13435984 | 974 | 988 | - | 9.2857 | 1.62E-05 | 0.515 | CATCTGGTCACACTG |
| 1 | GMTACGGTCACACTG | chr3R:18198595-18200595 | 976 | 990 | - | 9.2857 | 1.62E-05 | 0.515 | CAAACGGTCACACTC |
| 1 | GMTACGGTCACACTG | chrX:15577171-15579171 | 976 | 990 | - | 9.2857 | 1.62E-05 | 0.515 | GATATGGTCACTCTA |
| 1 | GMTACGGTCACACTG | chr3L:1664064-1666064 | 976 | 990 | - | 9.2857 | 1.62E-05 | 0.515 | CATCTGGTCACACTG |
| 1 | GMTACGGTCACACTG | chr3L:7322411-7324411 | 979 | 993 | - | 9.2857 | 1.62E-05 | 0.515 | CAAACGGTAACACTG |
| 1 | GMTACGGTCACACTG | chr2R:24047903-24049903 | 981 | 995 | - | 9.2857 | 1.62E-05 | 0.515 | CATTCGGTCACACTA |
| 1 | GMTACGGTCACACTG | chr3L:5607300-5609300 | 982 | 996 | - | 9.2857 | 1.62E-05 | 0.515 | TATCCGGTCACATTG |
| 1 | GMTACGGTCACACTG | chr3R:8268896-8270896 | 984 | 998 | - | 9.2857 | 1.62E-05 | 0.515 | GAAATGGTCACACTA |
| 1 | GMTACGGTCACACTG | chr3R:25234529-25236529 | 986 | 1000 | - | 9.2857 | 1.62E-05 | 0.515 | CCTACGGTCATACTA |
| 1 | GMTACGGTCACACTG | chr3R:5793317-5795317 | 986 | 1000 | - | 9.2857 | 1.62E-05 | 0.515 | CACATGGTCACACTG |
| 1 | GMTACGGTCACACTG | chr2R:12635141-12637141 | 987 | 1001 | - | 9.2857 | 1.62E-05 | 0.515 | AATACGGTCACACAA |
| 1 | GMTACGGTCACACTG | chr2R:12635141-12637141 | 987 | 1001 | - | 9.2857 | 1.62E-05 | 0.515 | AATACGGTCACACAA |
| 1 | GMTACGGTCACACTG | chr3L:1312696-1314696 | 987 | 1001 | - | 9.2857 | 1.62E-05 | 0.515 | GATAGGGCCACATTG |
| 1 | GMTACGGTCACACTG | chr2L:10857094-10859094 | 992 | 1006 | - | 9.2857 | 1.62E-05 | 0.515 | CATGCGGTCACACTA |
| 1 | GMTACGGTCACACTG | chr3R:22744581-22746581 | 993 | 1007 | - | 9.2857 | 1.62E-05 | 0.515 | AAAACGGTCACACTC |

| | | | | | | | | | |
|---|---|---|---|---|---|---|---|---|---|
| 1 | GMTACGGTCACACTG | chr2L:10986420-10988420 | 993 | 1007 | - | 9.2857 | 1.62E-05 | 0.515 | CATTTGGTCACACTG |
| 1 | GMTACGGTCACACTG | chrX:19578800-19580800 | 994 | 1008 | - | 9.2857 | 1.62E-05 | 0.515 | GATCTGGTAACACTG |
| 1 | GMTACGGTCACACTG | chr2L:20462444-20464444 | 995 | 1009 | - | 9.2857 | 1.62E-05 | 0.515 | GATCTGGTAACACTG |
| 1 | GMTACGGTCACACTG | chr3R:11219064-11221064 | 995 | 1009 | - | 9.2857 | 1.62E-05 | 0.515 | CATTTGGTCACACTG |
| 1 | GMTACGGTCACACTG | chr2R:17766070-17768070 | 997 | 1011 | - | 9.2857 | 1.62E-05 | 0.515 | CATACGGTCACTCTT |
| 1 | GMTACGGTCACACTG | chr2L:19180702-19182702 | 1001 | 1015 | - | 9.2857 | 1.62E-05 | 0.515 | TATCTGGTCACACTG |
| 1 | GMTACGGTCACACTG | chr2R:15531259-15533259 | 1002 | 1016 | - | 9.2857 | 1.62E-05 | 0.515 | CACACGGTCACACTC |
| 1 | GMTACGGTCACACTG | chr2L:18694664-18696664 | 1003 | 1017 | - | 9.2857 | 1.62E-05 | 0.515 | GATTCGGTCATACTA |
| 1 | GMTACGGTCACACTG | chr3L:14001127-14003127 | 1004 | 1018 | - | 9.2857 | 1.62E-05 | 0.515 | AAGACGGTCACACTA |
| 1 | GMTACGGTCACACTG | chr2L:868857-870857 | 1004 | 1018 | - | 9.2857 | 1.62E-05 | 0.515 | GATATGGTCACACGC |
| 1 | GMTACGGTCACACTG | chr2L:12691993-12693993 | 1006 | 1020 | - | 9.2857 | 1.62E-05 | 0.515 | AATACGGTCACACCA |
| 1 | GMTACGGTCACACTG | chr2R:23808004-23810004 | 1007 | 1021 | - | 9.2857 | 1.62E-05 | 0.515 | CATACGGTCACTCTA |
| 1 | GMTACGGTCACACTG | chrX:6364708-6366708 | 1014 | 1028 | - | 9.2857 | 1.62E-05 | 0.515 | GAAATGGTCACACTA |
| 1 | GMTACGGTCACACTG | chr3L:20518393-20520393 | 1026 | 1040 | - | 9.2857 | 1.62E-05 | 0.515 | CATCTGGTCACACTG |
| 1 | GMTACGGTCACACTG | chr3R:24914131-24916131 | 1031 | 1045 | - | 9.2857 | 1.62E-05 | 0.515 | GATACGGAAACACTT |
| 1 | GMTACGGTCACACTG | chr2R:16346500-16348500 | 1035 | 1049 | - | 9.2857 | 1.62E-05 | 0.515 | GATGTGGTCACACTT |
| 1 | GMTACGGTCACACTG | chr3L:11832578-11834578 | 1040 | 1054 | - | 9.2857 | 1.62E-05 | 0.515 | TATATGGCCACACTG |
| 1 | GMTACGGTCACACTG | chr3L:8412012-8414012 | 1040 | 1054 | - | 9.2857 | 1.62E-05 | 0.515 | GATACTGTGACACAG |
| 1 | GMTACGGTCACACTG | chr2L:6466698-6468698 | 1048 | 1062 | - | 9.2857 | 1.62E-05 | 0.515 | TCTACGGTAACACTT |
| 1 | GMTACGGTCACACTG | chr2R:22200508-22202508 | 1051 | 1065 | - | 9.2857 | 1.62E-05 | 0.515 | GATTCGGTGAGACTG |
| 1 | GMTACGGTCACACTG | chr3R:16025079-16027079 | 1052 | 1066 | - | 9.2857 | 1.62E-05 | 0.515 | TATACGGTCACACAC |
| 1 | GMTACGGTCACACTG | chr3L:539614-541614 | 1057 | 1071 | - | 9.2857 | 1.62E-05 | 0.515 | GAAATGGTCACACTC |
| 1 | GMTACGGTCACACTG | chr2L:18694602-18696602 | 1065 | 1079 | - | 9.2857 | 1.62E-05 | 0.515 | GATTCGGTCATACTA |
| 1 | GMTACGGTCACACTG | chr2L:12691934-12693934 | 1065 | 1079 | - | 9.2857 | 1.62E-05 | 0.515 | AATACGGTCACACCA |
| 1 | GMTACGGTCACACTG | chr2R:5662222-5664222 | 1092 | 1106 | - | 9.2857 | 1.62E-05 | 0.515 | GCTACGTTAACACTT |
| 1 | GMTACGGTCACACTG | chr3R:28338765-28340765 | 1098 | 1112 | - | 9.2857 | 1.62E-05 | 0.515 | AATACGGTCACACAA |
| 1 | GMTACGGTCACACTG | chr3L:16710677-16712677 | 1110 | 1124 | - | 9.2857 | 1.62E-05 | 0.515 | TAGATGGTCACACTG |
| 1 | GMTACGGTCACACTG | chr3L:16710677-16712677 | 1110 | 1124 | - | 9.2857 | 1.62E-05 | 0.515 | TAGATGGTCACACTG |
| 1 | GMTACGGTCACACTG | chr3L:13839611-13841611 | 1121 | 1135 | - | 9.2857 | 1.62E-05 | 0.515 | GATATGGGCAAACTG |
| 1 | GMTACGGTCACACTG | chr3L:21475479-21477479 | 1123 | 1137 | - | 9.2857 | 1.62E-05 | 0.515 | GATACGGATACATTG |
| 1 | GMTACGGTCACACTG | chr2L:22103401-22105401 | 1138 | 1152 | - | 9.2857 | 1.62E-05 | 0.515 | TATACGGTCAGACGG |
| 1 | GMTACGGTCACACTG | chr3L:19927508-19929508 | 1192 | 1206 | - | 9.2857 | 1.62E-05 | 0.515 | CATCCGTTCACACTG |
| 1 | GMTACGGTCACACTG | chr2R:17765865-17767865 | 1202 | 1216 | - | 9.2857 | 1.62E-05 | 0.515 | CATACGGTCACTCTT |

| | | | | | | | | | |
|---|---|---|---|---|---|---|---|---|---|
| 1 | GMTACGGTCACACTG | chr3L:8411825-8413825 | 1227 | 1241 | - | 9.2857 | 1.62E-05 | 0.515 | GATACTGTGACACAG |
| 1 | GMTACGGTCACACTG | chr2R:19304172-19306172 | 1227 | 1241 | - | 9.2857 | 1.62E-05 | 0.515 | GATACGGGAACACTT |
| 1 | GMTACGGTCACACTG | chr2L:8402573-8404573 | 1234 | 1248 | - | 9.2857 | 1.62E-05 | 0.515 | CATACGCACACACTG |
| 1 | GMTACGGTCACACTG | chr2R:8626611-8628611 | 1240 | 1254 | - | 9.2857 | 1.62E-05 | 0.515 | GATACGGTCTTACTT |
| 1 | GMTACGGTCACACTG | chr3R:11209247-11211247 | 1245 | 1259 | - | 9.2857 | 1.62E-05 | 0.515 | GATATCGTCACCCTG |
| 1 | GMTACGGTCACACTG | chr3R:9236682-9238682 | 1250 | 1264 | - | 9.2857 | 1.62E-05 | 0.515 | TATATGGTCACTCTG |
| 1 | GMTACGGTCACACTG | chr2R:21250150-21252150 | 1250 | 1264 | - | 9.2857 | 1.62E-05 | 0.515 | GATAACGACACACTG |
| 1 | GMTACGGTCACACTG | chr3L:14000880-14002880 | 1251 | 1265 | - | 9.2857 | 1.62E-05 | 0.515 | AAGACGGTCACACTA |
| 1 | GMTACGGTCACACTG | chrX:22489872-22491872 | 1281 | 1295 | - | 9.2857 | 1.62E-05 | 0.515 | AATACAGTCACACAG |
| 1 | GMTACGGTCACACTG | chr3L:1312377-1314377 | 1306 | 1320 | - | 9.2857 | 1.62E-05 | 0.515 | GATAGGGCCACATTG |
| 1 | GMTACGGTCACACTG | chr3R:18297879-18299879 | 1313 | 1327 | - | 9.2857 | 1.62E-05 | 0.515 | GATCTGGTCAGACTG |
| 1 | GMTACGGTCACACTG | chr3R:24913834-24915834 | 1328 | 1342 | - | 9.2857 | 1.62E-05 | 0.515 | GATACGGAAACACTT |
| 1 | GMTACGGTCACACTG | chr3L:19860458-19862458 | 1359 | 1373 | - | 9.2857 | 1.62E-05 | 0.515 | GATACTGTCCTACTG |
| 1 | GMTACGGTCACACTG | chr3L:8914233-8916233 | 1391 | 1405 | - | 9.2857 | 1.62E-05 | 0.515 | GATACAGATACACTG |
| 1 | GMTACGGTCACACTG | chr2L:6853142-6855142 | 1479 | 1493 | - | 9.2857 | 1.62E-05 | 0.515 | GCTACGGTTATACTC |
| 1 | GMTACGGTCACACTG | chrX:2065359-2067359 | 1500 | 1514 | - | 9.2857 | 1.62E-05 | 0.515 | GATACGGTGACAGCG |
| 1 | GMTACGGTCACACTG | chr2R:22316662-22318662 | 1521 | 1535 | - | 9.2857 | 1.62E-05 | 0.515 | GATACGCTCACACAC |
| 1 | GMTACGGTCACACTG | chr2L:8402280-8404280 | 1527 | 1541 | - | 9.2857 | 1.62E-05 | 0.515 | CATACGCACACACTG |
| 1 | GMTACGGTCACACTG | chr3L:11619101-11621101 | 1548 | 1562 | - | 9.2857 | 1.62E-05 | 0.515 | AATACGGTCACCCTC |
| 1 | GMTACGGTCACACTG | chrX:1860790-1862790 | 1556 | 1570 | - | 9.2857 | 1.62E-05 | 0.515 | GAAACGCTGACACTG |
| 1 | GMTACGGTCACACTG | chr2R:19443179-19445179 | 1564 | 1578 | - | 9.2857 | 1.62E-05 | 0.515 | GCTATTTTCACACTG |
| 1 | GMTACGGTCACACTG | chrX:15496862-15498862 | 1605 | 1619 | - | 9.2857 | 1.62E-05 | 0.515 | ACTATGGTCATACTG |
| 1 | GMTACGGTCACACTG | chr2R:17765460-17767460 | 1607 | 1621 | - | 9.2857 | 1.62E-05 | 0.515 | CATACGGTCACTCTT |
| 1 | GMTACGGTCACACTG | chrX:1482479-1484479 | 1618 | 1632 | - | 9.2857 | 1.62E-05 | 0.515 | CACACAGTCACACTG |
| 1 | GMTACGGTCACACTG | chr3R:7085599-7087599 | 1632 | 1646 | - | 9.2857 | 1.62E-05 | 0.515 | GATACATTCACGCTG |
| 1 | GMTACGGTCACACTG | chr3R:11208849-11210849 | 1643 | 1657 | - | 9.2857 | 1.62E-05 | 0.515 | GATATCGTCACCCTG |
| 1 | GMTACGGTCACACTG | chrX:1482413-1484413 | 1684 | 1698 | - | 9.2857 | 1.62E-05 | 0.515 | CACACAGTCACACTG |
| 1 | GMTACGGTCACACTG | chr3R:24391889-24393889 | 1704 | 1718 | - | 9.2857 | 1.62E-05 | 0.515 | GATTCGATCATACTG |
| 1 | GMTACGGTCACACTG | chr2L:7779085-7781085 | 1783 | 1797 | - | 9.2857 | 1.62E-05 | 0.515 | GATGCGATCATACTG |
| 1 | GMTACGGTCACACTG | chr3L:11937872-11939872 | 1785 | 1799 | - | 9.2857 | 1.62E-05 | 0.515 | GAAATGGTCACACTA |
| 1 | GMTACGGTCACACTG | chr3R:24140395-24142395 | 1796 | 1810 | - | 9.2857 | 1.62E-05 | 0.515 | GCTATGGTGACACTT |
| 1 | GMTACGGTCACACTG | chr3L:8752619-8754619 | 1797 | 1811 | - | 9.2857 | 1.62E-05 | 0.515 | AATTCGGTCACAATG |
| 1 | GMTACGGTCACACTG | chr3R:17732598-17734598 | 1818 | 1832 | - | 9.2857 | 1.62E-05 | 0.515 | GAAACAGTCACACTC |

| | | | | | | | | | |
|---|---|---|---|---|---|---|---|---|---|
| 1 | GMTACGGTCACACTG | chr3L:259280-261280 | 1868 | 1882 | - | 9.2857 | 1.62E-05 | 0.515 | GATAACGCCACACTG |
| 1 | GMTACGGTCACACTG | chr3L:13839611-13841611 | 1884 | 1898 | - | 9.2857 | 1.62E-05 | 0.515 | GATACGGTCATCTTG |
| 1 | GMTACGGTCACACTG | chr3L:803051-805051 | 1975 | 1989 | - | 9.2857 | 1.62E-05 | 0.515 | GATAGGGTTTCACTG |
| 1 | GMTACGGTCACACTG | chr3L:11060571-11062571 | 1 | 15 | + | 9.2857 | 1.62E-05 | 0.515 | ACTATGGTCACAGTG |
| 1 | GMTACGGTCACACTG | chr3L:16050959-16052959 | 87 | 101 | + | 9.2857 | 1.62E-05 | 0.515 | AATGGGGTCACACTG |
| 1 | GMTACGGTCACACTG | chr2R:23244399-23246399 | 101 | 115 | + | 9.2857 | 1.62E-05 | 0.515 | GACACGGTCACATTC |
| 1 | GMTACGGTCACACTG | chr2L:18112179-18114179 | 147 | 161 | + | 9.2857 | 1.62E-05 | 0.515 | GAAAAGTTCACACTG |
| 1 | GMTACGGTCACACTG | chr3R:11695509-11697509 | 163 | 177 | + | 9.2857 | 1.62E-05 | 0.515 | GATACTGTCAAACAG |
| 1 | GMTACGGTCACACTG | chr2R:23061593-23063593 | 188 | 202 | + | 9.2857 | 1.62E-05 | 0.515 | GCTACGATTACACTT |
| 1 | GMTACGGTCACACTG | chr3L:26164235-26166235 | 188 | 202 | + | 9.2857 | 1.62E-05 | 0.515 | GCTACGCTCACATTT |
| 1 | GMTACGGTCACACTG | chr3R:20677183-20679183 | 191 | 205 | + | 9.2857 | 1.62E-05 | 0.515 | TATACGGTCATACGG |
| 1 | GMTACGGTCACACTG | chr2R:15216617-15218617 | 193 | 207 | + | 9.2857 | 1.62E-05 | 0.515 | GAAACGGTCACAGTC |
| 1 | GMTACGGTCACACTG | chr3R:23588819-23590819 | 211 | 225 | + | 9.2857 | 1.62E-05 | 0.515 | GATAAGATCACTCTG |
| 1 | GMTACGGTCACACTG | chrX:1010643-1012643 | 237 | 251 | + | 9.2857 | 1.62E-05 | 0.515 | AATACGCCCACACTG |
| 1 | GMTACGGTCACACTG | chr2R:12435577-12437577 | 253 | 267 | + | 9.2857 | 1.62E-05 | 0.515 | GACACGTTCAGACTG |
| 1 | GMTACGGTCACACTG | chr3L:12652090-12654090 | 304 | 318 | + | 9.2857 | 1.62E-05 | 0.515 | GATACAGATACACTG |
| 1 | GMTACGGTCACACTG | chr3R:10878319-10880319 | 335 | 349 | + | 9.2857 | 1.62E-05 | 0.515 | TCTACGATCACAATG |
| 1 | GMTACGGTCACACTG | chr3L:11060224-11062224 | 348 | 362 | + | 9.2857 | 1.62E-05 | 0.515 | ACTATGGTCACAGTG |
| 1 | GMTACGGTCACACTG | chr3L:12266528-12268528 | 353 | 367 | + | 9.2857 | 1.62E-05 | 0.515 | CATACGGTCACACAC |
| 1 | GMTACGGTCACACTG | chr2R:22378637-22380637 | 371 | 385 | + | 9.2857 | 1.62E-05 | 0.515 | GATCCGATCACAATG |
| 1 | GMTACGGTCACACTG | chr3R:27259050-27261050 | 375 | 389 | + | 9.2857 | 1.62E-05 | 0.515 | GATATGGTGGCACTG |
| 1 | GMTACGGTCACACTG | chr2R:13482229-13484229 | 377 | 391 | + | 9.2857 | 1.62E-05 | 0.515 | GATAAGGTGACTCTG |
| 1 | GMTACGGTCACACTG | chr2R:23104169-23106169 | 387 | 401 | + | 9.2857 | 1.62E-05 | 0.515 | GATTCGGTCACAATC |
| 1 | GMTACGGTCACACTG | chr3R:27581241-27583241 | 400 | 414 | + | 9.2857 | 1.62E-05 | 0.515 | CACACGGTCACACTA |
| 1 | GMTACGGTCACACTG | chr2L:7453212-7455212 | 446 | 460 | + | 9.2857 | 1.62E-05 | 0.515 | GATATGGTGGCACTG |
| 1 | GMTACGGTCACACTG | chr3L:1830783-1832783 | 460 | 474 | + | 9.2857 | 1.62E-05 | 0.515 | GATGCAATCACACTG |
| 1 | GMTACGGTCACACTG | chr3L:769363-771363 | 482 | 496 | + | 9.2857 | 1.62E-05 | 0.515 | GCTACGTTTAAACTG |
| 1 | GMTACGGTCACACTG | chr3L:15096914-15098914 | 487 | 501 | + | 9.2857 | 1.62E-05 | 0.515 | CATACGGTGACTCTG |
| 1 | GMTACGGTCACACTG | chr2R:7781637-7783637 | 514 | 528 | + | 9.2857 | 1.62E-05 | 0.515 | GATACCGTTGCACTG |
| 1 | GMTACGGTCACACTG | chr2R:7781637-7783637 | 514 | 528 | + | 9.2857 | 1.62E-05 | 0.515 | GATACCGTTGCACTG |
| 1 | GMTACGGTCACACTG | chrX:15908676-15910676 | 535 | 549 | + | 9.2857 | 1.62E-05 | 0.515 | GATATGGGAACACTG |
| 1 | GMTACGGTCACACTG | chrX:9727488-9729488 | 535 | 549 | + | 9.2857 | 1.62E-05 | 0.515 | GATACGTGCATACTG |
| 1 | GMTACGGTCACACTG | chr2L:16531877-16533877 | 540 | 554 | + | 9.2857 | 1.62E-05 | 0.515 | GATACGTTTACACAG |

| | | | | | | | | | |
|---|---|---|---|---|---|---|---|---|---|
| 1 | GMTACGGTCACACTG | chr3R:7130560-7132560 | 562 | 576 | + | 9.2857 | 1.62E-05 | 0.515 | TATGCGGTCACACTC |
| 1 | GMTACGGTCACACTG | chr3R:15327729-15329729 | 569 | 583 | + | 9.2857 | 1.62E-05 | 0.515 | AATTCGGTCATACTG |
| 1 | GMTACGGTCACACTG | chr3L:17757112-17759112 | 594 | 608 | + | 9.2857 | 1.62E-05 | 0.515 | GCTAAGCTTACACTG |
| 1 | GMTACGGTCACACTG | chr3L:12137868-12139868 | 603 | 617 | + | 9.2857 | 1.62E-05 | 0.515 | CAAACGGTCACACTT |
| 1 | GMTACGGTCACACTG | chrX:15787897-15789897 | 604 | 618 | + | 9.2857 | 1.62E-05 | 0.515 | AATGCGGTTACACTG |
| 1 | GMTACGGTCACACTG | chr3R:4947096-4949096 | 632 | 646 | + | 9.2857 | 1.62E-05 | 0.515 | TATACGGGCAGACTG |
| 1 | GMTACGGTCACACTG | chr3R:7059767-7061767 | 661 | 675 | + | 9.2857 | 1.62E-05 | 0.515 | CATACGGACAGACTG |
| 1 | GMTACGGTCACACTG | chr2R:7915856-7917856 | 662 | 676 | + | 9.2857 | 1.62E-05 | 0.515 | TACACGGTCACATTG |
| 1 | GMTACGGTCACACTG | chr2R:9862564-9864564 | 677 | 691 | + | 9.2857 | 1.62E-05 | 0.515 | GAAACGGTAACACTT |
| 1 | GMTACGGTCACACTG | chrX:10771541-10773541 | 687 | 701 | + | 9.2857 | 1.62E-05 | 0.515 | AATACGGTCAATCTG |
| 1 | GMTACGGTCACACTG | chrX:8324189-8326189 | 687 | 701 | + | 9.2857 | 1.62E-05 | 0.515 | GATACGGCGAAACTG |
| 1 | GMTACGGTCACACTG | chr3L:15094142-15096142 | 695 | 709 | + | 9.2857 | 1.62E-05 | 0.515 | GACACGGTAACAGTG |
| 1 | GMTACGGTCACACTG | chr2R:12435120-12437120 | 710 | 724 | + | 9.2857 | 1.62E-05 | 0.515 | GACACGTTCAGACTG |
| 1 | GMTACGGTCACACTG | chr2L:5380074-5382074 | 750 | 764 | + | 9.2857 | 1.62E-05 | 0.515 | GATCCGGTCATATTG |
| 1 | GMTACGGTCACACTG | chrX:19565019-19567019 | 764 | 778 | + | 9.2857 | 1.62E-05 | 0.515 | GATGTGGTGACACTG |
| 1 | GMTACGGTCACACTG | chr3R:18219699-18221699 | 797 | 811 | + | 9.2857 | 1.62E-05 | 0.515 | AATACGGCCACACTA |
| 1 | GMTACGGTCACACTG | chrX:13781459-13783459 | 836 | 850 | + | 9.2857 | 1.62E-05 | 0.515 | ACTACGATCACACTA |
| 1 | GMTACGGTCACACTG | chrX:7880679-7882679 | 873 | 887 | + | 9.2857 | 1.62E-05 | 0.515 | GATGGGGTCACACTC |
| 1 | GMTACGGTCACACTG | chr3L:2256456-2258456 | 886 | 900 | + | 9.2857 | 1.62E-05 | 0.515 | GACACGGTCAAACTT |
| 1 | GMTACGGTCACACTG | chr3L:12137584-12139584 | 887 | 901 | + | 9.2857 | 1.62E-05 | 0.515 | CAAACGGTCACACTT |
| 1 | GMTACGGTCACACTG | chr2R:9838776-9840776 | 900 | 914 | + | 9.2857 | 1.62E-05 | 0.515 | GCTACTGTCAAAATG |
| 1 | GMTACGGTCACACTG | chr2R:14172693-14174693 | 909 | 923 | + | 9.2857 | 1.62E-05 | 0.515 | GATTGGGTCACACTA |
| 1 | GMTACGGTCACACTG | chr3L:2256423-2258423 | 919 | 933 | + | 9.2857 | 1.62E-05 | 0.515 | GACACGGTCAAACTT |
| 1 | GMTACGGTCACACTG | chrX:19604199-19606199 | 922 | 936 | + | 9.2857 | 1.62E-05 | 0.515 | GATATCGCCACACTG |
| 1 | GMTACGGTCACACTG | chr2L:19957713-19959713 | 933 | 947 | + | 9.2857 | 1.62E-05 | 0.515 | GATGTGGTCACACTT |
| 1 | GMTACGGTCACACTG | chr2R:10141338-10143338 | 934 | 948 | + | 9.2857 | 1.62E-05 | 0.515 | AAGACAGTCACACTG |
| 1 | GMTACGGTCACACTG | chr3R:8800564-8802564 | 940 | 954 | + | 9.2857 | 1.62E-05 | 0.515 | CAAACGTTCACACTG |
| 1 | GMTACGGTCACACTG | chr3R:18219526-18221526 | 970 | 984 | + | 9.2857 | 1.62E-05 | 0.515 | AATACGGCCACACTA |
| 1 | GMTACGGTCACACTG | chr2L:21572389-21574389 | 974 | 988 | + | 9.2857 | 1.62E-05 | 0.515 | AATTCGGTCACACTA |
| 1 | GMTACGGTCACACTG | chr2L:21309866-21311866 | 977 | 991 | + | 9.2857 | 1.62E-05 | 0.515 | GATCTGGTAACACTG |
| 1 | GMTACGGTCACACTG | chr3R:10707340-10709340 | 984 | 998 | + | 9.2857 | 1.62E-05 | 0.515 | AACACGGTCATACTG |
| 1 | GMTACGGTCACACTG | chr3R:15277668-15279668 | 985 | 999 | + | 9.2857 | 1.62E-05 | 0.515 | CATGCGGTTACACTG |
| 1 | GMTACGGTCACACTG | chr3R:19026368-19028368 | 985 | 999 | + | 9.2857 | 1.62E-05 | 0.515 | CAAACGTTCACACTG |

| | | | | | | | | | |
|---|---|---|---|---|---|---|---|---|---|
| 1 | GMTACGGTCACACTG | chr2L:6061989-6063989 | 990 | 1004 | + | 9.2857 | 1.62E-05 | 0.515 | AATACGTCCACACTG |
| 1 | GMTACGGTCACACTG | chr2L:16261141-16263141 | 990 | 1004 | + | 9.2857 | 1.62E-05 | 0.515 | CATCTGGTCACACTG |
| 1 | GMTACGGTCACACTG | chr2R:20666306-20668306 | 993 | 1007 | + | 9.2857 | 1.62E-05 | 0.515 | AATACGGTCACACCT |
| 1 | GMTACGGTCACACTG | chr3L:19091670-19093670 | 996 | 1010 | + | 9.2857 | 1.62E-05 | 0.515 | AATCTGGTCACACTG |
| 1 | GMTACGGTCACACTG | chr3R:23923399-23925399 | 997 | 1011 | + | 9.2857 | 1.62E-05 | 0.515 | CATGCGGTCACACTT |
| 1 | GMTACGGTCACACTG | chrX:8460479-8462479 | 998 | 1012 | + | 9.2857 | 1.62E-05 | 0.515 | AAAACGGTCACACTT |
| 1 | GMTACGGTCACACTG | chr3L:20312632-20314632 | 1001 | 1015 | + | 9.2857 | 1.62E-05 | 0.515 | GAGATGGTAACACTG |
| 1 | GMTACGGTCACACTG | chr2R:22378007-22380007 | 1001 | 1015 | + | 9.2857 | 1.62E-05 | 0.515 | GATCCGATCACAATG |
| 1 | GMTACGGTCACACTG | chr2R:14172596-14174596 | 1006 | 1020 | + | 9.2857 | 1.62E-05 | 0.515 | GATTGGGTCACACTA |
| 1 | GMTACGGTCACACTG | chr3L:4102564-4104564 | 1006 | 1020 | + | 9.2857 | 1.62E-05 | 0.515 | CAAACGGTCATACTG |
| 1 | GMTACGGTCACACTG | chr2L:4970702-4972702 | 1008 | 1022 | + | 9.2857 | 1.62E-05 | 0.515 | AATACGGCCACACTA |
| 1 | GMTACGGTCACACTG | chr3L:4291482-4293482 | 1009 | 1023 | + | 9.2857 | 1.62E-05 | 0.515 | AATACGGTCACACAC |
| 1 | GMTACGGTCACACTG | chr3L:16050034-16052034 | 1012 | 1026 | + | 9.2857 | 1.62E-05 | 0.515 | AATGGGGTCACACTG |
| 1 | GMTACGGTCACACTG | chr2R:22887928-22889928 | 1013 | 1027 | + | 9.2857 | 1.62E-05 | 0.515 | CATTTGGTCACACTG |
| 1 | GMTACGGTCACACTG | chr2L:13792521-13794521 | 1018 | 1032 | + | 9.2857 | 1.62E-05 | 0.515 | TACACGGTCACACTT |
| 1 | GMTACGGTCACACTG | chrX:19564764-19566764 | 1019 | 1033 | + | 9.2857 | 1.62E-05 | 0.515 | GATGTGGTGACACTG |
| 1 | GMTACGGTCACACTG | chr2R:24742749-24744749 | 1021 | 1035 | + | 9.2857 | 1.62E-05 | 0.515 | GAAACAGTCACACTC |
| 1 | GMTACGGTCACACTG | chr3R:20676351-20678351 | 1023 | 1037 | + | 9.2857 | 1.62E-05 | 0.515 | TATACGGTCATACGG |
| 1 | GMTACGGTCACACTG | chr2R:14756846-14758846 | 1024 | 1038 | + | 9.2857 | 1.62E-05 | 0.515 | GAAATGGTCACACTT |
| 1 | GMTACGGTCACACTG | chr3L:20797645-20799645 | 1024 | 1038 | + | 9.2857 | 1.62E-05 | 0.515 | CAAACGGTCACACTC |
| 1 | GMTACGGTCACACTG | chr3R:15327269-15329269 | 1029 | 1043 | + | 9.2857 | 1.62E-05 | 0.515 | AATTCGGTCATACTG |
| 1 | GMTACGGTCACACTG | chr2L:9789882-9791882 | 1032 | 1046 | + | 9.2857 | 1.62E-05 | 0.515 | CATCCGGTAACACTG |
| 1 | GMTACGGTCACACTG | chr3L:12265841-12267841 | 1040 | 1054 | + | 9.2857 | 1.62E-05 | 0.515 | CATACGGTCACACAC |
| 1 | GMTACGGTCACACTG | chr2R:7915478-7917478 | 1040 | 1054 | + | 9.2857 | 1.62E-05 | 0.515 | TACACGGTCACATTG |
| 1 | GMTACGGTCACACTG | chr2L:251373-253373 | 1044 | 1058 | + | 9.2857 | 1.62E-05 | 0.515 | AACACGGTCACACTA |
| 1 | GMTACGGTCACACTG | chr3L:276437-278437 | 1045 | 1059 | + | 9.2857 | 1.62E-05 | 0.515 | GATAAAGTCCCACTG |
| 1 | GMTACGGTCACACTG | chr2R:10121574-10123574 | 1049 | 1063 | + | 9.2857 | 1.62E-05 | 0.515 | GATAGGTTCACACGG |
| 1 | GMTACGGTCACACTG | chr2R:12137695-12139695 | 1051 | 1065 | + | 9.2857 | 1.62E-05 | 0.515 | AATGCGGTCACACTA |
| 1 | GMTACGGTCACACTG | chr2R:22887857-22889857 | 1084 | 1098 | + | 9.2857 | 1.62E-05 | 0.515 | CATTTGGTCACACTG |
| 1 | GMTACGGTCACACTG | chrX:12956462-12958462 | 1088 | 1102 | + | 9.2857 | 1.62E-05 | 0.515 | CATACTGGCACACTG |
| 1 | GMTACGGTCACACTG | chr3R:30629387-30631387 | 1091 | 1105 | + | 9.2857 | 1.62E-05 | 0.515 | GATCCGGTTACACTT |
| 1 | GMTACGGTCACACTG | chr3R:7130018-7132018 | 1104 | 1118 | + | 9.2857 | 1.62E-05 | 0.515 | TATGCGGTCACACTC |
| 1 | GMTACGGTCACACTG | chr3R:25684051-25686051 | 1104 | 1118 | + | 9.2857 | 1.62E-05 | 0.515 | TATCTGGTCACACTG |

| | | | | | | | | | |
|---|---|---|---|---|---|---|---|---|---|
| 1 | GMTACGGTCACACTG | chrX:21519017-21521017 | 1105 | 1119 | + | 9.2857 | 1.62E-05 | 0.515 | AAGACGGTCACACTT |
| 1 | GMTACGGTCACACTG | chr3L:627435-629435 | 1110 | 1124 | + | 9.2857 | 1.62E-05 | 0.515 | CATCTGGTCACACTG |
| 1 | GMTACGGTCACACTG | chr3L:15093724-15095724 | 1113 | 1127 | + | 9.2857 | 1.62E-05 | 0.515 | GACACGGTAACAGTG |
| 1 | GMTACGGTCACACTG | chr3L:15093719-15095719 | 1118 | 1132 | + | 9.2857 | 1.62E-05 | 0.515 | GACACGGTAACAGTG |
| 1 | GMTACGGTCACACTG | chr2R:11339132-11341132 | 1122 | 1136 | + | 9.2857 | 1.62E-05 | 0.515 | GCTACTATAACACTG |
| 1 | GMTACGGTCACACTG | chrX:13781170-13783170 | 1125 | 1139 | + | 9.2857 | 1.62E-05 | 0.515 | ACTACGATCACACTA |
| 1 | GMTACGGTCACACTG | chrX:1050525-1052525 | 1134 | 1148 | + | 9.2857 | 1.62E-05 | 0.515 | GCTAAGGTCAAACTA |
| 1 | GMTACGGTCACACTG | chr3L:6529729-6531729 | 1140 | 1154 | + | 9.2857 | 1.62E-05 | 0.515 | GAGATTGTCACACTG |
| 1 | GMTACGGTCACACTG | chr3R:21867003-21869003 | 1168 | 1182 | + | 9.2857 | 1.62E-05 | 0.515 | GATACGGTCTTACTA |
| 1 | GMTACGGTCACACTG | chr2L:9789745-9791745 | 1169 | 1183 | + | 9.2857 | 1.62E-05 | 0.515 | CATCCGGTAACACTG |
| 1 | GMTACGGTCACACTG | chr2R:9838500-9840500 | 1176 | 1190 | + | 9.2857 | 1.62E-05 | 0.515 | GCTACTGTCAAAATG |
| 1 | GMTACGGTCACACTG | chr3L:19518649-19520649 | 1182 | 1196 | + | 9.2857 | 1.62E-05 | 0.515 | GATACGATCACGATG |
| 1 | GMTACGGTCACACTG | chr3L:20344122-20346122 | 1198 | 1212 | + | 9.2857 | 1.62E-05 | 0.515 | GAGATGGTCACAGTG |
| 1 | GMTACGGTCACACTG | chr2R:10141074-10143074 | 1198 | 1212 | + | 9.2857 | 1.62E-05 | 0.515 | AAGACAGTCACACTG |
| 1 | GMTACGGTCACACTG | chr2R:14991230-14993230 | 1207 | 1221 | + | 9.2857 | 1.62E-05 | 0.515 | CAAAAGGTCACACTG |
| 1 | GMTACGGTCACACTG | chr2L:6061757-6063757 | 1222 | 1236 | + | 9.2857 | 1.62E-05 | 0.515 | AATACGTCCACACTG |
| 1 | GMTACGGTCACACTG | chr3L:1730107-1732107 | 1240 | 1254 | + | 9.2857 | 1.62E-05 | 0.515 | GACACGATCAAACTG |
| 1 | GMTACGGTCACACTG | chr3L:4291246-4293246 | 1245 | 1259 | + | 9.2857 | 1.62E-05 | 0.515 | AATACGGTCACACAC |
| 1 | GMTACGGTCACACTG | chr3R:21866919-21868919 | 1252 | 1266 | + | 9.2857 | 1.62E-05 | 0.515 | GATACGGTCTTACTA |
| 1 | GMTACGGTCACACTG | chr2L:16260877-16262877 | 1254 | 1268 | + | 9.2857 | 1.62E-05 | 0.515 | CATCTGGTCACACTG |
| 1 | GMTACGGTCACACTG | chr3L:20797375-20799375 | 1294 | 1308 | + | 9.2857 | 1.62E-05 | 0.515 | CAAACGGTCACACTC |
| 1 | GMTACGGTCACACTG | chr3R:20676080-20678080 | 1294 | 1308 | + | 9.2857 | 1.62E-05 | 0.515 | TATACGGTCATACGG |
| 1 | GMTACGGTCACACTG | chrX:10841205-10843205 | 1296 | 1310 | + | 9.2857 | 1.62E-05 | 0.515 | GATGCCGTCACATTG |
| 1 | GMTACGGTCACACTG | chr3L:4102256-4104256 | 1314 | 1328 | + | 9.2857 | 1.62E-05 | 0.515 | CAAACGGTCATACTG |
| 1 | GMTACGGTCACACTG | chr3R:8800176-8802176 | 1328 | 1342 | + | 9.2857 | 1.62E-05 | 0.515 | CAAACGTTCACACTG |
| 1 | GMTACGGTCACACTG | chr3R:8800176-8802176 | 1328 | 1342 | + | 9.2857 | 1.62E-05 | 0.515 | CAAACGTTCACACTG |
| 1 | GMTACGGTCACACTG | chr3L:768503-770503 | 1342 | 1356 | + | 9.2857 | 1.62E-05 | 0.515 | GCTACGTTTAAACTG |
| 1 | GMTACGGTCACACTG | chr2R:12080276-12082276 | 1348 | 1362 | + | 9.2857 | 1.62E-05 | 0.515 | GATAAGTTCCCACTG |
| 1 | GMTACGGTCACACTG | chr2R:23358215-23360215 | 1376 | 1390 | + | 9.2857 | 1.62E-05 | 0.515 | GCTAAGGTAACATTG |
| 1 | GMTACGGTCACACTG | chr2L:816950-818950 | 1387 | 1401 | + | 9.2857 | 1.62E-05 | 0.515 | GATCCTGTCATACTG |
| 1 | GMTACGGTCACACTG | chr2R:20908704-20910704 | 1408 | 1422 | + | 9.2857 | 1.62E-05 | 0.515 | GATATGGTCGCACTA |
| 1 | GMTACGGTCACACTG | chrX:11062170-11064170 | 1448 | 1462 | + | 9.2857 | 1.62E-05 | 0.515 | GATACAGTTCCACTG |
| 1 | GMTACGGTCACACTG | chr3L:2261919-2263919 | 1455 | 1469 | + | 9.2857 | 1.62E-05 | 0.515 | GATACGATCACATCG |

| | | | | | | | | | |
|---|---|---|---|---|---|---|---|---|---|
| 1 | GMTACGGTCACACTG | chr2L:3022226-3024226 | 1468 | 1482 | + | 9.2857 | 1.62E-05 | 0.515 | GATACGGGCAAAGTG |
| 1 | GMTACGGTCACACTG | chrX:4996441-4998441 | 1470 | 1484 | + | 9.2857 | 1.62E-05 | 0.515 | GAGACGATCAAACTG |
| 1 | GMTACGGTCACACTG | chr2L:9937863-9939863 | 1482 | 1496 | + | 9.2857 | 1.62E-05 | 0.515 | GATACCGCCACAGTG |
| 1 | GMTACGGTCACACTG | chrX:7825197-7827197 | 1533 | 1547 | + | 9.2857 | 1.62E-05 | 0.515 | GATACAGTCACATAG |
| 1 | GMTACGGTCACACTG | chr3R:15277109-15279109 | 1544 | 1558 | + | 9.2857 | 1.62E-05 | 0.515 | CATGCGGTTACACTG |
| 1 | GMTACGGTCACACTG | chr2L:3022141-3024141 | 1553 | 1567 | + | 9.2857 | 1.62E-05 | 0.515 | GATACGGGCAAAGTG |
| 1 | GMTACGGTCACACTG | chr2L:9937790-9939790 | 1555 | 1569 | + | 9.2857 | 1.62E-05 | 0.515 | GATACCGCCACAGTG |
| 1 | GMTACGGTCACACTG | chr2R:21865859-21867859 | 1571 | 1585 | + | 9.2857 | 1.62E-05 | 0.515 | GAGAGGGTGACACTG |
| 1 | GMTACGGTCACACTG | chrX:8459898-8461898 | 1579 | 1593 | + | 9.2857 | 1.62E-05 | 0.515 | AAAACGGTCACACTT |
| 1 | GMTACGGTCACACTG | chr2R:6136303-6138303 | 1595 | 1609 | + | 9.2857 | 1.62E-05 | 0.515 | GACACTGTTACACTG |
| 1 | GMTACGGTCACACTG | chr3R:13419279-13421279 | 1595 | 1609 | + | 9.2857 | 1.62E-05 | 0.515 | ACTACGGTAAAACTG |
| 1 | GMTACGGTCACACTG | chr2R:15215207-15217207 | 1603 | 1617 | + | 9.2857 | 1.62E-05 | 0.515 | GAAACGGTCACAGTC |
| 1 | GMTACGGTCACACTG | chr3R:19397567-19399567 | 1604 | 1618 | + | 9.2857 | 1.62E-05 | 0.515 | GCTATTCTCACACTG |
| 1 | GMTACGGTCACACTG | chr3R:20896320-20898320 | 1622 | 1636 | + | 9.2857 | 1.62E-05 | 0.515 | GACACCGTGACACTG |
| 1 | GMTACGGTCACACTG | chr2R:23357961-23359961 | 1630 | 1644 | + | 9.2857 | 1.62E-05 | 0.515 | GCTAAGGTAACATTG |
| 1 | GMTACGGTCACACTG | chr3L:3207985-3209985 | 1641 | 1655 | + | 9.2857 | 1.62E-05 | 0.515 | GATACGCTCACTGTG |
| 1 | GMTACGGTCACACTG | chr3L:3207985-3209985 | 1641 | 1655 | + | 9.2857 | 1.62E-05 | 0.515 | GATACGCTCACTGTG |
| 1 | GMTACGGTCACACTG | chr2R:13445309-13447309 | 1665 | 1679 | + | 9.2857 | 1.62E-05 | 0.515 | GCTAAAGTCACACTT |
| 1 | GMTACGGTCACACTG | chr3L:8603657-8605657 | 1674 | 1688 | + | 9.2857 | 1.62E-05 | 0.515 | GCTACAGTTACACTA |
| 1 | GMTACGGTCACACTG | chr3R:28922044-28924044 | 1694 | 1708 | + | 9.2857 | 1.62E-05 | 0.515 | GATAAGGGCAAACTG |
| 1 | GMTACGGTCACACTG | chr2L:16793211-16795211 | 1709 | 1723 | + | 9.2857 | 1.62E-05 | 0.515 | GATATGCCCACACTG |
| 1 | GMTACGGTCACACTG | chr3L:2261664-2263664 | 1710 | 1724 | + | 9.2857 | 1.62E-05 | 0.515 | GATACGATCACATCG |
| 1 | GMTACGGTCACACTG | chrX:15836830-15838830 | 1763 | 1777 | + | 9.2857 | 1.62E-05 | 0.515 | AATACGGTCAAACAG |
| 1 | GMTACGGTCACACTG | chrX:18355191-18357191 | 1770 | 1784 | + | 9.2857 | 1.62E-05 | 0.515 | CAAACGGTCACACTC |
| 1 | GMTACGGTCACACTG | chr2L:331335-333335 | 1784 | 1798 | + | 9.2857 | 1.62E-05 | 0.515 | GATAAGGTCACCCTA |
| 1 | GMTACGGTCACACTG | chr2L:21309050-21311050 | 1793 | 1807 | + | 9.2857 | 1.62E-05 | 0.515 | GATCTGGTAACACTG |
| 1 | GMTACGGTCACACTG | chr2R:23102733-23104733 | 1823 | 1837 | + | 9.2857 | 1.62E-05 | 0.515 | GATTCGGTCACAATC |
| 1 | GMTACGGTCACACTG | chr2L:20639917-20641917 | 1828 | 1842 | + | 9.2857 | 1.62E-05 | 0.515 | GATACTGTCACACCA |
| 1 | GMTACGGTCACACTG | chr3R:23780765-23782765 | 1833 | 1847 | + | 9.2857 | 1.62E-05 | 0.515 | GCTACGGTAAAACTT |
| 1 | GMTACGGTCACACTG | chr3R:18590027-18592027 | 1837 | 1851 | + | 9.2857 | 1.62E-05 | 0.515 | GATGCGGTCATACTC |
| 1 | GMTACGGTCACACTG | chrX:19520929-19522929 | 1859 | 1873 | + | 9.2857 | 1.62E-05 | 0.515 | CATACGCTCACACGG |
| 1 | GMTACGGTCACACTG | chr3R:10876783-10878783 | 1871 | 1885 | + | 9.2857 | 1.62E-05 | 0.515 | TCTACGATCACAATG |
| 1 | GMTACGGTCACACTG | chr3L:9628193-9630193 | 1887 | 1901 | + | 9.2857 | 1.62E-05 | 0.515 | CACACAGTCACACTG |

| | | | | | | | | | |
|---|---|---|---|---|---|---|---|---|---|
| 1 | GMTACGGTCACACTG | chr3L:1033875-1035875 | 1891 | 1905 | + | 9.2857 | 1.62E-05 | 0.515 | GATACGGTAAAGCTG |
| 1 | GMTACGGTCACACTG | chr3L:11097412-11099412 | 1895 | 1909 | + | 9.2857 | 1.62E-05 | 0.515 | AATACGGGAACACTG |
| 1 | GMTACGGTCACACTG | chrX:21518218-21520218 | 1904 | 1918 | + | 9.2857 | 1.62E-05 | 0.515 | AAGACGGTCACACTT |
| 1 | GMTACGGTCACACTG | chr2R:20985500-20987500 | 1952 | 1966 | + | 9.2857 | 1.62E-05 | 0.515 | TATATGGTCACGCTG |
| 1 | GMTACGGTCACACTG | chrX:4616564-4618564 | 1962 | 1976 | + | 9.2857 | 1.62E-05 | 0.515 | CCTACGATCACAATG |
| 1 | GMTACGGTCACACTG | chr3R:17137999-17139999 | 1972 | 1986 | + | 9.2857 | 1.62E-05 | 0.515 | GCTATGTTCACAGTG |
| 1 | GMTACGGTCACACTG | chr3R:23780621-23782621 | 1977 | 1991 | + | 9.2857 | 1.62E-05 | 0.515 | GCTACGGTAAAACTT |
| 1 | GMTACGGTCACACTG | chrX:19520810-19522810 | 1978 | 1992 | + | 9.2857 | 1.62E-05 | 0.515 | CATACGCTCACACGG |
| 1 | GMTACGGTCACACTG | chr2R:23357612-23359612 | 1979 | 1993 | + | 9.2857 | 1.62E-05 | 0.515 | GCTAAGGTAACATTG |
| 1 | GMTACGGTCACACTG | chr3L:18679409-18681409 | 24 | 38 | - | 9.0071 | 1.80E-05 | 0.546 | GATATGATCACATTG |
| 1 | GMTACGGTCACACTG | chr3R:29493782-29495782 | 30 | 44 | - | 9.0071 | 1.80E-05 | 0.546 | TATACGCTCACAGTG |
| 1 | GMTACGGTCACACTG | chr3R:7143295-7145295 | 266 | 280 | - | 9.0071 | 1.80E-05 | 0.546 | GATATGGTCACAATA |
| 1 | GMTACGGTCACACTG | chr3L:13964203-13966203 | 487 | 501 | - | 9.0071 | 1.80E-05 | 0.546 | AATACGTTCACATTG |
| 1 | GMTACGGTCACACTG | chr3L:13964203-13966203 | 487 | 501 | - | 9.0071 | 1.80E-05 | 0.546 | AATACGTTCACATTG |
| 1 | GMTACGGTCACACTG | chr3R:13836991-13838991 | 522 | 536 | - | 9.0071 | 1.80E-05 | 0.546 | GATATCATCACACTG |
| 1 | GMTACGGTCACACTG | chr2L:7408809-7410809 | 617 | 631 | - | 9.0071 | 1.80E-05 | 0.546 | GATATTGTCAGACTG |
| 1 | GMTACGGTCACACTG | chr2R:9867219-9869219 | 660 | 674 | - | 9.0071 | 1.80E-05 | 0.546 | GATACAGTCACAATT |
| 1 | GMTACGGTCACACTG | chrX:6065416-6067416 | 706 | 720 | - | 9.0071 | 1.80E-05 | 0.546 | GATAGTGTCACACTT |
| 1 | GMTACGGTCACACTG | chr3R:10261123-10263123 | 726 | 740 | - | 9.0071 | 1.80E-05 | 0.546 | GATACGGTCATATTA |
| 1 | GMTACGGTCACACTG | chr3R:14662450-14664450 | 729 | 743 | - | 9.0071 | 1.80E-05 | 0.546 | CATATGGTAACACTG |
| 1 | GMTACGGTCACACTG | chr3R:14131072-14133072 | 747 | 761 | - | 9.0071 | 1.80E-05 | 0.546 | TATAAAGTCACACTG |
| 1 | GMTACGGTCACACTG | chrX:8922960-8924960 | 891 | 905 | - | 9.0071 | 1.80E-05 | 0.546 | GATAGCTTCACACTG |
| 1 | GMTACGGTCACACTG | chr3R:16348404-16350404 | 898 | 912 | - | 9.0071 | 1.80E-05 | 0.546 | GATATGGTCATACTA |
| 1 | GMTACGGTCACACTG | chr3R:26535350-26537350 | 924 | 938 | - | 9.0071 | 1.80E-05 | 0.546 | CATACGTTCACACTT |
| 1 | GMTACGGTCACACTG | chr3R:9792295-9794295 | 927 | 941 | - | 9.0071 | 1.80E-05 | 0.546 | CATATGGTCACACTA |
| 1 | GMTACGGTCACACTG | chr3L:15497959-15499959 | 932 | 946 | - | 9.0071 | 1.80E-05 | 0.546 | AATAGGGTAACACTG |
| 1 | GMTACGGTCACACTG | chr3R:30045470-30047470 | 953 | 967 | - | 9.0071 | 1.80E-05 | 0.546 | AATATGTTCACACTG |
| 1 | GMTACGGTCACACTG | chr3R:29492852-29494852 | 960 | 974 | - | 9.0071 | 1.80E-05 | 0.546 | TATACGCTCACAGTG |
| 1 | GMTACGGTCACACTG | chr3R:14662197-14664197 | 982 | 996 | - | 9.0071 | 1.80E-05 | 0.546 | CATATGGTAACACTG |
| 1 | GMTACGGTCACACTG | chr3L:13963702-13965702 | 988 | 1002 | - | 9.0071 | 1.80E-05 | 0.546 | AATACGTTCACATTG |
| 1 | GMTACGGTCACACTG | chr3L:10660475-10662475 | 993 | 1007 | - | 9.0071 | 1.80E-05 | 0.546 | GATACTGTCACAGTT |
| 1 | GMTACGGTCACACTG | chr3R:10260849-10262849 | 1000 | 1014 | - | 9.0071 | 1.80E-05 | 0.546 | GATACGGTCATATTA |
| 1 | GMTACGGTCACACTG | chr2R:9191096-9193096 | 1006 | 1020 | - | 9.0071 | 1.80E-05 | 0.546 | CATATGGTAACACTG |

| | | | | | | | | | |
|---|---|---|---|---|---|---|---|---|---|
| 1 | GMTACGGTCACACTG | chr2R:9191096-9193096 | 1006 | 1020 | - | 9.0071 | 1.80E-05 | 0.546 | CATATGGTAACACTG |
| 1 | GMTACGGTCACACTG | chr2R:6014689-6016689 | 1009 | 1023 | - | 9.0071 | 1.80E-05 | 0.546 | TATATGGTCACACTA |
| 1 | GMTACGGTCACACTG | chr3R:7142487-7144487 | 1074 | 1088 | - | 9.0071 | 1.80E-05 | 0.546 | GATATGGTCACAATA |
| 1 | GMTACGGTCACACTG | chr3R:16348226-16350226 | 1076 | 1090 | - | 9.0071 | 1.80E-05 | 0.546 | GATATGGTCATACTA |
| 1 | GMTACGGTCACACTG | chr2L:20675045-20677045 | 1146 | 1160 | - | 9.0071 | 1.80E-05 | 0.546 | AATACGATAACACTG |
| 1 | GMTACGGTCACACTG | chrX:8922687-8924687 | 1164 | 1178 | - | 9.0071 | 1.80E-05 | 0.546 | GATAGCTTCACACTG |
| 1 | GMTACGGTCACACTG | chr3L:21130446-21132446 | 1167 | 1181 | - | 9.0071 | 1.80E-05 | 0.546 | TATAGGGTCACACTA |
| 1 | GMTACGGTCACACTG | chr3L:10693108-10695108 | 1170 | 1184 | - | 9.0071 | 1.80E-05 | 0.546 | GATACGTTCACATTA |
| 1 | GMTACGGTCACACTG | chr3R:23383777-23385777 | 1188 | 1202 | - | 9.0071 | 1.80E-05 | 0.546 | GATACTGTGATACTG |
| 1 | GMTACGGTCACACTG | chr3L:3393088-3395088 | 1188 | 1202 | - | 9.0071 | 1.80E-05 | 0.546 | TATACACTCACACTG |
| 1 | GMTACGGTCACACTG | chr3R:29923615-29925615 | 1228 | 1242 | - | 9.0071 | 1.80E-05 | 0.546 | GATACTATCATACTG |
| 1 | GMTACGGTCACACTG | chr2L:20674941-20676941 | 1250 | 1264 | - | 9.0071 | 1.80E-05 | 0.546 | AATACGATAACACTG |
| 1 | GMTACGGTCACACTG | chr3R:30045161-30047161 | 1262 | 1276 | - | 9.0071 | 1.80E-05 | 0.546 | AATATGTTCACACTG |
| 1 | GMTACGGTCACACTG | chr3R:29492518-29494518 | 1294 | 1308 | - | 9.0071 | 1.80E-05 | 0.546 | TATACGCTCACAGTG |
| 1 | GMTACGGTCACACTG | chr3L:16761683-16763683 | 1341 | 1355 | - | 9.0071 | 1.80E-05 | 0.546 | GATACTTTCAAACTG |
| 1 | GMTACGGTCACACTG | chr3L:10625499-10627499 | 1365 | 1379 | - | 9.0071 | 1.80E-05 | 0.546 | GATACCATTACACTG |
| 1 | GMTACGGTCACACTG | chr3L:21130159-21132159 | 1454 | 1468 | - | 9.0071 | 1.80E-05 | 0.546 | TATAGGGTCACACTA |
| 1 | GMTACGGTCACACTG | chr3L:10660013-10662013 | 1455 | 1469 | - | 9.0071 | 1.80E-05 | 0.546 | GATACTGTCACAGTT |
| 1 | GMTACGGTCACACTG | chr3L:15497202-15499202 | 1689 | 1703 | - | 9.0071 | 1.80E-05 | 0.546 | AATAGGGTAACACTG |
| 1 | GMTACGGTCACACTG | chr2L:6347427-6349427 | 1788 | 1802 | - | 9.0071 | 1.80E-05 | 0.546 | GATAAGATCACATTG |
| 1 | GMTACGGTCACACTG | chr3R:11821493-11823493 | 1794 | 1808 | - | 9.0071 | 1.80E-05 | 0.546 | AATAGGATCACACTG |
| 1 | GMTACGGTCACACTG | chrX:17888547-17890547 | 1808 | 1822 | - | 9.0071 | 1.80E-05 | 0.546 | TATACCGTCACATTG |
| 1 | GMTACGGTCACACTG | chrX:17898702-17900702 | 1808 | 1822 | - | 9.0071 | 1.80E-05 | 0.546 | TATACCGTCACATTG |
| 1 | GMTACGGTCACACTG | chr2R:9182395-9184395 | 1902 | 1916 | - | 9.0071 | 1.80E-05 | 0.546 | GATATGGTCATACTA |
| 1 | GMTACGGTCACACTG | chr3R:17412901-17414901 | 81 | 95 | + | 9.0071 | 1.80E-05 | 0.546 | CATATGCTCACACTG |
| 1 | GMTACGGTCACACTG | chr2R:14620782-14622782 | 84 | 98 | + | 9.0071 | 1.80E-05 | 0.546 | TATATGCTCACACTG |
| 1 | GMTACGGTCACACTG | chr2R:7400859-7402859 | 94 | 108 | + | 9.0071 | 1.80E-05 | 0.546 | GATAAGGTCATAATG |
| 1 | GMTACGGTCACACTG | chr2R:14620601-14622601 | 265 | 279 | + | 9.0071 | 1.80E-05 | 0.546 | TATATGCTCACACTG |
| 1 | GMTACGGTCACACTG | chr2R:22126808-22128808 | 293 | 307 | + | 9.0071 | 1.80E-05 | 0.546 | GATATGCTCACAGTG |
| 1 | GMTACGGTCACACTG | chr3L:15579800-15581800 | 395 | 409 | + | 9.0071 | 1.80E-05 | 0.546 | GATACAGTCATACTT |
| 1 | GMTACGGTCACACTG | chr2R:10417718-10419718 | 444 | 458 | + | 9.0071 | 1.80E-05 | 0.546 | AATACCATCACACTG |
| 1 | GMTACGGTCACACTG | chr2R:17019027-17021027 | 539 | 553 | + | 9.0071 | 1.80E-05 | 0.546 | AATATGGTCACACTT |
| 1 | GMTACGGTCACACTG | chr2R:14620320-14622320 | 546 | 560 | + | 9.0071 | 1.80E-05 | 0.546 | TATATGCTCACACTG |

| | | | | | | | | | |
|---|---|---|---|---|---|---|---|---|---|
| 1 | GMTACGGTCACACTG | chr2R:22126502-22128502 | 599 | 613 | + | 9.0071 | 1.80E-05 | 0.546 | GATATGCTCACAGTG |
| 1 | GMTACGGTCACACTG | chr2L:15057303-15059303 | 651 | 665 | + | 9.0071 | 1.80E-05 | 0.546 | TATATGGTCACATTG |
| 1 | GMTACGGTCACACTG | chr3L:15579537-15581537 | 658 | 672 | + | 9.0071 | 1.80E-05 | 0.546 | GATACAGTCATACTT |
| 1 | GMTACGGTCACACTG | chr3L:11053035-11055035 | 665 | 679 | + | 9.0071 | 1.80E-05 | 0.546 | GATAATGTCACACTT |
| 1 | GMTACGGTCACACTG | chr3L:9893743-9895743 | 683 | 697 | + | 9.0071 | 1.80E-05 | 0.546 | GATAGGTTCACAATG |
| 1 | GMTACGGTCACACTG | chr3L:9893743-9895743 | 683 | 697 | + | 9.0071 | 1.80E-05 | 0.546 | GATAGGTTCACAATG |
| 1 | GMTACGGTCACACTG | chrX:17214304-17216304 | 880 | 894 | + | 9.0071 | 1.80E-05 | 0.546 | GATATGGTCACAGTT |
| 1 | GMTACGGTCACACTG | chr3R:17412077-17414077 | 905 | 919 | + | 9.0071 | 1.80E-05 | 0.546 | CATATGCTCACACTG |
| 1 | GMTACGGTCACACTG | chr3R:17149587-17151587 | 963 | 977 | + | 9.0071 | 1.80E-05 | 0.546 | TATATGGTCATACTG |
| 1 | GMTACGGTCACACTG | chr3L:5764846-5766846 | 967 | 981 | + | 9.0071 | 1.80E-05 | 0.546 | GATACCATCACAGTG |
| 1 | GMTACGGTCACACTG | chr3R:28605079-28607079 | 992 | 1006 | + | 9.0071 | 1.80E-05 | 0.546 | AATACGGTCATACTA |
| 1 | GMTACGGTCACACTG | chr2R:19783294-19785294 | 997 | 1011 | + | 9.0071 | 1.80E-05 | 0.546 | GATACAGTTAAACTG |
| 1 | GMTACGGTCACACTG | chr2L:16826078-16828078 | 1000 | 1014 | + | 9.0071 | 1.80E-05 | 0.546 | CATATGGTCACACTT |
| 1 | GMTACGGTCACACTG | chr3R:22033455-22035455 | 1015 | 1029 | + | 9.0071 | 1.80E-05 | 0.546 | CATACGGTCATACTA |
| 1 | GMTACGGTCACACTG | chr2L:10330471-10332471 | 1015 | 1029 | + | 9.0071 | 1.80E-05 | 0.546 | GATATGGTAACACTT |
| 1 | GMTACGGTCACACTG | chr3L:19927743-19929743 | 1017 | 1031 | + | 9.0071 | 1.80E-05 | 0.546 | CATACGTTCACACTT |
| 1 | GMTACGGTCACACTG | chr2R:17018522-17020522 | 1044 | 1058 | + | 9.0071 | 1.80E-05 | 0.546 | AATATGGTCACACTT |
| 1 | GMTACGGTCACACTG | chr3R:11973910-11975910 | 1059 | 1073 | + | 9.0071 | 1.80E-05 | 0.546 | AATATGGTCACACTA |
| 1 | GMTACGGTCACACTG | chr3L:7725473-7727473 | 1069 | 1083 | + | 9.0071 | 1.80E-05 | 0.546 | AATATGGTCACATTG |
| 1 | GMTACGGTCACACTG | chr2L:15056862-15058862 | 1092 | 1106 | + | 9.0071 | 1.80E-05 | 0.546 | TATATGGTCACATTG |
| 1 | GMTACGGTCACACTG | chr3L:5764702-5766702 | 1111 | 1125 | + | 9.0071 | 1.80E-05 | 0.546 | GATACCATCACAGTG |
| 1 | GMTACGGTCACACTG | chr3R:11779641-11781641 | 1158 | 1172 | + | 9.0071 | 1.80E-05 | 0.546 | GATACTCTCACACTC |
| 1 | GMTACGGTCACACTG | chr3R:22033298-22035298 | 1172 | 1186 | + | 9.0071 | 1.80E-05 | 0.546 | CATACGGTCATACTA |
| 1 | GMTACGGTCACACTG | chr3R:22033286-22035286 | 1184 | 1198 | + | 9.0071 | 1.80E-05 | 0.546 | CATACGGTCATACTA |
| 1 | GMTACGGTCACACTG | chr3R:22033286-22035286 | 1184 | 1198 | + | 9.0071 | 1.80E-05 | 0.546 | CATACGGTCATACTA |
| 1 | GMTACGGTCACACTG | chrX:7900880-7902880 | 1208 | 1222 | + | 9.0071 | 1.80E-05 | 0.546 | GATAAAGTCAAACTG |
| 1 | GMTACGGTCACACTG | chr2R:7399743-7401743 | 1210 | 1224 | + | 9.0071 | 1.80E-05 | 0.546 | GATAAGGTCATAATG |
| 1 | GMTACGGTCACACTG | chr2R:19783043-19785043 | 1248 | 1262 | + | 9.0071 | 1.80E-05 | 0.546 | GATACAGTTAAACTG |
| 1 | GMTACGGTCACACTG | chr3L:19927508-19929508 | 1252 | 1266 | + | 9.0071 | 1.80E-05 | 0.546 | CATACGTTCACACTT |
| 1 | GMTACGGTCACACTG | chrX:5775647-5777647 | 1269 | 1283 | + | 9.0071 | 1.80E-05 | 0.546 | TATACTGTCATACTG |
| 1 | GMTACGGTCACACTG | chrX:3870939-3872939 | 1302 | 1316 | + | 9.0071 | 1.80E-05 | 0.546 | GATACGATAACATTG |
| 1 | GMTACGGTCACACTG | chr3L:8787545-8789545 | 1311 | 1325 | + | 9.0071 | 1.80E-05 | 0.546 | GATAAGGTCAGACTA |
| 1 | GMTACGGTCACACTG | chr2L:10330116-10332116 | 1370 | 1384 | + | 9.0071 | 1.80E-05 | 0.546 | GATATGGTAACACTT |

| | | | | | | | | | |
|---|---|---|---|---|---|---|---|---|---|
| 1 | GMTACGGTCACACTG | chr2R:15240589-15242589 | 1386 | 1400 | + | 9.0071 | 1.80E-05 | 0.546 | TATACTGTAACACTG |
| 1 | GMTACGGTCACACTG | chr2L:6307819-6309819 | 1466 | 1480 | + | 9.0071 | 1.80E-05 | 0.546 | GATACTGTTACAGTG |
| 1 | GMTACGGTCACACTG | chr2L:16836972-16838972 | 1534 | 1548 | + | 9.0071 | 1.80E-05 | 0.546 | GATATGCTCAAACTG |
| 1 | GMTACGGTCACACTG | chr2L:16836972-16838972 | 1534 | 1548 | + | 9.0071 | 1.80E-05 | 0.546 | GATATGCTCAAACTG |
| 1 | GMTACGGTCACACTG | chr2L:6307751-6309751 | 1534 | 1548 | + | 9.0071 | 1.80E-05 | 0.546 | GATACTGTTACAGTG |
| 1 | GMTACGGTCACACTG | chr3L:22895127-22897127 | 1552 | 1566 | + | 9.0071 | 1.80E-05 | 0.546 | GATACGGTTAAAGTG |
| 1 | GMTACGGTCACACTG | chr3L:1940068-1942068 | 1655 | 1669 | + | 9.0071 | 1.80E-05 | 0.546 | AATAAGATCACACTG |
| 1 | GMTACGGTCACACTG | chr3L:16496651-16498651 | 1687 | 1701 | + | 9.0071 | 1.80E-05 | 0.546 | GATAAGATTACACTG |
| 1 | GMTACGGTCACACTG | chrX:7900331-7902331 | 1757 | 1771 | + | 9.0071 | 1.80E-05 | 0.546 | GATAAAGTCAAACTG |
| 1 | GMTACGGTCACACTG | chr3L:5764046-5766046 | 1767 | 1781 | + | 9.0071 | 1.80E-05 | 0.546 | GATACCATCACAGTG |
| 1 | GMTACGGTCACACTG | chrX:7900302-7902302 | 1786 | 1800 | + | 9.0071 | 1.80E-05 | 0.546 | GATAAAGTCAAACTG |
| 1 | GMTACGGTCACACTG | chr3R:17148667-17150667 | 1883 | 1897 | + | 9.0071 | 1.80E-05 | 0.546 | TATATGGTCATACTG |
| 1 | GMTACGGTCACACTG | chrX:7900203-7902203 | 1885 | 1899 | + | 9.0071 | 1.80E-05 | 0.546 | GATAAAGTCAAACTG |
| 1 | GMTACGGTCACACTG | chr3R:31616977-31618977 | 75 | 89 | - | 4.6143 | 1.85E-05 | 0.551 | GTGCCGGCCACACTG |
| 1 | GMTACGGTCACACTG | chr2R:12325360-12327360 | 79 | 93 | - | 4.6143 | 1.85E-05 | 0.551 | GTTACGGCCACGCGG |
| 1 | GMTACGGTCACACTG | chr2R:12325258-12327258 | 181 | 195 | - | 4.6143 | 1.85E-05 | 0.551 | GTTACGGCCACGCGG |
| 1 | GMTACGGTCACACTG | chr3R:20733748-20735748 | 273 | 287 | - | 4.6143 | 1.85E-05 | 0.551 | GTTACGGCCACTCCG |
| 1 | GMTACGGTCACACTG | chr2R:14617771-14619771 | 384 | 398 | - | 4.6143 | 1.85E-05 | 0.551 | GGAACGGCCACACCG |
| 1 | GMTACGGTCACACTG | chr2L:285383-287383 | 388 | 402 | - | 4.6143 | 1.85E-05 | 0.551 | GGCTCGGGCACACTG |
| 1 | GMTACGGTCACACTG | chr3R:14654716-14656716 | 399 | 413 | - | 4.6143 | 1.85E-05 | 0.551 | GTTTCGGCCCCACTG |
| 1 | GMTACGGTCACACTG | chr2L:18091256-18093256 | 400 | 414 | - | 4.6143 | 1.85E-05 | 0.551 | GTTCCGGTCCCGCTG |
| 1 | GMTACGGTCACACTG | chrX:19642327-19644327 | 453 | 467 | - | 4.6143 | 1.85E-05 | 0.551 | GGCGCGGACACACTG |
| 1 | GMTACGGTCACACTG | chr2L:6045050-6047050 | 574 | 588 | - | 4.6143 | 1.85E-05 | 0.551 | GGCACGGACCCACTG |
| 1 | GMTACGGTCACACTG | chr3R:10042270-10044270 | 620 | 634 | - | 4.6143 | 1.85E-05 | 0.551 | GGCGCGGCCACACTG |
| 1 | GMTACGGTCACACTG | chrX:19641869-19643869 | 911 | 925 | - | 4.6143 | 1.85E-05 | 0.551 | GGCGCGGACACACTG |
| 1 | GMTACGGTCACACTG | chr2L:6044713-6046713 | 911 | 925 | - | 4.6143 | 1.85E-05 | 0.551 | GGCACGGACCCACTG |
| 1 | GMTACGGTCACACTG | chr2R:22798099-22800099 | 938 | 952 | - | 4.6143 | 1.85E-05 | 0.551 | GGTGCGGGCACACCG |
| 1 | GMTACGGTCACACTG | chr3R:10041895-10043895 | 995 | 1009 | - | 4.6143 | 1.85E-05 | 0.551 | GGCGCGGCCACACTG |
| 1 | GMTACGGTCACACTG | chr2R:14617118-14619118 | 1037 | 1051 | - | 4.6143 | 1.85E-05 | 0.551 | GGAACGGCCACACCG |
| 1 | GMTACGGTCACACTG | chrX:14265367-14267367 | 1039 | 1053 | - | 4.6143 | 1.85E-05 | 0.551 | GTTACGGCCACTCAG |
| 1 | GMTACGGTCACACTG | chrX:14922493-14924493 | 1140 | 1154 | - | 4.6143 | 1.85E-05 | 0.551 | GTAACGGTCCCGCTG |
| 1 | GMTACGGTCACACTG | chrX:14265250-14267250 | 1156 | 1170 | - | 4.6143 | 1.85E-05 | 0.551 | GTTACGGCCACTCAG |
| 1 | GMTACGGTCACACTG | chr3R:11215535-11217535 | 1165 | 1179 | - | 4.6143 | 1.85E-05 | 0.551 | GGAACGGACGCACTG |

| | | | | | | | | | |
|---|---|---|---|---|---|---|---|---|---|
| 1 | GMTACGGTCACACTG | chr2R:12152071-12154071 | 1200 | 1214 | - | 4.6143 | 1.85E-05 | 0.551 | GTTTCGGGCACACGG |
| 1 | GMTACGGTCACACTG | chr2L:6716625-6718625 | 1296 | 1310 | - | 4.6143 | 1.85E-05 | 0.551 | GTAGCGGTCACTCTG |
| 1 | GMTACGGTCACACTG | chr3R:11215287-11217287 | 1413 | 1427 | - | 4.6143 | 1.85E-05 | 0.551 | GGAACGGACGCACTG |
| 1 | GMTACGGTCACACTG | chr3R:15139121-15141121 | 1424 | 1438 | - | 4.6143 | 1.85E-05 | 0.551 | GGTCCGGGCACACAG |
| 1 | GMTACGGTCACACTG | chr3L:5712331-5714331 | 1519 | 1533 | - | 4.6143 | 1.85E-05 | 0.551 | GTCACGGTCTCGCTG |
| 1 | GMTACGGTCACACTG | chr3R:15138958-15140958 | 1587 | 1601 | - | 4.6143 | 1.85E-05 | 0.551 | GGTCCGGGCACACAG |
| 1 | GMTACGGTCACACTG | chr2R:22797289-22799289 | 1748 | 1762 | - | 4.6143 | 1.85E-05 | 0.551 | GGTGCGGGCACACCG |
| 1 | GMTACGGTCACACTG | chr3R:31615303-31617303 | 1749 | 1763 | - | 4.6143 | 1.85E-05 | 0.551 | GTGCCGGCCACACTG |
| 1 | GMTACGGTCACACTG | chr3R:31615160-31617160 | 1892 | 1906 | - | 4.6143 | 1.85E-05 | 0.551 | GTGCCGGCCACACTG |
| 1 | GMTACGGTCACACTG | chr2R:19095174-19097174 | 195 | 209 | + | 4.6143 | 1.85E-05 | 0.551 | GGGCCGGACACACTG |
| 1 | GMTACGGTCACACTG | chr3R:3586054-3588054 | 199 | 213 | + | 4.6143 | 1.85E-05 | 0.551 | GGGTCGGTCCCACTG |
| 1 | GMTACGGTCACACTG | chr2R:5979778-5981778 | 272 | 286 | + | 4.6143 | 1.85E-05 | 0.551 | GTTCCGGGCCCACTG |
| 1 | GMTACGGTCACACTG | chr3L:5602979-5604979 | 599 | 613 | + | 4.6143 | 1.85E-05 | 0.551 | GGTACGGGCCCACCG |
| 1 | GMTACGGTCACACTG | chrX:6300285-6302285 | 826 | 840 | + | 4.6143 | 1.85E-05 | 0.551 | GGTGCGGGCGCACTG |
| 1 | GMTACGGTCACACTG | chr3L:13449770-13451770 | 955 | 969 | + | 4.6143 | 1.85E-05 | 0.551 | GGCGCGGTCCCACTG |
| 1 | GMTACGGTCACACTG | chr3R:5127087-5129087 | 1011 | 1025 | + | 4.6143 | 1.85E-05 | 0.551 | GGCGCGGCCACACTG |
| 1 | GMTACGGTCACACTG | chr2L:8029721-8031721 | 1020 | 1034 | + | 4.6143 | 1.85E-05 | 0.551 | GGCGCGGTCACGCTG |
| 1 | GMTACGGTCACACTG | chrX:4548070-4550070 | 1064 | 1078 | + | 4.6143 | 1.85E-05 | 0.551 | GTAACGGACACGCTG |
| 1 | GMTACGGTCACACTG | chr3L:2275550-2277550 | 1318 | 1332 | + | 4.6143 | 1.85E-05 | 0.551 | GTTCCGGCCACGCTG |
| 1 | GMTACGGTCACACTG | chrX:4547794-4549794 | 1340 | 1354 | + | 4.6143 | 1.85E-05 | 0.551 | GTAACGGACACGCTG |
| 1 | GMTACGGTCACACTG | chr3L:13449283-13451283 | 1442 | 1456 | + | 4.6143 | 1.85E-05 | 0.551 | GGCGCGGTCCCACTG |
| 1 | GMTACGGTCACACTG | chr2R:13410739-13412739 | 1542 | 1556 | + | 4.6143 | 1.85E-05 | 0.551 | GTCCCGGTCGCACTG |
| 1 | GMTACGGTCACACTG | chr2R:13406011-13408011 | 1543 | 1557 | + | 4.6143 | 1.85E-05 | 0.551 | GTCCCGGTCGCACTG |
| 1 | GMTACGGTCACACTG | chr2R:13553077-13555077 | 1557 | 1571 | + | 4.6143 | 1.85E-05 | 0.551 | GTGGCGGACACACTG |
| 1 | GMTACGGTCACACTG | chr2L:10422291-10424291 | 1590 | 1604 | + | 4.6143 | 1.85E-05 | 0.551 | GGAACGGTCACCCAG |
| 1 | GMTACGGTCACACTG | chr3R:30882503-30884503 | 1823 | 1837 | + | 4.6143 | 1.85E-05 | 0.551 | GTGACGGCCACGCTG |
| 1 | GMTACGGTCACACTG | chr3R:16338575-16340575 | 3 | 17 | - | 4.3357 | 2.26E-05 | 0.59 | AGCACGGCCACACTG |
| 1 | GMTACGGTCACACTG | chr3R:4185963-4187963 | 7 | 21 | - | 4.3357 | 2.26E-05 | 0.59 | GTAACGGTCACAAAG |
| 1 | GMTACGGTCACACTG | chr2R:22419376-22421376 | 60 | 74 | - | 4.3357 | 2.26E-05 | 0.59 | GTCACGCTCACACAG |
| 1 | GMTACGGTCACACTG | chr2R:16571082-16573082 | 155 | 169 | - | 4.3357 | 2.26E-05 | 0.59 | TGCGCGGTCACACTG |
| 1 | GMTACGGTCACACTG | chr2R:6614752-6616752 | 196 | 210 | - | 4.3357 | 2.26E-05 | 0.59 | GGGACGGGCAAACTG |
| 1 | GMTACGGTCACACTG | chr2L:18968028-18970028 | 244 | 258 | - | 4.3357 | 2.26E-05 | 0.59 | GTCACGGTCACCGTG |
| 1 | GMTACGGTCACACTG | chr2R:22126808-22128808 | 266 | 280 | - | 4.3357 | 2.26E-05 | 0.59 | GGAACGGTCAGACAG |

| | | | | | | | | | |
|---|---|---|---|---|---|---|---|---|---|
| 1 | GMTACGGTCACACTG | chr2L:14348070-14350070 | 292 | 306 | - | 4.3357 | 2.26E-05 | 0.59 | GTCATGGACACACTG |
| 1 | GMTACGGTCACACTG | chr2L:2752068-2754068 | 348 | 362 | - | 4.3357 | 2.26E-05 | 0.59 | GGTGCAGTCACTCTG |
| 1 | GMTACGGTCACACTG | chrX:4803862-4805862 | 362 | 376 | - | 4.3357 | 2.26E-05 | 0.59 | GTGACGGCCACACTT |
| 1 | GMTACGGTCACACTG | chr3L:7311896-7313896 | 447 | 461 | - | 4.3357 | 2.26E-05 | 0.59 | AGGACGGACACACTG |
| 1 | GMTACGGTCACACTG | chr3L:14801168-14803168 | 461 | 475 | - | 4.3357 | 2.26E-05 | 0.59 | GGTACAGTCGCCCTG |
| 1 | GMTACGGTCACACTG | chrX:18439581-18441581 | 467 | 481 | - | 4.3357 | 2.26E-05 | 0.59 | GTTGCGGACACACTC |
| 1 | GMTACGGTCACACTG | chr3R:30038719-30040719 | 468 | 482 | - | 4.3357 | 2.26E-05 | 0.59 | GTTACTGTCCCACAG |
| 1 | GMTACGGTCACACTG | chr3R:30038719-30040719 | 468 | 482 | - | 4.3357 | 2.26E-05 | 0.59 | GTTACTGTCCCACAG |
| 1 | GMTACGGTCACACTG | chr3R:4196206-4198206 | 493 | 507 | - | 4.3357 | 2.26E-05 | 0.59 | GTCACGGCCACACTT |
| 1 | GMTACGGTCACACTG | chr3L:1638711-1640711 | 497 | 511 | - | 4.3357 | 2.26E-05 | 0.59 | TGCACGGCCACACTG |
| 1 | GMTACGGTCACACTG | chr2L:11943129-11945129 | 504 | 518 | - | 4.3357 | 2.26E-05 | 0.59 | GGAACGGTCACCCTA |
| 1 | GMTACGGTCACACTG | chr2L:18967744-18969744 | 528 | 542 | - | 4.3357 | 2.26E-05 | 0.59 | GTCACGGTCACCGTG |
| 1 | GMTACGGTCACACTG | chr2R:9206561-9208561 | 540 | 554 | - | 4.3357 | 2.26E-05 | 0.59 | AGCACGGCCACACTG |
| 1 | GMTACGGTCACACTG | chr3L:704330-706330 | 564 | 578 | - | 4.3357 | 2.26E-05 | 0.59 | GTTACTGCCACACGG |
| 1 | GMTACGGTCACACTG | chr2L:18986707-18988707 | 572 | 586 | - | 4.3357 | 2.26E-05 | 0.59 | GTCACGGTCGCACTT |
| 1 | GMTACGGTCACACTG | chr2L:14347790-14349790 | 572 | 586 | - | 4.3357 | 2.26E-05 | 0.59 | GTCATGGACACACTG |
| 1 | GMTACGGTCACACTG | chr2R:22126502-22128502 | 572 | 586 | - | 4.3357 | 2.26E-05 | 0.59 | GGAACGGTCAGACAG |
| 1 | GMTACGGTCACACTG | chr3L:11063991-11065991 | 577 | 591 | - | 4.3357 | 2.26E-05 | 0.59 | GTTCCTGTCACACCG |
| 1 | GMTACGGTCACACTG | chr3L:11063991-11065991 | 577 | 591 | - | 4.3357 | 2.26E-05 | 0.59 | GTTCCTGTCACACCG |
| 1 | GMTACGGTCACACTG | chrX:3066562-3068562 | 590 | 604 | - | 4.3357 | 2.26E-05 | 0.59 | GTTACGGGCACATCG |
| 1 | GMTACGGTCACACTG | chrX:22454803-22456803 | 595 | 609 | - | 4.3357 | 2.26E-05 | 0.59 | TGCGCGGTCACACTG |
| 1 | GMTACGGTCACACTG | chr2R:8890832-8892832 | 599 | 613 | - | 4.3357 | 2.26E-05 | 0.59 | GGCACGGTCCTACTG |
| 1 | GMTACGGTCACACTG | chrX:11579772-11581772 | 600 | 614 | - | 4.3357 | 2.26E-05 | 0.59 | GTTCTGGTCACCCTG |
| 1 | GMTACGGTCACACTG | chr3L:11063941-11065941 | 627 | 641 | - | 4.3357 | 2.26E-05 | 0.59 | GTTCCTGTCACACCG |
| 1 | GMTACGGTCACACTG | chr2L:3293706-3295706 | 639 | 653 | - | 4.3357 | 2.26E-05 | 0.59 | GTTCCGGGCAAACTG |
| 1 | GMTACGGTCACACTG | chr3R:30038533-30040533 | 654 | 668 | - | 4.3357 | 2.26E-05 | 0.59 | GTTACTGTCCCACAG |
| 1 | GMTACGGTCACACTG | chr3L:21087989-21089989 | 657 | 671 | - | 4.3357 | 2.26E-05 | 0.59 | GTTACGGACACGGTG |
| 1 | GMTACGGTCACACTG | chrX:16733947-16735947 | 659 | 673 | - | 4.3357 | 2.26E-05 | 0.59 | GGTGCGGTGACCCTG |
| 1 | GMTACGGTCACACTG | chr2L:16302264-16304264 | 659 | 673 | - | 4.3357 | 2.26E-05 | 0.59 | TGAACGGTCTCACTG |
| 1 | GMTACGGTCACACTG | chr2L:8437169-8439169 | 686 | 700 | - | 4.3357 | 2.26E-05 | 0.59 | GTTGCGGTGGCACTG |
| 1 | GMTACGGTCACACTG | chrX:20734960-20736960 | 747 | 761 | - | 4.3357 | 2.26E-05 | 0.59 | CGTACGGACCCACTG |
| 1 | GMTACGGTCACACTG | chr2R:23687077-23689077 | 764 | 778 | - | 4.3357 | 2.26E-05 | 0.59 | GTTTCGGTTTCACTG |
| 1 | GMTACGGTCACACTG | chr2L:8863864-8865864 | 780 | 794 | - | 4.3357 | 2.26E-05 | 0.59 | GTTTCGATCACACAG |

| | | | | | | | | | |
|---|---|---|---|---|---|---|---|---|---|
| 1 | GMTACGGTCACACTG | chr3L:7497831-7499831 | 816 | 830 | - | 4.3357 | 2.26E-05 | 0.59 | GTTCCCGGCACACTG |
| 1 | GMTACGGTCACACTG | chrX:1463220-1465220 | 823 | 837 | - | 4.3357 | 2.26E-05 | 0.59 | GTTTCGGACACACTC |
| 1 | GMTACGGTCACACTG | chr2R:11678492-11680492 | 830 | 844 | - | 4.3357 | 2.26E-05 | 0.59 | GGTGTGGCCACACTG |
| 1 | GMTACGGTCACACTG | chr3R:15224576-15226576 | 836 | 850 | - | 4.3357 | 2.26E-05 | 0.59 | GTTGCGGGCACACTA |
| 1 | GMTACGGTCACACTG | chr3R:23759600-23761600 | 838 | 852 | - | 4.3357 | 2.26E-05 | 0.59 | GGTAAGGACCCACTG |
| 1 | GMTACGGTCACACTG | chr3L:6148548-6150548 | 846 | 860 | - | 4.3357 | 2.26E-05 | 0.59 | GTTCCGGTCTAACTG |
| 1 | GMTACGGTCACACTG | chr3L:6145674-6147674 | 849 | 863 | - | 4.3357 | 2.26E-05 | 0.59 | GTTCCGGTCTAACTG |
| 1 | GMTACGGTCACACTG | chr3R:22447099-22449099 | 857 | 871 | - | 4.3357 | 2.26E-05 | 0.59 | CGAACGGTCTCACTG |
| 1 | GMTACGGTCACACTG | chr3R:19863496-19865496 | 893 | 907 | - | 4.3357 | 2.26E-05 | 0.59 | GGTGCGGATACACTG |
| 1 | GMTACGGTCACACTG | chr3R:19094000-19096000 | 893 | 907 | - | 4.3357 | 2.26E-05 | 0.59 | GGGTCGGTCACACTA |
| 1 | GMTACGGTCACACTG | chrX:16349315-16351315 | 911 | 925 | - | 4.3357 | 2.26E-05 | 0.59 | GGGGCGTTCACACTG |
| 1 | GMTACGGTCACACTG | chr3L:8108132-8110132 | 917 | 931 | - | 4.3357 | 2.26E-05 | 0.59 | GGTGTGGACACACTG |
| 1 | GMTACGGTCACACTG | chr2L:10305922-10307922 | 961 | 975 | - | 4.3357 | 2.26E-05 | 0.59 | CTCACGGACACACTG |
| 1 | GMTACGGTCACACTG | chrX:10847757-10849757 | 962 | 976 | - | 4.3357 | 2.26E-05 | 0.59 | AGTTCGGCCACACTG |
| 1 | GMTACGGTCACACTG | chr3R:29490150-29492150 | 965 | 979 | - | 4.3357 | 2.26E-05 | 0.59 | GGCACGGTCACACCA |
| 1 | GMTACGGTCACACTG | chr2R:11678354-11680354 | 968 | 982 | - | 4.3357 | 2.26E-05 | 0.59 | GGTGTGGCCACACTG |
| 1 | GMTACGGTCACACTG | chr2L:16301954-16303954 | 969 | 983 | - | 4.3357 | 2.26E-05 | 0.59 | TGAACGGTCTCACTG |
| 1 | GMTACGGTCACACTG | chr2L:2751445-2753445 | 971 | 985 | - | 4.3357 | 2.26E-05 | 0.59 | GGTGCAGTCACTCTG |
| 1 | GMTACGGTCACACTG | chr3L:1638235-1640235 | 973 | 987 | - | 4.3357 | 2.26E-05 | 0.59 | TGCACGGCCACACTG |
| 1 | GMTACGGTCACACTG | chr3L:9398227-9400227 | 974 | 988 | - | 4.3357 | 2.26E-05 | 0.59 | GGCGCAGTCACACTG |
| 1 | GMTACGGTCACACTG | chr3L:12192688-12194688 | 975 | 989 | - | 4.3357 | 2.26E-05 | 0.59 | GGAACGGTCACTCTA |
| 1 | GMTACGGTCACACTG | chr2L:4457364-4459364 | 975 | 989 | - | 4.3357 | 2.26E-05 | 0.59 | CGGGCGGTCACACTG |
| 1 | GMTACGGTCACACTG | chr3R:9792295-9794295 | 975 | 989 | - | 4.3357 | 2.26E-05 | 0.59 | GGTGCGGTCCTACTG |
| 1 | GMTACGGTCACACTG | chrX:22454422-22456422 | 976 | 990 | - | 4.3357 | 2.26E-05 | 0.59 | TGCGCGGTCACACTG |
| 1 | GMTACGGTCACACTG | chr2R:13493862-13495862 | 976 | 990 | - | 4.3357 | 2.26E-05 | 0.59 | ATCGCGGTCACACTG |
| 1 | GMTACGGTCACACTG | chr2R:9206124-9208124 | 977 | 991 | - | 4.3357 | 2.26E-05 | 0.59 | AGCACGGCCACACTG |
| 1 | GMTACGGTCACACTG | chr3R:9005456-9007456 | 977 | 991 | - | 4.3357 | 2.26E-05 | 0.59 | GGCCTGGTCACACTG |
| 1 | GMTACGGTCACACTG | chr2R:22881690-22883690 | 978 | 992 | - | 4.3357 | 2.26E-05 | 0.59 | GTCGCGGTCACACTT |
| 1 | GMTACGGTCACACTG | chr3L:5560598-5562598 | 979 | 993 | - | 4.3357 | 2.26E-05 | 0.59 | AGTGCGGCCACACTG |
| 1 | GMTACGGTCACACTG | chr3R:5753774-5755774 | 981 | 995 | - | 4.3357 | 2.26E-05 | 0.59 | GTCATGGTCACGCTG |
| 1 | GMTACGGTCACACTG | chr3R:28831024-28833024 | 986 | 1000 | - | 4.3357 | 2.26E-05 | 0.59 | CGGTCGGTCACACTG |
| 1 | GMTACGGTCACACTG | chr3R:19093905-19095905 | 988 | 1002 | - | 4.3357 | 2.26E-05 | 0.59 | GGGTCGGTCACACTA |
| 1 | GMTACGGTCACACTG | chrX:19275898-19277898 | 989 | 1003 | - | 4.3357 | 2.26E-05 | 0.59 | GTGGCGGTCACACTA |

| | | | | | | | | | |
|---|---|---|---|---|---|---|---|---|---|
| 1 | GMTACGGTCACACTG | chr3R:27558145-27560145 | 990 | 1004 | - | 4.3357 | 2.26E-05 | 0.59 | CGAACGGTCACACCG |
| 1 | GMTACGGTCACACTG | chr3R:20815190-20817190 | 992 | 1006 | - | 4.3357 | 2.26E-05 | 0.59 | TGCTCGGTCACACTG |
| 1 | GMTACGGTCACACTG | chrX:1857971-1859971 | 992 | 1006 | - | 4.3357 | 2.26E-05 | 0.59 | CGAGCGGTCACACTG |
| 1 | GMTACGGTCACACTG | chrX:21027361-21029361 | 993 | 1007 | - | 4.3357 | 2.26E-05 | 0.59 | CGCACGGCCACACTG |
| 1 | GMTACGGTCACACTG | chr3R:11973910-11975910 | 995 | 1009 | - | 4.3357 | 2.26E-05 | 0.59 | TGTGCGGTCTCACTG |
| 1 | GMTACGGTCACACTG | chr3L:16086525-16088525 | 995 | 1009 | - | 4.3357 | 2.26E-05 | 0.59 | CGCACGGTCACTCTG |
| 1 | GMTACGGTCACACTG | chr3R:19863393-19865393 | 996 | 1010 | - | 4.3357 | 2.26E-05 | 0.59 | GGTGCGGATACACTG |
| 1 | GMTACGGTCACACTG | chr2R:23861593-23863593 | 997 | 1011 | - | 4.3357 | 2.26E-05 | 0.59 | TGTTCGGCCACACTG |
| 1 | GMTACGGTCACACTG | chr2L:18986280-18988280 | 999 | 1013 | - | 4.3357 | 2.26E-05 | 0.59 | GTCACGGTCGCACTT |
| 1 | GMTACGGTCACACTG | chrX:22539383-22541383 | 1003 | 1017 | - | 4.3357 | 2.26E-05 | 0.59 | TGCACGGCCACACTG |
| 1 | GMTACGGTCACACTG | chr2R:16570230-16572230 | 1007 | 1021 | - | 4.3357 | 2.26E-05 | 0.59 | TGCGCGGTCACACTG |
| 1 | GMTACGGTCACACTG | chr2R:10074645-10076645 | 1012 | 1026 | - | 4.3357 | 2.26E-05 | 0.59 | CGAACGGCCACACTG |
| 1 | GMTACGGTCACACTG | chr2R:9692328-9694328 | 1014 | 1028 | - | 4.3357 | 2.26E-05 | 0.59 | AGGTCGGTCACACTG |
| 1 | GMTACGGTCACACTG | chr2L:11808438-11810438 | 1018 | 1032 | - | 4.3357 | 2.26E-05 | 0.59 | AGCCCGGTCACACTG |
| 1 | GMTACGGTCACACTG | chr2R:8890401-8892401 | 1030 | 1044 | - | 4.3357 | 2.26E-05 | 0.59 | GGCACGGTCCTACTG |
| 1 | GMTACGGTCACACTG | chr2R:6647654-6649654 | 1043 | 1057 | - | 4.3357 | 2.26E-05 | 0.59 | GTTACGGTTCCACGG |
| 1 | GMTACGGTCACACTG | chr3R:15224329-15226329 | 1083 | 1097 | - | 4.3357 | 2.26E-05 | 0.59 | GTTGCGGGCACACTA |
| 1 | GMTACGGTCACACTG | chr2L:1047053-1049053 | 1106 | 1120 | - | 4.3357 | 2.26E-05 | 0.59 | GGTGAGGGCACACTG |
| 1 | GMTACGGTCACACTG | chr3R:10124664-10126664 | 1138 | 1152 | - | 4.3357 | 2.26E-05 | 0.59 | GTTACGGCCACTCTT |
| 1 | GMTACGGTCACACTG | chr3R:16593414-16595414 | 1148 | 1162 | - | 4.3357 | 2.26E-05 | 0.59 | ATAACGGTCACTCTG |
| 1 | GMTACGGTCACACTG | chr2L:7780465-7782465 | 1160 | 1174 | - | 4.3357 | 2.26E-05 | 0.59 | GTCATGGTCACGCTG |
| 1 | GMTACGGTCACACTG | chr2R:11345325-11347325 | 1205 | 1219 | - | 4.3357 | 2.26E-05 | 0.59 | GTCACGATCACCCTG |
| 1 | GMTACGGTCACACTG | chr3R:9005226-9007226 | 1207 | 1221 | - | 4.3357 | 2.26E-05 | 0.59 | GGCCTGGTCACACTG |
| 1 | GMTACGGTCACACTG | chr2R:8086396-8088396 | 1229 | 1243 | - | 4.3357 | 2.26E-05 | 0.59 | GTTACGGTCAGGCAG |
| 1 | GMTACGGTCACACTG | chr3R:21096827-21098827 | 1244 | 1258 | - | 4.3357 | 2.26E-05 | 0.59 | GTGCTGGTCACACTG |
| 1 | GMTACGGTCACACTG | chr3L:16922252-16924252 | 1271 | 1285 | - | 4.3357 | 2.26E-05 | 0.59 | GTTACTGCCACTCTG |
| 1 | GMTACGGTCACACTG | chr2L:10305594-10307594 | 1289 | 1303 | - | 4.3357 | 2.26E-05 | 0.59 | CTCACGGACACACTG |
| 1 | GMTACGGTCACACTG | chr2L:17981472-17983472 | 1307 | 1321 | - | 4.3357 | 2.26E-05 | 0.59 | GGTACGGACAGACAG |
| 1 | GMTACGGTCACACTG | chr3L:10960500-10962500 | 1335 | 1349 | - | 4.3357 | 2.26E-05 | 0.59 | GGTACGGTGGCTCTG |
| 1 | GMTACGGTCACACTG | chr3R:27287769-27289769 | 1343 | 1357 | - | 4.3357 | 2.26E-05 | 0.59 | GGCGGGGTCACACTG |
| 1 | GMTACGGTCACACTG | chrX:19275539-19277539 | 1348 | 1362 | - | 4.3357 | 2.26E-05 | 0.59 | GTGGCGGTCACACTA |
| 1 | GMTACGGTCACACTG | chrX:18438667-18440667 | 1381 | 1395 | - | 4.3357 | 2.26E-05 | 0.59 | GTTGCGGACACACTC |
| 1 | GMTACGGTCACACTG | chr3R:10433789-10435789 | 1387 | 1401 | - | 4.3357 | 2.26E-05 | 0.59 | GTTCCTGCCACACTG |

| | | | | | | | | | |
|---|---|---|---|---|---|---|---|---|---|
| 1 | GMTACGGTCACACTG | chrX:10847757-10849757 | 1390 | 1404 | - | 4.3357 | 2.26E-05 | 0.59 | GTTTCGCTCGCACTG |
| 1 | GMTACGGTCACACTG | chr2R:7418350-7420350 | 1396 | 1410 | - | 4.3357 | 2.26E-05 | 0.59 | GTCGCGATCACACTG |
| 1 | GMTACGGTCACACTG | chr3L:21285099-21287099 | 1403 | 1417 | - | 4.3357 | 2.26E-05 | 0.59 | GGACCGGTCACAGTG |
| 1 | GMTACGGTCACACTG | chr2L:21142854-21144854 | 1423 | 1437 | - | 4.3357 | 2.26E-05 | 0.59 | GTCATGGTCGCACTG |
| 1 | GMTACGGTCACACTG | chr3R:7518456-7520456 | 1439 | 1453 | - | 4.3357 | 2.26E-05 | 0.59 | GGTATGGACACTCTG |
| 1 | GMTACGGTCACACTG | chr2L:8436405-8438405 | 1450 | 1464 | - | 4.3357 | 2.26E-05 | 0.59 | GTTGCGGTGGCACTG |
| 1 | GMTACGGTCACACTG | chr3R:30037728-30039728 | 1459 | 1473 | - | 4.3357 | 2.26E-05 | 0.59 | GTTACTGTCCCACAG |
| 1 | GMTACGGTCACACTG | chr2R:14617771-14619771 | 1460 | 1474 | - | 4.3357 | 2.26E-05 | 0.59 | GGTACGGGACCACTG |
| 1 | GMTACGGTCACACTG | chr2R:11345066-11347066 | 1464 | 1478 | - | 4.3357 | 2.26E-05 | 0.59 | GTCACGATCACCCTG |
| 1 | GMTACGGTCACACTG | chr3L:9728999-9730999 | 1467 | 1481 | - | 4.3357 | 2.26E-05 | 0.59 | GGTCCGGTGACACCG |
| 1 | GMTACGGTCACACTG | chr2L:8436365-8438365 | 1490 | 1504 | - | 4.3357 | 2.26E-05 | 0.59 | GTTGCGGTGGCACTG |
| 1 | GMTACGGTCACACTG | chr2R:18159720-18161720 | 1506 | 1520 | - | 4.3357 | 2.26E-05 | 0.59 | GTTGCGGTAGCACTG |
| 1 | GMTACGGTCACACTG | chr2R:7782334-7784334 | 1509 | 1523 | - | 4.3357 | 2.26E-05 | 0.59 | TTCACGGTCACGCTG |
| 1 | GMTACGGTCACACTG | chr3R:11677116-11679116 | 1517 | 1531 | - | 4.3357 | 2.26E-05 | 0.59 | GGTACGGACGAACTG |
| 1 | GMTACGGTCACACTG | chr3L:21284962-21286962 | 1540 | 1554 | - | 4.3357 | 2.26E-05 | 0.59 | GGACCGGTCACAGTG |
| 1 | GMTACGGTCACACTG | chr2R:13433207-13435207 | 1551 | 1565 | - | 4.3357 | 2.26E-05 | 0.59 | GGAACTGCCACACTG |
| 1 | GMTACGGTCACACTG | chr2R:18019268-18021268 | 1581 | 1595 | - | 4.3357 | 2.26E-05 | 0.59 | GGTAAGGACGCACTG |
| 1 | GMTACGGTCACACTG | chr2L:15284000-15286000 | 1584 | 1598 | - | 4.3357 | 2.26E-05 | 0.59 | GTGCTGGTCACACTG |
| 1 | GMTACGGTCACACTG | chr3R:27287526-27289526 | 1586 | 1600 | - | 4.3357 | 2.26E-05 | 0.59 | GGCGGGGTCACACTG |
| 1 | GMTACGGTCACACTG | chrX:9607946-9609946 | 1589 | 1603 | - | 4.3357 | 2.26E-05 | 0.59 | GGTAAGGGCACTCTG |
| 1 | GMTACGGTCACACTG | chr3R:17608359-17610359 | 1605 | 1619 | - | 4.3357 | 2.26E-05 | 0.59 | CGTACGGACGCACTG |
| 1 | GMTACGGTCACACTG | chr2R:8892613-8894613 | 1614 | 1628 | - | 4.3357 | 2.26E-05 | 0.59 | GGTGTGGTCCCACTG |
| 1 | GMTACGGTCACACTG | chr3R:19093270-19095270 | 1623 | 1637 | - | 4.3357 | 2.26E-05 | 0.59 | GGGTCGGTCACACTA |
| 1 | GMTACGGTCACACTG | chrX:9607884-9609884 | 1651 | 1665 | - | 4.3357 | 2.26E-05 | 0.59 | GGTAAGGGCACTCTG |
| 1 | GMTACGGTCACACTG | chr2L:4456674-4458674 | 1665 | 1679 | - | 4.3357 | 2.26E-05 | 0.59 | CGGGCGGTCACACTG |
| 1 | GMTACGGTCACACTG | chr3R:22446263-22448263 | 1693 | 1707 | - | 4.3357 | 2.26E-05 | 0.59 | CGAACGGTCTCACTG |
| 1 | GMTACGGTCACACTG | chr2L:13434622-13436622 | 1696 | 1710 | - | 4.3357 | 2.26E-05 | 0.59 | GTCACGTTCACGCTG |
| 1 | GMTACGGTCACACTG | chr2R:7392161-7394161 | 1724 | 1738 | - | 4.3357 | 2.26E-05 | 0.59 | GTTTCGGACACATTG |
| 1 | GMTACGGTCACACTG | chr3L:4147696-4149696 | 1813 | 1827 | - | 4.3357 | 2.26E-05 | 0.59 | GTACCGGTCACAATG |
| 1 | GMTACGGTCACACTG | chr2L:8862811-8864811 | 1833 | 1847 | - | 4.3357 | 2.26E-05 | 0.59 | GTTTCGATCACACAG |
| 1 | GMTACGGTCACACTG | chr3R:11327480-11329480 | 1859 | 1873 | - | 4.3357 | 2.26E-05 | 0.59 | GTTACGGACTGACTG |
| 1 | GMTACGGTCACACTG | chr2R:12205092-12207092 | 1951 | 1965 | - | 4.3357 | 2.26E-05 | 0.59 | GTCCCGGTCACATTG |
| 1 | GMTACGGTCACACTG | chr3R:11235286-11237286 | 17 | 31 | + | 4.3357 | 2.26E-05 | 0.59 | GTTGCGGTCCCACTA |

| | | | | | | | | | |
|---|---|---|---|---|---|---|---|---|---|
| 1 | GMTACGGTCACACTG | chr3R:14307589-14309589 | 26 | 40 | + | 4.3357 | 2.26E-05 | 0.59 | GTCACGGTAGCACTG |
| 1 | GMTACGGTCACACTG | chr2R:13963653-13965653 | 30 | 44 | + | 4.3357 | 2.26E-05 | 0.59 | CGCTCGGTCACACTG |
| 1 | GMTACGGTCACACTG | chr3L:9718144-9720144 | 52 | 66 | + | 4.3357 | 2.26E-05 | 0.59 | GTTACGGTCGCCGTG |
| 1 | GMTACGGTCACACTG | chrX:19189214-19191214 | 92 | 106 | + | 4.3357 | 2.26E-05 | 0.59 | GTGACGGTCACTGTG |
| 1 | GMTACGGTCACACTG | chr2L:2239593-2241593 | 197 | 211 | + | 4.3357 | 2.26E-05 | 0.59 | GGTACGGCCACCATG |
| 1 | GMTACGGTCACACTG | chr3R:28879122-28881122 | 232 | 246 | + | 4.3357 | 2.26E-05 | 0.59 | GTTACGGCCACTTTG |
| 1 | GMTACGGTCACACTG | chr3R:14150940-14152940 | 245 | 259 | + | 4.3357 | 2.26E-05 | 0.59 | GTTACGGGCGCACTC |
| 1 | GMTACGGTCACACTG | chr2L:20933113-20935113 | 249 | 263 | + | 4.3357 | 2.26E-05 | 0.59 | GGTAAGGTCGCTCTG |
| 1 | GMTACGGTCACACTG | chr2L:19489262-19491262 | 251 | 265 | + | 4.3357 | 2.26E-05 | 0.59 | GGCGCGGTCAAACTG |
| 1 | GMTACGGTCACACTG | chr2R:7784067-7786067 | 298 | 312 | + | 4.3357 | 2.26E-05 | 0.59 | CTTACGGCCACGCTG |
| 1 | GMTACGGTCACACTG | chr4:426331-428331 | 310 | 324 | + | 4.3357 | 2.26E-05 | 0.59 | GGTAGGGACACTCTG |
| 1 | GMTACGGTCACACTG | chr3L:16287422-16289422 | 339 | 353 | + | 4.3357 | 2.26E-05 | 0.59 | GTTCTGGGCACACTG |
| 1 | GMTACGGTCACACTG | chr3L:22266382-22268382 | 345 | 359 | + | 4.3357 | 2.26E-05 | 0.59 | GTTCCGGCCAAACTG |
| 1 | GMTACGGTCACACTG | chr3R:18725434-18727434 | 351 | 365 | + | 4.3357 | 2.26E-05 | 0.59 | GGTACGGGCACCGTG |
| 1 | GMTACGGTCACACTG | chr2L:19489104-19491104 | 409 | 423 | + | 4.3357 | 2.26E-05 | 0.59 | GGCGCGGTCAAACTG |
| 1 | GMTACGGTCACACTG | chr3L:15975870-15977870 | 422 | 436 | + | 4.3357 | 2.26E-05 | 0.59 | GGAACGGAGACACTG |
| 1 | GMTACGGTCACACTG | chrX:10767537-10769537 | 439 | 453 | + | 4.3357 | 2.26E-05 | 0.59 | CTTACGGGCACACGG |
| 1 | GMTACGGTCACACTG | chr2R:7578929-7580929 | 439 | 453 | + | 4.3357 | 2.26E-05 | 0.59 | TTTTCGGCCACACTG |
| 1 | GMTACGGTCACACTG | chrX:11763590-11765590 | 440 | 454 | + | 4.3357 | 2.26E-05 | 0.59 | TGTCCGGCCACACTG |
| 1 | GMTACGGTCACACTG | chr3R:9411548-9413548 | 443 | 457 | + | 4.3357 | 2.26E-05 | 0.59 | GGTACGGACATTCTG |
| 1 | GMTACGGTCACACTG | chr3R:8561673-8563673 | 470 | 484 | + | 4.3357 | 2.26E-05 | 0.59 | GGAACGGACATACTG |
| 1 | GMTACGGTCACACTG | chr3R:29458278-29460278 | 473 | 487 | + | 4.3357 | 2.26E-05 | 0.59 | TTTTCGGACACACTG |
| 1 | GMTACGGTCACACTG | chrX:17077836-17079836 | 479 | 493 | + | 4.3357 | 2.26E-05 | 0.59 | ATTTCGGGCACACTG |
| 1 | GMTACGGTCACACTG | chr3L:9385112-9387112 | 486 | 500 | + | 4.3357 | 2.26E-05 | 0.59 | GTTCCGCGCACACTG |
| 1 | GMTACGGTCACACTG | chr3L:9385112-9387112 | 486 | 500 | + | 4.3357 | 2.26E-05 | 0.59 | GTTCCGCGCACACTG |
| 1 | GMTACGGTCACACTG | chr3L:7362598-7364598 | 494 | 508 | + | 4.3357 | 2.26E-05 | 0.59 | GTCACGGTTACGCTG |
| 1 | GMTACGGTCACACTG | chr3R:21295127-21297127 | 502 | 516 | + | 4.3357 | 2.26E-05 | 0.59 | GTCACGGCCACACTT |
| 1 | GMTACGGTCACACTG | chr3R:6405855-6407855 | 517 | 531 | + | 4.3357 | 2.26E-05 | 0.59 | GGAATGGGCACACTG |
| 1 | GMTACGGTCACACTG | chr2R:24350389-24352389 | 526 | 540 | + | 4.3357 | 2.26E-05 | 0.59 | GTTGCGTTCACACCG |
| 1 | GMTACGGTCACACTG | chr3R:18572091-18574091 | 527 | 541 | + | 4.3357 | 2.26E-05 | 0.59 | GGTACGGTGACTCAG |
| 1 | GMTACGGTCACACTG | chr3R:18572091-18574091 | 527 | 541 | + | 4.3357 | 2.26E-05 | 0.59 | GGTACGGTGACTCAG |
| 1 | GMTACGGTCACACTG | chr2R:11356650-11358650 | 538 | 552 | + | 4.3357 | 2.26E-05 | 0.59 | GTTGGGGCCACACTG |
| 1 | GMTACGGTCACACTG | chr2L:6917743-6919743 | 539 | 553 | + | 4.3357 | 2.26E-05 | 0.59 | GTTGCAGACACACTG |

| | | | | | | | | | |
|---|---|---|---|---|---|---|---|---|---|
| 1 | GMTACGGTCACACTG | chr3R:8561593-8563593 | 550 | 564 | + | 4.3357 | 2.26E-05 | 0.59 | GGAACGGACATACTG |
| 1 | GMTACGGTCACACTG | chr3R:10876783-10878783 | 569 | 583 | + | 4.3357 | 2.26E-05 | 0.59 | GGTTTGGTCACCCTG |
| 1 | GMTACGGTCACACTG | chr3L:2599331-2601331 | 580 | 594 | + | 4.3357 | 2.26E-05 | 0.59 | GTTACGGCCAACCTG |
| 1 | GMTACGGTCACACTG | chr2L:6036015-6038015 | 592 | 606 | + | 4.3357 | 2.26E-05 | 0.59 | GGTACGGACACACGT |
| 1 | GMTACGGTCACACTG | chr2L:8082420-8084420 | 615 | 629 | + | 4.3357 | 2.26E-05 | 0.59 | TTAACGGTCGCACTG |
| 1 | GMTACGGTCACACTG | chr2L:6479930-6481930 | 616 | 630 | + | 4.3357 | 2.26E-05 | 0.59 | CGACCGGTCACACTG |
| 1 | GMTACGGTCACACTG | chr2L:6479929-6481929 | 617 | 631 | + | 4.3357 | 2.26E-05 | 0.59 | CGACCGGTCACACTG |
| 1 | GMTACGGTCACACTG | chrX:6049828-6051828 | 620 | 634 | + | 4.3357 | 2.26E-05 | 0.59 | GTGTTGGTCACACTG |
| 1 | GMTACGGTCACACTG | chrX:10767338-10769338 | 638 | 652 | + | 4.3357 | 2.26E-05 | 0.59 | CTTACGGGCACACGG |
| 1 | GMTACGGTCACACTG | chr2R:21177636-21179636 | 660 | 674 | + | 4.3357 | 2.26E-05 | 0.59 | GGTTCGGGCACATTG |
| 1 | GMTACGGTCACACTG | chrX:1353014-1355014 | 668 | 682 | + | 4.3357 | 2.26E-05 | 0.59 | GTTGCGCTCCCACTG |
| 1 | GMTACGGTCACACTG | chr3R:9411293-9413293 | 698 | 712 | + | 4.3357 | 2.26E-05 | 0.59 | GGTACGGACATTCTG |
| 1 | GMTACGGTCACACTG | chrX:1352981-1354981 | 701 | 715 | + | 4.3357 | 2.26E-05 | 0.59 | GTTGCGCTCCCACTG |
| 1 | GMTACGGTCACACTG | chr3R:12374733-12376733 | 715 | 729 | + | 4.3357 | 2.26E-05 | 0.59 | CGATCGGTCACACTG |
| 1 | GMTACGGTCACACTG | chr3L:13386907-13388907 | 727 | 741 | + | 4.3357 | 2.26E-05 | 0.59 | GTAACGTTCGCACTG |
| 1 | GMTACGGTCACACTG | chr3L:2881814-2883814 | 730 | 744 | + | 4.3357 | 2.26E-05 | 0.59 | GGTTCCGTCCCACTG |
| 1 | GMTACGGTCACACTG | chr3L:11952317-11954317 | 740 | 754 | + | 4.3357 | 2.26E-05 | 0.59 | GTTCCGGTCACAGCG |
| 1 | GMTACGGTCACACTG | chr3R:26020823-26022823 | 746 | 760 | + | 4.3357 | 2.26E-05 | 0.59 | GTGATGGTCCCACTG |
| 1 | GMTACGGTCACACTG | chr3L:3318555-3320555 | 758 | 772 | + | 4.3357 | 2.26E-05 | 0.59 | GTGCTGGTCACACTG |
| 1 | GMTACGGTCACACTG | chr3L:3318555-3320555 | 758 | 772 | + | 4.3357 | 2.26E-05 | 0.59 | GTGCTGGTCACACTG |
| 1 | GMTACGGTCACACTG | chr3R:27844089-27846089 | 798 | 812 | + | 4.3357 | 2.26E-05 | 0.59 | GTTGCGGTCACAAAG |
| 1 | GMTACGGTCACACTG | chr3R:31261598-31263598 | 802 | 816 | + | 4.3357 | 2.26E-05 | 0.59 | GTTACGGGCTCAGTG |
| 1 | GMTACGGTCACACTG | chrX:7963024-7965024 | 817 | 831 | + | 4.3357 | 2.26E-05 | 0.59 | GTTACGGTCACCGAG |
| 1 | GMTACGGTCACACTG | chr3R:10387258-10389258 | 833 | 847 | + | 4.3357 | 2.26E-05 | 0.59 | GTTTCTGTCCCACTG |
| 1 | GMTACGGTCACACTG | chr3R:22033455-22035455 | 853 | 867 | + | 4.3357 | 2.26E-05 | 0.59 | GTAACGGTCTCACTA |
| 1 | GMTACGGTCACACTG | chr2R:14218898-14220898 | 862 | 876 | + | 4.3357 | 2.26E-05 | 0.59 | GTTGCGCTCACACAG |
| 1 | GMTACGGTCACACTG | chr3R:12228413-12230413 | 874 | 888 | + | 4.3357 | 2.26E-05 | 0.59 | GGAAAGGTCACACCG |
| 1 | GMTACGGTCACACTG | chr3L:15746403-15748403 | 883 | 897 | + | 4.3357 | 2.26E-05 | 0.59 | GGTCCTGTCCCACTG |
| 1 | GMTACGGTCACACTG | chr3L:2272744-2274744 | 884 | 898 | + | 4.3357 | 2.26E-05 | 0.59 | GGAACAGTCACTCTG |
| 1 | GMTACGGTCACACTG | chr3R:23140906-23142906 | 894 | 908 | + | 4.3357 | 2.26E-05 | 0.59 | GTTGCGCTCGCACTG |
| 1 | GMTACGGTCACACTG | chr3R:20634480-20636480 | 914 | 928 | + | 4.3357 | 2.26E-05 | 0.59 | ATTTCGGTCACACAG |
| 1 | GMTACGGTCACACTG | chrX:9873057-9875057 | 920 | 934 | + | 4.3357 | 2.26E-05 | 0.59 | GTTTCAGTCGCACTG |
| 1 | GMTACGGTCACACTG | chr3R:24294078-24296078 | 926 | 940 | + | 4.3357 | 2.26E-05 | 0.59 | GGACTGGTCACACTG |

| | | | | | | | | | |
|---|---|---|---|---|---|---|---|---|---|
| 1 | GMTACGGTCACACTG | chrX:1482479-1484479 | 933 | 947 | + | 4.3357 | 2.26E-05 | 0.59 | AGAGCGGTCACACTG |
| 1 | GMTACGGTCACACTG | chr3R:18983025-18985025 | 934 | 948 | + | 4.3357 | 2.26E-05 | 0.59 | CGAACGGTCACACGG |
| 1 | GMTACGGTCACACTG | chr3R:8008400-8010400 | 963 | 977 | + | 4.3357 | 2.26E-05 | 0.59 | GTAAGGGTCACCCTG |
| 1 | GMTACGGTCACACTG | chr2L:9889885-9891885 | 965 | 979 | + | 4.3357 | 2.26E-05 | 0.59 | GGAATGGTCACTCTG |
| 1 | GMTACGGTCACACTG | chr2L:6479575-6481575 | 971 | 985 | + | 4.3357 | 2.26E-05 | 0.59 | CGACCGGTCACACTG |
| 1 | GMTACGGTCACACTG | chr3R:10349182-10351182 | 972 | 986 | + | 4.3357 | 2.26E-05 | 0.59 | GGTACGGTCACTCCA |
| 1 | GMTACGGTCACACTG | chr2L:19525686-19527686 | 974 | 988 | + | 4.3357 | 2.26E-05 | 0.59 | TGTGCGGTCACACAG |
| 1 | GMTACGGTCACACTG | chr3L:11719934-11721934 | 976 | 990 | + | 4.3357 | 2.26E-05 | 0.59 | GTTGCGACCACACTG |
| 1 | GMTACGGTCACACTG | chr2L:6963421-6965421 | 976 | 990 | + | 4.3357 | 2.26E-05 | 0.59 | ATCACGGTCACACCG |
| 1 | GMTACGGTCACACTG | chrX:15703125-15705125 | 979 | 993 | + | 4.3357 | 2.26E-05 | 0.59 | GGCATGGTCACACCG |
| 1 | GMTACGGTCACACTG | chr2L:11988936-11990936 | 981 | 995 | + | 4.3357 | 2.26E-05 | 0.59 | CTACCGGTCACACTG |
| 1 | GMTACGGTCACACTG | chr3L:15162628-15164628 | 984 | 998 | + | 4.3357 | 2.26E-05 | 0.59 | GGTACGGCCAAACAG |
| 1 | GMTACGGTCACACTG | chr2R:15681719-15683719 | 986 | 1000 | + | 4.3357 | 2.26E-05 | 0.59 | CGGTCGGTCACACTG |
| 1 | GMTACGGTCACACTG | chr2R:16149123-16151123 | 991 | 1005 | + | 4.3357 | 2.26E-05 | 0.59 | ATCACGGTCACTCTG |
| 1 | GMTACGGTCACACTG | chrX:1469666-1471666 | 991 | 1005 | + | 4.3357 | 2.26E-05 | 0.59 | GTACTGGTCACACTG |
| 1 | GMTACGGTCACACTG | chrX:13781459-13783459 | 993 | 1007 | + | 4.3357 | 2.26E-05 | 0.59 | GGTGTGGCCACACTG |
| 1 | GMTACGGTCACACTG | chr2L:2219748-2221748 | 993 | 1007 | + | 4.3357 | 2.26E-05 | 0.59 | ATAACGGTCACACCG |
| 1 | GMTACGGTCACACTG | chr3L:19914051-19916051 | 994 | 1008 | + | 4.3357 | 2.26E-05 | 0.59 | CGCGCGGTCACACTG |
| 1 | GMTACGGTCACACTG | chr3L:21822461-21824461 | 996 | 1010 | + | 4.3357 | 2.26E-05 | 0.59 | CTAACGGCCACACTG |
| 1 | GMTACGGTCACACTG | chrX:20390447-20392447 | 998 | 1012 | + | 4.3357 | 2.26E-05 | 0.59 | AGCGCGGTCACACTG |
| 1 | GMTACGGTCACACTG | chrX:1482413-1484413 | 999 | 1013 | + | 4.3357 | 2.26E-05 | 0.59 | AGAGCGGTCACACTG |
| 1 | GMTACGGTCACACTG | chr3L:12749119-12751119 | 1003 | 1017 | + | 4.3357 | 2.26E-05 | 0.59 | GTAGTGGTCACACTG |
| 1 | GMTACGGTCACACTG | chr2R:24021343-24023343 | 1004 | 1018 | + | 4.3357 | 2.26E-05 | 0.59 | ATAACGGTCTCACTG |
| 1 | GMTACGGTCACACTG | chr3R:17605106-17607106 | 1005 | 1019 | + | 4.3357 | 2.26E-05 | 0.59 | GGCGCGGTCATACTG |
| 1 | GMTACGGTCACACTG | chr2L:2380097-2382097 | 1006 | 1020 | + | 4.3357 | 2.26E-05 | 0.59 | CGAACGGCCACACTG |
| 1 | GMTACGGTCACACTG | chrX:7934896-7936896 | 1007 | 1021 | + | 4.3357 | 2.26E-05 | 0.59 | AGAGCGGTCACACTG |
| 1 | GMTACGGTCACACTG | chr3R:23766712-23768712 | 1009 | 1023 | + | 4.3357 | 2.26E-05 | 0.59 | CGCACGGTCACTCTG |
| 1 | GMTACGGTCACACTG | chr3R:22033298-22035298 | 1010 | 1024 | + | 4.3357 | 2.26E-05 | 0.59 | GTAACGGTCTCACTA |
| 1 | GMTACGGTCACACTG | chr3R:21748853-21750853 | 1013 | 1027 | + | 4.3357 | 2.26E-05 | 0.59 | GTAACGGTCATTCTG |
| 1 | GMTACGGTCACACTG | chr3R:20620938-20622938 | 1013 | 1027 | + | 4.3357 | 2.26E-05 | 0.59 | TTGACGGTCACACAG |
| 1 | GMTACGGTCACACTG | chr2L:7055301-7057301 | 1016 | 1030 | + | 4.3357 | 2.26E-05 | 0.59 | GGTGTGGACACACTG |
| 1 | GMTACGGTCACACTG | chr3R:29932287-29934287 | 1021 | 1035 | + | 4.3357 | 2.26E-05 | 0.59 | CTAACGGGCACACTG |
| 1 | GMTACGGTCACACTG | chr3R:29932287-29934287 | 1021 | 1035 | + | 4.3357 | 2.26E-05 | 0.59 | CTAACGGGCACACTG |

| | | | | | | | | | |
|---|---|---|---|---|---|---|---|---|---|
| 1 | GMTACGGTCACACTG | chr3R:22033286-22035286 | 1022 | 1036 | + | 4.3357 | 2.26E-05 | 0.59 | GTAACGGTCTCACTA |
| 1 | GMTACGGTCACACTG | chr3R:22033286-22035286 | 1022 | 1036 | + | 4.3357 | 2.26E-05 | 0.59 | GTAACGGTCTCACTA |
| 1 | GMTACGGTCACACTG | chr2L:14511083-14513083 | 1023 | 1037 | + | 4.3357 | 2.26E-05 | 0.59 | TGAACGGTCGCACTG |
| 1 | GMTACGGTCACACTG | chr3L:4138076-4140076 | 1026 | 1040 | + | 4.3357 | 2.26E-05 | 0.59 | AGAACGGTCACGCTG |
| 1 | GMTACGGTCACACTG | chr3R:13046731-13048731 | 1032 | 1046 | + | 4.3357 | 2.26E-05 | 0.59 | GTTATGGTCGCACCG |
| 1 | GMTACGGTCACACTG | chr2L:11988882-11990882 | 1035 | 1049 | + | 4.3357 | 2.26E-05 | 0.59 | CTACCGGTCACACTG |
| 1 | GMTACGGTCACACTG | chr3L:22265679-22267679 | 1048 | 1062 | + | 4.3357 | 2.26E-05 | 0.59 | GTTCCGGCCAAACTG |
| 1 | GMTACGGTCACACTG | chr2R:24021298-24023298 | 1049 | 1063 | + | 4.3357 | 2.26E-05 | 0.59 | ATAACGGTCTCACTG |
| 1 | GMTACGGTCACACTG | chrX:16072568-16074568 | 1053 | 1067 | + | 4.3357 | 2.26E-05 | 0.59 | TTTGCGGTCACTCTG |
| 1 | GMTACGGTCACACTG | chr2L:8081982-8083982 | 1053 | 1067 | + | 4.3357 | 2.26E-05 | 0.59 | TTAACGGTCGCACTG |
| 1 | GMTACGGTCACACTG | chr3R:8008297-8010297 | 1066 | 1080 | + | 4.3357 | 2.26E-05 | 0.59 | GTAAGGGTCACCCTG |
| 1 | GMTACGGTCACACTG | chr3L:12749056-12751056 | 1066 | 1080 | + | 4.3357 | 2.26E-05 | 0.59 | GTAGTGGTCACACTG |
| 1 | GMTACGGTCACACTG | chr2L:9889772-9891772 | 1078 | 1092 | + | 4.3357 | 2.26E-05 | 0.59 | GGAATGGTCACTCTG |
| 1 | GMTACGGTCACACTG | chr3L:3318232-3320232 | 1081 | 1095 | + | 4.3357 | 2.26E-05 | 0.59 | GTGCTGGTCACACTG |
| 1 | GMTACGGTCACACTG | chr3R:12915056-12917056 | 1087 | 1101 | + | 4.3357 | 2.26E-05 | 0.59 | GGTACGGTCCATCTG |
| 1 | GMTACGGTCACACTG | chr3L:4137996-4139996 | 1106 | 1120 | + | 4.3357 | 2.26E-05 | 0.59 | AGAACGGTCACGCTG |
| 1 | GMTACGGTCACACTG | chr3L:2553292-2555292 | 1106 | 1120 | + | 4.3357 | 2.26E-05 | 0.59 | AGTGCGGTCTCACTG |
| 1 | GMTACGGTCACACTG | chrX:5621648-5623648 | 1115 | 1129 | + | 4.3357 | 2.26E-05 | 0.59 | GGCCGGGTCACACTG |
| 1 | GMTACGGTCACACTG | chrX:19637767-19639767 | 1115 | 1129 | + | 4.3357 | 2.26E-05 | 0.59 | GTTCTGGCCACACTG |
| 1 | GMTACGGTCACACTG | chr3L:742794-744794 | 1123 | 1137 | + | 4.3357 | 2.26E-05 | 0.59 | GGTGTGGCCACACTG |
| 1 | GMTACGGTCACACTG | chr2R:16343377-16345377 | 1131 | 1145 | + | 4.3357 | 2.26E-05 | 0.59 | GGAGGGGTCACACTG |
| 1 | GMTACGGTCACACTG | chr3R:11234140-11236140 | 1163 | 1177 | + | 4.3357 | 2.26E-05 | 0.59 | GTTGCGGTCCCACTA |
| 1 | GMTACGGTCACACTG | chr2L:8081859-8083859 | 1176 | 1190 | + | 4.3357 | 2.26E-05 | 0.59 | TTAACGGTCGCACTG |
| 1 | GMTACGGTCACACTG | chr3L:3198090-3200090 | 1188 | 1202 | + | 4.3357 | 2.26E-05 | 0.59 | CGGACGGTCGCACTG |
| 1 | GMTACGGTCACACTG | chr3L:6639891-6641891 | 1197 | 1211 | + | 4.3357 | 2.26E-05 | 0.59 | GTCGCGGTCACAGTG |
| 1 | GMTACGGTCACACTG | chrX:14242686-14244686 | 1203 | 1217 | + | 4.3357 | 2.26E-05 | 0.59 | GGAACGGGCACAGTG |
| 1 | GMTACGGTCACACTG | chr2R:24631932-24633932 | 1205 | 1219 | + | 4.3357 | 2.26E-05 | 0.59 | GTTGCGGTTACACCG |
| 1 | GMTACGGTCACACTG | chr2R:6985043-6987043 | 1222 | 1236 | + | 4.3357 | 2.26E-05 | 0.59 | GGTTCGGTCAAACAG |
| 1 | GMTACGGTCACACTG | chrX:5621539-5623539 | 1224 | 1238 | + | 4.3357 | 2.26E-05 | 0.59 | GGCCGGGTCACACTG |
| 1 | GMTACGGTCACACTG | chr3L:3198047-3200047 | 1231 | 1245 | + | 4.3357 | 2.26E-05 | 0.59 | CGGACGGTCGCACTG |
| 1 | GMTACGGTCACACTG | chr3R:23766488-23768488 | 1233 | 1247 | + | 4.3357 | 2.26E-05 | 0.59 | CGCACGGTCACTCTG |
| 1 | GMTACGGTCACACTG | chr2L:17175618-17177618 | 1240 | 1254 | + | 4.3357 | 2.26E-05 | 0.59 | GTGATGGTCTCACTG |
| 1 | GMTACGGTCACACTG | chrX:11136859-11138859 | 1243 | 1257 | + | 4.3357 | 2.26E-05 | 0.59 | GTCACGGTGACACCG |

| | | | | | | | | | |
|---|---|---|---|---|---|---|---|---|---|
| 1 | GMTACGGTCACACTG | chr3R:29932057-29934057 | 1251 | 1265 | + | 4.3357 | 2.26E-05 | 0.59 | CTAACGGGCACACTG |
| 1 | GMTACGGTCACACTG | chr3R:27244333-27246333 | 1259 | 1273 | + | 4.3357 | 2.26E-05 | 0.59 | GGCACAGTCACACCG |
| 1 | GMTACGGTCACACTG | chrX:13781170-13783170 | 1282 | 1296 | + | 4.3357 | 2.26E-05 | 0.59 | GGTGTGGCCACACTG |
| 1 | GMTACGGTCACACTG | chr3R:11234013-11236013 | 1290 | 1304 | + | 4.3357 | 2.26E-05 | 0.59 | GTTGCGGTCCCACTA |
| 1 | GMTACGGTCACACTG | chr2L:19525368-19527368 | 1292 | 1306 | + | 4.3357 | 2.26E-05 | 0.59 | TGTGCGGTCACACAG |
| 1 | GMTACGGTCACACTG | chr3R:29932287-29934287 | 1292 | 1306 | + | 4.3357 | 2.26E-05 | 0.59 | GGTACGGGGACACAG |
| 1 | GMTACGGTCACACTG | chr3R:29932287-29934287 | 1292 | 1306 | + | 4.3357 | 2.26E-05 | 0.59 | GGTACGGGGACACAG |
| 1 | GMTACGGTCACACTG | chr3R:23923399-23925399 | 1295 | 1309 | + | 4.3357 | 2.26E-05 | 0.59 | GTGACGGGAACACTG |
| 1 | GMTACGGTCACACTG | chr3R:18982650-18984650 | 1309 | 1323 | + | 4.3357 | 2.26E-05 | 0.59 | CGAACGGTCACACGG |
| 1 | GMTACGGTCACACTG | chr2L:12433665-12435665 | 1312 | 1326 | + | 4.3357 | 2.26E-05 | 0.59 | ATTACGGACGCACTG |
| 1 | GMTACGGTCACACTG | chr2R:16148795-16150795 | 1319 | 1333 | + | 4.3357 | 2.26E-05 | 0.59 | ATCACGGTCACTCTG |
| 1 | GMTACGGTCACACTG | chr2L:6963074-6965074 | 1323 | 1337 | + | 4.3357 | 2.26E-05 | 0.59 | ATCACGGTCACACCG |
| 1 | GMTACGGTCACACTG | chr2R:16343182-16345182 | 1326 | 1340 | + | 4.3357 | 2.26E-05 | 0.59 | GGAGGGGTCACACTG |
| 1 | GMTACGGTCACACTG | chr3L:2598579-2600579 | 1332 | 1346 | + | 4.3357 | 2.26E-05 | 0.59 | GTTACGGCCAACCTG |
| 1 | GMTACGGTCACACTG | chrX:15702770-15704770 | 1334 | 1348 | + | 4.3357 | 2.26E-05 | 0.59 | GGCATGGTCACACCG |
| 1 | GMTACGGTCACACTG | chr3R:27244249-27246249 | 1343 | 1357 | + | 4.3357 | 2.26E-05 | 0.59 | GGCACAGTCACACCG |
| 1 | GMTACGGTCACACTG | chrX:5621416-5623416 | 1347 | 1361 | + | 4.3357 | 2.26E-05 | 0.59 | GGCCGGGTCACACTG |
| 1 | GMTACGGTCACACTG | chr3R:15289799-15291799 | 1348 | 1362 | + | 4.3357 | 2.26E-05 | 0.59 | GGTTCGTTCACTCTG |
| 1 | GMTACGGTCACACTG | chr3L:3188978-3190978 | 1350 | 1364 | + | 4.3357 | 2.26E-05 | 0.59 | GTTGCGGCGACACTG |
| 1 | GMTACGGTCACACTG | chr2R:6838055-6840055 | 1354 | 1368 | + | 4.3357 | 2.26E-05 | 0.59 | TTTGCGGCCACACTG |
| 1 | GMTACGGTCACACTG | chr3R:8560764-8562764 | 1379 | 1393 | + | 4.3357 | 2.26E-05 | 0.59 | GGAACGGACATACTG |
| 1 | GMTACGGTCACACTG | chr2L:2238384-2240384 | 1406 | 1420 | + | 4.3357 | 2.26E-05 | 0.59 | GGTACGGCCACCATG |
| 1 | GMTACGGTCACACTG | chrX:17076897-17078897 | 1418 | 1432 | + | 4.3357 | 2.26E-05 | 0.59 | ATTTCGGGCACACTG |
| 1 | GMTACGGTCACACTG | chr2R:11355720-11357720 | 1468 | 1482 | + | 4.3357 | 2.26E-05 | 0.59 | GTTGGGGCCACACTG |
| 1 | GMTACGGTCACACTG | chr2L:4351975-4353975 | 1470 | 1484 | + | 4.3357 | 2.26E-05 | 0.59 | GGCACGGTCACAAAG |
| 1 | GMTACGGTCACACTG | chr3R:6404895-6406895 | 1477 | 1491 | + | 4.3357 | 2.26E-05 | 0.59 | GGAATGGGCACACTG |
| 1 | GMTACGGTCACACTG | chr3R:26020076-26022076 | 1493 | 1507 | + | 4.3357 | 2.26E-05 | 0.59 | GTGATGGTCCCACTG |
| 1 | GMTACGGTCACACTG | chrX:19637369-19639369 | 1513 | 1527 | + | 4.3357 | 2.26E-05 | 0.59 | GTTCTGGCCACACTG |
| 1 | GMTACGGTCACACTG | chr3R:29932057-29934057 | 1522 | 1536 | + | 4.3357 | 2.26E-05 | 0.59 | GGTACGGGGACACAG |
| 1 | GMTACGGTCACACTG | chr2L:12433452-12435452 | 1525 | 1539 | + | 4.3357 | 2.26E-05 | 0.59 | ATTACGGACGCACTG |
| 1 | GMTACGGTCACACTG | chr3R:4505365-4507365 | 1528 | 1542 | + | 4.3357 | 2.26E-05 | 0.59 | GTTCCGGTCAGACAG |
| 1 | GMTACGGTCACACTG | chr2R:6984712-6986712 | 1553 | 1567 | + | 4.3357 | 2.26E-05 | 0.59 | GGTTCGGTCAAACAG |
| 1 | GMTACGGTCACACTG | chr2R:6984712-6986712 | 1553 | 1567 | + | 4.3357 | 2.26E-05 | 0.59 | GGTTCGGTCAAACAG |

| | | | | | | | | | |
|---|---|---|---|---|---|---|---|---|---|
| 1 | GMTACGGTCACACTG | chr4:425067-427067 | 1574 | 1588 | + | 4.3357 | 2.26E-05 | 0.59 | GGTAGGGACACTCTG |
| 1 | GMTACGGTCACACTG | chr3L:3188747-3190747 | 1581 | 1595 | + | 4.3357 | 2.26E-05 | 0.59 | GTTGCGGCGACACTG |
| 1 | GMTACGGTCACACTG | chr2R:11355600-11357600 | 1588 | 1602 | + | 4.3357 | 2.26E-05 | 0.59 | GTTGGGGCCACACTG |
| 1 | GMTACGGTCACACTG | chr3R:24293399-24295399 | 1605 | 1619 | + | 4.3357 | 2.26E-05 | 0.59 | GGACTGGTCACACTG |
| 1 | GMTACGGTCACACTG | chr3L:2598297-2600297 | 1614 | 1628 | + | 4.3357 | 2.26E-05 | 0.59 | GTTACGGCCAACCTG |
| 1 | GMTACGGTCACACTG | chr2R:20549117-20551117 | 1627 | 1641 | + | 4.3357 | 2.26E-05 | 0.59 | GTTTCTGGCACACTG |
| 1 | GMTACGGTCACACTG | chr2R:20549117-20551117 | 1627 | 1641 | + | 4.3357 | 2.26E-05 | 0.59 | GTTTCTGGCACACTG |
| 1 | GMTACGGTCACACTG | chrX:20389806-20391806 | 1639 | 1653 | + | 4.3357 | 2.26E-05 | 0.59 | AGCGCGGTCACACTG |
| 1 | GMTACGGTCACACTG | chr3R:20633732-20635732 | 1662 | 1676 | + | 4.3357 | 2.26E-05 | 0.59 | ATTTCGGTCACACAG |
| 1 | GMTACGGTCACACTG | chr3L:9385112-9387112 | 1667 | 1681 | + | 4.3357 | 2.26E-05 | 0.59 | GTTCCGCGCACACTG |
| 1 | GMTACGGTCACACTG | chr3L:9385112-9387112 | 1667 | 1681 | + | 4.3357 | 2.26E-05 | 0.59 | GTTCCGCGCACACTG |
| 1 | GMTACGGTCACACTG | chr2L:7753174-7755174 | 1684 | 1698 | + | 4.3357 | 2.26E-05 | 0.59 | GGTTCGGTGACTCTG |
| 1 | GMTACGGTCACACTG | chr3R:10386385-10388385 | 1706 | 1720 | + | 4.3357 | 2.26E-05 | 0.59 | GTTTCTGTCCCACTG |
| 1 | GMTACGGTCACACTG | chr2L:18184857-18186857 | 1736 | 1750 | + | 4.3357 | 2.26E-05 | 0.59 | GGTCGGGTCACACGG |
| 1 | GMTACGGTCACACTG | chrX:19696091-19698091 | 1742 | 1756 | + | 4.3357 | 2.26E-05 | 0.59 | CGAACGGTCACTCTG |
| 1 | GMTACGGTCACACTG | chr3R:31977121-31979121 | 1750 | 1764 | + | 4.3357 | 2.26E-05 | 0.59 | GGCACGGGCAAACTG |
| 1 | GMTACGGTCACACTG | chr3R:9049555-9051555 | 1753 | 1767 | + | 4.3357 | 2.26E-05 | 0.59 | GTTACGGGCGGACTG |
| 1 | GMTACGGTCACACTG | chr3R:4253708-4255708 | 1771 | 1785 | + | 4.3357 | 2.26E-05 | 0.59 | GTGACGGTCACATCG |
| 1 | GMTACGGTCACACTG | chrX:1481629-1483629 | 1783 | 1797 | + | 4.3357 | 2.26E-05 | 0.59 | AGAGCGGTCACACTG |
| 1 | GMTACGGTCACACTG | chr2L:6081603-6083603 | 1815 | 1829 | + | 4.3357 | 2.26E-05 | 0.59 | CTAACGGTCTCACTG |
| 1 | GMTACGGTCACACTG | chrX:1481590-1483590 | 1822 | 1836 | + | 4.3357 | 2.26E-05 | 0.59 | AGAGCGGTCACACTG |
| 1 | GMTACGGTCACACTG | chr2R:16487053-16489053 | 1834 | 1848 | + | 4.3357 | 2.26E-05 | 0.59 | GGTACGGACATTCTG |
| 1 | GMTACGGTCACACTG | chrX:1468817-1470817 | 1840 | 1854 | + | 4.3357 | 2.26E-05 | 0.59 | GTACTGGTCACACTG |
| 1 | GMTACGGTCACACTG | chr3R:11720734-11722734 | 1841 | 1855 | + | 4.3357 | 2.26E-05 | 0.59 | GTTAAGGTCCCTCTG |
| 1 | GMTACGGTCACACTG | chr3L:2271769-2273769 | 1859 | 1873 | + | 4.3357 | 2.26E-05 | 0.59 | GGAACAGTCACTCTG |
| 1 | GMTACGGTCACACTG | chr3L:15974368-15976368 | 1924 | 1938 | + | 4.3357 | 2.26E-05 | 0.59 | GGAACGGAGACACTG |
| 1 | GMTACGGTCACACTG | chr3R:10875384-10877384 | 1968 | 1982 | + | 4.3357 | 2.26E-05 | 0.59 | GGTTTGGTCACCCTG |
| 1 | GMTACGGTCACACTG | chr3R:4432965-4434965 | 1972 | 1986 | + | 4.3357 | 2.26E-05 | 0.59 | GGAACGGTCAATCTG |
| 1 | GMTACGGTCACACTG | chr2L:6916310-6918310 | 1972 | 1986 | + | 4.3357 | 2.26E-05 | 0.59 | GTTGCAGACACACTG |
| 1 | GMTACGGTCACACTG | chrX:11136118-11138118 | 1984 | 1998 | + | 4.3357 | 2.26E-05 | 0.59 | GTCACGGTGACACCG |
| 1 | GMTACGGTCACACTG | chrY:3211119-3213119 | 6 | 20 | - | 4.0571 | 3.01E-05 | 0.673 | GTAACTGTAACACTG |
| 1 | GMTACGGTCACACTG | chr3R:11821493-11823493 | 14 | 28 | - | 4.0571 | 3.01E-05 | 0.673 | GTTACGGTCACGTTC |
| 1 | GMTACGGTCACACTG | chr3R:11619090-11621090 | 30 | 44 | - | 4.0571 | 3.01E-05 | 0.673 | GTTAAGTGCACACTG |

| | | | | | | | | | |
|---|---|---|---|---|---|---|---|---|---|
| 1 | GMTACGGTCACACTG | chrX:15858487-15860487 | 37 | 51 | - | 4.0571 | 3.01E-05 | 0.673 | GTTTTGGTCAGACTG |
| 1 | GMTACGGTCACACTG | chr3R:29748163-29750163 | 44 | 58 | - | 4.0571 | 3.01E-05 | 0.673 | GTTACGATTACACCG |
| 1 | GMTACGGTCACACTG | chr2L:7332714-7334714 | 54 | 68 | - | 4.0571 | 3.01E-05 | 0.673 | GTTTCTGTGACACTG |
| 1 | GMTACGGTCACACTG | chr2R:17535213-17537213 | 57 | 71 | - | 4.0571 | 3.01E-05 | 0.673 | GTTATTGTCGCACTG |
| 1 | GMTACGGTCACACTG | chr2R:13429355-13431355 | 70 | 84 | - | 4.0571 | 3.01E-05 | 0.673 | TGTACGGGCACAGTG |
| 1 | GMTACGGTCACACTG | chr2L:2289738-2291738 | 75 | 89 | - | 4.0571 | 3.01E-05 | 0.673 | GGTAAAGTCACACGG |
| 1 | GMTACGGTCACACTG | chr3L:14408939-14410939 | 76 | 90 | - | 4.0571 | 3.01E-05 | 0.673 | GTTTTGGTAACACTG |
| 1 | GMTACGGTCACACTG | chr3L:9713324-9715324 | 79 | 93 | - | 4.0571 | 3.01E-05 | 0.673 | GTTACGATCAGACGG |
| 1 | GMTACGGTCACACTG | chr3R:13318270-13320270 | 115 | 129 | - | 4.0571 | 3.01E-05 | 0.673 | GTAACTGTAACACTG |
| 1 | GMTACGGTCACACTG | chr2R:19451504-19453504 | 132 | 146 | - | 4.0571 | 3.01E-05 | 0.673 | AGTATGGCCACACTG |
| 1 | GMTACGGTCACACTG | chr2R:22102177-22104177 | 161 | 175 | - | 4.0571 | 3.01E-05 | 0.673 | GTTAAGGTCACGGTG |
| 1 | GMTACGGTCACACTG | chr2L:1751122-1753122 | 168 | 182 | - | 4.0571 | 3.01E-05 | 0.673 | TGCACAGTCACACTG |
| 1 | GMTACGGTCACACTG | chrX:15687857-15689857 | 176 | 190 | - | 4.0571 | 3.01E-05 | 0.673 | GGTACCCCCACACTG |
| 1 | GMTACGGTCACACTG | chrY:2266259-2268259 | 180 | 194 | - | 4.0571 | 3.01E-05 | 0.673 | ATTACGGTTACGCTG |
| 1 | GMTACGGTCACACTG | chr2R:15988498-15990498 | 188 | 202 | - | 4.0571 | 3.01E-05 | 0.673 | TGTACGGCCACAGTG |
| 1 | GMTACGGTCACACTG | chr2R:7177390-7179390 | 191 | 205 | - | 4.0571 | 3.01E-05 | 0.673 | GGTAGGGACACAATG |
| 1 | GMTACGGTCACACTG | chrX:9873057-9875057 | 205 | 219 | - | 4.0571 | 3.01E-05 | 0.673 | GGTACGATCCCAATG |
| 1 | GMTACGGTCACACTG | chr3R:11618906-11620906 | 214 | 228 | - | 4.0571 | 3.01E-05 | 0.673 | GTTAAGTGCACACTG |
| 1 | GMTACGGTCACACTG | chr2R:18051107-18053107 | 234 | 248 | - | 4.0571 | 3.01E-05 | 0.673 | GTTATGTTCACCCTG |
| 1 | GMTACGGTCACACTG | chrX:21155621-21157621 | 236 | 250 | - | 4.0571 | 3.01E-05 | 0.673 | GTCACTGTCACAGTG |
| 1 | GMTACGGTCACACTG | chr2R:23385521-23387521 | 263 | 277 | - | 4.0571 | 3.01E-05 | 0.673 | AGTTCGGTCACACTA |
| 1 | GMTACGGTCACACTG | chrX:10054109-10056109 | 268 | 282 | - | 4.0571 | 3.01E-05 | 0.673 | AGTCCGGTCACAATG |
| 1 | GMTACGGTCACACTG | chr3R:26190205-26192205 | 271 | 285 | - | 4.0571 | 3.01E-05 | 0.673 | CTTCGGGTCACACTG |
| 1 | GMTACGGTCACACTG | chr2R:10810078-10812078 | 291 | 305 | - | 4.0571 | 3.01E-05 | 0.673 | GGCACGGTCATAATG |
| 1 | GMTACGGTCACACTG | chr3R:31122415-31124415 | 294 | 308 | - | 4.0571 | 3.01E-05 | 0.673 | GTTACGGTAACACCA |
| 1 | GMTACGGTCACACTG | chr2R:23385463-23387463 | 321 | 335 | - | 4.0571 | 3.01E-05 | 0.673 | AGTTCGGTCACACTA |
| 1 | GMTACGGTCACACTG | chrX:6066958-6068958 | 323 | 337 | - | 4.0571 | 3.01E-05 | 0.673 | ATTACGACCACACTG |
| 1 | GMTACGGTCACACTG | chr3R:25328316-25330316 | 336 | 350 | - | 4.0571 | 3.01E-05 | 0.673 | GTAACTGTAACACTG |
| 1 | GMTACGGTCACACTG | chr2R:16823664-16825664 | 345 | 359 | - | 4.0571 | 3.01E-05 | 0.673 | GTTTCGATAACACTG |
| 1 | GMTACGGTCACACTG | chr2R:8723905-8725905 | 345 | 359 | - | 4.0571 | 3.01E-05 | 0.673 | GTTATGGTCACCCTA |
| 1 | GMTACGGTCACACTG | chr3R:10872038-10874038 | 354 | 368 | - | 4.0571 | 3.01E-05 | 0.673 | TGTCTGGTCACACTG |
| 1 | GMTACGGTCACACTG | chr2R:17534866-17536866 | 404 | 418 | - | 4.0571 | 3.01E-05 | 0.673 | GTTATTGTCGCACTG |
| 1 | GMTACGGTCACACTG | chr2R:15694386-15696386 | 406 | 420 | - | 4.0571 | 3.01E-05 | 0.673 | GTTAAGGACACATTG |

| | | | | | | | | | |
|---|---|---|---|---|---|---|---|---|---|
| 1 | GMTACGGTCACACTG | chr2L:13366617-13368617 | 433 | 447 | - | 4.0571 | 3.01E-05 | 0.673 | GTTTCGTTCACAATG |
| 1 | GMTACGGTCACACTG | chr2R:10809936-10811936 | 433 | 447 | - | 4.0571 | 3.01E-05 | 0.673 | GGCACGGTCATAATG |
| 1 | GMTACGGTCACACTG | chr2L:6901062-6903062 | 457 | 471 | - | 4.0571 | 3.01E-05 | 0.673 | GTTTCGATTACACTG |
| 1 | GMTACGGTCACACTG | chr2L:22537230-22539230 | 464 | 478 | - | 4.0571 | 3.01E-05 | 0.673 | GTTTGGGTCACATTG |
| 1 | GMTACGGTCACACTG | chr3R:28844270-28846270 | 481 | 495 | - | 4.0571 | 3.01E-05 | 0.673 | ATAACAGTCACACTG |
| 1 | GMTACGGTCACACTG | chr2L:10326943-10328943 | 497 | 511 | - | 4.0571 | 3.01E-05 | 0.673 | GTTACTATCACCCTG |
| 1 | GMTACGGTCACACTG | chr2R:16872772-16874772 | 507 | 521 | - | 4.0571 | 3.01E-05 | 0.673 | GGTATGGTCAATCTG |
| 1 | GMTACGGTCACACTG | chr2L:11342318-11344318 | 526 | 540 | - | 4.0571 | 3.01E-05 | 0.673 | GTTACGGTCATTCTC |
| 1 | GMTACGGTCACACTG | chr2R:24774711-24776711 | 530 | 544 | - | 4.0571 | 3.01E-05 | 0.673 | CGAACGGTCACACTA |
| 1 | GMTACGGTCACACTG | chr2R:20973017-20975017 | 537 | 551 | - | 4.0571 | 3.01E-05 | 0.673 | ATTACGGAGACACTG |
| 1 | GMTACGGTCACACTG | chr2R:5753517-5755517 | 554 | 568 | - | 4.0571 | 3.01E-05 | 0.673 | TTTTGGGTCACACTG |
| 1 | GMTACGGTCACACTG | chr3R:22461455-22463455 | 572 | 586 | - | 4.0571 | 3.01E-05 | 0.673 | TTTGGGGTCACACTG |
| 1 | GMTACGGTCACACTG | chr3L:9712827-9714827 | 576 | 590 | - | 4.0571 | 3.01E-05 | 0.673 | GTTACGATCAGACGG |
| 1 | GMTACGGTCACACTG | chr3R:17810541-17812541 | 582 | 596 | - | 4.0571 | 3.01E-05 | 0.673 | AGTACGGTCCCACTA |
| 1 | GMTACGGTCACACTG | chr3L:20472783-20474783 | 582 | 596 | - | 4.0571 | 3.01E-05 | 0.673 | AGTGTGGTCACACTG |
| 1 | GMTACGGTCACACTG | chr3L:11073769-11075769 | 629 | 643 | - | 4.0571 | 3.01E-05 | 0.673 | TTTACGGTCTTACTG |
| 1 | GMTACGGTCACACTG | chr2R:15679545-15681545 | 635 | 649 | - | 4.0571 | 3.01E-05 | 0.673 | CTTACGGTCACACCA |
| 1 | GMTACGGTCACACTG | chr3L:20472723-20474723 | 642 | 656 | - | 4.0571 | 3.01E-05 | 0.673 | AGTGTGGTCACACTG |
| 1 | GMTACGGTCACACTG | chr3L:13928387-13930387 | 644 | 658 | - | 4.0571 | 3.01E-05 | 0.673 | GTTCCGGTCACAGTT |
| 1 | GMTACGGTCACACTG | chr3R:16341318-16343318 | 661 | 675 | - | 4.0571 | 3.01E-05 | 0.673 | CTAACGGTCATACTG |
| 1 | GMTACGGTCACACTG | chrX:21146041-21148041 | 665 | 679 | - | 4.0571 | 3.01E-05 | 0.673 | GTCACTGTCACAGTG |
| 1 | GMTACGGTCACACTG | chrX:147426-149426 | 671 | 685 | - | 4.0571 | 3.01E-05 | 0.673 | GTCACGCTCATACTG |
| 1 | GMTACGGTCACACTG | chr2R:10644735-10646735 | 682 | 696 | - | 4.0571 | 3.01E-05 | 0.673 | TGTTTGGTCACACTG |
| 1 | GMTACGGTCACACTG | chrX:21155173-21157173 | 684 | 698 | - | 4.0571 | 3.01E-05 | 0.673 | GTCACTGTCACAGTG |
| 1 | GMTACGGTCACACTG | chr3R:29798658-29800658 | 685 | 699 | - | 4.0571 | 3.01E-05 | 0.673 | TGCACGGTCACACTC |
| 1 | GMTACGGTCACACTG | chr3R:19904164-19906164 | 697 | 711 | - | 4.0571 | 3.01E-05 | 0.673 | TGTGGGGTCACACTG |
| 1 | GMTACGGTCACACTG | chr3L:19598088-19600088 | 699 | 713 | - | 4.0571 | 3.01E-05 | 0.673 | CGCAAGGTCACACTG |
| 1 | GMTACGGTCACACTG | chrX:5621648-5623648 | 718 | 732 | - | 4.0571 | 3.01E-05 | 0.673 | TTTACGATCGCACTG |
| 1 | GMTACGGTCACACTG | chrX:3484650-3486650 | 728 | 742 | - | 4.0571 | 3.01E-05 | 0.673 | CGCACAGTCACACTG |
| 1 | GMTACGGTCACACTG | chr2R:9890955-9892955 | 730 | 744 | - | 4.0571 | 3.01E-05 | 0.673 | CGCACGGTCACACTC |
| 1 | GMTACGGTCACACTG | chr2R:13857952-13859952 | 730 | 744 | - | 4.0571 | 3.01E-05 | 0.673 | TTAATGGTCACACTG |
| 1 | GMTACGGTCACACTG | chr3L:6192480-6194480 | 732 | 746 | - | 4.0571 | 3.01E-05 | 0.673 | TGCACGGTCACACTC |
| 1 | GMTACGGTCACACTG | chr3L:22497310-22499310 | 737 | 751 | - | 4.0571 | 3.01E-05 | 0.673 | ATTACTGTCACACGG |

| | | | | | | | | | |
|---|---|---|---|---|---|---|---|---|---|
| 1 | GMTACGGTCACACTG | chr2L:4970702-4972702 | 744 | 758 | - | 4.0571 | 3.01E-05 | 0.673 | ATTACAGTCACACCG |
| 1 | GMTACGGTCACACTG | chr2L:6490061-6492061 | 762 | 776 | - | 4.0571 | 3.01E-05 | 0.673 | GTAAAGGTCACACTA |
| 1 | GMTACGGTCACACTG | chr3L:12500513-12502513 | 791 | 805 | - | 4.0571 | 3.01E-05 | 0.673 | GTTTTGGTCACACTA |
| 1 | GMTACGGTCACACTG | chr3R:26880734-26882734 | 793 | 807 | - | 4.0571 | 3.01E-05 | 0.673 | TGTACGCACACACTG |
| 1 | GMTACGGTCACACTG | chr3L:8299064-8301064 | 801 | 815 | - | 4.0571 | 3.01E-05 | 0.673 | GTTACGGTCTCATTA |
| 1 | GMTACGGTCACACTG | chr3L:11168817-11170817 | 801 | 815 | - | 4.0571 | 3.01E-05 | 0.673 | CGCAAGGTCACACTG |
| 1 | GMTACGGTCACACTG | chrX:5621539-5623539 | 827 | 841 | - | 4.0571 | 3.01E-05 | 0.673 | TTTACGATCGCACTG |
| 1 | GMTACGGTCACACTG | chrX:16639968-16641968 | 830 | 844 | - | 4.0571 | 3.01E-05 | 0.673 | GGTATGTTCGCACTG |
| 1 | GMTACGGTCACACTG | chrX:7285358-7287358 | 830 | 844 | - | 4.0571 | 3.01E-05 | 0.673 | TGCACAGTCACACTG |
| 1 | GMTACGGTCACACTG | chr3L:23725829-23727829 | 844 | 858 | - | 4.0571 | 3.01E-05 | 0.673 | GTGACGGTCAGACTT |
| 1 | GMTACGGTCACACTG | chr2L:1499830-1501830 | 852 | 866 | - | 4.0571 | 3.01E-05 | 0.673 | TGAACGGTCACATTG |
| 1 | GMTACGGTCACACTG | chr2L:5042270-5044270 | 882 | 896 | - | 4.0571 | 3.01E-05 | 0.673 | CTGACGGTCACATTG |
| 1 | GMTACGGTCACACTG | chr2L:5042270-5044270 | 882 | 896 | - | 4.0571 | 3.01E-05 | 0.673 | CTGACGGTCACATTG |
| 1 | GMTACGGTCACACTG | chr2L:5042270-5044270 | 882 | 896 | - | 4.0571 | 3.01E-05 | 0.673 | CTGACGGTCACATTG |
| 1 | GMTACGGTCACACTG | chr2L:16728777-16730777 | 885 | 899 | - | 4.0571 | 3.01E-05 | 0.673 | GTCATGGTCACACTA |
| 1 | GMTACGGTCACACTG | chrX:3670801-3672801 | 889 | 903 | - | 4.0571 | 3.01E-05 | 0.673 | GTAATGGTCACACTT |
| 1 | GMTACGGTCACACTG | chr2L:8126001-8128001 | 907 | 921 | - | 4.0571 | 3.01E-05 | 0.673 | GTTATGGTCACCCTA |
| 1 | GMTACGGTCACACTG | chr2R:22798099-22800099 | 917 | 931 | - | 4.0571 | 3.01E-05 | 0.673 | AGTACGGTCACCCTC |
| 1 | GMTACGGTCACACTG | chr2R:14255493-14257493 | 929 | 943 | - | 4.0571 | 3.01E-05 | 0.673 | TGTACGGTCTCACTA |
| 1 | GMTACGGTCACACTG | chr3L:16231084-16233084 | 931 | 945 | - | 4.0571 | 3.01E-05 | 0.673 | AGCAAGGTCACACTG |
| 1 | GMTACGGTCACACTG | chr3L:11511221-11513221 | 939 | 953 | - | 4.0571 | 3.01E-05 | 0.673 | GGTACGCTTTCACTG |
| 1 | GMTACGGTCACACTG | chr2R:17000994-17002994 | 940 | 954 | - | 4.0571 | 3.01E-05 | 0.673 | ATGACGGTCACACTA |
| 1 | GMTACGGTCACACTG | chr3R:5567921-5569921 | 943 | 957 | - | 4.0571 | 3.01E-05 | 0.673 | GGAACCGTCACACTA |
| 1 | GMTACGGTCACACTG | chr2R:13845393-13847393 | 950 | 964 | - | 4.0571 | 3.01E-05 | 0.673 | ATGATGGTCACACTG |
| 1 | GMTACGGTCACACTG | chrX:5621416-5623416 | 950 | 964 | - | 4.0571 | 3.01E-05 | 0.673 | TTTACGATCGCACTG |
| 1 | GMTACGGTCACACTG | chr3R:14225762-14227762 | 952 | 966 | - | 4.0571 | 3.01E-05 | 0.673 | GTTACCGTCGCAATG |
| 1 | GMTACGGTCACACTG | chr3R:6656882-6658882 | 955 | 969 | - | 4.0571 | 3.01E-05 | 0.673 | TGTTCGGTCACACTA |
| 1 | GMTACGGTCACACTG | chr2L:6913138-6915138 | 957 | 971 | - | 4.0571 | 3.01E-05 | 0.673 | GGTACGGCCACATTA |
| 1 | GMTACGGTCACACTG | chr3L:3325265-3327265 | 960 | 974 | - | 4.0571 | 3.01E-05 | 0.673 | GGTCTGGTCACACTA |
| 1 | GMTACGGTCACACTG | chr3R:22546846-22548846 | 963 | 977 | - | 4.0571 | 3.01E-05 | 0.673 | CTTGCAGTCACACTG |
| 1 | GMTACGGTCACACTG | chr3L:9133560-9135560 | 967 | 981 | - | 4.0571 | 3.01E-05 | 0.673 | AGCAAGGTCACACTG |
| 1 | GMTACGGTCACACTG | chr3R:31052915-31054915 | 968 | 982 | - | 4.0571 | 3.01E-05 | 0.673 | TTTAAGGTCACTCTG |
| 1 | GMTACGGTCACACTG | chrX:11723246-11725246 | 968 | 982 | - | 4.0571 | 3.01E-05 | 0.673 | GGTACGGTAACACAA |

| | | | | | | | | | |
|---|---|---|---|---|---|---|---|---|---|
| 1 | GMTACGGTCACACTG | chr3R:27681029-27683029 | 968 | 982 | - | 4.0571 | 3.01E-05 | 0.673 | AGTGCGGTCACACTT |
| 1 | GMTACGGTCACACTG | chr3L:16121002-16123002 | 971 | 985 | - | 4.0571 | 3.01E-05 | 0.673 | AGTCCGGTCACACTA |
| 1 | GMTACGGTCACACTG | chr3R:22461054-22463054 | 973 | 987 | - | 4.0571 | 3.01E-05 | 0.673 | TTTGGGGTCACACTG |
| 1 | GMTACGGTCACACTG | chr2R:9102238-9104238 | 973 | 987 | - | 4.0571 | 3.01E-05 | 0.673 | AGCACGGTCACATTG |
| 1 | GMTACGGTCACACTG | chr2R:24774265-24776265 | 976 | 990 | - | 4.0571 | 3.01E-05 | 0.673 | CGAACGGTCACACTA |
| 1 | GMTACGGTCACACTG | chr3L:16231038-16233038 | 977 | 991 | - | 4.0571 | 3.01E-05 | 0.673 | AGCAAGGTCACACTG |
| 1 | GMTACGGTCACACTG | chr3R:15962884-15964884 | 979 | 993 | - | 4.0571 | 3.01E-05 | 0.673 | GGTATGGTCACCCTT |
| 1 | GMTACGGTCACACTG | chr2L:6489843-6491843 | 980 | 994 | - | 4.0571 | 3.01E-05 | 0.673 | GTAAAGGTCACACTA |
| 1 | GMTACGGTCACACTG | chr3L:18747687-18749687 | 981 | 995 | - | 4.0571 | 3.01E-05 | 0.673 | GGTGCGGTCATACTA |
| 1 | GMTACGGTCACACTG | chr2R:9816300-9818300 | 981 | 995 | - | 4.0571 | 3.01E-05 | 0.673 | TGTCCGGTCAAACTG |
| 1 | GMTACGGTCACACTG | chr2L:16728681-16730681 | 981 | 995 | - | 4.0571 | 3.01E-05 | 0.673 | GTCATGGTCACACTA |
| 1 | GMTACGGTCACACTG | chr3L:10898123-10900123 | 981 | 995 | - | 4.0571 | 3.01E-05 | 0.673 | CGCACGGTCACACTC |
| 1 | GMTACGGTCACACTG | chr2L:7408809-7410809 | 984 | 998 | - | 4.0571 | 3.01E-05 | 0.673 | GGTGAGGTCACACTA |
| 1 | GMTACGGTCACACTG | chr3R:29798354-29800354 | 989 | 1003 | - | 4.0571 | 3.01E-05 | 0.673 | TGCACGGTCACACTC |
| 1 | GMTACGGTCACACTG | chr3R:17810134-17812134 | 989 | 1003 | - | 4.0571 | 3.01E-05 | 0.673 | AGTACGGTCCCACTA |
| 1 | GMTACGGTCACACTG | chr2R:10644427-10646427 | 990 | 1004 | - | 4.0571 | 3.01E-05 | 0.673 | TGTTTGGTCACACTG |
| 1 | GMTACGGTCACACTG | chr2R:21659007-21661007 | 990 | 1004 | - | 4.0571 | 3.01E-05 | 0.673 | GGTGTGGTGACACTG |
| 1 | GMTACGGTCACACTG | chr3R:16340988-16342988 | 991 | 1005 | - | 4.0571 | 3.01E-05 | 0.673 | CTAACGGTCATACTG |
| 1 | GMTACGGTCACACTG | chr2L:5057248-5059248 | 992 | 1006 | - | 4.0571 | 3.01E-05 | 0.673 | TTTAGGGTCACACCG |
| 1 | GMTACGGTCACACTG | chr2R:8723258-8725258 | 992 | 1006 | - | 4.0571 | 3.01E-05 | 0.673 | GTTATGGTCACCCTA |
| 1 | GMTACGGTCACACTG | chr3L:12500311-12502311 | 993 | 1007 | - | 4.0571 | 3.01E-05 | 0.673 | GTTTTGGTCACACTA |
| 1 | GMTACGGTCACACTG | chr2R:13857689-13859689 | 993 | 1007 | - | 4.0571 | 3.01E-05 | 0.673 | TTAATGGTCACACTG |
| 1 | GMTACGGTCACACTG | chr3R:19903868-19905868 | 993 | 1007 | - | 4.0571 | 3.01E-05 | 0.673 | TGTGGGGTCACACTG |
| 1 | GMTACGGTCACACTG | chr3L:11168625-11170625 | 993 | 1007 | - | 4.0571 | 3.01E-05 | 0.673 | CGCAAGGTCACACTG |
| 1 | GMTACGGTCACACTG | chr2R:17000941-17002941 | 993 | 1007 | - | 4.0571 | 3.01E-05 | 0.673 | ATGACGGTCACACTA |
| 1 | GMTACGGTCACACTG | chr2R:13151677-13153677 | 995 | 1009 | - | 4.0571 | 3.01E-05 | 0.673 | GTGACGGTCATACTT |
| 1 | GMTACGGTCACACTG | chr3L:10224240-10226240 | 996 | 1010 | - | 4.0571 | 3.01E-05 | 0.673 | GTCACGGTTACACTC |
| 1 | GMTACGGTCACACTG | chr2R:9890689-9892689 | 996 | 1010 | - | 4.0571 | 3.01E-05 | 0.673 | CGCACGGTCACACTC |
| 1 | GMTACGGTCACACTG | chr3R:15962867-15964867 | 996 | 1010 | - | 4.0571 | 3.01E-05 | 0.673 | GGTATGGTCACCCTT |
| 1 | GMTACGGTCACACTG | chr3L:21026915-21028915 | 998 | 1012 | - | 4.0571 | 3.01E-05 | 0.673 | AGTACGGTCACTCTT |
| 1 | GMTACGGTCACACTG | chrX:3484378-3486378 | 1000 | 1014 | - | 4.0571 | 3.01E-05 | 0.673 | CGCACAGTCACACTG |
| 1 | GMTACGGTCACACTG | chr2L:4393196-4395196 | 1001 | 1015 | - | 4.0571 | 3.01E-05 | 0.673 | AGTACGGTCACTCTA |
| 1 | GMTACGGTCACACTG | chr3R:28843748-28845748 | 1003 | 1017 | - | 4.0571 | 3.01E-05 | 0.673 | ATAACAGTCACACTG |

| | | | | | | | | | |
|---|---|---|---|---|---|---|---|---|---|
| 1 | GMTACGGTCACACTG | chr2L:19999448-20001448 | 1004 | 1018 | - | 4.0571 | 3.01E-05 | 0.673 | GTTCCAGTCACACTC |
| 1 | GMTACGGTCACACTG | chr2L:12001266-12003266 | 1005 | 1019 | - | 4.0571 | 3.01E-05 | 0.673 | CGTGCGGTCATACTG |
| 1 | GMTACGGTCACACTG | chr3L:11073388-11075388 | 1010 | 1024 | - | 4.0571 | 3.01E-05 | 0.673 | TTTACGGTCTTACTG |
| 1 | GMTACGGTCACACTG | chr3R:25032465-25034465 | 1011 | 1025 | - | 4.0571 | 3.01E-05 | 0.673 | GGTCTGGTCACACTC |
| 1 | GMTACGGTCACACTG | chr3R:10871381-10873381 | 1011 | 1025 | - | 4.0571 | 3.01E-05 | 0.673 | TGTCTGGTCACACTG |
| 1 | GMTACGGTCACACTG | chr2R:6879995-6881995 | 1013 | 1027 | - | 4.0571 | 3.01E-05 | 0.673 | GTTTCCGTCAAACTG |
| 1 | GMTACGGTCACACTG | chr2L:8125890-8127890 | 1018 | 1032 | - | 4.0571 | 3.01E-05 | 0.673 | GTTATGGTCACCCTA |
| 1 | GMTACGGTCACACTG | chrX:1939307-1941307 | 1020 | 1034 | - | 4.0571 | 3.01E-05 | 0.673 | GTTCCGCTCACAATG |
| 1 | GMTACGGTCACACTG | chr2L:22536668-22538668 | 1026 | 1040 | - | 4.0571 | 3.01E-05 | 0.673 | GTTTGGGTCACATTG |
| 1 | GMTACGGTCACACTG | chr3L:8298839-8300839 | 1026 | 1040 | - | 4.0571 | 3.01E-05 | 0.673 | GTTACGGTCTCATTA |
| 1 | GMTACGGTCACACTG | chr3L:13928000-13930000 | 1031 | 1045 | - | 4.0571 | 3.01E-05 | 0.673 | GTTCCGGTCACAGTT |
| 1 | GMTACGGTCACACTG | chr3R:2156206-2158206 | 1032 | 1046 | - | 4.0571 | 3.01E-05 | 0.673 | GTTTCGGTCACATTC |
| 1 | GMTACGGTCACACTG | chr3L:19597755-19599755 | 1032 | 1046 | - | 4.0571 | 3.01E-05 | 0.673 | CGCAAGGTCACACTG |
| 1 | GMTACGGTCACACTG | chr3L:6192179-6194179 | 1033 | 1047 | - | 4.0571 | 3.01E-05 | 0.673 | TGCACGGTCACACTC |
| 1 | GMTACGGTCACACTG | chr2R:19450600-19452600 | 1036 | 1050 | - | 4.0571 | 3.01E-05 | 0.673 | AGTATGGCCACACTG |
| 1 | GMTACGGTCACACTG | chr3R:14696292-14698292 | 1037 | 1051 | - | 4.0571 | 3.01E-05 | 0.673 | CGTCTGGTCACACTG |
| 1 | GMTACGGTCACACTG | chr2L:11787334-11789334 | 1051 | 1065 | - | 4.0571 | 3.01E-05 | 0.673 | GTAATGGTCACACTT |
| 1 | GMTACGGTCACACTG | chr2L:1499608-1501608 | 1074 | 1088 | - | 4.0571 | 3.01E-05 | 0.673 | TGAACGGTCACATTG |
| 1 | GMTACGGTCACACTG | chr2R:23210539-23212539 | 1079 | 1093 | - | 4.0571 | 3.01E-05 | 0.673 | AGTGCGGTCACATTG |
| 1 | GMTACGGTCACACTG | chr2R:13845255-13847255 | 1088 | 1102 | - | 4.0571 | 3.01E-05 | 0.673 | ATGATGGTCACACTG |
| 1 | GMTACGGTCACACTG | chrX:19546482-19548482 | 1089 | 1103 | - | 4.0571 | 3.01E-05 | 0.673 | GTTACGGCCAAACTT |
| 1 | GMTACGGTCACACTG | chr2L:9527515-9529515 | 1091 | 1105 | - | 4.0571 | 3.01E-05 | 0.673 | AGTAGGGTCCCACTG |
| 1 | GMTACGGTCACACTG | chr2R:15679084-15681084 | 1096 | 1110 | - | 4.0571 | 3.01E-05 | 0.673 | CTTACGGTCACACCA |
| 1 | GMTACGGTCACACTG | chr3R:27676522-27678522 | 1098 | 1112 | - | 4.0571 | 3.01E-05 | 0.673 | GTAACGGTGACAGTG |
| 1 | GMTACGGTCACACTG | chr3R:9745367-9747367 | 1115 | 1129 | - | 4.0571 | 3.01E-05 | 0.673 | GTTACAGTCACCGTG |
| 1 | GMTACGGTCACACTG | chr3R:16974204-16976204 | 1121 | 1135 | - | 4.0571 | 3.01E-05 | 0.673 | GTTAGGATCACACAG |
| 1 | GMTACGGTCACACTG | chr3L:9133405-9135405 | 1122 | 1136 | - | 4.0571 | 3.01E-05 | 0.673 | AGCAAGGTCACACTG |
| 1 | GMTACGGTCACACTG | chr2R:9074951-9076951 | 1132 | 1146 | - | 4.0571 | 3.01E-05 | 0.673 | AGTACGGTCACACAC |
| 1 | GMTACGGTCACACTG | chr3L:13290770-13292770 | 1175 | 1189 | - | 4.0571 | 3.01E-05 | 0.673 | TGTTCTGTCACACTG |
| 1 | GMTACGGTCACACTG | chr2L:304935-306935 | 1180 | 1194 | - | 4.0571 | 3.01E-05 | 0.673 | GTTACTCTCACGCTG |
| 1 | GMTACGGTCACACTG | chrX:3670502-3672502 | 1188 | 1202 | - | 4.0571 | 3.01E-05 | 0.673 | GTAATGGTCACACTT |
| 1 | GMTACGGTCACACTG | chr3R:19161300-19163300 | 1199 | 1213 | - | 4.0571 | 3.01E-05 | 0.673 | CGTATGGCCACACTG |
| 1 | GMTACGGTCACACTG | chr3R:19161300-19163300 | 1199 | 1213 | - | 4.0571 | 3.01E-05 | 0.673 | CGTATGGCCACACTG |

| | | | | | | | | | |
|---|---|---|---|---|---|---|---|---|---|
| 1 | GMTACGGTCACACTG | chr3L:19263184-19265184 | 1214 | 1228 | - | 4.0571 | 3.01E-05 | 0.673 | AGTGTGGTCACACTG |
| 1 | GMTACGGTCACACTG | chr2R:14255192-14257192 | 1230 | 1244 | - | 4.0571 | 3.01E-05 | 0.673 | TGTACGGTCTCACTA |
| 1 | GMTACGGTCACACTG | chr2R:6879770-6881770 | 1238 | 1252 | - | 4.0571 | 3.01E-05 | 0.673 | GTTTCCGTCAAACTG |
| 1 | GMTACGGTCACACTG | chr3R:5567615-5569615 | 1249 | 1263 | - | 4.0571 | 3.01E-05 | 0.673 | GGAACCGTCACACTA |
| 1 | GMTACGGTCACACTG | chr3R:18198857-18200857 | 1260 | 1274 | - | 4.0571 | 3.01E-05 | 0.673 | CGTCTGGTCACACTG |
| 1 | GMTACGGTCACACTG | chr2L:12452664-12454664 | 1261 | 1275 | - | 4.0571 | 3.01E-05 | 0.673 | GTTATCGCCACACTG |
| 1 | GMTACGGTCACACTG | chr2R:16872015-16874015 | 1264 | 1278 | - | 4.0571 | 3.01E-05 | 0.673 | GGTATGGTCAATCTG |
| 1 | GMTACGGTCACACTG | chrX:1939040-1941040 | 1287 | 1301 | - | 4.0571 | 3.01E-05 | 0.673 | GTTCCGCTCACAATG |
| 1 | GMTACGGTCACACTG | chr3R:13278721-13280721 | 1290 | 1304 | - | 4.0571 | 3.01E-05 | 0.673 | GGTTCGCTCACACTA |
| 1 | GMTACGGTCACACTG | chr3L:18747367-18749367 | 1301 | 1315 | - | 4.0571 | 3.01E-05 | 0.673 | GGTGCGGTCATACTA |
| 1 | GMTACGGTCACACTG | chrX:4678537-4680537 | 1314 | 1328 | - | 4.0571 | 3.01E-05 | 0.673 | AGCATGGTCACACTG |
| 1 | GMTACGGTCACACTG | chr2R:23384468-23386468 | 1316 | 1330 | - | 4.0571 | 3.01E-05 | 0.673 | AGTTCGGTCACACTA |
| 1 | GMTACGGTCACACTG | chr2L:4392876-4394876 | 1321 | 1335 | - | 4.0571 | 3.01E-05 | 0.673 | AGTACGGTCACTCTA |
| 1 | GMTACGGTCACACTG | chr3R:26880171-26882171 | 1356 | 1370 | - | 4.0571 | 3.01E-05 | 0.673 | TGTACGCACACACTG |
| 1 | GMTACGGTCACACTG | chrX:16639419-16641419 | 1379 | 1393 | - | 4.0571 | 3.01E-05 | 0.673 | GGTATGTTCGCACTG |
| 1 | GMTACGGTCACACTG | chr3R:16369533-16371533 | 1388 | 1402 | - | 4.0571 | 3.01E-05 | 0.673 | GGTACGGTAATGCTG |
| 1 | GMTACGGTCACACTG | chr3R:6656442-6658442 | 1395 | 1409 | - | 4.0571 | 3.01E-05 | 0.673 | TGTTCGGTCACACTA |
| 1 | GMTACGGTCACACTG | chr2L:2288416-2290416 | 1397 | 1411 | - | 4.0571 | 3.01E-05 | 0.673 | GGTAAAGTCACACGG |
| 1 | GMTACGGTCACACTG | chr2R:8722839-8724839 | 1411 | 1425 | - | 4.0571 | 3.01E-05 | 0.673 | GTTATGGTCACCCTA |
| 1 | GMTACGGTCACACTG | chr3R:7109012-7111012 | 1426 | 1440 | - | 4.0571 | 3.01E-05 | 0.673 | ATTACGGTCGCAATG |
| 1 | GMTACGGTCACACTG | chr2R:8122036-8124036 | 1439 | 1453 | - | 4.0571 | 3.01E-05 | 0.673 | GGTGCCGTCACATTG |
| 1 | GMTACGGTCACACTG | chr2R:8122001-8124001 | 1474 | 1488 | - | 4.0571 | 3.01E-05 | 0.673 | GGTGCCGTCACATTG |
| 1 | GMTACGGTCACACTG | chr3L:21026434-21028434 | 1479 | 1493 | - | 4.0571 | 3.01E-05 | 0.673 | AGTACGGTCACTCTT |
| 1 | GMTACGGTCACACTG | chr3L:9008838-9010838 | 1494 | 1508 | - | 4.0571 | 3.01E-05 | 0.673 | GGTCCGTTCACATTG |
| 1 | GMTACGGTCACACTG | chr3R:26188977-26190977 | 1499 | 1513 | - | 4.0571 | 3.01E-05 | 0.673 | CTTCGGGTCACACTG |
| 1 | GMTACGGTCACACTG | chrX:3347400-3349400 | 1514 | 1528 | - | 4.0571 | 3.01E-05 | 0.673 | TGTGTGGTCACACTG |
| 1 | GMTACGGTCACACTG | chr3R:18198595-18200595 | 1522 | 1536 | - | 4.0571 | 3.01E-05 | 0.673 | CGTCTGGTCACACTG |
| 1 | GMTACGGTCACACTG | chr3L:890250-892250 | 1544 | 1558 | - | 4.0571 | 3.01E-05 | 0.673 | GGTACGGTAATGCTG |
| 1 | GMTACGGTCACACTG | chr2L:18697714-18699714 | 1560 | 1574 | - | 4.0571 | 3.01E-05 | 0.673 | GGTGCGGTCACAGTA |
| 1 | GMTACGGTCACACTG | chr3L:5881304-5883304 | 1561 | 1575 | - | 4.0571 | 3.01E-05 | 0.673 | GGTATGACCACACTG |
| 1 | GMTACGGTCACACTG | chrX:10052811-10054811 | 1566 | 1580 | - | 4.0571 | 3.01E-05 | 0.673 | AGTCCGGTCACAATG |
| 1 | GMTACGGTCACACTG | chr3R:24924344-24926344 | 1574 | 1588 | - | 4.0571 | 3.01E-05 | 0.673 | AGGACGGTCACAGTG |
| 1 | GMTACGGTCACACTG | chr2L:13962507-13964507 | 1594 | 1608 | - | 4.0571 | 3.01E-05 | 0.673 | GTTACGGACAAACTC |

| | | | | | | | | | |
|---|---|---|---|---|---|---|---|---|---|
| 1 | GMTACGGTCACACTG | chr3R:27999302-28001302 | 1597 | 1611 | - | 4.0571 | 3.01E-05 | 0.673 | GTTACTGTGCCACTG |
| 1 | GMTACGGTCACACTG | chr3R:8663115-8665115 | 1601 | 1615 | - | 4.0571 | 3.01E-05 | 0.673 | GGTATGGACACAATG |
| 1 | GMTACGGTCACACTG | chrX:18485921-18487921 | 1603 | 1617 | - | 4.0571 | 3.01E-05 | 0.673 | AGCATGGTCACACTG |
| 1 | GMTACGGTCACACTG | chr2L:13365447-13367447 | 1603 | 1617 | - | 4.0571 | 3.01E-05 | 0.673 | GTTTCGTTCACAATG |
| 1 | GMTACGGTCACACTG | chr3L:19262766-19264766 | 1632 | 1646 | - | 4.0571 | 3.01E-05 | 0.673 | AGTGTGGTCACACTG |
| 1 | GMTACGGTCACACTG | chr3L:19262766-19264766 | 1632 | 1646 | - | 4.0571 | 3.01E-05 | 0.673 | AGTGTGGTCACACTG |
| 1 | GMTACGGTCACACTG | chr3L:3325265-3327265 | 1677 | 1691 | - | 4.0571 | 3.01E-05 | 0.673 | GTTACAGTGACGCTG |
| 1 | GMTACGGTCACACTG | chr2L:12452242-12454242 | 1683 | 1697 | - | 4.0571 | 3.01E-05 | 0.673 | GTTATCGCCACACTG |
| 1 | GMTACGGTCACACTG | chr3L:21267601-21269601 | 1686 | 1700 | - | 4.0571 | 3.01E-05 | 0.673 | GGTATGGTGACCCTG |
| 1 | GMTACGGTCACACTG | chr3L:7087787-7089787 | 1689 | 1703 | - | 4.0571 | 3.01E-05 | 0.673 | GTTATGGGCAGACTG |
| 1 | GMTACGGTCACACTG | chr3R:8088757-8090757 | 1692 | 1706 | - | 4.0571 | 3.01E-05 | 0.673 | GGTACCGAAACACTG |
| 1 | GMTACGGTCACACTG | chr2R:15693092-15695092 | 1700 | 1714 | - | 4.0571 | 3.01E-05 | 0.673 | GTTAAGGACACATTG |
| 1 | GMTACGGTCACACTG | chrX:15856823-15858823 | 1701 | 1715 | - | 4.0571 | 3.01E-05 | 0.673 | GTTTTGGTCAGACTG |
| 1 | GMTACGGTCACACTG | chr3R:24924216-24926216 | 1702 | 1716 | - | 4.0571 | 3.01E-05 | 0.673 | AGGACGGTCACAGTG |
| 1 | GMTACGGTCACACTG | chr2R:19602295-19604295 | 1704 | 1718 | - | 4.0571 | 3.01E-05 | 0.673 | GTTGCGGTTACACTT |
| 1 | GMTACGGTCACACTG | chr3L:23125025-23127025 | 1712 | 1726 | - | 4.0571 | 3.01E-05 | 0.673 | ATTACCGTCACTCTG |
| 1 | GMTACGGTCACACTG | chr2R:22797289-22799289 | 1727 | 1741 | - | 4.0571 | 3.01E-05 | 0.673 | AGTACGGTCACCCTC |
| 1 | GMTACGGTCACACTG | chr3R:30730165-30732165 | 1733 | 1747 | - | 4.0571 | 3.01E-05 | 0.673 | TGTACGGGGACACTG |
| 1 | GMTACGGTCACACTG | chr3L:3892330-3894330 | 1751 | 1765 | - | 4.0571 | 3.01E-05 | 0.673 | GGTATGGCCACAGTG |
| 1 | GMTACGGTCACACTG | chr2L:11737948-11739948 | 1764 | 1778 | - | 4.0571 | 3.01E-05 | 0.673 | ATTACGGTTTCACTG |
| 1 | GMTACGGTCACACTG | chr2L:8304586-8306586 | 1771 | 1785 | - | 4.0571 | 3.01E-05 | 0.673 | GTTACTGGCATACTG |
| 1 | GMTACGGTCACACTG | chr2L:22535884-22537884 | 1810 | 1824 | - | 4.0571 | 3.01E-05 | 0.673 | GTTTGGGTCACATTG |
| 1 | GMTACGGTCACACTG | chr2R:11306879-11308879 | 1839 | 1853 | - | 4.0571 | 3.01E-05 | 0.673 | GTTGAGGTCAGACTG |
| 1 | GMTACGGTCACACTG | chr2L:13365189-13367189 | 1861 | 1875 | - | 4.0571 | 3.01E-05 | 0.673 | GTTTCGTTCACAATG |
| 1 | GMTACGGTCACACTG | chrX:6065416-6067416 | 1865 | 1879 | - | 4.0571 | 3.01E-05 | 0.673 | ATTACGACCACACTG |
| 1 | GMTACGGTCACACTG | chr3L:15534395-15536395 | 1893 | 1907 | - | 4.0571 | 3.01E-05 | 0.673 | GTTGCAGTCACACTA |
| 1 | GMTACGGTCACACTG | chr2R:15692887-15694887 | 1905 | 1919 | - | 4.0571 | 3.01E-05 | 0.673 | GTTAAGGACACATTG |
| 1 | GMTACGGTCACACTG | chr3L:22496114-22498114 | 1933 | 1947 | - | 4.0571 | 3.01E-05 | 0.673 | ATTACTGTCACACGG |
| 1 | GMTACGGTCACACTG | chr3L:21267335-21269335 | 1952 | 1966 | - | 4.0571 | 3.01E-05 | 0.673 | GGTATGGTGACCCTG |
| 1 | GMTACGGTCACACTG | chrX:6066958-6068958 | 1953 | 1967 | - | 4.0571 | 3.01E-05 | 0.673 | GGTACTGTCAATCTG |
| 1 | GMTACGGTCACACTG | chr3R:8725854-8727854 | 1957 | 1971 | - | 4.0571 | 3.01E-05 | 0.673 | GGTGCGGTCACAGTA |
| 1 | GMTACGGTCACACTG | chr2R:10808397-10810397 | 1972 | 1986 | - | 4.0571 | 3.01E-05 | 0.673 | GGCACGGTCATAATG |
| 1 | GMTACGGTCACACTG | chr3R:24825471-24827471 | 7 | 21 | + | 4.0571 | 3.01E-05 | 0.673 | GTTGTGGTCACAATG |

| | | | | | | | | | |
|---|---|---|---|---|---|---|---|---|---|
| 1 | GMTACGGTCACACTG | chr2L:21262933-21264933 | 7 | 21 | + | 4.0571 | 3.01E-05 | 0.673 | GGTACAGTCCTACTG |
| 1 | GMTACGGTCACACTG | chrX:2680723-2682723 | 20 | 34 | + | 4.0571 | 3.01E-05 | 0.673 | GGTGCGTTCACACTT |
| 1 | GMTACGGTCACACTG | chrX:7963024-7965024 | 27 | 41 | + | 4.0571 | 3.01E-05 | 0.673 | GGCATAGTCACACTG |
| 1 | GMTACGGTCACACTG | chr3R:30866521-30868521 | 38 | 52 | + | 4.0571 | 3.01E-05 | 0.673 | ATTGCGGTAACACTG |
| 1 | GMTACGGTCACACTG | chrX:9594082-9596082 | 47 | 61 | + | 4.0571 | 3.01E-05 | 0.673 | GTTACGTAAACACTG |
| 1 | GMTACGGTCACACTG | chr3R:13381106-13383106 | 61 | 75 | + | 4.0571 | 3.01E-05 | 0.673 | GGAAAGGTCACACTC |
| 1 | GMTACGGTCACACTG | chr3L:13449770-13451770 | 83 | 97 | + | 4.0571 | 3.01E-05 | 0.673 | GGTAGGTCCACACTG |
| 1 | GMTACGGTCACACTG | chr3L:11943852-11945852 | 104 | 118 | + | 4.0571 | 3.01E-05 | 0.673 | GTTCTCGTCACACTG |
| 1 | GMTACGGTCACACTG | chr2R:7785322-7787322 | 105 | 119 | + | 4.0571 | 3.01E-05 | 0.673 | GTTACGCGCACACTT |
| 1 | GMTACGGTCACACTG | chr3L:8305980-8307980 | 202 | 216 | + | 4.0571 | 3.01E-05 | 0.673 | GTTGCTGTCATACTG |
| 1 | GMTACGGTCACACTG | chr2L:11337920-11339920 | 216 | 230 | + | 4.0571 | 3.01E-05 | 0.673 | GTTAGGGTCACGGTG |
| 1 | GMTACGGTCACACTG | chr2R:16972927-16974927 | 221 | 235 | + | 4.0571 | 3.01E-05 | 0.673 | GTTTCGGTCAAAATG |
| 1 | GMTACGGTCACACTG | chr2L:7362379-7364379 | 246 | 260 | + | 4.0571 | 3.01E-05 | 0.673 | GGAATGGTGACACTG |
| 1 | GMTACGGTCACACTG | chr3L:7731191-7733191 | 274 | 288 | + | 4.0571 | 3.01E-05 | 0.673 | GGTACTGTTGCACTG |
| 1 | GMTACGGTCACACTG | chr4:24054-26054 | 286 | 300 | + | 4.0571 | 3.01E-05 | 0.673 | GTTACGCACAAACTG |
| 1 | GMTACGGTCACACTG | chr2R:19495856-19497856 | 297 | 311 | + | 4.0571 | 3.01E-05 | 0.673 | GTTCCGATCACAGTG |
| 1 | GMTACGGTCACACTG | chr3L:8068267-8070267 | 297 | 311 | + | 4.0571 | 3.01E-05 | 0.673 | GTTATGGACACATTG |
| 1 | GMTACGGTCACACTG | chr3R:31122415-31124415 | 298 | 312 | + | 4.0571 | 3.01E-05 | 0.673 | GTTACCGTAACGCTG |
| 1 | GMTACGGTCACACTG | chr4:10765-12765 | 303 | 317 | + | 4.0571 | 3.01E-05 | 0.673 | GTTACGCACAAACTG |
| 1 | GMTACGGTCACACTG | chr2L:1158355-1160355 | 316 | 330 | + | 4.0571 | 3.01E-05 | 0.673 | CGCATGGTCACACTG |
| 1 | GMTACGGTCACACTG | chr3R:24363980-24365980 | 319 | 333 | + | 4.0571 | 3.01E-05 | 0.673 | TTTACGATCTCACTG |
| 1 | GMTACGGTCACACTG | chrX:17378004-17380004 | 347 | 361 | + | 4.0571 | 3.01E-05 | 0.673 | GTTAAGGGTACACTG |
| 1 | GMTACGGTCACACTG | chr3R:21618321-21620321 | 349 | 363 | + | 4.0571 | 3.01E-05 | 0.673 | GTTACGTTCTCACTA |
| 1 | GMTACGGTCACACTG | chr3R:10082381-10084381 | 351 | 365 | + | 4.0571 | 3.01E-05 | 0.673 | GGTACAGTAACCCTG |
| 1 | GMTACGGTCACACTG | chr3R:13380809-13382809 | 358 | 372 | + | 4.0571 | 3.01E-05 | 0.673 | GGAAAGGTCACACTC |
| 1 | GMTACGGTCACACTG | chr3R:21568149-21570149 | 360 | 374 | + | 4.0571 | 3.01E-05 | 0.673 | GGTACGGTCATCCTC |
| 1 | GMTACGGTCACACTG | chr2R:7253602-7255602 | 401 | 415 | + | 4.0571 | 3.01E-05 | 0.673 | GGTACGGTCGAACTC |
| 1 | GMTACGGTCACACTG | chr2L:17962379-17964379 | 408 | 422 | + | 4.0571 | 3.01E-05 | 0.673 | GTTCCGGTCACAATA |
| 1 | GMTACGGTCACACTG | chr3R:5567921-5569921 | 413 | 427 | + | 4.0571 | 3.01E-05 | 0.673 | GGTACGGTCAGACGC |
| 1 | GMTACGGTCACACTG | chr2R:7785010-7787010 | 417 | 431 | + | 4.0571 | 3.01E-05 | 0.673 | GTTACGCGCACACTT |
| 1 | GMTACGGTCACACTG | chr2R:16666675-16668675 | 418 | 432 | + | 4.0571 | 3.01E-05 | 0.673 | GTTACGGTTACACAA |
| 1 | GMTACGGTCACACTG | chr3L:12692639-12694639 | 433 | 447 | + | 4.0571 | 3.01E-05 | 0.673 | TTTATGGTCACACAG |
| 1 | GMTACGGTCACACTG | chr3L:13845054-13847054 | 464 | 478 | + | 4.0571 | 3.01E-05 | 0.673 | TTAACGGTCACAATG |

| | | | | | | | | | |
|---|---|---|---|---|---|---|---|---|---|
| 1 | GMTACGGTCACACTG | chr3L:2265394-2267394 | 484 | 498 | + | 4.0571 | 3.01E-05 | 0.673 | AGTTCGTTCACACTG |
| 1 | GMTACGGTCACACTG | chr3L:7362598-7364598 | 488 | 502 | + | 4.0571 | 3.01E-05 | 0.673 | GTTACGGTCACGGTT |
| 1 | GMTACGGTCACACTG | chr2R:21537015-21539015 | 505 | 519 | + | 4.0571 | 3.01E-05 | 0.673 | GTTCTGGTCACATTG |
| 1 | GMTACGGTCACACTG | chr3R:11948680-11950680 | 545 | 559 | + | 4.0571 | 3.01E-05 | 0.673 | CGTACGGCCACACTT |
| 1 | GMTACGGTCACACTG | chr3L:13449283-13451283 | 570 | 584 | + | 4.0571 | 3.01E-05 | 0.673 | GGTAGGTCCACACTG |
| 1 | GMTACGGTCACACTG | chr2L:8010405-8012405 | 608 | 622 | + | 4.0571 | 3.01E-05 | 0.673 | GTCAAGGTCACAATG |
| 1 | GMTACGGTCACACTG | chr2R:16335637-16337637 | 631 | 645 | + | 4.0571 | 3.01E-05 | 0.673 | TTCAGGGTCACACTG |
| 1 | GMTACGGTCACACTG | chr3R:16341318-16343318 | 632 | 646 | + | 4.0571 | 3.01E-05 | 0.673 | GTCACGCTCACAATG |
| 1 | GMTACGGTCACACTG | chr3R:10872038-10874038 | 633 | 647 | + | 4.0571 | 3.01E-05 | 0.673 | TTTTTGGTCACACTG |
| 1 | GMTACGGTCACACTG | chr3L:6605032-6607032 | 636 | 650 | + | 4.0571 | 3.01E-05 | 0.673 | GTCACAGTCACATTG |
| 1 | GMTACGGTCACACTG | chrX:15908676-15910676 | 638 | 652 | + | 4.0571 | 3.01E-05 | 0.673 | AGTATGGGCACACTG |
| 1 | GMTACGGTCACACTG | chr3R:15666978-15668978 | 652 | 666 | + | 4.0571 | 3.01E-05 | 0.673 | CTTGCGGTCACAATG |
| 1 | GMTACGGTCACACTG | chr3L:22852512-22854512 | 668 | 682 | + | 4.0571 | 3.01E-05 | 0.673 | GGTACTGGCACACTC |
| 1 | GMTACGGTCACACTG | chr2L:9847289-9849289 | 671 | 685 | + | 4.0571 | 3.01E-05 | 0.673 | GTAACTGTCAAACTG |
| 1 | GMTACGGTCACACTG | chrX:21338741-21340741 | 700 | 714 | + | 4.0571 | 3.01E-05 | 0.673 | TTAACGGTCACACTC |
| 1 | GMTACGGTCACACTG | chr3R:5567615-5569615 | 719 | 733 | + | 4.0571 | 3.01E-05 | 0.673 | GGTACGGTCAGACGC |
| 1 | GMTACGGTCACACTG | chr2R:11277445-11279445 | 735 | 749 | + | 4.0571 | 3.01E-05 | 0.673 | GTTAAGGTCACCCTT |
| 1 | GMTACGGTCACACTG | chr3R:10754634-10756634 | 746 | 760 | + | 4.0571 | 3.01E-05 | 0.673 | GTCAAGGTCACACTC |
| 1 | GMTACGGTCACACTG | chr2R:12592827-12594827 | 747 | 761 | + | 4.0571 | 3.01E-05 | 0.673 | CGCACAGTCACACTG |
| 1 | GMTACGGTCACACTG | chr3R:13785583-13787583 | 751 | 765 | + | 4.0571 | 3.01E-05 | 0.673 | TTGACAGTCACACTG |
| 1 | GMTACGGTCACACTG | chrX:21338676-21340676 | 765 | 779 | + | 4.0571 | 3.01E-05 | 0.673 | TTAACGGTCACACTC |
| 1 | GMTACGGTCACACTG | chr3R:24187896-24189896 | 787 | 801 | + | 4.0571 | 3.01E-05 | 0.673 | TGCACGGTCAGACTG |
| 1 | GMTACGGTCACACTG | chr3L:7935373-7937373 | 789 | 803 | + | 4.0571 | 3.01E-05 | 0.673 | TTTACGGTCGCACTT |
| 1 | GMTACGGTCACACTG | chr3R:11749333-11751333 | 806 | 820 | + | 4.0571 | 3.01E-05 | 0.673 | AGTATGGCCACACTG |
| 1 | GMTACGGTCACACTG | chr2R:7564751-7566751 | 808 | 822 | + | 4.0571 | 3.01E-05 | 0.673 | CTTAAGGTCACACAG |
| 1 | GMTACGGTCACACTG | chr2L:13903661-13905661 | 827 | 841 | + | 4.0571 | 3.01E-05 | 0.673 | TGCACGGTCAGACTG |
| 1 | GMTACGGTCACACTG | chr3R:24824648-24826648 | 830 | 844 | + | 4.0571 | 3.01E-05 | 0.673 | GTTGTGGTCACAATG |
| 1 | GMTACGGTCACACTG | chr3R:25081684-25083684 | 840 | 854 | + | 4.0571 | 3.01E-05 | 0.673 | GGTCTGGTCAGACTG |
| 1 | GMTACGGTCACACTG | chr3L:14134102-14136102 | 856 | 870 | + | 4.0571 | 3.01E-05 | 0.673 | GGTAAGGACAGACTG |
| 1 | GMTACGGTCACACTG | chrX:9249420-9251420 | 904 | 918 | + | 4.0571 | 3.01E-05 | 0.673 | ATTTTGGTCACACTG |
| 1 | GMTACGGTCACACTG | chr2L:3264714-3266714 | 918 | 932 | + | 4.0571 | 3.01E-05 | 0.673 | GTTTCGCTCACACTT |
| 1 | GMTACGGTCACACTG | chr2R:24113398-24115398 | 918 | 932 | + | 4.0571 | 3.01E-05 | 0.673 | TGAACGGTCAGACTG |
| 1 | GMTACGGTCACACTG | chr3R:24517270-24519270 | 927 | 941 | + | 4.0571 | 3.01E-05 | 0.673 | ATTCCGGTCACACTA |

| | | | | | | | | | |
|---|---|---|---|---|---|---|---|---|---|
| 1 | GMTACGGTCACACTG | chrX:16447665-16449665 | 930 | 944 | + | 4.0571 | 3.01E-05 | 0.673 | GGTGTGGTGACACTG |
| 1 | GMTACGGTCACACTG | chr3L:1946565-1948565 | 958 | 972 | + | 4.0571 | 3.01E-05 | 0.673 | TGAACGGTCACACTT |
| 1 | GMTACGGTCACACTG | chr3R:16340988-16342988 | 962 | 976 | + | 4.0571 | 3.01E-05 | 0.673 | GTCACGCTCACAATG |
| 1 | GMTACGGTCACACTG | chr3R:20359308-20361308 | 965 | 979 | + | 4.0571 | 3.01E-05 | 0.673 | GGCACGATCAAACTG |
| 1 | GMTACGGTCACACTG | chr3R:20559865-20561865 | 968 | 982 | + | 4.0571 | 3.01E-05 | 0.673 | GTTCTGGTCACACTT |
| 1 | GMTACGGTCACACTG | chr2R:12592604-12594604 | 970 | 984 | + | 4.0571 | 3.01E-05 | 0.673 | CGCACAGTCACACTG |
| 1 | GMTACGGTCACACTG | chrX:18514428-18516428 | 971 | 985 | + | 4.0571 | 3.01E-05 | 0.673 | CGAACGGTCACATTG |
| 1 | GMTACGGTCACACTG | chrX:18514428-18516428 | 971 | 985 | + | 4.0571 | 3.01E-05 | 0.673 | CGAACGGTCACATTG |
| 1 | GMTACGGTCACACTG | chr2L:13178247-13180247 | 976 | 990 | + | 4.0571 | 3.01E-05 | 0.673 | AGTACGGGCACACTA |
| 1 | GMTACGGTCACACTG | chr3L:17907143-17909143 | 977 | 991 | + | 4.0571 | 3.01E-05 | 0.673 | TGTACGGCTACACTG |
| 1 | GMTACGGTCACACTG | chr3L:327097-329097 | 978 | 992 | + | 4.0571 | 3.01E-05 | 0.673 | AGCACGGTCACACTT |
| 1 | GMTACGGTCACACTG | chr3L:3902492-3904492 | 978 | 992 | + | 4.0571 | 3.01E-05 | 0.673 | CGCACGGTCACACTA |
| 1 | GMTACGGTCACACTG | chr2L:6907083-6909083 | 978 | 992 | + | 4.0571 | 3.01E-05 | 0.673 | TGTACGGTCCCATTG |
| 1 | GMTACGGTCACACTG | chr2L:21261956-21263956 | 984 | 998 | + | 4.0571 | 3.01E-05 | 0.673 | GGTACAGTCCTACTG |
| 1 | GMTACGGTCACACTG | chr2L:8399421-8401421 | 986 | 1000 | + | 4.0571 | 3.01E-05 | 0.673 | GGTGTGGTTACACTG |
| 1 | GMTACGGTCACACTG | chr3L:211709-213709 | 986 | 1000 | + | 4.0571 | 3.01E-05 | 0.673 | AGTACGGTCCTACTG |
| 1 | GMTACGGTCACACTG | chr3L:1341403-1343403 | 987 | 1001 | + | 4.0571 | 3.01E-05 | 0.673 | ATTCCGGTCACACTA |
| 1 | GMTACGGTCACACTG | chr3R:12412828-12414828 | 988 | 1002 | + | 4.0571 | 3.01E-05 | 0.673 | AGTCTGGTCACACTG |
| 1 | GMTACGGTCACACTG | chr3L:9863639-9865639 | 991 | 1005 | + | 4.0571 | 3.01E-05 | 0.673 | AGCACGGTCACACTT |
| 1 | GMTACGGTCACACTG | chr2R:16123981-16125981 | 992 | 1006 | + | 4.0571 | 3.01E-05 | 0.673 | CGGACGGTCACACTT |
| 1 | GMTACGGTCACACTG | chr2R:9113274-9115274 | 993 | 1007 | + | 4.0571 | 3.01E-05 | 0.673 | AGTATGGTCACTCTG |
| 1 | GMTACGGTCACACTG | chr3R:5793705-5795705 | 993 | 1007 | + | 4.0571 | 3.01E-05 | 0.673 | AGCACGGTCATACTG |
| 1 | GMTACGGTCACACTG | chr2R:9890955-9892955 | 993 | 1007 | + | 4.0571 | 3.01E-05 | 0.673 | AGCACAGTCACACTG |
| 1 | GMTACGGTCACACTG | chr3R:29999530-30001530 | 994 | 1008 | + | 4.0571 | 3.01E-05 | 0.673 | TGGACGGTTACACTG |
| 1 | GMTACGGTCACACTG | chr2L:158032-160032 | 994 | 1008 | + | 4.0571 | 3.01E-05 | 0.673 | CTAACGGTCACATTG |
| 1 | GMTACGGTCACACTG | chr3L:3054079-3056079 | 994 | 1008 | + | 4.0571 | 3.01E-05 | 0.673 | GGTGTGGTGACACTG |
| 1 | GMTACGGTCACACTG | chrX:7012887-7014887 | 994 | 1008 | + | 4.0571 | 3.01E-05 | 0.673 | CGTCTGGTCACACTG |
| 1 | GMTACGGTCACACTG | chr3R:23932912-23934912 | 995 | 1009 | + | 4.0571 | 3.01E-05 | 0.673 | GGGATGGTCACACTA |
| 1 | GMTACGGTCACACTG | chr3L:3460317-3462317 | 999 | 1013 | + | 4.0571 | 3.01E-05 | 0.673 | AGTCCGGTCACACTT |
| 1 | GMTACGGTCACACTG | chrX:7012881-7014881 | 1000 | 1014 | + | 4.0571 | 3.01E-05 | 0.673 | CGTCTGGTCACACTG |
| 1 | GMTACGGTCACACTG | chrX:15066007-15068007 | 1003 | 1017 | + | 4.0571 | 3.01E-05 | 0.673 | TGCAAGGTCACACTG |
| 1 | GMTACGGTCACACTG | chr2R:10818978-10820978 | 1003 | 1017 | + | 4.0571 | 3.01E-05 | 0.673 | AGTACGGTCACTCTA |
| 1 | GMTACGGTCACACTG | chr3L:18871747-18873747 | 1006 | 1020 | + | 4.0571 | 3.01E-05 | 0.673 | AGTCTGGTCACACTG |

| | | | | | | | | | |
|---|---|---|---|---|---|---|---|---|---|
| 1 | GMTACGGTCACACTG | chr3R:13029913-13031913 | 1006 | 1020 | + | 4.0571 | 3.01E-05 | 0.673 | GTCAGGGTCACACTT |
| 1 | GMTACGGTCACACTG | chr3L:211687-213687 | 1008 | 1022 | + | 4.0571 | 3.01E-05 | 0.673 | AGTACGGTCCTACTG |
| 1 | GMTACGGTCACACTG | chrX:17860002-17862002 | 1009 | 1023 | + | 4.0571 | 3.01E-05 | 0.673 | CGTCTGGTCACACTG |
| 1 | GMTACGGTCACACTG | chr2L:1157661-1159661 | 1010 | 1024 | + | 4.0571 | 3.01E-05 | 0.673 | CGCATGGTCACACTG |
| 1 | GMTACGGTCACACTG | chr2L:1079911-1081911 | 1011 | 1025 | + | 4.0571 | 3.01E-05 | 0.673 | GTTGCGGTCATACTA |
| 1 | GMTACGGTCACACTG | chr2L:6648388-6650388 | 1019 | 1033 | + | 4.0571 | 3.01E-05 | 0.673 | AGTACGGTCACACGA |
| 1 | GMTACGGTCACACTG | chr3R:13051245-13053245 | 1020 | 1034 | + | 4.0571 | 3.01E-05 | 0.673 | TGAACGGTCACACTA |
| 1 | GMTACGGTCACACTG | chr3L:3041359-3043359 | 1020 | 1034 | + | 4.0571 | 3.01E-05 | 0.673 | ATTACGGTCACACCA |
| 1 | GMTACGGTCACACTG | chr3L:2264857-2266857 | 1021 | 1035 | + | 4.0571 | 3.01E-05 | 0.673 | AGTTCGTTCACACTG |
| 1 | GMTACGGTCACACTG | chr2R:24113292-24115292 | 1024 | 1038 | + | 4.0571 | 3.01E-05 | 0.673 | TGAACGGTCAGACTG |
| 1 | GMTACGGTCACACTG | chr3R:24624671-24626671 | 1028 | 1042 | + | 4.0571 | 3.01E-05 | 0.673 | TGTACGGTCACACCA |
| 1 | GMTACGGTCACACTG | chr2L:19547507-19549507 | 1031 | 1045 | + | 4.0571 | 3.01E-05 | 0.673 | GTTCCGTTAACACTG |
| 1 | GMTACGGTCACACTG | chr3L:11112740-11114740 | 1033 | 1047 | + | 4.0571 | 3.01E-05 | 0.673 | GTTTCGTTCACACTA |
| 1 | GMTACGGTCACACTG | chr3R:12377500-12379500 | 1034 | 1048 | + | 4.0571 | 3.01E-05 | 0.673 | GTTTCAGTCACAATG |
| 1 | GMTACGGTCACACTG | chr3R:24356938-24358938 | 1037 | 1051 | + | 4.0571 | 3.01E-05 | 0.673 | GTTAAGGTCACTCTA |
| 1 | GMTACGGTCACACTG | chr3L:2771325-2773325 | 1037 | 1051 | + | 4.0571 | 3.01E-05 | 0.673 | GGTAAGTTCCCACTG |
| 1 | GMTACGGTCACACTG | chr3L:7935120-7937120 | 1042 | 1056 | + | 4.0571 | 3.01E-05 | 0.673 | TTTACGGTCGCACTT |
| 1 | GMTACGGTCACACTG | chr3R:12766627-12768627 | 1046 | 1060 | + | 4.0571 | 3.01E-05 | 0.673 | ATGACGGTCACATTG |
| 1 | GMTACGGTCACACTG | chr2L:16287290-16289290 | 1048 | 1062 | + | 4.0571 | 3.01E-05 | 0.673 | TTTACCGTCGCACTG |
| 1 | GMTACGGTCACACTG | chr3L:1946475-1948475 | 1048 | 1062 | + | 4.0571 | 3.01E-05 | 0.673 | TGAACGGTCACACTT |
| 1 | GMTACGGTCACACTG | chrX:16445993-16447993 | 1053 | 1067 | + | 4.0571 | 3.01E-05 | 0.673 | CTTCTGGTCACACTG |
| 1 | GMTACGGTCACACTG | chr3R:12412759-12414759 | 1057 | 1071 | + | 4.0571 | 3.01E-05 | 0.673 | AGTCTGGTCACACTG |
| 1 | GMTACGGTCACACTG | chr3R:30305640-30307640 | 1073 | 1087 | + | 4.0571 | 3.01E-05 | 0.673 | CGGAAGGTCACACTG |
| 1 | GMTACGGTCACACTG | chr3L:7629508-7631508 | 1074 | 1088 | + | 4.0571 | 3.01E-05 | 0.673 | GTAACTGTCACATTG |
| 1 | GMTACGGTCACACTG | chr2L:9792107-9794107 | 1079 | 1093 | + | 4.0571 | 3.01E-05 | 0.673 | TTTACGGTCGTACTG |
| 1 | GMTACGGTCACACTG | chr3L:326992-328992 | 1083 | 1097 | + | 4.0571 | 3.01E-05 | 0.673 | AGCACGGTCACACTT |
| 1 | GMTACGGTCACACTG | chr3L:16380570-16382570 | 1087 | 1101 | + | 4.0571 | 3.01E-05 | 0.673 | GTTAAGGTCACTCTT |
| 1 | GMTACGGTCACACTG | chr2R:24968678-24970678 | 1087 | 1101 | + | 4.0571 | 3.01E-05 | 0.673 | CTTGCGGTCACACTA |
| 1 | GMTACGGTCACACTG | chr2R:16588184-16590184 | 1087 | 1101 | + | 4.0571 | 3.01E-05 | 0.673 | GGTGAGGTCACACTA |
| 1 | GMTACGGTCACACTG | chr3R:11948129-11950129 | 1096 | 1110 | + | 4.0571 | 3.01E-05 | 0.673 | CGTACGGCCACACTT |
| 1 | GMTACGGTCACACTG | chr2R:10818884-10820884 | 1097 | 1111 | + | 4.0571 | 3.01E-05 | 0.673 | AGTACGGTCACTCTA |
| 1 | GMTACGGTCACACTG | chr3R:13085224-13087224 | 1102 | 1116 | + | 4.0571 | 3.01E-05 | 0.673 | GTTGCGGTCACATTT |
| 1 | GMTACGGTCACACTG | chrX:8080605-8082605 | 1104 | 1118 | + | 4.0571 | 3.01E-05 | 0.673 | GGTACTGGTACACTG |

| | | | | | | | | | |
|---|---|---|---|---|---|---|---|---|---|
| 1 | GMTACGGTCACACTG | chr3R:11749029-11751029 | 1110 | 1124 | + | 4.0571 | 3.01E-05 | 0.673 | AGTATGGCCACACTG |
| 1 | GMTACGGTCACACTG | chr3L:15508445-15510445 | 1122 | 1136 | + | 4.0571 | 3.01E-05 | 0.673 | ATTGTGGTCACACTG |
| 1 | GMTACGGTCACACTG | chr3L:15508445-15510445 | 1122 | 1136 | + | 4.0571 | 3.01E-05 | 0.673 | ATTGTGGTCACACTG |
| 1 | GMTACGGTCACACTG | chr3R:21214079-21216079 | 1126 | 1140 | + | 4.0571 | 3.01E-05 | 0.673 | GTGATCGTCACACTG |
| 1 | GMTACGGTCACACTG | chr3R:24624562-24626562 | 1137 | 1151 | + | 4.0571 | 3.01E-05 | 0.673 | TGTACGGTCACACCA |
| 1 | GMTACGGTCACACTG | chr2L:11987270-11989270 | 1168 | 1182 | + | 4.0571 | 3.01E-05 | 0.673 | GTTTCGCTCAAACTG |
| 1 | GMTACGGTCACACTG | chr2R:16335090-16337090 | 1178 | 1192 | + | 4.0571 | 3.01E-05 | 0.673 | TTCAGGGTCACACTG |
| 1 | GMTACGGTCACACTG | chr3L:18871553-18873553 | 1200 | 1214 | + | 4.0571 | 3.01E-05 | 0.673 | AGTCTGGTCACACTG |
| 1 | GMTACGGTCACACTG | chrX:14233066-14235066 | 1202 | 1216 | + | 4.0571 | 3.01E-05 | 0.673 | AGTGTGGTCACACTG |
| 1 | GMTACGGTCACACTG | chr3R:12766468-12768468 | 1205 | 1219 | + | 4.0571 | 3.01E-05 | 0.673 | ATGACGGTCACATTG |
| 1 | GMTACGGTCACACTG | chrX:17859800-17861800 | 1211 | 1225 | + | 4.0571 | 3.01E-05 | 0.673 | CGTCTGGTCACACTG |
| 1 | GMTACGGTCACACTG | chr3R:14480713-14482713 | 1213 | 1227 | + | 4.0571 | 3.01E-05 | 0.673 | GTTGCGCTCAAACTG |
| 1 | GMTACGGTCACACTG | chr2L:6906844-6908844 | 1217 | 1231 | + | 4.0571 | 3.01E-05 | 0.673 | TGTACGGTCCCATTG |
| 1 | GMTACGGTCACACTG | chr2R:24968547-24970547 | 1218 | 1232 | + | 4.0571 | 3.01E-05 | 0.673 | CTTGCGGTCACACTA |
| 1 | GMTACGGTCACACTG | chr2R:9890689-9892689 | 1259 | 1273 | + | 4.0571 | 3.01E-05 | 0.673 | AGCACAGTCACACTG |
| 1 | GMTACGGTCACACTG | chr3R:13029659-13031659 | 1260 | 1274 | + | 4.0571 | 3.01E-05 | 0.673 | GTCAGGGTCACACTT |
| 1 | GMTACGGTCACACTG | chr3L:8067290-8069290 | 1274 | 1288 | + | 4.0571 | 3.01E-05 | 0.673 | GTTATGGACACATTG |
| 1 | GMTACGGTCACACTG | chr3L:9863349-9865349 | 1281 | 1295 | + | 4.0571 | 3.01E-05 | 0.673 | AGCACGGTCACACTT |
| 1 | GMTACGGTCACACTG | chr3L:3902184-3904184 | 1286 | 1300 | + | 4.0571 | 3.01E-05 | 0.673 | CGCACGGTCACACTA |
| 1 | GMTACGGTCACACTG | chr3R:10871381-10873381 | 1290 | 1304 | + | 4.0571 | 3.01E-05 | 0.673 | TTTTTGGTCACACTG |
| 1 | GMTACGGTCACACTG | chrX:16447302-16449302 | 1293 | 1307 | + | 4.0571 | 3.01E-05 | 0.673 | GGTGTGGTGACACTG |
| 1 | GMTACGGTCACACTG | chr3R:3696640-3698640 | 1293 | 1307 | + | 4.0571 | 3.01E-05 | 0.673 | GTAACTGTAACACTG |
| 1 | GMTACGGTCACACTG | chr2L:6648114-6650114 | 1293 | 1307 | + | 4.0571 | 3.01E-05 | 0.673 | AGTACGGTCACACGA |
| 1 | GMTACGGTCACACTG | chr3L:17906812-17908812 | 1308 | 1322 | + | 4.0571 | 3.01E-05 | 0.673 | TGTACGGCTACACTG |
| 1 | GMTACGGTCACACTG | chr2R:9112951-9114951 | 1316 | 1330 | + | 4.0571 | 3.01E-05 | 0.673 | AGTATGGTCACTCTG |
| 1 | GMTACGGTCACACTG | chrX:15319037-15321037 | 1323 | 1337 | + | 4.0571 | 3.01E-05 | 0.673 | AGTTTGGTCACACTG |
| 1 | GMTACGGTCACACTG | chr3L:3459977-3461977 | 1339 | 1353 | + | 4.0571 | 3.01E-05 | 0.673 | AGTCCGGTCACACTT |
| 1 | GMTACGGTCACACTG | chrX:18782494-18784494 | 1345 | 1359 | + | 4.0571 | 3.01E-05 | 0.673 | GGTCTGGTAACACTG |
| 1 | GMTACGGTCACACTG | chr3L:2168108-2170108 | 1347 | 1361 | + | 4.0571 | 3.01E-05 | 0.673 | GGTATGGTCCCATTG |
| 1 | GMTACGGTCACACTG | chr2R:7784067-7786067 | 1360 | 1374 | + | 4.0571 | 3.01E-05 | 0.673 | GTTACGCGCACACTT |
| 1 | GMTACGGTCACACTG | chr2L:8009652-8011652 | 1361 | 1375 | + | 4.0571 | 3.01E-05 | 0.673 | GTCAAGGTCACAATG |
| 1 | GMTACGGTCACACTG | chr2L:157664-159664 | 1362 | 1376 | + | 4.0571 | 3.01E-05 | 0.673 | CTAACGGTCACATTG |
| 1 | GMTACGGTCACACTG | chr3R:12412446-12414446 | 1370 | 1384 | + | 4.0571 | 3.01E-05 | 0.673 | AGTCTGGTCACACTG |

| | | | | | | | | | |
|---|---|---|---|---|---|---|---|---|---|
| 1 | GMTACGGTCACACTG | chr3R:23932530-23934530 | 1377 | 1391 | + | 4.0571 | 3.01E-05 | 0.673 | GGGATGGTCACACTA |
| 1 | GMTACGGTCACACTG | chr3R:5793317-5795317 | 1381 | 1395 | + | 4.0571 | 3.01E-05 | 0.673 | AGCACGGTCATACTG |
| 1 | GMTACGGTCACACTG | chr2L:11987039-11989039 | 1399 | 1413 | + | 4.0571 | 3.01E-05 | 0.673 | GTTTCGCTCAAACTG |
| 1 | GMTACGGTCACACTG | chr2L:1157262-1159262 | 1409 | 1423 | + | 4.0571 | 3.01E-05 | 0.673 | CGCATGGTCACACTG |
| 1 | GMTACGGTCACACTG | chr3R:5735748-5737748 | 1424 | 1438 | + | 4.0571 | 3.01E-05 | 0.673 | GTTACGATCACATCG |
| 1 | GMTACGGTCACACTG | chr2L:8302429-8304429 | 1439 | 1453 | + | 4.0571 | 3.01E-05 | 0.673 | GTTGTGGTCACAATG |
| 1 | GMTACGGTCACACTG | chr2R:5764180-5766180 | 1439 | 1453 | + | 4.0571 | 3.01E-05 | 0.673 | GGTACGTTCAAGCTG |
| 1 | GMTACGGTCACACTG | chr2R:11755817-11757817 | 1452 | 1466 | + | 4.0571 | 3.01E-05 | 0.673 | GTTATGGGCACAGTG |
| 1 | GMTACGGTCACACTG | chr3L:15508061-15510061 | 1506 | 1520 | + | 4.0571 | 3.01E-05 | 0.673 | ATTGTGGTCACACTG |
| 1 | GMTACGGTCACACTG | chrX:18513867-18515867 | 1532 | 1546 | + | 4.0571 | 3.01E-05 | 0.673 | CGAACGGTCACATTG |
| 1 | GMTACGGTCACACTG | chr2R:21388405-21390405 | 1534 | 1548 | + | 4.0571 | 3.01E-05 | 0.673 | GGCACAGTCACACTT |
| 1 | GMTACGGTCACACTG | chr3L:18462163-18464163 | 1547 | 1561 | + | 4.0571 | 3.01E-05 | 0.673 | GTCACGCTCACACTT |
| 1 | GMTACGGTCACACTG | chr3R:20559280-20561280 | 1553 | 1567 | + | 4.0571 | 3.01E-05 | 0.673 | GTTCTGGTCACACTT |
| 1 | GMTACGGTCACACTG | chr2L:11129737-11131737 | 1563 | 1577 | + | 4.0571 | 3.01E-05 | 0.673 | CGCACCGTCACACTG |
| 1 | GMTACGGTCACACTG | chrX:15696494-15698494 | 1568 | 1582 | + | 4.0571 | 3.01E-05 | 0.673 | GTCAAAGTCACACTG |
| 1 | GMTACGGTCACACTG | chr2R:11851617-11853617 | 1575 | 1589 | + | 4.0571 | 3.01E-05 | 0.673 | GGTGTGGTAACACTG |
| 1 | GMTACGGTCACACTG | chr3L:16590050-16592050 | 1577 | 1591 | + | 4.0571 | 3.01E-05 | 0.673 | GTTACGGACAAAGTG |
| 1 | GMTACGGTCACACTG | chrX:15065432-15067432 | 1578 | 1592 | + | 4.0571 | 3.01E-05 | 0.673 | TGCAAGGTCACACTG |
| 1 | GMTACGGTCACACTG | chr3R:20208791-20210791 | 1582 | 1596 | + | 4.0571 | 3.01E-05 | 0.673 | GTTACGGTGACCTTG |
| 1 | GMTACGGTCACACTG | chr2R:24560607-24562607 | 1590 | 1604 | + | 4.0571 | 3.01E-05 | 0.673 | GTTTTGTTCACACTG |
| 1 | GMTACGGTCACACTG | chr3R:28805702-28807702 | 1600 | 1614 | + | 4.0571 | 3.01E-05 | 0.673 | GTTACCGTCCCACTT |
| 1 | GMTACGGTCACACTG | chr2L:16286730-16288730 | 1608 | 1622 | + | 4.0571 | 3.01E-05 | 0.673 | TTTACCGTCGCACTG |
| 1 | GMTACGGTCACACTG | chr3R:13029307-13031307 | 1612 | 1626 | + | 4.0571 | 3.01E-05 | 0.673 | GTCAGGGTCACACTT |
| 1 | GMTACGGTCACACTG | chr3L:550010-552010 | 1635 | 1649 | + | 4.0571 | 3.01E-05 | 0.673 | GGGAAGGTCAAACTG |
| 1 | GMTACGGTCACACTG | chr3L:13025437-13027437 | 1646 | 1660 | + | 4.0571 | 3.01E-05 | 0.673 | GTTGCGTTGACACTG |
| 1 | GMTACGGTCACACTG | chrX:16695131-16697131 | 1649 | 1663 | + | 4.0571 | 3.01E-05 | 0.673 | GTGAAGGTCACATTG |
| 1 | GMTACGGTCACACTG | chr3R:22341251-22343251 | 1655 | 1669 | + | 4.0571 | 3.01E-05 | 0.673 | GTGACTGTCAAACTG |
| 1 | GMTACGGTCACACTG | chr2L:11129645-11131645 | 1655 | 1669 | + | 4.0571 | 3.01E-05 | 0.673 | CGCACCGTCACACTG |
| 1 | GMTACGGTCACACTG | chr2R:12591917-12593917 | 1657 | 1671 | + | 4.0571 | 3.01E-05 | 0.673 | CGCACAGTCACACTG |
| 1 | GMTACGGTCACACTG | chr2R:19424113-19426113 | 1676 | 1690 | + | 4.0571 | 3.01E-05 | 0.673 | CTTCCAGTCACACTG |
| 1 | GMTACGGTCACACTG | chr2R:21535795-21537795 | 1725 | 1739 | + | 4.0571 | 3.01E-05 | 0.673 | GTTCTGGTCACATTG |
| 1 | GMTACGGTCACACTG | chr3R:29998795-30000795 | 1729 | 1743 | + | 4.0571 | 3.01E-05 | 0.673 | TGGACGGTTACACTG |
| 1 | GMTACGGTCACACTG | chr3R:5615308-5617308 | 1758 | 1772 | + | 4.0571 | 3.01E-05 | 0.673 | GGTCAGGTCAGACTG |

| | | | | | | | | | |
|---|---|---|---|---|---|---|---|---|---|
| 1 | GMTACGGTCACACTG | chr2L:14333158-14335158 | 1762 | 1776 | + | 4.0571 | 3.01E-05 | 0.673 | GGTACGGTCACCATA |
| 1 | GMTACGGTCACACTG | chr3L:4258430-4260430 | 1800 | 1814 | + | 4.0571 | 3.01E-05 | 0.673 | GGTTCGGTCAAACTA |
| 1 | GMTACGGTCACACTG | chrY:2272422-2274422 | 1806 | 1820 | + | 4.0571 | 3.01E-05 | 0.673 | ATTACGGTTACGCTG |
| 1 | GMTACGGTCACACTG | chr3R:13050453-13052453 | 1812 | 1826 | + | 4.0571 | 3.01E-05 | 0.673 | TGAACGGTCACACTA |
| 1 | GMTACGGTCACACTG | chr3R:5735355-5737355 | 1817 | 1831 | + | 4.0571 | 3.01E-05 | 0.673 | GTTACGATCACATCG |
| 1 | GMTACGGTCACACTG | chrX:19560837-19562837 | 1826 | 1840 | + | 4.0571 | 3.01E-05 | 0.673 | GGTACCCTCGCACTG |
| 1 | GMTACGGTCACACTG | chr3R:13854454-13856454 | 1832 | 1846 | + | 4.0571 | 3.01E-05 | 0.673 | GGTCTGGTCAGACTG |
| 1 | GMTACGGTCACACTG | chrX:15768664-15770664 | 1843 | 1857 | + | 4.0571 | 3.01E-05 | 0.673 | GTTACGGACACAGTT |
| 1 | GMTACGGTCACACTG | chr3R:13084478-13086478 | 1848 | 1862 | + | 4.0571 | 3.01E-05 | 0.673 | GTTGCGGTCACATTT |
| 1 | GMTACGGTCACACTG | chrY:3084676-3086676 | 1851 | 1865 | + | 4.0571 | 3.01E-05 | 0.673 | GTAACTGTAACACTG |
| 1 | GMTACGGTCACACTG | chr3L:21515306-21517306 | 1871 | 1885 | + | 4.0571 | 3.01E-05 | 0.673 | TTTTCGGTCACACTT |
| 1 | GMTACGGTCACACTG | chrX:15318479-15320479 | 1881 | 1895 | + | 4.0571 | 3.01E-05 | 0.673 | AGTTTGGTCACACTG |
| 1 | GMTACGGTCACACTG | chr2L:8301985-8303985 | 1883 | 1897 | + | 4.0571 | 3.01E-05 | 0.673 | GTTGTGGTCACAATG |
| 1 | GMTACGGTCACACTG | chr3L:14001127-14003127 | 1885 | 1899 | + | 4.0571 | 3.01E-05 | 0.673 | AGAACGGTCACACTA |
| 1 | GMTACGGTCACACTG | chr2R:19423903-19425903 | 1886 | 1900 | + | 4.0571 | 3.01E-05 | 0.673 | CTTCCAGTCACACTG |
| 1 | GMTACGGTCACACTG | chr2L:6359403-6361403 | 1892 | 1906 | + | 4.0571 | 3.01E-05 | 0.673 | GGTCCTGTCAGACTG |
| 1 | GMTACGGTCACACTG | chr2L:9962949-9964949 | 1893 | 1907 | + | 4.0571 | 3.01E-05 | 0.673 | GTTACTGTCAAACAG |
| 1 | GMTACGGTCACACTG | chr2L:1517148-1519148 | 1967 | 1981 | + | 4.0571 | 3.01E-05 | 0.673 | GTTTCGGTCACAATT |
| 1 | GMTACGGTCACACTG | chr3L:22406863-22408863 | 37 | 51 | - | 3.9143 | 3.06E-05 | 0.673 | GCAGCGGACACACGG |
| 1 | GMTACGGTCACACTG | chr2R:17771552-17773552 | 151 | 165 | - | 3.9143 | 3.06E-05 | 0.673 | GCAGCGGTCCCACCG |
| 1 | GMTACGGTCACACTG | chrX:125714-127714 | 206 | 220 | - | 3.9143 | 3.06E-05 | 0.673 | GCTGCGGCCACTCAG |
| 1 | GMTACGGTCACACTG | chr3L:9725895-9727895 | 252 | 266 | - | 3.9143 | 3.06E-05 | 0.673 | GCTGCGGACTCACCG |
| 1 | GMTACGGTCACACTG | chr3L:22406547-22408547 | 353 | 367 | - | 3.9143 | 3.06E-05 | 0.673 | GCAGCGGACACACGG |
| 1 | GMTACGGTCACACTG | chrX:5716778-5718778 | 379 | 393 | - | 3.9143 | 3.06E-05 | 0.673 | GCTTCGGTCGCTCAG |
| 1 | GMTACGGTCACACTG | chr3L:17627057-17629057 | 389 | 403 | - | 3.9143 | 3.06E-05 | 0.673 | GCGCCGGCCACGCTG |
| 1 | GMTACGGTCACACTG | chr3R:21063471-21065471 | 440 | 454 | - | 3.9143 | 3.06E-05 | 0.673 | GCAGCGGTCGCACAG |
| 1 | GMTACGGTCACACTG | chr2R:7386187-7388187 | 711 | 725 | - | 3.9143 | 3.06E-05 | 0.673 | GCTTCGGCCCCACAG |
| 1 | GMTACGGTCACACTG | chr3L:20132268-20134268 | 780 | 794 | - | 3.9143 | 3.06E-05 | 0.673 | GCGGCGGTCCCGCTG |
| 1 | GMTACGGTCACACTG | chr2R:7386115-7388115 | 783 | 797 | - | 3.9143 | 3.06E-05 | 0.673 | GCTTCGGCCCCACAG |
| 1 | GMTACGGTCACACTG | chr3R:19225651-19227651 | 923 | 937 | - | 3.9143 | 3.06E-05 | 0.673 | GCTCCGGACACGCAG |
| 1 | GMTACGGTCACACTG | chr3L:16711126-16713126 | 971 | 985 | - | 3.9143 | 3.06E-05 | 0.673 | GCCGCGGGCACCCTG |
| 1 | GMTACGGTCACACTG | chrX:17790010-17792010 | 1299 | 1313 | - | 3.9143 | 3.06E-05 | 0.673 | GCGGCGGCCACTCTG |
| 1 | GMTACGGTCACACTG | chr3L:16710677-16712677 | 1420 | 1434 | - | 3.9143 | 3.06E-05 | 0.673 | GCCGCGGGCACCCTG |

| | | | | | | | | | |
|---|---|---|---|---|---|---|---|---|---|
| 1 | GMTACGGTCACACTG | chr3L:16710677-16712677 | 1420 | 1434 | - | 3.9143 | 3.06E-05 | 0.673 | GCCGCGGGCACCCTG |
| 1 | GMTACGGTCACACTG | chr2R:13410739-13412739 | 1586 | 1600 | - | 3.9143 | 3.06E-05 | 0.673 | GCGTCGGCCACGCTG |
| 1 | GMTACGGTCACACTG | chr2R:13406011-13408011 | 1587 | 1601 | - | 3.9143 | 3.06E-05 | 0.673 | GCGTCGGCCACGCTG |
| 1 | GMTACGGTCACACTG | chr2L:5052094-5054094 | 1624 | 1638 | - | 3.9143 | 3.06E-05 | 0.673 | GCTGCGGACTCGCTG |
| 1 | GMTACGGTCACACTG | chr2R:23555672-23557672 | 1764 | 1778 | - | 3.9143 | 3.06E-05 | 0.673 | GCTACGGACTCCCGG |
| 1 | GMTACGGTCACACTG | chr2L:13373019-13375019 | 1845 | 1859 | - | 3.9143 | 3.06E-05 | 0.673 | GCAGCGGTCGCACAG |
| 1 | GMTACGGTCACACTG | chr2L:5051843-5053843 | 1875 | 1889 | - | 3.9143 | 3.06E-05 | 0.673 | GCTGCGGACTCGCTG |
| 1 | GMTACGGTCACACTG | chr2L:13372890-13374890 | 1974 | 1988 | - | 3.9143 | 3.06E-05 | 0.673 | GCAGCGGTCGCACAG |
| 1 | GMTACGGTCACACTG | chr2R:12335733-12337733 | 80 | 94 | + | 3.9143 | 3.06E-05 | 0.673 | GCCACGGTCGCCCAG |
| 1 | GMTACGGTCACACTG | chr3R:13544400-13546400 | 164 | 178 | + | 3.9143 | 3.06E-05 | 0.673 | GCAACGGACCCTCTG |
| 1 | GMTACGGTCACACTG | chr2R:12335418-12337418 | 395 | 409 | + | 3.9143 | 3.06E-05 | 0.673 | GCCACGGTCGCCCAG |
| 1 | GMTACGGTCACACTG | chr3L:21950548-21952548 | 477 | 491 | + | 3.9143 | 3.06E-05 | 0.673 | GCGACGGTCGCTCAG |
| 1 | GMTACGGTCACACTG | chr2R:24158384-24160384 | 497 | 511 | + | 3.9143 | 3.06E-05 | 0.673 | GCTCCGGACACGCAG |
| 1 | GMTACGGTCACACTG | chr2R:15694386-15696386 | 525 | 539 | + | 3.9143 | 3.06E-05 | 0.673 | GCGACGGCCTCACCG |
| 1 | GMTACGGTCACACTG | chr2R:24368689-24370689 | 538 | 552 | + | 3.9143 | 3.06E-05 | 0.673 | GCACCGGGCACACAG |
| 1 | GMTACGGTCACACTG | chr2R:24158312-24160312 | 569 | 583 | + | 3.9143 | 3.06E-05 | 0.673 | GCTCCGGACACGCAG |
| 1 | GMTACGGTCACACTG | chr2R:24158312-24160312 | 569 | 583 | + | 3.9143 | 3.06E-05 | 0.673 | GCTCCGGACACGCAG |
| 1 | GMTACGGTCACACTG | chr3R:14258991-14260991 | 597 | 611 | + | 3.9143 | 3.06E-05 | 0.673 | GCAACGGGCACCCCG |
| 1 | GMTACGGTCACACTG | chr3R:29488468-29490468 | 624 | 638 | + | 3.9143 | 3.06E-05 | 0.673 | GCTGCGGACACTCGG |
| 1 | GMTACGGTCACACTG | chrX:14321226-14323226 | 812 | 826 | + | 3.9143 | 3.06E-05 | 0.673 | GCAGCGGACACACGG |
| 1 | GMTACGGTCACACTG | chr3R:29488265-29490265 | 827 | 841 | + | 3.9143 | 3.06E-05 | 0.673 | GCTGCGGACACTCGG |
| 1 | GMTACGGTCACACTG | chr2L:6493532-6495532 | 859 | 873 | + | 3.9143 | 3.06E-05 | 0.673 | GCGGCGGGCACACGG |
| 1 | GMTACGGTCACACTG | chr2R:20660585-20662585 | 944 | 958 | + | 3.9143 | 3.06E-05 | 0.673 | GCGACGGTCCCTCGG |
| 1 | GMTACGGTCACACTG | chr3R:19032538-19034538 | 1023 | 1037 | + | 3.9143 | 3.06E-05 | 0.673 | GCAACGGTCGCGCCG |
| 1 | GMTACGGTCACACTG | chr2R:7673450-7675450 | 1130 | 1144 | + | 3.9143 | 3.06E-05 | 0.673 | GCAACGGCCACTCGG |
| 1 | GMTACGGTCACACTG | chr2R:7673400-7675400 | 1180 | 1194 | + | 3.9143 | 3.06E-05 | 0.673 | GCAACGGCCACTCGG |
| 1 | GMTACGGTCACACTG | chr2R:16142147-16144147 | 1222 | 1236 | + | 3.9143 | 3.06E-05 | 0.673 | GCTGCGGACCCACCG |
| 1 | GMTACGGTCACACTG | chr2R:7011254-7013254 | 1470 | 1484 | + | 3.9143 | 3.06E-05 | 0.673 | GCAGCGGGCACGCTG |
| 1 | GMTACGGTCACACTG | chr3L:11108690-11110690 | 1477 | 1491 | + | 3.9143 | 3.06E-05 | 0.673 | GCTGCGGTCCCGCGG |
| 1 | GMTACGGTCACACTG | chr2R:12334316-12336316 | 1497 | 1511 | + | 3.9143 | 3.06E-05 | 0.673 | GCCACGGTCGCCCAG |
| 1 | GMTACGGTCACACTG | chr3L:11108505-11110505 | 1662 | 1676 | + | 3.9143 | 3.06E-05 | 0.673 | GCTGCGGTCCCGCGG |
| 1 | GMTACGGTCACACTG | chr2R:12334066-12336066 | 1747 | 1761 | + | 3.9143 | 3.06E-05 | 0.673 | GCCACGGTCGCCCAG |
| 1 | GMTACGGTCACACTG | chr3L:11628505-11630505 | 1750 | 1764 | + | 3.9143 | 3.06E-05 | 0.673 | GCCGCGGCCCCACTG |

| | | | | | | | | | |
|---|---|---|---|---|---|---|---|---|---|
| 1 | GMTACGGTCACACTG | chr2R:15693092-15695092 | 1819 | 1833 | + | 3.9143 | 3.06E-05 | 0.673 | GCGACGGCCTCACCG |
| 1 | GMTACGGTCACACTG | chrX:16720411-16722411 | 1847 | 1861 | + | 3.9143 | 3.06E-05 | 0.673 | GCGCCGGCCACTCTG |
| 1 | GMTACGGTCACACTG | chr2R:7010739-7012739 | 1985 | 1999 | + | 3.9143 | 3.06E-05 | 0.673 | GCAGCGGGCACGCTG |
| 1 | GMTACGGTCACACTG | chr3R:20251770-20253770 | 69 | 83 | - | 3.7786 | 3.38E-05 | 0.71 | GGTAAGGTAACACTT |
| 1 | GMTACGGTCACACTG | chr2L:9332567-9334567 | 129 | 143 | - | 3.7786 | 3.38E-05 | 0.71 | GGTATGTTCAAACTG |
| 1 | GMTACGGTCACACTG | chr3R:20841140-20843140 | 145 | 159 | - | 3.7786 | 3.38E-05 | 0.71 | GGTAAATTCACACTG |
| 1 | GMTACGGTCACACTG | chr3R:18848224-18850224 | 165 | 179 | - | 3.7786 | 3.38E-05 | 0.71 | ATTACGATCACAATG |
| 1 | GMTACGGTCACACTG | chr3L:13299152-13301152 | 168 | 182 | - | 3.7786 | 3.38E-05 | 0.71 | GTTACGTTTACACTA |
| 1 | GMTACGGTCACACTG | chr3L:17796594-17798594 | 192 | 206 | - | 3.7786 | 3.38E-05 | 0.71 | CTTAAGGTGACACTG |
| 1 | GMTACGGTCACACTG | chr2L:2284163-2286163 | 198 | 212 | - | 3.7786 | 3.38E-05 | 0.71 | TGTACGGTCATAATG |
| 1 | GMTACGGTCACACTG | chr2R:22940877-22942877 | 221 | 235 | - | 3.7786 | 3.38E-05 | 0.71 | GTTACCGTTACACTT |
| 1 | GMTACGGTCACACTG | chr2L:10526572-10528572 | 238 | 252 | - | 3.7786 | 3.38E-05 | 0.71 | GTTACAGTCAAACTT |
| 1 | GMTACGGTCACACTG | chr2L:6366882-6368882 | 259 | 273 | - | 3.7786 | 3.38E-05 | 0.71 | GTTACAGTTACAGTG |
| 1 | GMTACGGTCACACTG | chr2R:25076048-25078048 | 275 | 289 | - | 3.7786 | 3.38E-05 | 0.71 | GGTACTGTCAAAATG |
| 1 | GMTACGGTCACACTG | chr2L:2284003-2286003 | 358 | 372 | - | 3.7786 | 3.38E-05 | 0.71 | TGTACGGTCATAATG |
| 1 | GMTACGGTCACACTG | chr3L:20960386-20962386 | 387 | 401 | - | 3.7786 | 3.38E-05 | 0.71 | CGTACGGTGACAGTG |
| 1 | GMTACGGTCACACTG | chr3R:23698027-23700027 | 401 | 415 | - | 3.7786 | 3.38E-05 | 0.71 | GTTAACTTCACACTG |
| 1 | GMTACGGTCACACTG | chrX:19815072-19817072 | 405 | 419 | - | 3.7786 | 3.38E-05 | 0.71 | GTTACCGTAAAACTG |
| 1 | GMTACGGTCACACTG | chr3L:20961555-20963555 | 427 | 441 | - | 3.7786 | 3.38E-05 | 0.71 | CGTACGGTGACAGTG |
| 1 | GMTACGGTCACACTG | chr2L:1554129-1556129 | 433 | 447 | - | 3.7786 | 3.38E-05 | 0.71 | GGTAACGTAACACTG |
| 1 | GMTACGGTCACACTG | chrX:15991468-15993468 | 481 | 495 | - | 3.7786 | 3.38E-05 | 0.71 | GTTATGGTCATATTG |
| 1 | GMTACGGTCACACTG | chr3L:9600523-9602523 | 495 | 509 | - | 3.7786 | 3.38E-05 | 0.71 | TTTACGGTGACAGTG |
| 1 | GMTACGGTCACACTG | chr3L:13296004-13298004 | 500 | 514 | - | 3.7786 | 3.38E-05 | 0.71 | GTTATGTTCAAACTG |
| 1 | GMTACGGTCACACTG | chr3R:5299145-5301145 | 539 | 553 | - | 3.7786 | 3.38E-05 | 0.71 | GTTATGGTCACAATT |
| 1 | GMTACGGTCACACTG | chr3R:21621571-21623571 | 558 | 572 | - | 3.7786 | 3.38E-05 | 0.71 | TTTACTTTCACACTG |
| 1 | GMTACGGTCACACTG | chr2R:25075765-25077765 | 558 | 572 | - | 3.7786 | 3.38E-05 | 0.71 | GGTACTGTCAAAATG |
| 1 | GMTACGGTCACACTG | chr3R:7073859-7075859 | 559 | 573 | - | 3.7786 | 3.38E-05 | 0.71 | AGTAAGGTCACACTT |
| 1 | GMTACGGTCACACTG | chrX:8456632-8458632 | 620 | 634 | - | 3.7786 | 3.38E-05 | 0.71 | GTTATGTTTACACTG |
| 1 | GMTACGGTCACACTG | chr3L:13295860-13297860 | 644 | 658 | - | 3.7786 | 3.38E-05 | 0.71 | GTTATGTTCAAACTG |
| 1 | GMTACGGTCACACTG | chr3L:9600325-9602325 | 693 | 707 | - | 3.7786 | 3.38E-05 | 0.71 | TTTACGGTGACAGTG |
| 1 | GMTACGGTCACACTG | chrX:16390698-16392698 | 724 | 738 | - | 3.7786 | 3.38E-05 | 0.71 | TTTACAGTCACACTA |
| 1 | GMTACGGTCACACTG | chr2R:8420033-8422033 | 732 | 746 | - | 3.7786 | 3.38E-05 | 0.71 | ATTAAGGTCACACTA |
| 1 | GMTACGGTCACACTG | chr3R:5469926-5471926 | 737 | 751 | - | 3.7786 | 3.38E-05 | 0.71 | TTTACGGTTAGACTG |

| | | | | | | | | | |
|---|---|---|---|---|---|---|---|---|---|
| 1 | GMTACGGTCACACTG | chr2R:9147689-9149689 | 788 | 802 | - | 3.7786 | 3.38E-05 | 0.71 | TTTATTGTCACACTG |
| 1 | GMTACGGTCACACTG | chr3R:20485679-20487679 | 836 | 850 | - | 3.7786 | 3.38E-05 | 0.71 | GTTACAGTCACAGTC |
| 1 | GMTACGGTCACACTG | chr2R:21480927-21482927 | 872 | 886 | - | 3.7786 | 3.38E-05 | 0.71 | GTTACTGTTACAATG |
| 1 | GMTACGGTCACACTG | chrX:14954477-14956477 | 876 | 890 | - | 3.7786 | 3.38E-05 | 0.71 | GTTACTTTCACATTG |
| 1 | GMTACGGTCACACTG | chrX:6673387-6675387 | 920 | 934 | - | 3.7786 | 3.38E-05 | 0.71 | CTTACGGTCACATTA |
| 1 | GMTACGGTCACACTG | chr2R:22940169-22942169 | 929 | 943 | - | 3.7786 | 3.38E-05 | 0.71 | GTTACCGTTACACTT |
| 1 | GMTACGGTCACACTG | chr3R:7073452-7075452 | 966 | 980 | - | 3.7786 | 3.38E-05 | 0.71 | AGTAAGGTCACACTT |
| 1 | GMTACGGTCACACTG | chrX:19207187-19209187 | 966 | 980 | - | 3.7786 | 3.38E-05 | 0.71 | TGTACGGTCATACTA |
| 1 | GMTACGGTCACACTG | chr3R:13365910-13367910 | 971 | 985 | - | 3.7786 | 3.38E-05 | 0.71 | TTTACGGTCATACTA |
| 1 | GMTACGGTCACACTG | chrX:15990974-15992974 | 975 | 989 | - | 3.7786 | 3.38E-05 | 0.71 | GTTATGGTCATATTG |
| 1 | GMTACGGTCACACTG | chrX:15990974-15992974 | 975 | 989 | - | 3.7786 | 3.38E-05 | 0.71 | GTTATGGTCATATTG |
| 1 | GMTACGGTCACACTG | chrX:16390438-16392438 | 984 | 998 | - | 3.7786 | 3.38E-05 | 0.71 | TTTACAGTCACACTA |
| 1 | GMTACGGTCACACTG | chr2R:11278725-11280725 | 998 | 1012 | - | 3.7786 | 3.38E-05 | 0.71 | GTTACGGTAACATTT |
| 1 | GMTACGGTCACACTG | chrX:2584980-2586980 | 1003 | 1017 | - | 3.7786 | 3.38E-05 | 0.71 | CTTAAGGTCACACTA |
| 1 | GMTACGGTCACACTG | chr3L:16010812-16012812 | 1021 | 1035 | - | 3.7786 | 3.38E-05 | 0.71 | CGTAACGTCACACTG |
| 1 | GMTACGGTCACACTG | chr3R:5469636-5471636 | 1027 | 1041 | - | 3.7786 | 3.38E-05 | 0.71 | TTTACGGTTAGACTG |
| 1 | GMTACGGTCACACTG | chr2R:11278683-11280683 | 1040 | 1054 | - | 3.7786 | 3.38E-05 | 0.71 | GTTACGGTAACATTT |
| 1 | GMTACGGTCACACTG | chr3R:20573906-20575906 | 1103 | 1117 | - | 3.7786 | 3.38E-05 | 0.71 | GTTAATATCACACTG |
| 1 | GMTACGGTCACACTG | chr3R:21621021-21623021 | 1108 | 1122 | - | 3.7786 | 3.38E-05 | 0.71 | TTTACTTTCACACTG |
| 1 | GMTACGGTCACACTG | chr3L:10793964-10795964 | 1170 | 1184 | - | 3.7786 | 3.38E-05 | 0.71 | GTTATTGTTACACTG |
| 1 | GMTACGGTCACACTG | chr2R:20505395-20507395 | 1192 | 1206 | - | 3.7786 | 3.38E-05 | 0.71 | GGTACGTTTACAATG |
| 1 | GMTACGGTCACACTG | chr2R:21795395-21797395 | 1246 | 1260 | - | 3.7786 | 3.38E-05 | 0.71 | GTTATTGTCATACTG |
| 1 | GMTACGGTCACACTG | chrX:14954100-14956100 | 1253 | 1267 | - | 3.7786 | 3.38E-05 | 0.71 | GTTACTTTCACATTG |
| 1 | GMTACGGTCACACTG | chr2R:18815452-18817452 | 1263 | 1277 | - | 3.7786 | 3.38E-05 | 0.71 | TTTACAGTCACATTG |
| 1 | GMTACGGTCACACTG | chrX:2584699-2586699 | 1284 | 1298 | - | 3.7786 | 3.38E-05 | 0.71 | CTTAAGGTCACACTA |
| 1 | GMTACGGTCACACTG | chrX:20109575-20111575 | 1293 | 1307 | - | 3.7786 | 3.38E-05 | 0.71 | ATTATGGTCACACTA |
| 1 | GMTACGGTCACACTG | chr3L:13298023-13300023 | 1297 | 1311 | - | 3.7786 | 3.38E-05 | 0.71 | GTTACGTTTACACTA |
| 1 | GMTACGGTCACACTG | chr2R:25075005-25077005 | 1318 | 1332 | - | 3.7786 | 3.38E-05 | 0.71 | GGTACTGTCAAAATG |
| 1 | GMTACGGTCACACTG | chrX:6672973-6674973 | 1334 | 1348 | - | 3.7786 | 3.38E-05 | 0.71 | CTTACGGTCACATTA |
| 1 | GMTACGGTCACACTG | chr3L:4291482-4293482 | 1346 | 1360 | - | 3.7786 | 3.38E-05 | 0.71 | GTTATGTTAACACTG |
| 1 | GMTACGGTCACACTG | chr2R:20505195-20507195 | 1392 | 1406 | - | 3.7786 | 3.38E-05 | 0.71 | GGTACGTTTACAATG |
| 1 | GMTACGGTCACACTG | chr3L:7694654-7696654 | 1402 | 1416 | - | 3.7786 | 3.38E-05 | 0.71 | ATTACGTTAACACTG |
| 1 | GMTACGGTCACACTG | chr2L:8448576-8450576 | 1488 | 1502 | - | 3.7786 | 3.38E-05 | 0.71 | GTTACGATCAAACTC |

| | | | | | | | | | |
|---|---|---|---|---|---|---|---|---|---|
| 1 | GMTACGGTCACACTG | chr3L:15843062-15845062 | 1525 | 1539 | - | 3.7786 | 3.38E-05 | 0.71 | GTTAAGATCACAATG |
| 1 | GMTACGGTCACACTG | chr3L:4291246-4293246 | 1582 | 1596 | - | 3.7786 | 3.38E-05 | 0.71 | GTTATGTTAACACTG |
| 1 | GMTACGGTCACACTG | chr3L:20960386-20962386 | 1596 | 1610 | - | 3.7786 | 3.38E-05 | 0.71 | CGTACGGTGACAGTG |
| 1 | GMTACGGTCACACTG | chr2R:21480176-21482176 | 1623 | 1637 | - | 3.7786 | 3.38E-05 | 0.71 | GTTACTGTTACAATG |
| 1 | GMTACGGTCACACTG | chr3R:15329738-15331738 | 1663 | 1677 | - | 3.7786 | 3.38E-05 | 0.71 | TTTATGGTCACAGTG |
| 1 | GMTACGGTCACACTG | chrX:7065052-7067052 | 1670 | 1684 | - | 3.7786 | 3.38E-05 | 0.71 | GTTACGTTCAAACTA |
| 1 | GMTACGGTCACACTG | chrX:3347400-3349400 | 1723 | 1737 | - | 3.7786 | 3.38E-05 | 0.71 | ATTAAGGTCACACTT |
| 1 | GMTACGGTCACACTG | chrX:8723674-8725674 | 1750 | 1764 | - | 3.7786 | 3.38E-05 | 0.71 | GTTAGTCTCACACTG |
| 1 | GMTACGGTCACACTG | chr2R:8126575-8128575 | 1875 | 1889 | - | 3.7786 | 3.38E-05 | 0.71 | GTTACGTTCACAATC |
| 1 | GMTACGGTCACACTG | chr3L:19254340-19256340 | 1882 | 1896 | - | 3.7786 | 3.38E-05 | 0.71 | TGTAGGGTCACAGTG |
| 1 | GMTACGGTCACACTG | chr2L:15901410-15903410 | 1960 | 1974 | - | 3.7786 | 3.38E-05 | 0.71 | TTTAAGGTTACACTG |
| 1 | GMTACGGTCACACTG | chr3R:25686370-25688370 | 60 | 74 | + | 3.7786 | 3.38E-05 | 0.71 | GGTATGGTCATAATG |
| 1 | GMTACGGTCACACTG | chr3L:1809588-1811588 | 67 | 81 | + | 3.7786 | 3.38E-05 | 0.71 | GGTACAGTCACATTC |
| 1 | GMTACGGTCACACTG | chrX:13358248-13360248 | 100 | 114 | + | 3.7786 | 3.38E-05 | 0.71 | AGTAAGGTCAAACTG |
| 1 | GMTACGGTCACACTG | chr2R:21696152-21698152 | 225 | 239 | + | 3.7786 | 3.38E-05 | 0.71 | GTTAAGGTCACAGTA |
| 1 | GMTACGGTCACACTG | chr3R:12004473-12006473 | 399 | 413 | + | 3.7786 | 3.38E-05 | 0.71 | TTTATGGTCAGACTG |
| 1 | GMTACGGTCACACTG | chr2R:23140999-23142999 | 414 | 428 | + | 3.7786 | 3.38E-05 | 0.71 | GTTACAGTCACAGTC |
| 1 | GMTACGGTCACACTG | chrX:7402174-7404174 | 414 | 428 | + | 3.7786 | 3.38E-05 | 0.71 | ATTAGGGTCATACTG |
| 1 | GMTACGGTCACACTG | chr3L:20973700-20975700 | 468 | 482 | + | 3.7786 | 3.38E-05 | 0.71 | GTTATGGTCATAGTG |
| 1 | GMTACGGTCACACTG | chr3L:19804181-19806181 | 474 | 488 | + | 3.7786 | 3.38E-05 | 0.71 | CTTAAAGTCACACTG |
| 1 | GMTACGGTCACACTG | chr3R:4983232-4985232 | 479 | 493 | + | 3.7786 | 3.38E-05 | 0.71 | GTTATGTTCACAGTG |
| 1 | GMTACGGTCACACTG | chr2R:13546577-13548577 | 483 | 497 | + | 3.7786 | 3.38E-05 | 0.71 | CGTATGGTCACACTA |
| 1 | GMTACGGTCACACTG | chrX:15795477-15797477 | 483 | 497 | + | 3.7786 | 3.38E-05 | 0.71 | GTTACGGTAAAATTG |
| 1 | GMTACGGTCACACTG | chr2R:10874748-10876748 | 495 | 509 | + | 3.7786 | 3.38E-05 | 0.71 | GTTACTCTCACATTG |
| 1 | GMTACGGTCACACTG | chr2R:14872953-14874953 | 526 | 540 | + | 3.7786 | 3.38E-05 | 0.71 | GGTACAGTAAAACTG |
| 1 | GMTACGGTCACACTG | chr3R:6656882-6658882 | 551 | 565 | + | 3.7786 | 3.38E-05 | 0.71 | GTTACTGTCAAATTG |
| 1 | GMTACGGTCACACTG | chr3R:20830127-20832127 | 555 | 569 | + | 3.7786 | 3.38E-05 | 0.71 | GGTACGGTCATATTA |
| 1 | GMTACGGTCACACTG | chr3L:5151678-5153678 | 573 | 587 | + | 3.7786 | 3.38E-05 | 0.71 | TTTACGTTCACAATG |
| 1 | GMTACGGTCACACTG | chr3L:19804080-19806080 | 575 | 589 | + | 3.7786 | 3.38E-05 | 0.71 | CTTAAAGTCACACTG |
| 1 | GMTACGGTCACACTG | chr2R:11699317-11701317 | 588 | 602 | + | 3.7786 | 3.38E-05 | 0.71 | GGTACAATCACATTG |
| 1 | GMTACGGTCACACTG | chr3R:14161709-14163709 | 602 | 616 | + | 3.7786 | 3.38E-05 | 0.71 | CTTACTTTCACACTG |
| 1 | GMTACGGTCACACTG | chrX:10060627-10062627 | 649 | 663 | + | 3.7786 | 3.38E-05 | 0.71 | GGTATGCTCACATTG |
| 1 | GMTACGGTCACACTG | chr3R:22465034-22467034 | 670 | 684 | + | 3.7786 | 3.38E-05 | 0.71 | TTTACTGTGACACTG |

| | | | | | | | | | |
|---|---|---|---|---|---|---|---|---|---|
| 1 | GMTACGGTCACACTG | chr2L:12084827-12086827 | 692 | 706 | + | 3.7786 | 3.38E-05 | 0.71 | GTTACGGTCATAGTT |
| 1 | GMTACGGTCACACTG | chr3R:11171664-11173664 | 764 | 778 | + | 3.7786 | 3.38E-05 | 0.71 | ATTACGGTGACATTG |
| 1 | GMTACGGTCACACTG | chr2R:10874392-10876392 | 851 | 865 | + | 3.7786 | 3.38E-05 | 0.71 | GTTACTCTCACATTG |
| 1 | GMTACGGTCACACTG | chr3R:13190599-13192599 | 864 | 878 | + | 3.7786 | 3.38E-05 | 0.71 | GTTACGGTGAAACTT |
| 1 | GMTACGGTCACACTG | chr2L:12084649-12086649 | 870 | 884 | + | 3.7786 | 3.38E-05 | 0.71 | GTTACGGTCATAGTT |
| 1 | GMTACGGTCACACTG | chr3L:5151376-5153376 | 875 | 889 | + | 3.7786 | 3.38E-05 | 0.71 | TTTACGTTCACAATG |
| 1 | GMTACGGTCACACTG | chrX:15795028-15797028 | 932 | 946 | + | 3.7786 | 3.38E-05 | 0.71 | GTTACGGTAAAATTG |
| 1 | GMTACGGTCACACTG | chrX:5322855-5324855 | 975 | 989 | + | 3.7786 | 3.38E-05 | 0.71 | AGTATGGTCACACTA |
| 1 | GMTACGGTCACACTG | chr3R:19772225-19774225 | 979 | 993 | + | 3.7786 | 3.38E-05 | 0.71 | GGTACGGTCAGATTT |
| 1 | GMTACGGTCACACTG | chrX:4701284-4703284 | 990 | 1004 | + | 3.7786 | 3.38E-05 | 0.71 | TTTAGGGTCACACTA |
| 1 | GMTACGGTCACACTG | chr3R:6656442-6658442 | 991 | 1005 | + | 3.7786 | 3.38E-05 | 0.71 | GTTACTGTCAAATTG |
| 1 | GMTACGGTCACACTG | chr2R:8162564-8164564 | 997 | 1011 | + | 3.7786 | 3.38E-05 | 0.71 | CGTATGGTCACACTC |
| 1 | GMTACGGTCACACTG | chr2L:5948450-5950450 | 1004 | 1018 | + | 3.7786 | 3.38E-05 | 0.71 | GTTACTGTCAAATTG |
| 1 | GMTACGGTCACACTG | chr2R:20643036-20645036 | 1018 | 1032 | + | 3.7786 | 3.38E-05 | 0.71 | TTTACGGTCATACTA |
| 1 | GMTACGGTCACACTG | chr3L:4555087-4557087 | 1027 | 1041 | + | 3.7786 | 3.38E-05 | 0.71 | GGTACGGTCATAGTT |
| 1 | GMTACGGTCACACTG | chr2R:11292174-11294174 | 1053 | 1067 | + | 3.7786 | 3.38E-05 | 0.71 | GTTATGGTAACACTA |
| 1 | GMTACGGTCACACTG | chr2R:8162500-8164500 | 1061 | 1075 | + | 3.7786 | 3.38E-05 | 0.71 | CGTATGGTCACACTC |
| 1 | GMTACGGTCACACTG | chr2L:2572037-2574037 | 1067 | 1081 | + | 3.7786 | 3.38E-05 | 0.71 | TTTACTTTCACACTG |
| 1 | GMTACGGTCACACTG | chr3L:14704464-14706464 | 1068 | 1082 | + | 3.7786 | 3.38E-05 | 0.71 | TGTACGCTCACAATG |
| 1 | GMTACGGTCACACTG | chr3R:23404509-23406509 | 1078 | 1092 | + | 3.7786 | 3.38E-05 | 0.71 | GGTACGTTGAAACTG |
| 1 | GMTACGGTCACACTG | chr2R:7949339-7951339 | 1082 | 1096 | + | 3.7786 | 3.38E-05 | 0.71 | TTTATGTTCACACTG |
| 1 | GMTACGGTCACACTG | chr3R:20829580-20831580 | 1102 | 1116 | + | 3.7786 | 3.38E-05 | 0.71 | GGTACGGTCATATTA |
| 1 | GMTACGGTCACACTG | chr2L:22174415-22176415 | 1182 | 1196 | + | 3.7786 | 3.38E-05 | 0.71 | GTTACGGTAAGATTG |
| 1 | GMTACGGTCACACTG | chr2R:8012755-8014755 | 1186 | 1200 | + | 3.7786 | 3.38E-05 | 0.71 | ATTAAGGTCACAGTG |
| 1 | GMTACGGTCACACTG | chr2R:20642785-20644785 | 1269 | 1283 | + | 3.7786 | 3.38E-05 | 0.71 | TTTACGGTCATACTA |
| 1 | GMTACGGTCACACTG | chr2R:20642785-20644785 | 1269 | 1283 | + | 3.7786 | 3.38E-05 | 0.71 | TTTACGGTCATACTA |
| 1 | GMTACGGTCACACTG | chr2R:5726525-5728525 | 1310 | 1324 | + | 3.7786 | 3.38E-05 | 0.71 | GTTACGGTTACATTT |
| 1 | GMTACGGTCACACTG | chr2L:9250459-9252459 | 1385 | 1399 | + | 3.7786 | 3.38E-05 | 0.71 | GGTATGTTCACATTG |
| 1 | GMTACGGTCACACTG | chr2R:12486391-12488391 | 1421 | 1435 | + | 3.7786 | 3.38E-05 | 0.71 | CTTAAGGTCACATTG |
| 1 | GMTACGGTCACACTG | chr3R:22701084-22703084 | 1430 | 1444 | + | 3.7786 | 3.38E-05 | 0.71 | TTTACTGTCACACTT |
| 1 | GMTACGGTCACACTG | chr2L:10645415-10647415 | 1434 | 1448 | + | 3.7786 | 3.38E-05 | 0.71 | GTTACAGTCACAGTC |
| 1 | GMTACGGTCACACTG | chr2R:10873771-10875771 | 1472 | 1486 | + | 3.7786 | 3.38E-05 | 0.71 | GTTACTCTCACATTG |
| 1 | GMTACGGTCACACTG | chr3R:7757122-7759122 | 1474 | 1488 | + | 3.7786 | 3.38E-05 | 0.71 | GTTACGGTCAAAATA |

| | | | | | | | | | |
|---|---|---|---|---|---|---|---|---|---|
| 1 | GMTACGGTCACACTG | chrX:12902516-12904516 | 1498 | 1512 | + | 3.7786 | 3.38E-05 | 0.71 | TGTACGTTCACACTT |
| 1 | GMTACGGTCACACTG | chr3L:15528243-15530243 | 1505 | 1519 | + | 3.7786 | 3.38E-05 | 0.71 | GTTATGATCAGACTG |
| 1 | GMTACGGTCACACTG | chr3R:22701007-22703007 | 1507 | 1521 | + | 3.7786 | 3.38E-05 | 0.71 | TTTACTGTCACACTT |
| 1 | GMTACGGTCACACTG | chrX:15452471-15454471 | 1566 | 1580 | + | 3.7786 | 3.38E-05 | 0.71 | GGTACGATCATACTA |
| 1 | GMTACGGTCACACTG | chr3R:31209730-31211730 | 1579 | 1593 | + | 3.7786 | 3.38E-05 | 0.71 | CTTACCATCACACTG |
| 1 | GMTACGGTCACACTG | chr2R:7948839-7950839 | 1582 | 1596 | + | 3.7786 | 3.38E-05 | 0.71 | TTTATGTTCACACTG |
| 1 | GMTACGGTCACACTG | chr2R:8416706-8418706 | 1682 | 1696 | + | 3.7786 | 3.38E-05 | 0.71 | GTTAAGATCAAACTG |
| 1 | GMTACGGTCACACTG | chr2R:11291531-11293531 | 1696 | 1710 | + | 3.7786 | 3.38E-05 | 0.71 | GTTATGGTAACACTA |
| 1 | GMTACGGTCACACTG | chr3R:22463985-22465985 | 1719 | 1733 | + | 3.7786 | 3.38E-05 | 0.71 | TTTACTGTGACACTG |
| 1 | GMTACGGTCACACTG | chr3R:20852358-20854358 | 1742 | 1756 | + | 3.7786 | 3.38E-05 | 0.71 | TTTATGGTCACAGTG |
| 1 | GMTACGGTCACACTG | chrX:16390698-16392698 | 1756 | 1770 | + | 3.7786 | 3.38E-05 | 0.71 | ATTACTGTCACAATG |
| 1 | GMTACGGTCACACTG | chr2L:9250042-9252042 | 1802 | 1816 | + | 3.7786 | 3.38E-05 | 0.71 | GGTATGTTCACATTG |
| 1 | GMTACGGTCACACTG | chr2R:23139577-23141577 | 1836 | 1850 | + | 3.7786 | 3.38E-05 | 0.71 | GTTACAGTCACAGTC |
| 1 | GMTACGGTCACACTG | chr3L:4554236-4556236 | 1878 | 1892 | + | 3.7786 | 3.38E-05 | 0.71 | GGTACGGTCATAGTT |
| 1 | GMTACGGTCACACTG | chr2L:20071195-20073195 | 1892 | 1906 | + | 3.7786 | 3.38E-05 | 0.71 | GTTACCATCAGACTG |
| 1 | GMTACGGTCACACTG | chr2R:22491197-22493197 | 3 | 17 | + | 3.6357 | 4.09E-05 | 0.71 | GCTCAGGTCACGCCG |
| 1 | GMTACGGTCACACTG | chr2L:19431574-19433574 | 7 | 21 | + | 3.6357 | 4.09E-05 | 0.71 | GCTACCGACGCCCTG |
| 1 | GMTACGGTCACACTG | chr2R:24638688-24640688 | 12 | 26 | + | 3.6357 | 4.09E-05 | 0.71 | GCCACTGGCACACGG |
| 1 | GMTACGGTCACACTG | chr3L:3373282-3375282 | 1 | 15 | - | 3.6357 | 4.09E-05 | 0.71 | GATTCGGGCACTCAG |
| 1 | GMTACGGTCACACTG | chr2L:2563830-2565830 | 1 | 15 | - | 3.6357 | 4.09E-05 | 0.71 | GCGAGGGTCGCACGG |
| 1 | GMTACGGTCACACTG | chr3L:3256878-3258878 | 2 | 16 | - | 3.6357 | 4.09E-05 | 0.71 | GCATCGCACACACTG |
| 1 | GMTACGGTCACACTG | chr2R:24902935-24904935 | 5 | 19 | - | 3.6357 | 4.09E-05 | 0.71 | GCCCAGGCCACACTG |
| 1 | GMTACGGTCACACTG | chrX:1202777-1204777 | 6 | 20 | - | 3.6357 | 4.09E-05 | 0.71 | GCTCCGGTCCCCGTG |
| 1 | GMTACGGTCACACTG | chr2L:10199974-10201974 | 7 | 21 | - | 3.6357 | 4.09E-05 | 0.71 | GCTTCGAGCACTCTG |
| 1 | GMTACGGTCACACTG | chrX:16786008-16788008 | 23 | 37 | + | 3.6357 | 4.09E-05 | 0.71 | GCAGCGGTCGCACTT |
| 1 | GMTACGGTCACACTG | chr3L:19804080-19806080 | 9 | 23 | - | 3.6357 | 4.09E-05 | 0.71 | GCCACGGATACTCTG |
| 1 | GMTACGGTCACACTG | chr3L:21766229-21768229 | 31 | 45 | + | 3.6357 | 4.09E-05 | 0.71 | GCACCGCCCACACTG |
| 1 | GMTACGGTCACACTG | chr3R:9746875-9748875 | 31 | 45 | + | 3.6357 | 4.09E-05 | 0.71 | GCGTCGATCTCACTG |
| 1 | GMTACGGTCACACTG | chr3R:24864350-24866350 | 32 | 46 | + | 3.6357 | 4.09E-05 | 0.71 | GCGAAGGACGCACTG |
| 1 | GMTACGGTCACACTG | chr2R:23844001-23846001 | 34 | 48 | + | 3.6357 | 4.09E-05 | 0.71 | GCCACAGGCACCCTG |
| 1 | GMTACGGTCACACTG | chr3R:15356229-15358229 | 21 | 35 | - | 3.6357 | 4.09E-05 | 0.71 | GCTGCGGTCACGTGG |
| 1 | GMTACGGTCACACTG | chrX:22528964-22530964 | 39 | 53 | + | 3.6357 | 4.09E-05 | 0.71 | GCAGCGCTCTCACTG |
| 1 | GMTACGGTCACACTG | chrX:6831684-6833684 | 39 | 53 | + | 3.6357 | 4.09E-05 | 0.71 | GCAATGGCCACACAG |

| | | | | | | | | | |
|---|---|---|---|---|---|---|---|---|---|
| 1 | GMTACGGTCACACTG | chr2R:21670700-21672700 | 32 | 46 | - | 3.6357 | 4.09E-05 | 0.71 | TCCTCGGCCACACTG |
| 1 | GMTACGGTCACACTG | chr3R:24814759-24816759 | 47 | 61 | + | 3.6357 | 4.09E-05 | 0.71 | GCTGCGGTGGCGCTG |
| 1 | GMTACGGTCACACTG | chr2R:18660548-18662548 | 38 | 52 | - | 3.6357 | 4.09E-05 | 0.71 | GCAACGGGCACACCA |
| 1 | GMTACGGTCACACTG | chr2R:18660548-18662548 | 38 | 52 | - | 3.6357 | 4.09E-05 | 0.71 | GCAACGGGCACACCA |
| 1 | GMTACGGTCACACTG | chr2L:10974293-10976293 | 53 | 67 | + | 3.6357 | 4.09E-05 | 0.71 | GCTAAGGACGCGCTG |
| 1 | GMTACGGTCACACTG | chr2R:18638557-18640557 | 41 | 55 | - | 3.6357 | 4.09E-05 | 0.71 | GCGCCGGTCACACCA |
| 1 | GMTACGGTCACACTG | chr3R:8544685-8546685 | 43 | 57 | - | 3.6357 | 4.09E-05 | 0.71 | GCGTCGGTCAGACAG |
| 1 | GMTACGGTCACACTG | chr3L:7346123-7348123 | 45 | 59 | - | 3.6357 | 4.09E-05 | 0.71 | GATTCGGCCACTCCG |
| 1 | GMTACGGTCACACTG | chrX:11347001-11349001 | 59 | 73 | + | 3.6357 | 4.09E-05 | 0.71 | GCGACGGTGCCACCG |
| 1 | GMTACGGTCACACTG | chr2R:24638688-24640688 | 45 | 59 | - | 3.6357 | 4.09E-05 | 0.71 | GATGCGGTCTCTCCG |
| 1 | GMTACGGTCACACTG | chr2L:778173-780173 | 46 | 60 | - | 3.6357 | 4.09E-05 | 0.71 | GCTCCAGCCACACCG |
| 1 | GMTACGGTCACACTG | chr2L:21033775-21035775 | 60 | 74 | + | 3.6357 | 4.09E-05 | 0.71 | GCTCCGCGCACACAG |
| 1 | GMTACGGTCACACTG | chrX:1907239-1909239 | 48 | 62 | - | 3.6357 | 4.09E-05 | 0.71 | GCTCTGGACACGCTG |
| 1 | GMTACGGTCACACTG | chrX:20418613-20420613 | 69 | 83 | + | 3.6357 | 4.09E-05 | 0.71 | GCCACGGACGCAGTG |
| 1 | GMTACGGTCACACTG | chrX:2725678-2727678 | 57 | 71 | - | 3.6357 | 4.09E-05 | 0.71 | GCACCGGTGACACGG |
| 1 | GMTACGGTCACACTG | chr3L:17314006-17316006 | 79 | 93 | + | 3.6357 | 4.09E-05 | 0.71 | GAAACGGCCACGCAG |
| 1 | GMTACGGTCACACTG | chr2R:7392161-7394161 | 66 | 80 | - | 3.6357 | 4.09E-05 | 0.71 | GCTGCCGTCTCTCTG |
| 1 | GMTACGGTCACACTG | chr2R:24596995-24598995 | 81 | 95 | + | 3.6357 | 4.09E-05 | 0.71 | GATTCGGTCGCGCAG |
| 1 | GMTACGGTCACACTG | chr3R:13944812-13946812 | 71 | 85 | - | 3.6357 | 4.09E-05 | 0.71 | GCCATGGACGCACTG |
| 1 | GMTACGGTCACACTG | chr2L:18675998-18677998 | 91 | 105 | + | 3.6357 | 4.09E-05 | 0.71 | GCGACTGACGCACTG |
| 1 | GMTACGGTCACACTG | chr3R:9046977-9048977 | 78 | 92 | - | 3.6357 | 4.09E-05 | 0.71 | GCTCCGGCCACTGTG |
| 1 | GMTACGGTCACACTG | chr2R:9560638-9562638 | 92 | 106 | + | 3.6357 | 4.09E-05 | 0.71 | GCTACGGAGGCACAG |
| 1 | GMTACGGTCACACTG | chr3L:7354717-7356717 | 78 | 92 | - | 3.6357 | 4.09E-05 | 0.71 | GCTTCGGTCACCCAA |
| 1 | GMTACGGTCACACTG | chrX:20418613-20420613 | 79 | 93 | - | 3.6357 | 4.09E-05 | 0.71 | GCTGCAGACGCACTG |
| 1 | GMTACGGTCACACTG | chr2R:12874164-12876164 | 93 | 107 | + | 3.6357 | 4.09E-05 | 0.71 | GCTGCGGTCGCACAC |
| 1 | GMTACGGTCACACTG | chr2L:13826772-13828772 | 83 | 97 | - | 3.6357 | 4.09E-05 | 0.71 | GACACGGGCACTCGG |
| 1 | GMTACGGTCACACTG | chr3R:21087792-21089792 | 98 | 112 | + | 3.6357 | 4.09E-05 | 0.71 | GCCACCGCCACGCTG |
| 1 | GMTACGGTCACACTG | chr3L:391633-393633 | 99 | 113 | + | 3.6357 | 4.09E-05 | 0.71 | GCCACGAACCCACTG |
| 1 | GMTACGGTCACACTG | chrX:1694889-1696889 | 86 | 100 | - | 3.6357 | 4.09E-05 | 0.71 | GCACCGCTCACACAG |
| 1 | GMTACGGTCACACTG | chr3R:22746599-22748599 | 86 | 100 | - | 3.6357 | 4.09E-05 | 0.71 | GCTCCGGACCCAGTG |
| 1 | GMTACGGTCACACTG | chr2R:20985500-20987500 | 102 | 116 | + | 3.6357 | 4.09E-05 | 0.71 | GCTCCGGCGGCACTG |
| 1 | GMTACGGTCACACTG | chr3L:7914437-7916437 | 90 | 104 | - | 3.6357 | 4.09E-05 | 0.71 | GCCACGGACACACGA |
| 1 | GMTACGGTCACACTG | chr3R:24160498-24162498 | 105 | 119 | + | 3.6357 | 4.09E-05 | 0.71 | GCTCCGCTCTCACCG |

| | | | | | | | | | |
|---|---|---|---|---|---|---|---|---|---|
| 1 | GMTACGGTCACACTG | chr3L:22842264-22844264 | 105 | 119 | + | 3.6357 | 4.09E-05 | 0.71 | GCTGCAGTCACGCAG |
| 1 | GMTACGGTCACACTG | chr3L:21743269-21745269 | 93 | 107 | - | 3.6357 | 4.09E-05 | 0.71 | GCCACGGGGACTCTG |
| 1 | GMTACGGTCACACTG | chr3R:5354092-5356092 | 108 | 122 | + | 3.6357 | 4.09E-05 | 0.71 | GCTCCAGTCGCACGG |
| 1 | GMTACGGTCACACTG | chr2R:22885398-22887398 | 99 | 113 | - | 3.6357 | 4.09E-05 | 0.71 | GAGTCGGCCACACAG |
| 1 | GMTACGGTCACACTG | chr3L:8955791-8957791 | 116 | 130 | + | 3.6357 | 4.09E-05 | 0.71 | GCCGCGGGCACAGTG |
| 1 | GMTACGGTCACACTG | chr2L:20646533-20648533 | 123 | 137 | + | 3.6357 | 4.09E-05 | 0.71 | GAGTCGGCCGCACTG |
| 1 | GMTACGGTCACACTG | chr3L:3373170-3375170 | 113 | 127 | - | 3.6357 | 4.09E-05 | 0.71 | GATTCGGGCACTCAG |
| 1 | GMTACGGTCACACTG | chr3R:7525369-7527369 | 130 | 144 | + | 3.6357 | 4.09E-05 | 0.71 | GCAACGGTCACGGGG |
| 1 | GMTACGGTCACACTG | chrX:18857203-18859203 | 130 | 144 | + | 3.6357 | 4.09E-05 | 0.71 | GCTCCGGTCACCAAG |
| 1 | GMTACGGTCACACTG | chrX:17855974-17857974 | 117 | 131 | - | 3.6357 | 4.09E-05 | 0.71 | GCCACGGCCACAAAG |
| 1 | GMTACGGTCACACTG | chr3L:19564131-19566131 | 133 | 147 | + | 3.6357 | 4.09E-05 | 0.71 | GCTAAGGCCACTCAG |
| 1 | GMTACGGTCACACTG | chr3L:8519552-8521552 | 119 | 133 | - | 3.6357 | 4.09E-05 | 0.71 | GCAACGGACTCACTT |
| 1 | GMTACGGTCACACTG | chr2L:6132674-6134674 | 134 | 148 | + | 3.6357 | 4.09E-05 | 0.71 | GCAATGGACACACAG |
| 1 | GMTACGGTCACACTG | chr3R:13259356-13261356 | 134 | 148 | - | 3.6357 | 4.09E-05 | 0.71 | TCGTCGGTCACGCTG |
| 1 | GMTACGGTCACACTG | chrX:16941871-16943871 | 136 | 150 | - | 3.6357 | 4.09E-05 | 0.71 | GCCACGGTCACCGCG |
| 1 | GMTACGGTCACACTG | chr2R:14215627-14217627 | 155 | 169 | + | 3.6357 | 4.09E-05 | 0.71 | GCTGCTGGCACGCTG |
| 1 | GMTACGGTCACACTG | chr3L:9445864-9447864 | 145 | 159 | - | 3.6357 | 4.09E-05 | 0.71 | TCGACGGTCCCGCTG |
| 1 | GMTACGGTCACACTG | chr3R:9746744-9748744 | 162 | 176 | + | 3.6357 | 4.09E-05 | 0.71 | GCGTCGATCTCACTG |
| 1 | GMTACGGTCACACTG | chr3L:3817679-3819679 | 166 | 180 | + | 3.6357 | 4.09E-05 | 0.71 | GCATGGGTCACGCTG |
| 1 | GMTACGGTCACACTG | chr3R:9573211-9575211 | 157 | 171 | - | 3.6357 | 4.09E-05 | 0.71 | GCTGCGGCCACTGTG |
| 1 | GMTACGGTCACACTG | chr2L:2213266-2215266 | 179 | 193 | + | 3.6357 | 4.09E-05 | 0.71 | GCGACGGTTCCACAG |
| 1 | GMTACGGTCACACTG | chr2L:4936409-4938409 | 180 | 194 | + | 3.6357 | 4.09E-05 | 0.71 | CCTTCGGTCCCACGG |
| 1 | GMTACGGTCACACTG | chr2R:16418398-16420398 | 184 | 198 | + | 3.6357 | 4.09E-05 | 0.71 | CCGTCGGTCACACAG |
| 1 | GMTACGGTCACACTG | chr2R:16418398-16420398 | 184 | 198 | + | 3.6357 | 4.09E-05 | 0.71 | CCGTCGGTCACACAG |
| 1 | GMTACGGTCACACTG | chr3L:16300453-16302453 | 186 | 200 | + | 3.6357 | 4.09E-05 | 0.71 | TCTGCGGGCACACAG |
| 1 | GMTACGGTCACACTG | chr2R:7792380-7794380 | 173 | 187 | - | 3.6357 | 4.09E-05 | 0.71 | TCGACGGTCCCTCTG |
| 1 | GMTACGGTCACACTG | chrX:13090145-13092145 | 188 | 202 | + | 3.6357 | 4.09E-05 | 0.71 | GCGGCGGTATCACTG |
| 1 | GMTACGGTCACACTG | chr2R:17664678-17666678 | 189 | 203 | + | 3.6357 | 4.09E-05 | 0.71 | GCCACGCACACACCG |
| 1 | GMTACGGTCACACTG | chr2L:1076948-1078948 | 191 | 205 | + | 3.6357 | 4.09E-05 | 0.71 | GCCTCGCACACACTG |
| 1 | GMTACGGTCACACTG | chr2L:8540761-8542761 | 191 | 205 | + | 3.6357 | 4.09E-05 | 0.71 | GCTCGGGCCACACAG |
| 1 | GMTACGGTCACACTG | chr2L:9575769-9577769 | 199 | 213 | + | 3.6357 | 4.09E-05 | 0.71 | GCTCCGGCCACAAAG |
| 1 | GMTACGGTCACACTG | chr3R:22467512-22469512 | 200 | 214 | + | 3.6357 | 4.09E-05 | 0.71 | GCTGCGGCGGCACTG |
| 1 | GMTACGGTCACACTG | chr3L:16328795-16330795 | 204 | 218 | + | 3.6357 | 4.09E-05 | 0.71 | GCCACAGTCACGCAG |

| | | | | | | | | | |
|---|---|---|---|---|---|---|---|---|---|
| 1 | GMTACGGTCACACTG | chr2R:7062569-7064569 | 191 | 205 | - | 3.6357 | 4.09E-05 | 0.71 | GCTGCGGACAGACGG |
| 1 | GMTACGGTCACACTG | chr3L:8179514-8181514 | 207 | 221 | + | 3.6357 | 4.09E-05 | 0.71 | GCAAAGGGCACACAG |
| 1 | GMTACGGTCACACTG | chr3R:15859196-15861196 | 207 | 221 | + | 3.6357 | 4.09E-05 | 0.71 | ACTCCGGCCACACAG |
| 1 | GMTACGGTCACACTG | chr2L:15627678-15629678 | 211 | 225 | + | 3.6357 | 4.09E-05 | 0.71 | GCCACGCCCACTCTG |
| 1 | GMTACGGTCACACTG | chr3R:11683342-11685342 | 218 | 232 | + | 3.6357 | 4.09E-05 | 0.71 | GCTGCGTTCTCGCTG |
| 1 | GMTACGGTCACACTG | chr3L:19519945-19521945 | 219 | 233 | + | 3.6357 | 4.09E-05 | 0.71 | GCTACGGACACGGCG |
| 1 | GMTACGGTCACACTG | chr2R:9094996-9096996 | 225 | 239 | + | 3.6357 | 4.09E-05 | 0.71 | GCAGCGTGCACACTG |
| 1 | GMTACGGTCACACTG | chr2L:2213211-2215211 | 234 | 248 | + | 3.6357 | 4.09E-05 | 0.71 | GCGACGGTTCCACAG |
| 1 | GMTACGGTCACACTG | chr3L:8835462-8837462 | 221 | 235 | - | 3.6357 | 4.09E-05 | 0.71 | GCCCCCGTCACGCTG |
| 1 | GMTACGGTCACACTG | chr2R:12874012-12876012 | 245 | 259 | + | 3.6357 | 4.09E-05 | 0.71 | GCTGCGGTCGCACAC |
| 1 | GMTACGGTCACACTG | chr2R:18225795-18227795 | 236 | 250 | - | 3.6357 | 4.09E-05 | 0.71 | GCTGCGGTCATTCGG |
| 1 | GMTACGGTCACACTG | chr2R:12678593-12680593 | 256 | 270 | + | 3.6357 | 4.09E-05 | 0.71 | GCGTCAGTCACGCTG |
| 1 | GMTACGGTCACACTG | chr2R:22885247-22887247 | 250 | 264 | - | 3.6357 | 4.09E-05 | 0.71 | GAGTCGGCCACACAG |
| 1 | GMTACGGTCACACTG | chr3L:6588479-6590479 | 269 | 283 | + | 3.6357 | 4.09E-05 | 0.71 | GAAACGGTCGCCCCG |
| 1 | GMTACGGTCACACTG | chr2R:22613328-22615328 | 270 | 284 | + | 3.6357 | 4.09E-05 | 0.71 | GCAACAGCCACACCG |
| 1 | GMTACGGTCACACTG | chr2R:23843763-23845763 | 272 | 286 | + | 3.6357 | 4.09E-05 | 0.71 | GCCACAGGCACCCTG |
| 1 | GMTACGGTCACACTG | chr2L:4954144-4956144 | 259 | 273 | - | 3.6357 | 4.09E-05 | 0.71 | GCAACGGTTGCACCG |
| 1 | GMTACGGTCACACTG | chr2R:23346641-23348641 | 282 | 296 | + | 3.6357 | 4.09E-05 | 0.71 | GCGGCGGTCACCATG |
| 1 | GMTACGGTCACACTG | chr3L:22423909-22425909 | 282 | 296 | + | 3.6357 | 4.09E-05 | 0.71 | GCAACAGCCACGCTG |
| 1 | GMTACGGTCACACTG | chrX:11871779-11873779 | 290 | 304 | + | 3.6357 | 4.09E-05 | 0.71 | GCTTCGGTTTCACGG |
| 1 | GMTACGGTCACACTG | chrX:12456763-12458763 | 278 | 292 | - | 3.6357 | 4.09E-05 | 0.71 | GCCACGGACACGGTG |
| 1 | GMTACGGTCACACTG | chr3L:1249696-1251696 | 293 | 307 | + | 3.6357 | 4.09E-05 | 0.71 | GCCACGGGCATCCTG |
| 1 | GMTACGGTCACACTG | chrX:7933898-7935898 | 282 | 296 | - | 3.6357 | 4.09E-05 | 0.71 | GCGATGGTCTCACGG |
| 1 | GMTACGGTCACACTG | chr2R:7038130-7040130 | 299 | 313 | + | 3.6357 | 4.09E-05 | 0.71 | GCGACGGACACAAAG |
| 1 | GMTACGGTCACACTG | chrX:18063483-18065483 | 289 | 303 | - | 3.6357 | 4.09E-05 | 0.71 | GCTTCGGACACATAG |
| 1 | GMTACGGTCACACTG | chr3R:29051680-29053680 | 305 | 319 | + | 3.6357 | 4.09E-05 | 0.71 | GCAACGGCGACTCTG |
| 1 | GMTACGGTCACACTG | chr2L:15037861-15039861 | 307 | 321 | + | 3.6357 | 4.09E-05 | 0.71 | GCTACAGACACGCAG |
| 1 | GMTACGGTCACACTG | chr2L:2551825-2553825 | 296 | 310 | - | 3.6357 | 4.09E-05 | 0.71 | GCGACGGCCACTATG |
| 1 | GMTACGGTCACACTG | chr3R:5968983-5970983 | 296 | 310 | - | 3.6357 | 4.09E-05 | 0.71 | GCGATGGGCGCACTG |
| 1 | GMTACGGTCACACTG | chr2R:17668395-17670395 | 299 | 313 | - | 3.6357 | 4.09E-05 | 0.71 | GCCACGGTCACCGCG |
| 1 | GMTACGGTCACACTG | chr3R:27947711-27949711 | 304 | 318 | - | 3.6357 | 4.09E-05 | 0.71 | GCGACGTTCCCGCTG |
| 1 | GMTACGGTCACACTG | chr3R:24658621-24660621 | 319 | 333 | + | 3.6357 | 4.09E-05 | 0.71 | GCGACGTTCCCACAG |
| 1 | GMTACGGTCACACTG | chr2L:320248-322248 | 310 | 324 | - | 3.6357 | 4.09E-05 | 0.71 | GCAACGGTGACTCCG |

| | | | | | | | | | |
|---|---|---|---|---|---|---|---|---|---|
| 1 | GMTACGGTCACACTG | chr2L:1076815-1078815 | 324 | 338 | + | 3.6357 | 4.09E-05 | 0.71 | GCCTCGCACACACTG |
| 1 | GMTACGGTCACACTG | chr3L:18147923-18149923 | 311 | 325 | - | 3.6357 | 4.09E-05 | 0.71 | GCTACTGCCTCTCTG |
| 1 | GMTACGGTCACACTG | chr2R:22895174-22897174 | 326 | 340 | + | 3.6357 | 4.09E-05 | 0.71 | GCAACGGTGACTCCG |
| 1 | GMTACGGTCACACTG | chr3L:836861-838861 | 312 | 326 | - | 3.6357 | 4.09E-05 | 0.71 | GAAACGGACTCCCTG |
| 1 | GMTACGGTCACACTG | chr2L:13394131-13396131 | 315 | 329 | - | 3.6357 | 4.09E-05 | 0.71 | GCCAAGGTCACGCAG |
| 1 | GMTACGGTCACACTG | chr2R:5726525-5728525 | 315 | 329 | - | 3.6357 | 4.09E-05 | 0.71 | GCTGCGCTCGCACAG |
| 1 | GMTACGGTCACACTG | chr2L:8540623-8542623 | 329 | 343 | + | 3.6357 | 4.09E-05 | 0.71 | GCTCGGGCCACACAG |
| 1 | GMTACGGTCACACTG | chr3R:17035045-17037045 | 330 | 344 | + | 3.6357 | 4.09E-05 | 0.71 | GCATCCGACACACTG |
| 1 | GMTACGGTCACACTG | chr3L:13434229-13436229 | 316 | 330 | - | 3.6357 | 4.09E-05 | 0.71 | CCGACGGTCACTCAG |
| 1 | GMTACGGTCACACTG | chr3R:16271452-16273452 | 317 | 331 | - | 3.6357 | 4.09E-05 | 0.71 | GCCACGGGCACTTTG |
| 1 | GMTACGGTCACACTG | chr3R:15355924-15357924 | 326 | 340 | - | 3.6357 | 4.09E-05 | 0.71 | GCTGCGGTCACGTGG |
| 1 | GMTACGGTCACACTG | chr3R:28539324-28541324 | 328 | 342 | - | 3.6357 | 4.09E-05 | 0.71 | GCTACGTGCTCGCTG |
| 1 | GMTACGGTCACACTG | chr2R:18638270-18640270 | 328 | 342 | - | 3.6357 | 4.09E-05 | 0.71 | GCGCCGGTCACACCA |
| 1 | GMTACGGTCACACTG | chr2R:12798259-12800259 | 343 | 357 | + | 3.6357 | 4.09E-05 | 0.71 | GCTGCTGTCACTCAG |
| 1 | GMTACGGTCACACTG | chr3L:5534759-5536759 | 332 | 346 | - | 3.6357 | 4.09E-05 | 0.71 | GCTACGGTTCCTCCG |
| 1 | GMTACGGTCACACTG | chr2L:4954144-4956144 | 334 | 348 | - | 3.6357 | 4.09E-05 | 0.71 | GCTCCGGTGACTCCG |
| 1 | GMTACGGTCACACTG | chr3R:20035056-20037056 | 353 | 367 | + | 3.6357 | 4.09E-05 | 0.71 | GCGATGGCCACGCTG |
| 1 | GMTACGGTCACACTG | chr3R:17419638-17421638 | 362 | 376 | + | 3.6357 | 4.09E-05 | 0.71 | GCACCGGTCAATCTG |
| 1 | GMTACGGTCACACTG | chrX:20393395-20395395 | 349 | 363 | - | 3.6357 | 4.09E-05 | 0.71 | TCAGCGGGCACACTG |
| 1 | GMTACGGTCACACTG | chr2R:10311023-10313023 | 350 | 364 | - | 3.6357 | 4.09E-05 | 0.71 | GCAGCGCTCTCACTG |
| 1 | GMTACGGTCACACTG | chr3L:836821-838821 | 352 | 366 | - | 3.6357 | 4.09E-05 | 0.71 | GAAACGGACTCCCTG |
| 1 | GMTACGGTCACACTG | chrX:2603197-2605197 | 367 | 381 | + | 3.6357 | 4.09E-05 | 0.71 | GCCACTGCCACACAG |
| 1 | GMTACGGTCACACTG | chr3R:16271415-16273415 | 354 | 368 | - | 3.6357 | 4.09E-05 | 0.71 | GCCACGGGCACTTTG |
| 1 | GMTACGGTCACACTG | chr2R:15567960-15569960 | 354 | 368 | - | 3.6357 | 4.09E-05 | 0.71 | GCCTTGGTCACCCTG |
| 1 | GMTACGGTCACACTG | chr2L:15006431-15008431 | 355 | 369 | - | 3.6357 | 4.09E-05 | 0.71 | GCCACGGTCCTTCTG |
| 1 | GMTACGGTCACACTG | chr2R:22482303-22484303 | 370 | 384 | + | 3.6357 | 4.09E-05 | 0.71 | CCTACGGTCTCGCAG |
| 1 | GMTACGGTCACACTG | chrX:19958403-19960403 | 357 | 371 | - | 3.6357 | 4.09E-05 | 0.71 | GCTACGGATACGCCG |
| 1 | GMTACGGTCACACTG | chrX:9273714-9275714 | 363 | 377 | - | 3.6357 | 4.09E-05 | 0.71 | GCAGTGGTCGCACTG |
| 1 | GMTACGGTCACACTG | chrX:20392751-20394751 | 378 | 392 | + | 3.6357 | 4.09E-05 | 0.71 | GCTGCGCACGCACTG |
| 1 | GMTACGGTCACACTG | chr2R:7038050-7040050 | 379 | 393 | + | 3.6357 | 4.09E-05 | 0.71 | GCGACGGACACAAAG |
| 1 | GMTACGGTCACACTG | chr3R:9732250-9734250 | 384 | 398 | + | 3.6357 | 4.09E-05 | 0.71 | GCTCCAGTCTCGCTG |
| 1 | GMTACGGTCACACTG | chr3R:28303605-28305605 | 377 | 391 | - | 3.6357 | 4.09E-05 | 0.71 | GCGATGGACCCACTG |
| 1 | GMTACGGTCACACTG | chr2L:115488-117488 | 393 | 407 | + | 3.6357 | 4.09E-05 | 0.71 | GCTGCGCTCGCACGG |

| | | | | | | | | | |
|---|---|---|---|---|---|---|---|---|---|
| 1 | GMTACGGTCACACTG | chrX:1408069-1410069 | 379 | 393 | - | 3.6357 | 4.09E-05 | 0.71 | GCTGCGGTCGCTTTG |
| 1 | GMTACGGTCACACTG | chrX:10898302-10900302 | 388 | 402 | - | 3.6357 | 4.09E-05 | 0.71 | GCTCCGGCGACACGG |
| 1 | GMTACGGTCACACTG | chr3R:30038719-30040719 | 390 | 404 | - | 3.6357 | 4.09E-05 | 0.71 | GCGGCGGTCGCATTG |
| 1 | GMTACGGTCACACTG | chr3R:30038719-30040719 | 390 | 404 | - | 3.6357 | 4.09E-05 | 0.71 | GCGGCGGTCGCATTG |
| 1 | GMTACGGTCACACTG | chr3L:7538836-7540836 | 391 | 405 | - | 3.6357 | 4.09E-05 | 0.71 | GCTACGGCGACTCAG |
| 1 | GMTACGGTCACACTG | chr3L:5354923-5356923 | 407 | 421 | + | 3.6357 | 4.09E-05 | 0.71 | GCAGCGGACAGACTG |
| 1 | GMTACGGTCACACTG | chr3R:27004469-27006469 | 415 | 429 | + | 3.6357 | 4.09E-05 | 0.71 | GCTCCGGCCACAGAG |
| 1 | GMTACGGTCACACTG | chr3L:9732807-9734807 | 403 | 417 | - | 3.6357 | 4.09E-05 | 0.71 | GCTGCGGTCTCGTTG |
| 1 | GMTACGGTCACACTG | chr3R:24133831-24135831 | 419 | 433 | + | 3.6357 | 4.09E-05 | 0.71 | GCATCGGTCACACGC |
| 1 | GMTACGGTCACACTG | chr2L:14801334-14803334 | 407 | 421 | - | 3.6357 | 4.09E-05 | 0.71 | GAACCGGACCCACTG |
| 1 | GMTACGGTCACACTG | chr3L:7914120-7916120 | 407 | 421 | - | 3.6357 | 4.09E-05 | 0.71 | GCCACGGACACACGA |
| 1 | GMTACGGTCACACTG | chr2L:5546086-5548086 | 427 | 441 | + | 3.6357 | 4.09E-05 | 0.71 | GCAACGCCCACACAG |
| 1 | GMTACGGTCACACTG | chr3R:22582225-22584225 | 431 | 445 | + | 3.6357 | 4.09E-05 | 0.71 | GAAACGGTCTCGCCG |
| 1 | GMTACGGTCACACTG | chrX:8572593-8574593 | 439 | 453 | + | 3.6357 | 4.09E-05 | 0.71 | CCAACGGACACACAG |
| 1 | GMTACGGTCACACTG | chrX:18847566-18849566 | 426 | 440 | - | 3.6357 | 4.09E-05 | 0.71 | GCAACGGCCCGACTG |
| 1 | GMTACGGTCACACTG | chr3L:9732780-9734780 | 430 | 444 | - | 3.6357 | 4.09E-05 | 0.71 | GCTGCGGTCTCGTTG |
| 1 | GMTACGGTCACACTG | chr2R:7069043-7071043 | 431 | 445 | - | 3.6357 | 4.09E-05 | 0.71 | GAAGCGGTCCCACAG |
| 1 | GMTACGGTCACACTG | chrX:6866194-6868194 | 431 | 445 | - | 3.6357 | 4.09E-05 | 0.71 | TCTACGGGCTCCCTG |
| 1 | GMTACGGTCACACTG | chr3R:28598834-28600834 | 433 | 447 | - | 3.6357 | 4.09E-05 | 0.71 | GCAGCAGCCACACTG |
| 1 | GMTACGGTCACACTG | chr2L:16287290-16289290 | 433 | 447 | - | 3.6357 | 4.09E-05 | 0.71 | GCCGTGGTCACTCTG |
| 1 | GMTACGGTCACACTG | chr3R:5353747-5355747 | 453 | 467 | + | 3.6357 | 4.09E-05 | 0.71 | GCTCCAGTCGCACGG |
| 1 | GMTACGGTCACACTG | chr3L:4239505-4241505 | 453 | 467 | + | 3.6357 | 4.09E-05 | 0.71 | GCCACGTACACTCTG |
| 1 | GMTACGGTCACACTG | chr3R:17013576-17015576 | 454 | 468 | + | 3.6357 | 4.09E-05 | 0.71 | ACGACGGTCACGCAG |
| 1 | GMTACGGTCACACTG | chr3R:14121654-14123654 | 458 | 472 | + | 3.6357 | 4.09E-05 | 0.71 | GCTGCGGGCAAACGG |
| 1 | GMTACGGTCACACTG | chr3R:30458521-30460521 | 447 | 461 | - | 3.6357 | 4.09E-05 | 0.71 | GCACCTGTCGCACTG |
| 1 | GMTACGGTCACACTG | chr2R:24598947-24600947 | 463 | 477 | + | 3.6357 | 4.09E-05 | 0.71 | GCTCCGGCCACAGCG |
| 1 | GMTACGGTCACACTG | chr3L:10888990-10890990 | 464 | 478 | + | 3.6357 | 4.09E-05 | 0.71 | GCAGCGGTCAAGCTG |
| 1 | GMTACGGTCACACTG | chr2R:10785493-10787493 | 464 | 478 | + | 3.6357 | 4.09E-05 | 0.71 | GCTCCGGCCTCAGTG |
| 1 | GMTACGGTCACACTG | chr2L:1999058-2001058 | 451 | 465 | - | 3.6357 | 4.09E-05 | 0.71 | GCTTCGGCTGCACTG |
| 1 | GMTACGGTCACACTG | chrX:15641108-15643108 | 452 | 466 | - | 3.6357 | 4.09E-05 | 0.71 | GCCACGCCCACACCG |
| 1 | GMTACGGTCACACTG | chrX:7934896-7936896 | 453 | 467 | - | 3.6357 | 4.09E-05 | 0.71 | GCGATGGTCTCACGG |
| 1 | GMTACGGTCACACTG | chr3R:12696322-12698322 | 468 | 482 | + | 3.6357 | 4.09E-05 | 0.71 | GCAGCGGTCATCCTG |
| 1 | GMTACGGTCACACTG | chr2L:276582-278582 | 456 | 470 | - | 3.6357 | 4.09E-05 | 0.71 | GCGCCGGTCGCACTC |

| | | | | | | | | | |
|---|---|---|---|---|---|---|---|---|---|
| 1 | GMTACGGTCACACTG | chr3L:20887002-20889002 | 456 | 470 | - | 3.6357 | 4.09E-05 | 0.71 | GCGCCGGGCACACTC |
| 1 | GMTACGGTCACACTG | chr3R:18254530-18256530 | 458 | 472 | - | 3.6357 | 4.09E-05 | 0.71 | GCTGCAGTCTCACAG |
| 1 | GMTACGGTCACACTG | chr3R:18254530-18256530 | 458 | 472 | - | 3.6357 | 4.09E-05 | 0.71 | GCTGCAGTCTCACAG |
| 1 | GMTACGGTCACACTG | chrX:2681932-2683932 | 472 | 486 | - | 3.6357 | 4.09E-05 | 0.71 | GCGAGGGTCACGCCG |
| 1 | GMTACGGTCACACTG | chrX:10325464-10327464 | 472 | 486 | - | 3.6357 | 4.09E-05 | 0.71 | GCGGCGGGCACATTG |
| 1 | GMTACGGTCACACTG | chr2R:13157666-13159666 | 474 | 488 | - | 3.6357 | 4.09E-05 | 0.71 | CCCACGGGCGCACTG |
| 1 | GMTACGGTCACACTG | chr3R:29376646-29378646 | 475 | 489 | - | 3.6357 | 4.09E-05 | 0.71 | GCTTCGGCCACACAT |
| 1 | GMTACGGTCACACTG | chr3R:8340100-8342100 | 490 | 504 | + | 3.6357 | 4.09E-05 | 0.71 | GCGACGGTGACGCGG |
| 1 | GMTACGGTCACACTG | chr2R:8134315-8136315 | 493 | 507 | + | 3.6357 | 4.09E-05 | 0.71 | GCCTCGGTCACCTTG |
| 1 | GMTACGGTCACACTG | chr3R:6220781-6222781 | 496 | 510 | + | 3.6357 | 4.09E-05 | 0.71 | CCCACGGGCACCCTG |
| 1 | GMTACGGTCACACTG | chr3L:16609418-16611418 | 500 | 514 | + | 3.6357 | 4.09E-05 | 0.71 | GCCACGGCCTCAGTG |
| 1 | GMTACGGTCACACTG | chr3L:1568181-1570181 | 500 | 514 | + | 3.6357 | 4.09E-05 | 0.71 | ACCACGGGCACCCTG |
| 1 | GMTACGGTCACACTG | chr3L:1237319-1239319 | 505 | 519 | + | 3.6357 | 4.09E-05 | 0.71 | GCTGCGGTCTCCCTA |
| 1 | GMTACGGTCACACTG | chr3L:13432048-13434048 | 507 | 521 | + | 3.6357 | 4.09E-05 | 0.71 | GCTCCTGCCACCCTG |
| 1 | GMTACGGTCACACTG | chr2R:9838776-9840776 | 510 | 524 | + | 3.6357 | 4.09E-05 | 0.71 | GCATCGGTCACCCTA |
| 1 | GMTACGGTCACACTG | chrX:2603052-2605052 | 512 | 526 | + | 3.6357 | 4.09E-05 | 0.71 | GCCACTGCCACACAG |
| 1 | GMTACGGTCACACTG | chr3R:10878319-10880319 | 499 | 513 | - | 3.6357 | 4.09E-05 | 0.71 | GCCACTGCCTCACTG |
| 1 | GMTACGGTCACACTG | chr3R:18254487-18256487 | 501 | 515 | - | 3.6357 | 4.09E-05 | 0.71 | GCTGCAGTCTCACAG |
| 1 | GMTACGGTCACACTG | chr3R:18254487-18256487 | 501 | 515 | - | 3.6357 | 4.09E-05 | 0.71 | GCTGCAGTCTCACAG |
| 1 | GMTACGGTCACACTG | chr3R:29147248-29149248 | 503 | 517 | - | 3.6357 | 4.09E-05 | 0.71 | GCCGCGCTCACCCTG |
| 1 | GMTACGGTCACACTG | chr3L:13441594-13443594 | 506 | 520 | - | 3.6357 | 4.09E-05 | 0.71 | TCTACGGCCTCGCTG |
| 1 | GMTACGGTCACACTG | chr2R:24410870-24412870 | 515 | 529 | - | 3.6357 | 4.09E-05 | 0.71 | GCCACAGACACTCTG |
| 1 | GMTACGGTCACACTG | chr2R:24410870-24412870 | 515 | 529 | - | 3.6357 | 4.09E-05 | 0.71 | GCCACAGACACTCTG |
| 1 | GMTACGGTCACACTG | chr3L:16294547-16296547 | 516 | 530 | - | 3.6357 | 4.09E-05 | 0.71 | GCTACGGACACGGAG |
| 1 | GMTACGGTCACACTG | chr3R:30048293-30050293 | 534 | 548 | + | 3.6357 | 4.09E-05 | 0.71 | GCTGCGGTAGCGCTG |
| 1 | GMTACGGTCACACTG | chr2R:9094685-9096685 | 536 | 550 | + | 3.6357 | 4.09E-05 | 0.71 | GCAGCGTGCACACTG |
| 1 | GMTACGGTCACACTG | chr2L:15424580-15426580 | 543 | 557 | + | 3.6357 | 4.09E-05 | 0.71 | GCGACGGGCCAACTG |
| 1 | GMTACGGTCACACTG | chr3R:9327511-9329511 | 529 | 543 | - | 3.6357 | 4.09E-05 | 0.71 | GCGAGGGCCACACGG |
| 1 | GMTACGGTCACACTG | chr3R:8340043-8342043 | 547 | 561 | + | 3.6357 | 4.09E-05 | 0.71 | GCGACGGTGACGCGG |
| 1 | GMTACGGTCACACTG | chrX:6329842-6331842 | 548 | 562 | + | 3.6357 | 4.09E-05 | 0.71 | GCGTCGCTCTCACTG |
| 1 | GMTACGGTCACACTG | chr2L:824969-826969 | 534 | 548 | - | 3.6357 | 4.09E-05 | 0.71 | GCTACGGGTTCACGG |
| 1 | GMTACGGTCACACTG | chr2R:15691448-15693448 | 549 | 563 | + | 3.6357 | 4.09E-05 | 0.71 | GCTCCGGTGACTCAG |
| 1 | GMTACGGTCACACTG | chr3L:1864430-1866430 | 537 | 551 | - | 3.6357 | 4.09E-05 | 0.71 | TCTACGGGCACTCCG |

| | | | | | | | | | |
|---|---|---|---|---|---|---|---|---|---|
| 1 | GMTACGGTCACACTG | chr3R:23025713-23027713 | 554 | 568 | + | 3.6357 | 4.09E-05 | 0.71 | GCTCCCGGCACACGG |
| 1 | GMTACGGTCACACTG | chr3R:22467157-22469157 | 555 | 569 | + | 3.6357 | 4.09E-05 | 0.71 | GCTGCGGCGGCACTG |
| 1 | GMTACGGTCACACTG | chr2L:16715195-16717195 | 545 | 559 | - | 3.6357 | 4.09E-05 | 0.71 | GCTACCGACACCCAG |
| 1 | GMTACGGTCACACTG | chr2L:13548328-13550328 | 561 | 575 | + | 3.6357 | 4.09E-05 | 0.71 | GCTTCGGTCTCTCTC |
| 1 | GMTACGGTCACACTG | chr3L:7975406-7977406 | 564 | 578 | + | 3.6357 | 4.09E-05 | 0.71 | GCTCCGTTCGCCCTG |
| 1 | GMTACGGTCACACTG | chr2R:7386187-7388187 | 566 | 580 | + | 3.6357 | 4.09E-05 | 0.71 | GAATCGGCCGCACTG |
| 1 | GMTACGGTCACACTG | chrX:4774232-4776232 | 555 | 569 | - | 3.6357 | 4.09E-05 | 0.71 | GCCACGGACACAGGG |
| 1 | GMTACGGTCACACTG | chr2R:7157241-7159241 | 556 | 570 | - | 3.6357 | 4.09E-05 | 0.71 | GAGCCGGTCACTCCG |
| 1 | GMTACGGTCACACTG | chr3L:16104852-16106852 | 557 | 571 | - | 3.6357 | 4.09E-05 | 0.71 | ACTGCGGTCTCACGG |
| 1 | GMTACGGTCACACTG | chr2R:17584692-17586692 | 571 | 585 | + | 3.6357 | 4.09E-05 | 0.71 | GCAACGGTCACCTGG |
| 1 | GMTACGGTCACACTG | chrX:18856757-18858757 | 576 | 590 | + | 3.6357 | 4.09E-05 | 0.71 | GCTCCGGTCACCAAG |
| 1 | GMTACGGTCACACTG | chrX:10330818-10332818 | 576 | 590 | + | 3.6357 | 4.09E-05 | 0.71 | GCCACGGGACCACTG |
| 1 | GMTACGGTCACACTG | chr2R:23384468-23386468 | 565 | 579 | - | 3.6357 | 4.09E-05 | 0.71 | GCTTCGTTCTCACGG |
| 1 | GMTACGGTCACACTG | chr2R:8088482-8090482 | 572 | 586 | - | 3.6357 | 4.09E-05 | 0.71 | GCTCCAGACACTCTG |
| 1 | GMTACGGTCACACTG | chr2R:12518166-12520166 | 589 | 603 | + | 3.6357 | 4.09E-05 | 0.71 | GCTCCGGCCACCGTG |
| 1 | GMTACGGTCACACTG | chr2R:24598821-24600821 | 589 | 603 | + | 3.6357 | 4.09E-05 | 0.71 | GCTCCGGCCACAGCG |
| 1 | GMTACGGTCACACTG | chr3R:30038533-30040533 | 576 | 590 | - | 3.6357 | 4.09E-05 | 0.71 | GCGGCGGTCGCATTG |
| 1 | GMTACGGTCACACTG | chr3R:3586054-3588054 | 592 | 606 | + | 3.6357 | 4.09E-05 | 0.71 | GCCACTGGCACCCTG |
| 1 | GMTACGGTCACACTG | chr3R:29147172-29149172 | 579 | 593 | - | 3.6357 | 4.09E-05 | 0.71 | GCCGCGCTCACCCTG |
| 1 | GMTACGGTCACACTG | chr3R:27953112-27955112 | 580 | 594 | - | 3.6357 | 4.09E-05 | 0.71 | GCAACGGTCGCTCTT |
| 1 | GMTACGGTCACACTG | chrX:13296082-13298082 | 609 | 623 | + | 3.6357 | 4.09E-05 | 0.71 | GCCCTGGTCACTCTG |
| 1 | GMTACGGTCACACTG | chr2R:13445309-13447309 | 616 | 630 | + | 3.6357 | 4.09E-05 | 0.71 | GAGCCGGGCGCACTG |
| 1 | GMTACGGTCACACTG | chr2R:10246814-10248814 | 603 | 617 | - | 3.6357 | 4.09E-05 | 0.71 | GCCACGCCCACACCG |
| 1 | GMTACGGTCACACTG | chrX:22656324-22658324 | 604 | 618 | - | 3.6357 | 4.09E-05 | 0.71 | GCTGCGGCTGCACTG |
| 1 | GMTACGGTCACACTG | chr3R:27217184-27219184 | 605 | 619 | - | 3.6357 | 4.09E-05 | 0.71 | GCCACCGCCACTCTG |
| 1 | GMTACGGTCACACTG | chr3L:18674800-18676800 | 615 | 629 | - | 3.6357 | 4.09E-05 | 0.71 | GCAACAGTCCCACCG |
| 1 | GMTACGGTCACACTG | chr2R:23385521-23387521 | 631 | 645 | + | 3.6357 | 4.09E-05 | 0.71 | GCCACGGCCGCAATG |
| 1 | GMTACGGTCACACTG | chr2L:4829693-4831693 | 622 | 636 | - | 3.6357 | 4.09E-05 | 0.71 | ACCACGGACGCACTG |
| 1 | GMTACGGTCACACTG | chr2R:18620541-18622541 | 636 | 650 | + | 3.6357 | 4.09E-05 | 0.71 | GCCACGCCCACTCTG |
| 1 | GMTACGGTCACACTG | chr2R:7386115-7388115 | 638 | 652 | + | 3.6357 | 4.09E-05 | 0.71 | GAATCGGCCGCACTG |
| 1 | GMTACGGTCACACTG | chr3R:14495378-14497378 | 631 | 645 | - | 3.6357 | 4.09E-05 | 0.71 | GCGACGGGAACTCTG |
| 1 | GMTACGGTCACACTG | chr3R:24863732-24865732 | 650 | 664 | + | 3.6357 | 4.09E-05 | 0.71 | GCGAAGGACGCACTG |
| 1 | GMTACGGTCACACTG | chr2L:7859839-7861839 | 636 | 650 | - | 3.6357 | 4.09E-05 | 0.71 | GCTACGGCGCCCCTG |

| | | | | | | | | | |
|---|---|---|---|---|---|---|---|---|---|
| 1 | GMTACGGTCACACTG | chr3L:691919-693919 | 653 | 667 | + | 3.6357 | 4.09E-05 | 0.71 | GCCTCGGGCAAACTG |
| 1 | GMTACGGTCACACTG | chr2R:13664114-13666114 | 653 | 667 | + | 3.6357 | 4.09E-05 | 0.71 | GCTGCGGTCGCGCTT |
| 1 | GMTACGGTCACACTG | chr3L:19792935-19794935 | 640 | 654 | - | 3.6357 | 4.09E-05 | 0.71 | GCACTGGCCACACTG |
| 1 | GMTACGGTCACACTG | chr2L:824861-826861 | 642 | 656 | - | 3.6357 | 4.09E-05 | 0.71 | GCTACGGGTTCACGG |
| 1 | GMTACGGTCACACTG | chr2L:13373019-13375019 | 643 | 657 | - | 3.6357 | 4.09E-05 | 0.71 | GCTCCGGCCAACCTG |
| 1 | GMTACGGTCACACTG | chr2R:20778487-20780487 | 658 | 672 | + | 3.6357 | 4.09E-05 | 0.71 | GCTGCGGTCGCGGTG |
| 1 | GMTACGGTCACACTG | chr3L:9378534-9380534 | 667 | 681 | + | 3.6357 | 4.09E-05 | 0.71 | GCTCCGGCCACACGA |
| 1 | GMTACGGTCACACTG | chr3L:7403181-7405181 | 673 | 687 | + | 3.6357 | 4.09E-05 | 0.71 | GCCAAGGCCACGCTG |
| 1 | GMTACGGTCACACTG | chr3R:26846809-26848809 | 677 | 691 | + | 3.6357 | 4.09E-05 | 0.71 | GCTTCGGTCGCAAAG |
| 1 | GMTACGGTCACACTG | chr3R:30458300-30460300 | 668 | 682 | - | 3.6357 | 4.09E-05 | 0.71 | GCACCTGTCGCACTG |
| 1 | GMTACGGTCACACTG | chr3R:19027931-19029931 | 682 | 696 | + | 3.6357 | 4.09E-05 | 0.71 | TCTTCGGTCACCCGG |
| 1 | GMTACGGTCACACTG | chr3L:19909300-19911300 | 682 | 696 | + | 3.6357 | 4.09E-05 | 0.71 | GCGACGGTCGGACCG |
| 1 | GMTACGGTCACACTG | chrX:17225867-17227867 | 673 | 687 | - | 3.6357 | 4.09E-05 | 0.71 | GCTCTGGTCGCTCTG |
| 1 | GMTACGGTCACACTG | chrX:10756666-10758666 | 688 | 702 | + | 3.6357 | 4.09E-05 | 0.71 | GCACCGTTCCCACTG |
| 1 | GMTACGGTCACACTG | chr2R:23385463-23387463 | 689 | 703 | + | 3.6357 | 4.09E-05 | 0.71 | GCCACGGCCGCAATG |
| 1 | GMTACGGTCACACTG | chr3L:8902349-8904349 | 675 | 689 | - | 3.6357 | 4.09E-05 | 0.71 | GATGCGGACACGCCG |
| 1 | GMTACGGTCACACTG | chr3L:11457490-11459490 | 692 | 706 | + | 3.6357 | 4.09E-05 | 0.71 | GCACCTGTCGCACTG |
| 1 | GMTACGGTCACACTG | chr3R:14121420-14123420 | 692 | 706 | + | 3.6357 | 4.09E-05 | 0.71 | GCTGCGGGCAAACGG |
| 1 | GMTACGGTCACACTG | chr2L:20413800-20415800 | 693 | 707 | + | 3.6357 | 4.09E-05 | 0.71 | GCGACGGGCGCACTC |
| 1 | GMTACGGTCACACTG | chr2L:7799147-7801147 | 702 | 716 | + | 3.6357 | 4.09E-05 | 0.71 | GCGTCGGTCACAGGG |
| 1 | GMTACGGTCACACTG | chr2R:13673633-13675633 | 691 | 705 | - | 3.6357 | 4.09E-05 | 0.71 | GCAACGGGCACAAGG |
| 1 | GMTACGGTCACACTG | chr3L:11457470-11459470 | 712 | 726 | + | 3.6357 | 4.09E-05 | 0.71 | GCACCTGTCGCACTG |
| 1 | GMTACGGTCACACTG | chrX:20417964-20419964 | 718 | 732 | + | 3.6357 | 4.09E-05 | 0.71 | GCCACGGACGCAGTG |
| 1 | GMTACGGTCACACTG | chr3L:11008188-11010188 | 726 | 740 | + | 3.6357 | 4.09E-05 | 0.71 | GCGACGGCAACACCG |
| 1 | GMTACGGTCACACTG | chr2R:17543190-17545190 | 714 | 728 | - | 3.6357 | 4.09E-05 | 0.71 | GCTGCGGACAAACGG |
| 1 | GMTACGGTCACACTG | chrX:1876317-1878317 | 715 | 729 | - | 3.6357 | 4.09E-05 | 0.71 | GCAATGGACACCCTG |
| 1 | GMTACGGTCACACTG | chr3L:210457-212457 | 730 | 744 | + | 3.6357 | 4.09E-05 | 0.71 | TCTTCGGTCTCGCTG |
| 1 | GMTACGGTCACACTG | chr3R:4189260-4191260 | 718 | 732 | - | 3.6357 | 4.09E-05 | 0.71 | GCTTCGGTCGCAGAG |
| 1 | GMTACGGTCACACTG | chrX:11416422-11418422 | 718 | 732 | - | 3.6357 | 4.09E-05 | 0.71 | TCGGCGGTCACGCTG |
| 1 | GMTACGGTCACACTG | chrX:11416422-11418422 | 718 | 732 | - | 3.6357 | 4.09E-05 | 0.71 | TCGGCGGTCACGCTG |
| 1 | GMTACGGTCACACTG | chr2L:4393196-4395196 | 725 | 739 | - | 3.6357 | 4.09E-05 | 0.71 | GCTACAGACTCACCG |
| 1 | GMTACGGTCACACTG | chr3L:1864240-1866240 | 727 | 741 | - | 3.6357 | 4.09E-05 | 0.71 | TCTACGGGCACTCCG |
| 1 | GMTACGGTCACACTG | chrX:20417964-20419964 | 728 | 742 | - | 3.6357 | 4.09E-05 | 0.71 | GCTGCAGACGCACTG |

| | | | | | | | | | |
|---|---|---|---|---|---|---|---|---|---|
| 1 | GMTACGGTCACACTG | chr3R:7966408-7968408 | 750 | 764 | + | 3.6357 | 4.09E-05 | 0.71 | GAGACGGGCCCACCG |
| 1 | GMTACGGTCACACTG | chr3R:19027857-19029857 | 756 | 770 | + | 3.6357 | 4.09E-05 | 0.71 | TCTTCGGTCACCCGG |
| 1 | GMTACGGTCACACTG | chr3R:26016470-26018470 | 745 | 759 | - | 3.6357 | 4.09E-05 | 0.71 | GCAAAGGTCGCACCG |
| 1 | GMTACGGTCACACTG | chr2R:11239114-11241114 | 763 | 777 | + | 3.6357 | 4.09E-05 | 0.71 | GCTTCTGGCGCACTG |
| 1 | GMTACGGTCACACTG | chr3R:9551034-9553034 | 752 | 766 | - | 3.6357 | 4.09E-05 | 0.71 | GCCGCGGCCACACTC |
| 1 | GMTACGGTCACACTG | chr3L:19909215-19911215 | 767 | 781 | + | 3.6357 | 4.09E-05 | 0.71 | GCGACGGTCGGACCG |
| 1 | GMTACGGTCACACTG | chr3R:23125422-23127422 | 756 | 770 | - | 3.6357 | 4.09E-05 | 0.71 | GCGACGGATACTCTG |
| 1 | GMTACGGTCACACTG | chrX:1876270-1878270 | 762 | 776 | - | 3.6357 | 4.09E-05 | 0.71 | GCAATGGACACCCTG |
| 1 | GMTACGGTCACACTG | chr2R:11348870-11350870 | 777 | 791 | + | 3.6357 | 4.09E-05 | 0.71 | GCGGCGGTCACGATG |
| 1 | GMTACGGTCACACTG | chr2R:11353373-11355373 | 778 | 792 | + | 3.6357 | 4.09E-05 | 0.71 | GCGGCGGTCACGATG |
| 1 | GMTACGGTCACACTG | chrX:2028094-2030094 | 764 | 778 | - | 3.6357 | 4.09E-05 | 0.71 | GCTCCGCTCGCTCTG |
| 1 | GMTACGGTCACACTG | chr2R:13128052-13130052 | 780 | 794 | + | 3.6357 | 4.09E-05 | 0.71 | GCCAAGGTCGCCCTG |
| 1 | GMTACGGTCACACTG | chr3L:4239173-4241173 | 785 | 799 | + | 3.6357 | 4.09E-05 | 0.71 | GCCACGTACACTCTG |
| 1 | GMTACGGTCACACTG | chr2L:13372890-13374890 | 772 | 786 | - | 3.6357 | 4.09E-05 | 0.71 | GCTCCGGCCAACCTG |
| 1 | GMTACGGTCACACTG | chr2R:9838500-9840500 | 786 | 800 | + | 3.6357 | 4.09E-05 | 0.71 | GCATCGGTCACCCTA |
| 1 | GMTACGGTCACACTG | chrX:20256199-20258199 | 790 | 804 | + | 3.6357 | 4.09E-05 | 0.71 | GCTCCGGAGACTCTG |
| 1 | GMTACGGTCACACTG | chr2L:7799058-7801058 | 791 | 805 | + | 3.6357 | 4.09E-05 | 0.71 | GCGTCGGTCACAGGG |
| 1 | GMTACGGTCACACTG | chrX:15440252-15442252 | 778 | 792 | - | 3.6357 | 4.09E-05 | 0.71 | GCAACGAACACTCTG |
| 1 | GMTACGGTCACACTG | chr3L:16104627-16106627 | 782 | 796 | - | 3.6357 | 4.09E-05 | 0.71 | ACTGCGGTCTCACGG |
| 1 | GMTACGGTCACACTG | chr2R:17664068-17666068 | 799 | 813 | + | 3.6357 | 4.09E-05 | 0.71 | GCCACGCACACACCG |
| 1 | GMTACGGTCACACTG | chr3L:16280930-16282930 | 790 | 804 | - | 3.6357 | 4.09E-05 | 0.71 | GCTCCGGCTCCACTG |
| 1 | GMTACGGTCACACTG | chr3L:7975154-7977154 | 816 | 830 | + | 3.6357 | 4.09E-05 | 0.71 | GCTCCGTTCGCCCTG |
| 1 | GMTACGGTCACACTG | chr3R:23042990-23044990 | 807 | 821 | - | 3.6357 | 4.09E-05 | 0.71 | GCCACGCCCACTCTG |
| 1 | GMTACGGTCACACTG | chrX:10255629-10257629 | 830 | 844 | + | 3.6357 | 4.09E-05 | 0.71 | GCGGTGGGCACACTG |
| 1 | GMTACGGTCACACTG | chr2R:12517912-12519912 | 843 | 857 | + | 3.6357 | 4.09E-05 | 0.71 | GCTCCGGCCACCGTG |
| 1 | GMTACGGTCACACTG | chrX:7880679-7882679 | 845 | 859 | + | 3.6357 | 4.09E-05 | 0.71 | GCTCCGGCAACACAG |
| 1 | GMTACGGTCACACTG | chr3L:16268392-16270392 | 832 | 846 | - | 3.6357 | 4.09E-05 | 0.71 | GCTACGTTCCCGCCG |
| 1 | GMTACGGTCACACTG | chr2R:9094372-9096372 | 849 | 863 | + | 3.6357 | 4.09E-05 | 0.71 | GCAGCGTGCACACTG |
| 1 | GMTACGGTCACACTG | chrX:13717285-13719285 | 854 | 868 | + | 3.6357 | 4.09E-05 | 0.71 | ACTACGGTCGCTCCG |
| 1 | GMTACGGTCACACTG | chr3R:29856491-29858491 | 862 | 876 | + | 3.6357 | 4.09E-05 | 0.71 | GAAGCGGGCGCACTG |
| 1 | GMTACGGTCACACTG | chr2L:16714891-16716891 | 849 | 863 | - | 3.6357 | 4.09E-05 | 0.71 | GCTACCGACACCCAG |
| 1 | GMTACGGTCACACTG | chr2R:19783294-19785294 | 863 | 877 | + | 3.6357 | 4.09E-05 | 0.71 | GCATTGGTCACGCTG |
| 1 | GMTACGGTCACACTG | chr2L:3769454-3771454 | 863 | 877 | + | 3.6357 | 4.09E-05 | 0.71 | GCTACGGACTCTCTT |

| | | | | | | | | | |
|---|---|---|---|---|---|---|---|---|---|
| 1 | GMTACGGTCACACTG | chrX:6012708-6014708 | 864 | 878 | + | 3.6357 | 4.09E-05 | 0.71 | GCCTCGCTCACACAG |
| 1 | GMTACGGTCACACTG | chr3L:21961401-21963401 | 851 | 865 | - | 3.6357 | 4.09E-05 | 0.71 | GCCACAGACCCACTG |
| 1 | GMTACGGTCACACTG | chr2L:1882214-1884214 | 869 | 883 | + | 3.6357 | 4.09E-05 | 0.71 | GCAGCAGGCACACTG |
| 1 | GMTACGGTCACACTG | chr2R:10246560-10248560 | 857 | 871 | - | 3.6357 | 4.09E-05 | 0.71 | GCCACGCCCACACCG |
| 1 | GMTACGGTCACACTG | chr2L:16548959-16550959 | 862 | 876 | - | 3.6357 | 4.09E-05 | 0.71 | GCGCCGGCGACACTG |
| 1 | GMTACGGTCACACTG | chr3R:27063199-27065199 | 880 | 894 | + | 3.6357 | 4.09E-05 | 0.71 | GCTAGGGTCGCTCCG |
| 1 | GMTACGGTCACACTG | chr2L:5622762-5624762 | 866 | 880 | - | 3.6357 | 4.09E-05 | 0.71 | GCTGCGGCCGCACTT |
| 1 | GMTACGGTCACACTG | chr2L:4462611-4464611 | 869 | 883 | - | 3.6357 | 4.09E-05 | 0.71 | GCAAGGGTCGCCCTG |
| 1 | GMTACGGTCACACTG | chr3R:30201998-30203998 | 887 | 901 | + | 3.6357 | 4.09E-05 | 0.71 | GATACGGACTCGCCG |
| 1 | GMTACGGTCACACTG | chrX:764288-766288 | 875 | 889 | - | 3.6357 | 4.09E-05 | 0.71 | GCTGCGGCCACACCA |
| 1 | GMTACGGTCACACTG | chr2L:8824645-8826645 | 890 | 904 | + | 3.6357 | 4.09E-05 | 0.71 | GCGACGCACACACAG |
| 1 | GMTACGGTCACACTG | chr2R:8018176-8020176 | 878 | 892 | - | 3.6357 | 4.09E-05 | 0.71 | GCCGCGCTCACACAG |
| 1 | GMTACGGTCACACTG | chr3L:9891109-9893109 | 893 | 907 | + | 3.6357 | 4.09E-05 | 0.71 | GCTCCGGTCGCCTTG |
| 1 | GMTACGGTCACACTG | chr2L:20561298-20563298 | 898 | 912 | + | 3.6357 | 4.09E-05 | 0.71 | GCTACGGCTCCGCTG |
| 1 | GMTACGGTCACACTG | chr2R:24985070-24987070 | 884 | 898 | - | 3.6357 | 4.09E-05 | 0.71 | GCTGCCGCCGCACTG |
| 1 | GMTACGGTCACACTG | chr3R:18089803-18091803 | 885 | 899 | - | 3.6357 | 4.09E-05 | 0.71 | GCCTCGGTTTCACTG |
| 1 | GMTACGGTCACACTG | chrX:13089434-13091434 | 899 | 913 | + | 3.6357 | 4.09E-05 | 0.71 | GCGGCGGTATCACTG |
| 1 | GMTACGGTCACACTG | chrX:10054109-10056109 | 886 | 900 | - | 3.6357 | 4.09E-05 | 0.71 | GCTCCGGGAACACCG |
| 1 | GMTACGGTCACACTG | chr3R:29860582-29862582 | 888 | 902 | - | 3.6357 | 4.09E-05 | 0.71 | GCTTCGCGCACACGG |
| 1 | GMTACGGTCACACTG | chr3R:19863496-19865496 | 903 | 917 | + | 3.6357 | 4.09E-05 | 0.71 | GCACCGGTCAAACGG |
| 1 | GMTACGGTCACACTG | chr3L:4141364-4143364 | 895 | 909 | - | 3.6357 | 4.09E-05 | 0.71 | GCCACTGACACACCG |
| 1 | GMTACGGTCACACTG | chr2L:20561298-20563298 | 912 | 926 | + | 3.6357 | 4.09E-05 | 0.71 | GCTACGGCTTCGCTG |
| 1 | GMTACGGTCACACTG | chr2R:14214864-14216864 | 918 | 932 | + | 3.6357 | 4.09E-05 | 0.71 | GCTGCTGGCACGCTG |
| 1 | GMTACGGTCACACTG | chr2L:277234-279234 | 904 | 918 | - | 3.6357 | 4.09E-05 | 0.71 | ACGTCGGACACACTG |
| 1 | GMTACGGTCACACTG | chr2R:24985044-24987044 | 910 | 924 | - | 3.6357 | 4.09E-05 | 0.71 | GCTGCCGCCGCACTG |
| 1 | GMTACGGTCACACTG | chr2R:18245763-18247763 | 926 | 940 | + | 3.6357 | 4.09E-05 | 0.71 | GCCCCGACCACACTG |
| 1 | GMTACGGTCACACTG | chr2L:20437701-20439701 | 913 | 927 | - | 3.6357 | 4.09E-05 | 0.71 | GCAACGGACACACAC |
| 1 | GMTACGGTCACACTG | chr2R:10785026-10787026 | 931 | 945 | + | 3.6357 | 4.09E-05 | 0.71 | GCTCCGGCCTCAGTG |
| 1 | GMTACGGTCACACTG | chr2R:7386187-7388187 | 919 | 933 | - | 3.6357 | 4.09E-05 | 0.71 | GCGTCGGCCACACTA |
| 1 | GMTACGGTCACACTG | chr3L:16482650-16484650 | 937 | 951 | + | 3.6357 | 4.09E-05 | 0.71 | GCCTCGGTTACTCTG |
| 1 | GMTACGGTCACACTG | chr2R:22701695-22703695 | 938 | 952 | + | 3.6357 | 4.09E-05 | 0.71 | GCGTCAGTCACTCTG |
| 1 | GMTACGGTCACACTG | chr3R:31804468-31806468 | 927 | 941 | - | 3.6357 | 4.09E-05 | 0.71 | GCCTTGGTCCCACTG |
| 1 | GMTACGGTCACACTG | chr3L:18592771-18594771 | 944 | 958 | + | 3.6357 | 4.09E-05 | 0.71 | GCTCCGGAAGCACTG |

| | | | | | | | | | |
|---|---|---|---|---|---|---|---|---|---|
| 1 | GMTACGGTCACACTG | chr2R:17053002-17055002 | 930 | 944 | - | 3.6357 | 4.09E-05 | 0.71 | GCTTCGGGAACTCTG |
| 1 | GMTACGGTCACACTG | chr2R:9094273-9096273 | 948 | 962 | + | 3.6357 | 4.09E-05 | 0.71 | GCAGCGTGCACACTG |
| 1 | GMTACGGTCACACTG | chr2L:20561247-20563247 | 949 | 963 | + | 3.6357 | 4.09E-05 | 0.71 | GCTACGGCTCCGCTG |
| 1 | GMTACGGTCACACTG | chr2L:16718932-16720932 | 952 | 966 | + | 3.6357 | 4.09E-05 | 0.71 | CCCCCGGTCACTCTG |
| 1 | GMTACGGTCACACTG | chr2R:12883632-12885632 | 954 | 968 | + | 3.6357 | 4.09E-05 | 0.71 | GCTGCCGTCGCGCTG |
| 1 | GMTACGGTCACACTG | chr2R:20776749-20778749 | 954 | 968 | + | 3.6357 | 4.09E-05 | 0.71 | GCTGCGGCCAGACAG |
| 1 | GMTACGGTCACACTG | chr2R:6806626-6808626 | 940 | 954 | - | 3.6357 | 4.09E-05 | 0.71 | GCACTGGTCACACAG |
| 1 | GMTACGGTCACACTG | chr3R:24289172-24291172 | 948 | 962 | - | 3.6357 | 4.09E-05 | 0.71 | GCCACGGTTACTCCG |
| 1 | GMTACGGTCACACTG | chr2L:20561247-20563247 | 963 | 977 | + | 3.6357 | 4.09E-05 | 0.71 | GCTACGGCTTCGCTG |
| 1 | GMTACGGTCACACTG | chr3L:10126635-10128635 | 965 | 979 | + | 3.6357 | 4.09E-05 | 0.71 | GCTCCGGTCCCACAC |
| 1 | GMTACGGTCACACTG | chr2R:21615907-21617907 | 956 | 970 | - | 3.6357 | 4.09E-05 | 0.71 | GCGGCGGTCACACCA |
| 1 | GMTACGGTCACACTG | chr3L:4379387-4381387 | 972 | 986 | + | 3.6357 | 4.09E-05 | 0.71 | GCGACGCGCTCACTG |
| 1 | GMTACGGTCACACTG | chrX:17855129-17857129 | 962 | 976 | - | 3.6357 | 4.09E-05 | 0.71 | GCCACGGCCACAAAG |
| 1 | GMTACGGTCACACTG | chrX:1163105-1165105 | 966 | 980 | - | 3.6357 | 4.09E-05 | 0.71 | GCCGCGGTGACTCTG |
| 1 | GMTACGGTCACACTG | chr2L:6337836-6339836 | 981 | 995 | + | 3.6357 | 4.09E-05 | 0.71 | ACGACGGTCCCTCTG |
| 1 | GMTACGGTCACACTG | chr3R:9808368-9810368 | 970 | 984 | - | 3.6357 | 4.09E-05 | 0.71 | GCAGCGGTCACTCTC |
| 1 | GMTACGGTCACACTG | chr3R:13258517-13260517 | 973 | 987 | - | 3.6357 | 4.09E-05 | 0.71 | TCGTCGGTCACGCTG |
| 1 | GMTACGGTCACACTG | chr2L:4226699-4228699 | 988 | 1002 | + | 3.6357 | 4.09E-05 | 0.71 | GCGGCGGCCACACTC |
| 1 | GMTACGGTCACACTG | chr2R:10074645-10076645 | 975 | 989 | - | 3.6357 | 4.09E-05 | 0.71 | GCCCTGGTCACTCTG |
| 1 | GMTACGGTCACACTG | chr3R:29860494-29862494 | 976 | 990 | - | 3.6357 | 4.09E-05 | 0.71 | GCTTCGCGCACACGG |
| 1 | GMTACGGTCACACTG | chr3R:27735538-27737538 | 976 | 990 | - | 3.6357 | 4.09E-05 | 0.71 | GCTGCGATCGCTCTG |
| 1 | GMTACGGTCACACTG | chr3L:17620824-17622824 | 991 | 1005 | + | 3.6357 | 4.09E-05 | 0.71 | GCTGCGGTCCCACCA |
| 1 | GMTACGGTCACACTG | chr2R:9692328-9694328 | 977 | 991 | - | 3.6357 | 4.09E-05 | 0.71 | GCCGCGGCCACACTA |
| 1 | GMTACGGTCACACTG | chrX:10324958-10326958 | 978 | 992 | - | 3.6357 | 4.09E-05 | 0.71 | GCGGCGGGCACATTG |
| 1 | GMTACGGTCACACTG | chr3R:28598288-28600288 | 979 | 993 | - | 3.6357 | 4.09E-05 | 0.71 | GCAGCAGCCACACTG |
| 1 | GMTACGGTCACACTG | chr2R:22997884-22999884 | 980 | 994 | - | 3.6357 | 4.09E-05 | 0.71 | GCTTCTGTCACTCGG |
| 1 | GMTACGGTCACACTG | chr3R:9550806-9552806 | 980 | 994 | - | 3.6357 | 4.09E-05 | 0.71 | GCCGCGGCCACACTC |
| 1 | GMTACGGTCACACTG | chr3R:9550806-9552806 | 980 | 994 | - | 3.6357 | 4.09E-05 | 0.71 | GCCGCGGCCACACTC |
| 1 | GMTACGGTCACACTG | chr3R:22582225-22584225 | 990 | 1004 | - | 3.6357 | 4.09E-05 | 0.71 | GCGGCGGTCACACAA |
| 1 | GMTACGGTCACACTG | chr3L:14107118-14109118 | 991 | 1005 | - | 3.6357 | 4.09E-05 | 0.71 | GAAACGGACGCGCTG |
| 1 | GMTACGGTCACACTG | chr2R:7386115-7388115 | 991 | 1005 | - | 3.6357 | 4.09E-05 | 0.71 | GCGTCGGCCACACTA |
| 1 | GMTACGGTCACACTG | chr3R:19863393-19865393 | 1006 | 1020 | + | 3.6357 | 4.09E-05 | 0.71 | GCACCGGTCAAACGG |
| 1 | GMTACGGTCACACTG | chr3L:21639995-21641995 | 1006 | 1020 | + | 3.6357 | 4.09E-05 | 0.71 | CCCTCGGTCACTCTG |

| | | | | | | | | | |
|---|---|---|---|---|---|---|---|---|---|
| 1 | GMTACGGTCACACTG | chr3R:11763457-11765457 | 992 | 1006 | - | 3.6357 | 4.09E-05 | 0.71 | CCTCCGGCCACACGG |
| 1 | GMTACGGTCACACTG | chr2L:16286730-16288730 | 993 | 1007 | - | 3.6357 | 4.09E-05 | 0.71 | GCCGTGGTCACTCTG |
| 1 | GMTACGGTCACACTG | chrX:20392751-20394751 | 993 | 1007 | - | 3.6357 | 4.09E-05 | 0.71 | TCAGCGGGCACACTG |
| 1 | GMTACGGTCACACTG | chr3R:27947021-27949021 | 994 | 1008 | - | 3.6357 | 4.09E-05 | 0.71 | GCGACGTTCCCGCTG |
| 1 | GMTACGGTCACACTG | chr2R:11378177-11380177 | 1008 | 1022 | + | 3.6357 | 4.09E-05 | 0.71 | GCGACGGTCCGGCTG |
| 1 | GMTACGGTCACACTG | chr2R:15537175-15539175 | 1009 | 1023 | + | 3.6357 | 4.09E-05 | 0.71 | CCAACGGACCCACTG |
| 1 | GMTACGGTCACACTG | chr2L:11812780-11814780 | 998 | 1012 | - | 3.6357 | 4.09E-05 | 0.71 | TCAACGGGCACACCG |
| 1 | GMTACGGTCACACTG | chr3L:1869019-1871019 | 998 | 1012 | - | 3.6357 | 4.09E-05 | 0.71 | GCTGCGGGCACATCG |
| 1 | GMTACGGTCACACTG | chr3L:8423954-8425954 | 1014 | 1028 | + | 3.6357 | 4.09E-05 | 0.71 | GCCGCGGTCATACCG |
| 1 | GMTACGGTCACACTG | chr3R:7548124-7550124 | 1001 | 1015 | - | 3.6357 | 4.09E-05 | 0.71 | GCTGCGGGCTTACTG |
| 1 | GMTACGGTCACACTG | chr3L:18674413-18676413 | 1002 | 1016 | - | 3.6357 | 4.09E-05 | 0.71 | GCAACAGTCCCACCG |
| 1 | GMTACGGTCACACTG | chr3L:19792569-19794569 | 1006 | 1020 | - | 3.6357 | 4.09E-05 | 0.71 | GCACTGGCCACACTG |
| 1 | GMTACGGTCACACTG | chr3R:31310157-31312157 | 1006 | 1020 | - | 3.6357 | 4.09E-05 | 0.71 | GCGACGGTCCAACAG |
| 1 | GMTACGGTCACACTG | chr3L:19792569-19794569 | 1006 | 1020 | - | 3.6357 | 4.09E-05 | 0.71 | GCACTGGCCACACTG |
| 1 | GMTACGGTCACACTG | chr2R:18626918-18628918 | 1020 | 1034 | + | 3.6357 | 4.09E-05 | 0.71 | GCGACGGTCTCTCTC |
| 1 | GMTACGGTCACACTG | chr3R:24140395-24142395 | 1022 | 1036 | + | 3.6357 | 4.09E-05 | 0.71 | GCAACGGTCGCACGA |
| 1 | GMTACGGTCACACTG | chrX:2587503-2589503 | 1023 | 1037 | + | 3.6357 | 4.09E-05 | 0.71 | GCGACGGTCACTTGG |
| 1 | GMTACGGTCACACTG | chr2R:20013495-20015495 | 1010 | 1024 | - | 3.6357 | 4.09E-05 | 0.71 | GCTGCGGTCTAGCTG |
| 1 | GMTACGGTCACACTG | chr3R:31595204-31597204 | 1012 | 1026 | - | 3.6357 | 4.09E-05 | 0.71 | GCCCTGGTCACACCG |
| 1 | GMTACGGTCACACTG | chrX:6862888-6864888 | 1014 | 1028 | - | 3.6357 | 4.09E-05 | 0.71 | GCTTCGATCTCTCTG |
| 1 | GMTACGGTCACACTG | chrX:1408069-1410069 | 1035 | 1049 | + | 3.6357 | 4.09E-05 | 0.71 | GCTGCGGCGCCACTG |
| 1 | GMTACGGTCACACTG | chrX:12431224-12433224 | 1023 | 1037 | - | 3.6357 | 4.09E-05 | 0.71 | GCTGCGTTCGCACGG |
| 1 | GMTACGGTCACACTG | chrX:16730091-16732091 | 1023 | 1037 | - | 3.6357 | 4.09E-05 | 0.71 | GCAACTGTCGCCCTG |
| 1 | GMTACGGTCACACTG | chr3R:30151059-30153059 | 1045 | 1059 | + | 3.6357 | 4.09E-05 | 0.71 | GCGGCGGTCTCAGTG |
| 1 | GMTACGGTCACACTG | chr3R:30212797-30214797 | 1046 | 1060 | + | 3.6357 | 4.09E-05 | 0.71 | GCTACGGCTGCACAG |
| 1 | GMTACGGTCACACTG | chrX:764130-766130 | 1033 | 1047 | - | 3.6357 | 4.09E-05 | 0.71 | GCTGCGGCCACACCA |
| 1 | GMTACGGTCACACTG | chr2L:4462447-4464447 | 1033 | 1047 | - | 3.6357 | 4.09E-05 | 0.71 | GCAAGGGTCGCCCTG |
| 1 | GMTACGGTCACACTG | chr2R:22200508-22202508 | 1047 | 1061 | + | 3.6357 | 4.09E-05 | 0.71 | GCTTCAGTCTCACCG |
| 1 | GMTACGGTCACACTG | chr2L:16487148-16489148 | 1036 | 1050 | - | 3.6357 | 4.09E-05 | 0.71 | GCTGCGAGCGCACTG |
| 1 | GMTACGGTCACACTG | chr2L:4392876-4394876 | 1045 | 1059 | - | 3.6357 | 4.09E-05 | 0.71 | GCTACAGACTCACCG |
| 1 | GMTACGGTCACACTG | chr2R:5081431-5083431 | 1066 | 1080 | + | 3.6357 | 4.09E-05 | 0.71 | GCCACTGGCACCCTG |
| 1 | GMTACGGTCACACTG | chr2R:15481834-15483834 | 1052 | 1066 | - | 3.6357 | 4.09E-05 | 0.71 | GCTTGGGTCTCTCTG |
| 1 | GMTACGGTCACACTG | chr3R:25244109-25246109 | 1053 | 1067 | - | 3.6357 | 4.09E-05 | 0.71 | GCGACGGCAACCCTG |

| | | | | | | | | | |
|---|---|---|---|---|---|---|---|---|---|
| 1 | GMTACGGTCACACTG | chr3R:7782446-7784446 | 1055 | 1069 | - | 3.6357 | 4.09E-05 | 0.71 | GACGCGGTCGCACCG |
| 1 | GMTACGGTCACACTG | chr2L:19569264-19571264 | 1069 | 1083 | + | 3.6357 | 4.09E-05 | 0.71 | GCAACGTTCTCACGG |
| 1 | GMTACGGTCACACTG | chr2R:7465759-7467759 | 1070 | 1084 | + | 3.6357 | 4.09E-05 | 0.71 | GCGCTGGTCACACCG |
| 1 | GMTACGGTCACACTG | chr3L:1567608-1569608 | 1073 | 1087 | + | 3.6357 | 4.09E-05 | 0.71 | ACCACGGGCACCCTG |
| 1 | GMTACGGTCACACTG | chr3R:12711299-12713299 | 1076 | 1090 | + | 3.6357 | 4.09E-05 | 0.71 | CCAGCGGTCCCACTG |
| 1 | GMTACGGTCACACTG | chrX:6997672-6999672 | 1068 | 1082 | - | 3.6357 | 4.09E-05 | 0.71 | GCGACGGTTACCCAG |
| 1 | GMTACGGTCACACTG | chr3L:6081883-6083883 | 1083 | 1097 | + | 3.6357 | 4.09E-05 | 0.71 | GCGACGGTCGCAGCG |
| 1 | GMTACGGTCACACTG | chr3L:11424158-11426158 | 1083 | 1097 | - | 3.6357 | 4.09E-05 | 0.71 | GCAAAGGTCACGCAG |
| 1 | GMTACGGTCACACTG | chr3L:9797420-9799420 | 1098 | 1112 | + | 3.6357 | 4.09E-05 | 0.71 | GCAATGGTCACGCCG |
| 1 | GMTACGGTCACACTG | chr2L:6037117-6039117 | 1103 | 1117 | + | 3.6357 | 4.09E-05 | 0.71 | GCCGCGGACACAATG |
| 1 | GMTACGGTCACACTG | chr2L:4800923-4802923 | 1103 | 1117 | + | 3.6357 | 4.09E-05 | 0.71 | GCGACGGCGACGCTG |
| 1 | GMTACGGTCACACTG | chr2R:24637588-24639588 | 1112 | 1126 | + | 3.6357 | 4.09E-05 | 0.71 | GCCACTGGCACACGG |
| 1 | GMTACGGTCACACTG | chr3L:11681038-11683038 | 1113 | 1127 | + | 3.6357 | 4.09E-05 | 0.71 | TCCACGGACCCACTG |
| 1 | GMTACGGTCACACTG | chr2R:19783043-19785043 | 1114 | 1128 | + | 3.6357 | 4.09E-05 | 0.71 | GCATTGGTCACGCTG |
| 1 | GMTACGGTCACACTG | chr3R:27105184-27107184 | 1119 | 1133 | + | 3.6357 | 4.09E-05 | 0.71 | GCACCGCTCACACAG |
| 1 | GMTACGGTCACACTG | chr2L:15922395-15924395 | 1105 | 1119 | - | 3.6357 | 4.09E-05 | 0.71 | GCAACGGTTTCGCTG |
| 1 | GMTACGGTCACACTG | chr3R:31804278-31806278 | 1117 | 1131 | - | 3.6357 | 4.09E-05 | 0.71 | GCCTTGGTCCCACTG |
| 1 | GMTACGGTCACACTG | chr2L:4935450-4937450 | 1139 | 1153 | + | 3.6357 | 4.09E-05 | 0.71 | CCTTCGGTCCCACGG |
| 1 | GMTACGGTCACACTG | chr3R:11430738-11432738 | 1145 | 1159 | + | 3.6357 | 4.09E-05 | 0.71 | GCGCTGGTCACACCG |
| 1 | GMTACGGTCACACTG | chr2L:6337671-6339671 | 1146 | 1160 | + | 3.6357 | 4.09E-05 | 0.71 | ACGACGGTCCCTCTG |
| 1 | GMTACGGTCACACTG | chr2R:6890760-6892760 | 1138 | 1152 | - | 3.6357 | 4.09E-05 | 0.71 | GCCACAGACACTCTG |
| 1 | GMTACGGTCACACTG | chr3R:30725087-30727087 | 1141 | 1155 | - | 3.6357 | 4.09E-05 | 0.71 | ACCACGGACACACGG |
| 1 | GMTACGGTCACACTG | chr3L:16327841-16329841 | 1158 | 1172 | + | 3.6357 | 4.09E-05 | 0.71 | GCCACAGTCACGCAG |
| 1 | GMTACGGTCACACTG | chr2R:24637588-24639588 | 1145 | 1159 | - | 3.6357 | 4.09E-05 | 0.71 | GATGCGGTCTCTCCG |
| 1 | GMTACGGTCACACTG | chr2L:10430338-10432338 | 1162 | 1176 | + | 3.6357 | 4.09E-05 | 0.71 | GCTACGCACACCCCG |
| 1 | GMTACGGTCACACTG | chr3R:12994620-12996620 | 1151 | 1165 | - | 3.6357 | 4.09E-05 | 0.71 | GCGCCGGTCAAGCTG |
| 1 | GMTACGGTCACACTG | chrX:19642327-19644327 | 1166 | 1180 | + | 3.6357 | 4.09E-05 | 0.71 | GCTGCGCTCGCACAG |
| 1 | GMTACGGTCACACTG | chr2L:5257942-5259942 | 1166 | 1180 | + | 3.6357 | 4.09E-05 | 0.71 | GCTGCGGCCACAGCG |
| 1 | GMTACGGTCACACTG | chr2R:14238932-14240932 | 1168 | 1182 | + | 3.6357 | 4.09E-05 | 0.71 | GCCAGGGCCACACGG |
| 1 | GMTACGGTCACACTG | chr3L:12490840-12492840 | 1154 | 1168 | - | 3.6357 | 4.09E-05 | 0.71 | ACTACGGTCTCGCGG |
| 1 | GMTACGGTCACACTG | chr2R:22070286-22072286 | 1173 | 1187 | + | 3.6357 | 4.09E-05 | 0.71 | GCATAGGTCACTCTG |
| 1 | GMTACGGTCACACTG | chr2R:16116109-16118109 | 1165 | 1179 | - | 3.6357 | 4.09E-05 | 0.71 | GCAGCGGTCCAACTG |
| 1 | GMTACGGTCACACTG | chr2R:22074988-22076988 | 1165 | 1179 | - | 3.6357 | 4.09E-05 | 0.71 | GCTGCGGTTGCCCTG |

| | | | | | | | | | |
|---|---|---|---|---|---|---|---|---|---|
| 1 | GMTACGGTCACACTG | chr3R:16222395-16224395 | 1172 | 1186 | - | 3.6357 | 4.09E-05 | 0.71 | GCAACGGCCACACCA |
| 1 | GMTACGGTCACACTG | chr3L:17620625-17622625 | 1190 | 1204 | + | 3.6357 | 4.09E-05 | 0.71 | GCTGCGGTCCCACCA |
| 1 | GMTACGGTCACACTG | chr3R:30629387-30631387 | 1177 | 1191 | - | 3.6357 | 4.09E-05 | 0.71 | CCTTCGGACTCACTG |
| 1 | GMTACGGTCACACTG | chrX:5780620-5782620 | 1178 | 1192 | - | 3.6357 | 4.09E-05 | 0.71 | GCTACGGGCACTTGG |
| 1 | GMTACGGTCACACTG | chr3R:31498235-31500235 | 1179 | 1193 | - | 3.6357 | 4.09E-05 | 0.71 | GCGACGTCCTCACTG |
| 1 | GMTACGGTCACACTG | chrX:5696123-5698123 | 1191 | 1205 | - | 3.6357 | 4.09E-05 | 0.71 | GCCAAGGACACACCG |
| 1 | GMTACGGTCACACTG | chr3L:16841294-16843294 | 1205 | 1219 | + | 3.6357 | 4.09E-05 | 0.71 | GCGTTGGCCACACTG |
| 1 | GMTACGGTCACACTG | chr2L:3589301-3591301 | 1208 | 1222 | + | 3.6357 | 4.09E-05 | 0.71 | GCAGCGGTCGCAGTG |
| 1 | GMTACGGTCACACTG | chrX:14168592-14170592 | 1211 | 1225 | + | 3.6357 | 4.09E-05 | 0.71 | GCTATGGACTCACCG |
| 1 | GMTACGGTCACACTG | chrX:6294389-6296389 | 1198 | 1212 | - | 3.6357 | 4.09E-05 | 0.71 | GCAGTGGACACACTG |
| 1 | GMTACGGTCACACTG | chr2L:4387327-4389327 | 1212 | 1226 | + | 3.6357 | 4.09E-05 | 0.71 | GCAACGGTCGCTCTC |
| 1 | GMTACGGTCACACTG | chr3R:17087280-17089280 | 1201 | 1215 | - | 3.6357 | 4.09E-05 | 0.71 | TCTACGGACTCACCG |
| 1 | GMTACGGTCACACTG | chr2R:22612376-22614376 | 1222 | 1236 | + | 3.6357 | 4.09E-05 | 0.71 | GCAACAGCCACACCG |
| 1 | GMTACGGTCACACTG | chr2R:11354319-11356319 | 1208 | 1222 | - | 3.6357 | 4.09E-05 | 0.71 | GCGGCGGTCACGATG |
| 1 | GMTACGGTCACACTG | chr2R:11349811-11351811 | 1210 | 1224 | - | 3.6357 | 4.09E-05 | 0.71 | GCGGCGGTCACGATG |
| 1 | GMTACGGTCACACTG | chr2R:13433207-13435207 | 1211 | 1225 | - | 3.6357 | 4.09E-05 | 0.71 | GCGGCGGTCACGATG |
| 1 | GMTACGGTCACACTG | chrX:15577205-15579205 | 1228 | 1242 | + | 3.6357 | 4.09E-05 | 0.71 | GATGCGGACGCACGG |
| 1 | GMTACGGTCACACTG | chr3R:9005456-9007456 | 1217 | 1231 | - | 3.6357 | 4.09E-05 | 0.71 | GCAACGGCTCCACTG |
| 1 | GMTACGGTCACACTG | chr2R:9765348-9767348 | 1237 | 1251 | + | 3.6357 | 4.09E-05 | 0.71 | GCTTCAGTCGCACGG |
| 1 | GMTACGGTCACACTG | chr2R:10546260-10548260 | 1238 | 1252 | + | 3.6357 | 4.09E-05 | 0.71 | GCTGCGATCTCACAG |
| 1 | GMTACGGTCACACTG | chr2L:2191386-2193386 | 1243 | 1257 | + | 3.6357 | 4.09E-05 | 0.71 | GCCAAGGCCACACAG |
| 1 | GMTACGGTCACACTG | chr3R:14656447-14658447 | 1232 | 1246 | - | 3.6357 | 4.09E-05 | 0.71 | GACACGGCCACTCGG |
| 1 | GMTACGGTCACACTG | chr3L:13556755-13558755 | 1253 | 1267 | + | 3.6357 | 4.09E-05 | 0.71 | GCAACGGACACCGTG |
| 1 | GMTACGGTCACACTG | chrX:6294345-6296345 | 1242 | 1256 | - | 3.6357 | 4.09E-05 | 0.71 | GCAGTGGACACACTG |
| 1 | GMTACGGTCACACTG | chr2R:22666155-22668155 | 1243 | 1257 | - | 3.6357 | 4.09E-05 | 0.71 | GATCCGGACCCACCG |
| 1 | GMTACGGTCACACTG | chr2R:22666155-22668155 | 1243 | 1257 | - | 3.6357 | 4.09E-05 | 0.71 | GATCCGGACCCACCG |
| 1 | GMTACGGTCACACTG | chrX:15577171-15579171 | 1262 | 1276 | + | 3.6357 | 4.09E-05 | 0.71 | GATGCGGACGCACGG |
| 1 | GMTACGGTCACACTG | chrX:8156628-8158628 | 1265 | 1279 | + | 3.6357 | 4.09E-05 | 0.71 | GCCACAGCCACACGG |
| 1 | GMTACGGTCACACTG | chrX:13954236-13956236 | 1257 | 1271 | - | 3.6357 | 4.09E-05 | 0.71 | GAGGCGGGCACGCTG |
| 1 | GMTACGGTCACACTG | chr3L:6369591-6371591 | 1261 | 1275 | - | 3.6357 | 4.09E-05 | 0.71 | GCATCGGACAAACTG |
| 1 | GMTACGGTCACACTG | chr3L:11294919-11296919 | 1265 | 1279 | - | 3.6357 | 4.09E-05 | 0.71 | GCAGCCGCCACACTG |
| 1 | GMTACGGTCACACTG | chrX:19049442-19051442 | 1281 | 1295 | + | 3.6357 | 4.09E-05 | 0.71 | GCGGCTGTCTCACTG |
| 1 | GMTACGGTCACACTG | chr2R:23345635-23347635 | 1288 | 1302 | + | 3.6357 | 4.09E-05 | 0.71 | GCGGCGGTCACCATG |

| | | | | | | | | | |
|---|---|---|---|---|---|---|---|---|---|
| 1 | GMTACGGTCACACTG | chr2L:4829039-4831039 | 1276 | 1290 | - | 3.6357 | 4.09E-05 | 0.71 | ACCACGGACGCACTG |
| 1 | GMTACGGTCACACTG | chr3R:21753324-21755324 | 1280 | 1294 | - | 3.6357 | 4.09E-05 | 0.71 | GCCAAGGACACACAG |
| 1 | GMTACGGTCACACTG | chr2R:12167731-12169731 | 1297 | 1311 | + | 3.6357 | 4.09E-05 | 0.71 | GAGACGGACGCACCG |
| 1 | GMTACGGTCACACTG | chr3R:11430582-11432582 | 1301 | 1315 | + | 3.6357 | 4.09E-05 | 0.71 | GCGCTGGTCACACCG |
| 1 | GMTACGGTCACACTG | chr3R:16224651-16226651 | 1309 | 1323 | + | 3.6357 | 4.09E-05 | 0.71 | GCGACGGCAACCCTG |
| 1 | GMTACGGTCACACTG | chr3R:12425578-12427578 | 1312 | 1326 | + | 3.6357 | 4.09E-05 | 0.71 | GCTGCAGGCCCACTG |
| 1 | GMTACGGTCACACTG | chr3R:24288818-24290818 | 1302 | 1316 | - | 3.6357 | 4.09E-05 | 0.71 | GCCACGGTTACTCCG |
| 1 | GMTACGGTCACACTG | chr2L:7022898-7024898 | 1302 | 1316 | - | 3.6357 | 4.09E-05 | 0.71 | GCAGCGGTTGCACTG |
| 1 | GMTACGGTCACACTG | chr2R:22660934-22662934 | 1317 | 1331 | + | 3.6357 | 4.09E-05 | 0.71 | GCGACGGACACACCC |
| 1 | GMTACGGTCACACTG | chr2L:6036902-6038902 | 1318 | 1332 | + | 3.6357 | 4.09E-05 | 0.71 | GCCGCGGACACAATG |
| 1 | GMTACGGTCACACTG | chr2R:24410074-24412074 | 1311 | 1325 | - | 3.6357 | 4.09E-05 | 0.71 | GCCACAGACACTCTG |
| 1 | GMTACGGTCACACTG | chr3R:30025218-30027218 | 1330 | 1344 | + | 3.6357 | 4.09E-05 | 0.71 | ACCACGGACACTCTG |
| 1 | GMTACGGTCACACTG | chr3R:24657605-24659605 | 1335 | 1349 | + | 3.6357 | 4.09E-05 | 0.71 | GCGACGTTCCCACAG |
| 1 | GMTACGGTCACACTG | chr2L:22339622-22341622 | 1322 | 1336 | - | 3.6357 | 4.09E-05 | 0.71 | GCTACGGATCCACCG |
| 1 | GMTACGGTCACACTG | chrX:15327342-15329342 | 1329 | 1343 | - | 3.6357 | 4.09E-05 | 0.71 | GCGACTGACTCACTG |
| 1 | GMTACGGTCACACTG | chr2L:10789222-10791222 | 1346 | 1360 | + | 3.6357 | 4.09E-05 | 0.71 | GCTACGGAAACGCGG |
| 1 | GMTACGGTCACACTG | chrX:7893031-7895031 | 1351 | 1365 | + | 3.6357 | 4.09E-05 | 0.71 | GCGCTGGACACACTG |
| 1 | GMTACGGTCACACTG | chr3L:3375757-3377757 | 1358 | 1372 | + | 3.6357 | 4.09E-05 | 0.71 | GCTCCGGTCGCTATG |
| 1 | GMTACGGTCACACTG | chr2L:19568972-19570972 | 1361 | 1375 | + | 3.6357 | 4.09E-05 | 0.71 | GCAACGTTCTCACGG |
| 1 | GMTACGGTCACACTG | chr3R:12016156-12018156 | 1361 | 1375 | + | 3.6357 | 4.09E-05 | 0.71 | GCACCTGTCTCACTG |
| 1 | GMTACGGTCACACTG | chrX:4550973-4552973 | 1363 | 1377 | + | 3.6357 | 4.09E-05 | 0.71 | GCGACGATCACGCGG |
| 1 | GMTACGGTCACACTG | chr3L:22464619-22466619 | 1352 | 1366 | - | 3.6357 | 4.09E-05 | 0.71 | GCTTCGATCGCACCG |
| 1 | GMTACGGTCACACTG | chr2L:20024811-20026811 | 1355 | 1369 | - | 3.6357 | 4.09E-05 | 0.71 | GCCACGCCCACTCTG |
| 1 | GMTACGGTCACACTG | chr3R:25236614-25238614 | 1370 | 1384 | + | 3.6357 | 4.09E-05 | 0.71 | CCGACGGACGCACTG |
| 1 | GMTACGGTCACACTG | chr3L:4166464-4168464 | 1367 | 1381 | - | 3.6357 | 4.09E-05 | 0.71 | GCCGCGGACACAATG |
| 1 | GMTACGGTCACACTG | chr3R:14527386-14529386 | 1382 | 1396 | + | 3.6357 | 4.09E-05 | 0.71 | GCTGGGGACCCACTG |
| 1 | GMTACGGTCACACTG | chr2L:11090444-11092444 | 1374 | 1388 | - | 3.6357 | 4.09E-05 | 0.71 | GCTACGGACACCTCG |
| 1 | GMTACGGTCACACTG | chr3L:1644467-1646467 | 1390 | 1404 | + | 3.6357 | 4.09E-05 | 0.71 | GCCCCGGTCATCCTG |
| 1 | GMTACGGTCACACTG | chrX:6012181-6014181 | 1391 | 1405 | + | 3.6357 | 4.09E-05 | 0.71 | GCCTCGCTCACACAG |
| 1 | GMTACGGTCACACTG | chr3L:20298410-20300410 | 1394 | 1408 | + | 3.6357 | 4.09E-05 | 0.71 | GCTCCGGCTTCACTG |
| 1 | GMTACGGTCACACTG | chr3L:20298410-20300410 | 1394 | 1408 | + | 3.6357 | 4.09E-05 | 0.71 | GCTCCGGCTTCACTG |
| 1 | GMTACGGTCACACTG | chr3R:30037728-30039728 | 1381 | 1395 | - | 3.6357 | 4.09E-05 | 0.71 | GCGGCGGTCGCATTG |
| 1 | GMTACGGTCACACTG | chr3L:21960871-21962871 | 1381 | 1395 | - | 3.6357 | 4.09E-05 | 0.71 | GCCACAGACCCACTG |

| | | | | | | | | | |
|---|---|---|---|---|---|---|---|---|---|
| 1 | GMTACGGTCACACTG | chr2R:17955570-17957570 | 1395 | 1409 | + | 3.6357 | 4.09E-05 | 0.71 | GCCACGCCCACACCG |
| 1 | GMTACGGTCACACTG | chr2R:14238703-14240703 | 1397 | 1411 | + | 3.6357 | 4.09E-05 | 0.71 | GCCAGGGCCACACGG |
| 1 | GMTACGGTCACACTG | chr2R:12685249-12687249 | 1384 | 1398 | - | 3.6357 | 4.09E-05 | 0.71 | GCCACAGCCTCACTG |
| 1 | GMTACGGTCACACTG | chr2L:814993-816993 | 1399 | 1413 | + | 3.6357 | 4.09E-05 | 0.71 | GCCTCTGCCACACTG |
| 1 | GMTACGGTCACACTG | chr3L:17032676-17034676 | 1402 | 1416 | + | 3.6357 | 4.09E-05 | 0.71 | GCCACGGGCACTCTC |
| 1 | GMTACGGTCACACTG | chr3R:16974204-16976204 | 1390 | 1404 | - | 3.6357 | 4.09E-05 | 0.71 | GCGACGGGGACACAG |
| 1 | GMTACGGTCACACTG | chr3R:30128119-30130119 | 1402 | 1416 | - | 3.6357 | 4.09E-05 | 0.71 | GCGACGTTCACGCGG |
| 1 | GMTACGGTCACACTG | chr3R:5370795-5372795 | 1418 | 1432 | + | 3.6357 | 4.09E-05 | 0.71 | GAAACGGACCCCCTG |
| 1 | GMTACGGTCACACTG | chr3R:28302571-28304571 | 1411 | 1425 | - | 3.6357 | 4.09E-05 | 0.71 | GCGATGGACCCACTG |
| 1 | GMTACGGTCACACTG | chr2L:4829693-4831693 | 1426 | 1440 | + | 3.6357 | 4.09E-05 | 0.71 | GCTTCGGCCACATCG |
| 1 | GMTACGGTCACACTG | chrX:18675761-18677761 | 1430 | 1444 | + | 3.6357 | 4.09E-05 | 0.71 | GCAACCGACTCACTG |
| 1 | GMTACGGTCACACTG | chrX:19957342-19959342 | 1418 | 1432 | - | 3.6357 | 4.09E-05 | 0.71 | GCTACGGATACGCCG |
| 1 | GMTACGGTCACACTG | chrX:17313581-17315581 | 1433 | 1447 | + | 3.6357 | 4.09E-05 | 0.71 | GCCACGGCCACAACG |
| 1 | GMTACGGTCACACTG | chr2L:814950-816950 | 1442 | 1456 | + | 3.6357 | 4.09E-05 | 0.71 | GCCTCTGCCACACTG |
| 1 | GMTACGGTCACACTG | chrX:12743212-12745212 | 1429 | 1443 | - | 3.6357 | 4.09E-05 | 0.71 | GCTGCTGTCCCACAG |
| 1 | GMTACGGTCACACTG | chr3R:9758622-9760622 | 1429 | 1443 | - | 3.6357 | 4.09E-05 | 0.71 | ACCACGGACACACCG |
| 1 | GMTACGGTCACACTG | chr3L:16082993-16084993 | 1433 | 1447 | - | 3.6357 | 4.09E-05 | 0.71 | GCAGCGGGCATACTG |
| 1 | GMTACGGTCACACTG | chr3L:826843-828843 | 1449 | 1463 | + | 3.6357 | 4.09E-05 | 0.71 | GCCACGATCCCACAG |
| 1 | GMTACGGTCACACTG | chr3L:1552493-1554493 | 1449 | 1463 | + | 3.6357 | 4.09E-05 | 0.71 | GCTCCTGCCACACCG |
| 1 | GMTACGGTCACACTG | chr2R:16000290-16002290 | 1451 | 1465 | + | 3.6357 | 4.09E-05 | 0.71 | ACGGCGGTCACACCG |
| 1 | GMTACGGTCACACTG | chrX:7932742-7934742 | 1438 | 1452 | - | 3.6357 | 4.09E-05 | 0.71 | GCGATGGTCTCACGG |
| 1 | GMTACGGTCACACTG | chr3L:16590050-16592050 | 1439 | 1453 | - | 3.6357 | 4.09E-05 | 0.71 | GCTCCGCTCACCCAG |
| 1 | GMTACGGTCACACTG | chrX:4550878-4552878 | 1458 | 1472 | + | 3.6357 | 4.09E-05 | 0.71 | GCGACGATCACGCGG |
| 1 | GMTACGGTCACACTG | chr3R:9005226-9007226 | 1447 | 1461 | - | 3.6357 | 4.09E-05 | 0.71 | GCAACGGCTCCACTG |
| 1 | GMTACGGTCACACTG | chr3L:11731141-11733141 | 1464 | 1478 | + | 3.6357 | 4.09E-05 | 0.71 | GCAGCAGTCACTCTG |
| 1 | GMTACGGTCACACTG | chr3R:21387880-21389880 | 1465 | 1479 | + | 3.6357 | 4.09E-05 | 0.71 | GCAACGGGCACATCG |
| 1 | GMTACGGTCACACTG | chr2R:9752697-9754697 | 1451 | 1465 | - | 3.6357 | 4.09E-05 | 0.71 | GCTCCGCACACACAG |
| 1 | GMTACGGTCACACTG | chrX:7933898-7935898 | 1451 | 1465 | - | 3.6357 | 4.09E-05 | 0.71 | GCGATGGTCTCACGG |
| 1 | GMTACGGTCACACTG | chr2L:19131323-19133323 | 1466 | 1480 | + | 3.6357 | 4.09E-05 | 0.71 | GCCCCGGTGTCACTG |
| 1 | GMTACGGTCACACTG | chr2R:12685179-12687179 | 1454 | 1468 | - | 3.6357 | 4.09E-05 | 0.71 | GCCACAGCCTCACTG |
| 1 | GMTACGGTCACACTG | chr2L:3814438-3816438 | 1469 | 1483 | + | 3.6357 | 4.09E-05 | 0.71 | GCTTCAGACACCCTG |
| 1 | GMTACGGTCACACTG | chr3R:24787054-24789054 | 1460 | 1474 | - | 3.6357 | 4.09E-05 | 0.71 | GCTCCAGTCCCACCG |
| 1 | GMTACGGTCACACTG | chrX:5312258-5314258 | 1477 | 1491 | + | 3.6357 | 4.09E-05 | 0.71 | GCAACAGGCACACGG |

| | | | | | | | | | |
|---|---|---|---|---|---|---|---|---|---|
| 1 | GMTACGGTCACACTG | chrX:14060817-14062817 | 1466 | 1480 | - | 3.6357 | 4.09E-05 | 0.71 | GCGATGGGCACCCTG |
| 1 | GMTACGGTCACACTG | chr3R:13614917-13616917 | 1481 | 1495 | + | 3.6357 | 4.09E-05 | 0.71 | GCTTCGGTGCCACCG |
| 1 | GMTACGGTCACACTG | chr3R:12016031-12018031 | 1486 | 1500 | + | 3.6357 | 4.09E-05 | 0.71 | GCACCTGTCTCACTG |
| 1 | GMTACGGTCACACTG | chr2R:16196042-16198042 | 1479 | 1493 | - | 3.6357 | 4.09E-05 | 0.71 | GCTTTGGTCGCTCTG |
| 1 | GMTACGGTCACACTG | chr2R:16196042-16198042 | 1479 | 1493 | - | 3.6357 | 4.09E-05 | 0.71 | GCTTTGGTCGCTCTG |
| 1 | GMTACGGTCACACTG | chr3R:13660311-13662311 | 1499 | 1513 | + | 3.6357 | 4.09E-05 | 0.71 | GCTACGGCCGCACAC |
| 1 | GMTACGGTCACACTG | chr3R:25122373-25124373 | 1500 | 1514 | + | 3.6357 | 4.09E-05 | 0.71 | GCAAGGGTCCCGCTG |
| 1 | GMTACGGTCACACTG | chr3R:15191357-15193357 | 1500 | 1514 | + | 3.6357 | 4.09E-05 | 0.71 | GCAGCGTTCACGCTG |
| 1 | GMTACGGTCACACTG | chr2R:10728203-10730203 | 1503 | 1517 | + | 3.6357 | 4.09E-05 | 0.71 | GCCGTGGTCACACCG |
| 1 | GMTACGGTCACACTG | chrY:3021289-3023289 | 1504 | 1518 | + | 3.6357 | 4.09E-05 | 0.71 | GCAAGGGTCCCGCTG |
| 1 | GMTACGGTCACACTG | chr2R:19451504-19453504 | 1490 | 1504 | - | 3.6357 | 4.09E-05 | 0.71 | GCTGCAGTCCCGCTG |
| 1 | GMTACGGTCACACTG | chrX:1451430-1453430 | 1505 | 1519 | + | 3.6357 | 4.09E-05 | 0.71 | GCATAGGTCACACGG |
| 1 | GMTACGGTCACACTG | chr2R:24937181-24939181 | 1491 | 1505 | - | 3.6357 | 4.09E-05 | 0.71 | GATTCGGACTCACCG |
| 1 | GMTACGGTCACACTG | chrX:8156386-8158386 | 1507 | 1521 | + | 3.6357 | 4.09E-05 | 0.71 | GCCACAGCCACACGG |
| 1 | GMTACGGTCACACTG | chr3L:19518649-19520649 | 1515 | 1529 | + | 3.6357 | 4.09E-05 | 0.71 | GCTACGGACACGGCG |
| 1 | GMTACGGTCACACTG | chr3L:1638711-1640711 | 1516 | 1530 | + | 3.6357 | 4.09E-05 | 0.71 | GCGAAGGACACACGG |
| 1 | GMTACGGTCACACTG | chr2R:21063986-21065986 | 1520 | 1534 | + | 3.6357 | 4.09E-05 | 0.71 | GCCACCGACGCACTG |
| 1 | GMTACGGTCACACTG | chrX:20821146-20823146 | 1524 | 1538 | + | 3.6357 | 4.09E-05 | 0.71 | GCTGCGGCGACACCG |
| 1 | GMTACGGTCACACTG | chr3R:17699741-17701741 | 1524 | 1538 | + | 3.6357 | 4.09E-05 | 0.71 | GCAACGGCGGCACTG |
| 1 | GMTACGGTCACACTG | chrX:21051361-21053361 | 1518 | 1532 | - | 3.6357 | 4.09E-05 | 0.71 | GCCACCGTCCCACCG |
| 1 | GMTACGGTCACACTG | chr3R:9745367-9747367 | 1539 | 1553 | + | 3.6357 | 4.09E-05 | 0.71 | GCGTCGATCTCACTG |
| 1 | GMTACGGTCACACTG | chrX:18062247-18064247 | 1525 | 1539 | - | 3.6357 | 4.09E-05 | 0.71 | GCTTCGGACACATAG |
| 1 | GMTACGGTCACACTG | chr3R:24813262-24815262 | 1544 | 1558 | + | 3.6357 | 4.09E-05 | 0.71 | GCTGCGGTGGCGCTG |
| 1 | GMTACGGTCACACTG | chr3L:6124405-6126405 | 1530 | 1544 | - | 3.6357 | 4.09E-05 | 0.71 | GCTCCGGTGGCTCTG |
| 1 | GMTACGGTCACACTG | chr2L:247909-249909 | 1536 | 1550 | - | 3.6357 | 4.09E-05 | 0.71 | GCTTCGGTCGCTGTG |
| 1 | GMTACGGTCACACTG | chr2R:16000181-16002181 | 1560 | 1574 | + | 3.6357 | 4.09E-05 | 0.71 | ACGGCGGTCACACCG |
| 1 | GMTACGGTCACACTG | chr2R:9093658-9095658 | 1563 | 1577 | + | 3.6357 | 4.09E-05 | 0.71 | GCAGCGTGCACACTG |
| 1 | GMTACGGTCACACTG | chr3R:15345205-15347205 | 1550 | 1564 | - | 3.6357 | 4.09E-05 | 0.71 | GCTTTGGTCTCTCTG |
| 1 | GMTACGGTCACACTG | chr2R:9564190-9566190 | 1564 | 1578 | + | 3.6357 | 4.09E-05 | 0.71 | GCGGCGGGGACACTG |
| 1 | GMTACGGTCACACTG | chr3R:7324597-7326597 | 1552 | 1566 | - | 3.6357 | 4.09E-05 | 0.71 | GCGCTGGTCCCACTG |
| 1 | GMTACGGTCACACTG | chr3L:3329063-3331063 | 1569 | 1583 | + | 3.6357 | 4.09E-05 | 0.71 | GCCATGGTCCCACCG |
| 1 | GMTACGGTCACACTG | chr2L:276582-278582 | 1556 | 1570 | - | 3.6357 | 4.09E-05 | 0.71 | ACGTCGGACACACTG |
| 1 | GMTACGGTCACACTG | chr3L:16608347-16610347 | 1571 | 1585 | + | 3.6357 | 4.09E-05 | 0.71 | GCCACGGCCTCAGTG |

| | | | | | | | | | |
|---|---|---|---|---|---|---|---|---|---|
| 1 | GMTACGGTCACACTG | chr3L:19908410-19910410 | 1572 | 1586 | + | 3.6357 | 4.09E-05 | 0.71 | GCGACGGTCGGACCG |
| 1 | GMTACGGTCACACTG | chr3R:20900956-20902956 | 1563 | 1577 | - | 3.6357 | 4.09E-05 | 0.71 | GAGACGGACACGCCG |
| 1 | GMTACGGTCACACTG | chr2L:10429918-10431918 | 1582 | 1596 | + | 3.6357 | 4.09E-05 | 0.71 | GCTACGCACACCCCG |
| 1 | GMTACGGTCACACTG | chr3R:31064253-31066253 | 1570 | 1584 | - | 3.6357 | 4.09E-05 | 0.71 | GCTTCGGATCCACTG |
| 1 | GMTACGGTCACACTG | chrX:7892798-7894798 | 1584 | 1598 | + | 3.6357 | 4.09E-05 | 0.71 | GCGCTGGACACACTG |
| 1 | GMTACGGTCACACTG | chrX:7892798-7894798 | 1584 | 1598 | + | 3.6357 | 4.09E-05 | 0.71 | GCGCTGGACACACTG |
| 1 | GMTACGGTCACACTG | chrX:2789567-2791567 | 1573 | 1587 | - | 3.6357 | 4.09E-05 | 0.71 | GCGCTGGTCACTCTG |
| 1 | GMTACGGTCACACTG | chr2L:4226097-4228097 | 1590 | 1604 | + | 3.6357 | 4.09E-05 | 0.71 | GCGGCGGCCACACTC |
| 1 | GMTACGGTCACACTG | chrX:7012887-7014887 | 1576 | 1590 | - | 3.6357 | 4.09E-05 | 0.71 | GCGACGCTCGCGCTG |
| 1 | GMTACGGTCACACTG | chr3R:15345176-15347176 | 1579 | 1593 | - | 3.6357 | 4.09E-05 | 0.71 | GCTTTGGTCTCTCTG |
| 1 | GMTACGGTCACACTG | chr2R:21669152-21671152 | 1580 | 1594 | - | 3.6357 | 4.09E-05 | 0.71 | TCCTCGGCCACACTG |
| 1 | GMTACGGTCACACTG | chrX:1709182-1711182 | 1596 | 1610 | + | 3.6357 | 4.09E-05 | 0.71 | GCAGCGAGCACACTG |
| 1 | GMTACGGTCACACTG | chrX:7012881-7014881 | 1582 | 1596 | - | 3.6357 | 4.09E-05 | 0.71 | GCGACGCTCGCGCTG |
| 1 | GMTACGGTCACACTG | chr3L:4806406-4808406 | 1583 | 1597 | - | 3.6357 | 4.09E-05 | 0.71 | GCTTCGGGGTCACTG |
| 1 | GMTACGGTCACACTG | chr3R:22744620-22746620 | 1594 | 1608 | - | 3.6357 | 4.09E-05 | 0.71 | TCGACGGGCGCACTG |
| 1 | GMTACGGTCACACTG | chr3L:1369628-1371628 | 1597 | 1611 | - | 3.6357 | 4.09E-05 | 0.71 | GCTGCTGCCGCACTG |
| 1 | GMTACGGTCACACTG | chr2L:16326069-16328069 | 1600 | 1614 | - | 3.6357 | 4.09E-05 | 0.71 | GCAACGTGCACGCTG |
| 1 | GMTACGGTCACACTG | chr3L:19908365-19910365 | 1617 | 1631 | + | 3.6357 | 4.09E-05 | 0.71 | GCGACGGTCGGACCG |
| 1 | GMTACGGTCACACTG | chrX:6266797-6268797 | 1605 | 1619 | - | 3.6357 | 4.09E-05 | 0.71 | GCCACGGTAGCACGG |
| 1 | GMTACGGTCACACTG | chr2R:10728203-10730203 | 1605 | 1619 | - | 3.6357 | 4.09E-05 | 0.71 | TCTGCGGCCACACCG |
| 1 | GMTACGGTCACACTG | chrX:19641869-19643869 | 1624 | 1638 | + | 3.6357 | 4.09E-05 | 0.71 | GCTGCGCTCGCACAG |
| 1 | GMTACGGTCACACTG | chr2R:21657078-21659078 | 1626 | 1640 | + | 3.6357 | 4.09E-05 | 0.71 | GCCACGGTCACGAGG |
| 1 | GMTACGGTCACACTG | chr2R:19398769-19400769 | 1616 | 1630 | - | 3.6357 | 4.09E-05 | 0.71 | CCGACGGCCACACAG |
| 1 | GMTACGGTCACACTG | chr2L:18399462-18401462 | 1631 | 1645 | + | 3.6357 | 4.09E-05 | 0.71 | GCTCAGGCCACACAG |
| 1 | GMTACGGTCACACTG | chr2L:18950837-18952837 | 1632 | 1646 | + | 3.6357 | 4.09E-05 | 0.71 | GCCAAGGGCACACAG |
| 1 | GMTACGGTCACACTG | chr3R:12994147-12996147 | 1624 | 1638 | - | 3.6357 | 4.09E-05 | 0.71 | GCGCCGGTCAAGCTG |
| 1 | GMTACGGTCACACTG | chr3R:22744581-22746581 | 1633 | 1647 | - | 3.6357 | 4.09E-05 | 0.71 | TCGACGGGCGCACTG |
| 1 | GMTACGGTCACACTG | chr3R:27516257-27518257 | 1635 | 1649 | - | 3.6357 | 4.09E-05 | 0.71 | GCTGCGGGCATGCTG |
| 1 | GMTACGGTCACACTG | chr3L:1644199-1646199 | 1658 | 1672 | + | 3.6357 | 4.09E-05 | 0.71 | GCCCCGGTCATCCTG |
| 1 | GMTACGGTCACACTG | chr2L:6138382-6140382 | 1647 | 1661 | - | 3.6357 | 4.09E-05 | 0.71 | GCCACTGTCGCTCTG |
| 1 | GMTACGGTCACACTG | chr3R:22927593-22929593 | 1664 | 1678 | + | 3.6357 | 4.09E-05 | 0.71 | GCATCGATCGCACTG |
| 1 | GMTACGGTCACACTG | chr3L:826620-828620 | 1672 | 1686 | + | 3.6357 | 4.09E-05 | 0.71 | GCCACGATCCCACAG |
| 1 | GMTACGGTCACACTG | chr3L:12187818-12189818 | 1677 | 1691 | + | 3.6357 | 4.09E-05 | 0.71 | GCTTCGGTCCATCTG |

| | | | | | | | | | |
|---|---|---|---|---|---|---|---|---|---|
| 1 | GMTACGGTCACACTG | chr3R:7529753-7531753 | 1664 | 1678 | - | 3.6357 | 4.09E-05 | 0.71 | GCTCCAGTCTCGCTG |
| 1 | GMTACGGTCACACTG | chrX:9272413-9274413 | 1664 | 1678 | - | 3.6357 | 4.09E-05 | 0.71 | GCAGTGGTCGCACTG |
| 1 | GMTACGGTCACACTG | chr2R:22121562-22123562 | 1668 | 1682 | - | 3.6357 | 4.09E-05 | 0.71 | GCTCCTGACGCACTG |
| 1 | GMTACGGTCACACTG | chr3R:18572091-18574091 | 1683 | 1697 | + | 3.6357 | 4.09E-05 | 0.71 | GCGACGGTGACTCAG |
| 1 | GMTACGGTCACACTG | chr3R:18572091-18574091 | 1683 | 1697 | + | 3.6357 | 4.09E-05 | 0.71 | GCGACGGTGACTCAG |
| 1 | GMTACGGTCACACTG | chr2R:23384468-23386468 | 1684 | 1698 | + | 3.6357 | 4.09E-05 | 0.71 | GCCACGGCCGCAATG |
| 1 | GMTACGGTCACACTG | chrX:2940408-2942408 | 1671 | 1685 | - | 3.6357 | 4.09E-05 | 0.71 | GCGACGGCGACGCTG |
| 1 | GMTACGGTCACACTG | chrX:19823080-19825080 | 1671 | 1685 | - | 3.6357 | 4.09E-05 | 0.71 | GCAACGGTCCCGCTT |
| 1 | GMTACGGTCACACTG | chr3R:10048199-10050199 | 1678 | 1692 | - | 3.6357 | 4.09E-05 | 0.71 | GCTACGCACTCGCTG |
| 1 | GMTACGGTCACACTG | chrX:1471522-1473522 | 1680 | 1694 | - | 3.6357 | 4.09E-05 | 0.71 | GCTGCGGGCTCAATG |
| 1 | GMTACGGTCACACTG | chrX:2680723-2682723 | 1681 | 1695 | - | 3.6357 | 4.09E-05 | 0.71 | GCGAGGGTCACGCCG |
| 1 | GMTACGGTCACACTG | chr2L:11290077-11292077 | 1682 | 1696 | - | 3.6357 | 4.09E-05 | 0.71 | GCAACGGTCGCAGCG |
| 1 | GMTACGGTCACACTG | chr3L:1369628-1371628 | 1701 | 1715 | + | 3.6357 | 4.09E-05 | 0.71 | GCAACGGGCCAACTG |
| 1 | GMTACGGTCACACTG | chr2R:8708733-8710733 | 1701 | 1715 | + | 3.6357 | 4.09E-05 | 0.71 | GCCATGGACCCACTG |
| 1 | GMTACGGTCACACTG | chrX:16324763-16326763 | 1706 | 1720 | + | 3.6357 | 4.09E-05 | 0.71 | CCGACGGACTCACTG |
| 1 | GMTACGGTCACACTG | chr3R:24162966-24164966 | 1707 | 1721 | + | 3.6357 | 4.09E-05 | 0.71 | GCTACGGCCTCATGG |
| 1 | GMTACGGTCACACTG | chr2L:19486490-19488490 | 1707 | 1721 | + | 3.6357 | 4.09E-05 | 0.71 | GCATCGAGCACACTG |
| 1 | GMTACGGTCACACTG | chr2R:14255493-14257493 | 1708 | 1722 | + | 3.6357 | 4.09E-05 | 0.71 | GCTCCGATCACGCGG |
| 1 | GMTACGGTCACACTG | chr2R:24474386-24476386 | 1695 | 1709 | - | 3.6357 | 4.09E-05 | 0.71 | GCTCTGGTCTCGCTG |
| 1 | GMTACGGTCACACTG | chr2R:24268179-24270179 | 1721 | 1735 | + | 3.6357 | 4.09E-05 | 0.71 | GCATTGGTCACTCTG |
| 1 | GMTACGGTCACACTG | chr3R:4210783-4212783 | 1725 | 1739 | + | 3.6357 | 4.09E-05 | 0.71 | GCCTCGGTCACGCTA |
| 1 | GMTACGGTCACACTG | chr3L:19895391-19897391 | 1714 | 1728 | - | 3.6357 | 4.09E-05 | 0.71 | GCGACTGGCACACGG |
| 1 | GMTACGGTCACACTG | chr2L:6138309-6140309 | 1720 | 1734 | - | 3.6357 | 4.09E-05 | 0.71 | GCCACTGTCGCTCTG |
| 1 | GMTACGGTCACACTG | chr3R:23706146-23708146 | 1737 | 1751 | + | 3.6357 | 4.09E-05 | 0.71 | GCTCCTGGCACACGG |
| 1 | GMTACGGTCACACTG | chr3R:11948680-11950680 | 1743 | 1757 | + | 3.6357 | 4.09E-05 | 0.71 | GCAGCTGTCACGCTG |
| 1 | GMTACGGTCACACTG | chr2L:5657560-5659560 | 1744 | 1758 | + | 3.6357 | 4.09E-05 | 0.71 | GCTTCGGTCTCTCTT |
| 1 | GMTACGGTCACACTG | chr3L:9429312-9431312 | 1745 | 1759 | + | 3.6357 | 4.09E-05 | 0.71 | GCTCCAGCCACTCTG |
| 1 | GMTACGGTCACACTG | chrX:19651305-19653305 | 1731 | 1745 | - | 3.6357 | 4.09E-05 | 0.71 | GCCACGGCCAGACAG |
| 1 | GMTACGGTCACACTG | chr2L:18278475-18280475 | 1731 | 1745 | - | 3.6357 | 4.09E-05 | 0.71 | GCGACGGTCACTCAA |
| 1 | GMTACGGTCACACTG | chr2R:23842286-23844286 | 1749 | 1763 | + | 3.6357 | 4.09E-05 | 0.71 | GCCACAGGCACCCTG |
| 1 | GMTACGGTCACACTG | chr3R:14216882-14218882 | 1737 | 1751 | - | 3.6357 | 4.09E-05 | 0.71 | GCACCGGCAACACTG |
| 1 | GMTACGGTCACACTG | chr3L:3555299-3557299 | 1762 | 1776 | + | 3.6357 | 4.09E-05 | 0.71 | GCGACGGCCACACGT |
| 1 | GMTACGGTCACACTG | chr3R:29798658-29800658 | 1768 | 1782 | + | 3.6357 | 4.09E-05 | 0.71 | GCCAAGGACACACAG |

| | | | | | | | | | |
|---|---|---|---|---|---|---|---|---|---|
| 1 | GMTACGGTCACACTG | chr3R:13617559-13619559 | 1756 | 1770 | - | 3.6357 | 4.09E-05 | 0.71 | GCTCCGGCCACTTTG |
| 1 | GMTACGGTCACACTG | chr2L:19568561-19570561 | 1772 | 1786 | + | 3.6357 | 4.09E-05 | 0.71 | GCAACGTTCTCACGG |
| 1 | GMTACGGTCACACTG | chr3L:6038155-6040155 | 1773 | 1787 | + | 3.6357 | 4.09E-05 | 0.71 | GCAACGGTGACTCCG |
| 1 | GMTACGGTCACACTG | chr3R:24230364-24232364 | 1774 | 1788 | + | 3.6357 | 4.09E-05 | 0.71 | GCCACGCCCACACCG |
| 1 | GMTACGGTCACACTG | chrX:19651269-19653269 | 1767 | 1781 | - | 3.6357 | 4.09E-05 | 0.71 | GCCACGGCCAGACAG |
| 1 | GMTACGGTCACACTG | chr3R:13573825-13575825 | 1784 | 1798 | + | 3.6357 | 4.09E-05 | 0.71 | TCCACGGCCACACCG |
| 1 | GMTACGGTCACACTG | chr3L:5809973-5811973 | 1785 | 1799 | + | 3.6357 | 4.09E-05 | 0.71 | GCTGCGGGAACACGG |
| 1 | GMTACGGTCACACTG | chr3L:20949783-20951783 | 1785 | 1799 | + | 3.6357 | 4.09E-05 | 0.71 | GCCACGGACTCAGTG |
| 1 | GMTACGGTCACACTG | chr2L:6731866-6733866 | 1772 | 1786 | - | 3.6357 | 4.09E-05 | 0.71 | GCATCGATCACACGG |
| 1 | GMTACGGTCACACTG | chr3R:24813012-24815012 | 1794 | 1808 | + | 3.6357 | 4.09E-05 | 0.71 | GCTGCGGTGGCGCTG |
| 1 | GMTACGGTCACACTG | chr3L:3279239-3281239 | 1782 | 1796 | - | 3.6357 | 4.09E-05 | 0.71 | TCTGCGGCCACTCTG |
| 1 | GMTACGGTCACACTG | chr3L:19600633-19602633 | 1783 | 1797 | - | 3.6357 | 4.09E-05 | 0.71 | ACAACGGTCGCCCTG |
| 1 | GMTACGGTCACACTG | chr3L:8517885-8519885 | 1786 | 1800 | - | 3.6357 | 4.09E-05 | 0.71 | GCAACGGACTCACTT |
| 1 | GMTACGGTCACACTG | chr3R:7085599-7087599 | 1790 | 1804 | - | 3.6357 | 4.09E-05 | 0.71 | GCTCCAGTCTCGCTG |
| 1 | GMTACGGTCACACTG | chr3L:4140467-4142467 | 1792 | 1806 | - | 3.6357 | 4.09E-05 | 0.71 | GCCACTGACACACCG |
| 1 | GMTACGGTCACACTG | chr2R:22665605-22667605 | 1793 | 1807 | - | 3.6357 | 4.09E-05 | 0.71 | GATCCGGACCCACCG |
| 1 | GMTACGGTCACACTG | chrX:10847757-10849757 | 1795 | 1809 | - | 3.6357 | 4.09E-05 | 0.71 | GCAATGGGCGCACTG |
| 1 | GMTACGGTCACACTG | chrX:16805068-16807068 | 1798 | 1812 | - | 3.6357 | 4.09E-05 | 0.71 | GCCCTGGTCACACAG |
| 1 | GMTACGGTCACACTG | chr3L:5809931-5811931 | 1827 | 1841 | + | 3.6357 | 4.09E-05 | 0.71 | GCTGCGGGAACACGG |
| 1 | GMTACGGTCACACTG | chr2R:21668916-21670916 | 1816 | 1830 | - | 3.6357 | 4.09E-05 | 0.71 | TCCTCGGCCACACTG |
| 1 | GMTACGGTCACACTG | chr2R:21668916-21670916 | 1816 | 1830 | - | 3.6357 | 4.09E-05 | 0.71 | TCCTCGGCCACACTG |
| 1 | GMTACGGTCACACTG | chr2R:21668916-21670916 | 1816 | 1830 | - | 3.6357 | 4.09E-05 | 0.71 | TCCTCGGCCACACTG |
| 1 | GMTACGGTCACACTG | chr2R:21668916-21670916 | 1816 | 1830 | - | 3.6357 | 4.09E-05 | 0.71 | TCCTCGGCCACACTG |
| 1 | GMTACGGTCACACTG | chr3L:9796687-9798687 | 1831 | 1845 | + | 3.6357 | 4.09E-05 | 0.71 | GCAATGGTCACGCCG |
| 1 | GMTACGGTCACACTG | chr2L:20644822-20646822 | 1834 | 1848 | + | 3.6357 | 4.09E-05 | 0.71 | GAGTCGGCCGCACTG |
| 1 | GMTACGGTCACACTG | chr2L:19130954-19132954 | 1835 | 1849 | + | 3.6357 | 4.09E-05 | 0.71 | GCCCCGGTGTCACTG |
| 1 | GMTACGGTCACACTG | chr3L:13348165-13350165 | 1840 | 1854 | + | 3.6357 | 4.09E-05 | 0.71 | GCCCCGCTCACACGG |
| 1 | GMTACGGTCACACTG | chr3L:8115179-8117179 | 1841 | 1855 | + | 3.6357 | 4.09E-05 | 0.71 | GCTTTGGCCCCACTG |
| 1 | GMTACGGTCACACTG | chr3L:13432718-13434718 | 1827 | 1841 | - | 3.6357 | 4.09E-05 | 0.71 | CCGACGGTCACTCAG |
| 1 | GMTACGGTCACACTG | chrX:7932351-7934351 | 1829 | 1843 | - | 3.6357 | 4.09E-05 | 0.71 | GCGATGGTCTCACGG |
| 1 | GMTACGGTCACACTG | chr2L:3824675-3826675 | 1848 | 1862 | + | 3.6357 | 4.09E-05 | 0.71 | GCTACGGATACCCGG |
| 1 | GMTACGGTCACACTG | chr2R:17036982-17038982 | 1839 | 1853 | - | 3.6357 | 4.09E-05 | 0.71 | GCCACGGATCCACTG |
| 1 | GMTACGGTCACACTG | chr2R:22074308-22076308 | 1845 | 1859 | - | 3.6357 | 4.09E-05 | 0.71 | GCTGCGGTTGCCCTG |

| | | | | | | | | | |
|---|---|---|---|---|---|---|---|---|---|
| 1 | GMTACGGTCACACTG | chr2L:20644791-20646791 | 1865 | 1879 | + | 3.6357 | 4.09E-05 | 0.71 | GAGTCGGCCGCACTG |
| 1 | GMTACGGTCACACTG | chr3L:9353033-9355033 | 1871 | 1885 | + | 3.6357 | 4.09E-05 | 0.71 | GCTACGGAATCCCTG |
| 1 | GMTACGGTCACACTG | chr3R:29145893-29147893 | 1858 | 1872 | - | 3.6357 | 4.09E-05 | 0.71 | GCCGCGCTCACCCTG |
| 1 | GMTACGGTCACACTG | chr2L:2211571-2213571 | 1874 | 1888 | + | 3.6357 | 4.09E-05 | 0.71 | GCGACGGTTCCACAG |
| 1 | GMTACGGTCACACTG | chr2L:17497821-17499821 | 1876 | 1890 | + | 3.6357 | 4.09E-05 | 0.71 | GCTACGATCGCTCCG |
| 1 | GMTACGGTCACACTG | chr2L:10198117-10200117 | 1864 | 1878 | - | 3.6357 | 4.09E-05 | 0.71 | GCTTCGAGCACTCTG |
| 1 | GMTACGGTCACACTG | chr2L:17476888-17478888 | 1868 | 1882 | - | 3.6357 | 4.09E-05 | 0.71 | TCTTCGGACACGCTG |
| 1 | GMTACGGTCACACTG | chr2R:11237995-11239995 | 1882 | 1896 | + | 3.6357 | 4.09E-05 | 0.71 | GCTTCTGGCGCACTG |
| 1 | GMTACGGTCACACTG | chr2L:6053302-6055302 | 1884 | 1898 | + | 3.6357 | 4.09E-05 | 0.71 | GCTACGGGCTGGCTG |
| 1 | GMTACGGTCACACTG | chr2R:15690113-15692113 | 1884 | 1898 | + | 3.6357 | 4.09E-05 | 0.71 | GCTCCGGTGACTCAG |
| 1 | GMTACGGTCACACTG | chr3L:8954021-8956021 | 1886 | 1900 | + | 3.6357 | 4.09E-05 | 0.71 | GCCGCGGGCACAGTG |
| 1 | GMTACGGTCACACTG | chr2L:19529305-19531305 | 1887 | 1901 | + | 3.6357 | 4.09E-05 | 0.71 | GCAACGAGCACGCTG |
| 1 | GMTACGGTCACACTG | chrX:13821826-13823826 | 1875 | 1889 | - | 3.6357 | 4.09E-05 | 0.71 | GCTACGGACTCATCG |
| 1 | GMTACGGTCACACTG | chrX:19207187-19209187 | 1896 | 1910 | + | 3.6357 | 4.09E-05 | 0.71 | ACGGCGGACACACTG |
| 1 | GMTACGGTCACACTG | chr3R:18273665-18275665 | 1884 | 1898 | - | 3.6357 | 4.09E-05 | 0.71 | TCGGCGGTCACCCTG |
| 1 | GMTACGGTCACACTG | chr3R:24787054-24789054 | 1889 | 1903 | - | 3.6357 | 4.09E-05 | 0.71 | GCCAGGGACCCACTG |
| 1 | GMTACGGTCACACTG | chr3L:8179923-8181923 | 1894 | 1908 | - | 3.6357 | 4.09E-05 | 0.71 | GCTGCGGATGCACTG |
| 1 | GMTACGGTCACACTG | chr3L:826383-828383 | 1909 | 1923 | + | 3.6357 | 4.09E-05 | 0.71 | GCCACGATCCCACAG |
| 1 | GMTACGGTCACACTG | chr3R:26716871-26718871 | 1899 | 1913 | - | 3.6357 | 4.09E-05 | 0.71 | GCTACGGGCACCCAC |
| 1 | GMTACGGTCACACTG | chrX:15023436-15025436 | 1915 | 1929 | + | 3.6357 | 4.09E-05 | 0.71 | ACTTCGGCCACTCTG |
| 1 | GMTACGGTCACACTG | chr3L:7349404-7351404 | 1901 | 1915 | - | 3.6357 | 4.09E-05 | 0.71 | GCCACGGGCACACCT |
| 1 | GMTACGGTCACACTG | chr2R:7067572-7069572 | 1902 | 1916 | - | 3.6357 | 4.09E-05 | 0.71 | GAAGCGGTCCCACAG |
| 1 | GMTACGGTCACACTG | chr3R:15224576-15226576 | 1918 | 1932 | + | 3.6357 | 4.09E-05 | 0.71 | GCTACGGACTCTCTT |
| 1 | GMTACGGTCACACTG | chr3R:29050056-29052056 | 1929 | 1943 | + | 3.6357 | 4.09E-05 | 0.71 | GCAACGGCGACTCTG |
| 1 | GMTACGGTCACACTG | chr3R:29691591-29693591 | 1934 | 1948 | + | 3.6357 | 4.09E-05 | 0.71 | GCTACGCCCACTCCG |
| 1 | GMTACGGTCACACTG | chr2L:19429643-19431643 | 1938 | 1952 | + | 3.6357 | 4.09E-05 | 0.71 | GCTACCGACGCCCTG |
| 1 | GMTACGGTCACACTG | chr3R:30211903-30213903 | 1940 | 1954 | + | 3.6357 | 4.09E-05 | 0.71 | GCTACGGCTGCACAG |
| 1 | GMTACGGTCACACTG | chrX:13758546-13760546 | 1933 | 1947 | - | 3.6357 | 4.09E-05 | 0.71 | GCTCCGGCCACATGG |
| 1 | GMTACGGTCACACTG | chr2R:8708486-8710486 | 1948 | 1962 | + | 3.6357 | 4.09E-05 | 0.71 | GCCATGGACCCACTG |
| 1 | GMTACGGTCACACTG | chr3R:24230186-24232186 | 1952 | 1966 | + | 3.6357 | 4.09E-05 | 0.71 | GCCACGCCCACACCG |
| 1 | GMTACGGTCACACTG | chr3R:25121920-25123920 | 1953 | 1967 | + | 3.6357 | 4.09E-05 | 0.71 | GCAAGGGTCCCGCTG |
| 1 | GMTACGGTCACACTG | chr3R:25121920-25123920 | 1953 | 1967 | + | 3.6357 | 4.09E-05 | 0.71 | GCAAGGGTCCCGCTG |
| 1 | GMTACGGTCACACTG | chrX:6996799-6998799 | 1941 | 1955 | - | 3.6357 | 4.09E-05 | 0.71 | GCGACGGTTACCCAG |

| | | | | | | | | | |
|---|---|---|---|---|---|---|---|---|---|
| 1 | GMTACGGTCACACTG | chr3L:1000027-1002027 | 1971 | 1985 | + | 3.6357 | 4.09E-05 | 0.71 | GCGACGGACACATCG |
| 1 | GMTACGGTCACACTG | chr3R:12424917-12426917 | 1973 | 1987 | + | 3.6357 | 4.09E-05 | 0.71 | GCTGCAGGCCCACTG |
| 1 | GMTACGGTCACACTG | chr3R:24371762-24373762 | 1961 | 1975 | - | 3.6357 | 4.09E-05 | 0.71 | GCTCCGCGCACTCTG |
| 1 | GMTACGGTCACACTG | chr3L:7973991-7975991 | 1979 | 1993 | + | 3.6357 | 4.09E-05 | 0.71 | GCTCCGTTCGCCCTG |
| 1 | GMTACGGTCACACTG | chr3L:7344183-7346183 | 1985 | 1999 | - | 3.6357 | 4.09E-05 | 0.71 | GATTCGGCCACTCCG |
| 1 | GMTACGGTCACACTG | chr3R:30038719-30040719 | 6 | 20 | + | 3.3571 | 7.12E-05 | 0.795 | GCTACAGACTAACTG |
| 1 | GMTACGGTCACACTG | chr3R:30038719-30040719 | 6 | 20 | + | 3.3571 | 7.12E-05 | 0.795 | GCTACAGACTAACTG |
| 1 | GMTACGGTCACACTG | chr2L:20416057-20418057 | 6 | 20 | + | 3.3571 | 7.12E-05 | 0.795 | GAAACGATCACGCAG |
| 1 | GMTACGGTCACACTG | chr3R:22582225-22584225 | 7 | 21 | + | 3.3571 | 7.12E-05 | 0.795 | GCTGGGGGCATACTG |
| 1 | GMTACGGTCACACTG | chr2R:9016195-9018195 | 8 | 22 | + | 3.3571 | 7.12E-05 | 0.795 | TCTGCGGTCGCAATG |
| 1 | GMTACGGTCACACTG | chr3L:15815979-15817979 | 11 | 25 | + | 3.3571 | 7.12E-05 | 0.795 | GCTCCGGTCCAATTG |
| 1 | GMTACGGTCACACTG | chr3L:14613458-14615458 | 13 | 27 | + | 3.3571 | 7.12E-05 | 0.795 | GCTACTCACGCACTG |
| 1 | GMTACGGTCACACTG | chr3R:4577644-4579644 | 14 | 28 | + | 3.3571 | 7.12E-05 | 0.795 | GCCACGCCCACACTT |
| 1 | GMTACGGTCACACTG | chr3L:17011565-17013565 | 16 | 30 | + | 3.3571 | 7.12E-05 | 0.795 | GCTGCGGTCAGCGTG |
| 1 | GMTACGGTCACACTG | chr3L:12413842-12415842 | 18 | 32 | + | 3.3571 | 7.12E-05 | 0.795 | GCTAAGGATGCACTG |
| 1 | GMTACGGTCACACTG | chr3L:19096709-19098709 | 18 | 32 | + | 3.3571 | 7.12E-05 | 0.795 | GCTGCGGCCAAACTT |
| 1 | GMTACGGTCACACTG | chr2L:10422291-10424291 | 19 | 33 | + | 3.3571 | 7.12E-05 | 0.795 | GCAACGATCGCATTG |
| 1 | GMTACGGTCACACTG | chr3L:11135049-11137049 | 20 | 34 | + | 3.3571 | 7.12E-05 | 0.795 | GCCACGGTCACTGTT |
| 1 | GMTACGGTCACACTG | chr3R:27238163-27240163 | 22 | 36 | + | 3.3571 | 7.12E-05 | 0.795 | TCGATGGTCACGCTG |
| 1 | GMTACGGTCACACTG | chr3R:8032301-8034301 | 23 | 37 | + | 3.3571 | 7.12E-05 | 0.795 | TATACGGCCACTCCG |
| 1 | GMTACGGTCACACTG | chr2L:453754-455754 | 24 | 38 | + | 3.3571 | 7.12E-05 | 0.795 | GAGATGGTCACCCGG |
| 1 | GMTACGGTCACACTG | chr2R:9251021-9253021 | 24 | 38 | + | 3.3571 | 7.12E-05 | 0.795 | GCTCCGTTCCAACTG |
| 1 | GMTACGGTCACACTG | chr3R:23679039-23681039 | 25 | 39 | + | 3.3571 | 7.12E-05 | 0.795 | GCGACGCTCCCATTG |
| 1 | GMTACGGTCACACTG | chr3R:28339800-28341800 | 25 | 39 | + | 3.3571 | 7.12E-05 | 0.795 | GCTACAGCCACACGC |
| 1 | GMTACGGTCACACTG | chrX:18514428-18516428 | 27 | 41 | + | 3.3571 | 7.12E-05 | 0.795 | GCGACGCTCACCATG |
| 1 | GMTACGGTCACACTG | chrX:18514428-18516428 | 27 | 41 | + | 3.3571 | 7.12E-05 | 0.795 | GCGACGCTCACCATG |
| 1 | GMTACGGTCACACTG | chr3R:5570108-5572108 | 28 | 42 | + | 3.3571 | 7.12E-05 | 0.795 | ACTACGGCCGAACTG |
| 1 | GMTACGGTCACACTG | chr3R:27687134-27689134 | 29 | 43 | + | 3.3571 | 7.12E-05 | 0.795 | GATGAGGTCGCTCTG |
| 1 | GMTACGGTCACACTG | chr2R:16773246-16775246 | 30 | 44 | + | 3.3571 | 7.12E-05 | 0.795 | GCCACGGTCAGCGTG |
| 1 | GMTACGGTCACACTG | chrX:13668699-13670699 | 31 | 45 | + | 3.3571 | 7.12E-05 | 0.795 | ACTGCGGCCACAGTG |
| 1 | GMTACGGTCACACTG | chr3L:16858080-16860080 | 32 | 46 | + | 3.3571 | 7.12E-05 | 0.795 | GCTAGGAACACACAG |
| 1 | GMTACGGTCACACTG | chr2R:6695661-6697661 | 32 | 46 | + | 3.3571 | 7.12E-05 | 0.795 | GAAACGGTGGCACAG |
| 1 | GMTACGGTCACACTG | chr2L:19131323-19133323 | 32 | 46 | + | 3.3571 | 7.12E-05 | 0.795 | GCCACCGGCAGACTG |

| | | | | | | | | | |
|---|---|---|---|---|---|---|---|---|---|
| 1 | GMTACGGTCACACTG | chr3R:18222722-18224722 | 33 | 47 | + | 3.3571 | 7.12E-05 | 0.795 | GAGACGTTCACGCAG |
| 1 | GMTACGGTCACACTG | chr3R:8113003-8115003 | 34 | 48 | + | 3.3571 | 7.12E-05 | 0.795 | ACTTTGGCCACACTG |
| 1 | GMTACGGTCACACTG | chrX:19192555-19194555 | 36 | 50 | + | 3.3571 | 7.12E-05 | 0.795 | GATGAGGACACGCTG |
| 1 | GMTACGGTCACACTG | chr2R:6161197-6163197 | 37 | 51 | + | 3.3571 | 7.12E-05 | 0.795 | GCCACGCCCACACTT |
| 1 | GMTACGGTCACACTG | chr3L:177963-179963 | 38 | 52 | + | 3.3571 | 7.12E-05 | 0.795 | ACTACGGTCTCACAT |
| 1 | GMTACGGTCACACTG | chr3R:22381035-22383035 | 41 | 55 | + | 3.3571 | 7.12E-05 | 0.795 | CAAACGGACGCACTG |
| 1 | GMTACGGTCACACTG | chr3L:4752738-4754738 | 42 | 56 | + | 3.3571 | 7.12E-05 | 0.795 | GCAACTGTCCAACTG |
| 1 | GMTACGGTCACACTG | chr2R:9108718-9110718 | 43 | 57 | + | 3.3571 | 7.12E-05 | 0.795 | GAGGCGGTCACCATG |
| 1 | GMTACGGTCACACTG | chr2R:23805249-23807249 | 47 | 61 | + | 3.3571 | 7.12E-05 | 0.795 | GCTTCGATGACACAG |
| 1 | GMTACGGTCACACTG | chr3R:31757710-31759710 | 52 | 66 | + | 3.3571 | 7.12E-05 | 0.795 | GCTACGCTCTCACCT |
| 1 | GMTACGGTCACACTG | chr3R:25357285-25359285 | 54 | 68 | + | 3.3571 | 7.12E-05 | 0.795 | ACTCCGCCCACACTG |
| 1 | GMTACGGTCACACTG | chr2L:8219741-8221741 | 54 | 68 | + | 3.3571 | 7.12E-05 | 0.795 | GCTAGGTACTCACTG |
| 1 | GMTACGGTCACACTG | chrX:6523464-6525464 | 56 | 70 | + | 3.3571 | 7.12E-05 | 0.795 | ACATTGGTCACACTG |
| 1 | GMTACGGTCACACTG | chr2R:7421994-7423994 | 56 | 70 | + | 3.3571 | 7.12E-05 | 0.795 | GAAACAGGCGCACTG |
| 1 | GMTACGGTCACACTG | chrX:19267896-19269896 | 57 | 71 | + | 3.3571 | 7.12E-05 | 0.795 | GATGCGGTACCGCTG |
| 1 | GMTACGGTCACACTG | chr3L:8728865-8730865 | 58 | 72 | + | 3.3571 | 7.12E-05 | 0.795 | GCTACGGCCAACGTG |
| 1 | GMTACGGTCACACTG | chr3R:24335922-24337922 | 59 | 73 | + | 3.3571 | 7.12E-05 | 0.795 | ACTACGGTCTGGCTG |
| 1 | GMTACGGTCACACTG | chr2L:10697043-10699043 | 59 | 73 | + | 3.3571 | 7.12E-05 | 0.795 | GATCCTGACCCACTG |
| 1 | GMTACGGTCACACTG | chr2R:13959675-13961675 | 60 | 74 | + | 3.3571 | 7.12E-05 | 0.795 | GATGCGGCGGCACTG |
| 1 | GMTACGGTCACACTG | chr2L:4453626-4455626 | 62 | 76 | + | 3.3571 | 7.12E-05 | 0.795 | GCTGCAGTCATACCG |
| 1 | GMTACGGTCACACTG | chr2L:18290236-18292236 | 62 | 76 | + | 3.3571 | 7.12E-05 | 0.795 | GCTAAGGATGCACTG |
| 1 | GMTACGGTCACACTG | chr2R:16293130-16295130 | 70 | 84 | + | 3.3571 | 7.12E-05 | 0.795 | GAAACGGGCACCATG |
| 1 | GMTACGGTCACACTG | chr2R:19669618-19671618 | 71 | 85 | + | 3.3571 | 7.12E-05 | 0.795 | GACGGGGGCACACTG |
| 1 | GMTACGGTCACACTG | chr2L:6130023-6132023 | 73 | 87 | + | 3.3571 | 7.12E-05 | 0.795 | GAAACGCTCACGCAG |
| 1 | GMTACGGTCACACTG | chr3R:18222682-18224682 | 73 | 87 | + | 3.3571 | 7.12E-05 | 0.795 | GAGACGTTCACGCAG |
| 1 | GMTACGGTCACACTG | chr3R:22381035-22383035 | 77 | 91 | + | 3.3571 | 7.12E-05 | 0.795 | GAATCGCTCGCACTG |
| 1 | GMTACGGTCACACTG | chr3R:5624367-5626367 | 78 | 92 | + | 3.3571 | 7.12E-05 | 0.795 | GCGATGGTCGAACTG |
| 1 | GMTACGGTCACACTG | chr3L:3177580-3179580 | 78 | 92 | + | 3.3571 | 7.12E-05 | 0.795 | GAGAAGGTCGCTCTG |
| 1 | GMTACGGTCACACTG | chr3R:8728693-8730693 | 79 | 93 | + | 3.3571 | 7.12E-05 | 0.795 | GCTGCGGGAACAGTG |
| 1 | GMTACGGTCACACTG | chrX:19179173-19181173 | 80 | 94 | + | 3.3571 | 7.12E-05 | 0.795 | GATACTGGCCCACGG |
| 1 | GMTACGGTCACACTG | chr3R:7116732-7118732 | 83 | 97 | + | 3.3571 | 7.12E-05 | 0.795 | GATGCGGCAACACGG |
| 1 | GMTACGGTCACACTG | chr2R:8939415-8941415 | 83 | 97 | + | 3.3571 | 7.12E-05 | 0.795 | GCTGCGGACACATTC |
| 1 | GMTACGGTCACACTG | chrX:4704309-4706309 | 86 | 100 | + | 3.3571 | 7.12E-05 | 0.795 | GATCCGGGCACATCG |

| | | | | | | | | | |
|---|---|---|---|---|---|---|---|---|---|
| 1 | GMTACGGTCACACTG | chr3L:11576044-11578044 | 88 | 102 | + | 3.3571 | 7.12E-05 | 0.795 | GAAACGGACAGACAG |
| 1 | GMTACGGTCACACTG | chrX:2926258-2928258 | 89 | 103 | + | 3.3571 | 7.12E-05 | 0.795 | GCTACGGCAATGCTG |
| 1 | GMTACGGTCACACTG | chr3R:24527047-24529047 | 89 | 103 | + | 3.3571 | 7.12E-05 | 0.795 | GCCAATGTCACGCTG |
| 1 | GMTACGGTCACACTG | chr2L:14517691-14519691 | 92 | 106 | + | 3.3571 | 7.12E-05 | 0.795 | GCTACTGTCAGTCGG |
| 1 | GMTACGGTCACACTG | chr3R:16079992-16081992 | 92 | 106 | + | 3.3571 | 7.12E-05 | 0.795 | GCTAAGATCACTCAG |
| 1 | GMTACGGTCACACTG | chrX:6673387-6675387 | 93 | 107 | + | 3.3571 | 7.12E-05 | 0.795 | GATGCGGGCACAGGG |
| 1 | GMTACGGTCACACTG | chr3L:8136808-8138808 | 94 | 108 | + | 3.3571 | 7.12E-05 | 0.795 | GCTACTGCCAGACGG |
| 1 | GMTACGGTCACACTG | chr3L:881013-883013 | 95 | 109 | + | 3.3571 | 7.12E-05 | 0.795 | GCTGCTCTCCCACTG |
| 1 | GMTACGGTCACACTG | chr2R:5803712-5805712 | 96 | 110 | + | 3.3571 | 7.12E-05 | 0.795 | GATGCAGGCACACAG |
| 1 | GMTACGGTCACACTG | chrX:10218139-10220139 | 96 | 110 | + | 3.3571 | 7.12E-05 | 0.795 | ACCACAGGCACACTG |
| 1 | GMTACGGTCACACTG | chr3R:27238085-27240085 | 100 | 114 | + | 3.3571 | 7.12E-05 | 0.795 | TCGATGGTCACGCTG |
| 1 | GMTACGGTCACACTG | chr2L:15761758-15763758 | 102 | 116 | + | 3.3571 | 7.12E-05 | 0.795 | GCAACGATAACGCTG |
| 1 | GMTACGGTCACACTG | chr3R:16453492-16455492 | 102 | 116 | + | 3.3571 | 7.12E-05 | 0.795 | GCTGCAATCGCACTG |
| 1 | GMTACGGTCACACTG | chr3R:6405855-6407855 | 105 | 119 | + | 3.3571 | 7.12E-05 | 0.795 | CAAACGGTCCCGCTG |
| 1 | GMTACGGTCACACTG | chr2L:21890095-21892095 | 107 | 121 | + | 3.3571 | 7.12E-05 | 0.795 | GCTTCGCTGCCACTG |
| 1 | GMTACGGTCACACTG | chrX:927139-929139 | 109 | 123 | + | 3.3571 | 7.12E-05 | 0.795 | GCTCCGGTCACAAAT |
| 1 | GMTACGGTCACACTG | chr3R:20578850-20580850 | 109 | 123 | + | 3.3571 | 7.12E-05 | 0.795 | GACACCGTCACGCCG |
| 1 | GMTACGGTCACACTG | chr3L:9837670-9839670 | 109 | 123 | + | 3.3571 | 7.12E-05 | 0.795 | GATGCGGTCTAGCTG |
| 1 | GMTACGGTCACACTG | chr3L:26164235-26166235 | 112 | 126 | + | 3.3571 | 7.12E-05 | 0.795 | GCCACGCACACACTT |
| 1 | GMTACGGTCACACTG | chr3L:5341084-5343084 | 112 | 126 | + | 3.3571 | 7.12E-05 | 0.795 | TACACGGCCACACAG |
| 1 | GMTACGGTCACACTG | chr3R:19876737-19878737 | 112 | 126 | + | 3.3571 | 7.12E-05 | 0.795 | GCTGTAGTCACACGG |
| 1 | GMTACGGTCACACTG | chr3L:18672324-18674324 | 112 | 126 | + | 3.3571 | 7.12E-05 | 0.795 | GATCCGGCAACCCTG |
| 1 | GMTACGGTCACACTG | chr2R:22070286-22072286 | 112 | 126 | + | 3.3571 | 7.12E-05 | 0.795 | TAGACGGGCGCACTG |
| 1 | GMTACGGTCACACTG | chrX:9910126-9912126 | 115 | 129 | + | 3.3571 | 7.12E-05 | 0.795 | GCCAAGGTCACGTTG |
| 1 | GMTACGGTCACACTG | chrX:10966134-10968134 | 116 | 130 | + | 3.3571 | 7.12E-05 | 0.795 | GCAAAGATCACCCTG |
| 1 | GMTACGGTCACACTG | chrX:4908102-4910102 | 116 | 130 | + | 3.3571 | 7.12E-05 | 0.795 | GATACCGGCTCGCTG |
| 1 | GMTACGGTCACACTG | chr2R:24056798-24058798 | 119 | 133 | + | 3.3571 | 7.12E-05 | 0.795 | ACTACGCCCACACCG |
| 1 | GMTACGGTCACACTG | chr3R:7405869-7407869 | 123 | 137 | + | 3.3571 | 7.12E-05 | 0.795 | GCTAGGTGCACACCG |
| 1 | GMTACGGTCACACTG | chr3R:31757637-31759637 | 125 | 139 | + | 3.3571 | 7.12E-05 | 0.795 | GCTACGCTCTCACCT |
| 1 | GMTACGGTCACACTG | chr3R:20559280-20561280 | 131 | 145 | + | 3.3571 | 7.12E-05 | 0.795 | GCTACGAGCACATCG |
| 1 | GMTACGGTCACACTG | chr3L:9637112-9639112 | 131 | 145 | + | 3.3571 | 7.12E-05 | 0.795 | GCAATGGGCATACTG |
| 1 | GMTACGGTCACACTG | chrX:16008114-16010114 | 135 | 149 | + | 3.3571 | 7.12E-05 | 0.795 | GAAACGGTCAGTCAG |
| 1 | GMTACGGTCACACTG | chr2L:20947170-20949170 | 136 | 150 | + | 3.3571 | 7.12E-05 | 0.795 | GCTCCGATGACGCTG |

| | | | | | | | | | |
|---|---|---|---|---|---|---|---|---|---|
| 1 | GMTACGGTCACACTG | chr2R:6161096-6163096 | 138 | 152 | + | 3.3571 | 7.12E-05 | 0.795 | GCCACGCCCACACTT |
| 1 | GMTACGGTCACACTG | chrX:8564178-8566178 | 141 | 155 | + | 3.3571 | 7.12E-05 | 0.795 | GCTCCTTTCGCACTG |
| 1 | GMTACGGTCACACTG | chr3L:13436136-13438136 | 142 | 156 | + | 3.3571 | 7.12E-05 | 0.795 | GCCAGGAGCACACTG |
| 1 | GMTACGGTCACACTG | chr2R:10426915-10428915 | 142 | 156 | + | 3.3571 | 7.12E-05 | 0.795 | TCATCGGTGACACTG |
| 1 | GMTACGGTCACACTG | chrX:19192448-19194448 | 143 | 157 | + | 3.3571 | 7.12E-05 | 0.795 | GATGAGGACACGCTG |
| 1 | GMTACGGTCACACTG | chr2L:20415918-20417918 | 145 | 159 | + | 3.3571 | 7.12E-05 | 0.795 | GAAACGATCACGCAG |
| 1 | GMTACGGTCACACTG | chr3R:16891410-16893410 | 145 | 159 | + | 3.3571 | 7.12E-05 | 0.795 | ACAAGGGTCACGCTG |
| 1 | GMTACGGTCACACTG | chr3L:22073523-22075523 | 146 | 160 | + | 3.3571 | 7.12E-05 | 0.795 | GCTAAGGACGAACTG |
| 1 | GMTACGGTCACACTG | chr3R:4228238-4230238 | 148 | 162 | + | 3.3571 | 7.12E-05 | 0.795 | GCCAGGGTTGCACTG |
| 1 | GMTACGGTCACACTG | chr3L:4092207-4094207 | 155 | 169 | + | 3.3571 | 7.12E-05 | 0.795 | GAACCGATCACACGG |
| 1 | GMTACGGTCACACTG | chrX:16650376-16652376 | 162 | 176 | + | 3.3571 | 7.12E-05 | 0.795 | GCCACTGGCACAGTG |
| 1 | GMTACGGTCACACTG | chrX:16650376-16652376 | 162 | 176 | + | 3.3571 | 7.12E-05 | 0.795 | GCCACTGGCACAGTG |
| 1 | GMTACGGTCACACTG | chr2R:5352994-5354994 | 166 | 180 | + | 3.3571 | 7.12E-05 | 0.795 | GCTGCGCTTACTCTG |
| 1 | GMTACGGTCACACTG | chr2R:14514211-14516211 | 168 | 182 | + | 3.3571 | 7.12E-05 | 0.795 | GCAACGGTCGCAGTT |
| 1 | GMTACGGTCACACTG | chr3L:19485498-19487498 | 169 | 183 | + | 3.3571 | 7.12E-05 | 0.795 | GCAACAGAAACACTG |
| 1 | GMTACGGTCACACTG | chr3L:8239601-8241601 | 170 | 184 | + | 3.3571 | 7.12E-05 | 0.795 | TCAACGGTCATCCTG |
| 1 | GMTACGGTCACACTG | chr3L:5629983-5631983 | 171 | 185 | + | 3.3571 | 7.12E-05 | 0.795 | ACTGCGGGTACACTG |
| 1 | GMTACGGTCACACTG | chr2L:14743421-14745421 | 172 | 186 | + | 3.3571 | 7.12E-05 | 0.795 | GATTCGGACACGGTG |
| 1 | GMTACGGTCACACTG | chr3R:17133197-17135197 | 173 | 187 | + | 3.3571 | 7.12E-05 | 0.795 | GCCATGGCGACACTG |
| 1 | GMTACGGTCACACTG | chr3L:25119709-25121709 | 174 | 188 | + | 3.3571 | 7.12E-05 | 0.795 | TCGACAGTCGCACTG |
| 1 | GMTACGGTCACACTG | chr3R:5959987-5961987 | 174 | 188 | + | 3.3571 | 7.12E-05 | 0.795 | GCTTCAGGCAAACTG |
| 1 | GMTACGGTCACACTG | chr2R:15339472-15341472 | 177 | 191 | + | 3.3571 | 7.12E-05 | 0.795 | GCTACCGTCCCATAG |
| 1 | GMTACGGTCACACTG | chr2R:15541661-15543661 | 179 | 193 | + | 3.3571 | 7.12E-05 | 0.795 | GCTGGTGTCACTCTG |
| 1 | GMTACGGTCACACTG | chrY:3202123-3204123 | 180 | 194 | + | 3.3571 | 7.12E-05 | 0.795 | GCTGCGCTCACACAA |
| 1 | GMTACGGTCACACTG | chr3R:24187896-24189896 | 181 | 195 | + | 3.3571 | 7.12E-05 | 0.795 | TCCACCGTCACACGG |
| 1 | GMTACGGTCACACTG | chr3R:9400495-9402495 | 182 | 196 | + | 3.3571 | 7.12E-05 | 0.795 | GATACGCTCGCGCAG |
| 1 | GMTACGGTCACACTG | chr2R:20551090-20553090 | 186 | 200 | + | 3.3571 | 7.12E-05 | 0.795 | GCTACTGCCAGTCTG |
| 1 | GMTACGGTCACACTG | chr2R:20551090-20553090 | 186 | 200 | + | 3.3571 | 7.12E-05 | 0.795 | GCTACTGCCAGTCTG |
| 1 | GMTACGGTCACACTG | chr3L:204844-206844 | 186 | 200 | + | 3.3571 | 7.12E-05 | 0.795 | GCCACCGTCACTGTG |
| 1 | GMTACGGTCACACTG | chr2L:8527618-8529618 | 187 | 201 | + | 3.3571 | 7.12E-05 | 0.795 | GCTACGGTCAGTGCG |
| 1 | GMTACGGTCACACTG | chr2L:8527618-8529618 | 187 | 201 | + | 3.3571 | 7.12E-05 | 0.795 | GCTACGGTCAGTGCG |
| 1 | GMTACGGTCACACTG | chr3R:30038533-30040533 | 192 | 206 | + | 3.3571 | 7.12E-05 | 0.795 | GCTACAGACTAACTG |
| 1 | GMTACGGTCACACTG | chr3R:24944237-24946237 | 194 | 208 | + | 3.3571 | 7.12E-05 | 0.795 | GAGCCCGCCACACTG |

| | | | | | | | | | |
|---|---|---|---|---|---|---|---|---|---|
| 1 | GMTACGGTCACACTG | chrX:6523323-6525323 | 197 | 211 | + | 3.3571 | 7.12E-05 | 0.795 | ACATTGGTCACACTG |
| 1 | GMTACGGTCACACTG | chr3L:17424477-17426477 | 198 | 212 | + | 3.3571 | 7.12E-05 | 0.795 | GCTGCGGGCACATTC |
| 1 | GMTACGGTCACACTG | chr2R:20246624-20248624 | 200 | 214 | + | 3.3571 | 7.12E-05 | 0.795 | GCTCTGGTTGCACTG |
| 1 | GMTACGGTCACACTG | chr3R:25677683-25679683 | 201 | 215 | + | 3.3571 | 7.12E-05 | 0.795 | AACCCGGTCGCACTG |
| 1 | GMTACGGTCACACTG | chr3R:25677682-25679682 | 202 | 216 | + | 3.3571 | 7.12E-05 | 0.795 | AACCCGGTCGCACTG |
| 1 | GMTACGGTCACACTG | chr2L:8437169-8439169 | 204 | 218 | + | 3.3571 | 7.12E-05 | 0.795 | GATGCGTGCACGCTG |
| 1 | GMTACGGTCACACTG | chrX:4116020-4118020 | 206 | 220 | + | 3.3571 | 7.12E-05 | 0.795 | GCTGCGGCCAGATTG |
| 1 | GMTACGGTCACACTG | chr3R:20483588-20485588 | 207 | 221 | + | 3.3571 | 7.12E-05 | 0.795 | GCTCCGGCCAGAGTG |
| 1 | GMTACGGTCACACTG | chr3L:18131689-18133689 | 209 | 223 | + | 3.3571 | 7.12E-05 | 0.795 | GATGCGGTCCCACCA |
| 1 | GMTACGGTCACACTG | chr3L:21820997-21822997 | 211 | 225 | + | 3.3571 | 7.12E-05 | 0.795 | GCTACTGTTCCGCTG |
| 1 | GMTACGGTCACACTG | chr2R:16866714-16868714 | 213 | 227 | + | 3.3571 | 7.12E-05 | 0.795 | GATTCGGGCAGACGG |
| 1 | GMTACGGTCACACTG | chrX:21250148-21252148 | 214 | 228 | + | 3.3571 | 7.12E-05 | 0.795 | GCTGGGTTCTCACTG |
| 1 | GMTACGGTCACACTG | chrX:18647182-18649182 | 214 | 228 | + | 3.3571 | 7.12E-05 | 0.795 | GCTGCTCGCACACTG |
| 1 | GMTACGGTCACACTG | chr3R:6126955-6128955 | 217 | 231 | + | 3.3571 | 7.12E-05 | 0.795 | GATACGTGCACTCAG |
| 1 | GMTACGGTCACACTG | chr2L:12924611-12926611 | 219 | 233 | + | 3.3571 | 7.12E-05 | 0.795 | GCTACGTTTGCTCTG |
| 1 | GMTACGGTCACACTG | chr2L:20952-22952 | 219 | 233 | + | 3.3571 | 7.12E-05 | 0.795 | GAAACGCCCGCACTG |
| 1 | GMTACGGTCACACTG | chr2R:24431733-24433733 | 219 | 233 | + | 3.3571 | 7.12E-05 | 0.795 | CAGACGGCCACACAG |
| 1 | GMTACGGTCACACTG | chr3L:12502894-12504894 | 220 | 234 | + | 3.3571 | 7.12E-05 | 0.795 | GCTCCAGGCACAATG |
| 1 | GMTACGGTCACACTG | chr3R:20713577-20715577 | 220 | 234 | + | 3.3571 | 7.12E-05 | 0.795 | GCAACGGCAACATTG |
| 1 | GMTACGGTCACACTG | chr3R:16430940-16432940 | 222 | 236 | + | 3.3571 | 7.12E-05 | 0.795 | GAAACTGTCTCCCTG |
| 1 | GMTACGGTCACACTG | chr2L:3465157-3467157 | 231 | 245 | + | 3.3571 | 7.12E-05 | 0.795 | GCAACCCTCGCACTG |
| 1 | GMTACGGTCACACTG | chr2R:13963653-13965653 | 234 | 248 | + | 3.3571 | 7.12E-05 | 0.795 | GCAACTGTGGCACTG |
| 1 | GMTACGGTCACACTG | chrX:10157434-10159434 | 234 | 248 | + | 3.3571 | 7.12E-05 | 0.795 | TCACAGGTCACACTG |
| 1 | GMTACGGTCACACTG | chr2L:4453453-4455453 | 235 | 249 | + | 3.3571 | 7.12E-05 | 0.795 | GCTGCAGTCATACCG |
| 1 | GMTACGGTCACACTG | chr2L:8399421-8401421 | 235 | 249 | + | 3.3571 | 7.12E-05 | 0.795 | GACCGGGTCACGCTG |
| 1 | GMTACGGTCACACTG | chr2R:23904334-23906334 | 237 | 251 | + | 3.3571 | 7.12E-05 | 0.795 | CCTACGGCCACACCA |
| 1 | GMTACGGTCACACTG | chr3R:21139739-21141739 | 238 | 252 | + | 3.3571 | 7.12E-05 | 0.795 | GCTACGGCCAAAAAG |
| 1 | GMTACGGTCACACTG | chr2L:2252045-2254045 | 242 | 256 | + | 3.3571 | 7.12E-05 | 0.795 | ACCACGGTGACGCTG |
| 1 | GMTACGGTCACACTG | chr3R:5636234-5638234 | 242 | 256 | + | 3.3571 | 7.12E-05 | 0.795 | GACGCGGTCGTACTG |
| 1 | GMTACGGTCACACTG | chr2R:18769131-18771131 | 243 | 257 | + | 3.3571 | 7.12E-05 | 0.795 | GCACTGGTCACAATG |
| 1 | GMTACGGTCACACTG | chr3L:17739540-17741540 | 244 | 258 | + | 3.3571 | 7.12E-05 | 0.795 | GCCACTGGCAAACTG |
| 1 | GMTACGGTCACACTG | chr3L:1841084-1843084 | 245 | 259 | + | 3.3571 | 7.12E-05 | 0.795 | GCCACAGTGACCCTG |
| 1 | GMTACGGTCACACTG | chr2R:13158141-13160141 | 245 | 259 | + | 3.3571 | 7.12E-05 | 0.795 | AAAACGGTCACGCAG |

| | | | | | | | | | |
|---|---|---|---|---|---|---|---|---|---|
| 1 | GMTACGGTCACACTG | chrX:15670029-15672029 | 246 | 260 | + | 3.3571 | 7.12E-05 | 0.795 | GCTGCCGGCACACTC |
| 1 | GMTACGGTCACACTG | chr3L:21135353-21137353 | 248 | 262 | + | 3.3571 | 7.12E-05 | 0.795 | GCTGCCGTCGCACTA |
| 1 | GMTACGGTCACACTG | chr3R:26534217-26536217 | 250 | 264 | + | 3.3571 | 7.12E-05 | 0.795 | GCTACGCTCTTGCTG |
| 1 | GMTACGGTCACACTG | chr2L:10869867-10871867 | 251 | 265 | + | 3.3571 | 7.12E-05 | 0.795 | CCAACGATCACACAG |
| 1 | GMTACGGTCACACTG | chr3R:6834869-6836869 | 253 | 267 | + | 3.3571 | 7.12E-05 | 0.795 | GACCAGGTCCCACTG |
| 1 | GMTACGGTCACACTG | chr3L:4364434-4366434 | 255 | 269 | + | 3.3571 | 7.12E-05 | 0.795 | TCACCGGTCACAATG |
| 1 | GMTACGGTCACACTG | chr2L:9896552-9898552 | 258 | 272 | + | 3.3571 | 7.12E-05 | 0.795 | GAAACGGGTACGCTG |
| 1 | GMTACGGTCACACTG | chr2L:4346501-4348501 | 259 | 273 | + | 3.3571 | 7.12E-05 | 0.795 | GCCACTCTCACTCTG |
| 1 | GMTACGGTCACACTG | chr2L:2517037-2519037 | 262 | 276 | + | 3.3571 | 7.12E-05 | 0.795 | GAGGCGTTCACACAG |
| 1 | GMTACGGTCACACTG | chrX:1205737-1207737 | 262 | 276 | + | 3.3571 | 7.12E-05 | 0.795 | GCTGAGGGCAAACTG |
| 1 | GMTACGGTCACACTG | chr2R:12933045-12935045 | 262 | 276 | + | 3.3571 | 7.12E-05 | 0.795 | GCTCCTGTCAACCTG |
| 1 | GMTACGGTCACACTG | chr2L:1988450-1990450 | 263 | 277 | + | 3.3571 | 7.12E-05 | 0.795 | TACACGGCCACCCTG |
| 1 | GMTACGGTCACACTG | chr3R:5624178-5626178 | 267 | 281 | + | 3.3571 | 7.12E-05 | 0.795 | GCGATGGTCGAACTG |
| 1 | GMTACGGTCACACTG | chr2L:18973710-18975710 | 267 | 281 | + | 3.3571 | 7.12E-05 | 0.795 | GCCACGCCCACATTG |
| 1 | GMTACGGTCACACTG | chr2R:13959675-13961675 | 267 | 281 | + | 3.3571 | 7.12E-05 | 0.795 | GCCATCGTCACTCTG |
| 1 | GMTACGGTCACACTG | chr3R:29840657-29842657 | 270 | 284 | + | 3.3571 | 7.12E-05 | 0.795 | GCAACTGTCTCATTG |
| 1 | GMTACGGTCACACTG | chrX:646304-648304 | 270 | 284 | + | 3.3571 | 7.12E-05 | 0.795 | GCTACGCGCGCACTT |
| 1 | GMTACGGTCACACTG | chr3L:204844-206844 | 271 | 285 | + | 3.3571 | 7.12E-05 | 0.795 | ACTTCGGGCACAATG |
| 1 | GMTACGGTCACACTG | chr3L:15815410-15817410 | 273 | 287 | + | 3.3571 | 7.12E-05 | 0.795 | TCTCCGCTCTCACTG |
| 1 | GMTACGGTCACACTG | chr3L:12813254-12815254 | 273 | 287 | + | 3.3571 | 7.12E-05 | 0.795 | GCTGAGTTCACGCTG |
| 1 | GMTACGGTCACACTG | chrX:926975-928975 | 273 | 287 | + | 3.3571 | 7.12E-05 | 0.795 | GCTCCGGTCACAAAT |
| 1 | GMTACGGTCACACTG | chr2R:23432354-23434354 | 276 | 290 | + | 3.3571 | 7.12E-05 | 0.795 | GATACCGACTCGCTG |
| 1 | GMTACGGTCACACTG | chr2L:3465112-3467112 | 276 | 290 | + | 3.3571 | 7.12E-05 | 0.795 | GCAACCCTCGCACTG |
| 1 | GMTACGGTCACACTG | chr2R:24555300-24557300 | 281 | 295 | + | 3.3571 | 7.12E-05 | 0.795 | CCTAGGGTCCCACAG |
| 1 | GMTACGGTCACACTG | chr3L:7087787-7089787 | 284 | 298 | + | 3.3571 | 7.12E-05 | 0.795 | GCTACGGATGCAATG |
| 1 | GMTACGGTCACACTG | chrX:19889060-19891060 | 289 | 303 | + | 3.3571 | 7.12E-05 | 0.795 | TACACGGCCACACGG |
| 1 | GMTACGGTCACACTG | chr3L:15049209-15051209 | 289 | 303 | + | 3.3571 | 7.12E-05 | 0.795 | GCTCGGAGCACACTG |
| 1 | GMTACGGTCACACTG | chrX:478309-480309 | 291 | 305 | + | 3.3571 | 7.12E-05 | 0.795 | GCTCCCTTCACGCTG |
| 1 | GMTACGGTCACACTG | chr3L:19519945-19521945 | 291 | 305 | + | 3.3571 | 7.12E-05 | 0.795 | GATACGGACACGGAG |
| 1 | GMTACGGTCACACTG | chr3L:11773501-11775501 | 292 | 306 | + | 3.3571 | 7.12E-05 | 0.795 | TCTCCGGGCAAACTG |
| 1 | GMTACGGTCACACTG | chr2L:19107427-19109427 | 296 | 310 | + | 3.3571 | 7.12E-05 | 0.795 | GACGCGGCAACACTG |
| 1 | GMTACGGTCACACTG | chr2R:19646645-19648645 | 297 | 311 | + | 3.3571 | 7.12E-05 | 0.795 | GCTGCAGCCATACTG |
| 1 | GMTACGGTCACACTG | chr3R:18664361-18666361 | 299 | 313 | + | 3.3571 | 7.12E-05 | 0.795 | GCCATGCACACACTG |

| | | | | | | | | | |
|---|---|---|---|---|---|---|---|---|---|
| 1 | GMTACGGTCACACTG | chr2R:22113997-22115997 | 303 | 317 | + | 3.3571 | 7.12E-05 | 0.795 | CCTCCGCTCACACCG |
| 1 | GMTACGGTCACACTG | chrX:12567452-12569452 | 306 | 320 | + | 3.3571 | 7.12E-05 | 0.795 | GACACTGTCTCTCTG |
| 1 | GMTACGGTCACACTG | chr3L:16136879-16138879 | 309 | 323 | + | 3.3571 | 7.12E-05 | 0.795 | GACTCGGAAACACTG |
| 1 | GMTACGGTCACACTG | chr3R:25183383-25185383 | 310 | 324 | + | 3.3571 | 7.12E-05 | 0.795 | GCTGCTCTCTCACTG |
| 1 | GMTACGGTCACACTG | chr3L:16662903-16664903 | 314 | 328 | + | 3.3571 | 7.12E-05 | 0.795 | GATTCTGCCACACGG |
| 1 | GMTACGGTCACACTG | chr3R:8032009-8034009 | 315 | 329 | + | 3.3571 | 7.12E-05 | 0.795 | TATACGGCCACTCCG |
| 1 | GMTACGGTCACACTG | chr3R:9676927-9678927 | 317 | 331 | + | 3.3571 | 7.12E-05 | 0.795 | GCAATGGTCACATGG |
| 1 | GMTACGGTCACACTG | chr3L:12953647-12955647 | 319 | 333 | + | 3.3571 | 7.12E-05 | 0.795 | GACTTGGTCACTCTG |
| 1 | GMTACGGTCACACTG | chr2L:17981472-17983472 | 322 | 336 | + | 3.3571 | 7.12E-05 | 0.795 | GCTGAGGTCACGATG |
| 1 | GMTACGGTCACACTG | chr3R:24332433-24334433 | 322 | 336 | + | 3.3571 | 7.12E-05 | 0.795 | GATGCGGCCCAACTG |
| 1 | GMTACGGTCACACTG | chr3L:20431468-20433468 | 322 | 336 | + | 3.3571 | 7.12E-05 | 0.795 | GCAAAGGTCACCATG |
| 1 | GMTACGGTCACACTG | chr3R:19633949-19635949 | 327 | 341 | + | 3.3571 | 7.12E-05 | 0.795 | GATTTGGTCACGCAG |
| 1 | GMTACGGTCACACTG | chr2L:10971280-10973280 | 330 | 344 | + | 3.3571 | 7.12E-05 | 0.795 | GACCCGATCACACAG |
| 1 | GMTACGGTCACACTG | chr2R:13959404-13961404 | 331 | 345 | + | 3.3571 | 7.12E-05 | 0.795 | GATGCGGCGGCACTG |
| 1 | GMTACGGTCACACTG | chr3R:22208477-22210477 | 331 | 345 | + | 3.3571 | 7.12E-05 | 0.795 | ACGCCGGTCACAATG |
| 1 | GMTACGGTCACACTG | chr3R:19876514-19878514 | 335 | 349 | + | 3.3571 | 7.12E-05 | 0.795 | GCTGTAGTCACACGG |
| 1 | GMTACGGTCACACTG | chr3L:23338326-23340326 | 335 | 349 | + | 3.3571 | 7.12E-05 | 0.795 | GCCACGCCCACACTT |
| 1 | GMTACGGTCACACTG | chr2L:20561298-20563298 | 335 | 349 | + | 3.3571 | 7.12E-05 | 0.795 | GCTTCGGTGATGCTG |
| 1 | GMTACGGTCACACTG | chr3L:19234373-19236373 | 337 | 351 | + | 3.3571 | 7.12E-05 | 0.795 | GCTGCGCTCACACAA |
| 1 | GMTACGGTCACACTG | chr2L:8070624-8072624 | 337 | 351 | + | 3.3571 | 7.12E-05 | 0.795 | TAGACGGACACCCTG |
| 1 | GMTACGGTCACACTG | chr3R:27581840-27583840 | 338 | 352 | + | 3.3571 | 7.12E-05 | 0.795 | GAATCGTTCACACAG |
| 1 | GMTACGGTCACACTG | chr3R:9704417-9706417 | 339 | 353 | + | 3.3571 | 7.12E-05 | 0.795 | GCTACTATCCCTCTG |
| 1 | GMTACGGTCACACTG | chrX:314123-316123 | 339 | 353 | + | 3.3571 | 7.12E-05 | 0.795 | GAGACGGACAGACGG |
| 1 | GMTACGGTCACACTG | chrX:13782714-13784714 | 340 | 354 | + | 3.3571 | 7.12E-05 | 0.795 | GCTGTGGGCAAACTG |
| 1 | GMTACGGTCACACTG | chrX:4557072-4559072 | 340 | 354 | + | 3.3571 | 7.12E-05 | 0.795 | GACACGGTCCCCATG |
| 1 | GMTACGGTCACACTG | chrX:7411668-7413668 | 342 | 356 | + | 3.3571 | 7.12E-05 | 0.795 | GCTCCGTTGACCCTG |
| 1 | GMTACGGTCACACTG | chr3L:14785791-14787791 | 343 | 357 | + | 3.3571 | 7.12E-05 | 0.795 | GCTACTGCGTCACTG |
| 1 | GMTACGGTCACACTG | chr2L:2213266-2215266 | 345 | 359 | + | 3.3571 | 7.12E-05 | 0.795 | CCTACGGTTTCGCTG |
| 1 | GMTACGGTCACACTG | chr2R:23701477-23703477 | 346 | 360 | + | 3.3571 | 7.12E-05 | 0.795 | GCTAAGGTCACGCGA |
| 1 | GMTACGGTCACACTG | chr3L:199174-201174 | 347 | 361 | + | 3.3571 | 7.12E-05 | 0.795 | GCCACTTTCGCACTG |
| 1 | GMTACGGTCACACTG | chr3L:15520554-15522554 | 350 | 364 | + | 3.3571 | 7.12E-05 | 0.795 | ACAACAGGCACACTG |
| 1 | GMTACGGTCACACTG | chr2L:18284720-18286720 | 352 | 366 | + | 3.3571 | 7.12E-05 | 0.795 | GATTCGGCCACGTTG |
| 1 | GMTACGGTCACACTG | chr3L:11521713-11523713 | 352 | 366 | + | 3.3571 | 7.12E-05 | 0.795 | GCGATGCGCACACTG |

| | | | | | | | | | |
|---|---|---|---|---|---|---|---|---|---|
| 1 | GMTACGGTCACACTG | chr3L:11521713-11523713 | 352 | 366 | + | 3.3571 | 7.12E-05 | 0.795 | GCGATGCGCACACTG |
| 1 | GMTACGGTCACACTG | chr3R:7074487-7076487 | 358 | 372 | + | 3.3571 | 7.12E-05 | 0.795 | GCTCCGGTCACTTTA |
| 1 | GMTACGGTCACACTG | chr3R:5595831-5597831 | 360 | 374 | + | 3.3571 | 7.12E-05 | 0.795 | GCTCCGGTTATGCTG |
| 1 | GMTACGGTCACACTG | chr3L:7438405-7440405 | 361 | 375 | + | 3.3571 | 7.12E-05 | 0.795 | GATTCGGCCCCATTG |
| 1 | GMTACGGTCACACTG | chr2R:16772915-16774915 | 361 | 375 | + | 3.3571 | 7.12E-05 | 0.795 | GCCACGGTCAGCGTG |
| 1 | GMTACGGTCACACTG | chr2R:6970637-6972637 | 363 | 377 | + | 3.3571 | 7.12E-05 | 0.795 | TCTGCGGTCACGGTG |
| 1 | GMTACGGTCACACTG | chr3R:12407316-12409316 | 363 | 377 | + | 3.3571 | 7.12E-05 | 0.795 | GCGAGGGTCCCAGTG |
| 1 | GMTACGGTCACACTG | chr2R:6970637-6972637 | 363 | 377 | + | 3.3571 | 7.12E-05 | 0.795 | TCTGCGGTCACGGTG |
| 1 | GMTACGGTCACACTG | chr2L:6347427-6349427 | 371 | 385 | + | 3.3571 | 7.12E-05 | 0.795 | TCTCCGGTTGCACTG |
| 1 | GMTACGGTCACACTG | chr2L:15961435-15963435 | 375 | 389 | + | 3.3571 | 7.12E-05 | 0.795 | GCTACGGGGCCAATG |
| 1 | GMTACGGTCACACTG | chr3L:12754049-12756049 | 375 | 389 | + | 3.3571 | 7.12E-05 | 0.795 | GCTTAGGACACACTT |
| 1 | GMTACGGTCACACTG | chrX:1680566-1682566 | 380 | 394 | + | 3.3571 | 7.12E-05 | 0.795 | GAGACGGACAGACGG |
| 1 | GMTACGGTCACACTG | chrX:17090854-17092854 | 385 | 399 | + | 3.3571 | 7.12E-05 | 0.795 | GCTATGCGCACTCTG |
| 1 | GMTACGGTCACACTG | chrX:17090854-17092854 | 385 | 399 | + | 3.3571 | 7.12E-05 | 0.795 | GCTATGCGCACTCTG |
| 1 | GMTACGGTCACACTG | chr2L:11828405-11830405 | 386 | 400 | + | 3.3571 | 7.12E-05 | 0.795 | TAGACGGTCACCCGG |
| 1 | GMTACGGTCACACTG | chr2L:20561247-20563247 | 386 | 400 | + | 3.3571 | 7.12E-05 | 0.795 | GCTTCGGTGATGCTG |
| 1 | GMTACGGTCACACTG | chr3R:28025615-28027615 | 390 | 404 | + | 3.3571 | 7.12E-05 | 0.795 | AACACGGGCGCACTG |
| 1 | GMTACGGTCACACTG | chrX:20393395-20395395 | 398 | 412 | + | 3.3571 | 7.12E-05 | 0.795 | GAACTGGTCACCCTG |
| 1 | GMTACGGTCACACTG | chr3R:16430940-16432940 | 399 | 413 | + | 3.3571 | 7.12E-05 | 0.795 | GCTACGGATGCAGTG |
| 1 | GMTACGGTCACACTG | chr3R:15551833-15553833 | 399 | 413 | + | 3.3571 | 7.12E-05 | 0.795 | TCCATGGTCACTCTG |
| 1 | GMTACGGTCACACTG | chr3R:30486928-30488928 | 400 | 414 | + | 3.3571 | 7.12E-05 | 0.795 | GCTGCAATCACTCTG |
| 1 | GMTACGGTCACACTG | chr2L:2213211-2215211 | 400 | 414 | + | 3.3571 | 7.12E-05 | 0.795 | CCTACGGTTTCGCTG |
| 1 | GMTACGGTCACACTG | chr2L:19130954-19132954 | 401 | 415 | + | 3.3571 | 7.12E-05 | 0.795 | GCCACCGGCAGACTG |
| 1 | GMTACGGTCACACTG | chr3L:8136498-8138498 | 404 | 418 | + | 3.3571 | 7.12E-05 | 0.795 | GCTACTGCCAGACGG |
| 1 | GMTACGGTCACACTG | chr3R:8771691-8773691 | 404 | 418 | + | 3.3571 | 7.12E-05 | 0.795 | CCTCCGACCACACTG |
| 1 | GMTACGGTCACACTG | chr3L:9797420-9799420 | 405 | 419 | + | 3.3571 | 7.12E-05 | 0.795 | GAGGCTGCCACACTG |
| 1 | GMTACGGTCACACTG | chr2L:14517377-14519377 | 406 | 420 | + | 3.3571 | 7.12E-05 | 0.795 | GCTACTGTCAGTCGG |
| 1 | GMTACGGTCACACTG | chr3R:22546846-22548846 | 409 | 423 | + | 3.3571 | 7.12E-05 | 0.795 | GCTCCGATCGCATTG |
| 1 | GMTACGGTCACACTG | chr3R:8032301-8034301 | 410 | 424 | + | 3.3571 | 7.12E-05 | 0.795 | GCTGCGTTCTCACTT |
| 1 | GMTACGGTCACACTG | chr3R:11948129-11950129 | 411 | 425 | + | 3.3571 | 7.12E-05 | 0.795 | GCAACAGTTGCACTG |
| 1 | GMTACGGTCACACTG | chr2R:24637588-24639588 | 411 | 425 | + | 3.3571 | 7.12E-05 | 0.795 | TCTCCGACCACACTG |
| 1 | GMTACGGTCACACTG | chr2R:23701406-23703406 | 417 | 431 | + | 3.3571 | 7.12E-05 | 0.795 | GCTAAGGTCACGCGA |
| 1 | GMTACGGTCACACTG | chr3L:20205046-20207046 | 418 | 432 | + | 3.3571 | 7.12E-05 | 0.795 | GCGACCGACACAGTG |

| | | | | | | | | | |
|---|---|---|---|---|---|---|---|---|---|
| 1 | GMTACGGTCACACTG | chrX:19178830-19180830 | 423 | 437 | + | 3.3571 | 7.12E-05 | 0.795 | GATACTGGCCCACGG |
| 1 | GMTACGGTCACACTG | chr3L:20431361-20433361 | 429 | 443 | + | 3.3571 | 7.12E-05 | 0.795 | GCAAAGGTCACCATG |
| 1 | GMTACGGTCACACTG | chr3R:6834690-6836690 | 432 | 446 | + | 3.3571 | 7.12E-05 | 0.795 | GACCAGGTCCCACTG |
| 1 | GMTACGGTCACACTG | chr2L:1728887-1730887 | 433 | 447 | + | 3.3571 | 7.12E-05 | 0.795 | GCTGCGGGAAAACTG |
| 1 | GMTACGGTCACACTG | chr3L:23338228-23340228 | 433 | 447 | + | 3.3571 | 7.12E-05 | 0.795 | GCCACGCCCACACTT |
| 1 | GMTACGGTCACACTG | chrX:7306939-7308939 | 435 | 449 | + | 3.3571 | 7.12E-05 | 0.795 | GAATTGGTCCCACTG |
| 1 | GMTACGGTCACACTG | chr3R:24332316-24334316 | 439 | 453 | + | 3.3571 | 7.12E-05 | 0.795 | GATGCGGCCCAACTG |
| 1 | GMTACGGTCACACTG | chr3R:28969541-28971541 | 440 | 454 | + | 3.3571 | 7.12E-05 | 0.795 | GCCAGTGCCACACTG |
| 1 | GMTACGGTCACACTG | chr2L:6129651-6131651 | 445 | 459 | + | 3.3571 | 7.12E-05 | 0.795 | GAAACGCTCACGCAG |
| 1 | GMTACGGTCACACTG | chr2L:6129651-6131651 | 445 | 459 | + | 3.3571 | 7.12E-05 | 0.795 | GAAACGCTCACGCAG |
| 1 | GMTACGGTCACACTG | chr2R:18186707-18188707 | 446 | 460 | + | 3.3571 | 7.12E-05 | 0.795 | GCTGTGGGAACACTG |
| 1 | GMTACGGTCACACTG | chr3R:17020120-17022120 | 446 | 460 | + | 3.3571 | 7.12E-05 | 0.795 | GCTCTGGGCAAACTG |
| 1 | GMTACGGTCACACTG | chr2R:8130431-8132431 | 449 | 463 | + | 3.3571 | 7.12E-05 | 0.795 | GCTGGCGACACACTG |
| 1 | GMTACGGTCACACTG | chr3R:9400226-9402226 | 451 | 465 | + | 3.3571 | 7.12E-05 | 0.795 | GATACGCTCGCGCAG |
| 1 | GMTACGGTCACACTG | chr2R:9890955-9892955 | 456 | 470 | + | 3.3571 | 7.12E-05 | 0.795 | GCTCCCGTGACACGG |
| 1 | GMTACGGTCACACTG | chr3L:17424217-17426217 | 458 | 472 | + | 3.3571 | 7.12E-05 | 0.795 | GCTGCGGGCACATTC |
| 1 | GMTACGGTCACACTG | chr3R:15792782-15794782 | 462 | 476 | + | 3.3571 | 7.12E-05 | 0.795 | GCTAAGGTCTCAAAG |
| 1 | GMTACGGTCACACTG | chr2L:20839750-20841750 | 464 | 478 | + | 3.3571 | 7.12E-05 | 0.795 | CATGCGGACACACGG |
| 1 | GMTACGGTCACACTG | chr2R:19897695-19899695 | 467 | 481 | + | 3.3571 | 7.12E-05 | 0.795 | GCAAGGGGCACACTA |
| 1 | GMTACGGTCACACTG | chr2L:2390660-2392660 | 469 | 483 | + | 3.3571 | 7.12E-05 | 0.795 | GCGACGGCGACACTC |
| 1 | GMTACGGTCACACTG | chrX:19734570-19736570 | 469 | 483 | + | 3.3571 | 7.12E-05 | 0.795 | GCCACGCCCACATTG |
| 1 | GMTACGGTCACACTG | chrX:8441348-8443348 | 472 | 486 | + | 3.3571 | 7.12E-05 | 0.795 | GATGCGCTCCCACAG |
| 1 | GMTACGGTCACACTG | chr3L:8674759-8676759 | 473 | 487 | + | 3.3571 | 7.12E-05 | 0.795 | GAACCTGTCGCACTG |
| 1 | GMTACGGTCACACTG | chr3R:27678291-27680291 | 473 | 487 | + | 3.3571 | 7.12E-05 | 0.795 | GCCACGCTCACCATG |
| 1 | GMTACGGTCACACTG | chr3R:13185496-13187496 | 473 | 487 | + | 3.3571 | 7.12E-05 | 0.795 | GCTCCCGTCGCATTG |
| 1 | GMTACGGTCACACTG | chr3R:24943956-24945956 | 475 | 489 | + | 3.3571 | 7.12E-05 | 0.795 | GAGCCCGCCACACTG |
| 1 | GMTACGGTCACACTG | chr2R:23751754-23753754 | 477 | 491 | + | 3.3571 | 7.12E-05 | 0.795 | GCTACGCCCACATCG |
| 1 | GMTACGGTCACACTG | chr3L:5776333-5778333 | 478 | 492 | + | 3.3571 | 7.12E-05 | 0.795 | GCTCTGGTCAGCCTG |
| 1 | GMTACGGTCACACTG | chr2L:19026710-19028710 | 480 | 494 | + | 3.3571 | 7.12E-05 | 0.795 | GACATGGTCACGCCG |
| 1 | GMTACGGTCACACTG | chr3R:24529576-24531576 | 486 | 500 | + | 3.3571 | 7.12E-05 | 0.795 | TCGACGGTCACACGC |
| 1 | GMTACGGTCACACTG | chrX:19207187-19209187 | 487 | 501 | + | 3.3571 | 7.12E-05 | 0.795 | GCTGCCGACACATTG |
| 1 | GMTACGGTCACACTG | chr3R:28339334-28341334 | 491 | 505 | + | 3.3571 | 7.12E-05 | 0.795 | GCTACAGCCACACGC |
| 1 | GMTACGGTCACACTG | chr2R:9312237-9314237 | 497 | 511 | + | 3.3571 | 7.12E-05 | 0.795 | TCTCCTGTCGCACTG |

| | | | | | | | | | |
|---|---|---|---|---|---|---|---|---|---|
| 1 | GMTACGGTCACACTG | chr3R:18725434-18727434 | 498 | 512 | + | 3.3571 | 7.12E-05 | 0.795 | GAGATGGGCACACAG |
| 1 | GMTACGGTCACACTG | chr3L:1436872-1438872 | 498 | 512 | + | 3.3571 | 7.12E-05 | 0.795 | GCCACGCTCTCAATG |
| 1 | GMTACGGTCACACTG | chr2L:856542-858542 | 499 | 513 | + | 3.3571 | 7.12E-05 | 0.795 | GCTCCGTTTCCACTG |
| 1 | GMTACGGTCACACTG | chr3L:22071369-22073369 | 502 | 516 | + | 3.3571 | 7.12E-05 | 0.795 | GCTATGGTCACGCGT |
| 1 | GMTACGGTCACACTG | chr3R:24529558-24531558 | 504 | 518 | + | 3.3571 | 7.12E-05 | 0.795 | TCGACGGTCACACGC |
| 1 | GMTACGGTCACACTG | chr2R:3525047-3527047 | 505 | 519 | + | 3.3571 | 7.12E-05 | 0.795 | GAGACGTTCACGCAG |
| 1 | GMTACGGTCACACTG | chr2R:20264145-20266145 | 506 | 520 | + | 3.3571 | 7.12E-05 | 0.795 | GATCCGCTCCCACCG |
| 1 | GMTACGGTCACACTG | chr3R:22766699-22768699 | 507 | 521 | + | 3.3571 | 7.12E-05 | 0.795 | GCGACGGTCAAGCTC |
| 1 | GMTACGGTCACACTG | chrX:6672973-6674973 | 507 | 521 | + | 3.3571 | 7.12E-05 | 0.795 | GATGCGGGCACAGGG |
| 1 | GMTACGGTCACACTG | chr3L:12813019-12815019 | 508 | 522 | + | 3.3571 | 7.12E-05 | 0.795 | GCTGAGTTCACGCTG |
| 1 | GMTACGGTCACACTG | chr2R:16765804-16767804 | 510 | 524 | + | 3.3571 | 7.12E-05 | 0.795 | GCTGCCGGCACAGTG |
| 1 | GMTACGGTCACACTG | chr3L:18800312-18802312 | 511 | 525 | + | 3.3571 | 7.12E-05 | 0.795 | GACACGCTCGCACCG |
| 1 | GMTACGGTCACACTG | chrX:2186820-2188820 | 512 | 526 | + | 3.3571 | 7.12E-05 | 0.795 | GACACGGTCACTACG |
| 1 | GMTACGGTCACACTG | chr3L:18800309-18802309 | 514 | 528 | + | 3.3571 | 7.12E-05 | 0.795 | GACACGCTCGCACCG |
| 1 | GMTACGGTCACACTG | chr3R:31394660-31396660 | 516 | 530 | + | 3.3571 | 7.12E-05 | 0.795 | GCAAAGGGAACACTG |
| 1 | GMTACGGTCACACTG | chr3L:1334359-1336359 | 517 | 531 | + | 3.3571 | 7.12E-05 | 0.795 | CCCGCGGTCACACTT |
| 1 | GMTACGGTCACACTG | chr2L:11497948-11499948 | 520 | 534 | + | 3.3571 | 7.12E-05 | 0.795 | GAAGCGTTCGCACTG |
| 1 | GMTACGGTCACACTG | chr3R:25995996-25997996 | 524 | 538 | + | 3.3571 | 7.12E-05 | 0.795 | GAAGAGGCCACACTG |
| 1 | GMTACGGTCACACTG | chr3R:18251734-18253734 | 526 | 540 | + | 3.3571 | 7.12E-05 | 0.795 | GACTCCGTCACGCTG |
| 1 | GMTACGGTCACACTG | chr3R:15381951-15383951 | 526 | 540 | + | 3.3571 | 7.12E-05 | 0.795 | GATGCGGGCATCCTG |
| 1 | GMTACGGTCACACTG | chr2L:18973448-18975448 | 529 | 543 | + | 3.3571 | 7.12E-05 | 0.795 | GCCACGCCCACATTG |
| 1 | GMTACGGTCACACTG | chr2L:10421780-10423780 | 530 | 544 | + | 3.3571 | 7.12E-05 | 0.795 | GCAACGATCGCATTG |
| 1 | GMTACGGTCACACTG | chr3R:23732462-23734462 | 531 | 545 | + | 3.3571 | 7.12E-05 | 0.795 | GCAGCTGTCACATTG |
| 1 | GMTACGGTCACACTG | chr3R:5595658-5597658 | 533 | 547 | + | 3.3571 | 7.12E-05 | 0.795 | GCTCCGGTTATGCTG |
| 1 | GMTACGGTCACACTG | chrX:478066-480066 | 534 | 548 | + | 3.3571 | 7.12E-05 | 0.795 | GCTCCCTTCACGCTG |
| 1 | GMTACGGTCACACTG | chr3L:5768002-5770002 | 536 | 550 | + | 3.3571 | 7.12E-05 | 0.795 | GCGAAGGTCCAACTG |
| 1 | GMTACGGTCACACTG | chr3R:10074373-10076373 | 536 | 550 | + | 3.3571 | 7.12E-05 | 0.795 | GCTCGGGAAACACTG |
| 1 | GMTACGGTCACACTG | chr3R:14723261-14725261 | 538 | 552 | + | 3.3571 | 7.12E-05 | 0.795 | AACACGGCCACACGG |
| 1 | GMTACGGTCACACTG | chr2R:13959404-13961404 | 538 | 552 | + | 3.3571 | 7.12E-05 | 0.795 | GCCATCGTCACTCTG |
| 1 | GMTACGGTCACACTG | chr2L:17177772-17179772 | 540 | 554 | + | 3.3571 | 7.12E-05 | 0.795 | GATACGGACTGACGG |
| 1 | GMTACGGTCACACTG | chr2L:9911911-9913911 | 540 | 554 | + | 3.3571 | 7.12E-05 | 0.795 | GATGCGGCCACTGTG |
| 1 | GMTACGGTCACACTG | chr2L:21631587-21633587 | 541 | 555 | + | 3.3571 | 7.12E-05 | 0.795 | GCCACGTCCACACTT |
| 1 | GMTACGGTCACACTG | chr3L:6138974-6140974 | 542 | 556 | + | 3.3571 | 7.12E-05 | 0.795 | GAACCGCGCACACTG |

| | | | | | | | | | |
|---|---|---|---|---|---|---|---|---|---|
| 1 | GMTACGGTCACACTG | chr3R:29992411-29994411 | 544 | 558 | + | 3.3571 | 7.12E-05 | 0.795 | GCTACGCACACATAG |
| 1 | GMTACGGTCACACTG | chr2R:22059950-22061950 | 544 | 558 | + | 3.3571 | 7.12E-05 | 0.795 | GATGCGTCCACTCTG |
| 1 | GMTACGGTCACACTG | chrX:8093064-8095064 | 545 | 559 | + | 3.3571 | 7.12E-05 | 0.795 | GCGGGGGTCAAACTG |
| 1 | GMTACGGTCACACTG | chrX:6036892-6038892 | 549 | 563 | + | 3.3571 | 7.12E-05 | 0.795 | GAACTGGTCACACCG |
| 1 | GMTACGGTCACACTG | chr2L:4220796-4222796 | 550 | 564 | + | 3.3571 | 7.12E-05 | 0.795 | GCACCACTCACACTG |
| 1 | GMTACGGTCACACTG | chrX:11791533-11793533 | 553 | 567 | + | 3.3571 | 7.12E-05 | 0.795 | GCTTCGCTCTCACTC |
| 1 | GMTACGGTCACACTG | chr2R:8130323-8132323 | 557 | 571 | + | 3.3571 | 7.12E-05 | 0.795 | GCTGGCGACACACTG |
| 1 | GMTACGGTCACACTG | chr2R:8130323-8132323 | 557 | 571 | + | 3.3571 | 7.12E-05 | 0.795 | GCTGGCGACACACTG |
| 1 | GMTACGGTCACACTG | chr3R:3586054-3588054 | 558 | 572 | + | 3.3571 | 7.12E-05 | 0.795 | GCCACACGCACACTG |
| 1 | GMTACGGTCACACTG | chr2R:15693092-15695092 | 568 | 582 | + | 3.3571 | 7.12E-05 | 0.795 | GCTATGGTCCAACGG |
| 1 | GMTACGGTCACACTG | chrX:19478411-19480411 | 568 | 582 | + | 3.3571 | 7.12E-05 | 0.795 | GCTGCGGTGATACGG |
| 1 | GMTACGGTCACACTG | chrX:7306939-7308939 | 569 | 583 | + | 3.3571 | 7.12E-05 | 0.795 | CATTCGGTCCCACCG |
| 1 | GMTACGGTCACACTG | chr3L:2247635-2249635 | 571 | 585 | + | 3.3571 | 7.12E-05 | 0.795 | TCTGGGGACACACTG |
| 1 | GMTACGGTCACACTG | chr2L:14871022-14873022 | 572 | 586 | + | 3.3571 | 7.12E-05 | 0.795 | GCTCCGGCAATACTG |
| 1 | GMTACGGTCACACTG | chr3R:4960351-4962351 | 575 | 589 | + | 3.3571 | 7.12E-05 | 0.795 | GCGACGAGCACAATG |
| 1 | GMTACGGTCACACTG | chr3L:15815410-15817410 | 580 | 594 | + | 3.3571 | 7.12E-05 | 0.795 | GCTCCGGTCCAATTG |
| 1 | GMTACGGTCACACTG | chr3R:28601556-28603556 | 583 | 597 | + | 3.3571 | 7.12E-05 | 0.795 | GCTACGATTCCACAG |
| 1 | GMTACGGTCACACTG | chr2R:23839577-23841577 | 587 | 601 | + | 3.3571 | 7.12E-05 | 0.795 | GCTATGGACGCACTT |
| 1 | GMTACGGTCACACTG | chr3R:30875186-30877186 | 588 | 602 | + | 3.3571 | 7.12E-05 | 0.795 | CCTGCGGTCTCACTT |
| 1 | GMTACGGTCACACTG | chr3R:10875384-10877384 | 588 | 602 | + | 3.3571 | 7.12E-05 | 0.795 | GAAACGCCCACACGG |
| 1 | GMTACGGTCACACTG | chrX:18513867-18515867 | 588 | 602 | + | 3.3571 | 7.12E-05 | 0.795 | GCGACGCTCACCATG |
| 1 | GMTACGGTCACACTG | chr3R:24849627-24851627 | 595 | 609 | + | 3.3571 | 7.12E-05 | 0.795 | GAGACGGCCTAACTG |
| 1 | GMTACGGTCACACTG | chr3L:7389742-7391742 | 595 | 609 | + | 3.3571 | 7.12E-05 | 0.795 | GCGGCGATCACAATG |
| 1 | GMTACGGTCACACTG | chrX:11728099-11730099 | 597 | 611 | + | 3.3571 | 7.12E-05 | 0.795 | GCGACGGTCAGGATG |
| 1 | GMTACGGTCACACTG | chr2R:17126023-17128023 | 608 | 622 | + | 3.3571 | 7.12E-05 | 0.795 | GCGTCGCTCACATTG |
| 1 | GMTACGGTCACACTG | chr3R:28969372-28971372 | 609 | 623 | + | 3.3571 | 7.12E-05 | 0.795 | GCCAGTGCCACACTG |
| 1 | GMTACGGTCACACTG | chrX:6275724-6277724 | 609 | 623 | + | 3.3571 | 7.12E-05 | 0.795 | GATCCAGGCGCACTG |
| 1 | GMTACGGTCACACTG | chr3L:24536236-24538236 | 612 | 626 | + | 3.3571 | 7.12E-05 | 0.795 | GCCACGCGCACAATG |
| 1 | GMTACGGTCACACTG | chr2L:11112061-11114061 | 613 | 627 | + | 3.3571 | 7.12E-05 | 0.795 | GCTGGGGTCGTACTG |
| 1 | GMTACGGTCACACTG | chrX:436448-438448 | 616 | 630 | + | 3.3571 | 7.12E-05 | 0.795 | GATCCTGACACGCTG |
| 1 | GMTACGGTCACACTG | chr3R:25476003-25478003 | 617 | 631 | + | 3.3571 | 7.12E-05 | 0.795 | GATGCGGTCGCGATG |
| 1 | GMTACGGTCACACTG | chr2R:12923567-12925567 | 617 | 631 | + | 3.3571 | 7.12E-05 | 0.795 | GACACGGGCAGACCG |
| 1 | GMTACGGTCACACTG | chr2R:21764489-21766489 | 617 | 631 | + | 3.3571 | 7.12E-05 | 0.795 | GCCATGGCCACACTT |

| | | | | | | | | | |
|---|---|---|---|---|---|---|---|---|---|
| 1 | GMTACGGTCACACTG | chr2R:9312116-9314116 | 618 | 632 | + | 3.3571 | 7.12E-05 | 0.795 | TCTCCTGTCGCACTG |
| 1 | GMTACGGTCACACTG | chr3L:14097053-14099053 | 619 | 633 | + | 3.3571 | 7.12E-05 | 0.795 | GCCACGATAACGCTG |
| 1 | GMTACGGTCACACTG | chrX:16705874-16707874 | 623 | 637 | + | 3.3571 | 7.12E-05 | 0.795 | CCTACGGTCACGGCG |
| 1 | GMTACGGTCACACTG | chr3R:5635852-5637852 | 624 | 638 | + | 3.3571 | 7.12E-05 | 0.795 | GACGCGGTCGTACTG |
| 1 | GMTACGGTCACACTG | chr2R:16968395-16970395 | 625 | 639 | + | 3.3571 | 7.12E-05 | 0.795 | GCTGCGTTCTTACTG |
| 1 | GMTACGGTCACACTG | chr2L:559310-561310 | 632 | 646 | + | 3.3571 | 7.12E-05 | 0.795 | GAGGCGGCCAAACTG |
| 1 | GMTACGGTCACACTG | chr2L:856403-858403 | 638 | 652 | + | 3.3571 | 7.12E-05 | 0.795 | GCTCCGTTTCCACTG |
| 1 | GMTACGGTCACACTG | chrX:6036800-6038800 | 641 | 655 | + | 3.3571 | 7.12E-05 | 0.795 | GAACTGGTCACACCG |
| 1 | GMTACGGTCACACTG | chr3L:9509017-9511017 | 643 | 657 | + | 3.3571 | 7.12E-05 | 0.795 | TCTCTGGGCACACTG |
| 1 | GMTACGGTCACACTG | chr3L:8371405-8373405 | 646 | 660 | + | 3.3571 | 7.12E-05 | 0.795 | GCAGCGGTCACATTT |
| 1 | GMTACGGTCACACTG | chr3L:11832578-11834578 | 648 | 662 | + | 3.3571 | 7.12E-05 | 0.795 | TCGACGCTCACACCG |
| 1 | GMTACGGTCACACTG | chr3L:12753774-12755774 | 650 | 664 | + | 3.3571 | 7.12E-05 | 0.795 | GCTTAGGACACACTT |
| 1 | GMTACGGTCACACTG | chr2R:7792625-7794625 | 652 | 666 | + | 3.3571 | 7.12E-05 | 0.795 | GCGTAGGTCACACTT |
| 1 | GMTACGGTCACACTG | chr3R:9503656-9505656 | 655 | 669 | + | 3.3571 | 7.12E-05 | 0.795 | ACAGCCGTCACACTG |
| 1 | GMTACGGTCACACTG | chr3L:2585540-2587540 | 656 | 670 | + | 3.3571 | 7.12E-05 | 0.795 | ACTTCGCACACACTG |
| 1 | GMTACGGTCACACTG | chr2R:18245763-18247763 | 656 | 670 | + | 3.3571 | 7.12E-05 | 0.795 | GATCCGGACTGACTG |
| 1 | GMTACGGTCACACTG | chr2R:20995220-20997220 | 659 | 673 | + | 3.3571 | 7.12E-05 | 0.795 | GAGATGGCCACGCTG |
| 1 | GMTACGGTCACACTG | chrX:11348049-11350049 | 660 | 674 | + | 3.3571 | 7.12E-05 | 0.795 | GATGCGGACGCAATG |
| 1 | GMTACGGTCACACTG | chrX:926582-928582 | 666 | 680 | + | 3.3571 | 7.12E-05 | 0.795 | GCTCCGGTCACAAAT |
| 1 | GMTACGGTCACACTG | chr3R:23732324-23734324 | 669 | 683 | + | 3.3571 | 7.12E-05 | 0.795 | GCAGCTGTCACATTG |
| 1 | GMTACGGTCACACTG | chr3L:17469127-17471127 | 676 | 690 | + | 3.3571 | 7.12E-05 | 0.795 | GATCCGAACCCACTG |
| 1 | GMTACGGTCACACTG | chr3R:13010649-13012649 | 679 | 693 | + | 3.3571 | 7.12E-05 | 0.795 | TCAGCGGTCACACTT |
| 1 | GMTACGGTCACACTG | chr2R:18108985-18110985 | 681 | 695 | + | 3.3571 | 7.12E-05 | 0.795 | GCCACGATTGCACTG |
| 1 | GMTACGGTCACACTG | chr2L:18158272-18160272 | 682 | 696 | + | 3.3571 | 7.12E-05 | 0.795 | GCTTCGAACACAATG |
| 1 | GMTACGGTCACACTG | chr3R:5595509-5597509 | 682 | 696 | + | 3.3571 | 7.12E-05 | 0.795 | GCTCCGGTTATGCTG |
| 1 | GMTACGGTCACACTG | chr2L:10846044-10848044 | 682 | 696 | + | 3.3571 | 7.12E-05 | 0.795 | GAGCCGGTCACCTTG |
| 1 | GMTACGGTCACACTG | chr2L:1728636-1730636 | 684 | 698 | + | 3.3571 | 7.12E-05 | 0.795 | GCTGCGGGAAAACTG |
| 1 | GMTACGGTCACACTG | chr3R:15366665-15368665 | 686 | 700 | + | 3.3571 | 7.12E-05 | 0.795 | GCTGGAGTCACGCTG |
| 1 | GMTACGGTCACACTG | chr2R:9097539-9099539 | 689 | 703 | + | 3.3571 | 7.12E-05 | 0.795 | GCTACGGCCACCTTC |
| 1 | GMTACGGTCACACTG | chr2R:11902918-11904918 | 690 | 704 | + | 3.3571 | 7.12E-05 | 0.795 | GAGACGGTGCCGCTG |
| 1 | GMTACGGTCACACTG | chr2R:18004395-18006395 | 690 | 704 | + | 3.3571 | 7.12E-05 | 0.795 | GACATGGTCGCCCTG |
| 1 | GMTACGGTCACACTG | chr2L:14517091-14519091 | 692 | 706 | + | 3.3571 | 7.12E-05 | 0.795 | GCTACTGTCAGTCGG |
| 1 | GMTACGGTCACACTG | chr3R:21117856-21119856 | 693 | 707 | + | 3.3571 | 7.12E-05 | 0.795 | ACTGCGGTCAGGCTG |

| | | | | | | | | | |
|---|---|---|---|---|---|---|---|---|---|
| 1 | GMTACGGTCACACTG | chr3L:9410038-9412038 | 696 | 710 | + | 3.3571 | 7.12E-05 | 0.795 | CAGACGGACACACGG |
| 1 | GMTACGGTCACACTG | chr2R:6160535-6162535 | 699 | 713 | + | 3.3571 | 7.12E-05 | 0.795 | GCCACGCCCACACTT |
| 1 | GMTACGGTCACACTG | chr3L:12523014-12525014 | 699 | 713 | + | 3.3571 | 7.12E-05 | 0.795 | TCTGCCGGCACACTG |
| 1 | GMTACGGTCACACTG | chr3R:8032009-8034009 | 702 | 716 | + | 3.3571 | 7.12E-05 | 0.795 | GCTGCGTTCTCACTT |
| 1 | GMTACGGTCACACTG | chr2R:9097525-9099525 | 703 | 717 | + | 3.3571 | 7.12E-05 | 0.795 | GCTACGGCCACCTTC |
| 1 | GMTACGGTCACACTG | chr3L:10638740-10640740 | 704 | 718 | + | 3.3571 | 7.12E-05 | 0.795 | GATTCGGACACCCTC |
| 1 | GMTACGGTCACACTG | chr3R:25475915-25477915 | 705 | 719 | + | 3.3571 | 7.12E-05 | 0.795 | GATGCGGTCGCGATG |
| 1 | GMTACGGTCACACTG | chrX:20258239-20260239 | 709 | 723 | + | 3.3571 | 7.12E-05 | 0.795 | GCCACGCCCACACTT |
| 1 | GMTACGGTCACACTG | chr2R:8938789-8940789 | 709 | 723 | + | 3.3571 | 7.12E-05 | 0.795 | GCTGCGGACACATTC |
| 1 | GMTACGGTCACACTG | chr2L:401198-403198 | 712 | 726 | + | 3.3571 | 7.12E-05 | 0.795 | GCGACGATCACACAC |
| 1 | GMTACGGTCACACTG | chr3L:7346123-7348123 | 713 | 727 | + | 3.3571 | 7.12E-05 | 0.795 | GCAATGCTCACACAG |
| 1 | GMTACGGTCACACTG | chr4:582108-584108 | 718 | 732 | + | 3.3571 | 7.12E-05 | 0.795 | GACACGGACAGACGG |
| 1 | GMTACGGTCACACTG | chr3L:16609418-16611418 | 719 | 733 | + | 3.3571 | 7.12E-05 | 0.795 | GCGATGGCCACAATG |
| 1 | GMTACGGTCACACTG | chr2R:13157666-13159666 | 720 | 734 | + | 3.3571 | 7.12E-05 | 0.795 | AAAACGGTCACGCAG |
| 1 | GMTACGGTCACACTG | chr2L:16981483-16983483 | 720 | 734 | + | 3.3571 | 7.12E-05 | 0.795 | GCAATTGTCCCACTG |
| 1 | GMTACGGTCACACTG | chr3R:28592488-28594488 | 720 | 734 | + | 3.3571 | 7.12E-05 | 0.795 | GATGCGGCCACAAAG |
| 1 | GMTACGGTCACACTG | chr3R:17421202-17423202 | 721 | 735 | + | 3.3571 | 7.12E-05 | 0.795 | GATCCGGTCACTCCA |
| 1 | GMTACGGTCACACTG | chr3L:14785412-14787412 | 722 | 736 | + | 3.3571 | 7.12E-05 | 0.795 | GCTACTGCGTCACTG |
| 1 | GMTACGGTCACACTG | chr2R:9890689-9892689 | 722 | 736 | + | 3.3571 | 7.12E-05 | 0.795 | GCTCCCGTGACACGG |
| 1 | GMTACGGTCACACTG | chr3L:12522987-12524987 | 726 | 740 | + | 3.3571 | 7.12E-05 | 0.795 | TCTGCCGGCACACTG |
| 1 | GMTACGGTCACACTG | chr2R:19946196-19948196 | 729 | 743 | + | 3.3571 | 7.12E-05 | 0.795 | GCCACGCCCACACTT |
| 1 | GMTACGGTCACACTG | chr2R:20263920-20265920 | 731 | 745 | + | 3.3571 | 7.12E-05 | 0.795 | GATCCGCTCCCACCG |
| 1 | GMTACGGTCACACTG | chr2R:14214864-14216864 | 733 | 747 | + | 3.3571 | 7.12E-05 | 0.795 | GCTACCGTCACGGGG |
| 1 | GMTACGGTCACACTG | chr3L:1834164-1836164 | 733 | 747 | + | 3.3571 | 7.12E-05 | 0.795 | GCCACAGCAACACTG |
| 1 | GMTACGGTCACACTG | chr3R:6010932-6012932 | 735 | 749 | + | 3.3571 | 7.12E-05 | 0.795 | CCTTCCGCCACACTG |
| 1 | GMTACGGTCACACTG | chr3R:22380340-22382340 | 736 | 750 | + | 3.3571 | 7.12E-05 | 0.795 | CAAACGGACGCACTG |
| 1 | GMTACGGTCACACTG | chr2R:13942377-13944377 | 736 | 750 | + | 3.3571 | 7.12E-05 | 0.795 | GCCACGGTAACAAGG |
| 1 | GMTACGGTCACACTG | chr2L:5891862-5893862 | 742 | 756 | + | 3.3571 | 7.12E-05 | 0.795 | GCATCGTTCACACTA |
| 1 | GMTACGGTCACACTG | chr3L:5767794-5769794 | 744 | 758 | + | 3.3571 | 7.12E-05 | 0.795 | GCGAAGGTCCAACTG |
| 1 | GMTACGGTCACACTG | chrX:18646648-18648648 | 748 | 762 | + | 3.3571 | 7.12E-05 | 0.795 | GCTGCTCGCACACTG |
| 1 | GMTACGGTCACACTG | chr3L:6724279-6726279 | 750 | 764 | + | 3.3571 | 7.12E-05 | 0.795 | GCTGCGGTCAATCTA |
| 1 | GMTACGGTCACACTG | chr2L:9165778-9167778 | 755 | 769 | + | 3.3571 | 7.12E-05 | 0.795 | GCTGCAGTAGCACTG |
| 1 | GMTACGGTCACACTG | chr3R:21879810-21881810 | 756 | 770 | + | 3.3571 | 7.12E-05 | 0.795 | GATATGGTCTCCCAG |

| | | | | | | | | | |
|---|---|---|---|---|---|---|---|---|---|
| 1 | GMTACGGTCACACTG | chrX:13053218-13055218 | 756 | 770 | + | 3.3571 | 7.12E-05 | 0.795 | TCTGCGGTCTCATTG |
| 1 | GMTACGGTCACACTG | chrY:1882972-1884972 | 758 | 772 | + | 3.3571 | 7.12E-05 | 0.795 | TCTGCAGCCACACTG |
| 1 | GMTACGGTCACACTG | chr3L:198762-200762 | 759 | 773 | + | 3.3571 | 7.12E-05 | 0.795 | GCCACTTTCGCACTG |
| 1 | GMTACGGTCACACTG | chr3L:9858889-9860889 | 762 | 776 | + | 3.3571 | 7.12E-05 | 0.795 | ACTTTGGTCACTCTG |
| 1 | GMTACGGTCACACTG | chr2L:11011135-11013135 | 763 | 777 | + | 3.3571 | 7.12E-05 | 0.795 | GATTCGGTTCCACCG |
| 1 | GMTACGGTCACACTG | chr2R:20848939-20850939 | 764 | 778 | + | 3.3571 | 7.12E-05 | 0.795 | TATGCGGGCACTCTG |
| 1 | GMTACGGTCACACTG | chr3R:18282660-18284660 | 766 | 780 | + | 3.3571 | 7.12E-05 | 0.795 | GATGCGGAAGCACTG |
| 1 | GMTACGGTCACACTG | chr3L:17885645-17887645 | 768 | 782 | + | 3.3571 | 7.12E-05 | 0.795 | GCTCCGGTACCACTC |
| 1 | GMTACGGTCACACTG | chr2L:18707846-18709846 | 770 | 784 | + | 3.3571 | 7.12E-05 | 0.795 | GACGCGTTCTCACTG |
| 1 | GMTACGGTCACACTG | chr3R:22380340-22382340 | 772 | 786 | + | 3.3571 | 7.12E-05 | 0.795 | GAATCGCTCGCACTG |
| 1 | GMTACGGTCACACTG | chr3R:4161235-4163235 | 772 | 786 | + | 3.3571 | 7.12E-05 | 0.795 | GCGATGCTCCCACTG |
| 1 | GMTACGGTCACACTG | chr3R:16452821-16454821 | 773 | 787 | + | 3.3571 | 7.12E-05 | 0.795 | GCTGCAATCGCACTG |
| 1 | GMTACGGTCACACTG | chr2R:15692887-15694887 | 773 | 787 | + | 3.3571 | 7.12E-05 | 0.795 | GCTATGGTCCAACGG |
| 1 | GMTACGGTCACACTG | chr3L:13035821-13037821 | 779 | 793 | + | 3.3571 | 7.12E-05 | 0.795 | GCTACGCACATACCG |
| 1 | GMTACGGTCACACTG | chr2R:24056137-24058137 | 780 | 794 | + | 3.3571 | 7.12E-05 | 0.795 | ACTACGCCCACACCG |
| 1 | GMTACGGTCACACTG | chr3R:6233708-6235708 | 781 | 795 | + | 3.3571 | 7.12E-05 | 0.795 | GCTGCGGTCAGAAAG |
| 1 | GMTACGGTCACACTG | chr3L:19686637-19688637 | 787 | 801 | + | 3.3571 | 7.12E-05 | 0.795 | GACCTGGTCACCCTG |
| 1 | GMTACGGTCACACTG | chr2R:13666736-13668736 | 790 | 804 | + | 3.3571 | 7.12E-05 | 0.795 | GCCACCGTCACCGTG |
| 1 | GMTACGGTCACACTG | chr3R:27092143-27094143 | 791 | 805 | + | 3.3571 | 7.12E-05 | 0.795 | GATTTGGGCACCCTG |
| 1 | GMTACGGTCACACTG | chr3L:12470728-12472728 | 791 | 805 | + | 3.3571 | 7.12E-05 | 0.795 | CCTTCGGTCACACCA |
| 1 | GMTACGGTCACACTG | chr2R:11340423-11342423 | 792 | 806 | + | 3.3571 | 7.12E-05 | 0.795 | GCTATGGTCACCGCG |
| 1 | GMTACGGTCACACTG | chr3L:19585253-19587253 | 793 | 807 | + | 3.3571 | 7.12E-05 | 0.795 | TCTTTGGTCACCCTG |
| 1 | GMTACGGTCACACTG | chr2L:9704954-9706954 | 794 | 808 | + | 3.3571 | 7.12E-05 | 0.795 | GCAACTTTCACTCTG |
| 1 | GMTACGGTCACACTG | chr2L:20376-22376 | 795 | 809 | + | 3.3571 | 7.12E-05 | 0.795 | GAAACGCCCGCACTG |
| 1 | GMTACGGTCACACTG | chr3R:28601340-28603340 | 799 | 813 | + | 3.3571 | 7.12E-05 | 0.795 | GCTACGATTCCACAG |
| 1 | GMTACGGTCACACTG | chr3L:11561233-11563233 | 801 | 815 | + | 3.3571 | 7.12E-05 | 0.795 | GACACAGACACACGG |
| 1 | GMTACGGTCACACTG | chr2L:4700485-4702485 | 804 | 818 | + | 3.3571 | 7.12E-05 | 0.795 | GCCAGGGACAGACTG |
| 1 | GMTACGGTCACACTG | chr3R:23051916-23053916 | 809 | 823 | + | 3.3571 | 7.12E-05 | 0.795 | GCTACCGTCAGGCAG |
| 1 | GMTACGGTCACACTG | chr3L:5785718-5787718 | 810 | 824 | + | 3.3571 | 7.12E-05 | 0.795 | GATACGGGTTCTCTG |
| 1 | GMTACGGTCACACTG | chrX:18847566-18849566 | 811 | 825 | + | 3.3571 | 7.12E-05 | 0.795 | GCTCCGGTGAATCTG |
| 1 | GMTACGGTCACACTG | chr2R:21764295-21766295 | 811 | 825 | + | 3.3571 | 7.12E-05 | 0.795 | GCCATGGCCACACTT |
| 1 | GMTACGGTCACACTG | chr2L:9371102-9373102 | 811 | 825 | + | 3.3571 | 7.12E-05 | 0.795 | GATCCGGGCAAACGG |
| 1 | GMTACGGTCACACTG | chr2R:23377573-23379573 | 815 | 829 | + | 3.3571 | 7.12E-05 | 0.795 | GAACCCGTCACACGG |

| | | | | | | | | | |
|---|---|---|---|---|---|---|---|---|---|
| 1 | GMTACGGTCACACTG | chr3R:4428335-4430335 | 817 | 831 | + | 3.3571 | 7.12E-05 | 0.795 | GCGGCAGTCAAACTG |
| 1 | GMTACGGTCACACTG | chr3L:12626934-12628934 | 822 | 836 | + | 3.3571 | 7.12E-05 | 0.795 | GCAACTGTCTCACTT |
| 1 | GMTACGGTCACACTG | chrX:14783052-14785052 | 823 | 837 | + | 3.3571 | 7.12E-05 | 0.795 | GCTACTGTCCCACCA |
| 1 | GMTACGGTCACACTG | chr3R:24859366-24861366 | 841 | 855 | + | 3.3571 | 7.12E-05 | 0.795 | GCGACCGTCACGCTT |
| 1 | GMTACGGTCACACTG | chr2L:9911608-9913608 | 843 | 857 | + | 3.3571 | 7.12E-05 | 0.795 | GATGCGGCCACTGTG |
| 1 | GMTACGGTCACACTG | chrX:6632959-6634959 | 849 | 863 | + | 3.3571 | 7.12E-05 | 0.795 | GAGGAGGACACACTG |
| 1 | GMTACGGTCACACTG | chr3R:5595340-5597340 | 851 | 865 | + | 3.3571 | 7.12E-05 | 0.795 | GCTCCGGTTATGCTG |
| 1 | GMTACGGTCACACTG | chr3R:23172873-23174873 | 851 | 865 | + | 3.3571 | 7.12E-05 | 0.795 | GCTGTTGTCACTCTG |
| 1 | GMTACGGTCACACTG | chr2R:24858260-24860260 | 852 | 866 | + | 3.3571 | 7.12E-05 | 0.795 | GCTACGCCCACACCA |
| 1 | GMTACGGTCACACTG | chr2R:19668832-19670832 | 857 | 871 | + | 3.3571 | 7.12E-05 | 0.795 | GACGGGGGCACACTG |
| 1 | GMTACGGTCACACTG | chr2R:10457892-10459892 | 858 | 872 | + | 3.3571 | 7.12E-05 | 0.795 | TCTGCGGTAACTCTG |
| 1 | GMTACGGTCACACTG | chr3R:9632076-9634076 | 858 | 872 | + | 3.3571 | 7.12E-05 | 0.795 | CCTGCGGCCACATTG |
| 1 | GMTACGGTCACACTG | chr2L:6715186-6717186 | 860 | 874 | + | 3.3571 | 7.12E-05 | 0.795 | ACACCGTTCACACTG |
| 1 | GMTACGGTCACACTG | chr2L:8593489-8595489 | 869 | 883 | + | 3.3571 | 7.12E-05 | 0.795 | GCCACTGTCCAACTG |
| 1 | GMTACGGTCACACTG | chr2L:249823-251823 | 870 | 884 | + | 3.3571 | 7.12E-05 | 0.795 | GATGCCGGCACCCTG |
| 1 | GMTACGGTCACACTG | chr2L:3913288-3915288 | 871 | 885 | + | 3.3571 | 7.12E-05 | 0.795 | TCAACGGCAACACTG |
| 1 | GMTACGGTCACACTG | chr3R:31394304-31396304 | 872 | 886 | + | 3.3571 | 7.12E-05 | 0.795 | GCAAAGGGAACACTG |
| 1 | GMTACGGTCACACTG | chr3R:25675778-25677778 | 872 | 886 | + | 3.3571 | 7.12E-05 | 0.795 | ACTCCGCTCACTCTG |
| 1 | GMTACGGTCACACTG | chr2L:9425784-9427784 | 874 | 888 | + | 3.3571 | 7.12E-05 | 0.795 | GCTACGGTCTTGTTG |
| 1 | GMTACGGTCACACTG | chr2R:7792380-7794380 | 897 | 911 | + | 3.3571 | 7.12E-05 | 0.795 | GCGTAGGTCACACTT |
| 1 | GMTACGGTCACACTG | chr2R:13294927-13296927 | 898 | 912 | + | 3.3571 | 7.12E-05 | 0.795 | GATTCGCGCACACCG |
| 1 | GMTACGGTCACACTG | chr3R:18282528-18284528 | 898 | 912 | + | 3.3571 | 7.12E-05 | 0.795 | GATGCGGAAGCACTG |
| 1 | GMTACGGTCACACTG | chr3R:17724777-17726777 | 898 | 912 | + | 3.3571 | 7.12E-05 | 0.795 | GCAACGGCAACAATG |
| 1 | GMTACGGTCACACTG | chr2R:6682749-6684749 | 898 | 912 | + | 3.3571 | 7.12E-05 | 0.795 | TCAACGGGCACAATG |
| 1 | GMTACGGTCACACTG | chr3L:9409832-9411832 | 902 | 916 | + | 3.3571 | 7.12E-05 | 0.795 | CAGACGGACACACGG |
| 1 | GMTACGGTCACACTG | chr2L:16981301-16983301 | 902 | 916 | + | 3.3571 | 7.12E-05 | 0.795 | GCAATTGTCCCACTG |
| 1 | GMTACGGTCACACTG | chr2L:12973322-12975322 | 903 | 917 | + | 3.3571 | 7.12E-05 | 0.795 | GCTTCGCTCGCACTC |
| 1 | GMTACGGTCACACTG | chr3R:18359983-18361983 | 905 | 919 | + | 3.3571 | 7.12E-05 | 0.795 | GATTCGGGAGCACTG |
| 1 | GMTACGGTCACACTG | chr3R:22760076-22762076 | 910 | 924 | + | 3.3571 | 7.12E-05 | 0.795 | GCTTCAGTCACATCG |
| 1 | GMTACGGTCACACTG | chr3L:11772878-11774878 | 915 | 929 | + | 3.3571 | 7.12E-05 | 0.795 | TCTCCGGGCAAACTG |
| 1 | GMTACGGTCACACTG | chrX:3967933-3969933 | 916 | 930 | + | 3.3571 | 7.12E-05 | 0.795 | GATGCGGCAACGCTG |
| 1 | GMTACGGTCACACTG | chr2L:18805205-18807205 | 916 | 930 | + | 3.3571 | 7.12E-05 | 0.795 | GCTACCACCACACAG |
| 1 | GMTACGGTCACACTG | chr3L:21522186-21524186 | 919 | 933 | + | 3.3571 | 7.12E-05 | 0.795 | GATACGGCCACTCAA |

| | | | | | | | | | |
|---|---|---|---|---|---|---|---|---|---|
| 1 | GMTACGGTCACACTG | chr2R:9307722-9309722 | 921 | 935 | + | 3.3571 | 7.12E-05 | 0.795 | GCTGCGTTCCCACTT |
| 1 | GMTACGGTCACACTG | chr3R:24056310-24058310 | 922 | 936 | + | 3.3571 | 7.12E-05 | 0.795 | TCAACTGTCACTCTG |
| 1 | GMTACGGTCACACTG | chr2L:11014620-11016620 | 927 | 941 | + | 3.3571 | 7.12E-05 | 0.795 | GCCACGGCCACAGTC |
| 1 | GMTACGGTCACACTG | chr2L:18707688-18709688 | 928 | 942 | + | 3.3571 | 7.12E-05 | 0.795 | GACGCGTTCTCACTG |
| 1 | GMTACGGTCACACTG | chr3R:6683445-6685445 | 931 | 945 | + | 3.3571 | 7.12E-05 | 0.795 | GCCACTGTGGCACTG |
| 1 | GMTACGGTCACACTG | chr3R:24384052-24386052 | 932 | 946 | + | 3.3571 | 7.12E-05 | 0.795 | GCCTCGCTCACATTG |
| 1 | GMTACGGTCACACTG | chr3R:27581241-27583241 | 937 | 951 | + | 3.3571 | 7.12E-05 | 0.795 | GAATCGTTCACACAG |
| 1 | GMTACGGTCACACTG | chr3R:23518445-23520445 | 940 | 954 | + | 3.3571 | 7.12E-05 | 0.795 | GCAACTGTCACAGCG |
| 1 | GMTACGGTCACACTG | chr2R:20976056-20978056 | 940 | 954 | + | 3.3571 | 7.12E-05 | 0.795 | GCAATGGTCACATCG |
| 1 | GMTACGGTCACACTG | chr2L:2251341-2253341 | 946 | 960 | + | 3.3571 | 7.12E-05 | 0.795 | ACCACGGTGACGCTG |
| 1 | GMTACGGTCACACTG | chr3R:20251770-20253770 | 947 | 961 | + | 3.3571 | 7.12E-05 | 0.795 | CCAACGGTCCCACTT |
| 1 | GMTACGGTCACACTG | chr3R:20030823-20032823 | 951 | 965 | + | 3.3571 | 7.12E-05 | 0.795 | ACTGTGGTCACCCTG |
| 1 | GMTACGGTCACACTG | chr2L:10869167-10871167 | 951 | 965 | + | 3.3571 | 7.12E-05 | 0.795 | CCAACGATCACACAG |
| 1 | GMTACGGTCACACTG | chrX:15761956-15763956 | 954 | 968 | + | 3.3571 | 7.12E-05 | 0.795 | ACGTGGGTCACACTG |
| 1 | GMTACGGTCACACTG | chr2L:158032-160032 | 957 | 971 | + | 3.3571 | 7.12E-05 | 0.795 | GCTGCGACCACATTG |
| 1 | GMTACGGTCACACTG | chrX:11301783-11303783 | 957 | 971 | + | 3.3571 | 7.12E-05 | 0.795 | ACTGCGCGCACACTG |
| 1 | GMTACGGTCACACTG | chr2R:20318180-20320180 | 960 | 974 | + | 3.3571 | 7.12E-05 | 0.795 | CCCACGGTCTCACTA |
| 1 | GMTACGGTCACACTG | chr3R:23762857-23764857 | 962 | 976 | + | 3.3571 | 7.12E-05 | 0.795 | GAAACGGGCACATCG |
| 1 | GMTACGGTCACACTG | chr2L:6771755-6773755 | 962 | 976 | + | 3.3571 | 7.12E-05 | 0.795 | GCTACGGTCTCGTTC |
| 1 | GMTACGGTCACACTG | chr2R:15330325-15332325 | 963 | 977 | + | 3.3571 | 7.12E-05 | 0.795 | GCTCCTGTCACGCTT |
| 1 | GMTACGGTCACACTG | chr2L:17584095-17586095 | 963 | 977 | + | 3.3571 | 7.12E-05 | 0.795 | CACACGGCCACCCTG |
| 1 | GMTACGGTCACACTG | chr3R:11304608-11306608 | 964 | 978 | + | 3.3571 | 7.12E-05 | 0.795 | GCTTCGATCACATCG |
| 1 | GMTACGGTCACACTG | chrX:2928796-2930796 | 965 | 979 | + | 3.3571 | 7.12E-05 | 0.795 | GCTCGGGTCGCAATG |
| 1 | GMTACGGTCACACTG | chr2L:10267872-10269872 | 965 | 979 | + | 3.3571 | 7.12E-05 | 0.795 | GCGCCGGTAACACTA |
| 1 | GMTACGGTCACACTG | chr2L:4613300-4615300 | 967 | 981 | + | 3.3571 | 7.12E-05 | 0.795 | GCGACGATCACACCT |
| 1 | GMTACGGTCACACTG | chr2R:5696872-5698872 | 968 | 982 | + | 3.3571 | 7.12E-05 | 0.795 | TCCGCGGTCACACTA |
| 1 | GMTACGGTCACACTG | chr2L:8436405-8438405 | 968 | 982 | + | 3.3571 | 7.12E-05 | 0.795 | GATGCGTGCACGCTG |
| 1 | GMTACGGTCACACTG | chr3L:3164181-3166181 | 968 | 982 | + | 3.3571 | 7.12E-05 | 0.795 | GCTACGGTATTTCTG |
| 1 | GMTACGGTCACACTG | chr2R:7406819-7408819 | 970 | 984 | + | 3.3571 | 7.12E-05 | 0.795 | AATGCGGCCGCACTG |
| 1 | GMTACGGTCACACTG | chr3R:5623474-5625474 | 971 | 985 | + | 3.3571 | 7.12E-05 | 0.795 | GCGATGGTCGAACTG |
| 1 | GMTACGGTCACACTG | chr2L:20065391-20067391 | 972 | 986 | + | 3.3571 | 7.12E-05 | 0.795 | CAAGCGGTCACACAG |
| 1 | GMTACGGTCACACTG | chr2R:11767453-11769453 | 972 | 986 | + | 3.3571 | 7.12E-05 | 0.795 | GAGCTGGTCGCACTG |
| 1 | GMTACGGTCACACTG | chr2R:12984367-12986367 | 972 | 986 | + | 3.3571 | 7.12E-05 | 0.795 | TCTACGCGCACACAG |

| | | | | | | | | | |
|---|---|---|---|---|---|---|---|---|---|
| 1 | GMTACGGTCACACTG | chr3L:24536236-24538236 | 973 | 987 | + | 3.3571 | 7.12E-05 | 0.795 | GAGACGGACAGACGG |
| 1 | GMTACGGTCACACTG | chrX:21357286-21359286 | 973 | 987 | + | 3.3571 | 7.12E-05 | 0.795 | GAACTGGTCCCACTG |
| 1 | GMTACGGTCACACTG | chr2R:16582369-16584369 | 973 | 987 | + | 3.3571 | 7.12E-05 | 0.795 | GATCCGCTCCCGCTG |
| 1 | GMTACGGTCACACTG | chr2R:17245277-17247277 | 976 | 990 | + | 3.3571 | 7.12E-05 | 0.795 | GATGCGGACTCACTC |
| 1 | GMTACGGTCACACTG | chr3R:30057311-30059311 | 976 | 990 | + | 3.3571 | 7.12E-05 | 0.795 | CCTGCTGCCACACTG |
| 1 | GMTACGGTCACACTG | chr3R:14219205-14221205 | 977 | 991 | + | 3.3571 | 7.12E-05 | 0.795 | GATGCGGACTCATTG |
| 1 | GMTACGGTCACACTG | chr3R:16782497-16784497 | 977 | 991 | + | 3.3571 | 7.12E-05 | 0.795 | GCTGCTGTCACCATG |
| 1 | GMTACGGTCACACTG | chr2L:18731524-18733524 | 978 | 992 | + | 3.3571 | 7.12E-05 | 0.795 | GAGACCGTCTCACGG |
| 1 | GMTACGGTCACACTG | chr3R:8216957-8218957 | 979 | 993 | + | 3.3571 | 7.12E-05 | 0.795 | GATACGGGCGAACAG |
| 1 | GMTACGGTCACACTG | chr3L:18063696-18065696 | 980 | 994 | + | 3.3571 | 7.12E-05 | 0.795 | GCGACGGTGACGGTG |
| 1 | GMTACGGTCACACTG | chr3L:5078033-5080033 | 980 | 994 | + | 3.3571 | 7.12E-05 | 0.795 | GCTTAGGACAAACTG |
| 1 | GMTACGGTCACACTG | chr2L:2289738-2291738 | 982 | 996 | + | 3.3571 | 7.12E-05 | 0.795 | GCTCCGGGCATACTT |
| 1 | GMTACGGTCACACTG | chr3L:9546220-9548220 | 983 | 997 | + | 3.3571 | 7.12E-05 | 0.795 | GAAACGGCCGAACTG |
| 1 | GMTACGGTCACACTG | chr3R:7073859-7075859 | 986 | 1000 | + | 3.3571 | 7.12E-05 | 0.795 | GCTCCGGTCACTTTA |
| 1 | GMTACGGTCACACTG | chr3L:5562774-5564774 | 986 | 1000 | + | 3.3571 | 7.12E-05 | 0.795 | GCAATGGTCTTACTG |
| 1 | GMTACGGTCACACTG | chr3L:1333889-1335889 | 987 | 1001 | + | 3.3571 | 7.12E-05 | 0.795 | CCCGCGGTCACACTT |
| 1 | GMTACGGTCACACTG | chr2R:14480187-14482187 | 987 | 1001 | + | 3.3571 | 7.12E-05 | 0.795 | ACCTCGGTCATACTG |
| 1 | GMTACGGTCACACTG | chr2R:23963772-23965772 | 990 | 1004 | + | 3.3571 | 7.12E-05 | 0.795 | GCTCTGGCCACACTC |
| 1 | GMTACGGTCACACTG | chr2R:13505430-13507430 | 991 | 1005 | + | 3.3571 | 7.12E-05 | 0.795 | GACACGGTCTCACCC |
| 1 | GMTACGGTCACACTG | chr2L:9612166-9614166 | 991 | 1005 | + | 3.3571 | 7.12E-05 | 0.795 | GACACAGACACACAG |
| 1 | GMTACGGTCACACTG | chr2R:17468624-17470624 | 991 | 1005 | + | 3.3571 | 7.12E-05 | 0.795 | TCCGCGGTCACACTT |
| 1 | GMTACGGTCACACTG | chr3R:28601147-28603147 | 992 | 1006 | + | 3.3571 | 7.12E-05 | 0.795 | GCTACGATTCCACAG |
| 1 | GMTACGGTCACACTG | chr3L:1330744-1332744 | 992 | 1006 | + | 3.3571 | 7.12E-05 | 0.795 | GCGACGGTCACGTTA |
| 1 | GMTACGGTCACACTG | chr3R:7081182-7083182 | 993 | 1007 | + | 3.3571 | 7.12E-05 | 0.795 | CCCTTGGTCACACTG |
| 1 | GMTACGGTCACACTG | chr3R:7529753-7531753 | 994 | 1008 | + | 3.3571 | 7.12E-05 | 0.795 | CCGGCGGTCACACTT |
| 1 | GMTACGGTCACACTG | chr3R:19835817-19837817 | 995 | 1009 | + | 3.3571 | 7.12E-05 | 0.795 | GCTCCGCTCACTGTG |
| 1 | GMTACGGTCACACTG | chr2L:10031690-10033690 | 995 | 1009 | + | 3.3571 | 7.12E-05 | 0.795 | GATCCGATCGCACAG |
| 1 | GMTACGGTCACACTG | chr3L:13802400-13804400 | 995 | 1009 | + | 3.3571 | 7.12E-05 | 0.795 | GCTACTGCCACACAA |
| 1 | GMTACGGTCACACTG | chr3R:18662738-18664738 | 996 | 1010 | + | 3.3571 | 7.12E-05 | 0.795 | GACCTGGTCACACCG |
| 1 | GMTACGGTCACACTG | chr3R:11649971-11651971 | 996 | 1010 | + | 3.3571 | 7.12E-05 | 0.795 | CCTCTGGTCACACAG |
| 1 | GMTACGGTCACACTG | chr3R:30037728-30039728 | 997 | 1011 | + | 3.3571 | 7.12E-05 | 0.795 | GCTACAGACTAACTG |
| 1 | GMTACGGTCACACTG | chr2L:1183065-1185065 | 1001 | 1015 | + | 3.3571 | 7.12E-05 | 0.795 | GAAGTGGTCACACAG |
| 1 | GMTACGGTCACACTG | chr3L:15098502-15100502 | 1001 | 1015 | + | 3.3571 | 7.12E-05 | 0.795 | GATCCGAACCCACTG |

| | | | | | | | | | |
|---|---|---|---|---|---|---|---|---|---|
| 1 | GMTACGGTCACACTG | chr3R:24348182-24350182 | 1002 | 1016 | + | 3.3571 | 7.12E-05 | 0.795 | ACCACGGCCACACTT |
| 1 | GMTACGGTCACACTG | chr3L:11937872-11939872 | 1002 | 1016 | + | 3.3571 | 7.12E-05 | 0.795 | GCTGCCAACACACTG |
| 1 | GMTACGGTCACACTG | chr3L:15048495-15050495 | 1003 | 1017 | + | 3.3571 | 7.12E-05 | 0.795 | GCTCGGAGCACACTG |
| 1 | GMTACGGTCACACTG | chr2L:6610011-6612011 | 1006 | 1020 | + | 3.3571 | 7.12E-05 | 0.795 | GACCTGGTCACACCG |
| 1 | GMTACGGTCACACTG | chr2L:8436365-8438365 | 1008 | 1022 | + | 3.3571 | 7.12E-05 | 0.795 | GATGCGTGCACGCTG |
| 1 | GMTACGGTCACACTG | chr3L:689517-691517 | 1009 | 1023 | + | 3.3571 | 7.12E-05 | 0.795 | GCTGTTGTCGCACTG |
| 1 | GMTACGGTCACACTG | chr2R:23864414-23866414 | 1010 | 1024 | + | 3.3571 | 7.12E-05 | 0.795 | CCTCCGGTCACTCTA |
| 1 | GMTACGGTCACACTG | chr2L:10969443-10971443 | 1011 | 1025 | + | 3.3571 | 7.12E-05 | 0.795 | ACATTGGTCACACTG |
| 1 | GMTACGGTCACACTG | chr2R:10656545-10658545 | 1016 | 1030 | + | 3.3571 | 7.12E-05 | 0.795 | GCTAAGGATACTCTG |
| 1 | GMTACGGTCACACTG | chr3R:23679039-23681039 | 1018 | 1032 | + | 3.3571 | 7.12E-05 | 0.795 | ACAACGGTCCCACTC |
| 1 | GMTACGGTCACACTG | chr3R:13010310-13012310 | 1018 | 1032 | + | 3.3571 | 7.12E-05 | 0.795 | TCAGCGGTCACACTT |
| 1 | GMTACGGTCACACTG | chr2L:9165514-9167514 | 1019 | 1033 | + | 3.3571 | 7.12E-05 | 0.795 | GCTGCAGTAGCACTG |
| 1 | GMTACGGTCACACTG | chr2R:14854429-14856429 | 1020 | 1034 | + | 3.3571 | 7.12E-05 | 0.795 | GCACCAGTGACACTG |
| 1 | GMTACGGTCACACTG | chr3R:6126152-6128152 | 1020 | 1034 | + | 3.3571 | 7.12E-05 | 0.795 | GATACGTGCACTCAG |
| 1 | GMTACGGTCACACTG | chr2L:13812816-13814816 | 1022 | 1036 | + | 3.3571 | 7.12E-05 | 0.795 | ACTCTGGTCACACCG |
| 1 | GMTACGGTCACACTG | chrX:1879282-1881282 | 1022 | 1036 | + | 3.3571 | 7.12E-05 | 0.795 | GCTTTGGTCTAACTG |
| 1 | GMTACGGTCACACTG | chr2L:13810522-13812522 | 1023 | 1037 | + | 3.3571 | 7.12E-05 | 0.795 | ACTCTGGTCTCACTG |
| 1 | GMTACGGTCACACTG | chr3R:19924371-19926371 | 1023 | 1037 | + | 3.3571 | 7.12E-05 | 0.795 | GCCAATGTCGCACTG |
| 1 | GMTACGGTCACACTG | chr3L:22070848-22072848 | 1023 | 1037 | + | 3.3571 | 7.12E-05 | 0.795 | GCTATGGTCACGCGT |
| 1 | GMTACGGTCACACTG | chr3L:20312632-20314632 | 1024 | 1038 | + | 3.3571 | 7.12E-05 | 0.795 | TCTGCGGTCACACCA |
| 1 | GMTACGGTCACACTG | chr2R:20273597-20275597 | 1024 | 1038 | + | 3.3571 | 7.12E-05 | 0.795 | GCTTTGGTCCCACTA |
| 1 | GMTACGGTCACACTG | chr3L:17464945-17466945 | 1024 | 1038 | + | 3.3571 | 7.12E-05 | 0.795 | GCTACGTAGACACGG |
| 1 | GMTACGGTCACACTG | chr2R:8927169-8929169 | 1025 | 1039 | + | 3.3571 | 7.12E-05 | 0.795 | GCCAGGGTCACTCTA |
| 1 | GMTACGGTCACACTG | chr3R:25334257-25336257 | 1026 | 1040 | + | 3.3571 | 7.12E-05 | 0.795 | ACATCGGTCACACTC |
| 1 | GMTACGGTCACACTG | chr2R:23864398-23866398 | 1026 | 1040 | + | 3.3571 | 7.12E-05 | 0.795 | CCTCCGGTCACTCTA |
| 1 | GMTACGGTCACACTG | chr3R:26716871-26718871 | 1029 | 1043 | + | 3.3571 | 7.12E-05 | 0.795 | ACTCCGGTCACACAA |
| 1 | GMTACGGTCACACTG | chrX:14060817-14062817 | 1031 | 1045 | + | 3.3571 | 7.12E-05 | 0.795 | GATGCGGCCAAGCTG |
| 1 | GMTACGGTCACACTG | chr2R:7925653-7927653 | 1031 | 1045 | + | 3.3571 | 7.12E-05 | 0.795 | ACTTCGGACAGACTG |
| 1 | GMTACGGTCACACTG | chrX:16782933-16784933 | 1033 | 1047 | + | 3.3571 | 7.12E-05 | 0.795 | CCATTGGTCACACTG |
| 1 | GMTACGGTCACACTG | chrX:19306925-19308925 | 1036 | 1050 | + | 3.3571 | 7.12E-05 | 0.795 | ACAACGCACACACTG |
| 1 | GMTACGGTCACACTG | chr3L:11560996-11562996 | 1038 | 1052 | + | 3.3571 | 7.12E-05 | 0.795 | GACACAGACACACGG |
| 1 | GMTACGGTCACACTG | chr2L:18731464-18733464 | 1038 | 1052 | + | 3.3571 | 7.12E-05 | 0.795 | GAGACCGTCTCACGG |
| 1 | GMTACGGTCACACTG | chr3R:25334243-25336243 | 1040 | 1054 | + | 3.3571 | 7.12E-05 | 0.795 | ACATCGGTCACACTC |

| | | | | | | | | | |
|---|---|---|---|---|---|---|---|---|---|
| 1 | GMTACGGTCACACTG | chr3R:14285903-14287903 | 1040 | 1054 | + | 3.3571 | 7.12E-05 | 0.795 | GACCTGGTCACACCG |
| 1 | GMTACGGTCACACTG | chr3L:15534395-15536395 | 1041 | 1055 | + | 3.3571 | 7.12E-05 | 0.795 | GCTGCAGTCGCACTA |
| 1 | GMTACGGTCACACTG | chrX:20392751-20394751 | 1042 | 1056 | + | 3.3571 | 7.12E-05 | 0.795 | GAACTGGTCACCCTG |
| 1 | GMTACGGTCACACTG | chr3L:19585003-19587003 | 1043 | 1057 | + | 3.3571 | 7.12E-05 | 0.795 | TCTTTGGTCACCCTG |
| 1 | GMTACGGTCACACTG | chr2L:1183020-1185020 | 1046 | 1060 | + | 3.3571 | 7.12E-05 | 0.795 | GAAGTGGTCACACAG |
| 1 | GMTACGGTCACACTG | chr3R:22542278-22544278 | 1051 | 1065 | + | 3.3571 | 7.12E-05 | 0.795 | GAAACTGGCACACGG |
| 1 | GMTACGGTCACACTG | chr3L:8575808-8577808 | 1052 | 1066 | + | 3.3571 | 7.12E-05 | 0.795 | GCTTCGGTCTAAATG |
| 1 | GMTACGGTCACACTG | chr2L:22537230-22539230 | 1052 | 1066 | + | 3.3571 | 7.12E-05 | 0.795 | GCCAAGGTCGCACTT |
| 1 | GMTACGGTCACACTG | chrX:5754574-5756574 | 1056 | 1070 | + | 3.3571 | 7.12E-05 | 0.795 | GCTCCGCTCTCACTC |
| 1 | GMTACGGTCACACTG | chrX:3239571-3241571 | 1058 | 1072 | + | 3.3571 | 7.12E-05 | 0.795 | TCTTCGGCCAAACTG |
| 1 | GMTACGGTCACACTG | chr2L:18694664-18696664 | 1059 | 1073 | + | 3.3571 | 7.12E-05 | 0.795 | GAAACGCACCCACTG |
| 1 | GMTACGGTCACACTG | chr3R:28338765-28340765 | 1060 | 1074 | + | 3.3571 | 7.12E-05 | 0.795 | GCTACAGCCACACGC |
| 1 | GMTACGGTCACACTG | chr3R:10142070-10144070 | 1063 | 1077 | + | 3.3571 | 7.12E-05 | 0.795 | CCTACGGACAACCTG |
| 1 | GMTACGGTCACACTG | chr2R:15427023-15429023 | 1063 | 1077 | + | 3.3571 | 7.12E-05 | 0.795 | GATTCGGTCAATCGG |
| 1 | GMTACGGTCACACTG | chr2R:24941413-24943413 | 1063 | 1077 | + | 3.3571 | 7.12E-05 | 0.795 | GCTTCTGGCACAGTG |
| 1 | GMTACGGTCACACTG | chr3R:6404895-6406895 | 1065 | 1079 | + | 3.3571 | 7.12E-05 | 0.795 | CAAACGGTCCCGCTG |
| 1 | GMTACGGTCACACTG | chrX:1709182-1711182 | 1065 | 1079 | + | 3.3571 | 7.12E-05 | 0.795 | GAAATGGACGCACTG |
| 1 | GMTACGGTCACACTG | chr3R:25684051-25686051 | 1068 | 1082 | + | 3.3571 | 7.12E-05 | 0.795 | GAGGCGGTCACTCTA |
| 1 | GMTACGGTCACACTG | chr4:500810-502810 | 1069 | 1083 | + | 3.3571 | 7.12E-05 | 0.795 | GATGCGGTTTCGCTG |
| 1 | GMTACGGTCACACTG | chr3L:5562691-5564691 | 1069 | 1083 | + | 3.3571 | 7.12E-05 | 0.795 | GCAATGGTCTTACTG |
| 1 | GMTACGGTCACACTG | chr2L:13810474-13812474 | 1071 | 1085 | + | 3.3571 | 7.12E-05 | 0.795 | ACTCTGGTCTCACTG |
| 1 | GMTACGGTCACACTG | chr3L:19686353-19688353 | 1071 | 1085 | + | 3.3571 | 7.12E-05 | 0.795 | GACCTGGTCACCCTG |
| 1 | GMTACGGTCACACTG | chr2L:3835840-3837840 | 1073 | 1087 | + | 3.3571 | 7.12E-05 | 0.795 | GCGTCGGTAAAACTG |
| 1 | GMTACGGTCACACTG | chr2R:10546260-10548260 | 1075 | 1089 | + | 3.3571 | 7.12E-05 | 0.795 | GCGAGGGTGGCACTG |
| 1 | GMTACGGTCACACTG | chr3L:9508583-9510583 | 1077 | 1091 | + | 3.3571 | 7.12E-05 | 0.795 | TCTCTGGGCACACTG |
| 1 | GMTACGGTCACACTG | chr3L:1644467-1646467 | 1077 | 1091 | + | 3.3571 | 7.12E-05 | 0.795 | GACACTGGCACACCG |
| 1 | GMTACGGTCACACTG | chr2L:23167825-23169825 | 1078 | 1092 | + | 3.3571 | 7.12E-05 | 0.795 | GCTTCGCTCTCACTC |
| 1 | GMTACGGTCACACTG | chr3R:16222395-16224395 | 1080 | 1094 | + | 3.3571 | 7.12E-05 | 0.795 | TCGACGGTGACGCTG |
| 1 | GMTACGGTCACACTG | chr3L:11133987-11135987 | 1082 | 1096 | + | 3.3571 | 7.12E-05 | 0.795 | GCCACGGTCACTGTT |
| 1 | GMTACGGTCACACTG | chr2R:8927105-8929105 | 1089 | 1103 | + | 3.3571 | 7.12E-05 | 0.795 | GCCAGGGTCACTCTA |
| 1 | GMTACGGTCACACTG | chr3R:12766627-12768627 | 1089 | 1103 | + | 3.3571 | 7.12E-05 | 0.795 | GCCGCCGTCACACTT |
| 1 | GMTACGGTCACACTG | chrX:16215750-16217750 | 1091 | 1105 | + | 3.3571 | 7.12E-05 | 0.795 | GACACCGACACACCG |
| 1 | GMTACGGTCACACTG | chr2L:1496243-1498243 | 1091 | 1105 | + | 3.3571 | 7.12E-05 | 0.795 | GAGACGATCCCACCG |

| | | | | | | | | | |
|---|---|---|---|---|---|---|---|---|---|
| 1 | GMTACGGTCACACTG | chr2L:13608974-13610974 | 1092 | 1106 | + | 3.3571 | 7.12E-05 | 0.795 | GCAAAGGTCCCATTG |
| 1 | GMTACGGTCACACTG | chr3R:23780765-23782765 | 1093 | 1107 | + | 3.3571 | 7.12E-05 | 0.795 | GCAACTGTCACACAC |
| 1 | GMTACGGTCACACTG | chr2R:15330195-15332195 | 1093 | 1107 | + | 3.3571 | 7.12E-05 | 0.795 | GCTCCTGTCACGCTT |
| 1 | GMTACGGTCACACTG | chr2L:2516205-2518205 | 1094 | 1108 | + | 3.3571 | 7.12E-05 | 0.795 | GAGGCGTTCACACAG |
| 1 | GMTACGGTCACACTG | chr3R:17222074-17224074 | 1095 | 1109 | + | 3.3571 | 7.12E-05 | 0.795 | GCCACCGACAAACTG |
| 1 | GMTACGGTCACACTG | chr2R:11767453-11769453 | 1098 | 1112 | + | 3.3571 | 7.12E-05 | 0.795 | GATACGGACTGACAG |
| 1 | GMTACGGTCACACTG | chrX:13781459-13783459 | 1099 | 1113 | + | 3.3571 | 7.12E-05 | 0.795 | GATACAGGCACGCAG |
| 1 | GMTACGGTCACACTG | chr3L:15658707-15660707 | 1099 | 1113 | + | 3.3571 | 7.12E-05 | 0.795 | GCTACTTTCTCACAG |
| 1 | GMTACGGTCACACTG | chr2R:23839064-23841064 | 1100 | 1114 | + | 3.3571 | 7.12E-05 | 0.795 | GCTATGGACGCACTT |
| 1 | GMTACGGTCACACTG | chr2R:15816004-15818004 | 1101 | 1115 | + | 3.3571 | 7.12E-05 | 0.795 | GCTGCGCTCTCACTC |
| 1 | GMTACGGTCACACTG | chr2L:5924961-5926961 | 1103 | 1117 | + | 3.3571 | 7.12E-05 | 0.795 | GCACCGGTTACACTA |
| 1 | GMTACGGTCACACTG | chr3R:27111566-27113566 | 1104 | 1118 | + | 3.3571 | 7.12E-05 | 0.795 | GATGCGGTCACCAAG |
| 1 | GMTACGGTCACACTG | chrX:7410906-7412906 | 1104 | 1118 | + | 3.3571 | 7.12E-05 | 0.795 | GCTCCGTTGACCCTG |
| 1 | GMTACGGTCACACTG | chr2R:19945814-19947814 | 1111 | 1125 | + | 3.3571 | 7.12E-05 | 0.795 | GCCACGCCCACACTT |
| 1 | GMTACGGTCACACTG | chr3R:30305640-30307640 | 1111 | 1125 | + | 3.3571 | 7.12E-05 | 0.795 | GCCATGGTCACTCTA |
| 1 | GMTACGGTCACACTG | chr2R:10198711-10200711 | 1111 | 1125 | + | 3.3571 | 7.12E-05 | 0.795 | GCGACGGTGACGGTG |
| 1 | GMTACGGTCACACTG | chrX:15733270-15735270 | 1117 | 1131 | + | 3.3571 | 7.12E-05 | 0.795 | GCAACGGCGACAGTG |
| 1 | GMTACGGTCACACTG | chr3R:9492677-9494677 | 1117 | 1131 | + | 3.3571 | 7.12E-05 | 0.795 | GATACGCTCTCTCAG |
| 1 | GMTACGGTCACACTG | chr3R:7721417-7723417 | 1118 | 1132 | + | 3.3571 | 7.12E-05 | 0.795 | GAGACGGTCGCCATG |
| 1 | GMTACGGTCACACTG | chr3R:18188072-18190072 | 1119 | 1133 | + | 3.3571 | 7.12E-05 | 0.795 | GCTACGATGGCACCG |
| 1 | GMTACGGTCACACTG | chr2L:18694602-18696602 | 1121 | 1135 | + | 3.3571 | 7.12E-05 | 0.795 | GAAACGCACCCACTG |
| 1 | GMTACGGTCACACTG | chrX:1926683-1928683 | 1121 | 1135 | + | 3.3571 | 7.12E-05 | 0.795 | GCCACGTTCCCAATG |
| 1 | GMTACGGTCACACTG | chr2R:20273499-20275499 | 1122 | 1136 | + | 3.3571 | 7.12E-05 | 0.795 | GCTTTGGTCCCACTA |
| 1 | GMTACGGTCACACTG | chr2L:14557852-14559852 | 1123 | 1137 | + | 3.3571 | 7.12E-05 | 0.795 | GAAACGATCGCTCTG |
| 1 | GMTACGGTCACACTG | chr3L:20459683-20461683 | 1125 | 1139 | + | 3.3571 | 7.12E-05 | 0.795 | GATCCGTTCGCACGG |
| 1 | GMTACGGTCACACTG | chr2L:5924937-5926937 | 1127 | 1141 | + | 3.3571 | 7.12E-05 | 0.795 | GCACCGGTTACACTA |
| 1 | GMTACGGTCACACTG | chr2L:11090444-11092444 | 1129 | 1143 | + | 3.3571 | 7.12E-05 | 0.795 | CCCACGGAGACACTG |
| 1 | GMTACGGTCACACTG | chr3R:30051514-30053514 | 1130 | 1144 | + | 3.3571 | 7.12E-05 | 0.795 | GAGCTGGCCACACTG |
| 1 | GMTACGGTCACACTG | chr3R:25677683-25679683 | 1130 | 1144 | + | 3.3571 | 7.12E-05 | 0.795 | GAACCGGACATACTG |
| 1 | GMTACGGTCACACTG | chr3R:25677682-25679682 | 1131 | 1145 | + | 3.3571 | 7.12E-05 | 0.795 | GAACCGGACATACTG |
| 1 | GMTACGGTCACACTG | chr2L:155029-157029 | 1132 | 1146 | + | 3.3571 | 7.12E-05 | 0.795 | ACTCTGGTCACACAG |
| 1 | GMTACGGTCACACTG | chr3R:29748219-29750219 | 1136 | 1150 | + | 3.3571 | 7.12E-05 | 0.795 | GACACGATCGCCCTG |
| 1 | GMTACGGTCACACTG | chr3L:9796687-9798687 | 1138 | 1152 | + | 3.3571 | 7.12E-05 | 0.795 | GAGGCTGCCACACTG |

| | | | | | | | | | |
|---|---|---|---|---|---|---|---|---|---|
| 1 | GMTACGGTCACACTG | chr3L:4363550-4365550 | 1139 | 1153 | + | 3.3571 | 7.12E-05 | 0.795 | TCACCGGTCACAATG |
| 1 | GMTACGGTCACACTG | chr3R:21014949-21016949 | 1141 | 1155 | + | 3.3571 | 7.12E-05 | 0.795 | GCTACGGCAACTTTG |
| 1 | GMTACGGTCACACTG | chr2L:7826650-7828650 | 1146 | 1160 | + | 3.3571 | 7.12E-05 | 0.795 | CAATCGGTCACACCG |
| 1 | GMTACGGTCACACTG | chr3R:22608485-22610485 | 1153 | 1167 | + | 3.3571 | 7.12E-05 | 0.795 | TCTACGGCCACAACG |
| 1 | GMTACGGTCACACTG | chr3R:6833967-6835967 | 1155 | 1169 | + | 3.3571 | 7.12E-05 | 0.795 | GACCAGGTCCCACTG |
| 1 | GMTACGGTCACACTG | chrX:16275270-16277270 | 1158 | 1172 | + | 3.3571 | 7.12E-05 | 0.795 | GCTACGATCGTGCTG |
| 1 | GMTACGGTCACACTG | chr3R:18359728-18361728 | 1160 | 1174 | + | 3.3571 | 7.12E-05 | 0.795 | GATTCGGGAGCACTG |
| 1 | GMTACGGTCACACTG | chr3R:17539421-17541421 | 1161 | 1175 | + | 3.3571 | 7.12E-05 | 0.795 | GCTACAGTCAGTCAG |
| 1 | GMTACGGTCACACTG | chr3L:3555299-3557299 | 1162 | 1176 | + | 3.3571 | 7.12E-05 | 0.795 | GCTGCTGACACAGTG |
| 1 | GMTACGGTCACACTG | chr3R:14035398-14037398 | 1163 | 1177 | + | 3.3571 | 7.12E-05 | 0.795 | GCTGCGGTCGAAATG |
| 1 | GMTACGGTCACACTG | chr3R:23677895-23679895 | 1169 | 1183 | + | 3.3571 | 7.12E-05 | 0.795 | GCGACGCTCCCATTG |
| 1 | GMTACGGTCACACTG | chr2R:8906518-8908518 | 1172 | 1186 | + | 3.3571 | 7.12E-05 | 0.795 | GAGACGAACACGCTG |
| 1 | GMTACGGTCACACTG | chrY:572088-574088 | 1175 | 1189 | + | 3.3571 | 7.12E-05 | 0.795 | GCGATGTTCTCACTG |
| 1 | GMTACGGTCACACTG | chr2R:13410739-13412739 | 1177 | 1191 | + | 3.3571 | 7.12E-05 | 0.795 | ACGACGGTCGCAATG |
| 1 | GMTACGGTCACACTG | chr2R:13406011-13408011 | 1178 | 1192 | + | 3.3571 | 7.12E-05 | 0.795 | ACGACGGTCGCAATG |
| 1 | GMTACGGTCACACTG | chr2R:11317676-11319676 | 1179 | 1193 | + | 3.3571 | 7.12E-05 | 0.795 | GCCATGGACACAATG |
| 1 | GMTACGGTCACACTG | chr2L:10969275-10971275 | 1179 | 1193 | + | 3.3571 | 7.12E-05 | 0.795 | ACATTGGTCACACTG |
| 1 | GMTACGGTCACACTG | chr3R:4137652-4139652 | 1180 | 1194 | + | 3.3571 | 7.12E-05 | 0.795 | GCCACGCCCACACTT |
| 1 | GMTACGGTCACACTG | chr3R:18662552-18664552 | 1182 | 1196 | + | 3.3571 | 7.12E-05 | 0.795 | GACCTGGTCACACCG |
| 1 | GMTACGGTCACACTG | chrX:1557400-1559400 | 1182 | 1196 | + | 3.3571 | 7.12E-05 | 0.795 | GCTACCGTATCACGG |
| 1 | GMTACGGTCACACTG | chr3L:21756157-21758157 | 1183 | 1197 | + | 3.3571 | 7.12E-05 | 0.795 | GAGACGGAACCACTG |
| 1 | GMTACGGTCACACTG | chr2R:10359213-10361213 | 1184 | 1198 | + | 3.3571 | 7.12E-05 | 0.795 | GCTGCGCTCACTTTG |
| 1 | GMTACGGTCACACTG | chrX:8478319-8480319 | 1188 | 1202 | + | 3.3571 | 7.12E-05 | 0.795 | ACTATGGTCCCGCTG |
| 1 | GMTACGGTCACACTG | chrX:13721246-13723246 | 1189 | 1203 | + | 3.3571 | 7.12E-05 | 0.795 | GAGGCCGACACACTG |
| 1 | GMTACGGTCACACTG | chr2R:20273597-20275597 | 1190 | 1204 | + | 3.3571 | 7.12E-05 | 0.795 | GCTACGTTCGCACGC |
| 1 | GMTACGGTCACACTG | chr3L:4039529-4041529 | 1190 | 1204 | + | 3.3571 | 7.12E-05 | 0.795 | GCTGCGAGCACACTA |
| 1 | GMTACGGTCACACTG | chr3R:29748163-29750163 | 1192 | 1206 | + | 3.3571 | 7.12E-05 | 0.795 | GACACGATCGCCCTG |
| 1 | GMTACGGTCACACTG | chrX:3967655-3969655 | 1194 | 1208 | + | 3.3571 | 7.12E-05 | 0.795 | GATGCGGCAACGCTG |
| 1 | GMTACGGTCACACTG | chr3L:19600633-19602633 | 1197 | 1211 | + | 3.3571 | 7.12E-05 | 0.795 | GAAACGGAGTCACTG |
| 1 | GMTACGGTCACACTG | chr3L:11383258-11385258 | 1206 | 1220 | + | 3.3571 | 7.12E-05 | 0.795 | GCTACAGCCTCAGTG |
| 1 | GMTACGGTCACACTG | chrX:15452471-15454471 | 1206 | 1220 | + | 3.3571 | 7.12E-05 | 0.795 | GCTGCGCTCACGTTG |
| 1 | GMTACGGTCACACTG | chr2L:21928064-21930064 | 1207 | 1221 | + | 3.3571 | 7.12E-05 | 0.795 | GCTTCTTTCTCACTG |
| 1 | GMTACGGTCACACTG | chrX:18100970-18102970 | 1210 | 1224 | + | 3.3571 | 7.12E-05 | 0.795 | GAGACGCTCACCCAG |

| | | | | | | | | | |
|---|---|---|---|---|---|---|---|---|---|
| 1 | GMTACGGTCACACTG | chr3R:22740150-22742150 | 1217 | 1231 | + | 3.3571 | 7.12E-05 | 0.795 | ACCACAGTCACCCTG |
| 1 | GMTACGGTCACACTG | chr2R:11320312-11322312 | 1224 | 1238 | + | 3.3571 | 7.12E-05 | 0.795 | CCTTCGGTCACATCG |
| 1 | GMTACGGTCACACTG | chr3R:9792295-9794295 | 1226 | 1240 | + | 3.3571 | 7.12E-05 | 0.795 | GCGACCGTCAGCCTG |
| 1 | GMTACGGTCACACTG | chrX:19703259-19705259 | 1228 | 1242 | + | 3.3571 | 7.12E-05 | 0.795 | ACTACGGTCACGTAG |
| 1 | GMTACGGTCACACTG | chrX:12407861-12409861 | 1230 | 1244 | + | 3.3571 | 7.12E-05 | 0.795 | GCACCAGTCACAGTG |
| 1 | GMTACGGTCACACTG | chr2L:19735044-19737044 | 1231 | 1245 | + | 3.3571 | 7.12E-05 | 0.795 | GATTCGGACAAACAG |
| 1 | GMTACGGTCACACTG | chr3L:12884039-12886039 | 1235 | 1249 | + | 3.3571 | 7.12E-05 | 0.795 | CCCACGGACACAGTG |
| 1 | GMTACGGTCACACTG | chrX:2294466-2296466 | 1236 | 1250 | + | 3.3571 | 7.12E-05 | 0.795 | GAAACGGACACACAT |
| 1 | GMTACGGTCACACTG | chr3R:23780621-23782621 | 1237 | 1251 | + | 3.3571 | 7.12E-05 | 0.795 | GCAACTGTCACACAC |
| 1 | GMTACGGTCACACTG | chr2R:17747935-17749935 | 1237 | 1251 | + | 3.3571 | 7.12E-05 | 0.795 | TCCTCGGTCACAGTG |
| 1 | GMTACGGTCACACTG | chr3L:4039482-4041482 | 1237 | 1251 | + | 3.3571 | 7.12E-05 | 0.795 | GCTGCGAGCACACTA |
| 1 | GMTACGGTCACACTG | chrX:19242631-19244631 | 1243 | 1257 | + | 3.3571 | 7.12E-05 | 0.795 | CAGTCGGTCACGCTG |
| 1 | GMTACGGTCACACTG | chr3R:16639205-16641205 | 1244 | 1258 | + | 3.3571 | 7.12E-05 | 0.795 | GCGACGGTAATCCTG |
| 1 | GMTACGGTCACACTG | chr3R:6998228-7000228 | 1247 | 1261 | + | 3.3571 | 7.12E-05 | 0.795 | GCTCCGGTCATAAAG |
| 1 | GMTACGGTCACACTG | chrX:16275180-16277180 | 1248 | 1262 | + | 3.3571 | 7.12E-05 | 0.795 | GCTACGATCGTGCTG |
| 1 | GMTACGGTCACACTG | chr3R:12766468-12768468 | 1248 | 1262 | + | 3.3571 | 7.12E-05 | 0.795 | GCCGCCGTCACACTT |
| 1 | GMTACGGTCACACTG | chr2L:9341832-9343832 | 1251 | 1265 | + | 3.3571 | 7.12E-05 | 0.795 | GCCACGGTCTCATTC |
| 1 | GMTACGGTCACACTG | chrX:1879282-1881282 | 1252 | 1266 | + | 3.3571 | 7.12E-05 | 0.795 | CAGTCGGCCACACTG |
| 1 | GMTACGGTCACACTG | chr2R:12377440-12379440 | 1254 | 1268 | + | 3.3571 | 7.12E-05 | 0.795 | GCTACGTGCACATGG |
| 1 | GMTACGGTCACACTG | chr2R:21170001-21172001 | 1256 | 1270 | + | 3.3571 | 7.12E-05 | 0.795 | AAGACGGCCACGCTG |
| 1 | GMTACGGTCACACTG | chr3R:11649706-11651706 | 1261 | 1275 | + | 3.3571 | 7.12E-05 | 0.795 | CCTCTGGTCACACAG |
| 1 | GMTACGGTCACACTG | chr3R:5829062-5831062 | 1261 | 1275 | + | 3.3571 | 7.12E-05 | 0.795 | GCTACGGTCAGGACG |
| 1 | GMTACGGTCACACTG | chr3R:28600877-28602877 | 1262 | 1276 | + | 3.3571 | 7.12E-05 | 0.795 | GCTACGATTCCACAG |
| 1 | GMTACGGTCACACTG | chr3R:5042025-5044025 | 1264 | 1278 | + | 3.3571 | 7.12E-05 | 0.795 | GATACGGAGCCACAG |
| 1 | GMTACGGTCACACTG | chr2L:4957411-4959411 | 1268 | 1282 | + | 3.3571 | 7.12E-05 | 0.795 | GATTCAGACACGCTG |
| 1 | GMTACGGTCACACTG | chr3R:25490460-25492460 | 1274 | 1288 | + | 3.3571 | 7.12E-05 | 0.795 | GATACGGGAACGCAG |
| 1 | GMTACGGTCACACTG | chr3R:7912851-7914851 | 1275 | 1289 | + | 3.3571 | 7.12E-05 | 0.795 | GAGACCGGCACCCTG |
| 1 | GMTACGGTCACACTG | chr3R:22740091-22742091 | 1276 | 1290 | + | 3.3571 | 7.12E-05 | 0.795 | ACCACAGTCACCCTG |
| 1 | GMTACGGTCACACTG | chr2R:11705222-11707222 | 1277 | 1291 | + | 3.3571 | 7.12E-05 | 0.795 | ACTTCGGGCACATTG |
| 1 | GMTACGGTCACACTG | chrX:493614-495614 | 1279 | 1293 | + | 3.3571 | 7.12E-05 | 0.795 | GACTCGCTCACTCTG |
| 1 | GMTACGGTCACACTG | chr2R:12922904-12924904 | 1280 | 1294 | + | 3.3571 | 7.12E-05 | 0.795 | GACACGGGCAGACCG |
| 1 | GMTACGGTCACACTG | chr3L:15519620-15521620 | 1284 | 1298 | + | 3.3571 | 7.12E-05 | 0.795 | ACAACAGGCACACTG |
| 1 | GMTACGGTCACACTG | chrX:6968918-6970918 | 1284 | 1298 | + | 3.3571 | 7.12E-05 | 0.795 | GCTACGAACAGACCG |

| | | | | | | | | | |
|---|---|---|---|---|---|---|---|---|---|
| 1 | GMTACGGTCACACTG | chr2R:9142956-9144956 | 1287 | 1301 | + | 3.3571 | 7.12E-05 | 0.795 | GCTTTGCTCTCACTG |
| 1 | GMTACGGTCACACTG | chr2R:20273499-20275499 | 1288 | 1302 | + | 3.3571 | 7.12E-05 | 0.795 | GCTACGTTCGCACGC |
| 1 | GMTACGGTCACACTG | chr2L:5715701-5717701 | 1289 | 1303 | + | 3.3571 | 7.12E-05 | 0.795 | GCTTCGGTACGACTG |
| 1 | GMTACGGTCACACTG | chr3L:19096999-19098999 | 1290 | 1304 | + | 3.3571 | 7.12E-05 | 0.795 | GCTTTGTTCACACGG |
| 1 | GMTACGGTCACACTG | chr2L:6053302-6055302 | 1294 | 1308 | + | 3.3571 | 7.12E-05 | 0.795 | GCAACGATCTAACTG |
| 1 | GMTACGGTCACACTG | chr2L:19734980-19736980 | 1295 | 1309 | + | 3.3571 | 7.12E-05 | 0.795 | GATTCGGACAAACAG |
| 1 | GMTACGGTCACACTG | chr2L:19537425-19539425 | 1299 | 1313 | + | 3.3571 | 7.12E-05 | 0.795 | GATGAGGACGCACTG |
| 1 | GMTACGGTCACACTG | chrX:16440433-16442433 | 1299 | 1313 | + | 3.3571 | 7.12E-05 | 0.795 | ACCACGGTAACACAG |
| 1 | GMTACGGTCACACTG | chr2R:20273597-20275597 | 1300 | 1314 | + | 3.3571 | 7.12E-05 | 0.795 | GATTCGGACAATCTG |
| 1 | GMTACGGTCACACTG | chr3R:29798658-29800658 | 1307 | 1321 | + | 3.3571 | 7.12E-05 | 0.795 | CCAACGATCACACAG |
| 1 | GMTACGGTCACACTG | chr2L:5554072-5556072 | 1308 | 1322 | + | 3.3571 | 7.12E-05 | 0.795 | GCGACGATCGCAATG |
| 1 | GMTACGGTCACACTG | chrX:11790777-11792777 | 1309 | 1323 | + | 3.3571 | 7.12E-05 | 0.795 | GCTTCGCTCTCACTC |
| 1 | GMTACGGTCACACTG | chr2L:5715680-5717680 | 1310 | 1324 | + | 3.3571 | 7.12E-05 | 0.795 | GCTTCGGTACGACTG |
| 1 | GMTACGGTCACACTG | chrX:19703176-19705176 | 1311 | 1325 | + | 3.3571 | 7.12E-05 | 0.795 | ACTACGGTCACGTAG |
| 1 | GMTACGGTCACACTG | chr2R:16765000-16767000 | 1314 | 1328 | + | 3.3571 | 7.12E-05 | 0.795 | GCTGCCGGCACAGTG |
| 1 | GMTACGGTCACACTG | chr2R:23750915-23752915 | 1316 | 1330 | + | 3.3571 | 7.12E-05 | 0.795 | GCTACGCCCACATCG |
| 1 | GMTACGGTCACACTG | chr2L:19534448-19536448 | 1316 | 1330 | + | 3.3571 | 7.12E-05 | 0.795 | GACCCGGTGTCACTG |
| 1 | GMTACGGTCACACTG | chrX:4927133-4929133 | 1317 | 1331 | + | 3.3571 | 7.12E-05 | 0.795 | GCTTCGGCGACAGTG |
| 1 | GMTACGGTCACACTG | chr3R:5623125-5625125 | 1320 | 1334 | + | 3.3571 | 7.12E-05 | 0.795 | GCGATGGTCGAACTG |
| 1 | GMTACGGTCACACTG | chr3R:5623125-5625125 | 1320 | 1334 | + | 3.3571 | 7.12E-05 | 0.795 | GCGATGGTCGAACTG |
| 1 | GMTACGGTCACACTG | chr3L:5493740-5495740 | 1322 | 1336 | + | 3.3571 | 7.12E-05 | 0.795 | GCAACGGACACATTT |
| 1 | GMTACGGTCACACTG | chr2L:157664-159664 | 1325 | 1339 | + | 3.3571 | 7.12E-05 | 0.795 | GCTGCGACCACATTG |
| 1 | GMTACGGTCACACTG | chr3R:12379008-12381008 | 1325 | 1339 | + | 3.3571 | 7.12E-05 | 0.795 | GACATGGGCACCCTG |
| 1 | GMTACGGTCACACTG | chr2R:9142914-9144914 | 1329 | 1343 | + | 3.3571 | 7.12E-05 | 0.795 | GCTTTGCTCTCACTG |
| 1 | GMTACGGTCACACTG | chr2L:558612-560612 | 1330 | 1344 | + | 3.3571 | 7.12E-05 | 0.795 | GAGGCGGCCAAACTG |
| 1 | GMTACGGTCACACTG | chr2R:22329200-22331200 | 1331 | 1345 | + | 3.3571 | 7.12E-05 | 0.795 | GATCCGGACAAGCTG |
| 1 | GMTACGGTCACACTG | chr3R:28750661-28752661 | 1331 | 1345 | + | 3.3571 | 7.12E-05 | 0.795 | GCTACGGGGGCAATG |
| 1 | GMTACGGTCACACTG | chr3R:18365638-18367638 | 1331 | 1345 | + | 3.3571 | 7.12E-05 | 0.795 | GACAAGGACACGCTG |
| 1 | GMTACGGTCACACTG | chr2L:16909418-16911418 | 1333 | 1347 | + | 3.3571 | 7.12E-05 | 0.795 | GCCAGGATCACACGG |
| 1 | GMTACGGTCACACTG | chr2R:22211835-22213835 | 1337 | 1351 | + | 3.3571 | 7.12E-05 | 0.795 | GAATCGGTGGCACTG |
| 1 | GMTACGGTCACACTG | chr3L:894951-896951 | 1343 | 1357 | + | 3.3571 | 7.12E-05 | 0.795 | GACACGGGTACTCTG |
| 1 | GMTACGGTCACACTG | chr3R:27735538-27737538 | 1344 | 1358 | + | 3.3571 | 7.12E-05 | 0.795 | GAGACGGCCACAGCG |
| 1 | GMTACGGTCACACTG | chr2R:24196824-24198824 | 1344 | 1358 | + | 3.3571 | 7.12E-05 | 0.795 | GCGACCGGGACACTG |

| | | | | | | | | | |
|---|---|---|---|---|---|---|---|---|---|
| 1 | GMTACGGTCACACTG | chr2R:22121562-22123562 | 1345 | 1359 | + | 3.3571 | 7.12E-05 | 0.795 | ACAACGGTCACTCTA |
| 1 | GMTACGGTCACACTG | chr3L:1644199-1646199 | 1345 | 1359 | + | 3.3571 | 7.12E-05 | 0.795 | GACACTGGCACACCG |
| 1 | GMTACGGTCACACTG | chr2L:12026438-12028438 | 1349 | 1363 | + | 3.3571 | 7.12E-05 | 0.795 | GCAACAGCCACAATG |
| 1 | GMTACGGTCACACTG | chr2R:14513030-14515030 | 1349 | 1363 | + | 3.3571 | 7.12E-05 | 0.795 | GCAACGGTCGCAGTT |
| 1 | GMTACGGTCACACTG | chr3R:24917106-24919106 | 1350 | 1364 | + | 3.3571 | 7.12E-05 | 0.795 | GCTAGGGCCAAACAG |
| 1 | GMTACGGTCACACTG | chr3R:17027693-17029693 | 1352 | 1366 | + | 3.3571 | 7.12E-05 | 0.795 | CCAAGGGACACACTG |
| 1 | GMTACGGTCACACTG | chr3R:31231336-31233336 | 1353 | 1367 | + | 3.3571 | 7.12E-05 | 0.795 | GCAACTGCAACACTG |
| 1 | GMTACGGTCACACTG | chr2R:12325360-12327360 | 1353 | 1367 | + | 3.3571 | 7.12E-05 | 0.795 | GAAATGGCCACCCTG |
| 1 | GMTACGGTCACACTG | chr2R:12162675-12164675 | 1358 | 1372 | + | 3.3571 | 7.12E-05 | 0.795 | GCTACGGCTACTATG |
| 1 | GMTACGGTCACACTG | chr3L:20308654-20310654 | 1358 | 1372 | + | 3.3571 | 7.12E-05 | 0.795 | GAAACGGGCACACCA |
| 1 | GMTACGGTCACACTG | chr2R:7670103-7672103 | 1362 | 1376 | + | 3.3571 | 7.12E-05 | 0.795 | GATACGCTCCCCCGG |
| 1 | GMTACGGTCACACTG | chrX:783746-785746 | 1363 | 1377 | + | 3.3571 | 7.12E-05 | 0.795 | GCTACCGTCTTCCTG |
| 1 | GMTACGGTCACACTG | chr3R:6643434-6645434 | 1365 | 1379 | + | 3.3571 | 7.12E-05 | 0.795 | GCCACGACCACACTT |
| 1 | GMTACGGTCACACTG | chr3L:15658441-15660441 | 1365 | 1379 | + | 3.3571 | 7.12E-05 | 0.795 | GCTACTTTCTCACAG |
| 1 | GMTACGGTCACACTG | chr3L:11247129-11249129 | 1371 | 1385 | + | 3.3571 | 7.12E-05 | 0.795 | GATTCGGTCACGTCG |
| 1 | GMTACGGTCACACTG | chr3R:5218657-5220657 | 1380 | 1394 | + | 3.3571 | 7.12E-05 | 0.795 | GATGCGGTGACGCAG |
| 1 | GMTACGGTCACACTG | chrX:1456054-1458054 | 1381 | 1395 | + | 3.3571 | 7.12E-05 | 0.795 | GCTACGGCAGAACTG |
| 1 | GMTACGGTCACACTG | chr2R:21294652-21296652 | 1383 | 1397 | + | 3.3571 | 7.12E-05 | 0.795 | GATACGGCTACGCGG |
| 1 | GMTACGGTCACACTG | chrX:15205636-15207636 | 1383 | 1397 | + | 3.3571 | 7.12E-05 | 0.795 | GACACGGAGACACAG |
| 1 | GMTACGGTCACACTG | chr3L:4241734-4243734 | 1383 | 1397 | + | 3.3571 | 7.12E-05 | 0.795 | ACGACGATCGCACTG |
| 1 | GMTACGGTCACACTG | chr2L:20084070-20086070 | 1383 | 1397 | + | 3.3571 | 7.12E-05 | 0.795 | GCTTGGGGCACATTG |
| 1 | GMTACGGTCACACTG | chr3R:7240760-7242760 | 1385 | 1399 | + | 3.3571 | 7.12E-05 | 0.795 | GCAACTGGCACAGTG |
| 1 | GMTACGGTCACACTG | chr3L:9363350-9365350 | 1385 | 1399 | + | 3.3571 | 7.12E-05 | 0.795 | GCCATCGACACACTG |
| 1 | GMTACGGTCACACTG | chrX:13781170-13783170 | 1388 | 1402 | + | 3.3571 | 7.12E-05 | 0.795 | GATACAGGCACGCAG |
| 1 | GMTACGGTCACACTG | chr3R:20324529-20326529 | 1389 | 1403 | + | 3.3571 | 7.12E-05 | 0.795 | ACTACGGTCATCCGG |
| 1 | GMTACGGTCACACTG | chrX:15732996-15734996 | 1391 | 1405 | + | 3.3571 | 7.12E-05 | 0.795 | GCAACGGCGACAGTG |
| 1 | GMTACGGTCACACTG | chr3R:7073452-7075452 | 1393 | 1407 | + | 3.3571 | 7.12E-05 | 0.795 | GCTCCGGTCACTTTA |
| 1 | GMTACGGTCACACTG | chr3R:24347791-24349791 | 1393 | 1407 | + | 3.3571 | 7.12E-05 | 0.795 | ACCACGGCCACACTT |
| 1 | GMTACGGTCACACTG | chr2L:1107540-1109540 | 1395 | 1409 | + | 3.3571 | 7.12E-05 | 0.795 | CCCACGTTCACGCTG |
| 1 | GMTACGGTCACACTG | chr2R:20273499-20275499 | 1398 | 1412 | + | 3.3571 | 7.12E-05 | 0.795 | GATTCGGACAATCTG |
| 1 | GMTACGGTCACACTG | chr2R:17430032-17432032 | 1399 | 1413 | + | 3.3571 | 7.12E-05 | 0.795 | GCTACGAGCACGTTG |
| 1 | GMTACGGTCACACTG | chr3R:27111270-27113270 | 1400 | 1414 | + | 3.3571 | 7.12E-05 | 0.795 | GATGCGGTCACCAAG |
| 1 | GMTACGGTCACACTG | chr2L:14742193-14744193 | 1400 | 1414 | + | 3.3571 | 7.12E-05 | 0.795 | GATTCGGACACGGTG |

| | | | | | | | | | |
|---|---|---|---|---|---|---|---|---|---|
| 1 | GMTACGGTCACACTG | chr2R:14854048-14856048 | 1401 | 1415 | + | 3.3571 | 7.12E-05 | 0.795 | GCACCAGTGACACTG |
| 1 | GMTACGGTCACACTG | chr3L:11247097-11249097 | 1403 | 1417 | + | 3.3571 | 7.12E-05 | 0.795 | GATTCGGTCACGTCG |
| 1 | GMTACGGTCACACTG | chr2R:5879395-5881395 | 1404 | 1418 | + | 3.3571 | 7.12E-05 | 0.795 | GCTTCAGTCGCAGTG |
| 1 | GMTACGGTCACACTG | chr3R:24944237-24946237 | 1405 | 1419 | + | 3.3571 | 7.12E-05 | 0.795 | GCCATGGTCAAGCTG |
| 1 | GMTACGGTCACACTG | chr2R:15540435-15542435 | 1405 | 1419 | + | 3.3571 | 7.12E-05 | 0.795 | GCTGGTGTCACTCTG |
| 1 | GMTACGGTCACACTG | chr3R:24858801-24860801 | 1406 | 1420 | + | 3.3571 | 7.12E-05 | 0.795 | GCGACCGTCACGCTT |
| 1 | GMTACGGTCACACTG | chr2R:5797512-5799512 | 1414 | 1428 | + | 3.3571 | 7.12E-05 | 0.795 | CCTGGGGCCACACTG |
| 1 | GMTACGGTCACACTG | chr2L:1987290-1989290 | 1423 | 1437 | + | 3.3571 | 7.12E-05 | 0.795 | TACACGGCCACCCTG |
| 1 | GMTACGGTCACACTG | chr2R:23086177-23088177 | 1424 | 1438 | + | 3.3571 | 7.12E-05 | 0.795 | GCTAAAGGCGCACTG |
| 1 | GMTACGGTCACACTG | chr3L:7935373-7937373 | 1426 | 1440 | + | 3.3571 | 7.12E-05 | 0.795 | GCCACGGCGACACTT |
| 1 | GMTACGGTCACACTG | chr2L:9575769-9577769 | 1426 | 1440 | + | 3.3571 | 7.12E-05 | 0.795 | GAATCTGTCACGCTG |
| 1 | GMTACGGTCACACTG | chrX:19610526-19612526 | 1434 | 1448 | + | 3.3571 | 7.12E-05 | 0.795 | GATTTGGTCACTCCG |
| 1 | GMTACGGTCACACTG | chrX:19610526-19612526 | 1434 | 1448 | + | 3.3571 | 7.12E-05 | 0.795 | GATTTGGTCACTCCG |
| 1 | GMTACGGTCACACTG | chrX:16440296-16442296 | 1436 | 1450 | + | 3.3571 | 7.12E-05 | 0.795 | ACCACGGTAACACAG |
| 1 | GMTACGGTCACACTG | chr2R:17468177-17470177 | 1438 | 1452 | + | 3.3571 | 7.12E-05 | 0.795 | TCCGCGGTCACACTT |
| 1 | GMTACGGTCACACTG | chr3R:13051245-13053245 | 1440 | 1454 | + | 3.3571 | 7.12E-05 | 0.795 | GATCTGGTCACGCCG |
| 1 | GMTACGGTCACACTG | chr2L:2389689-2391689 | 1440 | 1454 | + | 3.3571 | 7.12E-05 | 0.795 | GCGACGGCGACACTC |
| 1 | GMTACGGTCACACTG | chr3R:28600698-28602698 | 1441 | 1455 | + | 3.3571 | 7.12E-05 | 0.795 | GCTACGATTCCACAG |
| 1 | GMTACGGTCACACTG | chr2L:15763220-15765220 | 1441 | 1455 | + | 3.3571 | 7.12E-05 | 0.795 | GCCAAGGTCATCCTG |
| 1 | GMTACGGTCACACTG | chr3L:8727481-8729481 | 1442 | 1456 | + | 3.3571 | 7.12E-05 | 0.795 | GCTACGGCCAACGTG |
| 1 | GMTACGGTCACACTG | chr3L:689081-691081 | 1445 | 1459 | + | 3.3571 | 7.12E-05 | 0.795 | GCTGTTGTCGCACTG |
| 1 | GMTACGGTCACACTG | chr2R:21294588-21296588 | 1447 | 1461 | + | 3.3571 | 7.12E-05 | 0.795 | GATACGGCTACGCGG |
| 1 | GMTACGGTCACACTG | chrX:10009481-10011481 | 1447 | 1461 | + | 3.3571 | 7.12E-05 | 0.795 | GACACTGTCTCCCTG |
| 1 | GMTACGGTCACACTG | chr3R:20359308-20361308 | 1448 | 1462 | + | 3.3571 | 7.12E-05 | 0.795 | GCCAGGGTCGCATTG |
| 1 | GMTACGGTCACACTG | chr2L:12110110-12112110 | 1449 | 1463 | + | 3.3571 | 7.12E-05 | 0.795 | GCTCCGGTGAAGCTG |
| 1 | GMTACGGTCACACTG | chr2R:12933045-12935045 | 1452 | 1466 | + | 3.3571 | 7.12E-05 | 0.795 | GCTCCGGCCAGAATG |
| 1 | GMTACGGTCACACTG | chr2R:20253670-20255670 | 1452 | 1466 | + | 3.3571 | 7.12E-05 | 0.795 | GAGACGGTCCCACGC |
| 1 | GMTACGGTCACACTG | chr2R:12325258-12327258 | 1455 | 1469 | + | 3.3571 | 7.12E-05 | 0.795 | GAAATGGCCACCCTG |
| 1 | GMTACGGTCACACTG | chrX:13720977-13722977 | 1458 | 1472 | + | 3.3571 | 7.12E-05 | 0.795 | GAGGCCGACACACTG |
| 1 | GMTACGGTCACACTG | chr2R:16343377-16345377 | 1459 | 1473 | + | 3.3571 | 7.12E-05 | 0.795 | GCTCCCCTCACGCTG |
| 1 | GMTACGGTCACACTG | chr3R:10212007-10214007 | 1460 | 1474 | + | 3.3571 | 7.12E-05 | 0.795 | GCTCCGGTCATGCTT |
| 1 | GMTACGGTCACACTG | chr2R:6682186-6684186 | 1461 | 1475 | + | 3.3571 | 7.12E-05 | 0.795 | TCAACGGGCACAATG |
| 1 | GMTACGGTCACACTG | chr3R:10760797-10762797 | 1461 | 1475 | + | 3.3571 | 7.12E-05 | 0.795 | GCTACTGCGCCACTG |

| | | | | | | | | | |
|---|---|---|---|---|---|---|---|---|---|
| 1 | GMTACGGTCACACTG | chr3R:20030309-20032309 | 1465 | 1479 | + | 3.3571 | 7.12E-05 | 0.795 | ACTGTGGTCACCCTG |
| 1 | GMTACGGTCACACTG | chrX:6680767-6682767 | 1467 | 1481 | + | 3.3571 | 7.12E-05 | 0.795 | GCCATGGGCATACTG |
| 1 | GMTACGGTCACACTG | chr2R:7177390-7179390 | 1468 | 1482 | + | 3.3571 | 7.12E-05 | 0.795 | GAAACGGGCAAGCTG |
| 1 | GMTACGGTCACACTG | chr2R:17429963-17431963 | 1468 | 1482 | + | 3.3571 | 7.12E-05 | 0.795 | GCTACGAGCACGTTG |
| 1 | GMTACGGTCACACTG | chrX:11579772-11581772 | 1469 | 1483 | + | 3.3571 | 7.12E-05 | 0.795 | CCTACGGCCAGACGG |
| 1 | GMTACGGTCACACTG | chr3L:12266528-12268528 | 1471 | 1485 | + | 3.3571 | 7.12E-05 | 0.795 | GCTACGGACACTTTC |
| 1 | GMTACGGTCACACTG | chrX:18499127-18501127 | 1472 | 1486 | + | 3.3571 | 7.12E-05 | 0.795 | GCCATGGACACACTC |
| 1 | GMTACGGTCACACTG | chr3R:6348243-6350243 | 1476 | 1490 | + | 3.3571 | 7.12E-05 | 0.795 | GCGAGCGTCCCACTG |
| 1 | GMTACGGTCACACTG | chrX:2185855-2187855 | 1477 | 1491 | + | 3.3571 | 7.12E-05 | 0.795 | GACACGGTCACTACG |
| 1 | GMTACGGTCACACTG | chr2R:15540363-15542363 | 1477 | 1491 | + | 3.3571 | 7.12E-05 | 0.795 | GCTGGTGTCACTCTG |
| 1 | GMTACGGTCACACTG | chr3R:16429681-16431681 | 1481 | 1495 | + | 3.3571 | 7.12E-05 | 0.795 | GAAACTGTCTCCCTG |
| 1 | GMTACGGTCACACTG | chr2L:7826315-7828315 | 1481 | 1495 | + | 3.3571 | 7.12E-05 | 0.795 | CAATCGGTCACACCG |
| 1 | GMTACGGTCACACTG | chr2R:7386187-7388187 | 1482 | 1496 | + | 3.3571 | 7.12E-05 | 0.795 | GCTGCCGGCACAGTG |
| 1 | GMTACGGTCACACTG | chr3R:18187707-18189707 | 1484 | 1498 | + | 3.3571 | 7.12E-05 | 0.795 | GCTACGATGGCACCG |
| 1 | GMTACGGTCACACTG | chr2R:23803811-23805811 | 1485 | 1499 | + | 3.3571 | 7.12E-05 | 0.795 | GCTTCGATGACACAG |
| 1 | GMTACGGTCACACTG | chr2L:478688-480688 | 1486 | 1500 | + | 3.3571 | 7.12E-05 | 0.795 | GACACAGACACACAG |
| 1 | GMTACGGTCACACTG | chr3L:12398322-12400322 | 1486 | 1500 | + | 3.3571 | 7.12E-05 | 0.795 | GCTACGCTCTCCTTG |
| 1 | GMTACGGTCACACTG | chr3R:12985041-12987041 | 1487 | 1501 | + | 3.3571 | 7.12E-05 | 0.795 | GATACGGAAGCTCTG |
| 1 | GMTACGGTCACACTG | chr3R:21785208-21787208 | 1488 | 1502 | + | 3.3571 | 7.12E-05 | 0.795 | GATACGGGCACCCAC |
| 1 | GMTACGGTCACACTG | chr3R:22725967-22727967 | 1493 | 1507 | + | 3.3571 | 7.12E-05 | 0.795 | GAATCGGTAACCCTG |
| 1 | GMTACGGTCACACTG | chrX:10767537-10769537 | 1493 | 1507 | + | 3.3571 | 7.12E-05 | 0.795 | TACTCGGGCACACTG |
| 1 | GMTACGGTCACACTG | chr2R:8932423-8934423 | 1495 | 1509 | + | 3.3571 | 7.12E-05 | 0.795 | GAAGTGGTCGCACTG |
| 1 | GMTACGGTCACACTG | chr3R:10055083-10057083 | 1499 | 1513 | + | 3.3571 | 7.12E-05 | 0.795 | GATCCGGGCACAAAG |
| 1 | GMTACGGTCACACTG | chr3R:7529241-7531241 | 1506 | 1520 | + | 3.3571 | 7.12E-05 | 0.795 | CCGGCGGTCACACTT |
| 1 | GMTACGGTCACACTG | chr3L:16170380-16172380 | 1507 | 1521 | + | 3.3571 | 7.12E-05 | 0.795 | TCAACGGGCACAATG |
| 1 | GMTACGGTCACACTG | chr3R:5218528-5220528 | 1509 | 1523 | + | 3.3571 | 7.12E-05 | 0.795 | GATGCGGTGACGCAG |
| 1 | GMTACGGTCACACTG | chr3L:19095217-19097217 | 1510 | 1524 | + | 3.3571 | 7.12E-05 | 0.795 | GCTGCGGCCAAACTT |
| 1 | GMTACGGTCACACTG | chr3R:26716389-26718389 | 1511 | 1525 | + | 3.3571 | 7.12E-05 | 0.795 | ACTCCGGTCACACAA |
| 1 | GMTACGGTCACACTG | chrX:1369580-1371580 | 1518 | 1532 | + | 3.3571 | 7.12E-05 | 0.795 | GCTGCGCTCAATCTG |
| 1 | GMTACGGTCACACTG | chr2R:16097948-16099948 | 1519 | 1533 | + | 3.3571 | 7.12E-05 | 0.795 | ACGAAGGGCACACTG |
| 1 | GMTACGGTCACACTG | chr3R:18182479-18184479 | 1521 | 1535 | + | 3.3571 | 7.12E-05 | 0.795 | GCGAAGATCACACAG |
| 1 | GMTACGGTCACACTG | chr3L:1033875-1035875 | 1529 | 1543 | + | 3.3571 | 7.12E-05 | 0.795 | GAATCGGCCAAACTG |
| 1 | GMTACGGTCACACTG | chrX:1878774-1880774 | 1530 | 1544 | + | 3.3571 | 7.12E-05 | 0.795 | GCTTTGGTCTAACTG |

| | | | | | | | | | |
|---|---|---|---|---|---|---|---|---|---|
| 1 | GMTACGGTCACACTG | chr3R:30051112-30053112 | 1532 | 1546 | + | 3.3571 | 7.12E-05 | 0.795 | GAGCTGGCCACACTG |
| 1 | GMTACGGTCACACTG | chrX:20821146-20823146 | 1532 | 1546 | + | 3.3571 | 7.12E-05 | 0.795 | GACACCGACACACGG |
| 1 | GMTACGGTCACACTG | chrX:16782434-16784434 | 1532 | 1546 | + | 3.3571 | 7.12E-05 | 0.795 | CCATTGGTCACACTG |
| 1 | GMTACGGTCACACTG | chrX:21604264-21606264 | 1533 | 1547 | + | 3.3571 | 7.12E-05 | 0.795 | GCGGCGGTCACATTA |
| 1 | GMTACGGTCACACTG | chrX:22472698-22474698 | 1536 | 1550 | + | 3.3571 | 7.12E-05 | 0.795 | GCAACGGTCACATCC |
| 1 | GMTACGGTCACACTG | chr3L:10558082-10560082 | 1536 | 1550 | + | 3.3571 | 7.12E-05 | 0.795 | GCCACGATGGCACTG |
| 1 | GMTACGGTCACACTG | chrX:5868095-5870095 | 1537 | 1551 | + | 3.3571 | 7.12E-05 | 0.795 | GAGACGGATGCACTG |
| 1 | GMTACGGTCACACTG | chrX:19851214-19853214 | 1540 | 1554 | + | 3.3571 | 7.12E-05 | 0.795 | GAGGCGGTCATTCTG |
| 1 | GMTACGGTCACACTG | chr3R:18398169-18400169 | 1542 | 1556 | + | 3.3571 | 7.12E-05 | 0.795 | ACCACGGCCACACTC |
| 1 | GMTACGGTCACACTG | chr3R:6267757-6269757 | 1546 | 1560 | + | 3.3571 | 7.12E-05 | 0.795 | GCAACCGTCAGGCTG |
| 1 | GMTACGGTCACACTG | chr3R:31219702-31221702 | 1548 | 1562 | + | 3.3571 | 7.12E-05 | 0.795 | GACCTGGCCACACTG |
| 1 | GMTACGGTCACACTG | chr2R:7386115-7388115 | 1554 | 1568 | + | 3.3571 | 7.12E-05 | 0.795 | GCTGCCGGCACAGTG |
| 1 | GMTACGGTCACACTG | chr3R:23762263-23764263 | 1556 | 1570 | + | 3.3571 | 7.12E-05 | 0.795 | GAAACGGGCACATCG |
| 1 | GMTACGGTCACACTG | chr3R:15791688-15793688 | 1556 | 1570 | + | 3.3571 | 7.12E-05 | 0.795 | GCTAAGGTCTCAAAG |
| 1 | GMTACGGTCACACTG | chr2L:18672285-18674285 | 1557 | 1571 | + | 3.3571 | 7.12E-05 | 0.795 | GCTCCTGCCACACTC |
| 1 | GMTACGGTCACACTG | chr3L:13456238-13458238 | 1558 | 1572 | + | 3.3571 | 7.12E-05 | 0.795 | GCTACAGCTTCACTG |
| 1 | GMTACGGTCACACTG | chr3R:8320747-8322747 | 1561 | 1575 | + | 3.3571 | 7.12E-05 | 0.795 | GAATCGGTCACCATG |
| 1 | GMTACGGTCACACTG | chr3L:23337100-23339100 | 1561 | 1575 | + | 3.3571 | 7.12E-05 | 0.795 | GCCACGCCCACACTT |
| 1 | GMTACGGTCACACTG | chr2L:1107373-1109373 | 1562 | 1576 | + | 3.3571 | 7.12E-05 | 0.795 | CCCACGTTCACGCTG |
| 1 | GMTACGGTCACACTG | chr3R:11998278-12000278 | 1563 | 1577 | + | 3.3571 | 7.12E-05 | 0.795 | GATACGGAATCACAG |
| 1 | GMTACGGTCACACTG | chr2R:12588239-12590239 | 1564 | 1578 | + | 3.3571 | 7.12E-05 | 0.795 | GCCACGGCTACACTT |
| 1 | GMTACGGTCACACTG | chr3R:23398778-23400778 | 1565 | 1579 | + | 3.3571 | 7.12E-05 | 0.795 | GCTGCGGAAAAACTG |
| 1 | GMTACGGTCACACTG | chr3R:24924344-24926344 | 1566 | 1580 | + | 3.3571 | 7.12E-05 | 0.795 | GATCCGGACACTGTG |
| 1 | GMTACGGTCACACTG | chr3R:17626127-17628127 | 1568 | 1582 | + | 3.3571 | 7.12E-05 | 0.795 | GCTACGGCCACCGTT |
| 1 | GMTACGGTCACACTG | chr3L:223106-225106 | 1572 | 1586 | + | 3.3571 | 7.12E-05 | 0.795 | GCAACAGTTACACGG |
| 1 | GMTACGGTCACACTG | chr3R:27111566-27113566 | 1572 | 1586 | + | 3.3571 | 7.12E-05 | 0.795 | GCTACGTGTCCACTG |
| 1 | GMTACGGTCACACTG | chr3L:19096709-19098709 | 1580 | 1594 | + | 3.3571 | 7.12E-05 | 0.795 | GCTTTGTTCACACGG |
| 1 | GMTACGGTCACACTG | chr2L:13812256-13814256 | 1582 | 1596 | + | 3.3571 | 7.12E-05 | 0.795 | ACTCTGGTCACACCG |
| 1 | GMTACGGTCACACTG | chr2R:21169672-21171672 | 1585 | 1599 | + | 3.3571 | 7.12E-05 | 0.795 | AAGACGGCCACGCTG |
| 1 | GMTACGGTCACACTG | chrX:384558-386558 | 1585 | 1599 | + | 3.3571 | 7.12E-05 | 0.795 | GCTAAGGAGACACGG |
| 1 | GMTACGGTCACACTG | chr2L:11010312-11012312 | 1586 | 1600 | + | 3.3571 | 7.12E-05 | 0.795 | GATTCGGTTCCACCG |
| 1 | GMTACGGTCACACTG | chr2L:12427028-12429028 | 1586 | 1600 | + | 3.3571 | 7.12E-05 | 0.795 | GCTATTGGCACACGG |
| 1 | GMTACGGTCACACTG | chr3L:19518649-19520649 | 1587 | 1601 | + | 3.3571 | 7.12E-05 | 0.795 | GATACGGACACGGAG |

| | | | | | | | | | |
|---|---|---|---|---|---|---|---|---|---|
| 1 | GMTACGGTCACACTG | chr3L:4363102-4365102 | 1587 | 1601 | + | 3.3571 | 7.12E-05 | 0.795 | TCACCGGTCACAATG |
| 1 | GMTACGGTCACACTG | chr2L:1727732-1729732 | 1588 | 1602 | + | 3.3571 | 7.12E-05 | 0.795 | GCTGCGGGAAAACTG |
| 1 | GMTACGGTCACACTG | chrX:11347120-11349120 | 1589 | 1603 | + | 3.3571 | 7.12E-05 | 0.795 | GATGCGGACGCAATG |
| 1 | GMTACGGTCACACTG | chr3R:24916864-24918864 | 1592 | 1606 | + | 3.3571 | 7.12E-05 | 0.795 | GCTAGGGCCAAACAG |
| 1 | GMTACGGTCACACTG | chr3R:24916864-24918864 | 1592 | 1606 | + | 3.3571 | 7.12E-05 | 0.795 | GCTAGGGCCAAACAG |
| 1 | GMTACGGTCACACTG | chrX:16883581-16885581 | 1594 | 1608 | + | 3.3571 | 7.12E-05 | 0.795 | GCTAAGGAAACACCG |
| 1 | GMTACGGTCACACTG | chrY:3021289-3023289 | 1594 | 1608 | + | 3.3571 | 7.12E-05 | 0.795 | GCTACGGAAACAAGG |
| 1 | GMTACGGTCACACTG | chrX:13781459-13783459 | 1595 | 1609 | + | 3.3571 | 7.12E-05 | 0.795 | GCTGTGGGCAAACTG |
| 1 | GMTACGGTCACACTG | chr3R:10463978-10465978 | 1595 | 1609 | + | 3.3571 | 7.12E-05 | 0.795 | GATCTGGGCACACAG |
| 1 | GMTACGGTCACACTG | chr2L:12930551-12932551 | 1600 | 1614 | + | 3.3571 | 7.12E-05 | 0.795 | GCCACAGTCCGACTG |
| 1 | GMTACGGTCACACTG | chr2L:6959377-6961377 | 1600 | 1614 | + | 3.3571 | 7.12E-05 | 0.795 | GCTACGTTTCCACCG |
| 1 | GMTACGGTCACACTG | chr3R:22579773-22581773 | 1601 | 1615 | + | 3.3571 | 7.12E-05 | 0.795 | GCGACAATCCCACTG |
| 1 | GMTACGGTCACACTG | chr3R:15264591-15266591 | 1603 | 1617 | + | 3.3571 | 7.12E-05 | 0.795 | GCCACCGTCAATCTG |
| 1 | GMTACGGTCACACTG | chr2R:16771671-16773671 | 1605 | 1619 | + | 3.3571 | 7.12E-05 | 0.795 | GCCACGGTCAGCGTG |
| 1 | GMTACGGTCACACTG | chr3R:5958555-5960555 | 1606 | 1620 | + | 3.3571 | 7.12E-05 | 0.795 | GCTTCAGGCAAACTG |
| 1 | GMTACGGTCACACTG | chrX:1341106-1343106 | 1606 | 1620 | + | 3.3571 | 7.12E-05 | 0.795 | GCCATGGTCACAAGG |
| 1 | GMTACGGTCACACTG | chr2L:11735402-11737402 | 1606 | 1620 | + | 3.3571 | 7.12E-05 | 0.795 | GCAACAAACACACTG |
| 1 | GMTACGGTCACACTG | chr2L:6488090-6490090 | 1608 | 1622 | + | 3.3571 | 7.12E-05 | 0.795 | GCTACGCTGACTCAG |
| 1 | GMTACGGTCACACTG | chrX:10216627-10218627 | 1608 | 1622 | + | 3.3571 | 7.12E-05 | 0.795 | ACCACAGGCACACTG |
| 1 | GMTACGGTCACACTG | chr3R:28600529-28602529 | 1610 | 1624 | + | 3.3571 | 7.12E-05 | 0.795 | GCTACGATTCCACAG |
| 1 | GMTACGGTCACACTG | chr3R:29798354-29800354 | 1611 | 1625 | + | 3.3571 | 7.12E-05 | 0.795 | CCAACGATCACACAG |
| 1 | GMTACGGTCACACTG | chr3R:29688311-29690311 | 1611 | 1625 | + | 3.3571 | 7.12E-05 | 0.795 | TCTACTGACACACGG |
| 1 | GMTACGGTCACACTG | chr3L:23985410-23987410 | 1612 | 1626 | + | 3.3571 | 7.12E-05 | 0.795 | GCCACGCCCACACTT |
| 1 | GMTACGGTCACACTG | chr3R:17724063-17726063 | 1612 | 1626 | + | 3.3571 | 7.12E-05 | 0.795 | GCAACGGCAACAATG |
| 1 | GMTACGGTCACACTG | chrX:18498986-18500986 | 1613 | 1627 | + | 3.3571 | 7.12E-05 | 0.795 | GCCATGGACACACTC |
| 1 | GMTACGGTCACACTG | chr2L:22536668-22538668 | 1614 | 1628 | + | 3.3571 | 7.12E-05 | 0.795 | GCCAAGGTCGCACTT |
| 1 | GMTACGGTCACACTG | chrX:12407477-12409477 | 1614 | 1628 | + | 3.3571 | 7.12E-05 | 0.795 | GCACCAGTCACAGTG |
| 1 | GMTACGGTCACACTG | chrX:5746747-5748747 | 1616 | 1630 | + | 3.3571 | 7.12E-05 | 0.795 | GCTACGGGTGCACTC |
| 1 | GMTACGGTCACACTG | chr3R:23537908-23539908 | 1618 | 1632 | + | 3.3571 | 7.12E-05 | 0.795 | GCAATGGCCAGACTG |
| 1 | GMTACGGTCACACTG | chr2R:7394932-7396932 | 1618 | 1632 | + | 3.3571 | 7.12E-05 | 0.795 | GCCACGCCCACACTT |
| 1 | GMTACGGTCACACTG | chr2L:18256978-18258978 | 1619 | 1633 | + | 3.3571 | 7.12E-05 | 0.795 | GCAACGGTCATGCTC |
| 1 | GMTACGGTCACACTG | chr3R:17538963-17540963 | 1619 | 1633 | + | 3.3571 | 7.12E-05 | 0.795 | GCTACAGTCAGTCAG |
| 1 | GMTACGGTCACACTG | chr3L:22224878-22226878 | 1621 | 1635 | + | 3.3571 | 7.12E-05 | 0.795 | GCTGCTGCCACAATG |

| | | | | | | | | | |
|---|---|---|---|---|---|---|---|---|---|
| 1 | GMTACGGTCACACTG | chr2R:7931429-7933429 | 1622 | 1636 | + | 3.3571 | 7.12E-05 | 0.795 | GACACGGACACTATG |
| 1 | GMTACGGTCACACTG | chr3R:9399053-9401053 | 1624 | 1638 | + | 3.3571 | 7.12E-05 | 0.795 | GATACGCTCGCGCAG |
| 1 | GMTACGGTCACACTG | chr3R:29514717-29516717 | 1630 | 1644 | + | 3.3571 | 7.12E-05 | 0.795 | GCTGCGATCGCATTG |
| 1 | GMTACGGTCACACTG | chr3L:5872311-5874311 | 1632 | 1646 | + | 3.3571 | 7.12E-05 | 0.795 | GCTTGGGGCAAACTG |
| 1 | GMTACGGTCACACTG | chr3L:2994172-2996172 | 1635 | 1649 | + | 3.3571 | 7.12E-05 | 0.795 | GCTGCGATTTCACTG |
| 1 | GMTACGGTCACACTG | chr2L:8398021-8400021 | 1635 | 1649 | + | 3.3571 | 7.12E-05 | 0.795 | GACCGGGTCACGCTG |
| 1 | GMTACGGTCACACTG | chr2L:5922653-5924653 | 1641 | 1655 | + | 3.3571 | 7.12E-05 | 0.795 | CCCCCGCTCACACTG |
| 1 | GMTACGGTCACACTG | chr2L:4120702-4122702 | 1641 | 1655 | + | 3.3571 | 7.12E-05 | 0.795 | GCCACGATCACACGT |
| 1 | GMTACGGTCACACTG | chr2L:13444651-13446651 | 1642 | 1656 | + | 3.3571 | 7.12E-05 | 0.795 | GACATGGCCGCACTG |
| 1 | GMTACGGTCACACTG | chr3R:21785208-21787208 | 1643 | 1657 | + | 3.3571 | 7.12E-05 | 0.795 | GCTGCCGTCGTACTG |
| 1 | GMTACGGTCACACTG | chr2R:24194498-24196498 | 1644 | 1658 | + | 3.3571 | 7.12E-05 | 0.795 | GAGACGGACAAGCTG |
| 1 | GMTACGGTCACACTG | chr3L:17548826-17550826 | 1645 | 1659 | + | 3.3571 | 7.12E-05 | 0.795 | GCCATGGTGACTCTG |
| 1 | GMTACGGTCACACTG | chr2L:6959328-6961328 | 1649 | 1663 | + | 3.3571 | 7.12E-05 | 0.795 | GCTACGTTTCCACCG |
| 1 | GMTACGGTCACACTG | chr2L:4452038-4454038 | 1650 | 1664 | + | 3.3571 | 7.12E-05 | 0.795 | GCTGCAGTCATACCG |
| 1 | GMTACGGTCACACTG | chrX:9255168-9257168 | 1652 | 1666 | + | 3.3571 | 7.12E-05 | 0.795 | CCTGCGGCAACACTG |
| 1 | GMTACGGTCACACTG | chrX:9148309-9150309 | 1653 | 1667 | + | 3.3571 | 7.12E-05 | 0.795 | GACACCGTCACGCGG |
| 1 | GMTACGGTCACACTG | chr2R:16343182-16345182 | 1654 | 1668 | + | 3.3571 | 7.12E-05 | 0.795 | GCTCCCCTCACGCTG |
| 1 | GMTACGGTCACACTG | chr3R:16429681-16431681 | 1658 | 1672 | + | 3.3571 | 7.12E-05 | 0.795 | GCTACGGATGCAGTG |
| 1 | GMTACGGTCACACTG | chr2R:11981956-11983956 | 1660 | 1674 | + | 3.3571 | 7.12E-05 | 0.795 | GCTACGAACTCAATG |
| 1 | GMTACGGTCACACTG | chr2L:19053078-19055078 | 1663 | 1677 | + | 3.3571 | 7.12E-05 | 0.795 | GAGACGGGCCAACTG |
| 1 | GMTACGGTCACACTG | chr3R:15791579-15793579 | 1665 | 1679 | + | 3.3571 | 7.12E-05 | 0.795 | GCTAAGGTCTCAAAG |
| 1 | GMTACGGTCACACTG | chr2R:17544222-17546222 | 1669 | 1683 | + | 3.3571 | 7.12E-05 | 0.795 | GCTACGGCTACAAAG |
| 1 | GMTACGGTCACACTG | chr2L:452108-454108 | 1670 | 1684 | + | 3.3571 | 7.12E-05 | 0.795 | GAGATGGTCACCCGG |
| 1 | GMTACGGTCACACTG | chr2R:7931380-7933380 | 1671 | 1685 | + | 3.3571 | 7.12E-05 | 0.795 | GACACGGACACTATG |
| 1 | GMTACGGTCACACTG | chrX:5753958-5755958 | 1672 | 1686 | + | 3.3571 | 7.12E-05 | 0.795 | GCTCCGCTCTCACTC |
| 1 | GMTACGGTCACACTG | chr2R:20262979-20264979 | 1672 | 1686 | + | 3.3571 | 7.12E-05 | 0.795 | GATCCGCTCCCACCG |
| 1 | GMTACGGTCACACTG | chrX:19938396-19940396 | 1673 | 1687 | + | 3.3571 | 7.12E-05 | 0.795 | GCTGCGAGCACACTT |
| 1 | GMTACGGTCACACTG | chrX:19928580-19930580 | 1673 | 1687 | + | 3.3571 | 7.12E-05 | 0.795 | GCTGCGAGCACACTT |
| 1 | GMTACGGTCACACTG | chrX:19914685-19916685 | 1673 | 1687 | + | 3.3571 | 7.12E-05 | 0.795 | GCTGCGAGCACACTT |
| 1 | GMTACGGTCACACTG | chrX:19909521-19911521 | 1673 | 1687 | + | 3.3571 | 7.12E-05 | 0.795 | GCTGCGAGCACACTT |
| 1 | GMTACGGTCACACTG | chr2R:24598947-24600947 | 1674 | 1688 | + | 3.3571 | 7.12E-05 | 0.795 | GATCCGGTCACCTCG |
| 1 | GMTACGGTCACACTG | chrX:7148850-7150850 | 1678 | 1692 | + | 3.3571 | 7.12E-05 | 0.795 | CCTACGTTCGCTCTG |
| 1 | GMTACGGTCACACTG | chr3L:7935120-7937120 | 1679 | 1693 | + | 3.3571 | 7.12E-05 | 0.795 | GCCACGGCGACACTT |

| | | | | | | | | | |
|---|---|---|---|---|---|---|---|---|---|
| 1 | GMTACGGTCACACTG | chr2L:20064679-20066679 | 1684 | 1698 | + | 3.3571 | 7.12E-05 | 0.795 | CAAGCGGTCACACAG |
| 1 | GMTACGGTCACACTG | chr3R:24943956-24945956 | 1686 | 1700 | + | 3.3571 | 7.12E-05 | 0.795 | GCCATGGTCAAGCTG |
| 1 | GMTACGGTCACACTG | chrX:10767338-10769338 | 1692 | 1706 | + | 3.3571 | 7.12E-05 | 0.795 | TACTCGGGCACACTG |
| 1 | GMTACGGTCACACTG | chr3R:24924216-24926216 | 1694 | 1708 | + | 3.3571 | 7.12E-05 | 0.795 | GATCCGGACACTGTG |
| 1 | GMTACGGTCACACTG | chr3R:17133197-17135197 | 1694 | 1708 | + | 3.3571 | 7.12E-05 | 0.795 | GATACGGCAGCACGG |
| 1 | GMTACGGTCACACTG | chrX:19851059-19853059 | 1695 | 1709 | + | 3.3571 | 7.12E-05 | 0.795 | GAGGCGGTCATTCTG |
| 1 | GMTACGGTCACACTG | chr3L:17041797-17043797 | 1695 | 1709 | + | 3.3571 | 7.12E-05 | 0.795 | GCTGCTCTCACGCTG |
| 1 | GMTACGGTCACACTG | chr2R:8905990-8907990 | 1700 | 1714 | + | 3.3571 | 7.12E-05 | 0.795 | GAGACGAACACGCTG |
| 1 | GMTACGGTCACACTG | chr3R:20576053-20578053 | 1701 | 1715 | + | 3.3571 | 7.12E-05 | 0.795 | TCTACGGACAGACGG |
| 1 | GMTACGGTCACACTG | chr3L:8911671-8913671 | 1705 | 1719 | + | 3.3571 | 7.12E-05 | 0.795 | GCTACGGTACCGTTG |
| 1 | GMTACGGTCACACTG | chr3L:1033697-1035697 | 1707 | 1721 | + | 3.3571 | 7.12E-05 | 0.795 | GAATCGGCCAAACTG |
| 1 | GMTACGGTCACACTG | chr3R:22024506-22026506 | 1708 | 1722 | + | 3.3571 | 7.12E-05 | 0.795 | GCTACGCGCATTCTG |
| 1 | GMTACGGTCACACTG | chr3R:17131662-17133662 | 1708 | 1722 | + | 3.3571 | 7.12E-05 | 0.795 | GCCATGGCGACACTG |
| 1 | GMTACGGTCACACTG | chrX:11347001-11349001 | 1708 | 1722 | + | 3.3571 | 7.12E-05 | 0.795 | GATGCGGACGCAATG |
| 1 | GMTACGGTCACACTG | chrX:19682808-19684808 | 1710 | 1724 | + | 3.3571 | 7.12E-05 | 0.795 | GACACGGACACCCTT |
| 1 | GMTACGGTCACACTG | chr2R:22112589-22114589 | 1711 | 1725 | + | 3.3571 | 7.12E-05 | 0.795 | CCTCCGCTCACACCG |
| 1 | GMTACGGTCACACTG | chr2R:24268179-24270179 | 1712 | 1726 | + | 3.3571 | 7.12E-05 | 0.795 | GCCACCGTCGCATTG |
| 1 | GMTACGGTCACACTG | chrX:13667014-13669014 | 1716 | 1730 | + | 3.3571 | 7.12E-05 | 0.795 | ACTGCGGCCACAGTG |
| 1 | GMTACGGTCACACTG | chr2R:12756546-12758546 | 1717 | 1731 | + | 3.3571 | 7.12E-05 | 0.795 | GCTACACTCGCGCTG |
| 1 | GMTACGGTCACACTG | chr3R:10704075-10706075 | 1719 | 1733 | + | 3.3571 | 7.12E-05 | 0.795 | GAACCGGGCACAATG |
| 1 | GMTACGGTCACACTG | chrX:7716932-7718932 | 1720 | 1734 | + | 3.3571 | 7.12E-05 | 0.795 | GCTACGGGCAACGTG |
| 1 | GMTACGGTCACACTG | chrX:10347170-10349170 | 1722 | 1736 | + | 3.3571 | 7.12E-05 | 0.795 | GCACCGGTCAAATTG |
| 1 | GMTACGGTCACACTG | chr3L:16170380-16172380 | 1723 | 1737 | + | 3.3571 | 7.12E-05 | 0.795 | GCGAAAGTCGCACTG |
| 1 | GMTACGGTCACACTG | chr3R:16603861-16605861 | 1731 | 1745 | + | 3.3571 | 7.12E-05 | 0.795 | GATCCGATCGCGCTG |
| 1 | GMTACGGTCACACTG | chr2R:12756529-12758529 | 1734 | 1748 | + | 3.3571 | 7.12E-05 | 0.795 | GCTACACTCGCGCTG |
| 1 | GMTACGGTCACACTG | chr3L:2009751-2011751 | 1737 | 1751 | + | 3.3571 | 7.12E-05 | 0.795 | GCAACCGTCATTCTG |
| 1 | GMTACGGTCACACTG | chr2R:23365866-23367866 | 1738 | 1752 | + | 3.3571 | 7.12E-05 | 0.795 | GCTAAGGGAGCACTG |
| 1 | GMTACGGTCACACTG | chr2L:20074757-20076757 | 1743 | 1757 | + | 3.3571 | 7.12E-05 | 0.795 | CCTACGGCCAGACGG |
| 1 | GMTACGGTCACACTG | chr3R:20576010-20578010 | 1744 | 1758 | + | 3.3571 | 7.12E-05 | 0.795 | TCTACGGACAGACGG |
| 1 | GMTACGGTCACACTG | chr2R:10810078-10812078 | 1746 | 1760 | + | 3.3571 | 7.12E-05 | 0.795 | CCAATGGTCACGCTG |
| 1 | GMTACGGTCACACTG | chr2R:6688974-6690974 | 1750 | 1764 | + | 3.3571 | 7.12E-05 | 0.795 | GATCCGGCTGCACTG |
| 1 | GMTACGGTCACACTG | chr3L:11681038-11683038 | 1751 | 1765 | + | 3.3571 | 7.12E-05 | 0.795 | GACATGGCCACGCTG |
| 1 | GMTACGGTCACACTG | chr2R:17244501-17246501 | 1752 | 1766 | + | 3.3571 | 7.12E-05 | 0.795 | GATGCGGACTCACTC |

| | | | | | | | | | |
|---|---|---|---|---|---|---|---|---|---|
| 1 | GMTACGGTCACACTG | chrX:126449-128449 | 1753 | 1767 | + | 3.3571 | 7.12E-05 | 0.795 | GCTACTGTCGCACCT |
| 1 | GMTACGGTCACACTG | chr2L:452024-454024 | 1754 | 1768 | + | 3.3571 | 7.12E-05 | 0.795 | GAGATGGTCACCCGG |
| 1 | GMTACGGTCACACTG | chr2L:19536965-19538965 | 1759 | 1773 | + | 3.3571 | 7.12E-05 | 0.795 | GATGAGGACGCACTG |
| 1 | GMTACGGTCACACTG | chrX:1878774-1880774 | 1760 | 1774 | + | 3.3571 | 7.12E-05 | 0.795 | CAGTCGGCCACACTG |
| 1 | GMTACGGTCACACTG | chr2R:9888878-9890878 | 1763 | 1777 | + | 3.3571 | 7.12E-05 | 0.795 | TCCACGGTAACGCTG |
| 1 | GMTACGGTCACACTG | chr3R:23695245-23697245 | 1765 | 1779 | + | 3.3571 | 7.12E-05 | 0.795 | GCTGCCCTCACCCTG |
| 1 | GMTACGGTCACACTG | chr2R:23231626-23233626 | 1773 | 1787 | + | 3.3571 | 7.12E-05 | 0.795 | GATACGGCCACCGAG |
| 1 | GMTACGGTCACACTG | chr3L:14700678-14702678 | 1773 | 1787 | + | 3.3571 | 7.12E-05 | 0.795 | GCTACGCAAACACGG |
| 1 | GMTACGGTCACACTG | chr2L:18804347-18806347 | 1774 | 1788 | + | 3.3571 | 7.12E-05 | 0.795 | GCTACCACCACACAG |
| 1 | GMTACGGTCACACTG | chr3L:23173446-23175446 | 1777 | 1791 | + | 3.3571 | 7.12E-05 | 0.795 | GCCACGCCCACACTT |
| 1 | GMTACGGTCACACTG | chr2R:7518280-7520280 | 1779 | 1793 | + | 3.3571 | 7.12E-05 | 0.795 | GCCTCGATCACATTG |
| 1 | GMTACGGTCACACTG | chr2L:6224048-6226048 | 1780 | 1794 | + | 3.3571 | 7.12E-05 | 0.795 | GCGACGATCCCACTC |
| 1 | GMTACGGTCACACTG | chrX:4701284-4703284 | 1780 | 1794 | + | 3.3571 | 7.12E-05 | 0.795 | GACAAGGTCACGCAG |
| 1 | GMTACGGTCACACTG | chr3R:29368338-29370338 | 1780 | 1794 | + | 3.3571 | 7.12E-05 | 0.795 | GCCACGCCCACAGTG |
| 1 | GMTACGGTCACACTG | chr3L:11772012-11774012 | 1781 | 1795 | + | 3.3571 | 7.12E-05 | 0.795 | TCTCCGGGCAAACTG |
| 1 | GMTACGGTCACACTG | chr3L:8575078-8577078 | 1782 | 1796 | + | 3.3571 | 7.12E-05 | 0.795 | GCTTCGGTCTAAATG |
| 1 | GMTACGGTCACACTG | chr3R:18977809-18979809 | 1782 | 1796 | + | 3.3571 | 7.12E-05 | 0.795 | GCTTCGTTCACAAAG |
| 1 | GMTACGGTCACACTG | chrX:6236315-6238315 | 1787 | 1801 | + | 3.3571 | 7.12E-05 | 0.795 | GCTACAGGCATTCTG |
| 1 | GMTACGGTCACACTG | chr3L:16608347-16610347 | 1790 | 1804 | + | 3.3571 | 7.12E-05 | 0.795 | GCGATGGCCACAATG |
| 1 | GMTACGGTCACACTG | chr3R:30958700-30960700 | 1793 | 1807 | + | 3.3571 | 7.12E-05 | 0.795 | CCTCTGGTCACGCTG |
| 1 | GMTACGGTCACACTG | chr3R:11153626-11155626 | 1794 | 1808 | + | 3.3571 | 7.12E-05 | 0.795 | GCTTGGGTGCCACTG |
| 1 | GMTACGGTCACACTG | chrX:1144644-1146644 | 1794 | 1808 | + | 3.3571 | 7.12E-05 | 0.795 | GCAACGCCCACACTT |
| 1 | GMTACGGTCACACTG | chr3R:25489936-25491936 | 1798 | 1812 | + | 3.3571 | 7.12E-05 | 0.795 | GATACGGGAACGCAG |
| 1 | GMTACGGTCACACTG | chr2R:24598821-24600821 | 1800 | 1814 | + | 3.3571 | 7.12E-05 | 0.795 | GATCCGGTCACCTCG |
| 1 | GMTACGGTCACACTG | chr3L:7237542-7239542 | 1803 | 1817 | + | 3.3571 | 7.12E-05 | 0.795 | GCTACCGCCACAAAG |
| 1 | GMTACGGTCACACTG | chr2R:6599978-6601978 | 1805 | 1819 | + | 3.3571 | 7.12E-05 | 0.795 | GAATCTGCCACACTG |
| 1 | GMTACGGTCACACTG | chr2R:22121101-22123101 | 1806 | 1820 | + | 3.3571 | 7.12E-05 | 0.795 | ACAACGGTCACTCTA |
| 1 | GMTACGGTCACACTG | chr3R:18365162-18367162 | 1807 | 1821 | + | 3.3571 | 7.12E-05 | 0.795 | GACAAGGACACGCTG |
| 1 | GMTACGGTCACACTG | chr3L:18916581-18918581 | 1808 | 1822 | + | 3.3571 | 7.12E-05 | 0.795 | GCAACGGTGGCATTG |
| 1 | GMTACGGTCACACTG | chrX:8408272-8410272 | 1810 | 1824 | + | 3.3571 | 7.12E-05 | 0.795 | GCGACTCACACACTG |
| 1 | GMTACGGTCACACTG | chrX:16883365-16885365 | 1810 | 1824 | + | 3.3571 | 7.12E-05 | 0.795 | GCTAAGGAAACACCG |
| 1 | GMTACGGTCACACTG | chrX:9604969-9606969 | 1810 | 1824 | + | 3.3571 | 7.12E-05 | 0.795 | GAAACGCTCGCGCTG |
| 1 | GMTACGGTCACACTG | chr2L:16972091-16974091 | 1811 | 1825 | + | 3.3571 | 7.12E-05 | 0.795 | GATACGGGCCAACAG |

| | | | | | | | | | |
|---|---|---|---|---|---|---|---|---|---|
| 1 | GMTACGGTCACACTG | chr2R:22328720-22330720 | 1811 | 1825 | + | 3.3571 | 7.12E-05 | 0.795 | GATCCGGACAAGCTG |
| 1 | GMTACGGTCACACTG | chr2L:11010086-11012086 | 1812 | 1826 | + | 3.3571 | 7.12E-05 | 0.795 | GATTCGGTTCCACCG |
| 1 | GMTACGGTCACACTG | chrX:15471164-15473164 | 1812 | 1826 | + | 3.3571 | 7.12E-05 | 0.795 | CCTACGGTCTCTTTG |
| 1 | GMTACGGTCACACTG | chr2R:16013908-16015908 | 1814 | 1828 | + | 3.3571 | 7.12E-05 | 0.795 | GAGACGGTTCCGCTG |
| 1 | GMTACGGTCACACTG | chr2R:23902750-23904750 | 1821 | 1835 | + | 3.3571 | 7.12E-05 | 0.795 | CCTACGGCCACACCA |
| 1 | GMTACGGTCACACTG | chr3R:19397567-19399567 | 1826 | 1840 | + | 3.3571 | 7.12E-05 | 0.795 | GATGCGGAAACTCTG |
| 1 | GMTACGGTCACACTG | chr3L:222852-224852 | 1826 | 1840 | + | 3.3571 | 7.12E-05 | 0.795 | GCAACAGTTACACGG |
| 1 | GMTACGGTCACACTG | chr2L:15886905-15888905 | 1827 | 1841 | + | 3.3571 | 7.12E-05 | 0.795 | GCAAATGTCACACAG |
| 1 | GMTACGGTCACACTG | chr2L:23024915-23026915 | 1828 | 1842 | + | 3.3571 | 7.12E-05 | 0.795 | GCTACAGCCACTCTA |
| 1 | GMTACGGTCACACTG | chr2L:8520933-8522933 | 1829 | 1843 | + | 3.3571 | 7.12E-05 | 0.795 | GCTACCGCCACCGTG |
| 1 | GMTACGGTCACACTG | chr2L:16336885-16338885 | 1831 | 1845 | + | 3.3571 | 7.12E-05 | 0.795 | GCATCGGTCACAGTT |
| 1 | GMTACGGTCACACTG | chr2R:12841665-12843665 | 1831 | 1845 | + | 3.3571 | 7.12E-05 | 0.795 | GCTAGGGGCAATCTG |
| 1 | GMTACGGTCACACTG | chr2L:6722903-6724903 | 1831 | 1845 | + | 3.3571 | 7.12E-05 | 0.795 | GCTTCGGTAACAGCG |
| 1 | GMTACGGTCACACTG | chr2L:14007101-14009101 | 1833 | 1847 | + | 3.3571 | 7.12E-05 | 0.795 | GCAACTGTCGCATTG |
| 1 | GMTACGGTCACACTG | chr3R:7912290-7914290 | 1836 | 1850 | + | 3.3571 | 7.12E-05 | 0.795 | GAGACCGGCACCCTG |
| 1 | GMTACGGTCACACTG | chrX:16885818-16887818 | 1836 | 1850 | + | 3.3571 | 7.12E-05 | 0.795 | GCAAGAGCCACACTG |
| 1 | GMTACGGTCACACTG | chr3R:27962415-27964415 | 1837 | 1851 | + | 3.3571 | 7.12E-05 | 0.795 | TACACGGACACCCTG |
| 1 | GMTACGGTCACACTG | chrX:13542643-13544643 | 1838 | 1852 | + | 3.3571 | 7.12E-05 | 0.795 | GCTACGCCCACAGCG |
| 1 | GMTACGGTCACACTG | chr2R:24602572-24604572 | 1839 | 1853 | + | 3.3571 | 7.12E-05 | 0.795 | GCTACGGGTACGCTT |
| 1 | GMTACGGTCACACTG | chr2R:6035388-6037388 | 1841 | 1855 | + | 3.3571 | 7.12E-05 | 0.795 | GATCCGGTCAGGCCG |
| 1 | GMTACGGTCACACTG | chr3L:908112-910112 | 1841 | 1855 | + | 3.3571 | 7.12E-05 | 0.795 | GCAACGGCGACATTG |
| 1 | GMTACGGTCACACTG | chr3R:24900116-24902116 | 1842 | 1856 | + | 3.3571 | 7.12E-05 | 0.795 | GACCCGGTGACACCG |
| 1 | GMTACGGTCACACTG | chrX:1574132-1576132 | 1842 | 1856 | + | 3.3571 | 7.12E-05 | 0.795 | GCTTAGCACACACTG |
| 1 | GMTACGGTCACACTG | chr2R:8678953-8680953 | 1844 | 1858 | + | 3.3571 | 7.12E-05 | 0.795 | CCTATGGACGCACTG |
| 1 | GMTACGGTCACACTG | chrX:6195439-6197439 | 1845 | 1859 | + | 3.3571 | 7.12E-05 | 0.795 | GCCACCGTGACACAG |
| 1 | GMTACGGTCACACTG | chr2L:12977786-12979786 | 1846 | 1860 | + | 3.3571 | 7.12E-05 | 0.795 | GATACGCCCTCCCTG |
| 1 | GMTACGGTCACACTG | chr3R:22030900-22032900 | 1848 | 1862 | + | 3.3571 | 7.12E-05 | 0.795 | GCGAGTGTCCCACTG |
| 1 | GMTACGGTCACACTG | chr3R:5634624-5636624 | 1852 | 1866 | + | 3.3571 | 7.12E-05 | 0.795 | GACGCGGTCGTACTG |
| 1 | GMTACGGTCACACTG | chr2L:154305-156305 | 1856 | 1870 | + | 3.3571 | 7.12E-05 | 0.795 | ACTCTGGTCACACAG |
| 1 | GMTACGGTCACACTG | chr3L:7338340-7340340 | 1856 | 1870 | + | 3.3571 | 7.12E-05 | 0.795 | GCTTCGTTCACATGG |
| 1 | GMTACGGTCACACTG | chrX:15471119-15473119 | 1857 | 1871 | + | 3.3571 | 7.12E-05 | 0.795 | CCTACGGTCTCTTTG |
| 1 | GMTACGGTCACACTG | chr2R:8984900-8986900 | 1858 | 1872 | + | 3.3571 | 7.12E-05 | 0.795 | GCCACGGTGACCATG |
| 1 | GMTACGGTCACACTG | chr2R:22196374-22198374 | 1862 | 1876 | + | 3.3571 | 7.12E-05 | 0.795 | CCTACGGTCTCAGGG |

| | | | | | | | | | |
|---|---|---|---|---|---|---|---|---|---|
| 1 | GMTACGGTCACACTG | chr3L:21958254-21960254 | 1863 | 1877 | + | 3.3571 | 7.12E-05 | 0.795 | GCTACCTTCGCCCTG |
| 1 | GMTACGGTCACACTG | chr3L:1617414-1619414 | 1863 | 1877 | + | 3.3571 | 7.12E-05 | 0.795 | GCTAGGGACTCACTC |
| 1 | GMTACGGTCACACTG | chr3L:1617414-1619414 | 1863 | 1877 | + | 3.3571 | 7.12E-05 | 0.795 | GCTAGGGACTCACTC |
| 1 | GMTACGGTCACACTG | chrX:18100314-18102314 | 1866 | 1880 | + | 3.3571 | 7.12E-05 | 0.795 | GAGACGCTCACCCAG |
| 1 | GMTACGGTCACACTG | chr3R:27111270-27113270 | 1868 | 1882 | + | 3.3571 | 7.12E-05 | 0.795 | GCTACGTGTCCACTG |
| 1 | GMTACGGTCACACTG | chr3L:8911499-8913499 | 1877 | 1891 | + | 3.3571 | 7.12E-05 | 0.795 | GCTACGGTACCGTTG |
| 1 | GMTACGGTCACACTG | chrX:23282731-23284731 | 1883 | 1897 | + | 3.3571 | 7.12E-05 | 0.795 | GACACGGCCATTCTG |
| 1 | GMTACGGTCACACTG | chr3L:12660826-12662826 | 1883 | 1897 | + | 3.3571 | 7.12E-05 | 0.795 | GCTCAAGTCTCACTG |
| 1 | GMTACGGTCACACTG | chr3R:4265770-4267770 | 1884 | 1898 | + | 3.3571 | 7.12E-05 | 0.795 | GCTACGAACACAACG |
| 1 | GMTACGGTCACACTG | chrX:13781170-13783170 | 1884 | 1898 | + | 3.3571 | 7.12E-05 | 0.795 | GCTGTGGGCAAACTG |
| 1 | GMTACGGTCACACTG | chr3R:9407169-9409169 | 1884 | 1898 | + | 3.3571 | 7.12E-05 | 0.795 | GCGACGGTCAGATAG |
| 1 | GMTACGGTCACACTG | chr3L:17884528-17886528 | 1885 | 1899 | + | 3.3571 | 7.12E-05 | 0.795 | GCTCCGGTACCACTC |
| 1 | GMTACGGTCACACTG | chrY:191020-193020 | 1885 | 1899 | + | 3.3571 | 7.12E-05 | 0.795 | GCAACGCTCACTCTA |
| 1 | GMTACGGTCACACTG | chr3R:14922761-14924761 | 1886 | 1900 | + | 3.3571 | 7.12E-05 | 0.795 | GAAACGGTCACTTAG |
| 1 | GMTACGGTCACACTG | chr2R:22217500-22219500 | 1887 | 1901 | + | 3.3571 | 7.12E-05 | 0.795 | GCCACCGTCACAAAG |
| 1 | GMTACGGTCACACTG | chr3R:27962415-27964415 | 1888 | 1902 | + | 3.3571 | 7.12E-05 | 0.795 | ACGCCGGTCACACTT |
| 1 | GMTACGGTCACACTG | chr2R:10809936-10811936 | 1888 | 1902 | + | 3.3571 | 7.12E-05 | 0.795 | CCAATGGTCACGCTG |
| 1 | GMTACGGTCACACTG | chrX:9148071-9150071 | 1891 | 1905 | + | 3.3571 | 7.12E-05 | 0.795 | GACACCGTCACGCGG |
| 1 | GMTACGGTCACACTG | chr3R:29724830-29726830 | 1899 | 1913 | + | 3.3571 | 7.12E-05 | 0.795 | GATGCGAGCACACGG |
| 1 | GMTACGGTCACACTG | chr3R:18757873-18759873 | 1901 | 1915 | + | 3.3571 | 7.12E-05 | 0.795 | GAAACGCTCGCACAG |
| 1 | GMTACGGTCACACTG | chr2L:14026001-14028001 | 1901 | 1915 | + | 3.3571 | 7.12E-05 | 0.795 | GACAAGGACGCACTG |
| 1 | GMTACGGTCACACTG | chr3L:23205669-23207669 | 1901 | 1915 | + | 3.3571 | 7.12E-05 | 0.795 | GCCACGTCCACACTT |
| 1 | GMTACGGTCACACTG | chr2R:16097564-16099564 | 1903 | 1917 | + | 3.3571 | 7.12E-05 | 0.795 | ACGAAGGGCACACTG |
| 1 | GMTACGGTCACACTG | chr2L:12426709-12428709 | 1905 | 1919 | + | 3.3571 | 7.12E-05 | 0.795 | GCTATTGGCACACGG |
| 1 | GMTACGGTCACACTG | chr2L:7083635-7085635 | 1905 | 1919 | + | 3.3571 | 7.12E-05 | 0.795 | GCTACCGTCAATCAG |
| 1 | GMTACGGTCACACTG | chr3R:10224690-10226690 | 1908 | 1922 | + | 3.3571 | 7.12E-05 | 0.795 | GCTACCGTCACGCAC |
| 1 | GMTACGGTCACACTG | chr3R:8768394-8770394 | 1909 | 1923 | + | 3.3571 | 7.12E-05 | 0.795 | TCCATGGTCACGCTG |
| 1 | GMTACGGTCACACTG | chr2L:13722642-13724642 | 1910 | 1924 | + | 3.3571 | 7.12E-05 | 0.795 | GCAACTGTCAGTCTG |
| 1 | GMTACGGTCACACTG | chr2L:19182787-19184787 | 1913 | 1927 | + | 3.3571 | 7.12E-05 | 0.795 | GAGGCGGTCAGACCG |
| 1 | GMTACGGTCACACTG | chr3L:22842264-22844264 | 1914 | 1928 | + | 3.3571 | 7.12E-05 | 0.795 | GCTGCAATCGCACTG |
| 1 | GMTACGGTCACACTG | chr2L:5768752-5770752 | 1915 | 1929 | + | 3.3571 | 7.12E-05 | 0.795 | GCCGCCGTCAGACTG |
| 1 | GMTACGGTCACACTG | chr3R:26965930-26967930 | 1915 | 1929 | + | 3.3571 | 7.12E-05 | 0.795 | GCTCCGCTCTCAATG |
| 1 | GMTACGGTCACACTG | chr3L:17041574-17043574 | 1918 | 1932 | + | 3.3571 | 7.12E-05 | 0.795 | GCTGCTCTCACGCTG |

| | | | | | | | | | |
|---|---|---|---|---|---|---|---|---|---|
| 1 | GMTACGGTCACACTG | chr2L:13792521-13794521 | 1918 | 1932 | + | 3.3571 | 7.12E-05 | 0.795 | GCTACGCTCACCCAA |
| 1 | GMTACGGTCACACTG | chrX:13666811-13668811 | 1919 | 1933 | + | 3.3571 | 7.12E-05 | 0.795 | ACTGCGGCCACAGTG |
| 1 | GMTACGGTCACACTG | chrX:13188164-13190164 | 1920 | 1934 | + | 3.3571 | 7.12E-05 | 0.795 | GATTCGGCGACGCTG |
| 1 | GMTACGGTCACACTG | chr3R:18662738-18664738 | 1922 | 1936 | + | 3.3571 | 7.12E-05 | 0.795 | GCCATGCACACACTG |
| 1 | GMTACGGTCACACTG | chr3L:11163971-11165971 | 1928 | 1942 | + | 3.3571 | 7.12E-05 | 0.795 | GCTTCGATCATTCTG |
| 1 | GMTACGGTCACACTG | chr3L:7338266-7340266 | 1930 | 1944 | + | 3.3571 | 7.12E-05 | 0.795 | GCTTCGTTCACATGG |
| 1 | GMTACGGTCACACTG | chr3R:20407265-20409265 | 1935 | 1949 | + | 3.3571 | 7.12E-05 | 0.795 | GCTGCGGTGGCAGTG |
| 1 | GMTACGGTCACACTG | chr2R:16013787-16015787 | 1935 | 1949 | + | 3.3571 | 7.12E-05 | 0.795 | GAGACGGTTCCGCTG |
| 1 | GMTACGGTCACACTG | chr2R:12841558-12843558 | 1938 | 1952 | + | 3.3571 | 7.12E-05 | 0.795 | GCTAGGGGCAATCTG |
| 1 | GMTACGGTCACACTG | chr2R:5069210-5071210 | 1940 | 1954 | + | 3.3571 | 7.12E-05 | 0.795 | CCTTTGGACACACTG |
| 1 | GMTACGGTCACACTG | chr2L:6487756-6489756 | 1942 | 1956 | + | 3.3571 | 7.12E-05 | 0.795 | GCTACGCTGACTCAG |
| 1 | GMTACGGTCACACTG | chrX:8480837-8482837 | 1946 | 1960 | + | 3.3571 | 7.12E-05 | 0.795 | GACACGGACACAGCG |
| 1 | GMTACGGTCACACTG | chr3R:22379128-22381128 | 1948 | 1962 | + | 3.3571 | 7.12E-05 | 0.795 | CAAACGGACGCACTG |
| 1 | GMTACGGTCACACTG | chr3R:10760310-10762310 | 1948 | 1962 | + | 3.3571 | 7.12E-05 | 0.795 | GCTACTGCGCCACTG |
| 1 | GMTACGGTCACACTG | chr3R:8219146-8221146 | 1949 | 1963 | + | 3.3571 | 7.12E-05 | 0.795 | GATACGGCCGCCCTT |
| 1 | GMTACGGTCACACTG | chr2R:7405837-7407837 | 1952 | 1966 | + | 3.3571 | 7.12E-05 | 0.795 | AATGCGGCCGCACTG |
| 1 | GMTACGGTCACACTG | chr2R:15539887-15541887 | 1953 | 1967 | + | 3.3571 | 7.12E-05 | 0.795 | GCTGGTGTCACTCTG |
| 1 | GMTACGGTCACACTG | chr3L:7517997-7519997 | 1954 | 1968 | + | 3.3571 | 7.12E-05 | 0.795 | GCTACGGATGCAGTG |
| 1 | GMTACGGTCACACTG | chr2L:1986758-1988758 | 1955 | 1969 | + | 3.3571 | 7.12E-05 | 0.795 | TACACGGCCACCCTG |
| 1 | GMTACGGTCACACTG | chr2R:21010970-21012970 | 1957 | 1971 | + | 3.3571 | 7.12E-05 | 0.795 | GATACGGTCCCTTGG |
| 1 | GMTACGGTCACACTG | chr3L:5872311-5874311 | 1959 | 1973 | + | 3.3571 | 7.12E-05 | 0.795 | GCATCGATCACACTC |
| 1 | GMTACGGTCACACTG | chrX:8480837-8482837 | 1961 | 1975 | + | 3.3571 | 7.12E-05 | 0.795 | GACACGGACACGGTG |
| 1 | GMTACGGTCACACTG | chr3R:19389270-19391270 | 1963 | 1977 | + | 3.3571 | 7.12E-05 | 0.795 | GCTAGGGCCATGCTG |
| 1 | GMTACGGTCACACTG | chrX:16539852-16541852 | 1967 | 1981 | + | 3.3571 | 7.12E-05 | 0.795 | GAAGCGGCCACACTT |
| 1 | GMTACGGTCACACTG | chr2R:20262684-20264684 | 1967 | 1981 | + | 3.3571 | 7.12E-05 | 0.795 | GATCCGCTCCCACCG |
| 1 | GMTACGGTCACACTG | chrX:8922960-8924960 | 1969 | 1983 | + | 3.3571 | 7.12E-05 | 0.795 | GATACGGCCACTCGC |
| 1 | GMTACGGTCACACTG | chr2L:5201383-5203383 | 1974 | 1988 | + | 3.3571 | 7.12E-05 | 0.795 | GCTACGCACACTTTG |
| 1 | GMTACGGTCACACTG | chrX:2245231-2247231 | 1974 | 1988 | + | 3.3571 | 7.12E-05 | 0.795 | GATGCGGACACAGAG |
| 1 | GMTACGGTCACACTG | chr3L:9835805-9837805 | 1974 | 1988 | + | 3.3571 | 7.12E-05 | 0.795 | GATGCGGTCTAGCTG |
| 1 | GMTACGGTCACACTG | chrX:12900114-12902114 | 1975 | 1989 | + | 3.3571 | 7.12E-05 | 0.795 | GCCACTGTCACTGTG |
| 1 | GMTACGGTCACACTG | chr2R:20993900-20995900 | 1979 | 1993 | + | 3.3571 | 7.12E-05 | 0.795 | GAGATGGCCACGCTG |
| 1 | GMTACGGTCACACTG | chr3R:8340100-8342100 | 1979 | 1993 | + | 3.3571 | 7.12E-05 | 0.795 | GCGACGGGCACATTC |
| 1 | GMTACGGTCACACTG | chrX:22472698-22474698 | 1980 | 1994 | + | 3.3571 | 7.12E-05 | 0.795 | ACTACGGCCATTCTG |

| | | | | | | | | | |
|---|---|---|---|---|---|---|---|---|---|
| 1 | GMTACGGTCACACTG | chr3R:22379128-22381128 | 1984 | 1998 | + | 3.3571 | 7.12E-05 | 0.795 | GAATCGCTCGCACTG |
| 1 | GMTACGGTCACACTG | chr3R:12890956-12892956 | 2 | 16 | - | 3.3571 | 7.12E-05 | 0.795 | GCTGCATTCCCACTG |
| 1 | GMTACGGTCACACTG | chr2R:21706861-21708861 | 4 | 18 | - | 3.3571 | 7.12E-05 | 0.795 | GCTATGGCTACACAG |
| 1 | GMTACGGTCACACTG | chr3R:8139295-8141295 | 11 | 25 | - | 3.3571 | 7.12E-05 | 0.795 | ACAATGGCCACACTG |
| 1 | GMTACGGTCACACTG | chr3R:7284962-7286962 | 11 | 25 | - | 3.3571 | 7.12E-05 | 0.795 | GCCACTGCCACATTG |
| 1 | GMTACGGTCACACTG | chr2L:15128701-15130701 | 12 | 26 | - | 3.3571 | 7.12E-05 | 0.795 | GCTGCCGTCACGGTG |
| 1 | GMTACGGTCACACTG | chr2L:16731550-16733550 | 14 | 28 | - | 3.3571 | 7.12E-05 | 0.795 | GCTACGATCCCAACG |
| 1 | GMTACGGTCACACTG | chr2R:13445309-13447309 | 17 | 31 | - | 3.3571 | 7.12E-05 | 0.795 | TATTCGGCCGCACTG |
| 1 | GMTACGGTCACACTG | chr2R:24333307-24335307 | 17 | 31 | - | 3.3571 | 7.12E-05 | 0.795 | GACACGGAAACGCTG |
| 1 | GMTACGGTCACACTG | chr3L:18871553-18873553 | 24 | 38 | - | 3.3571 | 7.12E-05 | 0.795 | GCTTCGCTGACCCTG |
| 1 | GMTACGGTCACACTG | chr3L:8603149-8605149 | 26 | 40 | - | 3.3571 | 7.12E-05 | 0.795 | GCTCCTTTCTCACTG |
| 1 | GMTACGGTCACACTG | chr3L:13005817-13007817 | 26 | 40 | - | 3.3571 | 7.12E-05 | 0.795 | GATGCGGGCCCAGTG |
| 1 | GMTACGGTCACACTG | chr3L:1776439-1778439 | 27 | 41 | - | 3.3571 | 7.12E-05 | 0.795 | TCGACGGGCAGACTG |
| 1 | GMTACGGTCACACTG | chr3R:9988120-9990120 | 28 | 42 | - | 3.3571 | 7.12E-05 | 0.795 | GAGGTGGCCACACTG |
| 1 | GMTACGGTCACACTG | chr2L:19044355-19046355 | 29 | 43 | - | 3.3571 | 7.12E-05 | 0.795 | CCGAGGGTCACACCG |
| 1 | GMTACGGTCACACTG | chr3L:5598692-5600692 | 29 | 43 | - | 3.3571 | 7.12E-05 | 0.795 | GCTTTGGTCGAACTG |
| 1 | GMTACGGTCACACTG | chrX:1574132-1576132 | 29 | 43 | - | 3.3571 | 7.12E-05 | 0.795 | CACACGGACACACGG |
| 1 | GMTACGGTCACACTG | chr2L:6556800-6558800 | 32 | 46 | - | 3.3571 | 7.12E-05 | 0.795 | TCAACGGCGACACTG |
| 1 | GMTACGGTCACACTG | chr3R:8286169-8288169 | 32 | 46 | - | 3.3571 | 7.12E-05 | 0.795 | GAAAAGGACCCACTG |
| 1 | GMTACGGTCACACTG | chr2L:4256295-4258295 | 34 | 48 | - | 3.3571 | 7.12E-05 | 0.795 | GCAACTGCAACACTG |
| 1 | GMTACGGTCACACTG | chrX:2702852-2704852 | 36 | 50 | - | 3.3571 | 7.12E-05 | 0.795 | GCTTTGGTGACTCTG |
| 1 | GMTACGGTCACACTG | chrX:620215-622215 | 36 | 50 | - | 3.3571 | 7.12E-05 | 0.795 | GCGACAGCCACACTA |
| 1 | GMTACGGTCACACTG | chr3L:19062165-19064165 | 36 | 50 | - | 3.3571 | 7.12E-05 | 0.795 | GCTGCGGGAAAACTG |
| 1 | GMTACGGTCACACTG | chrX:9271960-9273960 | 36 | 50 | - | 3.3571 | 7.12E-05 | 0.795 | GCTACGGACCCAATT |
| 1 | GMTACGGTCACACTG | chrX:1574132-1576132 | 37 | 51 | - | 3.3571 | 7.12E-05 | 0.795 | CAAACGGACACACGG |
| 1 | GMTACGGTCACACTG | chr3L:4039529-4041529 | 37 | 51 | - | 3.3571 | 7.12E-05 | 0.795 | GCCAAGGTCAGCCTG |
| 1 | GMTACGGTCACACTG | chr3L:19423183-19425183 | 39 | 53 | - | 3.3571 | 7.12E-05 | 0.795 | ACTCCGGTCTGACTG |
| 1 | GMTACGGTCACACTG | chr3R:23957391-23959391 | 41 | 55 | - | 3.3571 | 7.12E-05 | 0.795 | GCTAAGGCGGCACTG |
| 1 | GMTACGGTCACACTG | chr2L:4226699-4228699 | 44 | 58 | - | 3.3571 | 7.12E-05 | 0.795 | GCAATGGTAACGCTG |
| 1 | GMTACGGTCACACTG | chr3L:15234619-15236619 | 44 | 58 | - | 3.3571 | 7.12E-05 | 0.795 | GATTTGGGCCCACTG |
| 1 | GMTACGGTCACACTG | chr2L:20647282-20649282 | 45 | 59 | - | 3.3571 | 7.12E-05 | 0.795 | GCTACGGACACATAA |
| 1 | GMTACGGTCACACTG | chr3L:19496723-19498723 | 46 | 60 | - | 3.3571 | 7.12E-05 | 0.795 | GATTCGGTGACTCGG |
| 1 | GMTACGGTCACACTG | chr2R:24782555-24784555 | 47 | 61 | - | 3.3571 | 7.12E-05 | 0.795 | GCTCCGATCAGACGG |

| | | | | | | | | | |
|---|---|---|---|---|---|---|---|---|---|
| 1 | GMTACGGTCACACTG | chr3L:9718144-9720144 | 48 | 62 | - | 3.3571 | 7.12E-05 | 0.795 | GCGACCGTAACCCTG |
| 1 | GMTACGGTCACACTG | chrX:6286990-6288990 | 52 | 66 | - | 3.3571 | 7.12E-05 | 0.795 | GCTACGGTGACCTGG |
| 1 | GMTACGGTCACACTG | chr3L:1776411-1778411 | 55 | 69 | - | 3.3571 | 7.12E-05 | 0.795 | TCGACGGGCAGACTG |
| 1 | GMTACGGTCACACTG | chr3R:22815178-22817178 | 57 | 71 | - | 3.3571 | 7.12E-05 | 0.795 | GCAACAGCCACACTT |
| 1 | GMTACGGTCACACTG | chr2L:13828837-13830837 | 60 | 74 | - | 3.3571 | 7.12E-05 | 0.795 | TCTACGGTCACTTCG |
| 1 | GMTACGGTCACACTG | chrX:7932742-7934742 | 60 | 74 | - | 3.3571 | 7.12E-05 | 0.795 | GCTACTGCCACACGA |
| 1 | GMTACGGTCACACTG | chr2R:21535795-21537795 | 63 | 77 | - | 3.3571 | 7.12E-05 | 0.795 | GCTATGGCAACGCTG |
| 1 | GMTACGGTCACACTG | chrX:4686800-4688800 | 66 | 80 | - | 3.3571 | 7.12E-05 | 0.795 | ACAACGGTCACACGC |
| 1 | GMTACGGTCACACTG | chrX:1030496-1032496 | 69 | 83 | - | 3.3571 | 7.12E-05 | 0.795 | ACACTGGTCACACTG |
| 1 | GMTACGGTCACACTG | chrX:9242467-9244467 | 84 | 98 | - | 3.3571 | 7.12E-05 | 0.795 | GCTGCCGCCACACTT |
| 1 | GMTACGGTCACACTG | chr3L:4039482-4041482 | 84 | 98 | - | 3.3571 | 7.12E-05 | 0.795 | GCCAAGGTCAGCCTG |
| 1 | GMTACGGTCACACTG | chr2R:8262278-8264278 | 85 | 99 | - | 3.3571 | 7.12E-05 | 0.795 | GCTCGGGTGGCACTG |
| 1 | GMTACGGTCACACTG | chr2L:249823-251823 | 85 | 99 | - | 3.3571 | 7.12E-05 | 0.795 | GCTACAGTTGCTCTG |
| 1 | GMTACGGTCACACTG | chrX:11052556-11054556 | 87 | 101 | - | 3.3571 | 7.12E-05 | 0.795 | GCGACGGTCCCAATA |
| 1 | GMTACGGTCACACTG | chr3L:15982135-15984135 | 89 | 103 | - | 3.3571 | 7.12E-05 | 0.795 | GCTCCAGTCACACGT |
| 1 | GMTACGGTCACACTG | chr3L:3393088-3395088 | 92 | 106 | - | 3.3571 | 7.12E-05 | 0.795 | GAGGCGGGCACATTG |
| 1 | GMTACGGTCACACTG | chr2R:7157241-7159241 | 94 | 108 | - | 3.3571 | 7.12E-05 | 0.795 | GACCTGGTCACACCG |
| 1 | GMTACGGTCACACTG | chrX:13781459-13783459 | 96 | 110 | - | 3.3571 | 7.12E-05 | 0.795 | GCAATGGCCAAACTG |
| 1 | GMTACGGTCACACTG | chr2R:14238932-14240932 | 96 | 110 | - | 3.3571 | 7.12E-05 | 0.795 | GATCCGGACACTTTG |
| 1 | GMTACGGTCACACTG | chrX:17849460-17851460 | 97 | 111 | - | 3.3571 | 7.12E-05 | 0.795 | GCTTTCGTCACCCTG |
| 1 | GMTACGGTCACACTG | chr2R:12335733-12337733 | 97 | 111 | - | 3.3571 | 7.12E-05 | 0.795 | GCCACGCTCACCTTG |
| 1 | GMTACGGTCACACTG | chr3R:19402455-19404455 | 100 | 114 | - | 3.3571 | 7.12E-05 | 0.795 | GCGACGATCCCACTA |
| 1 | GMTACGGTCACACTG | chr3L:598399-600399 | 102 | 116 | - | 3.3571 | 7.12E-05 | 0.795 | CCTGCGGTCCCACTT |
| 1 | GMTACGGTCACACTG | chr2L:7725298-7727298 | 102 | 116 | - | 3.3571 | 7.12E-05 | 0.795 | GCTGCGGTTCCACTT |
| 1 | GMTACGGTCACACTG | chr2R:24782500-24784500 | 102 | 116 | - | 3.3571 | 7.12E-05 | 0.795 | GCTCCGATCAGACGG |
| 1 | GMTACGGTCACACTG | chr2R:17475272-17477272 | 105 | 119 | - | 3.3571 | 7.12E-05 | 0.795 | GCTGCGATCTCATTG |
| 1 | GMTACGGTCACACTG | chr3L:13512112-13514112 | 106 | 120 | - | 3.3571 | 7.12E-05 | 0.795 | GATCCCGTCACGCGG |
| 1 | GMTACGGTCACACTG | chr2L:20638015-20640015 | 108 | 122 | - | 3.3571 | 7.12E-05 | 0.795 | GATCCGGTCCCACAT |
| 1 | GMTACGGTCACACTG | chr3R:25220589-25222589 | 110 | 124 | - | 3.3571 | 7.12E-05 | 0.795 | TCTACGGGTACGCTG |
| 1 | GMTACGGTCACACTG | chr3L:14050206-14052206 | 111 | 125 | - | 3.3571 | 7.12E-05 | 0.795 | TCTCCGGTGACACGG |
| 1 | GMTACGGTCACACTG | chr3L:7910954-7912954 | 112 | 126 | - | 3.3571 | 7.12E-05 | 0.795 | TCTTCGGCGACACTG |
| 1 | GMTACGGTCACACTG | chr2L:12095525-12097525 | 112 | 126 | - | 3.3571 | 7.12E-05 | 0.795 | CCAATGGTCCCACTG |
| 1 | GMTACGGTCACACTG | chr3L:1501843-1503843 | 113 | 127 | - | 3.3571 | 7.12E-05 | 0.795 | GCTAGGGTCTCTCTA |

| | | | | | | | | | |
|---|---|---|---|---|---|---|---|---|---|
| 1 | GMTACGGTCACACTG | chrX:2632952-2634952 | 113 | 127 | - | 3.3571 | 7.12E-05 | 0.795 | GATACGGACGAACGG |
| 1 | GMTACGGTCACACTG | chrX:21480124-21482124 | 114 | 128 | - | 3.3571 | 7.12E-05 | 0.795 | GAAAAGGACACTCTG |
| 1 | GMTACGGTCACACTG | chr3R:26865807-26867807 | 115 | 129 | - | 3.3571 | 7.12E-05 | 0.795 | GCTACGTTCATGCCG |
| 1 | GMTACGGTCACACTG | chr2L:8682877-8684877 | 117 | 131 | - | 3.3571 | 7.12E-05 | 0.795 | ACGACGGTGACTCTG |
| 1 | GMTACGGTCACACTG | chr2L:13398831-13400831 | 117 | 131 | - | 3.3571 | 7.12E-05 | 0.795 | GCAACAGGCACACTA |
| 1 | GMTACGGTCACACTG | chrX:5613467-5615467 | 119 | 133 | - | 3.3571 | 7.12E-05 | 0.795 | GCTCCGGTTTCATTG |
| 1 | GMTACGGTCACACTG | chrX:18847566-18849566 | 121 | 135 | - | 3.3571 | 7.12E-05 | 0.795 | GCTGGGCTCACCCTG |
| 1 | GMTACGGTCACACTG | chr2L:12095511-12097511 | 126 | 140 | - | 3.3571 | 7.12E-05 | 0.795 | CCAATGGTCCCACTG |
| 1 | GMTACGGTCACACTG | chr2L:22923795-22925795 | 129 | 143 | - | 3.3571 | 7.12E-05 | 0.795 | GATCCGCACACACGG |
| 1 | GMTACGGTCACACTG | chr2L:9919160-9921160 | 129 | 143 | - | 3.3571 | 7.12E-05 | 0.795 | GCTGTGGGAACACTG |
| 1 | GMTACGGTCACACTG | chrX:19911655-19913655 | 129 | 143 | - | 3.3571 | 7.12E-05 | 0.795 | GCTACGGTGGCACCC |
| 1 | GMTACGGTCACACTG | chrX:19940530-19942530 | 129 | 143 | - | 3.3571 | 7.12E-05 | 0.795 | GCTACGGTGGCACCC |
| 1 | GMTACGGTCACACTG | chrX:19930714-19932714 | 129 | 143 | - | 3.3571 | 7.12E-05 | 0.795 | GCTACGGTGGCACCC |
| 1 | GMTACGGTCACACTG | chrX:19916819-19918819 | 129 | 143 | - | 3.3571 | 7.12E-05 | 0.795 | GCTACGGTGGCACCC |
| 1 | GMTACGGTCACACTG | chr2L:8238878-8240878 | 129 | 143 | - | 3.3571 | 7.12E-05 | 0.795 | GCTGCGATCAAACGG |
| 1 | GMTACGGTCACACTG | chr2L:6561325-6563325 | 129 | 143 | - | 3.3571 | 7.12E-05 | 0.795 | ACCACGGCCACATTG |
| 1 | GMTACGGTCACACTG | chr2L:2009717-2011717 | 131 | 145 | - | 3.3571 | 7.12E-05 | 0.795 | GCAACTGTCAATCTG |
| 1 | GMTACGGTCACACTG | chrX:16080056-16082056 | 132 | 146 | - | 3.3571 | 7.12E-05 | 0.795 | GCTTTGGTCTCAATG |
| 1 | GMTACGGTCACACTG | chr3R:9257501-9259501 | 133 | 147 | - | 3.3571 | 7.12E-05 | 0.795 | ACTACGGCCCAACTG |
| 1 | GMTACGGTCACACTG | chrX:12969711-12971711 | 136 | 150 | - | 3.3571 | 7.12E-05 | 0.795 | GCTACGGTGGTGCTG |
| 1 | GMTACGGTCACACTG | chrX:9070408-9072408 | 136 | 150 | - | 3.3571 | 7.12E-05 | 0.795 | GATACGGAGCCGCTG |
| 1 | GMTACGGTCACACTG | chr3L:21192363-21194363 | 137 | 151 | - | 3.3571 | 7.12E-05 | 0.795 | GCTCGGGACACATTG |
| 1 | GMTACGGTCACACTG | chr3R:24864350-24866350 | 137 | 151 | - | 3.3571 | 7.12E-05 | 0.795 | GCTACGGCTACGGTG |
| 1 | GMTACGGTCACACTG | chr3L:13062046-13064046 | 139 | 153 | - | 3.3571 | 7.12E-05 | 0.795 | GCTCCGGTAACACCC |
| 1 | GMTACGGTCACACTG | chrX:634890-636890 | 139 | 153 | - | 3.3571 | 7.12E-05 | 0.795 | GCTGGGATCACGCTG |
| 1 | GMTACGGTCACACTG | chr2R:1344119-1346119 | 141 | 155 | - | 3.3571 | 7.12E-05 | 0.795 | GACACGGTAGCGCTG |
| 1 | GMTACGGTCACACTG | chr3R:4540356-4542356 | 142 | 156 | - | 3.3571 | 7.12E-05 | 0.795 | GCTACGTTCAGTCCG |
| 1 | GMTACGGTCACACTG | chr2L:3509102-3511102 | 144 | 158 | - | 3.3571 | 7.12E-05 | 0.795 | GCTACGGTTGCGCTA |
| 1 | GMTACGGTCACACTG | chr2R:21291181-21293181 | 146 | 160 | - | 3.3571 | 7.12E-05 | 0.795 | ACTACGATCGCACAG |
| 1 | GMTACGGTCACACTG | chrX:8713936-8715936 | 147 | 161 | - | 3.3571 | 7.12E-05 | 0.795 | GCGACAGACACACTT |
| 1 | GMTACGGTCACACTG | chr3R:15228408-15230408 | 148 | 162 | - | 3.3571 | 7.12E-05 | 0.795 | GCTACGCCCACTTTG |
| 1 | GMTACGGTCACACTG | chr2R:17018522-17020522 | 150 | 164 | - | 3.3571 | 7.12E-05 | 0.795 | GCCACGGCCACAATT |
| 1 | GMTACGGTCACACTG | chrX:2551206-2553206 | 151 | 165 | - | 3.3571 | 7.12E-05 | 0.795 | CCTACGGTCACTGCG |

| | | | | | | | | | |
|---|---|---|---|---|---|---|---|---|---|
| 1 | GMTACGGTCACACTG | chr2L:13211116-13213116 | 152 | 166 | - | 3.3571 | 7.12E-05 | 0.795 | GCTATCGGCGCACTG |
| 1 | GMTACGGTCACACTG | chr2R:10885810-10887810 | 153 | 167 | - | 3.3571 | 7.12E-05 | 0.795 | GCGAAGGTGACACCG |
| 1 | GMTACGGTCACACTG | chr2L:5519135-5521135 | 154 | 168 | - | 3.3571 | 7.12E-05 | 0.795 | GCTACGAGCGCAATG |
| 1 | GMTACGGTCACACTG | chr2L:13828741-13830741 | 156 | 170 | - | 3.3571 | 7.12E-05 | 0.795 | TCTACGGTCACTTCG |
| 1 | GMTACGGTCACACTG | chr2R:10312891-10314891 | 158 | 172 | - | 3.3571 | 7.12E-05 | 0.795 | TCAACAGTCACACCG |
| 1 | GMTACGGTCACACTG | chr2R:15539887-15541887 | 158 | 172 | - | 3.3571 | 7.12E-05 | 0.795 | GCTACCGGCACCTTG |
| 1 | GMTACGGTCACACTG | chr2R:16830432-16832432 | 159 | 173 | - | 3.3571 | 7.12E-05 | 0.795 | GACACGAGCACGCTG |
| 1 | GMTACGGTCACACTG | chrX:2221567-2223567 | 159 | 173 | - | 3.3571 | 7.12E-05 | 0.795 | GCTCCGATCAACCTG |
| 1 | GMTACGGTCACACTG | chrX:8713924-8715924 | 159 | 173 | - | 3.3571 | 7.12E-05 | 0.795 | GCGACAGACACACTT |
| 1 | GMTACGGTCACACTG | chr2R:20261883-20263883 | 164 | 178 | - | 3.3571 | 7.12E-05 | 0.795 | GAGCCGGTGACGCTG |
| 1 | GMTACGGTCACACTG | chr2L:4849641-4851641 | 165 | 179 | - | 3.3571 | 7.12E-05 | 0.795 | GATGCGGTCACCAGG |
| 1 | GMTACGGTCACACTG | chrX:4317801-4319801 | 165 | 179 | - | 3.3571 | 7.12E-05 | 0.795 | GACACCGTCCCACAG |
| 1 | GMTACGGTCACACTG | chr2L:6561289-6563289 | 165 | 179 | - | 3.3571 | 7.12E-05 | 0.795 | ACCACGGCCACATTG |
| 1 | GMTACGGTCACACTG | chr3R:25998851-26000851 | 172 | 186 | - | 3.3571 | 7.12E-05 | 0.795 | GCTGTGGACACACTT |
| 1 | GMTACGGTCACACTG | chr3R:9554408-9556408 | 172 | 186 | - | 3.3571 | 7.12E-05 | 0.795 | GCTATGGTCCCTCTA |
| 1 | GMTACGGTCACACTG | chr2R:18864218-18866218 | 173 | 187 | - | 3.3571 | 7.12E-05 | 0.795 | GAGACAGACACACGG |
| 1 | GMTACGGTCACACTG | chr3R:4333071-4335071 | 175 | 189 | - | 3.3571 | 7.12E-05 | 0.795 | GCTACGGTCCGCCTC |
| 1 | GMTACGGTCACACTG | chr3L:3147358-3149358 | 178 | 192 | - | 3.3571 | 7.12E-05 | 0.795 | GAGAAGGCCACGCTG |
| 1 | GMTACGGTCACACTG | chr2R:5965856-5967856 | 179 | 193 | - | 3.3571 | 7.12E-05 | 0.795 | TCTACGGACAGACGG |
| 1 | GMTACGGTCACACTG | chrX:18692042-18694042 | 179 | 193 | - | 3.3571 | 7.12E-05 | 0.795 | GCTACAGCTACACCG |
| 1 | GMTACGGTCACACTG | chr3L:11472291-11474291 | 181 | 195 | - | 3.3571 | 7.12E-05 | 0.795 | GCTCCAGTCGCACTC |
| 1 | GMTACGGTCACACTG | chr3L:25842821-25844821 | 182 | 196 | - | 3.3571 | 7.12E-05 | 0.795 | GCCACGCCCACACTT |
| 1 | GMTACGGTCACACTG | chrX:15785047-15787047 | 183 | 197 | - | 3.3571 | 7.12E-05 | 0.795 | GCAACGCTGTCACTG |
| 1 | GMTACGGTCACACTG | chr3R:12421768-12423768 | 191 | 205 | - | 3.3571 | 7.12E-05 | 0.795 | TCTACGGTACCACGG |
| 1 | GMTACGGTCACACTG | chr2R:7524716-7526716 | 192 | 206 | - | 3.3571 | 7.12E-05 | 0.795 | GCCACGATGACACAG |
| 1 | GMTACGGTCACACTG | chr3L:5576348-5578348 | 194 | 208 | - | 3.3571 | 7.12E-05 | 0.795 | GCCAGGGTCACCCTT |
| 1 | GMTACGGTCACACTG | chrX:15643222-15645222 | 195 | 209 | - | 3.3571 | 7.12E-05 | 0.795 | GCTTTCGTCCCACTG |
| 1 | GMTACGGTCACACTG | chr3L:11464864-11466864 | 198 | 212 | - | 3.3571 | 7.12E-05 | 0.795 | GCCATGGGCATACTG |
| 1 | GMTACGGTCACACTG | chr4:866824-868824 | 200 | 214 | - | 3.3571 | 7.12E-05 | 0.795 | AAGACGGCCACGCTG |
| 1 | GMTACGGTCACACTG | chrX:4100031-4102031 | 203 | 217 | - | 3.3571 | 7.12E-05 | 0.795 | GATCCGGCCATCCTG |
| 1 | GMTACGGTCACACTG | chr2R:7154424-7156424 | 205 | 219 | - | 3.3571 | 7.12E-05 | 0.795 | GCTACAGGTACGCTG |
| 1 | GMTACGGTCACACTG | chr2R:13484391-13486391 | 205 | 219 | - | 3.3571 | 7.12E-05 | 0.795 | GCAGAGGTAACACTG |
| 1 | GMTACGGTCACACTG | chrX:495446-497446 | 209 | 223 | - | 3.3571 | 7.12E-05 | 0.795 | GCTGACGTCACGCTG |

| | | | | | | | | | |
|---|---|---|---|---|---|---|---|---|---|
| 1 | GMTACGGTCACACTG | chr3L:27134184-27136184 | 212 | 226 | - | 3.3571 | 7.12E-05 | 0.795 | GCCGCGTTCACAATG |
| 1 | GMTACGGTCACACTG | chr2R:10311023-10313023 | 214 | 228 | - | 3.3571 | 7.12E-05 | 0.795 | GCTTCCGTCAAACAG |
| 1 | GMTACGGTCACACTG | chrX:17849342-17851342 | 215 | 229 | - | 3.3571 | 7.12E-05 | 0.795 | GCTTTCGTCACCCTG |
| 1 | GMTACGGTCACACTG | chr3R:29599862-29601862 | 218 | 232 | - | 3.3571 | 7.12E-05 | 0.795 | TCTATGGACACACCG |
| 1 | GMTACGGTCACACTG | chr2R:20642785-20644785 | 223 | 237 | - | 3.3571 | 7.12E-05 | 0.795 | GCTTTGGTCAGGCTG |
| 1 | GMTACGGTCACACTG | chr2R:20642785-20644785 | 223 | 237 | - | 3.3571 | 7.12E-05 | 0.795 | GCTTTGGTCAGGCTG |
| 1 | GMTACGGTCACACTG | chr2R:11134923-11136923 | 224 | 238 | - | 3.3571 | 7.12E-05 | 0.795 | GCTACGACCACAACG |
| 1 | GMTACGGTCACACTG | chr3R:16787311-16789311 | 228 | 242 | - | 3.3571 | 7.12E-05 | 0.795 | CCTCCGGACAAACTG |
| 1 | GMTACGGTCACACTG | chr2R:9847061-9849061 | 232 | 246 | - | 3.3571 | 7.12E-05 | 0.795 | GCTACGTTCACGCCT |
| 1 | GMTACGGTCACACTG | chr3R:31763772-31765772 | 234 | 248 | - | 3.3571 | 7.12E-05 | 0.795 | GCTACGGTCCACCTT |
| 1 | GMTACGGTCACACTG | chr2R:21710613-21712613 | 235 | 249 | - | 3.3571 | 7.12E-05 | 0.795 | ACTGCGGTCACAGCG |
| 1 | GMTACGGTCACACTG | chr2L:19044148-19046148 | 236 | 250 | - | 3.3571 | 7.12E-05 | 0.795 | CCGAGGGTCACACCG |
| 1 | GMTACGGTCACACTG | chr2R:13553077-13555077 | 237 | 251 | - | 3.3571 | 7.12E-05 | 0.795 | TCTCCGATCGCACTG |
| 1 | GMTACGGTCACACTG | chrX:8146949-8148949 | 238 | 252 | - | 3.3571 | 7.12E-05 | 0.795 | GAGAAGGTCGCACCG |
| 1 | GMTACGGTCACACTG | chr2L:9792107-9794107 | 247 | 261 | - | 3.3571 | 7.12E-05 | 0.795 | GCTGCCGTCGAACTG |
| 1 | GMTACGGTCACACTG | chr2R:19441342-19443342 | 251 | 265 | - | 3.3571 | 7.12E-05 | 0.795 | ACGATGGACACACTG |
| 1 | GMTACGGTCACACTG | chr3L:16086525-16088525 | 252 | 266 | - | 3.3571 | 7.12E-05 | 0.795 | GCTTTGGACACAGTG |
| 1 | GMTACGGTCACACTG | chr3R:18410631-18412631 | 253 | 267 | - | 3.3571 | 7.12E-05 | 0.795 | GCTACGGCAACATGG |
| 1 | GMTACGGTCACACTG | chr3R:6677590-6679590 | 256 | 270 | - | 3.3571 | 7.12E-05 | 0.795 | GCTACCGCCGCAGTG |
| 1 | GMTACGGTCACACTG | chr3R:18273665-18275665 | 256 | 270 | - | 3.3571 | 7.12E-05 | 0.795 | GCCACCGTCATACGG |
| 1 | GMTACGGTCACACTG | chr2R:12144710-12146710 | 258 | 272 | - | 3.3571 | 7.12E-05 | 0.795 | GCCACGGACAGACTT |
| 1 | GMTACGGTCACACTG | chr3L:1567608-1569608 | 258 | 272 | - | 3.3571 | 7.12E-05 | 0.795 | GCTAGGATCACTCAG |
| 1 | GMTACGGTCACACTG | chr3R:31108191-31110191 | 259 | 273 | - | 3.3571 | 7.12E-05 | 0.795 | GCCACGGTCAGTGTG |
| 1 | GMTACGGTCACACTG | chr3R:20870036-20872036 | 261 | 275 | - | 3.3571 | 7.12E-05 | 0.795 | GACAAGGACACACAG |
| 1 | GMTACGGTCACACTG | chr3R:19094000-19096000 | 261 | 275 | - | 3.3571 | 7.12E-05 | 0.795 | ACAACGGTCACTCTA |
| 1 | GMTACGGTCACACTG | chr3R:26240291-26242291 | 265 | 279 | - | 3.3571 | 7.12E-05 | 0.795 | GCTATGGTGGCTCTG |
| 1 | GMTACGGTCACACTG | chr2R:21537015-21539015 | 270 | 284 | - | 3.3571 | 7.12E-05 | 0.795 | GATGGGGACGCACTG |
| 1 | GMTACGGTCACACTG | chr3L:12266528-12268528 | 272 | 286 | - | 3.3571 | 7.12E-05 | 0.795 | AATTCGGACACACCG |
| 1 | GMTACGGTCACACTG | chr2R:14258030-14260030 | 273 | 287 | - | 3.3571 | 7.12E-05 | 0.795 | GCTACGGTGACGTCG |
| 1 | GMTACGGTCACACTG | chr2R:17955570-17957570 | 273 | 287 | - | 3.3571 | 7.12E-05 | 0.795 | GCTGGGGTCACAACG |
| 1 | GMTACGGTCACACTG | chr2L:20647051-20649051 | 276 | 290 | - | 3.3571 | 7.12E-05 | 0.795 | GCTACGGACACATAA |
| 1 | GMTACGGTCACACTG | chr3L:2950618-2952618 | 278 | 292 | - | 3.3571 | 7.12E-05 | 0.795 | GCGAGGGTCCAACTG |
| 1 | GMTACGGTCACACTG | chr3R:8139023-8141023 | 283 | 297 | - | 3.3571 | 7.12E-05 | 0.795 | ACAATGGCCACACTG |

| | | | | | | | | | |
|---|---|---|---|---|---|---|---|---|---|
| 1 | GMTACGGTCACACTG | chrX:16445993-16447993 | 283 | 297 | - | 3.3571 | 7.12E-05 | 0.795 | GCTACGGTTACGAGG |
| 1 | GMTACGGTCACACTG | chr2L:2035041-2037041 | 288 | 302 | - | 3.3571 | 7.12E-05 | 0.795 | GACGCGGACATACTG |
| 1 | GMTACGGTCACACTG | chr2R:8613155-8615155 | 291 | 305 | - | 3.3571 | 7.12E-05 | 0.795 | GCGAGGGGCAAACTG |
| 1 | GMTACGGTCACACTG | chr2L:1427474-1429474 | 293 | 307 | - | 3.3571 | 7.12E-05 | 0.795 | GCTGCGATCACAGGG |
| 1 | GMTACGGTCACACTG | chrX:17910462-17912462 | 293 | 307 | - | 3.3571 | 7.12E-05 | 0.795 | GCGACGATTCCACTG |
| 1 | GMTACGGTCACACTG | chrX:6677424-6679424 | 293 | 307 | - | 3.3571 | 7.12E-05 | 0.795 | GCTACGGTCTGGGTG |
| 1 | GMTACGGTCACACTG | chr3R:27017287-27019287 | 294 | 308 | - | 3.3571 | 7.12E-05 | 0.795 | GCTACGATAACCCGG |
| 1 | GMTACGGTCACACTG | chr3R:24840935-24842935 | 296 | 310 | - | 3.3571 | 7.12E-05 | 0.795 | GCTTCAGTCTCACTT |
| 1 | GMTACGGTCACACTG | chrX:6236315-6238315 | 299 | 313 | - | 3.3571 | 7.12E-05 | 0.795 | GCCAAGATCACGCTG |
| 1 | GMTACGGTCACACTG | chr2R:20605067-20607067 | 301 | 315 | - | 3.3571 | 7.12E-05 | 0.795 | GCTAGTGCCACCCTG |
| 1 | GMTACGGTCACACTG | chr4:1184768-1186768 | 302 | 316 | - | 3.3571 | 7.12E-05 | 0.795 | GCTACGCCCACTCTT |
| 1 | GMTACGGTCACACTG | chr3L:65430-67430 | 304 | 318 | - | 3.3571 | 7.12E-05 | 0.795 | GCCACGTTCGCACTT |
| 1 | GMTACGGTCACACTG | chr2R:9586193-9588193 | 305 | 319 | - | 3.3571 | 7.12E-05 | 0.795 | GATACGGGCTCAAAG |
| 1 | GMTACGGTCACACTG | chr3L:3939519-3941519 | 305 | 319 | - | 3.3571 | 7.12E-05 | 0.795 | GCTACCGGCACATAG |
| 1 | GMTACGGTCACACTG | chrX:20375522-20377522 | 306 | 320 | - | 3.3571 | 7.12E-05 | 0.795 | CCTACGCACTCACTG |
| 1 | GMTACGGTCACACTG | chr2R:15498632-15500632 | 308 | 322 | - | 3.3571 | 7.12E-05 | 0.795 | GCAAAGGCGACACTG |
| 1 | GMTACGGTCACACTG | chr2L:19044355-19046355 | 309 | 323 | - | 3.3571 | 7.12E-05 | 0.795 | GCTGCTTTCACTCTG |
| 1 | GMTACGGTCACACTG | chr3L:17565162-17567162 | 309 | 323 | - | 3.3571 | 7.12E-05 | 0.795 | GCAACGGCAACAATG |
| 1 | GMTACGGTCACACTG | chr3R:19889689-19891689 | 309 | 323 | - | 3.3571 | 7.12E-05 | 0.795 | GCGTCACTCACACTG |
| 1 | GMTACGGTCACACTG | chr3R:23722331-23724331 | 310 | 324 | - | 3.3571 | 7.12E-05 | 0.795 | GCAACAGACAAACTG |
| 1 | GMTACGGTCACACTG | chr2R:18398486-18400486 | 314 | 328 | - | 3.3571 | 7.12E-05 | 0.795 | ACTTCGGTAACACCG |
| 1 | GMTACGGTCACACTG | chr2L:318259-320259 | 317 | 331 | - | 3.3571 | 7.12E-05 | 0.795 | GCAACGGAGACAGTG |
| 1 | GMTACGGTCACACTG | chrX:7907286-7909286 | 317 | 331 | - | 3.3571 | 7.12E-05 | 0.795 | GAAAAGGACACTCTG |
| 1 | GMTACGGTCACACTG | chr2L:22106304-22108304 | 319 | 333 | - | 3.3571 | 7.12E-05 | 0.795 | GCGACGTTCTCACTC |
| 1 | GMTACGGTCACACTG | chr2R:16578470-16580470 | 319 | 333 | - | 3.3571 | 7.12E-05 | 0.795 | GATCTGGCCGCACTG |
| 1 | GMTACGGTCACACTG | chr3L:1834164-1836164 | 323 | 337 | - | 3.3571 | 7.12E-05 | 0.795 | GCTCCATTCGCACTG |
| 1 | GMTACGGTCACACTG | chr3L:21495393-21497393 | 323 | 337 | - | 3.3571 | 7.12E-05 | 0.795 | GCAAGGGGCACAGTG |
| 1 | GMTACGGTCACACTG | chr2L:20655426-20657426 | 324 | 338 | - | 3.3571 | 7.12E-05 | 0.795 | GATGCTGTCGCACGG |
| 1 | GMTACGGTCACACTG | chr2L:15892811-15894811 | 324 | 338 | - | 3.3571 | 7.12E-05 | 0.795 | CCTACGGTCTAACAG |
| 1 | GMTACGGTCACACTG | chr2R:14238703-14240703 | 325 | 339 | - | 3.3571 | 7.12E-05 | 0.795 | GATCCGGACACTTTG |
| 1 | GMTACGGTCACACTG | chr2L:6556502-6558502 | 330 | 344 | - | 3.3571 | 7.12E-05 | 0.795 | TCAACGGCGACACTG |
| 1 | GMTACGGTCACACTG | chr2R:8946344-8948344 | 333 | 347 | - | 3.3571 | 7.12E-05 | 0.795 | GCTGCGGTTCCAGTG |
| 1 | GMTACGGTCACACTG | chr2L:4457364-4459364 | 334 | 348 | - | 3.3571 | 7.12E-05 | 0.795 | GCTACGCGAACTCTG |

| | | | | | | | | | |
|---|---|---|---|---|---|---|---|---|---|
| 1 | GMTACGGTCACACTG | chrX:16312138-16314138 | 337 | 351 | - | 3.3571 | 7.12E-05 | 0.795 | GCTCCCGTCATTCTG |
| 1 | GMTACGGTCACACTG | chr2L:3318872-3320872 | 339 | 353 | - | 3.3571 | 7.12E-05 | 0.795 | GAGATGGACACTCTG |
| 1 | GMTACGGTCACACTG | chr3R:24787054-24789054 | 340 | 354 | - | 3.3571 | 7.12E-05 | 0.795 | ACTCTGGCCACACTG |
| 1 | GMTACGGTCACACTG | chr2L:20483702-20485702 | 342 | 356 | - | 3.3571 | 7.12E-05 | 0.795 | GATCGGGCCACACGG |
| 1 | GMTACGGTCACACTG | chr3R:27519866-27521866 | 343 | 357 | - | 3.3571 | 7.12E-05 | 0.795 | GCCGCAGTTACACTG |
| 1 | GMTACGGTCACACTG | chrX:13337318-13339318 | 345 | 359 | - | 3.3571 | 7.12E-05 | 0.795 | AACGCGGCCACACTG |
| 1 | GMTACGGTCACACTG | chrX:655114-657114 | 348 | 362 | - | 3.3571 | 7.12E-05 | 0.795 | GAGACGGACACACGC |
| 1 | GMTACGGTCACACTG | chr2L:13448497-13450497 | 349 | 363 | - | 3.3571 | 7.12E-05 | 0.795 | GCCAAGGTGGCACTG |
| 1 | GMTACGGTCACACTG | chr2R:12051944-12053944 | 351 | 365 | - | 3.3571 | 7.12E-05 | 0.795 | GCTAAGGACATCCTG |
| 1 | GMTACGGTCACACTG | chr3L:21267601-21269601 | 353 | 367 | - | 3.3571 | 7.12E-05 | 0.795 | GCTTTGGTCACTTTG |
| 1 | GMTACGGTCACACTG | chr3R:17702695-17704695 | 354 | 368 | - | 3.3571 | 7.12E-05 | 0.795 | GCGACAGTCGGACTG |
| 1 | GMTACGGTCACACTG | chr3L:21679050-21681050 | 354 | 368 | - | 3.3571 | 7.12E-05 | 0.795 | GCTACAGCTTCACTG |
| 1 | GMTACGGTCACACTG | chr3L:15981869-15983869 | 355 | 369 | - | 3.3571 | 7.12E-05 | 0.795 | GCTCCAGTCACACGT |
| 1 | GMTACGGTCACACTG | chr3R:19093905-19095905 | 356 | 370 | - | 3.3571 | 7.12E-05 | 0.795 | ACAACGGTCACTCTA |
| 1 | GMTACGGTCACACTG | chr3L:5498594-5500594 | 356 | 370 | - | 3.3571 | 7.12E-05 | 0.795 | GCTACGGTCTCTTTA |
| 1 | GMTACGGTCACACTG | chr3R:25339252-25341252 | 361 | 375 | - | 3.3571 | 7.12E-05 | 0.795 | CCTTCTGGCACACTG |
| 1 | GMTACGGTCACACTG | chr2L:11788919-11790919 | 361 | 375 | - | 3.3571 | 7.12E-05 | 0.795 | GAGACGGCGACGCTG |
| 1 | GMTACGGTCACACTG | chrX:18399065-18401065 | 362 | 376 | - | 3.3571 | 7.12E-05 | 0.795 | GATGCGGGCCCATTG |
| 1 | GMTACGGTCACACTG | chr2L:9885250-9887250 | 362 | 376 | - | 3.3571 | 7.12E-05 | 0.795 | GCTACGGTGCAACAG |
| 1 | GMTACGGTCACACTG | chr2L:10693103-10695103 | 363 | 377 | - | 3.3571 | 7.12E-05 | 0.795 | GATCCGGTAGCTCTG |
| 1 | GMTACGGTCACACTG | chr3R:29193994-29195994 | 366 | 380 | - | 3.3571 | 7.12E-05 | 0.795 | GCAACAGTAACACGG |
| 1 | GMTACGGTCACACTG | chr3L:1501590-1503590 | 366 | 380 | - | 3.3571 | 7.12E-05 | 0.795 | GCTAGGGTCTCTCTA |
| 1 | GMTACGGTCACACTG | chr2R:21289343-21291343 | 367 | 381 | - | 3.3571 | 7.12E-05 | 0.795 | CCTACAGCCCCACTG |
| 1 | GMTACGGTCACACTG | chr3R:19228463-19230463 | 368 | 382 | - | 3.3571 | 7.12E-05 | 0.795 | GATACGGTCTCGAAG |
| 1 | GMTACGGTCACACTG | chr2L:10412463-10414463 | 368 | 382 | - | 3.3571 | 7.12E-05 | 0.795 | GATCCGAACACCCTG |
| 1 | GMTACGGTCACACTG | chr3L:7438405-7440405 | 371 | 385 | - | 3.3571 | 7.12E-05 | 0.795 | GCGGTGGTCACAATG |
| 1 | GMTACGGTCACACTG | chr3L:7439733-7441733 | 371 | 385 | - | 3.3571 | 7.12E-05 | 0.795 | GCGGTGGTCACAATG |
| 1 | GMTACGGTCACACTG | chr2R:8230588-8232588 | 371 | 385 | - | 3.3571 | 7.12E-05 | 0.795 | GCGAAGATCGCACTG |
| 1 | GMTACGGTCACACTG | chr3R:11353348-11355348 | 373 | 387 | - | 3.3571 | 7.12E-05 | 0.795 | GCTACTCCCACACCG |
| 1 | GMTACGGTCACACTG | chr2L:2584430-2586430 | 375 | 389 | - | 3.3571 | 7.12E-05 | 0.795 | CCCACGAACACACTG |
| 1 | GMTACGGTCACACTG | chr2L:5580116-5582116 | 377 | 391 | - | 3.3571 | 7.12E-05 | 0.795 | GAGACGGTCTGCCTG |
| 1 | GMTACGGTCACACTG | chr3R:7284595-7286595 | 378 | 392 | - | 3.3571 | 7.12E-05 | 0.795 | GCCACTGCCACATTG |
| 1 | GMTACGGTCACACTG | chr3R:26005023-26007023 | 379 | 393 | - | 3.3571 | 7.12E-05 | 0.795 | GCTATGGTCACTCCA |

| | | | | | | | | | |
|---|---|---|---|---|---|---|---|---|---|
| 1 | GMTACGGTCACACTG | chr2R:21618079-21620079 | 379 | 393 | - | 3.3571 | 7.12E-05 | 0.795 | GCAACGGCAACAGTG |
| 1 | GMTACGGTCACACTG | chr3R:7085083-7087083 | 385 | 399 | - | 3.3571 | 7.12E-05 | 0.795 | GCTCCGGTAACAAAG |
| 1 | GMTACGGTCACACTG | chr3L:19061816-19063816 | 385 | 399 | - | 3.3571 | 7.12E-05 | 0.795 | GCTGCGGGAAAACTG |
| 1 | GMTACGGTCACACTG | chrX:13781170-13783170 | 385 | 399 | - | 3.3571 | 7.12E-05 | 0.795 | GCAATGGCCAAACTG |
| 1 | GMTACGGTCACACTG | chr2L:4849419-4851419 | 387 | 401 | - | 3.3571 | 7.12E-05 | 0.795 | GATGCGGTCACCAGG |
| 1 | GMTACGGTCACACTG | chr3L:1461801-1463801 | 389 | 403 | - | 3.3571 | 7.12E-05 | 0.795 | GACATGGACCCACTG |
| 1 | GMTACGGTCACACTG | chrX:6266797-6268797 | 392 | 406 | - | 3.3571 | 7.12E-05 | 0.795 | TCTCTGGGCACACTG |
| 1 | GMTACGGTCACACTG | chr2L:5518896-5520896 | 393 | 407 | - | 3.3571 | 7.12E-05 | 0.795 | GCTACGAGCGCAATG |
| 1 | GMTACGGTCACACTG | chr2L:12965590-12967590 | 396 | 410 | - | 3.3571 | 7.12E-05 | 0.795 | GCCACTGTCCCATTG |
| 1 | GMTACGGTCACACTG | chr3R:5668713-5670713 | 400 | 414 | - | 3.3571 | 7.12E-05 | 0.795 | GATACGGGCACCCAT |
| 1 | GMTACGGTCACACTG | chr3R:19402153-19404153 | 402 | 416 | - | 3.3571 | 7.12E-05 | 0.795 | GCGACGATCCCACTA |
| 1 | GMTACGGTCACACTG | chr2R:20326270-20328270 | 402 | 416 | - | 3.3571 | 7.12E-05 | 0.795 | GATACGGTCCTTCAG |
| 1 | GMTACGGTCACACTG | chr2L:14497116-14499116 | 403 | 417 | - | 3.3571 | 7.12E-05 | 0.795 | GCGACGACAACACTG |
| 1 | GMTACGGTCACACTG | chr3R:27259050-27261050 | 403 | 417 | - | 3.3571 | 7.12E-05 | 0.795 | GCTACGGTAACGGAG |
| 1 | GMTACGGTCACACTG | chr2R:12335418-12337418 | 412 | 426 | - | 3.3571 | 7.12E-05 | 0.795 | GCCACGCTCACCTTG |
| 1 | GMTACGGTCACACTG | chr2R:9819053-9821053 | 419 | 433 | - | 3.3571 | 7.12E-05 | 0.795 | GCTACCGCGACTCTG |
| 1 | GMTACGGTCACACTG | chr3R:28152301-28154301 | 419 | 433 | - | 3.3571 | 7.12E-05 | 0.795 | GACACGGGCACATAG |
| 1 | GMTACGGTCACACTG | chr3L:689517-691517 | 419 | 433 | - | 3.3571 | 7.12E-05 | 0.795 | GATACGGTCCCGGAG |
| 1 | GMTACGGTCACACTG | chr2L:8363855-8365855 | 420 | 434 | - | 3.3571 | 7.12E-05 | 0.795 | CCTACAGACGCACTG |
| 1 | GMTACGGTCACACTG | chrX:15062733-15064733 | 420 | 434 | - | 3.3571 | 7.12E-05 | 0.795 | GCTACGGTAACGGAG |
| 1 | GMTACGGTCACACTG | chr2L:4342696-4344696 | 422 | 436 | - | 3.3571 | 7.12E-05 | 0.795 | GCTCCGCTGCCACTG |
| 1 | GMTACGGTCACACTG | chr3R:27086778-27088778 | 426 | 440 | - | 3.3571 | 7.12E-05 | 0.795 | ACTACGACCACCCTG |
| 1 | GMTACGGTCACACTG | chr2L:9667501-9669501 | 426 | 440 | - | 3.3571 | 7.12E-05 | 0.795 | GCTGCTGACACATTG |
| 1 | GMTACGGTCACACTG | chrX:655033-657033 | 429 | 443 | - | 3.3571 | 7.12E-05 | 0.795 | GAGACGGACACACGC |
| 1 | GMTACGGTCACACTG | chrX:11728099-11730099 | 430 | 444 | - | 3.3571 | 7.12E-05 | 0.795 | TCAACGGCCAAACTG |
| 1 | GMTACGGTCACACTG | chr2L:16731133-16733133 | 431 | 445 | - | 3.3571 | 7.12E-05 | 0.795 | GCTACGATCCCAACG |
| 1 | GMTACGGTCACACTG | chrX:5963247-5965247 | 431 | 445 | - | 3.3571 | 7.12E-05 | 0.795 | GATCCAGCCACACAG |
| 1 | GMTACGGTCACACTG | chr2L:5903689-5905689 | 432 | 446 | - | 3.3571 | 7.12E-05 | 0.795 | CCTGCTGCCACACTG |
| 1 | GMTACGGTCACACTG | chr2L:11113014-11115014 | 433 | 447 | - | 3.3571 | 7.12E-05 | 0.795 | TAAACGGGCACGCTG |
| 1 | GMTACGGTCACACTG | chr3R:6381914-6383914 | 437 | 451 | - | 3.3571 | 7.12E-05 | 0.795 | GATACCGACGCACCG |
| 1 | GMTACGGTCACACTG | chr3R:20580522-20582522 | 439 | 453 | - | 3.3571 | 7.12E-05 | 0.795 | GCTCCCTTCACACGG |
| 1 | GMTACGGTCACACTG | chr2R:7056548-7058548 | 442 | 456 | - | 3.3571 | 7.12E-05 | 0.795 | GCTGCGGCTACAATG |
| 1 | GMTACGGTCACACTG | chrX:8922960-8924960 | 444 | 458 | - | 3.3571 | 7.12E-05 | 0.795 | GACCCGATCACGCTG |

| | | | | | | | | | |
|---|---|---|---|---|---|---|---|---|---|
| 1 | GMTACGGTCACACTG | chr2L:10731653-10733653 | 449 | 463 | - | 3.3571 | 7.12E-05 | 0.795 | GATCCGGGCACGCTT |
| 1 | GMTACGGTCACACTG | chrX:7932351-7934351 | 451 | 465 | - | 3.3571 | 7.12E-05 | 0.795 | GCTACTGCCACACGA |
| 1 | GMTACGGTCACACTG | chr2R:14756846-14758846 | 452 | 466 | - | 3.3571 | 7.12E-05 | 0.795 | GATAAGGGCTCGCTG |
| 1 | GMTACGGTCACACTG | chr2L:13210814-13212814 | 454 | 468 | - | 3.3571 | 7.12E-05 | 0.795 | GCTATCGGCGCACTG |
| 1 | GMTACGGTCACACTG | chr2R:9846836-9848836 | 457 | 471 | - | 3.3571 | 7.12E-05 | 0.795 | GCTACGTTCACGCCT |
| 1 | GMTACGGTCACACTG | chr3L:22605369-22607369 | 457 | 471 | - | 3.3571 | 7.12E-05 | 0.795 | ACTCCGCTCGCACTG |
| 1 | GMTACGGTCACACTG | chr2L:14614552-14616552 | 461 | 475 | - | 3.3571 | 7.12E-05 | 0.795 | GCTGCGGTCGCAATT |
| 1 | GMTACGGTCACACTG | chr2L:14614552-14616552 | 461 | 475 | - | 3.3571 | 7.12E-05 | 0.795 | GCTGCGGTCGCAATT |
| 1 | GMTACGGTCACACTG | chrX:21386946-21388946 | 462 | 476 | - | 3.3571 | 7.12E-05 | 0.795 | GACACGGTGACCCAG |
| 1 | GMTACGGTCACACTG | chr3L:19422758-19424758 | 464 | 478 | - | 3.3571 | 7.12E-05 | 0.795 | ACTCCGGTCTGACTG |
| 1 | GMTACGGTCACACTG | chr3L:1776439-1778439 | 468 | 482 | - | 3.3571 | 7.12E-05 | 0.795 | ACGACGGCCACACTC |
| 1 | GMTACGGTCACACTG | chr2R:23555672-23557672 | 469 | 483 | - | 3.3571 | 7.12E-05 | 0.795 | GCTTTGGCCAAACTG |
| 1 | GMTACGGTCACACTG | chr3R:15275217-15277217 | 469 | 483 | - | 3.3571 | 7.12E-05 | 0.795 | GCTGCGGGCACAATT |
| 1 | GMTACGGTCACACTG | chr2L:6971243-6973243 | 470 | 484 | - | 3.3571 | 7.12E-05 | 0.795 | GCGGGGGTCACACTT |
| 1 | GMTACGGTCACACTG | chr3R:30165419-30167419 | 471 | 485 | - | 3.3571 | 7.12E-05 | 0.795 | CCTACGGTCGCGGTG |
| 1 | GMTACGGTCACACTG | chr3R:30489074-30491074 | 475 | 489 | - | 3.3571 | 7.12E-05 | 0.795 | GCTGCAATCACTCTG |
| 1 | GMTACGGTCACACTG | chr3R:18273446-18275446 | 475 | 489 | - | 3.3571 | 7.12E-05 | 0.795 | GCCACCGTCATACGG |
| 1 | GMTACGGTCACACTG | chr3L:3470309-3472309 | 475 | 489 | - | 3.3571 | 7.12E-05 | 0.795 | GAAACGGCCACGTTG |
| 1 | GMTACGGTCACACTG | chrX:15836830-15838830 | 480 | 494 | - | 3.3571 | 7.12E-05 | 0.795 | GATCCTGTCGCCCTG |
| 1 | GMTACGGTCACACTG | chrX:634545-636545 | 484 | 498 | - | 3.3571 | 7.12E-05 | 0.795 | GCTGGGATCACGCTG |
| 1 | GMTACGGTCACACTG | chrX:19580981-19582981 | 484 | 498 | - | 3.3571 | 7.12E-05 | 0.795 | GCTAAGGCCAATCTG |
| 1 | GMTACGGTCACACTG | chr3R:5820440-5822440 | 487 | 501 | - | 3.3571 | 7.12E-05 | 0.795 | GACGTGGTCACTCTG |
| 1 | GMTACGGTCACACTG | chrX:8144839-8146839 | 489 | 503 | - | 3.3571 | 7.12E-05 | 0.795 | GAAACGGTCACCCAC |
| 1 | GMTACGGTCACACTG | chr2R:23924236-23926236 | 493 | 507 | - | 3.3571 | 7.12E-05 | 0.795 | GCTCCGCTCTCACTT |
| 1 | GMTACGGTCACACTG | chr3L:1776411-1778411 | 496 | 510 | - | 3.3571 | 7.12E-05 | 0.795 | ACGACGGCCACACTC |
| 1 | GMTACGGTCACACTG | chr3R:5632627-5634627 | 497 | 511 | - | 3.3571 | 7.12E-05 | 0.795 | TCGACGGGCAGACTG |
| 1 | GMTACGGTCACACTG | chr3L:3941710-3943710 | 497 | 511 | - | 3.3571 | 7.12E-05 | 0.795 | GCTACCGGCACATAG |
| 1 | GMTACGGTCACACTG | chr2R:20561331-20563331 | 497 | 511 | - | 3.3571 | 7.12E-05 | 0.795 | GCTAGCGACGCACTG |
| 1 | GMTACGGTCACACTG | chr3R:20742869-20744869 | 501 | 515 | - | 3.3571 | 7.12E-05 | 0.795 | GCTCCGGTCGGAGTG |
| 1 | GMTACGGTCACACTG | chr2L:5531223-5533223 | 502 | 516 | - | 3.3571 | 7.12E-05 | 0.795 | AACACGGACGCACTG |
| 1 | GMTACGGTCACACTG | chr3R:4947096-4949096 | 503 | 517 | - | 3.3571 | 7.12E-05 | 0.795 | GCCACGCCCACACTT |
| 1 | GMTACGGTCACACTG | chr2R:23712900-23714900 | 504 | 518 | - | 3.3571 | 7.12E-05 | 0.795 | GAGACGGTCGCTATG |
| 1 | GMTACGGTCACACTG | chrX:18493168-18495168 | 505 | 519 | - | 3.3571 | 7.12E-05 | 0.795 | GCAAGGGTGCCACTG |

| | | | | | | | | | |
|---|---|---|---|---|---|---|---|---|---|
| 1 | GMTACGGTCACACTG | chr3L:9434399-9436399 | 509 | 523 | - | 3.3571 | 7.12E-05 | 0.795 | GAAACGGCATCACTG |
| 1 | GMTACGGTCACACTG | chr2L:20646817-20648817 | 510 | 524 | - | 3.3571 | 7.12E-05 | 0.795 | GCTACGGACACATAA |
| 1 | GMTACGGTCACACTG | chr2L:19044148-19046148 | 516 | 530 | - | 3.3571 | 7.12E-05 | 0.795 | GCTGCTTTCACTCTG |
| 1 | GMTACGGTCACACTG | chr3L:331309-333309 | 525 | 539 | - | 3.3571 | 7.12E-05 | 0.795 | CCTACGCCCACACGG |
| 1 | GMTACGGTCACACTG | chr2L:20733609-20735609 | 526 | 540 | - | 3.3571 | 7.12E-05 | 0.795 | GACACGGCCACGTTG |
| 1 | GMTACGGTCACACTG | chr3R:15013770-15015770 | 528 | 542 | - | 3.3571 | 7.12E-05 | 0.795 | ACGACGGTCTGACTG |
| 1 | GMTACGGTCACACTG | chr3R:6695271-6697271 | 529 | 543 | - | 3.3571 | 7.12E-05 | 0.795 | GCTTCCGTCGTACTG |
| 1 | GMTACGGTCACACTG | chr3R:12421429-12423429 | 530 | 544 | - | 3.3571 | 7.12E-05 | 0.795 | TCTACGGTACCACGG |
| 1 | GMTACGGTCACACTG | chr2R:7713923-7715923 | 532 | 546 | - | 3.3571 | 7.12E-05 | 0.795 | GCTGAGGTCACACGT |
| 1 | GMTACGGTCACACTG | chr3R:30035870-30037870 | 533 | 547 | - | 3.3571 | 7.12E-05 | 0.795 | ACCACCGTCACCCTG |
| 1 | GMTACGGTCACACTG | chr3L:5358392-5360392 | 535 | 549 | - | 3.3571 | 7.12E-05 | 0.795 | CCTACGGACAAACGG |
| 1 | GMTACGGTCACACTG | chrX:940075-942075 | 536 | 550 | - | 3.3571 | 7.12E-05 | 0.795 | ACCACGTTCACACAG |
| 1 | GMTACGGTCACACTG | chr2R:21716804-21718804 | 537 | 551 | - | 3.3571 | 7.12E-05 | 0.795 | GCTATGGACACCCTT |
| 1 | GMTACGGTCACACTG | chr2R:14876786-14878786 | 543 | 557 | - | 3.3571 | 7.12E-05 | 0.795 | TCTGTGGACACACTG |
| 1 | GMTACGGTCACACTG | chr2L:8682451-8684451 | 543 | 557 | - | 3.3571 | 7.12E-05 | 0.795 | ACGACGGTGACTCTG |
| 1 | GMTACGGTCACACTG | chr3L:16004193-16006193 | 546 | 560 | - | 3.3571 | 7.12E-05 | 0.795 | ACCGCGGTCACACTA |
| 1 | GMTACGGTCACACTG | chr2R:17738901-17740901 | 546 | 560 | - | 3.3571 | 7.12E-05 | 0.795 | GCTTGGGTCACAAGG |
| 1 | GMTACGGTCACACTG | chr2R:8310645-8312645 | 549 | 563 | - | 3.3571 | 7.12E-05 | 0.795 | GCCACACTCACACAG |
| 1 | GMTACGGTCACACTG | chr3R:19228276-19230276 | 555 | 569 | - | 3.3571 | 7.12E-05 | 0.795 | GATACGGTCTCGAAG |
| 1 | GMTACGGTCACACTG | chr3L:21954110-21956110 | 558 | 572 | - | 3.3571 | 7.12E-05 | 0.795 | GCAACGCCCACACTT |
| 1 | GMTACGGTCACACTG | chr2L:2364345-2366345 | 562 | 576 | - | 3.3571 | 7.12E-05 | 0.795 | CCTGCGGTCACACAT |
| 1 | GMTACGGTCACACTG | chr2L:4683461-4685461 | 565 | 579 | - | 3.3571 | 7.12E-05 | 0.795 | GCTAAGGACACAGCG |
| 1 | GMTACGGTCACACTG | chr3R:5741510-5743510 | 568 | 582 | - | 3.3571 | 7.12E-05 | 0.795 | GAGACGGCCAATCTG |
| 1 | GMTACGGTCACACTG | chr3R:25399645-25401645 | 570 | 584 | - | 3.3571 | 7.12E-05 | 0.795 | GCTACCACCACACCG |
| 1 | GMTACGGTCACACTG | chr3R:10853501-10855501 | 570 | 584 | - | 3.3571 | 7.12E-05 | 0.795 | GATGCGGTCAATCAG |
| 1 | GMTACGGTCACACTG | chr3R:22922352-22924352 | 571 | 585 | - | 3.3571 | 7.12E-05 | 0.795 | GCTACGGAAACTTTG |
| 1 | GMTACGGTCACACTG | chr2L:20655178-20657178 | 572 | 586 | - | 3.3571 | 7.12E-05 | 0.795 | GATGCTGTCGCACGG |
| 1 | GMTACGGTCACACTG | chrX:2060915-2062915 | 573 | 587 | - | 3.3571 | 7.12E-05 | 0.795 | GATGCGGTTGCTCTG |
| 1 | GMTACGGTCACACTG | chr3R:10433789-10435789 | 580 | 594 | - | 3.3571 | 7.12E-05 | 0.795 | GCAACGTTGACCCTG |
| 1 | GMTACGGTCACACTG | chr2L:20530066-20532066 | 580 | 594 | - | 3.3571 | 7.12E-05 | 0.795 | GCAAAGGTCAAACGG |
| 1 | GMTACGGTCACACTG | chr3L:7380614-7382614 | 581 | 595 | - | 3.3571 | 7.12E-05 | 0.795 | GATCCGGTGACTCAG |
| 1 | GMTACGGTCACACTG | chr2R:20561245-20563245 | 583 | 597 | - | 3.3571 | 7.12E-05 | 0.795 | GCTAGCGACGCACTG |
| 1 | GMTACGGTCACACTG | chr2R:6058558-6060558 | 584 | 598 | - | 3.3571 | 7.12E-05 | 0.795 | GATACGGCAGCCCTG |

| | | | | | | | | | |
|---|---|---|---|---|---|---|---|---|---|
| 1 | GMTACGGTCACACTG | chr3L:17840671-17842671 | 585 | 599 | - | 3.3571 | 7.12E-05 | 0.795 | GCTAAGGTGCCACAG |
| 1 | GMTACGGTCACACTG | chr3L:15715453-15717453 | 586 | 600 | - | 3.3571 | 7.12E-05 | 0.795 | GCTCCGTTCACACAC |
| 1 | GMTACGGTCACACTG | chr2L:14424189-14426189 | 589 | 603 | - | 3.3571 | 7.12E-05 | 0.795 | GACATGGCCACACGG |
| 1 | GMTACGGTCACACTG | chrX:8144839-8146839 | 591 | 605 | - | 3.3571 | 7.12E-05 | 0.795 | AAGGCGGACACACTG |
| 1 | GMTACGGTCACACTG | chrX:7103885-7105885 | 591 | 605 | - | 3.3571 | 7.12E-05 | 0.795 | GCTCTGGTCACACCA |
| 1 | GMTACGGTCACACTG | chr3L:20403270-20405270 | 592 | 606 | - | 3.3571 | 7.12E-05 | 0.795 | GCTTGGGGCACAGTG |
| 1 | GMTACGGTCACACTG | chr2R:6197266-6199266 | 595 | 609 | - | 3.3571 | 7.12E-05 | 0.795 | GCTACGATCCCAAGG |
| 1 | GMTACGGTCACACTG | chr3L:8228224-8230224 | 597 | 611 | - | 3.3571 | 7.12E-05 | 0.795 | GACACGGCCACACAA |
| 1 | GMTACGGTCACACTG | chrX:18773027-18775027 | 599 | 613 | - | 3.3571 | 7.12E-05 | 0.795 | GAAACGGAGGCACTG |
| 1 | GMTACGGTCACACTG | chrX:16699170-16701170 | 599 | 613 | - | 3.3571 | 7.12E-05 | 0.795 | GCCACCGTCACCGTG |
| 1 | GMTACGGTCACACTG | chr3R:28302571-28304571 | 599 | 613 | - | 3.3571 | 7.12E-05 | 0.795 | GATCCGGGAGCACTG |
| 1 | GMTACGGTCACACTG | chr3R:5741476-5743476 | 602 | 616 | - | 3.3571 | 7.12E-05 | 0.795 | GAGACGGCCAATCTG |
| 1 | GMTACGGTCACACTG | chr2L:20638015-20640015 | 605 | 619 | - | 3.3571 | 7.12E-05 | 0.795 | CCGACGGGCACACTA |
| 1 | GMTACGGTCACACTG | chr2R:24782555-24784555 | 606 | 620 | - | 3.3571 | 7.12E-05 | 0.795 | GATTCGATCCCACCG |
| 1 | GMTACGGTCACACTG | chr3R:12425578-12427578 | 611 | 625 | - | 3.3571 | 7.12E-05 | 0.795 | GCTAGTGTCCCCCTG |
| 1 | GMTACGGTCACACTG | chr2L:20725551-20727551 | 612 | 626 | - | 3.3571 | 7.12E-05 | 0.795 | GAGACGGTGCCTCTG |
| 1 | GMTACGGTCACACTG | chr2L:8363662-8365662 | 613 | 627 | - | 3.3571 | 7.12E-05 | 0.795 | CCTACAGACGCACTG |
| 1 | GMTACGGTCACACTG | chr3L:9426010-9428010 | 614 | 628 | - | 3.3571 | 7.12E-05 | 0.795 | GCTACGGCCACAAAT |
| 1 | GMTACGGTCACACTG | chr3L:13389006-13391006 | 617 | 631 | - | 3.3571 | 7.12E-05 | 0.795 | GACACGATCGCACCG |
| 1 | GMTACGGTCACACTG | chr3L:21267335-21269335 | 619 | 633 | - | 3.3571 | 7.12E-05 | 0.795 | GCTTTGGTCACTTTG |
| 1 | GMTACGGTCACACTG | chr3R:10853451-10855451 | 620 | 634 | - | 3.3571 | 7.12E-05 | 0.795 | GATGCGGTCAATCAG |
| 1 | GMTACGGTCACACTG | chrX:3463044-3465044 | 622 | 636 | - | 3.3571 | 7.12E-05 | 0.795 | GCCGCGGTGACACTA |
| 1 | GMTACGGTCACACTG | chr2L:22755349-22757349 | 623 | 637 | - | 3.3571 | 7.12E-05 | 0.795 | GCTCAGGTAGCACTG |
| 1 | GMTACGGTCACACTG | chr2L:6475975-6477975 | 627 | 641 | - | 3.3571 | 7.12E-05 | 0.795 | GAGACGGCCATCCTG |
| 1 | GMTACGGTCACACTG | chr2R:22206987-22208987 | 628 | 642 | - | 3.3571 | 7.12E-05 | 0.795 | GCCAAGGAAACACTG |
| 1 | GMTACGGTCACACTG | chrX:15703125-15705125 | 629 | 643 | - | 3.3571 | 7.12E-05 | 0.795 | CCAAAGGTCACGCTG |
| 1 | GMTACGGTCACACTG | chr2L:10392434-10394434 | 633 | 647 | - | 3.3571 | 7.12E-05 | 0.795 | GCTGCGTTGTCACTG |
| 1 | GMTACGGTCACACTG | chr2L:2364273-2366273 | 634 | 648 | - | 3.3571 | 7.12E-05 | 0.795 | CCTGCGGTCACACAT |
| 1 | GMTACGGTCACACTG | chr3R:17610511-17612511 | 639 | 653 | - | 3.3571 | 7.12E-05 | 0.795 | GCTACGGAGGCAGTG |
| 1 | GMTACGGTCACACTG | chrX:2060915-2062915 | 642 | 656 | - | 3.3571 | 7.12E-05 | 0.795 | GAATCGGTTGCACTG |
| 1 | GMTACGGTCACACTG | chr2L:12999635-13001635 | 643 | 657 | - | 3.3571 | 7.12E-05 | 0.795 | TCTACGATCCCACGG |
| 1 | GMTACGGTCACACTG | chr3R:6834869-6836869 | 644 | 658 | - | 3.3571 | 7.12E-05 | 0.795 | GAACCGGACACAGTG |
| 1 | GMTACGGTCACACTG | chr2L:7408809-7410809 | 644 | 658 | - | 3.3571 | 7.12E-05 | 0.795 | GAGCCGGGCAAACTG |

| | | | | | | | | | |
|---|---|---|---|---|---|---|---|---|---|
| 1 | GMTACGGTCACACTG | chr3R:12633917-12635917 | 645 | 659 | - | 3.3571 | 7.12E-05 | 0.795 | GCGACGGTCATAGAG |
| 1 | GMTACGGTCACACTG | chr2L:20483399-20485399 | 645 | 659 | - | 3.3571 | 7.12E-05 | 0.795 | GATCGGGCCACACGG |
| 1 | GMTACGGTCACACTG | chr2L:4226097-4228097 | 646 | 660 | - | 3.3571 | 7.12E-05 | 0.795 | GCAATGGTAACGCTG |
| 1 | GMTACGGTCACACTG | chr2L:16836972-16838972 | 646 | 660 | - | 3.3571 | 7.12E-05 | 0.795 | GCTGTGGCCACATTG |
| 1 | GMTACGGTCACACTG | chr2L:16836972-16838972 | 646 | 660 | - | 3.3571 | 7.12E-05 | 0.795 | GCTGTGGCCACATTG |
| 1 | GMTACGGTCACACTG | chrX:15784583-15786583 | 647 | 661 | - | 3.3571 | 7.12E-05 | 0.795 | GCAACGCTGTCACTG |
| 1 | GMTACGGTCACACTG | chr3R:21269126-21271126 | 649 | 663 | - | 3.3571 | 7.12E-05 | 0.795 | GCCGTGGTCACAGTG |
| 1 | GMTACGGTCACACTG | chr3L:21678755-21680755 | 649 | 663 | - | 3.3571 | 7.12E-05 | 0.795 | GCTACAGCTTCACTG |
| 1 | GMTACGGTCACACTG | chr3R:10042270-10044270 | 656 | 670 | - | 3.3571 | 7.12E-05 | 0.795 | GCCATGGTCACACCA |
| 1 | GMTACGGTCACACTG | chrX:7460088-7462088 | 660 | 674 | - | 3.3571 | 7.12E-05 | 0.795 | GCAACGACCAAACTG |
| 1 | GMTACGGTCACACTG | chr2R:24782500-24784500 | 661 | 675 | - | 3.3571 | 7.12E-05 | 0.795 | GATTCGATCCCACCG |
| 1 | GMTACGGTCACACTG | chr2R:19455170-19457170 | 666 | 680 | - | 3.3571 | 7.12E-05 | 0.795 | GAAACGGCCACTTTG |
| 1 | GMTACGGTCACACTG | chr2L:11011135-11013135 | 666 | 680 | - | 3.3571 | 7.12E-05 | 0.795 | GATGCGGTCGCCATG |
| 1 | GMTACGGTCACACTG | chrX:3867973-3869973 | 666 | 680 | - | 3.3571 | 7.12E-05 | 0.795 | GATGCTGTCGCTCTG |
| 1 | GMTACGGTCACACTG | chr2R:19455170-19457170 | 666 | 680 | - | 3.3571 | 7.12E-05 | 0.795 | GAAACGGCCACTTTG |
| 1 | GMTACGGTCACACTG | chr3L:3142808-3144808 | 668 | 682 | - | 3.3571 | 7.12E-05 | 0.795 | GCTGCGGTAACCCTA |
| 1 | GMTACGGTCACACTG | chr3R:11243302-11245302 | 671 | 685 | - | 3.3571 | 7.12E-05 | 0.795 | GCTCTGGTCACACAT |
| 1 | GMTACGGTCACACTG | chr2R:7524716-7526716 | 673 | 687 | - | 3.3571 | 7.12E-05 | 0.795 | GAGCCGGACACAGTG |
| 1 | GMTACGGTCACACTG | chr3R:4377600-4379600 | 675 | 689 | - | 3.3571 | 7.12E-05 | 0.795 | GCCTCGCTCACACTT |
| 1 | GMTACGGTCACACTG | chr3L:22547437-22549437 | 675 | 689 | - | 3.3571 | 7.12E-05 | 0.795 | GCCACGCCCACACTT |
| 1 | GMTACGGTCACACTG | chr3R:9632076-9634076 | 675 | 689 | - | 3.3571 | 7.12E-05 | 0.795 | GAAACAGGCTCACTG |
| 1 | GMTACGGTCACACTG | chr3L:21343033-21345033 | 676 | 690 | - | 3.3571 | 7.12E-05 | 0.795 | CCAACGGGCACACTT |
| 1 | GMTACGGTCACACTG | chr2L:10731423-10733423 | 679 | 693 | - | 3.3571 | 7.12E-05 | 0.795 | GATCCGGGCACGCTT |
| 1 | GMTACGGTCACACTG | chr2L:1165506-1167506 | 681 | 695 | - | 3.3571 | 7.12E-05 | 0.795 | GCTCCGTTCAGGCTG |
| 1 | GMTACGGTCACACTG | chr3R:5222365-5224365 | 683 | 697 | - | 3.3571 | 7.12E-05 | 0.795 | GAGGCGGTCACGGTG |
| 1 | GMTACGGTCACACTG | chrX:16414046-16416046 | 684 | 698 | - | 3.3571 | 7.12E-05 | 0.795 | GCAACGGCTAAACTG |
| 1 | GMTACGGTCACACTG | chr2L:16742405-16744405 | 686 | 700 | - | 3.3571 | 7.12E-05 | 0.795 | GCTTTGGTCACTCTT |
| 1 | GMTACGGTCACACTG | chr3R:23780765-23782765 | 688 | 702 | - | 3.3571 | 7.12E-05 | 0.795 | GCCTTGGTCACACTT |
| 1 | GMTACGGTCACACTG | chr2L:4633342-4635342 | 689 | 703 | - | 3.3571 | 7.12E-05 | 0.795 | GCTATGGCCACTTTG |
| 1 | GMTACGGTCACACTG | chr3R:8779789-8781789 | 692 | 706 | - | 3.3571 | 7.12E-05 | 0.795 | TATCCGGCCACCCTG |
| 1 | GMTACGGTCACACTG | chr2L:19581474-19583474 | 693 | 707 | - | 3.3571 | 7.12E-05 | 0.795 | GATTCGGTCATGCAG |
| 1 | GMTACGGTCACACTG | chr3L:1601954-1603954 | 697 | 711 | - | 3.3571 | 7.12E-05 | 0.795 | CCGGCGGTCACACTA |
| 1 | GMTACGGTCACACTG | chr3L:1601954-1603954 | 697 | 711 | - | 3.3571 | 7.12E-05 | 0.795 | CCGGCGGTCACACTA |

| | | | | | | | | | |
|---|---|---|---|---|---|---|---|---|---|
| 1 | GMTACGGTCACACTG | chr3L:14802360-14804360 | 697 | 711 | - | 3.3571 | 7.12E-05 | 0.795 | GATACGGACTCTTTG |
| 1 | GMTACGGTCACACTG | chrX:2680723-2682723 | 698 | 712 | - | 3.3571 | 7.12E-05 | 0.795 | GAGACGGTGACGCGG |
| 1 | GMTACGGTCACACTG | chr3L:3939125-3941125 | 699 | 713 | - | 3.3571 | 7.12E-05 | 0.795 | GCTACCGGCACATAG |
| 1 | GMTACGGTCACACTG | chr2R:14870366-14872366 | 699 | 713 | - | 3.3571 | 7.12E-05 | 0.795 | TCGACGGCCACACTT |
| 1 | GMTACGGTCACACTG | chr3R:8138606-8140606 | 700 | 714 | - | 3.3571 | 7.12E-05 | 0.795 | ACAATGGCCACACTG |
| 1 | GMTACGGTCACACTG | chr2L:12108417-12110417 | 700 | 714 | - | 3.3571 | 7.12E-05 | 0.795 | ACTCTGGTCACACAG |
| 1 | GMTACGGTCACACTG | chr3R:21269075-21271075 | 700 | 714 | - | 3.3571 | 7.12E-05 | 0.795 | GCCGTGGTCACAGTG |
| 1 | GMTACGGTCACACTG | chr3L:21679050-21681050 | 700 | 714 | - | 3.3571 | 7.12E-05 | 0.795 | GCGCTGGTCACACTA |
| 1 | GMTACGGTCACACTG | chr2L:2766890-2768890 | 705 | 719 | - | 3.3571 | 7.12E-05 | 0.795 | GATGCGGTCACCTCG |
| 1 | GMTACGGTCACACTG | chrX:17065732-17067732 | 706 | 720 | - | 3.3571 | 7.12E-05 | 0.795 | GCAACGAACACAATG |
| 1 | GMTACGGTCACACTG | chr2L:6154141-6156141 | 707 | 721 | - | 3.3571 | 7.12E-05 | 0.795 | GATGCGGTCCCTATG |
| 1 | GMTACGGTCACACTG | chrX:21622750-21624750 | 713 | 727 | - | 3.3571 | 7.12E-05 | 0.795 | GATGTGGTCACTCAG |
| 1 | GMTACGGTCACACTG | chr2R:8362218-8364218 | 716 | 730 | - | 3.3571 | 7.12E-05 | 0.795 | GAATCAGTCCCACTG |
| 1 | GMTACGGTCACACTG | chrX:8922687-8924687 | 717 | 731 | - | 3.3571 | 7.12E-05 | 0.795 | GACCCGATCACGCTG |
| 1 | GMTACGGTCACACTG | chr3L:7380474-7382474 | 721 | 735 | - | 3.3571 | 7.12E-05 | 0.795 | GATCCGGTGACTCAG |
| 1 | GMTACGGTCACACTG | chr3R:12690743-12692743 | 722 | 736 | - | 3.3571 | 7.12E-05 | 0.795 | GCAACGTTCCAACTG |
| 1 | GMTACGGTCACACTG | chrX:15062429-15064429 | 724 | 738 | - | 3.3571 | 7.12E-05 | 0.795 | GCTACGGTAACGGAG |
| 1 | GMTACGGTCACACTG | chr3L:65430-67430 | 724 | 738 | - | 3.3571 | 7.12E-05 | 0.795 | GCGACGGTGACTGTG |
| 1 | GMTACGGTCACACTG | chr3L:3046087-3048087 | 725 | 739 | - | 3.3571 | 7.12E-05 | 0.795 | AACGCGGTCACCCTG |
| 1 | GMTACGGTCACACTG | chrX:8144602-8146602 | 726 | 740 | - | 3.3571 | 7.12E-05 | 0.795 | GAAACGGTCACCCAC |
| 1 | GMTACGGTCACACTG | chr2L:15922395-15924395 | 729 | 743 | - | 3.3571 | 7.12E-05 | 0.795 | GCCTCGCTCACACTT |
| 1 | GMTACGGTCACACTG | chr3L:6141413-6143413 | 729 | 743 | - | 3.3571 | 7.12E-05 | 0.795 | GAGAGGGCCACTCTG |
| 1 | GMTACGGTCACACTG | chr3R:30744263-30746263 | 731 | 745 | - | 3.3571 | 7.12E-05 | 0.795 | GATGCCGACACCCTG |
| 1 | GMTACGGTCACACTG | chr2L:1728887-1730887 | 732 | 746 | - | 3.3571 | 7.12E-05 | 0.795 | GCTTCCAACACACTG |
| 1 | GMTACGGTCACACTG | chr3L:8294327-8296327 | 734 | 748 | - | 3.3571 | 7.12E-05 | 0.795 | GACACGGGCAGCCTG |
| 1 | GMTACGGTCACACTG | chrX:21610526-21612526 | 737 | 751 | - | 3.3571 | 7.12E-05 | 0.795 | GATGTGGTCACTCAG |
| 1 | GMTACGGTCACACTG | chr2L:8516411-8518411 | 737 | 751 | - | 3.3571 | 7.12E-05 | 0.795 | CACGCGGTCCCACTG |
| 1 | GMTACGGTCACACTG | chrX:21616617-21618617 | 738 | 752 | - | 3.3571 | 7.12E-05 | 0.795 | GATGTGGTCACTCAG |
| 1 | GMTACGGTCACACTG | chr3L:3941465-3943465 | 742 | 756 | - | 3.3571 | 7.12E-05 | 0.795 | GCTACCGGCACATAG |
| 1 | GMTACGGTCACACTG | chr2R:14238932-14240932 | 743 | 757 | - | 3.3571 | 7.12E-05 | 0.795 | CCTTCGGCCACACTT |
| 1 | GMTACGGTCACACTG | chr3L:7436466-7438466 | 744 | 758 | - | 3.3571 | 7.12E-05 | 0.795 | CCAGCTGTCACACTG |
| 1 | GMTACGGTCACACTG | chr2L:2766845-2768845 | 750 | 764 | - | 3.3571 | 7.12E-05 | 0.795 | GATGCGGTCACCTCG |
| 1 | GMTACGGTCACACTG | chr3R:18402026-18404026 | 753 | 767 | - | 3.3571 | 7.12E-05 | 0.795 | GCCACAGTCCGACTG |

| | | | | | | | | | |
|---|---|---|---|---|---|---|---|---|---|
| 1 | GMTACGGTCACACTG | chr3R:24863732-24865732 | 755 | 769 | - | 3.3571 | 7.12E-05 | 0.795 | GCTACGGCTACGGTG |
| 1 | GMTACGGTCACACTG | chr2L:10740885-10742885 | 755 | 769 | - | 3.3571 | 7.12E-05 | 0.795 | GCTTGGTCCACACTG |
| 1 | GMTACGGTCACACTG | chr2R:17738685-17740685 | 762 | 776 | - | 3.3571 | 7.12E-05 | 0.795 | GCTTGGGTCACAAGG |
| 1 | GMTACGGTCACACTG | chr2L:22750795-22752795 | 764 | 778 | - | 3.3571 | 7.12E-05 | 0.795 | GCAACGCGCACACTT |
| 1 | GMTACGGTCACACTG | chrX:1680566-1682566 | 765 | 779 | - | 3.3571 | 7.12E-05 | 0.795 | GCGACAGTTACACAG |
| 1 | GMTACGGTCACACTG | chr2L:13772997-13774997 | 768 | 782 | - | 3.3571 | 7.12E-05 | 0.795 | GCAACGCTCACGCTC |
| 1 | GMTACGGTCACACTG | chr2L:3513872-3515872 | 768 | 782 | - | 3.3571 | 7.12E-05 | 0.795 | TCTGCTGCCACACTG |
| 1 | GMTACGGTCACACTG | chr2L:8516380-8518380 | 768 | 782 | - | 3.3571 | 7.12E-05 | 0.795 | CACGCGGTCCCACTG |
| 1 | GMTACGGTCACACTG | chr3R:26344473-26346473 | 772 | 786 | - | 3.3571 | 7.12E-05 | 0.795 | TCGATGGTCACGCTG |
| 1 | GMTACGGTCACACTG | chr3R:25045323-25047323 | 774 | 788 | - | 3.3571 | 7.12E-05 | 0.795 | GCTACCGTCTCACGT |
| 1 | GMTACGGTCACACTG | chr2R:16347527-16349527 | 776 | 790 | - | 3.3571 | 7.12E-05 | 0.795 | GCAACGCTTCCACTG |
| 1 | GMTACGGTCACACTG | chr3R:5668334-5670334 | 779 | 793 | - | 3.3571 | 7.12E-05 | 0.795 | GATACGGGCACCCAT |
| 1 | GMTACGGTCACACTG | chr3R:10853501-10855501 | 779 | 793 | - | 3.3571 | 7.12E-05 | 0.795 | GATCCGGCGACACCG |
| 1 | GMTACGGTCACACTG | chr2L:6556800-6558800 | 781 | 795 | - | 3.3571 | 7.12E-05 | 0.795 | ACAACGGTCACCCTA |
| 1 | GMTACGGTCACACTG | chr3L:6262399-6264399 | 782 | 796 | - | 3.3571 | 7.12E-05 | 0.795 | GCAAAGGTCACCTTG |
| 1 | GMTACGGTCACACTG | chr2R:13854778-13856778 | 783 | 797 | - | 3.3571 | 7.12E-05 | 0.795 | GAGACGGTCGCCATG |
| 1 | GMTACGGTCACACTG | chr3R:4964445-4966445 | 784 | 798 | - | 3.3571 | 7.12E-05 | 0.795 | GCTACTGACTGACTG |
| 1 | GMTACGGTCACACTG | chrX:16705874-16707874 | 784 | 798 | - | 3.3571 | 7.12E-05 | 0.795 | GATACGCCCTCACCG |
| 1 | GMTACGGTCACACTG | chr2R:9585714-9587714 | 784 | 798 | - | 3.3571 | 7.12E-05 | 0.795 | GATACGGGCTCAAAG |
| 1 | GMTACGGTCACACTG | chr2R:9144260-9146260 | 789 | 803 | - | 3.3571 | 7.12E-05 | 0.795 | GCCACGGTCACGTTC |
| 1 | GMTACGGTCACACTG | chr2L:2219748-2221748 | 791 | 805 | - | 3.3571 | 7.12E-05 | 0.795 | GCTACGGGCACATCC |
| 1 | GMTACGGTCACACTG | chr3R:11892297-11894297 | 791 | 805 | - | 3.3571 | 7.12E-05 | 0.795 | GCTCCGCTCACAGAG |
| 1 | GMTACGGTCACACTG | chr2L:14746080-14748080 | 792 | 806 | - | 3.3571 | 7.12E-05 | 0.795 | GCTACGTTCATTCAG |
| 1 | GMTACGGTCACACTG | chr2L:20646533-20648533 | 794 | 808 | - | 3.3571 | 7.12E-05 | 0.795 | GCTACGGACACATAA |
| 1 | GMTACGGTCACACTG | chr2R:23547239-23549239 | 795 | 809 | - | 3.3571 | 7.12E-05 | 0.795 | GCTGCCGTGACACAG |
| 1 | GMTACGGTCACACTG | chr3L:21515306-21517306 | 795 | 809 | - | 3.3571 | 7.12E-05 | 0.795 | GCTCTGGCAACACTG |
| 1 | GMTACGGTCACACTG | chr3R:5597777-5599777 | 799 | 813 | - | 3.3571 | 7.12E-05 | 0.795 | GCGAAGGTCACGCTA |
| 1 | GMTACGGTCACACTG | chrX:15932867-15934867 | 801 | 815 | - | 3.3571 | 7.12E-05 | 0.795 | GCCTTGGTCACACTT |
| 1 | GMTACGGTCACACTG | chrX:939805-941805 | 806 | 820 | - | 3.3571 | 7.12E-05 | 0.795 | ACCACGTTCACACAG |
| 1 | GMTACGGTCACACTG | chrX:1029758-1031758 | 807 | 821 | - | 3.3571 | 7.12E-05 | 0.795 | ACACTGGTCACACTG |
| 1 | GMTACGGTCACACTG | chr3R:7084660-7086660 | 808 | 822 | - | 3.3571 | 7.12E-05 | 0.795 | GCTCCGGTAACAAAG |
| 1 | GMTACGGTCACACTG | chr2L:4255519-4257519 | 810 | 824 | - | 3.3571 | 7.12E-05 | 0.795 | GCAACTGCAACACTG |
| 1 | GMTACGGTCACACTG | chr3L:16338003-16340003 | 814 | 828 | - | 3.3571 | 7.12E-05 | 0.795 | GCGACGGCGACAATG |

| | | | | | | | | | |
|---|---|---|---|---|---|---|---|---|---|
| 1 | GMTACGGTCACACTG | chr3R:12633744-12635744 | 818 | 832 | - | 3.3571 | 7.12E-05 | 0.795 | GCGACGGTCATAGAG |
| 1 | GMTACGGTCACACTG | chr3R:6834690-6836690 | 823 | 837 | - | 3.3571 | 7.12E-05 | 0.795 | GAACCGGACACAGTG |
| 1 | GMTACGGTCACACTG | chr2L:15462254-15464254 | 824 | 838 | - | 3.3571 | 7.12E-05 | 0.795 | GAATCGGACACACTC |
| 1 | GMTACGGTCACACTG | chr2L:20733309-20735309 | 826 | 840 | - | 3.3571 | 7.12E-05 | 0.795 | GACACGGCCACGTTG |
| 1 | GMTACGGTCACACTG | chr3R:27291209-27293209 | 826 | 840 | - | 3.3571 | 7.12E-05 | 0.795 | GCGACGATCACGCTC |
| 1 | GMTACGGTCACACTG | chrX:8144602-8146602 | 828 | 842 | - | 3.3571 | 7.12E-05 | 0.795 | AAGGCGGACACACTG |
| 1 | GMTACGGTCACACTG | chr3L:17840427-17842427 | 829 | 843 | - | 3.3571 | 7.12E-05 | 0.795 | GCTAAGGTGCCACAG |
| 1 | GMTACGGTCACACTG | chr3R:10853451-10855451 | 829 | 843 | - | 3.3571 | 7.12E-05 | 0.795 | GATCCGGCGACACCG |
| 1 | GMTACGGTCACACTG | chr3R:5482906-5484906 | 831 | 845 | - | 3.3571 | 7.12E-05 | 0.795 | GCGAGGGCCACAATG |
| 1 | GMTACGGTCACACTG | chr3R:23780621-23782621 | 832 | 846 | - | 3.3571 | 7.12E-05 | 0.795 | GCCTTGGTCACACTT |
| 1 | GMTACGGTCACACTG | chr2L:6064997-6066997 | 834 | 848 | - | 3.3571 | 7.12E-05 | 0.795 | CCCAAGGGCACACTG |
| 1 | GMTACGGTCACACTG | chr2R:24237477-24239477 | 842 | 856 | - | 3.3571 | 7.12E-05 | 0.795 | GCCACACACACACTG |
| 1 | GMTACGGTCACACTG | chr3R:9645154-9647154 | 844 | 858 | - | 3.3571 | 7.12E-05 | 0.795 | GCTCCGGTGACGATG |
| 1 | GMTACGGTCACACTG | chr2R:23139577-23141577 | 845 | 859 | - | 3.3571 | 7.12E-05 | 0.795 | GCCACGGTTCCACTA |
| 1 | GMTACGGTCACACTG | chr2R:13162948-13164948 | 845 | 859 | - | 3.3571 | 7.12E-05 | 0.795 | GAGGCGGTCATGCTG |
| 1 | GMTACGGTCACACTG | chrX:4960128-4962128 | 847 | 861 | - | 3.3571 | 7.12E-05 | 0.795 | GCGACTGTCACAAGG |
| 1 | GMTACGGTCACACTG | chr2R:18672536-18674536 | 849 | 863 | - | 3.3571 | 7.12E-05 | 0.795 | GCAACTGGCAAACTG |
| 1 | GMTACGGTCACACTG | chr2R:21935863-21937863 | 851 | 865 | - | 3.3571 | 7.12E-05 | 0.795 | CAAACGGACACACGG |
| 1 | GMTACGGTCACACTG | chr3R:20303977-20305977 | 852 | 866 | - | 3.3571 | 7.12E-05 | 0.795 | GCTACGATAACTCGG |
| 1 | GMTACGGTCACACTG | chrX:16421831-16423831 | 853 | 867 | - | 3.3571 | 7.12E-05 | 0.795 | TACCCGGTCACTCTG |
| 1 | GMTACGGTCACACTG | chr3L:689081-691081 | 855 | 869 | - | 3.3571 | 7.12E-05 | 0.795 | GATACGGTCCCGGAG |
| 1 | GMTACGGTCACACTG | chr2L:1792268-1794268 | 858 | 872 | - | 3.3571 | 7.12E-05 | 0.795 | GAGACGATCCCGCTG |
| 1 | GMTACGGTCACACTG | chrX:16358143-16360143 | 861 | 875 | - | 3.3571 | 7.12E-05 | 0.795 | GCCAGTGTCGCACTG |
| 1 | GMTACGGTCACACTG | chr3L:14052215-14054215 | 865 | 879 | - | 3.3571 | 7.12E-05 | 0.795 | CCAACGGTCGCACTT |
| 1 | GMTACGGTCACACTG | chr2R:15114236-15116236 | 865 | 879 | - | 3.3571 | 7.12E-05 | 0.795 | GCAACGCTCACGCTC |
| 1 | GMTACGGTCACACTG | chr3L:9425758-9427758 | 866 | 880 | - | 3.3571 | 7.12E-05 | 0.795 | GCTACGGCCACAAAT |
| 1 | GMTACGGTCACACTG | chr2R:11678492-11680492 | 866 | 880 | - | 3.3571 | 7.12E-05 | 0.795 | GCTGTGGCAACACTG |
| 1 | GMTACGGTCACACTG | chr2R:20325806-20327806 | 866 | 880 | - | 3.3571 | 7.12E-05 | 0.795 | GATACGGTCCTTCAG |
| 1 | GMTACGGTCACACTG | chr3R:21831503-21833503 | 867 | 881 | - | 3.3571 | 7.12E-05 | 0.795 | CCTTCAGGCACACTG |
| 1 | GMTACGGTCACACTG | chr3L:7538836-7540836 | 869 | 883 | - | 3.3571 | 7.12E-05 | 0.795 | GCTTCGATCACGGTG |
| 1 | GMTACGGTCACACTG | chr2L:4458729-4460729 | 873 | 887 | - | 3.3571 | 7.12E-05 | 0.795 | TCGGGGGTCACACTG |
| 1 | GMTACGGTCACACTG | chrX:16080056-16082056 | 875 | 889 | - | 3.3571 | 7.12E-05 | 0.795 | GAAACGGTGCCACCG |
| 1 | GMTACGGTCACACTG | chr2R:17543190-17545190 | 876 | 890 | - | 3.3571 | 7.12E-05 | 0.795 | GAAACAGCCACACGG |

| | | | | | | | | | |
|---|---|---|---|---|---|---|---|---|---|
| 1 | GMTACGGTCACACTG | chr2L:3415693-3417693 | 877 | 891 | - | 3.3571 | 7.12E-05 | 0.795 | AAACCGGTCACGCTG |
| 1 | GMTACGGTCACACTG | chr2R:11341760-11343760 | 877 | 891 | - | 3.3571 | 7.12E-05 | 0.795 | GCGAAGGTCGCATTG |
| 1 | GMTACGGTCACACTG | chr2R:20006095-20008095 | 877 | 891 | - | 3.3571 | 7.12E-05 | 0.795 | GCTAGGGCGCCACTG |
| 1 | GMTACGGTCACACTG | chr2L:6064953-6066953 | 878 | 892 | - | 3.3571 | 7.12E-05 | 0.795 | CCCAAGGGCACACTG |
| 1 | GMTACGGTCACACTG | chr2R:10945309-10947309 | 878 | 892 | - | 3.3571 | 7.12E-05 | 0.795 | GATCCGGTCACTGCG |
| 1 | GMTACGGTCACACTG | chr3R:29994357-29996357 | 880 | 894 | - | 3.3571 | 7.12E-05 | 0.795 | TCGCTGGTCACACTG |
| 1 | GMTACGGTCACACTG | chr2R:23547151-23549151 | 883 | 897 | - | 3.3571 | 7.12E-05 | 0.795 | GCTGCCGTGACACAG |
| 1 | GMTACGGTCACACTG | chr2R:24501277-24503277 | 886 | 900 | - | 3.3571 | 7.12E-05 | 0.795 | GATACGATCGCGCGG |
| 1 | GMTACGGTCACACTG | chr3R:12675260-12677260 | 892 | 906 | - | 3.3571 | 7.12E-05 | 0.795 | GCCATGGTGACCCTG |
| 1 | GMTACGGTCACACTG | chr3L:9542999-9544999 | 893 | 907 | - | 3.3571 | 7.12E-05 | 0.795 | GAAACGGTCCTTCTG |
| 1 | GMTACGGTCACACTG | chr2R:8230065-8232065 | 894 | 908 | - | 3.3571 | 7.12E-05 | 0.795 | GCGAAGATCGCACTG |
| 1 | GMTACGGTCACACTG | chr2R:10379109-10381109 | 900 | 914 | - | 3.3571 | 7.12E-05 | 0.795 | GCTTGGGGCACAATG |
| 1 | GMTACGGTCACACTG | chr2R:6241004-6243004 | 902 | 916 | - | 3.3571 | 7.12E-05 | 0.795 | GCTTCGATCAAACAG |
| 1 | GMTACGGTCACACTG | chrX:4781789-4783789 | 903 | 917 | - | 3.3571 | 7.12E-05 | 0.795 | GCTATGGCCAGCCTG |
| 1 | GMTACGGTCACACTG | chr3R:31304384-31306384 | 905 | 919 | - | 3.3571 | 7.12E-05 | 0.795 | GCTACTGGCGGACTG |
| 1 | GMTACGGTCACACTG | chr3L:11131322-11133322 | 909 | 923 | - | 3.3571 | 7.12E-05 | 0.795 | GCCACGGTCACTGTT |
| 1 | GMTACGGTCACACTG | chrX:5868095-5870095 | 909 | 923 | - | 3.3571 | 7.12E-05 | 0.795 | GAGACGGGCCTACTG |
| 1 | GMTACGGTCACACTG | chrX:15778146-15780146 | 914 | 928 | - | 3.3571 | 7.12E-05 | 0.795 | GCGACGGTTACATGG |
| 1 | GMTACGGTCACACTG | chr3R:9256716-9258716 | 918 | 932 | - | 3.3571 | 7.12E-05 | 0.795 | ACTACGGCCCAACTG |
| 1 | GMTACGGTCACACTG | chr2R:16293400-16295400 | 919 | 933 | - | 3.3571 | 7.12E-05 | 0.795 | GCAGTGTTCACACTG |
| 1 | GMTACGGTCACACTG | chr3R:29876059-29878059 | 920 | 934 | - | 3.3571 | 7.12E-05 | 0.795 | TACACGGCCGCACTG |
| 1 | GMTACGGTCACACTG | chr3R:15803248-15805248 | 920 | 934 | - | 3.3571 | 7.12E-05 | 0.795 | TCTGCGGCCACATTG |
| 1 | GMTACGGTCACACTG | chr3R:20748074-20750074 | 920 | 934 | - | 3.3571 | 7.12E-05 | 0.795 | GCTACGGTCGAGCTC |
| 1 | GMTACGGTCACACTG | chr2L:20425199-20427199 | 922 | 936 | - | 3.3571 | 7.12E-05 | 0.795 | GCTGGGGCGACACTG |
| 1 | GMTACGGTCACACTG | chr3R:26977541-26979541 | 923 | 937 | - | 3.3571 | 7.12E-05 | 0.795 | GCTATGGCGTCACTG |
| 1 | GMTACGGTCACACTG | chr3R:26977541-26979541 | 923 | 937 | - | 3.3571 | 7.12E-05 | 0.795 | GCTATGGCGTCACTG |
| 1 | GMTACGGTCACACTG | chr2R:23555217-23557217 | 924 | 938 | - | 3.3571 | 7.12E-05 | 0.795 | GCTTTGGCCAAACTG |
| 1 | GMTACGGTCACACTG | chr3L:3946891-3948891 | 924 | 938 | - | 3.3571 | 7.12E-05 | 0.795 | GCAACGGTATCAATG |
| 1 | GMTACGGTCACACTG | chr3L:20969663-20971663 | 929 | 943 | - | 3.3571 | 7.12E-05 | 0.795 | ACTACGGAAACACCG |
| 1 | GMTACGGTCACACTG | chrX:16390698-16392698 | 930 | 944 | - | 3.3571 | 7.12E-05 | 0.795 | GCCACGTTCCAACTG |
| 1 | GMTACGGTCACACTG | chrX:3372159-3374159 | 931 | 945 | - | 3.3571 | 7.12E-05 | 0.795 | TCTGCGCTCTCACTG |
| 1 | GMTACGGTCACACTG | chrX:3372159-3374159 | 931 | 945 | - | 3.3571 | 7.12E-05 | 0.795 | TCTGCGCTCTCACTG |
| 1 | GMTACGGTCACACTG | chr2L:856542-858542 | 938 | 952 | - | 3.3571 | 7.12E-05 | 0.795 | GCCACCTCCACACTG |

| | | | | | | | | | |
|---|---|---|---|---|---|---|---|---|---|
| 1 | GMTACGGTCACACTG | chrX:9249420-9251420 | 941 | 955 | - | 3.3571 | 7.12E-05 | 0.795 | ACTCTGGCCACACTG |
| 1 | GMTACGGTCACACTG | chrX:11492135-11494135 | 946 | 960 | - | 3.3571 | 7.12E-05 | 0.795 | TCTCTGGTCACTCTG |
| 1 | GMTACGGTCACACTG | chr2L:6049800-6051800 | 947 | 961 | - | 3.3571 | 7.12E-05 | 0.795 | TCGCCGTTCACACTG |
| 1 | GMTACGGTCACACTG | chrX:6302975-6304975 | 947 | 961 | - | 3.3571 | 7.12E-05 | 0.795 | GCCACTGGAACACTG |
| 1 | GMTACGGTCACACTG | chr3R:27904843-27906843 | 949 | 963 | - | 3.3571 | 7.12E-05 | 0.795 | GCTCTCGTCCCACTG |
| 1 | GMTACGGTCACACTG | chr3L:22708918-22710918 | 949 | 963 | - | 3.3571 | 7.12E-05 | 0.795 | CCTACGGTCCCGCTT |
| 1 | GMTACGGTCACACTG | chrX:17906692-17908692 | 951 | 965 | - | 3.3571 | 7.12E-05 | 0.795 | GCAATGGTCACACAT |
| 1 | GMTACGGTCACACTG | chr3R:8012791-8014791 | 954 | 968 | - | 3.3571 | 7.12E-05 | 0.795 | GCAATGGTCGAACTG |
| 1 | GMTACGGTCACACTG | chr2L:2250275-2252275 | 954 | 968 | - | 3.3571 | 7.12E-05 | 0.795 | GCCAGCGTCACCCTG |
| 1 | GMTACGGTCACACTG | chr2L:13397991-13399991 | 957 | 971 | - | 3.3571 | 7.12E-05 | 0.795 | GCAACAGGCACACTA |
| 1 | GMTACGGTCACACTG | chrX:13336705-13338705 | 958 | 972 | - | 3.3571 | 7.12E-05 | 0.795 | AACGCGGCCACACTG |
| 1 | GMTACGGTCACACTG | chr3L:12265841-12267841 | 959 | 973 | - | 3.3571 | 7.12E-05 | 0.795 | AATTCGGACACACCG |
| 1 | GMTACGGTCACACTG | chrX:21338741-21340741 | 960 | 974 | - | 3.3571 | 7.12E-05 | 0.795 | GAACTGGCCACACTG |
| 1 | GMTACGGTCACACTG | chr2R:8310234-8312234 | 960 | 974 | - | 3.3571 | 7.12E-05 | 0.795 | GCCACACTCACACAG |
| 1 | GMTACGGTCACACTG | chr3L:19812339-19814339 | 961 | 975 | - | 3.3571 | 7.12E-05 | 0.795 | CATACGGACGCTCTG |
| 1 | GMTACGGTCACACTG | chrX:6266228-6268228 | 961 | 975 | - | 3.3571 | 7.12E-05 | 0.795 | TCTCTGGGCACACTG |
| 1 | GMTACGGTCACACTG | chr3L:12660826-12662826 | 962 | 976 | - | 3.3571 | 7.12E-05 | 0.795 | GACACAGACACGCTG |
| 1 | GMTACGGTCACACTG | chrX:8093064-8095064 | 962 | 976 | - | 3.3571 | 7.12E-05 | 0.795 | GCGCTGGTCACACTA |
| 1 | GMTACGGTCACACTG | chr3L:691919-693919 | 963 | 977 | - | 3.3571 | 7.12E-05 | 0.795 | GACTCGGCCACACTT |
| 1 | GMTACGGTCACACTG | chr3R:28641891-28643891 | 963 | 977 | - | 3.3571 | 7.12E-05 | 0.795 | GCTACTGGCATTCTG |
| 1 | GMTACGGTCACACTG | chrX:4685903-4687903 | 963 | 977 | - | 3.3571 | 7.12E-05 | 0.795 | ACAACGGTCACACGC |
| 1 | GMTACGGTCACACTG | chr2L:4458638-4460638 | 964 | 978 | - | 3.3571 | 7.12E-05 | 0.795 | TCGGGGGTCACACTG |
| 1 | GMTACGGTCACACTG | chrX:16358039-16360039 | 965 | 979 | - | 3.3571 | 7.12E-05 | 0.795 | GCCAGTGTCGCACTG |
| 1 | GMTACGGTCACACTG | chr3R:22740150-22742150 | 967 | 981 | - | 3.3571 | 7.12E-05 | 0.795 | ACCCTGGTCACACTG |
| 1 | GMTACGGTCACACTG | chr2L:18858248-18860248 | 968 | 982 | - | 3.3571 | 7.12E-05 | 0.795 | ACCTTGGTCACACTG |
| 1 | GMTACGGTCACACTG | chr2L:18858248-18860248 | 968 | 982 | - | 3.3571 | 7.12E-05 | 0.795 | ACCTTGGTCACACTG |
| 1 | GMTACGGTCACACTG | chrX:619282-621282 | 969 | 983 | - | 3.3571 | 7.12E-05 | 0.795 | GCGACAGCCACACTA |
| 1 | GMTACGGTCACACTG | chr3L:12529171-12531171 | 970 | 984 | - | 3.3571 | 7.12E-05 | 0.795 | ACGACGGTCACACCC |
| 1 | GMTACGGTCACACTG | chr3L:16785941-16787941 | 971 | 985 | - | 3.3571 | 7.12E-05 | 0.795 | TCTCTGGGCACACTG |
| 1 | GMTACGGTCACACTG | chrX:4251790-4253790 | 971 | 985 | - | 3.3571 | 7.12E-05 | 0.795 | GCTCAGGTCTCATTG |
| 1 | GMTACGGTCACACTG | chr3R:7912851-7914851 | 972 | 986 | - | 3.3571 | 7.12E-05 | 0.795 | CCAACCGTCGCACTG |
| 1 | GMTACGGTCACACTG | chr2R:14238703-14240703 | 972 | 986 | - | 3.3571 | 7.12E-05 | 0.795 | CCTTCGGCCACACTT |
| 1 | GMTACGGTCACACTG | chr2R:17671640-17673640 | 972 | 986 | - | 3.3571 | 7.12E-05 | 0.795 | TCAACGGTCACACCA |

| | | | | | | | | | |
|---|---|---|---|---|---|---|---|---|---|
| 1 | GMTACGGTCACACTG | chr3R:24786421-24788421 | 973 | 987 | - | 3.3571 | 7.12E-05 | 0.795 | ACTCTGGCCACACTG |
| 1 | GMTACGGTCACACTG | chr3L:12525731-12527731 | 973 | 987 | - | 3.3571 | 7.12E-05 | 0.795 | CAGCCGGTCACACCG |
| 1 | GMTACGGTCACACTG | chr2L:18731524-18733524 | 974 | 988 | - | 3.3571 | 7.12E-05 | 0.795 | GAGACGGTCTCAAAG |
| 1 | GMTACGGTCACACTG | chr3R:11242999-11244999 | 974 | 988 | - | 3.3571 | 7.12E-05 | 0.795 | GCTCTGGTCACACAT |
| 1 | GMTACGGTCACACTG | chr2R:14234449-14236449 | 975 | 989 | - | 3.3571 | 7.12E-05 | 0.795 | ACTCTGGTCACACAG |
| 1 | GMTACGGTCACACTG | chr3R:5223139-5225139 | 976 | 990 | - | 3.3571 | 7.12E-05 | 0.795 | GCTGGTGTCCCACTG |
| 1 | GMTACGGTCACACTG | chr2L:6970737-6972737 | 976 | 990 | - | 3.3571 | 7.12E-05 | 0.795 | GCGGGGGTCACACTT |
| 1 | GMTACGGTCACACTG | chr3L:1601674-1603674 | 977 | 991 | - | 3.3571 | 7.12E-05 | 0.795 | CCGGCGGTCACACTA |
| 1 | GMTACGGTCACACTG | chr2R:22378637-22380637 | 979 | 993 | - | 3.3571 | 7.12E-05 | 0.795 | GAGCCGCACACACTG |
| 1 | GMTACGGTCACACTG | chr3R:18182479-18184479 | 979 | 993 | - | 3.3571 | 7.12E-05 | 0.795 | GCGTTGGTCACACTC |
| 1 | GMTACGGTCACACTG | chr3R:24359144-24361144 | 979 | 993 | - | 3.3571 | 7.12E-05 | 0.795 | TCCGCGGTCACACTT |
| 1 | GMTACGGTCACACTG | chr3L:4133550-4135550 | 979 | 993 | - | 3.3571 | 7.12E-05 | 0.795 | GAGCTGGCCACACTG |
| 1 | GMTACGGTCACACTG | chr2R:9425439-9427439 | 980 | 994 | - | 3.3571 | 7.12E-05 | 0.795 | GATCCGGTAGCACCG |
| 1 | GMTACGGTCACACTG | chr2L:1728636-1730636 | 983 | 997 | - | 3.3571 | 7.12E-05 | 0.795 | GCTTCCAACACACTG |
| 1 | GMTACGGTCACACTG | chrX:15702770-15704770 | 984 | 998 | - | 3.3571 | 7.12E-05 | 0.795 | CCAAAGGTCACGCTG |
| 1 | GMTACGGTCACACTG | chr2L:3513656-3515656 | 984 | 998 | - | 3.3571 | 7.12E-05 | 0.795 | TCTGCTGCCACACTG |
| 1 | GMTACGGTCACACTG | chr3R:25894822-25896822 | 985 | 999 | - | 3.3571 | 7.12E-05 | 0.795 | GCTTCGGTTGAACTG |
| 1 | GMTACGGTCACACTG | chrX:7103491-7105491 | 985 | 999 | - | 3.3571 | 7.12E-05 | 0.795 | GCTCTGGTCACACCA |
| 1 | GMTACGGTCACACTG | chr3L:316084-318084 | 986 | 1000 | - | 3.3571 | 7.12E-05 | 0.795 | GCTTCGATTTCACTG |
| 1 | GMTACGGTCACACTG | chr3R:28811568-28813568 | 988 | 1002 | - | 3.3571 | 7.12E-05 | 0.795 | GCCACACCCACACTG |
| 1 | GMTACGGTCACACTG | chr3R:18874528-18876528 | 988 | 1002 | - | 3.3571 | 7.12E-05 | 0.795 | GATGCGAACTCACTG |
| 1 | GMTACGGTCACACTG | chr3R:24335922-24337922 | 989 | 1003 | - | 3.3571 | 7.12E-05 | 0.795 | ACAACCGTCGCACTG |
| 1 | GMTACGGTCACACTG | chr2L:2008859-2010859 | 989 | 1003 | - | 3.3571 | 7.12E-05 | 0.795 | GCAACTGTCAATCTG |
| 1 | GMTACGGTCACACTG | chr2L:16742100-16744100 | 991 | 1005 | - | 3.3571 | 7.12E-05 | 0.795 | GCTTTGGTCACTCTT |
| 1 | GMTACGGTCACACTG | chr3R:19093270-19095270 | 991 | 1005 | - | 3.3571 | 7.12E-05 | 0.795 | ACAACGGTCACTCTA |
| 1 | GMTACGGTCACACTG | chrX:12582193-12584193 | 991 | 1005 | - | 3.3571 | 7.12E-05 | 0.795 | GCCAAGGCCACACTC |
| 1 | GMTACGGTCACACTG | chrX:9102739-9104739 | 991 | 1005 | - | 3.3571 | 7.12E-05 | 0.795 | ACGCTGGTCACACTG |
| 1 | GMTACGGTCACACTG | chr3L:5497958-5499958 | 992 | 1006 | - | 3.3571 | 7.12E-05 | 0.795 | GCTACGGTCTCTTTA |
| 1 | GMTACGGTCACACTG | chr2R:15144519-15146519 | 993 | 1007 | - | 3.3571 | 7.12E-05 | 0.795 | ACAACGGTCACTCTA |
| 1 | GMTACGGTCACACTG | chr3R:19631718-19633718 | 993 | 1007 | - | 3.3571 | 7.12E-05 | 0.795 | GCTACGCGCACATGG |
| 1 | GMTACGGTCACACTG | chr3L:21342716-21344716 | 993 | 1007 | - | 3.3571 | 7.12E-05 | 0.795 | CCAACGGGCACACTT |
| 1 | GMTACGGTCACACTG | chr3R:4428335-4430335 | 994 | 1008 | - | 3.3571 | 7.12E-05 | 0.795 | CCCCTGGTCACACTG |
| 1 | GMTACGGTCACACTG | chr3L:21440363-21442363 | 994 | 1008 | - | 3.3571 | 7.12E-05 | 0.795 | TCAGCGGTCACACTT |

| | | | | | | | | | |
|---|---|---|---|---|---|---|---|---|---|
| 1 | GMTACGGTCACACTG | chr2R:14870071-14872071 | 994 | 1008 | - | 3.3571 | 7.12E-05 | 0.795 | TCGACGGCCACACTT |
| 1 | GMTACGGTCACACTG | chrX:1860790-1862790 | 995 | 1009 | - | 3.3571 | 7.12E-05 | 0.795 | GCGCTGGTCACACTA |
| 1 | GMTACGGTCACACTG | chr3L:21678755-21680755 | 995 | 1009 | - | 3.3571 | 7.12E-05 | 0.795 | GCGCTGGTCACACTA |
| 1 | GMTACGGTCACACTG | chrX:10626737-10628737 | 996 | 1010 | - | 3.3571 | 7.12E-05 | 0.795 | GCTGAGAACACACTG |
| 1 | GMTACGGTCACACTG | chrX:18649444-18651444 | 997 | 1011 | - | 3.3571 | 7.12E-05 | 0.795 | ACTCTGGTCACGCTG |
| 1 | GMTACGGTCACACTG | chr3R:12690468-12692468 | 997 | 1011 | - | 3.3571 | 7.12E-05 | 0.795 | GCAACGTTCCAACTG |
| 1 | GMTACGGTCACACTG | chr4:866025-868025 | 999 | 1013 | - | 3.3571 | 7.12E-05 | 0.795 | AAGACGGCCACGCTG |
| 1 | GMTACGGTCACACTG | chr3L:10322501-10324501 | 999 | 1013 | - | 3.3571 | 7.12E-05 | 0.795 | GACGCGGGCAAACTG |
| 1 | GMTACGGTCACACTG | chr3R:11876235-11878235 | 999 | 1013 | - | 3.3571 | 7.12E-05 | 0.795 | GCCAGGGGCAGACTG |
| 1 | GMTACGGTCACACTG | chr3L:3045812-3047812 | 1000 | 1014 | - | 3.3571 | 7.12E-05 | 0.795 | AACGCGGTCACCCTG |
| 1 | GMTACGGTCACACTG | chr2L:12108116-12110116 | 1001 | 1015 | - | 3.3571 | 7.12E-05 | 0.795 | ACTCTGGTCACACAG |
| 1 | GMTACGGTCACACTG | chr3L:14052079-14054079 | 1001 | 1015 | - | 3.3571 | 7.12E-05 | 0.795 | CCAACGGTCGCACTT |
| 1 | GMTACGGTCACACTG | chr2R:10781895-10783895 | 1001 | 1015 | - | 3.3571 | 7.12E-05 | 0.795 | TCAAAGGCCACACTG |
| 1 | GMTACGGTCACACTG | chr3L:16003736-16005736 | 1003 | 1017 | - | 3.3571 | 7.12E-05 | 0.795 | ACCGCGGTCACACTA |
| 1 | GMTACGGTCACACTG | chr3R:15856470-15858470 | 1003 | 1017 | - | 3.3571 | 7.12E-05 | 0.795 | GAAACGGCCATACCG |
| 1 | GMTACGGTCACACTG | chr2R:11678354-11680354 | 1004 | 1018 | - | 3.3571 | 7.12E-05 | 0.795 | GCTGTGGCAACACTG |
| 1 | GMTACGGTCACACTG | chr3R:29994231-29996231 | 1006 | 1020 | - | 3.3571 | 7.12E-05 | 0.795 | TCGCTGGTCACACTG |
| 1 | GMTACGGTCACACTG | chr2R:22476295-22478295 | 1007 | 1021 | - | 3.3571 | 7.12E-05 | 0.795 | GCTCCTGAAACACTG |
| 1 | GMTACGGTCACACTG | chr2R:15656066-15658066 | 1009 | 1023 | - | 3.3571 | 7.12E-05 | 0.795 | CATCCGGTCGCACCG |
| 1 | GMTACGGTCACACTG | chrX:1029556-1031556 | 1009 | 1023 | - | 3.3571 | 7.12E-05 | 0.795 | ACACTGGTCACACTG |
| 1 | GMTACGGTCACACTG | chr3R:31712867-31714867 | 1010 | 1024 | - | 3.3571 | 7.12E-05 | 0.795 | GCTTTGATCACACGG |
| 1 | GMTACGGTCACACTG | chr3R:5597566-5599566 | 1010 | 1024 | - | 3.3571 | 7.12E-05 | 0.795 | GCGAAGGTCACGCTA |
| 1 | GMTACGGTCACACTG | chr3R:22393104-22395104 | 1012 | 1026 | - | 3.3571 | 7.12E-05 | 0.795 | GCCGCGATCACACTC |
| 1 | GMTACGGTCACACTG | chr2L:11497948-11499948 | 1013 | 1027 | - | 3.3571 | 7.12E-05 | 0.795 | GCTACCGTCGATCTG |
| 1 | GMTACGGTCACACTG | chr3L:7639446-7641446 | 1016 | 1030 | - | 3.3571 | 7.12E-05 | 0.795 | GACGCGGTCACACAT |
| 1 | GMTACGGTCACACTG | chr2L:12720843-12722843 | 1017 | 1031 | - | 3.3571 | 7.12E-05 | 0.795 | CCGGCGTTCACACTG |
| 1 | GMTACGGTCACACTG | chr3L:3142457-3144457 | 1019 | 1033 | - | 3.3571 | 7.12E-05 | 0.795 | GCTGCGGTAACCCTA |
| 1 | GMTACGGTCACACTG | chr2L:9013068-9015068 | 1021 | 1035 | - | 3.3571 | 7.12E-05 | 0.795 | GCTCTGGCCACATTG |
| 1 | GMTACGGTCACACTG | chr3L:19440195-19442195 | 1022 | 1036 | - | 3.3571 | 7.12E-05 | 0.795 | GACACGGACACAGAG |
| 1 | GMTACGGTCACACTG | chr2L:4456674-4458674 | 1024 | 1038 | - | 3.3571 | 7.12E-05 | 0.795 | GCTACGCGAACTCTG |
| 1 | GMTACGGTCACACTG | chr2L:9914253-9916253 | 1024 | 1038 | - | 3.3571 | 7.12E-05 | 0.795 | GCGACTATCGCACTG |
| 1 | GMTACGGTCACACTG | chr2L:541580-543580 | 1025 | 1039 | - | 3.3571 | 7.12E-05 | 0.795 | TCCACGGCCACACTA |
| 1 | GMTACGGTCACACTG | chrX:21338676-21340676 | 1025 | 1039 | - | 3.3571 | 7.12E-05 | 0.795 | GAACTGGCCACACTG |

| | | | | | | | | | |
|---|---|---|---|---|---|---|---|---|---|
| 1 | GMTACGGTCACACTG | chr3L:19641816-19643816 | 1025 | 1039 | - | 3.3571 | 7.12E-05 | 0.795 | TCAACGGTCATACCG |
| 1 | GMTACGGTCACACTG | chr3R:22740091-22742091 | 1026 | 1040 | - | 3.3571 | 7.12E-05 | 0.795 | ACCCTGGTCACACTG |
| 1 | GMTACGGTCACACTG | chr2R:6834635-6836635 | 1029 | 1043 | - | 3.3571 | 7.12E-05 | 0.795 | GCTCCTGTCACTCTT |
| 1 | GMTACGGTCACACTG | chr3R:15634468-15636468 | 1031 | 1045 | - | 3.3571 | 7.12E-05 | 0.795 | GCCACTGGCATACTG |
| 1 | GMTACGGTCACACTG | chr3R:10041895-10043895 | 1031 | 1045 | - | 3.3571 | 7.12E-05 | 0.795 | GCCATGGTCACACCA |
| 1 | GMTACGGTCACACTG | chrX:11585228-11587228 | 1033 | 1047 | - | 3.3571 | 7.12E-05 | 0.795 | GCTAAGTTCTCGCTG |
| 1 | GMTACGGTCACACTG | chrX:5682240-5684240 | 1033 | 1047 | - | 3.3571 | 7.12E-05 | 0.795 | GCTAAGGCTACTCTG |
| 1 | GMTACGGTCACACTG | chr2L:7222081-7224081 | 1034 | 1048 | - | 3.3571 | 7.12E-05 | 0.795 | GCCACGGTCTTACTA |
| 1 | GMTACGGTCACACTG | chr2L:18731464-18733464 | 1034 | 1048 | - | 3.3571 | 7.12E-05 | 0.795 | GAGACGGTCTCAAAG |
| 1 | GMTACGGTCACACTG | chr2R:16123981-16125981 | 1035 | 1049 | - | 3.3571 | 7.12E-05 | 0.795 | GCGACGCGAACACTG |
| 1 | GMTACGGTCACACTG | chr3R:9996163-9998163 | 1035 | 1049 | - | 3.3571 | 7.12E-05 | 0.795 | GCTGCTGCCAAACTG |
| 1 | GMTACGGTCACACTG | chr2R:10728203-10730203 | 1038 | 1052 | - | 3.3571 | 7.12E-05 | 0.795 | GCTACGCCCACACAC |
| 1 | GMTACGGTCACACTG | chr3L:8575808-8577808 | 1039 | 1053 | - | 3.3571 | 7.12E-05 | 0.795 | GCTCCTGGCAGACTG |
| 1 | GMTACGGTCACACTG | chr2L:22042877-22044877 | 1040 | 1054 | - | 3.3571 | 7.12E-05 | 0.795 | TCTTCGGTAACACCG |
| 1 | GMTACGGTCACACTG | chr2R:18386895-18388895 | 1041 | 1055 | - | 3.3571 | 7.12E-05 | 0.795 | GCGACGTTCTCAATG |
| 1 | GMTACGGTCACACTG | chr3L:4654405-4656405 | 1043 | 1057 | - | 3.3571 | 7.12E-05 | 0.795 | GCGACCGACACACTT |
| 1 | GMTACGGTCACACTG | chr2R:13825371-13827371 | 1044 | 1058 | - | 3.3571 | 7.12E-05 | 0.795 | GATGCGGTCACGCGA |
| 1 | GMTACGGTCACACTG | chrX:8239684-8241684 | 1045 | 1059 | - | 3.3571 | 7.12E-05 | 0.795 | GCTCCCTTCTCACTG |
| 1 | GMTACGGTCACACTG | chr2R:17800707-17802707 | 1046 | 1060 | - | 3.3571 | 7.12E-05 | 0.795 | GACACGGGCAAACCG |
| 1 | GMTACGGTCACACTG | chrX:12900114-12902114 | 1049 | 1063 | - | 3.3571 | 7.12E-05 | 0.795 | GCCATGGTGCCACTG |
| 1 | GMTACGGTCACACTG | chr3R:20592139-20594139 | 1051 | 1065 | - | 3.3571 | 7.12E-05 | 0.795 | CACACGGTCTCACGG |
| 1 | GMTACGGTCACACTG | chr3R:12691597-12693597 | 1051 | 1065 | - | 3.3571 | 7.12E-05 | 0.795 | GCTTTGATCACACCG |
| 1 | GMTACGGTCACACTG | chr3L:21683042-21685042 | 1056 | 1070 | - | 3.3571 | 7.12E-05 | 0.795 | GCTTAAGTCACCCTG |
| 1 | GMTACGGTCACACTG | chrX:619195-621195 | 1056 | 1070 | - | 3.3571 | 7.12E-05 | 0.795 | GCGACAGCCACACTA |
| 1 | GMTACGGTCACACTG | chr3L:8305980-8307980 | 1063 | 1077 | - | 3.3571 | 7.12E-05 | 0.795 | GCTAAGTGCACTCTG |
| 1 | GMTACGGTCACACTG | chr3R:18401716-18403716 | 1063 | 1077 | - | 3.3571 | 7.12E-05 | 0.795 | GCCACAGTCCGACTG |
| 1 | GMTACGGTCACACTG | chr3R:24840163-24842163 | 1068 | 1082 | - | 3.3571 | 7.12E-05 | 0.795 | GCTTCAGTCTCACTT |
| 1 | GMTACGGTCACACTG | chr2R:13552240-13554240 | 1074 | 1088 | - | 3.3571 | 7.12E-05 | 0.795 | TCTCCGATCGCACTG |
| 1 | GMTACGGTCACACTG | chr2R:9711321-9713321 | 1075 | 1089 | - | 3.3571 | 7.12E-05 | 0.795 | GCTATGAGCGCACTG |
| 1 | GMTACGGTCACACTG | chrX:15378064-15380064 | 1077 | 1091 | - | 3.3571 | 7.12E-05 | 0.795 | GACACGCCCACGCTG |
| 1 | GMTACGGTCACACTG | chr2L:856403-858403 | 1077 | 1091 | - | 3.3571 | 7.12E-05 | 0.795 | GCCACCTCCACACTG |
| 1 | GMTACGGTCACACTG | chr2R:18804595-18806595 | 1077 | 1091 | - | 3.3571 | 7.12E-05 | 0.795 | TCTGCAGTCGCACTG |
| 1 | GMTACGGTCACACTG | chr2L:6556502-6558502 | 1079 | 1093 | - | 3.3571 | 7.12E-05 | 0.795 | ACAACGGTCACCCTA |

| | | | | | | | | | |
|---|---|---|---|---|---|---|---|---|---|
| 1 | GMTACGGTCACACTG | chrX:11491998-11493998 | 1083 | 1097 | - | 3.3571 | 7.12E-05 | 0.795 | TCTCTGGTCACTCTG |
| 1 | GMTACGGTCACACTG | chr3R:6262976-6264976 | 1087 | 1101 | - | 3.3571 | 7.12E-05 | 0.795 | TCCACGAACACACTG |
| 1 | GMTACGGTCACACTG | chr3R:23932912-23934912 | 1089 | 1103 | - | 3.3571 | 7.12E-05 | 0.795 | GACACGGTCAACCGG |
| 1 | GMTACGGTCACACTG | chr3R:5632627-5634627 | 1091 | 1105 | - | 3.3571 | 7.12E-05 | 0.795 | GCTCCGGTCACGATT |
| 1 | GMTACGGTCACACTG | chr3R:18250115-18252115 | 1093 | 1107 | - | 3.3571 | 7.12E-05 | 0.795 | GATACGCCCGCACCG |
| 1 | GMTACGGTCACACTG | chr3R:4173632-4175632 | 1094 | 1108 | - | 3.3571 | 7.12E-05 | 0.795 | GCTACGTTTCCCCTG |
| 1 | GMTACGGTCACACTG | chr2L:4349999-4351999 | 1097 | 1111 | - | 3.3571 | 7.12E-05 | 0.795 | GCGAGGGGCACAATG |
| 1 | GMTACGGTCACACTG | chr3L:5562774-5564774 | 1098 | 1112 | - | 3.3571 | 7.12E-05 | 0.795 | GCTTCCGTCTCACTC |
| 1 | GMTACGGTCACACTG | chr3L:21683000-21685000 | 1098 | 1112 | - | 3.3571 | 7.12E-05 | 0.795 | GCTTAAGTCACCCTG |
| 1 | GMTACGGTCACACTG | chr2L:18739358-18741358 | 1111 | 1125 | - | 3.3571 | 7.12E-05 | 0.795 | TCTTCGGGCACACTC |
| 1 | GMTACGGTCACACTG | chr3R:30526037-30528037 | 1114 | 1128 | - | 3.3571 | 7.12E-05 | 0.795 | GCTCAGCTCACTCTG |
| 1 | GMTACGGTCACACTG | chr3R:18949196-18951196 | 1115 | 1129 | - | 3.3571 | 7.12E-05 | 0.795 | GCTCCTGTCACACGA |
| 1 | GMTACGGTCACACTG | chr3L:4211915-4213915 | 1116 | 1130 | - | 3.3571 | 7.12E-05 | 0.795 | GCAATTGTCACTCTG |
| 1 | GMTACGGTCACACTG | chr3L:8324859-8326859 | 1127 | 1141 | - | 3.3571 | 7.12E-05 | 0.795 | CAAACGGACACACGG |
| 1 | GMTACGGTCACACTG | chr3R:23956305-23958305 | 1127 | 1141 | - | 3.3571 | 7.12E-05 | 0.795 | GCTAAGGCGGCACTG |
| 1 | GMTACGGTCACACTG | chrX:9584675-9586675 | 1128 | 1142 | - | 3.3571 | 7.12E-05 | 0.795 | GCCGCGCTCACACTC |
| 1 | GMTACGGTCACACTG | chr2L:18676221-18678221 | 1130 | 1144 | - | 3.3571 | 7.12E-05 | 0.795 | GATTCGGTCCCACCT |
| 1 | GMTACGGTCACACTG | chr3R:23930639-23932639 | 1130 | 1144 | - | 3.3571 | 7.12E-05 | 0.795 | GCTGTTGTCCCACTG |
| 1 | GMTACGGTCACACTG | chr2L:20757144-20759144 | 1132 | 1146 | - | 3.3571 | 7.12E-05 | 0.795 | GCAACGCAAACACTG |
| 1 | GMTACGGTCACACTG | chrX:11892943-11894943 | 1133 | 1147 | - | 3.3571 | 7.12E-05 | 0.795 | GCGATCGTCGCACTG |
| 1 | GMTACGGTCACACTG | chrX:12983524-12985524 | 1134 | 1148 | - | 3.3571 | 7.12E-05 | 0.795 | GCTTCGGGAAAACTG |
| 1 | GMTACGGTCACACTG | chr2R:24869108-24871108 | 1134 | 1148 | - | 3.3571 | 7.12E-05 | 0.795 | CCGAAGGACACACTG |
| 1 | GMTACGGTCACACTG | chr3R:24149203-24151203 | 1138 | 1152 | - | 3.3571 | 7.12E-05 | 0.795 | TCTCCGGTCATACAG |
| 1 | GMTACGGTCACACTG | chr2L:3508099-3510099 | 1147 | 1161 | - | 3.3571 | 7.12E-05 | 0.795 | GCTACGGTTGCGCTA |
| 1 | GMTACGGTCACACTG | chrX:8177549-8179549 | 1149 | 1163 | - | 3.3571 | 7.12E-05 | 0.795 | GATGCTGGCACACAG |
| 1 | GMTACGGTCACACTG | chr2R:10420706-10422706 | 1151 | 1165 | - | 3.3571 | 7.12E-05 | 0.795 | GCAACGCACAGACTG |
| 1 | GMTACGGTCACACTG | chr2L:15127559-15129559 | 1154 | 1168 | - | 3.3571 | 7.12E-05 | 0.795 | GCTGCCGTCACGGTG |
| 1 | GMTACGGTCACACTG | chr3R:15274522-15276522 | 1164 | 1178 | - | 3.3571 | 7.12E-05 | 0.795 | GCTGCGGGCACAATT |
| 1 | GMTACGGTCACACTG | chr3R:27259050-27261050 | 1167 | 1181 | - | 3.3571 | 7.12E-05 | 0.795 | GCTCCCCTCACGCTG |
| 1 | GMTACGGTCACACTG | chr2R:9131480-9133480 | 1169 | 1183 | - | 3.3571 | 7.12E-05 | 0.795 | GCTTCGGTTACACGC |
| 1 | GMTACGGTCACACTG | chr2L:1203215-1205215 | 1173 | 1187 | - | 3.3571 | 7.12E-05 | 0.795 | GACGCGGTACCACTG |
| 1 | GMTACGGTCACACTG | chr2R:10311023-10313023 | 1173 | 1187 | - | 3.3571 | 7.12E-05 | 0.795 | TCGACGGTCACGATG |
| 1 | GMTACGGTCACACTG | chr3L:5562691-5564691 | 1181 | 1195 | - | 3.3571 | 7.12E-05 | 0.795 | GCTTCCGTCTCACTC |

| | | | | | | | | | |
|---|---|---|---|---|---|---|---|---|---|
| 1 | GMTACGGTCACACTG | chr2R:15541661-15543661 | 1181 | 1195 | - | 3.3571 | 7.12E-05 | 0.795 | GATATGGACCCACCG |
| 1 | GMTACGGTCACACTG | chr2R:22669587-22671587 | 1182 | 1196 | - | 3.3571 | 7.12E-05 | 0.795 | GCTGCGGCAACATTG |
| 1 | GMTACGGTCACACTG | chr3R:31038374-31040374 | 1183 | 1197 | - | 3.3571 | 7.12E-05 | 0.795 | GAGACGGCCAGACAG |
| 1 | GMTACGGTCACACTG | chrX:4770588-4772588 | 1188 | 1202 | - | 3.3571 | 7.12E-05 | 0.795 | GCCACGATCGCAATG |
| 1 | GMTACGGTCACACTG | chr2R:16293130-16295130 | 1189 | 1203 | - | 3.3571 | 7.12E-05 | 0.795 | GCAGTGTTCACACTG |
| 1 | GMTACGGTCACACTG | chrX:16390438-16392438 | 1190 | 1204 | - | 3.3571 | 7.12E-05 | 0.795 | GCCACGTTCCAACTG |
| 1 | GMTACGGTCACACTG | chr3L:2022391-2024391 | 1192 | 1206 | - | 3.3571 | 7.12E-05 | 0.795 | GCAACATTCACCCTG |
| 1 | GMTACGGTCACACTG | chr3L:21122753-21124753 | 1193 | 1207 | - | 3.3571 | 7.12E-05 | 0.795 | GATTCGGCGACTCTG |
| 1 | GMTACGGTCACACTG | chrX:12407861-12409861 | 1195 | 1209 | - | 3.3571 | 7.12E-05 | 0.795 | GAGCTGGCCACACTG |
| 1 | GMTACGGTCACACTG | chr3L:5949658-5951658 | 1196 | 1210 | - | 3.3571 | 7.12E-05 | 0.795 | TAAACGGGCCCACTG |
| 1 | GMTACGGTCACACTG | chr2R:22839739-22841739 | 1200 | 1214 | - | 3.3571 | 7.12E-05 | 0.795 | ACAATGGCCACACTG |
| 1 | GMTACGGTCACACTG | chr2L:6490061-6492061 | 1200 | 1214 | - | 3.3571 | 7.12E-05 | 0.795 | GCAGCGGTTACACTT |
| 1 | GMTACGGTCACACTG | chrX:9088513-9090513 | 1210 | 1224 | - | 3.3571 | 7.12E-05 | 0.795 | GCTTCGTTCACACAC |
| 1 | GMTACGGTCACACTG | chr3R:23930558-23932558 | 1211 | 1225 | - | 3.3571 | 7.12E-05 | 0.795 | GCTGTTGTCCCACTG |
| 1 | GMTACGGTCACACTG | chr3L:21514887-21516887 | 1214 | 1228 | - | 3.3571 | 7.12E-05 | 0.795 | GCTCTGGCAACACTG |
| 1 | GMTACGGTCACACTG | chrX:6308212-6310212 | 1222 | 1236 | - | 3.3571 | 7.12E-05 | 0.795 | GAGTCGGTGACTCTG |
| 1 | GMTACGGTCACACTG | chr3L:202730-204730 | 1222 | 1236 | - | 3.3571 | 7.12E-05 | 0.795 | GCAATGGTCGCACTT |
| 1 | GMTACGGTCACACTG | chrX:1030496-1032496 | 1226 | 1240 | - | 3.3571 | 7.12E-05 | 0.795 | GCCACGATCCCAGTG |
| 1 | GMTACGGTCACACTG | chr2L:19641601-19643601 | 1233 | 1247 | - | 3.3571 | 7.12E-05 | 0.795 | GCCACAGTCAGTCTG |
| 1 | GMTACGGTCACACTG | chr2R:8204706-8206706 | 1238 | 1252 | - | 3.3571 | 7.12E-05 | 0.795 | GACTCGGGAACACTG |
| 1 | GMTACGGTCACACTG | chr2R:11777526-11779526 | 1239 | 1253 | - | 3.3571 | 7.12E-05 | 0.795 | AATCCGGACACACAG |
| 1 | GMTACGGTCACACTG | chr2L:8395649-8397649 | 1239 | 1253 | - | 3.3571 | 7.12E-05 | 0.795 | GACAAGGTCGCTCTG |
| 1 | GMTACGGTCACACTG | chr3R:17378944-17380944 | 1242 | 1256 | - | 3.3571 | 7.12E-05 | 0.795 | GCTACGCGCACTTTG |
| 1 | GMTACGGTCACACTG | chr2L:4255085-4257085 | 1244 | 1258 | - | 3.3571 | 7.12E-05 | 0.795 | GCAACTGCAACACTG |
| 1 | GMTACGGTCACACTG | chrX:6539771-6541771 | 1246 | 1260 | - | 3.3571 | 7.12E-05 | 0.795 | GCTGCTGTCACACGC |
| 1 | GMTACGGTCACACTG | chrX:5613467-5615467 | 1247 | 1261 | - | 3.3571 | 7.12E-05 | 0.795 | GATACGTCCACGCCG |
| 1 | GMTACGGTCACACTG | chrX:6823496-6825496 | 1253 | 1267 | - | 3.3571 | 7.12E-05 | 0.795 | GCTGCTGTCACCGTG |
| 1 | GMTACGGTCACACTG | chr2L:13386571-13388571 | 1255 | 1269 | - | 3.3571 | 7.12E-05 | 0.795 | GATGCGGTCAGGCAG |
| 1 | GMTACGGTCACACTG | chr3R:21645301-21647301 | 1258 | 1272 | - | 3.3571 | 7.12E-05 | 0.795 | GATCTGGTCCCACGG |
| 1 | GMTACGGTCACACTG | chr3L:11727348-11729348 | 1260 | 1274 | - | 3.3571 | 7.12E-05 | 0.795 | GACACGGAAGCACTG |
| 1 | GMTACGGTCACACTG | chr2L:13641821-13643821 | 1262 | 1276 | - | 3.3571 | 7.12E-05 | 0.795 | GATTCAGGCACACAG |
| 1 | GMTACGGTCACACTG | chr3L:19727994-19729994 | 1263 | 1277 | - | 3.3571 | 7.12E-05 | 0.795 | GCCACTGTCACATAG |
| 1 | GMTACGGTCACACTG | chr3L:13456238-13458238 | 1263 | 1277 | - | 3.3571 | 7.12E-05 | 0.795 | GCTACGCTGCCGCTG |

| | | | | | | | | | |
|---|---|---|---|---|---|---|---|---|---|
| 1 | GMTACGGTCACACTG | chr3L:19759409-19761409 | 1263 | 1277 | - | 3.3571 | 7.12E-05 | 0.795 | GCCACTGTCACATAG |
| 1 | GMTACGGTCACACTG | chrX:13190883-13192883 | 1264 | 1278 | - | 3.3571 | 7.12E-05 | 0.795 | TCTACGGTACCACAG |
| 1 | GMTACGGTCACACTG | chr2R:14165982-14167982 | 1265 | 1279 | - | 3.3571 | 7.12E-05 | 0.795 | GCCACAGTCACAAAG |
| 1 | GMTACGGTCACACTG | chr3R:12424917-12426917 | 1272 | 1286 | - | 3.3571 | 7.12E-05 | 0.795 | GCTAGTGTCCCCCTG |
| 1 | GMTACGGTCACACTG | chr3R:24364554-24366554 | 1273 | 1287 | - | 3.3571 | 7.12E-05 | 0.795 | GCTGTGGTCCAACTG |
| 1 | GMTACGGTCACACTG | chr3R:6300804-6302804 | 1274 | 1288 | - | 3.3571 | 7.12E-05 | 0.795 | GCTACGGCCCGACTC |
| 1 | GMTACGGTCACACTG | chr3L:5562774-5564774 | 1281 | 1295 | - | 3.3571 | 7.12E-05 | 0.795 | ACCATGGACACACTG |
| 1 | GMTACGGTCACACTG | chr2R:18397519-18399519 | 1281 | 1295 | - | 3.3571 | 7.12E-05 | 0.795 | ACTTCGGTAACACCG |
| 1 | GMTACGGTCACACTG | chr2R:16737038-16739038 | 1282 | 1296 | - | 3.3571 | 7.12E-05 | 0.795 | GCTACGCATGCACTG |
| 1 | GMTACGGTCACACTG | chr2L:13397664-13399664 | 1284 | 1298 | - | 3.3571 | 7.12E-05 | 0.795 | GCAACAGGCACACTA |
| 1 | GMTACGGTCACACTG | chr2L:13202762-13204762 | 1284 | 1298 | - | 3.3571 | 7.12E-05 | 0.795 | GCGACGTTCACGCTT |
| 1 | GMTACGGTCACACTG | chr4:625126-627126 | 1286 | 1300 | - | 3.3571 | 7.12E-05 | 0.795 | GCCACGCCCACACTT |
| 1 | GMTACGGTCACACTG | chrX:21478949-21480949 | 1289 | 1303 | - | 3.3571 | 7.12E-05 | 0.795 | GAAAAGGACACTCTG |
| 1 | GMTACGGTCACACTG | chr2L:20756985-20758985 | 1291 | 1305 | - | 3.3571 | 7.12E-05 | 0.795 | GCAACGCAAACACTG |
| 1 | GMTACGGTCACACTG | chr2L:4700485-4702485 | 1293 | 1307 | - | 3.3571 | 7.12E-05 | 0.795 | GCTGCGGCAACAATG |
| 1 | GMTACGGTCACACTG | chr3R:5392922-5394922 | 1295 | 1309 | - | 3.3571 | 7.12E-05 | 0.795 | TCTACGGGCAAACCG |
| 1 | GMTACGGTCACACTG | chr2R:14590203-14592203 | 1296 | 1310 | - | 3.3571 | 7.12E-05 | 0.795 | GAGTCGGTGACTCTG |
| 1 | GMTACGGTCACACTG | chrX:5432439-5434439 | 1296 | 1310 | - | 3.3571 | 7.12E-05 | 0.795 | GCGCCTGTCACATTG |
| 1 | GMTACGGTCACACTG | chr2L:16351934-16353934 | 1298 | 1312 | - | 3.3571 | 7.12E-05 | 0.795 | GCTGCACTCGCACTG |
| 1 | GMTACGGTCACACTG | chr2R:14854429-14856429 | 1299 | 1313 | - | 3.3571 | 7.12E-05 | 0.795 | GCCTTGCTCACACTG |
| 1 | GMTACGGTCACACTG | chrX:7346506-7348506 | 1300 | 1314 | - | 3.3571 | 7.12E-05 | 0.795 | GCTACGGTAACCCAC |
| 1 | GMTACGGTCACACTG | chr2R:12945347-12947347 | 1301 | 1315 | - | 3.3571 | 7.12E-05 | 0.795 | TCTATGGACACGCTG |
| 1 | GMTACGGTCACACTG | chr3L:5233483-5235483 | 1304 | 1318 | - | 3.3571 | 7.12E-05 | 0.795 | GCTCCGGCCACAGTT |
| 1 | GMTACGGTCACACTG | chr2R:7662786-7664786 | 1304 | 1318 | - | 3.3571 | 7.12E-05 | 0.795 | GCACCGGTGACAATG |
| 1 | GMTACGGTCACACTG | chr2L:14358252-14360252 | 1307 | 1321 | - | 3.3571 | 7.12E-05 | 0.795 | CCGACGGTGACCCTG |
| 1 | GMTACGGTCACACTG | chr2L:13922553-13924553 | 1307 | 1321 | - | 3.3571 | 7.12E-05 | 0.795 | GAGACGGTGGCGCTG |
| 1 | GMTACGGTCACACTG | chr2L:2008538-2010538 | 1310 | 1324 | - | 3.3571 | 7.12E-05 | 0.795 | GCAACTGTCAATCTG |
| 1 | GMTACGGTCACACTG | chr2L:5345200-5347200 | 1322 | 1336 | - | 3.3571 | 7.12E-05 | 0.795 | GCAACAGACACACTA |
| 1 | GMTACGGTCACACTG | chr2L:10422291-10424291 | 1327 | 1341 | - | 3.3571 | 7.12E-05 | 0.795 | GCTGCGTTCACATCG |
| 1 | GMTACGGTCACACTG | chr2L:8476349-8478349 | 1333 | 1347 | - | 3.3571 | 7.12E-05 | 0.795 | GCTTCGTTCGCACTT |
| 1 | GMTACGGTCACACTG | chr3L:7362598-7364598 | 1335 | 1349 | - | 3.3571 | 7.12E-05 | 0.795 | GCTCAGATCCCACTG |
| 1 | GMTACGGTCACACTG | chr3R:27006055-27008055 | 1340 | 1354 | - | 3.3571 | 7.12E-05 | 0.795 | GACACAGACACTCTG |
| 1 | GMTACGGTCACACTG | chr2L:2180962-2182962 | 1340 | 1354 | - | 3.3571 | 7.12E-05 | 0.795 | GACACGCACACTCTG |

| | | | | | | | | | |
|---|---|---|---|---|---|---|---|---|---|
| 1 | GMTACGGTCACACTG | chr2R:22654635-22656635 | 1346 | 1360 | - | 3.3571 | 7.12E-05 | 0.795 | ACCACTGTCACGCTG |
| 1 | GMTACGGTCACACTG | chr2L:8383455-8385455 | 1347 | 1361 | - | 3.3571 | 7.12E-05 | 0.795 | GCGAAGGTCACAACG |
| 1 | GMTACGGTCACACTG | chr2L:5558342-5560342 | 1350 | 1364 | - | 3.3571 | 7.12E-05 | 0.795 | TCGACGGTCACATAG |
| 1 | GMTACGGTCACACTG | chr2R:16582369-16584369 | 1352 | 1366 | - | 3.3571 | 7.12E-05 | 0.795 | GCAACTGGCACATTG |
| 1 | GMTACGGTCACACTG | chr2L:18675998-18677998 | 1353 | 1367 | - | 3.3571 | 7.12E-05 | 0.795 | GATTCGGTCCCACCT |
| 1 | GMTACGGTCACACTG | chr3R:8545910-8547910 | 1355 | 1369 | - | 3.3571 | 7.12E-05 | 0.795 | GCTCCTGTCACGTTG |
| 1 | GMTACGGTCACACTG | chr2L:18739110-18741110 | 1359 | 1373 | - | 3.3571 | 7.12E-05 | 0.795 | TCTTCGGGCACACTC |
| 1 | GMTACGGTCACACTG | chr2L:16836258-16838258 | 1360 | 1374 | - | 3.3571 | 7.12E-05 | 0.795 | GCTGTGGCCACATTG |
| 1 | GMTACGGTCACACTG | chr3R:23890060-23892060 | 1361 | 1375 | - | 3.3571 | 7.12E-05 | 0.795 | GCCACGCCCACACTT |
| 1 | GMTACGGTCACACTG | chr2L:10363471-10365471 | 1363 | 1377 | - | 3.3571 | 7.12E-05 | 0.795 | GCACTGGTGACACTG |
| 1 | GMTACGGTCACACTG | chr3L:5562691-5564691 | 1364 | 1378 | - | 3.3571 | 7.12E-05 | 0.795 | ACCATGGACACACTG |
| 1 | GMTACGGTCACACTG | chr3R:19876737-19878737 | 1365 | 1379 | - | 3.3571 | 7.12E-05 | 0.795 | GCTGCGTTTCCACTG |
| 1 | GMTACGGTCACACTG | chr2L:6741502-6743502 | 1365 | 1379 | - | 3.3571 | 7.12E-05 | 0.795 | GCTGCGATCGCACTT |
| 1 | GMTACGGTCACACTG | chr2R:15498632-15500632 | 1368 | 1382 | - | 3.3571 | 7.12E-05 | 0.795 | GATTCGGGCTCACTA |
| 1 | GMTACGGTCACACTG | chr3R:4418578-4420578 | 1370 | 1384 | - | 3.3571 | 7.12E-05 | 0.795 | GATCCGGACACACGA |
| 1 | GMTACGGTCACACTG | chr2R:7622242-7624242 | 1375 | 1389 | - | 3.3571 | 7.12E-05 | 0.795 | GCTGCTGTCACTTTG |
| 1 | GMTACGGTCACACTG | chr2R:23443137-23445137 | 1375 | 1389 | - | 3.3571 | 7.12E-05 | 0.795 | GAGTGGGGCACACTG |
| 1 | GMTACGGTCACACTG | chr3R:9416940-9418940 | 1380 | 1394 | - | 3.3571 | 7.12E-05 | 0.795 | GACACGCGCACACAG |
| 1 | GMTACGGTCACACTG | chr3R:6753194-6755194 | 1381 | 1395 | - | 3.3571 | 7.12E-05 | 0.795 | GCTTCGGTTTCACTT |
| 1 | GMTACGGTCACACTG | chr4:625031-627031 | 1381 | 1395 | - | 3.3571 | 7.12E-05 | 0.795 | GCCACGCCCACACTT |
| 1 | GMTACGGTCACACTG | chr3R:21748853-21750853 | 1382 | 1396 | - | 3.3571 | 7.12E-05 | 0.795 | GCCACGGTACCATTG |
| 1 | GMTACGGTCACACTG | chrX:17906261-17908261 | 1382 | 1396 | - | 3.3571 | 7.12E-05 | 0.795 | GCAATGGTCACACAT |
| 1 | GMTACGGTCACACTG | chr2R:13554949-13556949 | 1383 | 1397 | - | 3.3571 | 7.12E-05 | 0.795 | TCTATGGTCTCGCTG |
| 1 | GMTACGGTCACACTG | chr2L:5783031-5785031 | 1385 | 1399 | - | 3.3571 | 7.12E-05 | 0.795 | TAGACGGTCCCCCTG |
| 1 | GMTACGGTCACACTG | chr2L:11112061-11114061 | 1386 | 1400 | - | 3.3571 | 7.12E-05 | 0.795 | TAAACGGGCACGCTG |
| 1 | GMTACGGTCACACTG | chr2L:3451466-3453466 | 1391 | 1405 | - | 3.3571 | 7.12E-05 | 0.795 | GCTCCCGACACAGTG |
| 1 | GMTACGGTCACACTG | chr3L:18105579-18107579 | 1402 | 1416 | - | 3.3571 | 7.12E-05 | 0.795 | GAAACTGACGCACTG |
| 1 | GMTACGGTCACACTG | chr3L:20225381-20227381 | 1403 | 1417 | - | 3.3571 | 7.12E-05 | 0.795 | GCCACCGGGACACTG |
| 1 | GMTACGGTCACACTG | chr2R:12440320-12442320 | 1405 | 1419 | - | 3.3571 | 7.12E-05 | 0.795 | GCAACTGTCACACAT |
| 1 | GMTACGGTCACACTG | chr2R:12424483-12426483 | 1406 | 1420 | - | 3.3571 | 7.12E-05 | 0.795 | GCTCCGGTGACATGG |
| 1 | GMTACGGTCACACTG | chrX:1872853-1874853 | 1407 | 1421 | - | 3.3571 | 7.12E-05 | 0.795 | GATACCGTCTCCCCG |
| 1 | GMTACGGTCACACTG | chr2R:18397393-18399393 | 1407 | 1421 | - | 3.3571 | 7.12E-05 | 0.795 | ACTTCGGTAACACCG |
| 1 | GMTACGGTCACACTG | chr2L:6254859-6256859 | 1409 | 1423 | - | 3.3571 | 7.12E-05 | 0.795 | GACACGTCCACACCG |

| | | | | | | | | | |
|---|---|---|---|---|---|---|---|---|---|
| 1 | GMTACGGTCACACTG | chrX:17790010-17792010 | 1409 | 1423 | - | 3.3571 | 7.12E-05 | 0.795 | GCGTGGGTGACACTG |
| 1 | GMTACGGTCACACTG | chr3R:9995789-9997789 | 1409 | 1423 | - | 3.3571 | 7.12E-05 | 0.795 | GCTGCTGCCAAACTG |
| 1 | GMTACGGTCACACTG | chrX:1408069-1410069 | 1411 | 1425 | - | 3.3571 | 7.12E-05 | 0.795 | TCTCCTGTCGCACTG |
| 1 | GMTACGGTCACACTG | chr3R:17910376-17912376 | 1415 | 1429 | - | 3.3571 | 7.12E-05 | 0.795 | GCCACGTTGACGCTG |
| 1 | GMTACGGTCACACTG | chr3R:4161235-4163235 | 1416 | 1430 | - | 3.3571 | 7.12E-05 | 0.795 | GCTACGCACTCACTT |
| 1 | GMTACGGTCACACTG | chr3L:8533977-8535977 | 1416 | 1430 | - | 3.3571 | 7.12E-05 | 0.795 | GCCACGCGCACACTT |
| 1 | GMTACGGTCACACTG | chr2R:12143552-12145552 | 1416 | 1430 | - | 3.3571 | 7.12E-05 | 0.795 | GCCACGGACAGACTT |
| 1 | GMTACGGTCACACTG | chr2L:6489843-6491843 | 1418 | 1432 | - | 3.3571 | 7.12E-05 | 0.795 | GCAGCGGTTACACTT |
| 1 | GMTACGGTCACACTG | chr3R:5597157-5599157 | 1419 | 1433 | - | 3.3571 | 7.12E-05 | 0.795 | GCGAAGGTCACGCTA |
| 1 | GMTACGGTCACACTG | chrX:16421265-16423265 | 1419 | 1433 | - | 3.3571 | 7.12E-05 | 0.795 | TACCCGGTCACTCTG |
| 1 | GMTACGGTCACACTG | chr2L:7404227-7406227 | 1419 | 1433 | - | 3.3571 | 7.12E-05 | 0.795 | GCATAGGTCACATTG |
| 1 | GMTACGGTCACACTG | chr2R:12494672-12496672 | 1420 | 1434 | - | 3.3571 | 7.12E-05 | 0.795 | GATCCGGAGCCACTG |
| 1 | GMTACGGTCACACTG | chr3R:25969080-25971080 | 1422 | 1436 | - | 3.3571 | 7.12E-05 | 0.795 | GCTAAGGACACGCTC |
| 1 | GMTACGGTCACACTG | chr3L:8807544-8809544 | 1423 | 1437 | - | 3.3571 | 7.12E-05 | 0.795 | ACTACCGTCACCCGG |
| 1 | GMTACGGTCACACTG | chr3L:14048892-14050892 | 1425 | 1439 | - | 3.3571 | 7.12E-05 | 0.795 | TCTCCGGTGACACGG |
| 1 | GMTACGGTCACACTG | chr3R:6764485-6766485 | 1425 | 1439 | - | 3.3571 | 7.12E-05 | 0.795 | GCTCCGGCCATAATG |
| 1 | GMTACGGTCACACTG | chr3L:13060759-13062759 | 1426 | 1440 | - | 3.3571 | 7.12E-05 | 0.795 | GCTCCGGTAACACCC |
| 1 | GMTACGGTCACACTG | chr3R:23357812-23359812 | 1428 | 1442 | - | 3.3571 | 7.12E-05 | 0.795 | GAGGGGGTCACACGG |
| 1 | GMTACGGTCACACTG | chr3L:2228646-2230646 | 1429 | 1443 | - | 3.3571 | 7.12E-05 | 0.795 | GATGCTGTCGCTCTG |
| 1 | GMTACGGTCACACTG | chr2R:10884533-10886533 | 1430 | 1444 | - | 3.3571 | 7.12E-05 | 0.795 | GCGAAGGTGACACCG |
| 1 | GMTACGGTCACACTG | chr3L:14704464-14706464 | 1434 | 1448 | - | 3.3571 | 7.12E-05 | 0.795 | ACTACGGTCATTCAG |
| 1 | GMTACGGTCACACTG | chr2L:10894344-10896344 | 1434 | 1448 | - | 3.3571 | 7.12E-05 | 0.795 | GCCACGGCAATACTG |
| 1 | GMTACGGTCACACTG | chr2R:16736885-16738885 | 1435 | 1449 | - | 3.3571 | 7.12E-05 | 0.795 | GCTACGCATGCACTG |
| 1 | GMTACGGTCACACTG | chrX:9231378-9233378 | 1435 | 1449 | - | 3.3571 | 7.12E-05 | 0.795 | GCCAGGATCACTCTG |
| 1 | GMTACGGTCACACTG | chr2R:8913524-8915524 | 1437 | 1451 | - | 3.3571 | 7.12E-05 | 0.795 | GCTACGCGCACACAA |
| 1 | GMTACGGTCACACTG | chrX:23105003-23107003 | 1439 | 1453 | - | 3.3571 | 7.12E-05 | 0.795 | GCCAGAGGCACACTG |
| 1 | GMTACGGTCACACTG | chr3L:270200-272200 | 1439 | 1453 | - | 3.3571 | 7.12E-05 | 0.795 | GCTGTGGGCAAACTG |
| 1 | GMTACGGTCACACTG | chrX:9607946-9609946 | 1440 | 1454 | - | 3.3571 | 7.12E-05 | 0.795 | GCTGCAGTCTGACTG |
| 1 | GMTACGGTCACACTG | chr3R:721621-723621 | 1440 | 1454 | - | 3.3571 | 7.12E-05 | 0.795 | GCTCCGGCCAAATTG |
| 1 | GMTACGGTCACACTG | chrX:18084761-18086761 | 1443 | 1457 | - | 3.3571 | 7.12E-05 | 0.795 | GCTCCAGTTGCACTG |
| 1 | GMTACGGTCACACTG | chr3R:15274241-15276241 | 1445 | 1459 | - | 3.3571 | 7.12E-05 | 0.795 | GCTGCGGGCACAATT |
| 1 | GMTACGGTCACACTG | chr3R:6247994-6249994 | 1446 | 1460 | - | 3.3571 | 7.12E-05 | 0.795 | ACCACGGTCACACCT |
| 1 | GMTACGGTCACACTG | chr3L:812300-814300 | 1447 | 1461 | - | 3.3571 | 7.12E-05 | 0.795 | GCTGCAGACAAACTG |

| | | | | | | | | | |
|---|---|---|---|---|---|---|---|---|---|
| 1 | GMTACGGTCACACTG | chr3R:28811102-28813102 | 1454 | 1468 | - | 3.3571 | 7.12E-05 | 0.795 | GCCACACCCACACTG |
| 1 | GMTACGGTCACACTG | chr2L:1499830-1501830 | 1454 | 1468 | - | 3.3571 | 7.12E-05 | 0.795 | GCTACGGTCAGCTGG |
| 1 | GMTACGGTCACACTG | chr3R:6892462-6894462 | 1461 | 1475 | - | 3.3571 | 7.12E-05 | 0.795 | GCTTGGTTCCCACTG |
| 1 | GMTACGGTCACACTG | chr3R:26618118-26620118 | 1464 | 1478 | - | 3.3571 | 7.12E-05 | 0.795 | GAGACGGGGGCACTG |
| 1 | GMTACGGTCACACTG | chr3L:1033875-1035875 | 1464 | 1478 | - | 3.3571 | 7.12E-05 | 0.795 | GATCGGGGCGCACTG |
| 1 | GMTACGGTCACACTG | chr3L:8754989-8756989 | 1465 | 1479 | - | 3.3571 | 7.12E-05 | 0.795 | GCTCCTGCAACACTG |
| 1 | GMTACGGTCACACTG | chr2L:13283927-13285927 | 1466 | 1480 | - | 3.3571 | 7.12E-05 | 0.795 | GCTATGGCCACACGT |
| 1 | GMTACGGTCACACTG | chr3R:29674651-29676651 | 1467 | 1481 | - | 3.3571 | 7.12E-05 | 0.795 | GACATGGGCTCACTG |
| 1 | GMTACGGTCACACTG | chr3R:25231321-25233321 | 1469 | 1483 | - | 3.3571 | 7.12E-05 | 0.795 | GCAGCGGTCAAAATG |
| 1 | GMTACGGTCACACTG | chr3R:6762478-6764478 | 1469 | 1483 | - | 3.3571 | 7.12E-05 | 0.795 | GAGTCGCCCACACTG |
| 1 | GMTACGGTCACACTG | chr3R:23932530-23934530 | 1471 | 1485 | - | 3.3571 | 7.12E-05 | 0.795 | GACACGGTCAACCGG |
| 1 | GMTACGGTCACACTG | chrX:21385935-21387935 | 1473 | 1487 | - | 3.3571 | 7.12E-05 | 0.795 | GACACGGTGACCCAG |
| 1 | GMTACGGTCACACTG | chr3L:3105958-3107958 | 1474 | 1488 | - | 3.3571 | 7.12E-05 | 0.795 | GCTACTGCCCCATTG |
| 1 | GMTACGGTCACACTG | chr2R:19687078-19689078 | 1474 | 1488 | - | 3.3571 | 7.12E-05 | 0.795 | GCTCGGGTGCCACTG |
| 1 | GMTACGGTCACACTG | chr2R:11865943-11867943 | 1476 | 1490 | - | 3.3571 | 7.12E-05 | 0.795 | CCTGCTGCCACACTG |
| 1 | GMTACGGTCACACTG | chr3L:7455149-7457149 | 1476 | 1490 | - | 3.3571 | 7.12E-05 | 0.795 | TCTCCGGACAAACTG |
| 1 | GMTACGGTCACACTG | chr3R:5823615-5825615 | 1480 | 1494 | - | 3.3571 | 7.12E-05 | 0.795 | GAGTCGGATACACTG |
| 1 | GMTACGGTCACACTG | chr2L:13641602-13643602 | 1481 | 1495 | - | 3.3571 | 7.12E-05 | 0.795 | GATTCAGGCACACAG |
| 1 | GMTACGGTCACACTG | chr3L:6261694-6263694 | 1487 | 1501 | - | 3.3571 | 7.12E-05 | 0.795 | GCAAAGGTCACCTTG |
| 1 | GMTACGGTCACACTG | chr3L:11102873-11104873 | 1487 | 1501 | - | 3.3571 | 7.12E-05 | 0.795 | CCTACGGACAGACAG |
| 1 | GMTACGGTCACACTG | chrX:4938128-4940128 | 1487 | 1501 | - | 3.3571 | 7.12E-05 | 0.795 | CACTCGGACACACTG |
| 1 | GMTACGGTCACACTG | chr2L:11010312-11012312 | 1489 | 1503 | - | 3.3571 | 7.12E-05 | 0.795 | GATGCGGTCGCCATG |
| 1 | GMTACGGTCACACTG | chrX:19530742-19532742 | 1490 | 1504 | - | 3.3571 | 7.12E-05 | 0.795 | CCTACTGACCCACTG |
| 1 | GMTACGGTCACACTG | chr2R:21535795-21537795 | 1490 | 1504 | - | 3.3571 | 7.12E-05 | 0.795 | GATGGGGACGCACTG |
| 1 | GMTACGGTCACACTG | chr3L:20298410-20300410 | 1491 | 1505 | - | 3.3571 | 7.12E-05 | 0.795 | GATCCGGGGACTCTG |
| 1 | GMTACGGTCACACTG | chr3L:20298410-20300410 | 1491 | 1505 | - | 3.3571 | 7.12E-05 | 0.795 | GATCCGGGGACTCTG |
| 1 | GMTACGGTCACACTG | chr3R:20849516-20851516 | 1492 | 1506 | - | 3.3571 | 7.12E-05 | 0.795 | GCGATGGCCACAATG |
| 1 | GMTACGGTCACACTG | chrX:1379914-1381914 | 1494 | 1508 | - | 3.3571 | 7.12E-05 | 0.795 | GCGACGGTCACAAGC |
| 1 | GMTACGGTCACACTG | chr2R:18862896-18864896 | 1495 | 1509 | - | 3.3571 | 7.12E-05 | 0.795 | GAGACAGACACACGG |
| 1 | GMTACGGTCACACTG | chr2L:2758924-2760924 | 1499 | 1513 | - | 3.3571 | 7.12E-05 | 0.795 | GCAACCGCCAGACTG |
| 1 | GMTACGGTCACACTG | chr3R:13900853-13902853 | 1499 | 1513 | - | 3.3571 | 7.12E-05 | 0.795 | GAACCGGGCACATTG |
| 1 | GMTACGGTCACACTG | chrX:9607884-9609884 | 1502 | 1516 | - | 3.3571 | 7.12E-05 | 0.795 | GCTGCAGTCTGACTG |
| 1 | GMTACGGTCACACTG | chr2L:3712360-3714360 | 1503 | 1517 | - | 3.3571 | 7.12E-05 | 0.795 | GCTGCGGTCACAAGT |

| | | | | | | | | | |
|---|---|---|---|---|---|---|---|---|---|
| 1 | GMTACGGTCACACTG | chrX:12058795-12060795 | 1503 | 1517 | - | 3.3571 | 7.12E-05 | 0.795 | GATGCGGTCTCCATG |
| 1 | GMTACGGTCACACTG | chrX:19730004-19732004 | 1512 | 1526 | - | 3.3571 | 7.12E-05 | 0.795 | CCCACGGTCATGCTG |
| 1 | GMTACGGTCACACTG | chr2L:13386312-13388312 | 1514 | 1528 | - | 3.3571 | 7.12E-05 | 0.795 | GATGCGGTCAGGCAG |
| 1 | GMTACGGTCACACTG | chr2R:12334316-12336316 | 1514 | 1528 | - | 3.3571 | 7.12E-05 | 0.795 | GCCACGCTCACCTTG |
| 1 | GMTACGGTCACACTG | chr3R:14800773-14802773 | 1516 | 1530 | - | 3.3571 | 7.12E-05 | 0.795 | GATCCTGTCACCCAG |
| 1 | GMTACGGTCACACTG | chr3L:5509860-5511860 | 1519 | 1533 | - | 3.3571 | 7.12E-05 | 0.795 | GACACGCACACACGG |
| 1 | GMTACGGTCACACTG | chr3R:20591671-20593671 | 1519 | 1533 | - | 3.3571 | 7.12E-05 | 0.795 | CACACGGTCTCACGG |
| 1 | GMTACGGTCACACTG | chr3R:11689051-11691051 | 1520 | 1534 | - | 3.3571 | 7.12E-05 | 0.795 | GCCACGGAAATACTG |
| 1 | GMTACGGTCACACTG | chr2R:19269078-19271078 | 1521 | 1535 | - | 3.3571 | 7.12E-05 | 0.795 | GCCACGTTTTCACTG |
| 1 | GMTACGGTCACACTG | chr2L:16730039-16732039 | 1525 | 1539 | - | 3.3571 | 7.12E-05 | 0.795 | GCTACGATCCCAACG |
| 1 | GMTACGGTCACACTG | chr2L:1728887-1730887 | 1525 | 1539 | - | 3.3571 | 7.12E-05 | 0.795 | ACTTCGGTCCAACTG |
| 1 | GMTACGGTCACACTG | chrX:12489291-12491291 | 1530 | 1544 | - | 3.3571 | 7.12E-05 | 0.795 | GCGAAGGCCACATTG |
| 1 | GMTACGGTCACACTG | chr2R:20057095-20059095 | 1531 | 1545 | - | 3.3571 | 7.12E-05 | 0.795 | ACTAGGGCCACACAG |
| 1 | GMTACGGTCACACTG | chr3R:7912290-7914290 | 1533 | 1547 | - | 3.3571 | 7.12E-05 | 0.795 | CCAACCGTCGCACTG |
| 1 | GMTACGGTCACACTG | chr3L:14968344-14970344 | 1533 | 1547 | - | 3.3571 | 7.12E-05 | 0.795 | CCAACGGTCGAACTG |
| 1 | GMTACGGTCACACTG | chr2L:21083326-21085326 | 1535 | 1549 | - | 3.3571 | 7.12E-05 | 0.795 | GCTACGGTCGTAAAG |
| 1 | GMTACGGTCACACTG | chr2R:12050755-12052755 | 1540 | 1554 | - | 3.3571 | 7.12E-05 | 0.795 | GCTAAGGACATCCTG |
| 1 | GMTACGGTCACACTG | chr3R:8012201-8014201 | 1544 | 1558 | - | 3.3571 | 7.12E-05 | 0.795 | GCAATGGTCGAACTG |
| 1 | GMTACGGTCACACTG | chr3R:8291335-8293335 | 1546 | 1560 | - | 3.3571 | 7.12E-05 | 0.795 | GCTGCGGTCACGATC |
| 1 | GMTACGGTCACACTG | chr3R:6833967-6835967 | 1546 | 1560 | - | 3.3571 | 7.12E-05 | 0.795 | GAACCGGACACAGTG |
| 1 | GMTACGGTCACACTG | chr2L:14875832-14877832 | 1547 | 1561 | - | 3.3571 | 7.12E-05 | 0.795 | GATTCGGTAGCCCTG |
| 1 | GMTACGGTCACACTG | chrX:21216878-21218878 | 1551 | 1565 | - | 3.3571 | 7.12E-05 | 0.795 | TCTACGGCGACACAG |
| 1 | GMTACGGTCACACTG | chr3R:12157595-12159595 | 1556 | 1570 | - | 3.3571 | 7.12E-05 | 0.795 | GCTGCTGTCCCACTT |
| 1 | GMTACGGTCACACTG | chr3R:8778923-8780923 | 1558 | 1572 | - | 3.3571 | 7.12E-05 | 0.795 | TATCCGGCCACCCTG |
| 1 | GMTACGGTCACACTG | chr2R:21309146-21311146 | 1559 | 1573 | - | 3.3571 | 7.12E-05 | 0.795 | GAAACGGCTACGCTG |
| 1 | GMTACGGTCACACTG | chrX:18084761-18086761 | 1561 | 1575 | - | 3.3571 | 7.12E-05 | 0.795 | GATCCAGGCACACGG |
| 1 | GMTACGGTCACACTG | chr2L:10530821-10532821 | 1562 | 1576 | - | 3.3571 | 7.12E-05 | 0.795 | GCCATCGTCACTCTG |
| 1 | GMTACGGTCACACTG | chr3R:5823532-5825532 | 1563 | 1577 | - | 3.3571 | 7.12E-05 | 0.795 | GAGTCGGATACACTG |
| 1 | GMTACGGTCACACTG | chr3L:689517-691517 | 1572 | 1586 | - | 3.3571 | 7.12E-05 | 0.795 | GCATAGGTCACAATG |
| 1 | GMTACGGTCACACTG | chrX:10951486-10953486 | 1575 | 1589 | - | 3.3571 | 7.12E-05 | 0.795 | ACCAAGGACACACTG |
| 1 | GMTACGGTCACACTG | chr3R:10734957-10736957 | 1575 | 1589 | - | 3.3571 | 7.12E-05 | 0.795 | GCTACGGGAACTGTG |
| 1 | GMTACGGTCACACTG | chrX:12856722-12858722 | 1576 | 1590 | - | 3.3571 | 7.12E-05 | 0.795 | CCTACGCACACACAG |
| 1 | GMTACGGTCACACTG | chrX:12407477-12409477 | 1579 | 1593 | - | 3.3571 | 7.12E-05 | 0.795 | GAGCTGGCCACACTG |

| | | | | | | | | | |
|---|---|---|---|---|---|---|---|---|---|
| 1 | GMTACGGTCACACTG | chr2R:17563340-17565340 | 1579 | 1593 | - | 3.3571 | 7.12E-05 | 0.795 | GCTTCAGTCACAAAG |
| 1 | GMTACGGTCACACTG | chr3R:4851813-4853813 | 1580 | 1594 | - | 3.3571 | 7.12E-05 | 0.795 | GATCCTGCCACACCG |
| 1 | GMTACGGTCACACTG | chr3R:15275217-15277217 | 1581 | 1595 | - | 3.3571 | 7.12E-05 | 0.795 | CATCCGGACACACCG |
| 1 | GMTACGGTCACACTG | chr3R:28805702-28807702 | 1584 | 1598 | - | 3.3571 | 7.12E-05 | 0.795 | GAGGCGGACAAACTG |
| 1 | GMTACGGTCACACTG | chr2L:10650613-10652613 | 1585 | 1599 | - | 3.3571 | 7.12E-05 | 0.795 | GCCACGCGCACAATG |
| 1 | GMTACGGTCACACTG | chr3R:30487963-30489963 | 1586 | 1600 | - | 3.3571 | 7.12E-05 | 0.795 | GCTGCAATCACTCTG |
| 1 | GMTACGGTCACACTG | chr3R:19876514-19878514 | 1588 | 1602 | - | 3.3571 | 7.12E-05 | 0.795 | GCTGCGTTTCCACTG |
| 1 | GMTACGGTCACACTG | chr2R:22654392-22656392 | 1589 | 1603 | - | 3.3571 | 7.12E-05 | 0.795 | ACCACTGTCACGCTG |
| 1 | GMTACGGTCACACTG | chr3R:11352131-11354131 | 1590 | 1604 | - | 3.3571 | 7.12E-05 | 0.795 | GCTACTCCCACACCG |
| 1 | GMTACGGTCACACTG | chr3R:11352131-11354131 | 1590 | 1604 | - | 3.3571 | 7.12E-05 | 0.795 | GCTACTCCCACACCG |
| 1 | GMTACGGTCACACTG | chrX:16296718-16298718 | 1591 | 1605 | - | 3.3571 | 7.12E-05 | 0.795 | GAGATGGTCTCGCTG |
| 1 | GMTACGGTCACACTG | chr3L:9725895-9727895 | 1592 | 1606 | - | 3.3571 | 7.12E-05 | 0.795 | CCTCCGTTCCCACTG |
| 1 | GMTACGGTCACACTG | chr3L:21886039-21888039 | 1596 | 1610 | - | 3.3571 | 7.12E-05 | 0.795 | GCTCAGCTCACACCG |
| 1 | GMTACGGTCACACTG | chrX:768945-770945 | 1598 | 1612 | - | 3.3571 | 7.12E-05 | 0.795 | GCGACGGAAACACTT |
| 1 | GMTACGGTCACACTG | chr2L:2744098-2746098 | 1601 | 1615 | - | 3.3571 | 7.12E-05 | 0.795 | GAGAAGGGCACACAG |
| 1 | GMTACGGTCACACTG | chr2R:24113398-24115398 | 1602 | 1616 | - | 3.3571 | 7.12E-05 | 0.795 | GCTAGGGACACTCTA |
| 1 | GMTACGGTCACACTG | chr2L:16351628-16353628 | 1604 | 1618 | - | 3.3571 | 7.12E-05 | 0.795 | GCTGCACTCGCACTG |
| 1 | GMTACGGTCACACTG | chr2L:16351628-16353628 | 1604 | 1618 | - | 3.3571 | 7.12E-05 | 0.795 | GCTGCACTCGCACTG |
| 1 | GMTACGGTCACACTG | chr2R:9137486-9139486 | 1605 | 1619 | - | 3.3571 | 7.12E-05 | 0.795 | TCTGCGATCGCACTG |
| 1 | GMTACGGTCACACTG | chr2R:9137486-9139486 | 1605 | 1619 | - | 3.3571 | 7.12E-05 | 0.795 | TCTGCGATCGCACTG |
| 1 | GMTACGGTCACACTG | chr2R:22378007-22380007 | 1609 | 1623 | - | 3.3571 | 7.12E-05 | 0.795 | GAGCCGCACACACTG |
| 1 | GMTACGGTCACACTG | chr2R:18612090-18614090 | 1612 | 1626 | - | 3.3571 | 7.12E-05 | 0.795 | GCAATGGACACAATG |
| 1 | GMTACGGTCACACTG | chr2L:8214915-8216915 | 1615 | 1629 | - | 3.3571 | 7.12E-05 | 0.795 | GCTACGTTCTCGATG |
| 1 | GMTACGGTCACACTG | chr3R:31746643-31748643 | 1616 | 1630 | - | 3.3571 | 7.12E-05 | 0.795 | GAAACGGACACAGCG |
| 1 | GMTACGGTCACACTG | chr2R:9131030-9133030 | 1619 | 1633 | - | 3.3571 | 7.12E-05 | 0.795 | GCTTCGGTTACACGC |
| 1 | GMTACGGTCACACTG | chr2R:17800132-17802132 | 1621 | 1635 | - | 3.3571 | 7.12E-05 | 0.795 | GACACGGGCAAACCG |
| 1 | GMTACGGTCACACTG | chrX:10626112-10628112 | 1621 | 1635 | - | 3.3571 | 7.12E-05 | 0.795 | GCTGAGAACACACTG |
| 1 | GMTACGGTCACACTG | chr2R:17868027-17870027 | 1621 | 1635 | - | 3.3571 | 7.12E-05 | 0.795 | GAGACGGACAGACGG |
| 1 | GMTACGGTCACACTG | chr3L:3270036-3272036 | 1624 | 1638 | - | 3.3571 | 7.12E-05 | 0.795 | GCATTTGTCACACTG |
| 1 | GMTACGGTCACACTG | chr3L:11789013-11791013 | 1628 | 1642 | - | 3.3571 | 7.12E-05 | 0.795 | GCTTTGGTCAATCTG |
| 1 | GMTACGGTCACACTG | chr2L:7710476-7712476 | 1633 | 1647 | - | 3.3571 | 7.12E-05 | 0.795 | GACGCGGTTACACAG |
| 1 | GMTACGGTCACACTG | chr3R:30741880-30743880 | 1638 | 1652 | - | 3.3571 | 7.12E-05 | 0.795 | GATTCGGTGGCACAG |
| 1 | GMTACGGTCACACTG | chr2R:23971431-23973431 | 1639 | 1653 | - | 3.3571 | 7.12E-05 | 0.795 | CCAACGACCACACTG |

| | | | | | | | | | |
|---|---|---|---|---|---|---|---|---|---|
| 1 | GMTACGGTCACACTG | chr2L:5009922-5011922 | 1640 | 1654 | - | 3.3571 | 7.12E-05 | 0.795 | ACCACGGACACAATG |
| 1 | GMTACGGTCACACTG | chr2L:18155043-18157043 | 1641 | 1655 | - | 3.3571 | 7.12E-05 | 0.795 | GATTGGGCCACTCTG |
| 1 | GMTACGGTCACACTG | chr3L:1033697-1035697 | 1642 | 1656 | - | 3.3571 | 7.12E-05 | 0.795 | GATCGGGGCGCACTG |
| 1 | GMTACGGTCACACTG | chr2L:567340-569340 | 1643 | 1657 | - | 3.3571 | 7.12E-05 | 0.795 | GATCCGCTCGCTCTG |
| 1 | GMTACGGTCACACTG | chrX:22854194-22856194 | 1643 | 1657 | - | 3.3571 | 7.12E-05 | 0.795 | GCCACGCTCACACCT |
| 1 | GMTACGGTCACACTG | chr3R:29216184-29218184 | 1646 | 1660 | - | 3.3571 | 7.12E-05 | 0.795 | GCCACGGTAACAACG |
| 1 | GMTACGGTCACACTG | chr3R:24171628-24173628 | 1648 | 1662 | - | 3.3571 | 7.12E-05 | 0.795 | GCAACTGTCAAACCG |
| 1 | GMTACGGTCACACTG | chr2L:3507598-3509598 | 1648 | 1662 | - | 3.3571 | 7.12E-05 | 0.795 | GCTACGGTTGCGCTA |
| 1 | GMTACGGTCACACTG | chr3R:19227180-19229180 | 1651 | 1665 | - | 3.3571 | 7.12E-05 | 0.795 | GATACGGTCTCGAAG |
| 1 | GMTACGGTCACACTG | chr3L:2272744-2274744 | 1651 | 1665 | - | 3.3571 | 7.12E-05 | 0.795 | GATTCGGCCCGACTG |
| 1 | GMTACGGTCACACTG | chrX:5621648-5623648 | 1655 | 1669 | - | 3.3571 | 7.12E-05 | 0.795 | CCCAAGGCCACACTG |
| 1 | GMTACGGTCACACTG | chr3R:22030900-22032900 | 1659 | 1673 | - | 3.3571 | 7.12E-05 | 0.795 | CCGGCGGTCACATTG |
| 1 | GMTACGGTCACACTG | chr3R:11979390-11981390 | 1659 | 1673 | - | 3.3571 | 7.12E-05 | 0.795 | TCGACGGTCTCACTT |
| 1 | GMTACGGTCACACTG | chr2R:10647888-10649888 | 1664 | 1678 | - | 3.3571 | 7.12E-05 | 0.795 | GCTTCTGTTCCACTG |
| 1 | GMTACGGTCACACTG | chr2L:8383137-8385137 | 1665 | 1679 | - | 3.3571 | 7.12E-05 | 0.795 | GCGAAGGTCACAACG |
| 1 | GMTACGGTCACACTG | chr3R:9996163-9998163 | 1666 | 1680 | - | 3.3571 | 7.12E-05 | 0.795 | GCCACCAGCACACTG |
| 1 | GMTACGGTCACACTG | chrX:6290303-6292303 | 1667 | 1681 | - | 3.3571 | 7.12E-05 | 0.795 | TAGGCGGCCACACTG |
| 1 | GMTACGGTCACACTG | chr3L:5712331-5714331 | 1667 | 1681 | - | 3.3571 | 7.12E-05 | 0.795 | GCAACTTGCACACTG |
| 1 | GMTACGGTCACACTG | chr3L:6865439-6867439 | 1669 | 1683 | - | 3.3571 | 7.12E-05 | 0.795 | GATACGGATACGCAG |
| 1 | GMTACGGTCACACTG | chr2L:9917618-9919618 | 1671 | 1685 | - | 3.3571 | 7.12E-05 | 0.795 | GCTGTGGGAACACTG |
| 1 | GMTACGGTCACACTG | chr3L:3904344-3906344 | 1671 | 1685 | - | 3.3571 | 7.12E-05 | 0.795 | GCTACGGCCATTCTT |
| 1 | GMTACGGTCACACTG | chr3R:15802495-15804495 | 1673 | 1687 | - | 3.3571 | 7.12E-05 | 0.795 | TCTGCGGCCACATTG |
| 1 | GMTACGGTCACACTG | chr2L:1499608-1501608 | 1676 | 1690 | - | 3.3571 | 7.12E-05 | 0.795 | GCTACGGTCAGCTGG |
| 1 | GMTACGGTCACACTG | chr2R:10810078-10812078 | 1680 | 1694 | - | 3.3571 | 7.12E-05 | 0.795 | CACACGGACGCACTG |
| 1 | GMTACGGTCACACTG | chr2R:14854048-14856048 | 1680 | 1694 | - | 3.3571 | 7.12E-05 | 0.795 | GCCTTGCTCACACTG |
| 1 | GMTACGGTCACACTG | chr2L:19261470-19263470 | 1685 | 1699 | - | 3.3571 | 7.12E-05 | 0.795 | GCTCGGATCACACAG |
| 1 | GMTACGGTCACACTG | chr3L:14536651-14538651 | 1692 | 1706 | - | 3.3571 | 7.12E-05 | 0.795 | GCCACGGTGACCTTG |
| 1 | GMTACGGTCACACTG | chr2L:2008155-2010155 | 1693 | 1707 | - | 3.3571 | 7.12E-05 | 0.795 | GCAACTGTCAATCTG |
| 1 | GMTACGGTCACACTG | chr3L:25842821-25844821 | 1694 | 1708 | - | 3.3571 | 7.12E-05 | 0.795 | GCCACGCCCACACTT |
| 1 | GMTACGGTCACACTG | chrX:16598241-16600241 | 1695 | 1709 | - | 3.3571 | 7.12E-05 | 0.795 | GCTTCGTTCTCACTC |
| 1 | GMTACGGTCACACTG | chrX:1872565-1874565 | 1695 | 1709 | - | 3.3571 | 7.12E-05 | 0.795 | GATACCGTCTCCCCG |
| 1 | GMTACGGTCACACTG | chr3R:29674421-29676421 | 1697 | 1711 | - | 3.3571 | 7.12E-05 | 0.795 | GACATGGGCTCACTG |
| 1 | GMTACGGTCACACTG | chr3L:7438405-7440405 | 1699 | 1713 | - | 3.3571 | 7.12E-05 | 0.795 | GCGGTGGTCACAATG |

| | | | | | | | | |
|---|---|---|---|---|---|---|---|---|
| 1 | GMTACGGTCACACTG | chr2R:8086396-8088396 | 1699 | 1713 | - | 3.3571 | 7.12E-05 | 0.795 CCGACGCTCCCACTG |
| 1 | GMTACGGTCACACTG | chr3R:31746558-31748558 | 1701 | 1715 | - | 3.3571 | 7.12E-05 | 0.795 GAAACGGACACAGCG |
| 1 | GMTACGGTCACACTG | chr2L:6019835-6021835 | 1701 | 1715 | - | 3.3571 | 7.12E-05 | 0.795 GATACCGCCTCACCG |
| 1 | GMTACGGTCACACTG | chr3L:19611455-19613455 | 1701 | 1715 | - | 3.3571 | 7.12E-05 | 0.795 GCTCCGATCACGTTG |
| 1 | GMTACGGTCACACTG | chrX:11583175-11585175 | 1701 | 1715 | - | 3.3571 | 7.12E-05 | 0.795 GCTAAGTTCTCGCTG |
| 1 | GMTACGGTCACACTG | chr3L:20651906-20653906 | 1706 | 1720 | - | 3.3571 | 7.12E-05 | 0.795 GCTGCGGCCAAATTG |
| 1 | GMTACGGTCACACTG | chr2R:24113292-24115292 | 1708 | 1722 | - | 3.3571 | 7.12E-05 | 0.795 GCTAGGGACACTCTA |
| 1 | GMTACGGTCACACTG | chr4:1083796-1085796 | 1712 | 1726 | - | 3.3571 | 7.12E-05 | 0.795 GCCACGTCCACACTT |
| 1 | GMTACGGTCACACTG | chr2R:13557545-13559545 | 1714 | 1728 | - | 3.3571 | 7.12E-05 | 0.795 GCAATGGAGACACTG |
| 1 | GMTACGGTCACACTG | chr2L:11010086-11012086 | 1715 | 1729 | - | 3.3571 | 7.12E-05 | 0.795 GATGCGGTCGCCATG |
| 1 | GMTACGGTCACACTG | chr3R:21268060-21270060 | 1715 | 1729 | - | 3.3571 | 7.12E-05 | 0.795 GCCGTGGTCACAGTG |
| 1 | GMTACGGTCACACTG | chr3R:8934287-8936287 | 1715 | 1729 | - | 3.3571 | 7.12E-05 | 0.795 TCAGCGGTCACATTG |
| 1 | GMTACGGTCACACTG | chr3R:16098228-16100228 | 1718 | 1732 | - | 3.3571 | 7.12E-05 | 0.795 GAGGCAGTCACACAG |
| 1 | GMTACGGTCACACTG | chrX:16598215-16600215 | 1721 | 1735 | - | 3.3571 | 7.12E-05 | 0.795 GCTTCGTTCTCACTC |
| 1 | GMTACGGTCACACTG | chr3R:29674651-29676651 | 1722 | 1736 | - | 3.3571 | 7.12E-05 | 0.795 GCTACGGGCGGACTC |
| 1 | GMTACGGTCACACTG | chr3L:18916581-18918581 | 1722 | 1736 | - | 3.3571 | 7.12E-05 | 0.795 GATCCGCCCACGCTG |
| 1 | GMTACGGTCACACTG | chrX:6290245-6292245 | 1725 | 1739 | - | 3.3571 | 7.12E-05 | 0.795 TAGGCGGCCACACTG |
| 1 | GMTACGGTCACACTG | chr3L:2168108-2170108 | 1726 | 1740 | - | 3.3571 | 7.12E-05 | 0.795 GCGCCGGTTACATTG |
| 1 | GMTACGGTCACACTG | chr3L:2660622-2662622 | 1727 | 1741 | - | 3.3571 | 7.12E-05 | 0.795 GCTCCGTTCGCAATG |
| 1 | GMTACGGTCACACTG | chr2L:5009834-5011834 | 1728 | 1742 | - | 3.3571 | 7.12E-05 | 0.795 ACCACGGACACAATG |
| 1 | GMTACGGTCACACTG | chr2L:5009834-5011834 | 1728 | 1742 | - | 3.3571 | 7.12E-05 | 0.795 ACCACGGACACAATG |
| 1 | GMTACGGTCACACTG | chrX:20510544-20512544 | 1730 | 1744 | - | 3.3571 | 7.12E-05 | 0.795 GCTATGCGCACTCTG |
| 1 | GMTACGGTCACACTG | chr3R:30513777-30515777 | 1735 | 1749 | - | 3.3571 | 7.12E-05 | 0.795 GCTATGTCCACACAG |
| 1 | GMTACGGTCACACTG | chr3L:3279239-3281239 | 1739 | 1753 | - | 3.3571 | 7.12E-05 | 0.795 GCTTCGGTGGCAGTG |
| 1 | GMTACGGTCACACTG | chr2R:5759862-5761862 | 1740 | 1754 | - | 3.3571 | 7.12E-05 | 0.795 GCGATGGTCACCTTG |
| 1 | GMTACGGTCACACTG | chr3L:7638717-7640717 | 1745 | 1759 | - | 3.3571 | 7.12E-05 | 0.795 GACGCGGTCACACAT |
| 1 | GMTACGGTCACACTG | chr2L:2758677-2760677 | 1746 | 1760 | - | 3.3571 | 7.12E-05 | 0.795 GCAACCGCCAGACTG |
| 1 | GMTACGGTCACACTG | chr2L:2758677-2760677 | 1746 | 1760 | - | 3.3571 | 7.12E-05 | 0.795 GCAACCGCCAGACTG |
| 1 | GMTACGGTCACACTG | chrX:126449-128449 | 1749 | 1763 | - | 3.3571 | 7.12E-05 | 0.795 GCGACAGTAGCACTG |
| 1 | GMTACGGTCACACTG | chr3R:5222365-5224365 | 1750 | 1764 | - | 3.3571 | 7.12E-05 | 0.795 GCTGGTGTCCCACTG |
| 1 | GMTACGGTCACACTG | chr3L:7527780-7529780 | 1751 | 1765 | - | 3.3571 | 7.12E-05 | 0.795 CACACGGACGCACTG |
| 1 | GMTACGGTCACACTG | chr2L:13202295-13204295 | 1751 | 1765 | - | 3.3571 | 7.12E-05 | 0.795 GCGACGTTCACGCTT |
| 1 | GMTACGGTCACACTG | chrX:23290700-23292700 | 1753 | 1767 | - | 3.3571 | 7.12E-05 | 0.795 GCTACGATCTCCATG |

| | | | | | | | | | |
|---|---|---|---|---|---|---|---|---|---|
| 1 | GMTACGGTCACACTG | chr2L:19579351-19581351 | 1755 | 1769 | - | 3.3571 | 7.12E-05 | 0.795 | TCGACGCTCACGCTG |
| 1 | GMTACGGTCACACTG | chr2L:3712105-3714105 | 1758 | 1772 | - | 3.3571 | 7.12E-05 | 0.795 | GCTGCGGTCACAAGT |
| 1 | GMTACGGTCACACTG | chr2L:7362379-7364379 | 1759 | 1773 | - | 3.3571 | 7.12E-05 | 0.795 | GCTACTGGCGGACTG |
| 1 | GMTACGGTCACACTG | chr2R:19661436-19663436 | 1759 | 1773 | - | 3.3571 | 7.12E-05 | 0.795 | GCTACGGCGAAACGG |
| 1 | GMTACGGTCACACTG | chr2R:6509295-6511295 | 1761 | 1775 | - | 3.3571 | 7.12E-05 | 0.795 | GCTTCGTTTGCACTG |
| 1 | GMTACGGTCACACTG | chr3L:7344183-7346183 | 1762 | 1776 | - | 3.3571 | 7.12E-05 | 0.795 | GAAGCCGTCACCCTG |
| 1 | GMTACGGTCACACTG | chr2L:18670246-18672246 | 1763 | 1777 | - | 3.3571 | 7.12E-05 | 0.795 | GCCAGGCTCGCACTG |
| 1 | GMTACGGTCACACTG | chr2L:1496243-1498243 | 1763 | 1777 | - | 3.3571 | 7.12E-05 | 0.795 | GCCTCGGTCACATTT |
| 1 | GMTACGGTCACACTG | chr3L:13060422-13062422 | 1763 | 1777 | - | 3.3571 | 7.12E-05 | 0.795 | GCTCCGGTAACACCC |
| 1 | GMTACGGTCACACTG | chr2R:12334066-12336066 | 1764 | 1778 | - | 3.3571 | 7.12E-05 | 0.795 | GCCACGCTCACCTTG |
| 1 | GMTACGGTCACACTG | chrX:5621539-5623539 | 1764 | 1778 | - | 3.3571 | 7.12E-05 | 0.795 | CCCAAGGCCACACTG |
| 1 | GMTACGGTCACACTG | chr2L:2743934-2745934 | 1765 | 1779 | - | 3.3571 | 7.12E-05 | 0.795 | GAGAAGGGCACACAG |
| 1 | GMTACGGTCACACTG | chr2L:10819705-10821705 | 1765 | 1779 | - | 3.3571 | 7.12E-05 | 0.795 | GAAACGGGCACATAG |
| 1 | GMTACGGTCACACTG | chr3L:3110227-3112227 | 1765 | 1779 | - | 3.3571 | 7.12E-05 | 0.795 | GCTACGGTAGCTTTG |
| 1 | GMTACGGTCACACTG | chr2R:19269078-19271078 | 1765 | 1779 | - | 3.3571 | 7.12E-05 | 0.795 | GCCATGGTCACATGG |
| 1 | GMTACGGTCACACTG | chr3L:8575078-8577078 | 1769 | 1783 | - | 3.3571 | 7.12E-05 | 0.795 | GCTCCTGGCAGACTG |
| 1 | GMTACGGTCACACTG | chrX:18090259-18092259 | 1770 | 1784 | - | 3.3571 | 7.12E-05 | 0.795 | GCCAGGATCACGCTG |
| 1 | GMTACGGTCACACTG | chr2R:14859758-14861758 | 1771 | 1785 | - | 3.3571 | 7.12E-05 | 0.795 | GAGACGGGCAGCCTG |
| 1 | GMTACGGTCACACTG | chr2L:1728636-1730636 | 1776 | 1790 | - | 3.3571 | 7.12E-05 | 0.795 | ACTTCGGTCCAACTG |
| 1 | GMTACGGTCACACTG | chr2R:8361156-8363156 | 1778 | 1792 | - | 3.3571 | 7.12E-05 | 0.795 | GAATCAGTCCCACTG |
| 1 | GMTACGGTCACACTG | chr2R:8361156-8363156 | 1778 | 1792 | - | 3.3571 | 7.12E-05 | 0.795 | GAATCAGTCCCACTG |
| 1 | GMTACGGTCACACTG | chrX:9594082-9596082 | 1779 | 1793 | - | 3.3571 | 7.12E-05 | 0.795 | GCATGGGTCACACTC |
| 1 | GMTACGGTCACACTG | chr3L:3471447-3473447 | 1782 | 1796 | - | 3.3571 | 7.12E-05 | 0.795 | GCACTTGTCACACTG |
| 1 | GMTACGGTCACACTG | chr2R:14755516-14757516 | 1782 | 1796 | - | 3.3571 | 7.12E-05 | 0.795 | GATAAGGGCTCGCTG |
| 1 | GMTACGGTCACACTG | chr3L:8967528-8969528 | 1786 | 1800 | - | 3.3571 | 7.12E-05 | 0.795 | GATGCGGCCACGTTG |
| 1 | GMTACGGTCACACTG | chr3L:1869019-1871019 | 1786 | 1800 | - | 3.3571 | 7.12E-05 | 0.795 | GCTCCCGTCAACCTG |
| 1 | GMTACGGTCACACTG | chr3L:20105094-20107094 | 1791 | 1805 | - | 3.3571 | 7.12E-05 | 0.795 | GCTGCGGGCACATTC |
| 1 | GMTACGGTCACACTG | chr2L:8395096-8397096 | 1792 | 1806 | - | 3.3571 | 7.12E-05 | 0.795 | GACAAGGTCGCTCTG |
| 1 | GMTACGGTCACACTG | chr3L:11457490-11459490 | 1796 | 1810 | - | 3.3571 | 7.12E-05 | 0.795 | GATGCCGGCACCCTG |
| 1 | GMTACGGTCACACTG | chrX:14175923-14177923 | 1797 | 1811 | - | 3.3571 | 7.12E-05 | 0.795 | CCTGCGGGCACATTG |
| 1 | GMTACGGTCACACTG | chr2L:4443732-4445732 | 1799 | 1813 | - | 3.3571 | 7.12E-05 | 0.795 | GCTTCCGCCACATTG |
| 1 | GMTACGGTCACACTG | chr2R:22227574-22229574 | 1800 | 1814 | - | 3.3571 | 7.12E-05 | 0.795 | GCCGCGGTCACATTT |
| 1 | GMTACGGTCACACTG | chrX:19300873-19302873 | 1801 | 1815 | - | 3.3571 | 7.12E-05 | 0.795 | GATGCGGAGGCACTG |

| | | | | | | | | | |
|---|---|---|---|---|---|---|---|---|---|
| 1 | GMTACGGTCACACTG | chr3L:21515306-21517306 | 1801 | 1815 | - | 3.3571 | 7.12E-05 | 0.795 | ACGCTGGTCACACTG |
| 1 | GMTACGGTCACACTG | chr3R:32037330-32039330 | 1802 | 1816 | - | 3.3571 | 7.12E-05 | 0.795 | GCTCCGGTTCCAGTG |
| 1 | GMTACGGTCACACTG | chr2R:15612854-15614854 | 1802 | 1816 | - | 3.3571 | 7.12E-05 | 0.795 | GCGACCCTCCCACTG |
| 1 | GMTACGGTCACACTG | chr2R:16346500-16348500 | 1803 | 1817 | - | 3.3571 | 7.12E-05 | 0.795 | GCAACGCTTCCACTG |
| 1 | GMTACGGTCACACTG | chrX:9068741-9070741 | 1803 | 1817 | - | 3.3571 | 7.12E-05 | 0.795 | GATACGGAGCCGCTG |
| 1 | GMTACGGTCACACTG | chr2R:12751936-12753936 | 1804 | 1818 | - | 3.3571 | 7.12E-05 | 0.795 | GAGAAGGCCGCACTG |
| 1 | GMTACGGTCACACTG | chrY:3093575-3095575 | 1806 | 1820 | - | 3.3571 | 7.12E-05 | 0.795 | GCTGCGCTCACACAA |
| 1 | GMTACGGTCACACTG | chr2R:9845485-9847485 | 1808 | 1822 | - | 3.3571 | 7.12E-05 | 0.795 | GCTACGTTCACGCCT |
| 1 | GMTACGGTCACACTG | chrX:15378064-15380064 | 1810 | 1824 | - | 3.3571 | 7.12E-05 | 0.795 | GATCGGGACACCCTG |
| 1 | GMTACGGTCACACTG | chrX:6399100-6401100 | 1811 | 1825 | - | 3.3571 | 7.12E-05 | 0.795 | GAATCGGCCACACTC |
| 1 | GMTACGGTCACACTG | chr3R:8332792-8334792 | 1811 | 1825 | - | 3.3571 | 7.12E-05 | 0.795 | GCTACGGATACGCTT |
| 1 | GMTACGGTCACACTG | chrX:12028032-12030032 | 1814 | 1828 | - | 3.3571 | 7.12E-05 | 0.795 | GCTCCGCTCAGGCTG |
| 1 | GMTACGGTCACACTG | chr3R:31799963-31801963 | 1815 | 1829 | - | 3.3571 | 7.12E-05 | 0.795 | TAAACGGCCGCACTG |
| 1 | GMTACGGTCACACTG | chr3L:11457470-11459470 | 1816 | 1830 | - | 3.3571 | 7.12E-05 | 0.795 | GATGCCGGCACCCTG |
| 1 | GMTACGGTCACACTG | chr3L:22696230-22698230 | 1816 | 1830 | - | 3.3571 | 7.12E-05 | 0.795 | GCCACGCCCACACTA |
| 1 | GMTACGGTCACACTG | chr3R:6762478-6764478 | 1816 | 1830 | - | 3.3571 | 7.12E-05 | 0.795 | GATTCTGTCCCTCTG |
| 1 | GMTACGGTCACACTG | chr3L:276437-278437 | 1821 | 1835 | - | 3.3571 | 7.12E-05 | 0.795 | GCTTTGGTCGCAGTG |
| 1 | GMTACGGTCACACTG | chr2R:10809936-10811936 | 1822 | 1836 | - | 3.3571 | 7.12E-05 | 0.795 | CACACGGACGCACTG |
| 1 | GMTACGGTCACACTG | chr2L:7723576-7725576 | 1824 | 1838 | - | 3.3571 | 7.12E-05 | 0.795 | GCTGCGGTTCCACTT |
| 1 | GMTACGGTCACACTG | chr2R:16206706-16208706 | 1827 | 1841 | - | 3.3571 | 7.12E-05 | 0.795 | GATATGGTCTCGCAG |
| 1 | GMTACGGTCACACTG | chr2R:21182322-21184322 | 1828 | 1842 | - | 3.3571 | 7.12E-05 | 0.795 | GATACGGACCCACGT |
| 1 | GMTACGGTCACACTG | chr2L:8189781-8191781 | 1832 | 1846 | - | 3.3571 | 7.12E-05 | 0.795 | GCCACGCCCACACTA |
| 1 | GMTACGGTCACACTG | chr3L:18105149-18107149 | 1832 | 1846 | - | 3.3571 | 7.12E-05 | 0.795 | GAAACTGACGCACTG |
| 1 | GMTACGGTCACACTG | chr3L:1774633-1776633 | 1833 | 1847 | - | 3.3571 | 7.12E-05 | 0.795 | TCGACGGGCAGACTG |
| 1 | GMTACGGTCACACTG | chr2R:12184667-12186667 | 1834 | 1848 | - | 3.3571 | 7.12E-05 | 0.795 | GCTATGGTTGCGCTG |
| 1 | GMTACGGTCACACTG | chr3R:10261123-10263123 | 1834 | 1848 | - | 3.3571 | 7.12E-05 | 0.795 | GCAAAGGTGTCACTG |
| 1 | GMTACGGTCACACTG | chr2R:13963653-13965653 | 1836 | 1850 | - | 3.3571 | 7.12E-05 | 0.795 | CCTCCCGGCACACTG |
| 1 | GMTACGGTCACACTG | chr3R:5596739-5598739 | 1837 | 1851 | - | 3.3571 | 7.12E-05 | 0.795 | GCGAAGGTCACGCTA |
| 1 | GMTACGGTCACACTG | chr2L:10421780-10423780 | 1838 | 1852 | - | 3.3571 | 7.12E-05 | 0.795 | GCTGCGTTCACATCG |
| 1 | GMTACGGTCACACTG | chr2R:5791967-5793967 | 1842 | 1856 | - | 3.3571 | 7.12E-05 | 0.795 | GCCACGCCCACACTT |
| 1 | GMTACGGTCACACTG | chr2L:17409641-17411641 | 1845 | 1859 | - | 3.3571 | 7.12E-05 | 0.795 | GCTGCGGTCCAACTC |
| 1 | GMTACGGTCACACTG | chr2L:17588700-17590700 | 1845 | 1859 | - | 3.3571 | 7.12E-05 | 0.795 | GACACGGCCACCGTG |
| 1 | GMTACGGTCACACTG | chr3L:20493921-20495921 | 1846 | 1860 | - | 3.3571 | 7.12E-05 | 0.795 | GCCACGATCCCATTG |

| | | | | | | | | | |
|---|---|---|---|---|---|---|---|---|---|
| 1 | GMTACGGTCACACTG | chr3R:20579115-20581115 | 1846 | 1860 | - | 3.3571 | 7.12E-05 | 0.795 | GCTCCCTTCACACGG |
| 1 | GMTACGGTCACACTG | chr3R:24363980-24365980 | 1847 | 1861 | - | 3.3571 | 7.12E-05 | 0.795 | GCTGTGGTCCAACTG |
| 1 | GMTACGGTCACACTG | chrX:13284707-13286707 | 1848 | 1862 | - | 3.3571 | 7.12E-05 | 0.795 | GAGACCGCCACGCTG |
| 1 | GMTACGGTCACACTG | chr2L:4443680-4445680 | 1851 | 1865 | - | 3.3571 | 7.12E-05 | 0.795 | GCTTCCGCCACATTG |
| 1 | GMTACGGTCACACTG | chr3R:25198818-25200818 | 1858 | 1872 | - | 3.3571 | 7.12E-05 | 0.795 | GCCATGGTCACGATG |
| 1 | GMTACGGTCACACTG | chr2L:567121-569121 | 1862 | 1876 | - | 3.3571 | 7.12E-05 | 0.795 | GATCCGCTCGCTCTG |
| 1 | GMTACGGTCACACTG | chr2R:23693003-23695003 | 1867 | 1881 | - | 3.3571 | 7.12E-05 | 0.795 | GAGTCGGGCAGACTG |
| 1 | GMTACGGTCACACTG | chr3L:20969663-20971663 | 1868 | 1882 | - | 3.3571 | 7.12E-05 | 0.795 | GCAAGGGTTACACGG |
| 1 | GMTACGGTCACACTG | chr3R:8934126-8936126 | 1876 | 1890 | - | 3.3571 | 7.12E-05 | 0.795 | TCAGCGGTCACATTG |
| 1 | GMTACGGTCACACTG | chrX:9584675-9586675 | 1877 | 1891 | - | 3.3571 | 7.12E-05 | 0.795 | GATGCGGTAGCACAG |
| 1 | GMTACGGTCACACTG | chr3L:10770925-10772925 | 1878 | 1892 | - | 3.3571 | 7.12E-05 | 0.795 | GACTCAGTCGCACTG |
| 1 | GMTACGGTCACACTG | chr2L:6853142-6855142 | 1880 | 1894 | - | 3.3571 | 7.12E-05 | 0.795 | GCAACGATCACATAG |
| 1 | GMTACGGTCACACTG | chrX:8719520-8721520 | 1881 | 1895 | - | 3.3571 | 7.12E-05 | 0.795 | TAGACGGGCGCACTG |
| 1 | GMTACGGTCACACTG | chr3L:20200638-20202638 | 1883 | 1897 | - | 3.3571 | 7.12E-05 | 0.795 | CCTACTGCCCCACTG |
| 1 | GMTACGGTCACACTG | chr2L:1727732-1729732 | 1887 | 1901 | - | 3.3571 | 7.12E-05 | 0.795 | GCTTCCAACACACTG |
| 1 | GMTACGGTCACACTG | chrX:5621416-5623416 | 1887 | 1901 | - | 3.3571 | 7.12E-05 | 0.795 | CCCAAGGCCACACTG |
| 1 | GMTACGGTCACACTG | chr3L:14801168-14803168 | 1889 | 1903 | - | 3.3571 | 7.12E-05 | 0.795 | GATACGGACTCTTTG |
| 1 | GMTACGGTCACACTG | chr2L:4455807-4457807 | 1891 | 1905 | - | 3.3571 | 7.12E-05 | 0.795 | GCTACGCGAACTCTG |
| 1 | GMTACGGTCACACTG | chr2R:8913069-8915069 | 1892 | 1906 | - | 3.3571 | 7.12E-05 | 0.795 | GCTACGCGCACACAA |
| 1 | GMTACGGTCACACTG | chr2L:10059816-10061816 | 1892 | 1906 | - | 3.3571 | 7.12E-05 | 0.795 | GCGACGCGCACACTT |
| 1 | GMTACGGTCACACTG | chr2R:8743376-8745376 | 1894 | 1908 | - | 3.3571 | 7.12E-05 | 0.795 | TCGACGCTCACTCTG |
| 1 | GMTACGGTCACACTG | chr3L:4555087-4557087 | 1897 | 1911 | - | 3.3571 | 7.12E-05 | 0.795 | GAAAGGGGCACACGG |
| 1 | GMTACGGTCACACTG | chr2L:7403747-7405747 | 1899 | 1913 | - | 3.3571 | 7.12E-05 | 0.795 | GCATAGGTCACATTG |
| 1 | GMTACGGTCACACTG | chr2R:8260463-8262463 | 1900 | 1914 | - | 3.3571 | 7.12E-05 | 0.795 | GCTCGGGTGGCACTG |
| 1 | GMTACGGTCACACTG | chr3R:11981622-11983622 | 1901 | 1915 | - | 3.3571 | 7.12E-05 | 0.795 | TCGACGGTCTCACTT |
| 1 | GMTACGGTCACACTG | chr2R:13853659-13855659 | 1902 | 1916 | - | 3.3571 | 7.12E-05 | 0.795 | GAGACGGTCGCCATG |
| 1 | GMTACGGTCACACTG | chr3L:3218156-3220156 | 1903 | 1917 | - | 3.3571 | 7.12E-05 | 0.795 | GATACGGATACCCAG |
| 1 | GMTACGGTCACACTG | chrX:17080447-17082447 | 1904 | 1918 | - | 3.3571 | 7.12E-05 | 0.795 | GCAACGCCCAGACTG |
| 1 | GMTACGGTCACACTG | chrX:23290549-23292549 | 1904 | 1918 | - | 3.3571 | 7.12E-05 | 0.795 | GCTACGATCTCCATG |
| 1 | GMTACGGTCACACTG | chr2L:13826772-13828772 | 1905 | 1919 | - | 3.3571 | 7.12E-05 | 0.795 | GCGACGGTAACAAGG |
| 1 | GMTACGGTCACACTG | chr3R:12690743-12692743 | 1905 | 1919 | - | 3.3571 | 7.12E-05 | 0.795 | GCTTTGATCACACCG |
| 1 | GMTACGGTCACACTG | chrX:15582159-15584159 | 1908 | 1922 | - | 3.3571 | 7.12E-05 | 0.795 | TCGACAGTCACACAG |
| 1 | GMTACGGTCACACTG | chr2R:23720007-23722007 | 1908 | 1922 | - | 3.3571 | 7.12E-05 | 0.795 | ACAACGGTCACAAAG |

| | | | | | | | | | |
|---|---|---|---|---|---|---|---|---|---|
| 1 | GMTACGGTCACACTG | chrX:5852761-5854761 | 1909 | 1923 | - | 3.3571 | 7.12E-05 | 0.795 | GCCACAGTCACAAGG |
| 1 | GMTACGGTCACACTG | chr3R:27735538-27737538 | 1912 | 1926 | - | 3.3571 | 7.12E-05 | 0.795 | GCTACTCTCACCCAG |
| 1 | GMTACGGTCACACTG | chr2L:1868803-1870803 | 1913 | 1927 | - | 3.3571 | 7.12E-05 | 0.795 | ACTACGATCCCCCTG |
| 1 | GMTACGGTCACACTG | chr2L:19261240-19263240 | 1915 | 1929 | - | 3.3571 | 7.12E-05 | 0.795 | GCTCGGATCACACAG |
| 1 | GMTACGGTCACACTG | chrX:15466891-15468891 | 1919 | 1933 | - | 3.3571 | 7.12E-05 | 0.795 | GCTGAAGACACACTG |
| 1 | GMTACGGTCACACTG | chr3L:9398227-9400227 | 1921 | 1935 | - | 3.3571 | 7.12E-05 | 0.795 | CCTTCGATCACTCTG |
| 1 | GMTACGGTCACACTG | chr2R:20246624-20248624 | 1922 | 1936 | - | 3.3571 | 7.12E-05 | 0.795 | TATGCGGGCACGCTG |
| 1 | GMTACGGTCACACTG | chr2R:7621695-7623695 | 1922 | 1936 | - | 3.3571 | 7.12E-05 | 0.795 | GCTGCTGTCACTTTG |
| 1 | GMTACGGTCACACTG | chr3R:14495378-14497378 | 1922 | 1936 | - | 3.3571 | 7.12E-05 | 0.795 | GCTCCGGTCAGCGTG |
| 1 | GMTACGGTCACACTG | chrX:13284632-13286632 | 1923 | 1937 | - | 3.3571 | 7.12E-05 | 0.795 | GAGACCGCCACGCTG |
| 1 | GMTACGGTCACACTG | chrX:1379480-1381480 | 1928 | 1942 | - | 3.3571 | 7.12E-05 | 0.795 | GCGACGGTCACAAGC |
| 1 | GMTACGGTCACACTG | chr3R:23357812-23359812 | 1929 | 1943 | - | 3.3571 | 7.12E-05 | 0.795 | ACAACGGACACATTG |
| 1 | GMTACGGTCACACTG | chrX:14764787-14766787 | 1930 | 1944 | - | 3.3571 | 7.12E-05 | 0.795 | GCCACGTCCACACTT |
| 1 | GMTACGGTCACACTG | chr3L:21679050-21681050 | 1931 | 1945 | - | 3.3571 | 7.12E-05 | 0.795 | TCCGCGGTGACACTG |
| 1 | GMTACGGTCACACTG | chr3R:5631192-5633192 | 1932 | 1946 | - | 3.3571 | 7.12E-05 | 0.795 | TCGACGGGCAGACTG |
| 1 | GMTACGGTCACACTG | chrX:11560643-11562643 | 1938 | 1952 | - | 3.3571 | 7.12E-05 | 0.795 | GCTCAGCTCACCCTG |
| 1 | GMTACGGTCACACTG | chr2L:6037117-6039117 | 1939 | 1953 | - | 3.3571 | 7.12E-05 | 0.795 | TCTGCGACCACACTG |
| 1 | GMTACGGTCACACTG | chr2R:20562001-20564001 | 1941 | 1955 | - | 3.3571 | 7.12E-05 | 0.795 | GCCACGGTCAACGTG |
| 1 | GMTACGGTCACACTG | chr2R:10884019-10886019 | 1944 | 1958 | - | 3.3571 | 7.12E-05 | 0.795 | GCGAAGGTGACACCG |
| 1 | GMTACGGTCACACTG | chr3R:25044151-25046151 | 1946 | 1960 | - | 3.3571 | 7.12E-05 | 0.795 | GCTACCGTCTCACGT |
| 1 | GMTACGGTCACACTG | chr2L:11787334-11789334 | 1946 | 1960 | - | 3.3571 | 7.12E-05 | 0.795 | GAGACGGCGACGCTG |
| 1 | GMTACGGTCACACTG | chr3R:29674421-29676421 | 1952 | 1966 | - | 3.3571 | 7.12E-05 | 0.795 | GCTACGGGCGGACTC |
| 1 | GMTACGGTCACACTG | chr3L:19611199-19613199 | 1957 | 1971 | - | 3.3571 | 7.12E-05 | 0.795 | GCTCCGATCACGTTG |
| 1 | GMTACGGTCACACTG | chr2L:9917330-9919330 | 1959 | 1973 | - | 3.3571 | 7.12E-05 | 0.795 | GCTGTGGGAACACTG |
| 1 | GMTACGGTCACACTG | chr3R:18020600-18022600 | 1960 | 1974 | - | 3.3571 | 7.12E-05 | 0.795 | GCGACGCCCACAATG |
| 1 | GMTACGGTCACACTG | chrX:15205636-15207636 | 1961 | 1975 | - | 3.3571 | 7.12E-05 | 0.795 | TCCACGGCCACATTG |
| 1 | GMTACGGTCACACTG | chr3L:7947186-7949186 | 1962 | 1976 | - | 3.3571 | 7.12E-05 | 0.795 | GAGGCGGTCACAAAG |
| 1 | GMTACGGTCACACTG | chr3R:6254175-6256175 | 1962 | 1976 | - | 3.3571 | 7.12E-05 | 0.795 | GCCAGGGGCACAATG |
| 1 | GMTACGGTCACACTG | chrX:1029758-1031758 | 1964 | 1978 | - | 3.3571 | 7.12E-05 | 0.795 | GCCACGATCCCAGTG |
| 1 | GMTACGGTCACACTG | chrX:18225076-18227076 | 1967 | 1981 | - | 3.3571 | 7.12E-05 | 0.795 | GAGGCGGCCACATTG |
| 1 | GMTACGGTCACACTG | chr3R:6026767-6028767 | 1967 | 1981 | - | 3.3571 | 7.12E-05 | 0.795 | GCTTCGGTAAGACAG |
| 1 | GMTACGGTCACACTG | chr3R:4276928-4278928 | 1969 | 1983 | - | 3.3571 | 7.12E-05 | 0.795 | GCCACGCCCACACTA |
| 1 | GMTACGGTCACACTG | chr2R:9601901-9603901 | 1971 | 1985 | - | 3.3571 | 7.12E-05 | 0.795 | GCCATGGTCACAGGG |

| | | | | | | | | | |
|---|---|---|---|---|---|---|---|---|---|
| 1 | GMTACGGTCACACTG | chr2R:18611730-18613730 | 1972 | 1986 | - | 3.3571 | 7.12E-05 | 0.795 | GCAATGGACACAATG |
| 1 | GMTACGGTCACACTG | chr2R:21633679-21635679 | 1973 | 1987 | - | 3.3571 | 7.12E-05 | 0.795 | GCCACGGTAAGACCG |
| 1 | GMTACGGTCACACTG | chr4:1087798-1089798 | 1980 | 1994 | - | 3.3571 | 7.12E-05 | 0.795 | CAGACGGACACACGG |
| 1 | GMTACGGTCACACTG | chr2L:17531234-17533234 | 1982 | 1996 | - | 3.3571 | 7.12E-05 | 0.795 | GATGGGGTCACGCAG |
| 1 | GMTACGGTCACACTG | chrX:19700755-19702755 | 1983 | 1997 | - | 3.3571 | 7.12E-05 | 0.795 | GACACGGCGACACGG |
| 1 | GMTACGGTCACACTG | chr2R:21289343-21291343 | 1984 | 1998 | - | 3.3571 | 7.12E-05 | 0.795 | ACTACGATCGCACAG |
| 1 | GMTACGGTCACACTG | chrX:19321106-19323106 | 1986 | 2000 | - | 3.3571 | 7.12E-05 | 0.795 | GCTACTGTCTACCTG |

**Supplementary Table 5**. Enrichment of Motif 1 at +/- 1kb of all TSS at or below *p* = 7.12E-05 analyzed by FIMO (Find Individual Motif Occurrences). The format of this file is described at http://meme-suite.org/doc/fimo-output-format.html.
